# Finding the LMA needle in the wheat proteome haystack

**DOI:** 10.1101/2023.01.22.525108

**Authors:** Delphine Vincent, AnhDuyen Bui, Vilnis Ezernieks, Saleh Shahinfar, Timothy Luke, Doris Ram, Nicholas Rigas, Joe Panozzo, Simone Rochfort, Hans Daetwyler, Matthew Hayden

## Abstract

Late maturity alpha-amylase (LMA) is a wheat genetic defect causing the synthesis of high isoelectric point (pI) alpha-amylase in the aleurone as a result of a temperature shock during mid-grain development or prolonged cold throughout grain development leading to an unacceptable low falling numbers (FN) at harvest or during storage. High pI alpha-amylase is normally not synthesized until after maturity in seeds when they may sprout in response to rain or germinate following sowing the next season’s crop. Whilst the physiology is well understood, the biochemical mechanisms involved in grain LMA response remain unclear. We have employed high-throughput proteomics to analyse thousands of wheat flours displaying a range of LMA values. We have applied an array of statistical analyses to select LMA-responsive biomarkers and we have mined them using a suite of tools applicable to wheat proteins. To our knowledge, this is not only the first proteomics study tackling the wheat LMA issue, but also the largest plant-based proteomics study published to date. Logistics, technicalities, requirements, and bottlenecks of such an ambitious large-scale high-throughput proteomics experiment along with the challenges associated with big data analyses are discussed. We observed that stored LMA-affected grains activated their primary metabolisms such as glycolysis and gluconeogenesis, TCA cycle, along with DNA- and RNA binding mechanisms, as well as protein translation. This logically transitioned to protein folding activities driven by chaperones and protein disulfide isomerase, as well as protein assembly via dimerisation and complexing. The secondary metabolism was also mobilised with the up-regulation of phytohormones, chemical and defense responses. LMA further invoked cellular structures among which ribosomes, microtubules, and chromatin. Finally, and unsurprisingly, LMA expression greatly impacted grain starch and other carbohydrates with the up-regulation of alpha-gliadins and starch metabolism, whereas LMW glutenin, stachyose, sucrose, UDP-galactose and UDP-glucose were down-regulated. This work demonstrates that proteomics deserves to be part of the wheat LMA molecular toolkit and should be adopted by LMA scientists and breeders in the future.

## Introduction

Common bread wheat (*Triticum aestivum L*.) is the dominant crop in temperate regions, currently covering more than 220 million hectares worldwide, exceeding 749 million tons in production annually (1) and predicted to reach 835 million tons by 2030 (2). From the most primitive form of wheat 10,000 years ago in the Fertile Crescent to the species currently grown all over the world, desirable characteristics have been selected and improved upon by human societies (3). Hexaploid *T. aestivum* (AABBDD; 2n = 6x = 42) originated from two polyploidization events. The first event associated diploid *Triticum urartu* (AA; 2n = 2x = 14) which provided the A genome with the other yet unknown species from the *Sitopsis* section of *Triticum* genus which provided the B genome to produce the allotetraploid wild emmer wheat (*Triticum turgidum*; AABB; 2n = 4x = 28). The second event associated *T. turgidum* with *Aegilops tauschii* (DD) (4, 5). The chromosomes from each closely related progenitor are grouped into homeologous groups. Because of the shared ancestry, genes may be common to all members of a homeologous group, albeit exhibiting high allelic variation and differences in gene count due gene duplication or silencing (2). Millenia of domestication have accrued an enormous genetic diversity in this species, with potentially more than 50,000 *T. aestivum* cultivars (6). Wheat owes its success to adaptability to temperate, Mediterranean, and subtropical climates, high yields, storability, but above all to the unique properties of doughs, which can be processed into a vast range of foods (3, 5). Wheat grains are not only a major source of carbohydrate in the form of starch, but also a great source of protein. The endosperm prolamins proteins comprise gliadins and glutenins; they are the main components of gluten and together confer unique viscoelastic and rheological properties to flour mixed with water. Indeed, hydrated gliadins largely determine the viscosity and extensibility of the dough, while the cohesive properties of hydrated glutenins essentially govern the strength and elasticity of the dough (5). Wheat seeds also contribute essential amino acids, minerals, vitamins, beneficial phytochemicals and dietary fibre components to the human diet. Beside nutritional benefits, different parts of the wheat plant confer advantageous medicinal uses such as anticancer properties of wheat bran and antimicrobial activities of wheat sprouts (7).

Currentbreedingprograms mainly aim at sustainingwheatproduction and quality with reduced agrochemical inputs, as well as developing new disease-resistant and stress-tolerant varieties with enhanced quality for specific end-uses (8). Wheat research and breeding must accelerate genetic gain to keep augmenting crop yield while maintaining or improving grain quality traits if the demands of the growing human population are to be met (9). A critical element in the equation was the sequencing and functional annotation of the genome. Sequencing the hexaploid bread wheat genome was a gigantic achievement proportionate to its large size, abundance of repetitive DNA and the immense difficulty of discerning homoeologs from subgenomes A, B and D. Whilst this required the commitment of 20 countries collaborating as a consortium (International Wheat Genome Sequencing Consortium IWGSC) and a lot of strategizing from 2005 onward, including sequencing diploid and tetraploid ancestors, it was the advent of next generation sequencing technologies producing long but error-prone or accurate yet short reads that made this massive endeavour successful (10). A 13-year effort ensued, drafting *T. aestivum* genome in 2014 based on key breakthrough short read technologies by NRGene (4), and culminating in 2018 with the release of the long-awaited fully assembled and annotated 14.5 Gb reference genome, cataloguing 107,891 high-confidence genes along 21 chromosome-like sequence assemblies (IWGSC RefSeq v1.0) (9). This helped bridge the gap on wheat research relative to other cereal model species such as rice, sorghum, corn and barley whose genomes had been sequenced years ago, and propelled wheat post-genomics studies forward with a continuous increase in publications since 2011. Both the numbers of high confidence protein-coding genes from subgenomes A, B, and D and their composition were largely similar (9). Transcriptomics analyses of genes present in all three subgenomes notonly showedcomparable expressionlevels for 72.5% of them, especially those located in syntenic regions, but also unveiled the lack of significant subgenome expression dominance (11, 12). As valuable such an asset was, it did not capture the extent of the wheat genomic diversity as only one cultivar, Chinese Spring, was chosen as a template. In fact, no single genome assembly can be sufficient to model the wheat proteome due to the high allelic and gene copy number variability (2). This shortfall was addressed in 2020 when 15 hexaploid wheat lines from different regions, growth habits and breeding programs were sequenced and annotated against IWGSC reference genome (13). Such pan-genomic comparative analysis outlined extensive structural rearrangements, introgressions from wild relatives and differences in gene content arising from complex breeding events to boost resistance to biotic and abiotic stresses, as well as grain yield and quality. Unfortunately, fasta sequences of annotated proteins are not publicly available for these 15 assemblies. A refined version of the reference genome using optical mapping and long sequence reads was recently released (IWGSC RefSeq v2.1) (14). With such worthwhile genomic resources in store, wheat can now be instated as a model for plant genetic research and employed to tackle complex biological questions on evolution, domestication, polyploidization, as well as genetic and epigenetic interaction between homoeologous genes and genomes (10). Moreover, genome annotations paves the way to investigate pathways and biochemical attributes behind bread wheat quality using transcriptomics (15) or proteomics (2) approaches.

The industry will equally benefit from these latest scientific developments since processing companies, markets and food industries demand not only high yielding and resistant varieties, but also those with specific end-use qualities (1, 3). Marketrequirements haveinfluencedwheat breeding as not to neglect essential protein content and quality. Because wheat is generally traded according to grain protein content and hardness, standards must be abided to by producers and distributors. Intact starch polymers provide the gelatinization and retrogradation needed for an acceptable product. Failure to meet receival standards for milling grades due to starch degradation measured in the wheat industry using the Hagberg–Perten falling number (FN) method (16) leads to grain discount and downgrading to animal feed, which incurs a loss of profit (17). The low FN values manifests as a loss of viscosity upon mixing starch-degraded flour with water can alter appearance and texture of end-products (18), however, it might not deteriorate baking functionality (19) and could be used instead in alternate preparations (20). There are multiple causes of low FN symptomatic of starch degradation including preharvest sprouting, late maturity alpha-amylase (LMA), and variation in kernel starch and protein (21). LMA is a wheat genetic defect causing the synthesis of high isoelectric point (pI) alpha-amylase in the aleurone as a result of a temperature shock during mid-grain development or prolonged cold throughout grain development leading to an unacceptable low FN at harvest or during storage (22–24). High pI alpha-amylase is normally not synthesized until after maturity in seeds when they may sprout in response to rain or germinate following sowing the next season’s crop (25).

Four alpha-amylase isoforms have been identified to date in wheat. Several α-amylase 1 (TaAMY1) loci have been localized on the long arm of group 6 chromosomes (26). In LMA-prone wheat genotypes and under given temperatures, Amy-1 genes are transcribed in isolated cells or cell islands distributed throughout the aleurone system of grains with a 50-60% moisture content before they have reached physiological maturity (25). Appearance of high pI a-amylase protein is preceded by a short-lived transient period of mRNA synthesis leading to a stable enzyme and retained through to seed maturity (22, 27). Multiple alpha-amylase 2 (TaAMY2) loci are positioned on the long arm of the group 7 chromosomes and produce a low pI alpha-amylase in the pericarp of the developing grain (28). A single locus encodes alpha-amylase 3 (TaAMY3) on group 5 chromosomes and is transcribed throughout the grain development suggesting a role in grain development and maturation (29). Similar to TaAMY2, TaAMY3 enzyme mainly appears during grain development in the pericarp and would be the predominant alpha-amylase enzyme throughout grain development (30). Despite its shorter length and elevated pI, TaAMY3 displays equal numbers of calcium-binding and active sites relative to the other three isoforms; however, the distance between key AA residues and the last two active site residues is shortened (31). Overexpressing TaAMY3 in the endosperm of developing grain to levels of up to 100-fold higher than the wild-type results in low FN similar to those seen in LMA-affected grains, yet has no detrimental effect neither on starch structure, flour composition and baking quality of bread (32), nor on noodle colour or firmness (33). A fourth isoform alpha-amylase 4 (TaAMY4) is also encoded by a single locus on group 5 chromosomes and is co-expressed with TaAMY1 in LMA-affected grains (31). Comparison of the four isoforms revealed that they contain 385-439 AAs, with a molecular mass between 45.4-48.3 kD, and a pI ranging from 5.5 to 8.6. All isoforms differ slightly in their 3-D protein structure including the presence of additional sugar binding domains hinting to various enzymatic properties (31, 34).

Although LMA expression correlates with measurable changes in both hormone content and transcript profiles during grain maturation, there are no obvious visual effects on grain appearance, development, or morphology (24), hence the need to perform assays to test for its activity (16). ELISA (35) and RT-qPCR (36) assays were developed to specifically target TaAMY1, the main enzyme involved in LMA. One limitation to the RT-qPCR method relates to the apparent short life of the high pI a-amylase mRNA (22). Commonly employed is the colorimetric Ceralpha assay (37) whereby the alpha-amylase activity is expressed in terms of Ceralpha units per gram of flour (u/g). A single unit corresponds to the amount of enzyme required to release 1 µMp-nitrophenylin the presence of excess quantities of alpha-glucosidase in 1 min at 40°C (38). Such measurements have revealed that LMA is more prevalent than originally thought, with reports arising from North America, Australia, Japan, Canada, South Africa, China, Mexico, Germany, and the United Kingdom (39). The presence of LMA in breeding populations could be attributed to unexplained positive effects on grain production/quality or alternately simply manifest the lack of significant selection pressure against this trait (24). Both a cool temperature shock near physiological maturity or continuous cool maximum temperatures during grain development can induce LMA synthesis in wheat (23). The prediction of LMA occurrence during LMA dedicated field trial is impeded by the stochastic nature of LMA expression resulting from specific genetics, climatic conditions, and developmental stages.

LMA has a genetic (G) component (alpha-amylase gene required), yet it is only expressed and enzymatically active under particular environmental (E) conditions (temperature shock) at a given developmental stage making it the product of a GxE interaction, which lends itself to post-genomic quantitative studies to shed some lights into the biological mechanisms underpinning LMA expression. Yet, to date, only one LMA-related transcriptomics study has been published and no proteomics work has been attempted despite the potential this technology offers to help improve bread wheat quality (2). Using microarray technology, Barrero and colleagues reported that LMA resulted from very narrow and transitory peak of expression of genes encoding high-pI alpha-amylase during grain development (22). Furthermore, the LMA phenotype triggered elevated levels of gibberellins such as GA19 and much lower levels of auxin in the de-embryonated fraction of grains sampled shortly after the initiation of LMA synthesis. A recent report questions this hormonal response since, unlike alpha-amylase synthesis by aleurone during germination or following treatment with exogenous GA, alpha-amylase synthesis by wheat aleurone during grain development appears to be independent of gibberellin (40). Even though on one hand genomics can catalogue genes present in a sample and possibly the biological context of their expression and on the other hand transcriptomics can validate expression levels, only proteomics can measure the actual protein abundance, recordpost-translationalmodification (PTM), as well as identify interacting proteins (2). We have developed a high-throughput proteomics method to rapidly profile *T. aestivum* grains and data mine their proteome (41). In the present study, we have applied our optimised procedure to a collection of in excess of 4,000 wheat cultivars and germplasm whose LMA content ranged from 0 to 8 u/g of flour. We have applied an array of statistical analyses to our big data to select LMA-responsive biomarkers and we have mined them using a suite of tools applicable to wheat proteins, yet not necessarily embraced by grain scientists. To our knowledge, this is not only the first proteomics study tackling the wheat LMA issue but also the largest plant-based proteomics study published to date. Logistics, technicalities, requirements, and bottlenecks of such an ambitious large-scale high-throughput proteomics experiment along with the challenges associated with big data analyses are discussed.

## 2. Materials and Methods

### 2.1. Wheat Cultivation, Sampling, and Storage

The wheat collection used in this study represents a diverse range of cultivars and germplasm sourced through the Australian Grains Genebank and representingglobalgenetic diversity. The wheat was grown in field trials at Horsham Victoria and harvested using a mechanical small-plot harvester. The threshed grain was stored in seal containers at 20°C. The environmental conditions (rain and temperature) at the trial site were monitored throughout the growing season. No preharvest rainfall was recorded and therefore any α-amylase activity was non-germinative but associated with LMA.

The list of wheat samples is supplied in Supplementary Table S1.

### 2.2. LMA assay

The alpha-amylase assay was performed usingthe Megazyme assay accordingto the procedure reported by McCleary and Sheehan (42) on 3,773 grain samples (Supplementary Table S1). The distribution of LMA values was plotted as a histogram in Microsoft Excel. Various transformations were performed to achieve a normal distribution such as standardisation, log natural, log 2, inverse and standardisationof inversedvalues (data notshown).The transformed values were also plotted as histograms to check for gaussian distribution.

### 2.3. Wheat Grain Processing for proteomics analyses

Sample preparation was optimised and thoroughly described (41); it is schematised in Figure 1. All sample packages were mixed together in a box for randomisation and assigned a unique number as they were processed. QR codes on sample bags and tubes were scanned and consigned to the Excel spreadsheet using a handheld barcode scanner (model 1902 GHD-2, Honeywell Australia, Matraville, NSW). All microtubes were pre-labelled with unique numbers and sample IDs, both also consigned to a QR code, usinga handheld label maker (PT-E550WVP, Brother, Australia) controlled by the P-touch editor software (Brother, Australia) fitted with 12mm white laminated tape.

**Figure 1.**
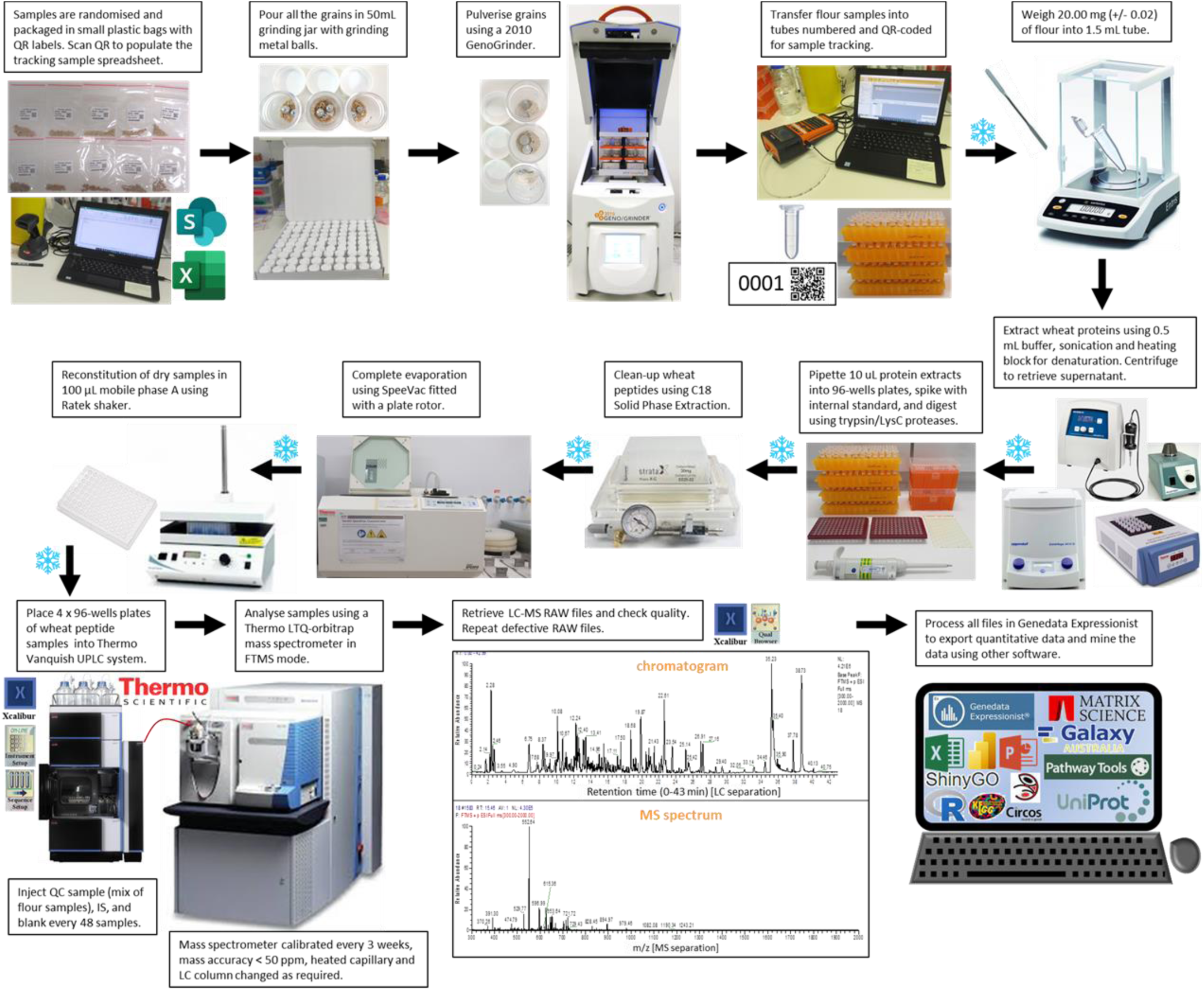
High-throughput workflow used on the 4061 wheat samples. The snowflakes indicate storage in-80°C freezers.

The grains were ground in 50 mL jars containing two 8 mm and two 3 mm metal grinding balls using an automated tissue homogeniser and cell lyser (Geno/Grinder® 2010, SPEX SamplePrep, Metuchen, NJ, USA) and pulverised into fine flour twice for 2 min at 1,500 rpm with a 15 s break in between. A total of 600 jars were employed in a rotation. Dirty jars and balls were rinsed to remove excess flour and soaked in 1% decon 90 surfactant (Decon, Hove, UK) for 2 hours followed by a thorough wash in a dishwasher with RO water and left to air dry prior to being used again. A wheat quality control (QC) sample was prepared by sampling 50 mg (±0.05 mg) from each of the 96 flour samples described in (41) and mixing them all thoroughly.

A 20 mg (±0.2 mg) aliquot of flour was weighed in a 1.5 mL microtube and resuspended in 0.5 mL Gnd-HCl buffer (6 M Guanidine hydrochloride, 0.1 M Bis-Tris, 10 mM DTT, 5.37 mM sodium citrate tribasic dihydrate) using a MS 1.5 sonicator probe (Ultrasonic Homogeniser SONOPULS mini 20, Bandelin, Berlin, Germany) for 30 s with 90% amplitude. The tubes were briefly vortexed (5 sec each, RAVM1 Ratek Vortex Mixers, Ratek, Boronia, VIC, Australia) and incubated for 60 min in a thermoblock (Digital Dry Bath/Block Heater, Thermo Scientific, Scoresby, VIC, Australia) at 60°C. The tubes were left to cool to room temperature for 5 min and 10 µL of 1 M iodoacetamide was added to each tube. The tubes were thoroughly mixed for 30 s using a rack vortex mixer (MTV1 Multi Tube Vortex Mixer, Ratek, Boronia, VIC, Australia) at high speed and left to incubate at room temperature in the dark for 30 min. The tubes were centrifuged usinga benchtop centrifuge (5415D Digital Microfuge, Eppendorf, Macquarie Park, NSW, Australia) at 13,000 rpm for 15 min at room temperature and the supernatant was transferred into a fresh 1.5 mL microtube pre-labelled with the QR code.

Two vials of trypsin/Lys-C mix (100µg, V5078, Promega, Alexandria, NSW, Australia) were dissolved into 1 mL of the resuspension buffer (50mM acetic acid) supplied by the manufacturer and kept on ice until use to digest 192 wheat samples at a time. Aliquots of 10 µL aliquotof protein extracts were transferredinto two 96-well plates (Strata 96-wellcollection plate, 350 µL conical polypropylene, Phenomenex, Lane Cove, NSW, Australia), diluted 6 times with 50 mM ammonium bicarbonate and digested with 5 µL aliquots of the trypsin/Lys-C solution prepared earlier. Plates were sealed with silicone covers (pierceable sealing mats, 96-square well, Phenomenex, Lane Cove, NSW, Australia) and vortexed for 30 s using a rack vortex mixer (MTV1 Multi Tube Vortex Mixer, Ratek, Boronia, VIC, Australia) at high speed. Plates were incubated at 37°C for 17 hours. Aliquots of 7 µL 10% formic acid (FA)/water were added to stop the digestion. An internal standard (IS, [Glu1]-fibrinopeptide B human, F3261, Sigma, Port Melbourne, VIC, Australia) was added at a final concentration of 1 µg.

Protein digests were cleaned using 96-wells solid phase extraction (SPE) plates (Strata C18-E 100 mg P/N 8E-S001-EGB, Phenomenex, Lane Cove, NSW, Australia) and fully evaporated as described in (Vincent, Bui et al. 2022). Peptide digests were reconstituted by adding 70 µL of 0.1% FA/water to each well. The digests were dissolved by shaking the plates for 50 min at medium speed using a rack vortex mixer (MTV1 Multi Tube Vortex Mixer, Ratek, Boronia, VIC, Australia) at room temperature. The collection plates were sealed with a silicone lid and stored at −80 °C until LC-MS analysis.

### 2.4. LC-MS analyses

All 4,061 wheat and QC samples were processed using the LC-MS method listed below. Liquid chromatography (LC) was optimised (41). Our chosen LC method applied 0.2 mL/min flow rate, 38 min LC run duration, 6% B for 2.5 min, 6 –36% B gradient for 30.5 min, increased up to 98% B gradient for 0.1 min, 98% B for 5 min, drop down to 3% B in 0.1 min, 6% B for 5 min. The LC system used was a Vanquish Flex Binary UHPLC System (Vanquish UHPLC+ focused, ThermoFisher Scientific, Scoresby, VIC, Australia). Mobile phase A was 0.1% FA/water and mobile phase B was 0.1% FA/acetonitrile (ACN). The needle wash solution was 80% isopropanol (IPA)/water, and the rear seal wash solution was 10% IPA/water. The needle wash solution was 10% IPA/water. The needle was washed after each injection. The rack types were specified as DeepWell96 in the LC-MS method and the SamplerModule tab of Xcalibur Direct Control software (version 3.0.63, ThermoFisher Scientific, Scoresby, VIC, Australia) with a 29,000 µm injection depth. Blanks (0.1% FA/water) and QC were injected from two 10 mL vials. Peptides were separated using a RP–LC column (bioZen 1.7 um Peptide XB-C18, 100 Å, LC column 150 × 2.1 mm, Phenomenex, Lane Cove, NSW, Australia) using a 60°C oven temperature. The blank, IS and QC samples were injected every 48 samples for normalisation purposes. The IS was used to check for mass accuracy (<50ppm). The LC separation column was changed with a new one when peak resolution degraded (every 1000 samples or so).

The UHPLC was online with an Orbitrap Velos hybrid ion trap–Orbitrap mass spectrometer (ThermoFisher Scientific, Scoresby, VIC, Australia) fitted with a heated electrospray ionisation (HESI) source. Every three weeks, the instrument was mass calibrated, and the source sweeping cone and the heated capillary were cleaned. HESI parameters were: needle at 3.9 kV, 100 µA, sheath gas flow 20, auxiliary gas flow 7, sweep gas flow 2, source heated to 200°C, capillary heated to 275°C, and S-Lens RF level 55%. Spectra were acquired using the full MS scan mode of the Fourier transform (FT) orbitrap mass analyser (FTMS) in positive ion mode at a resolution of 15,000 along a 300–2000 m/z mass window in profile mode with 3 microscans.

The sequence lists were prepared in advance in Excel as .cvs files and imported into Xcalibur data acquisition software (version 3.0.63); five sequences were needed as Xcalibur only accommodated a maximum of 1000 lines. Because samples had been randomised, 96-well plates were analysed consecutively. Throughout the LC-MS run, the RAW files were individually visualised using Xcalibur Qual Browser (version 3.0.63,). Files that failed to pass our check (loss of peak resolution, incomplete run, no signal, mass accuracy > 50 ppm, etc…) were rerun concomitantly to when LC-MS was interrupted for maintenance.

### 2.5. LC-MS/MS analyses

For protein identification, 400 random samples (10% samples, 4 plates) were used following the LC-MS1 analysis. LC, HESI and full scan FTMS parameters were as indicated above. MS2 data was acquired using ITMS in positive mode as centroid values and applied various methods summarised below. In an attempt to maximise the number of peptides sequenced, several passes were performed with inclusion and exclusion lists, and various parameters.

Pass 1: FTMS parameters were as specified above. Using the Nth order double play method, MS/MS spectra were acquired in data-dependent mode. Singly charged peptides were ignored. In the linear ion trap, the 10 most abundant peaks with charge state >2 and a minimum signal threshold of 3,000 were fragmented using collision-induced dissociation (CID) with a normalised collision energy of 35%, 0.25 activation Q, and activation time of 10 ms. The precursor isolation width was 2 m/z. Dynamic exclusion was activated, and peptides selected for fragmentation more than once within 30 s were excluded from selection for 180 s. No inclusion or exclusion list was used; however, a list of MS2 event was produced by exporting the “Scan Filters” of the RAW file in Xcalibur Qual Browser (ThermoFisher Scientific, Scoresby, VIC, Australia) and to be used in Pass 2 as an exclusion list containing 2,000 unique *m/z* values (maximum number allowed in Xcalibur). This method was run in duplicate.

Pass 2: Same method as Pass 1, except that the list of MS2 events generated in Pass 1 was uploaded in the Data Dependent Settings as a Reject Mass List. Like in Pass 1, a list of MS2 event was produced by exporting the “Scan Filters” of the RAW file and to be used in Pass 3 as an exclusion list containing 1,997 unique *m/z* values. This method was run in triplicate.

Pass 3: Same method as Pass 2, except that the list of MS2 events generated in Pass 2 was uploaded in the Data Dependent Settings as a Reject Mass List. Like in Pass 2, a list of MS2 event was produced by exporting the “Scan Filters” of the RAW file and to be used in Pass 4 as an exclusion list containing 1,998 unique *m/z* values. This method was run in duplicate.

Pass 4: Same method as Pass 3, except that the list of MS2 events generated in Pass 3 was uploaded in the Data Dependent Settings as a Reject Mass List. This was the last exclusion list used in this study. This method was run in duplicate.

Pass 5: Same method as Pass 1, except that the threshold was dropped from 3,000 down to 500 to perform MS2 on peptides of low abundance. This method was run in duplicate.

Pass 6: Same method as Pass 1, except with a Parent Mass List (i.e. an inclusion list) made out of the 2,000 most abundant peptides. This method was run in duplicate.

For the following methods, LC-MS1 reproducible peptides for which intensity exceeded 0.0001 (19,956 peptides in total) were randomised along retention time (RT) and divided into 10 lists (inclusion lists 1 to 10 containing <2,000 *m/z* values each).

Pass 7: FTMS parameters were as specified above. Using the global MS/MSn method, MS/MS spectra were acquired in non-data dependent mode. ITMS parameters were: CID with a normalised collision energy of 35%, 0.25 activation Q, isolation width of 1. and activation time of 10 ms. Inclusion list 1 was uploaded in the inclusion global MS/MS mass list tab of the Global Non-Data Dependent Settings. All remainingnine parentlists were loaded to individual pass 7 methods.

Pass 8: FTMS parameters were as specified above. ITMS parameters were: CID with a normalised collision energy of 35%, 0.25 activation Q, and activation time of 10 ms. The precursor isolation width was 2 m/z. The signal threshold was 500. Inclusion list 1 was uploaded in the parent mass list of the data-dependent settings. All remaining nine parent lists were loaded to individual pass 8 methods.

Pass 9: Same method as Pass 8, except that the precursor isolation width was 1 m/z to increase the mass accuracy the m/z values targeted in the parent mass list. All remaining nine parent lists were loaded to individual pass 9 methods.

Pass 10: Same method as Pass 8, exceptthatthe precursor isolation width was 0.5 m/z to further increase the mass accuracy the m/z values targeted in the parent mass list. All remaining nine parent lists were loaded to individual pass 10 methods.

Pass 11: Same method as Pass 8, except that the precursor isolation width was 0.2 m/z to target the parent masses as accurately as possible. All remaining nine parent lists were loaded to individual pass 11 methods.

All the Xcalibur parameters of the various MS/MS methods can be found in Supplementary File SF1. Exclusion and inclusion lists can be found in Supplementary File SF2. A total of 63 LC-MS2 files were thus acquired; they are available from the MassIVE repository ((https://massive.ucsd.edu/ProteoSAFe/static/massive.jsp, MSV000090572).

### 2.6. Proteomics data processing

The LC–MS RAW files of the 4,061 wheat samples along with the 86 QC and IS replicates (injected once every 48 what samples) were processed in the Refiner MS module of Genedata Expressionist® 13.0 (Genedata AG, Basel, Switzerland. To process all files in one batch, a stepwise workflow was devised (Supplementary Figure S1A-B).

In the first step, a repetition activity was used (processing one file at a time) in which the consecutive sub-activities were performed: 1/ Load from File, 2/ RT Structure Removal with a minimum of 4 scans and m/z Structure Removal with a minimum of 8 points, 3/ Chromatogram Smoothing using a 3 scan RT window and a Moving Average estimator, 4/ RT Structure Removal with a minimum of 5 scans, and 5/ Save Snapshot to export all the processed files individually. The files were individually checked for inconsistencies that would invalidate the subsequentquantitativeanalyses. Inadequate files were removed from the dataset leaving 3,990 reproducible wheat files. In the second step (Supplementary Figure S1C), the activities applied were: 1/ Load from File on the left for all the samples and on the right for the QCs, 2/ Adaptative Grid with 10 *m/z* scan counts, 3/ Average across Experiments (files) using the arithmetic mean, 4/ Reference Grid joining both sides, 5/ Chromatogram RT Alignment applying a maximum RT shift of 50 scans (30 s), 6/ Chromatogram Peak Detection using a 12 scan Summation Window, Minimum Peak Size of 8 scans, Maximum Merge Distance of 5 points and Boundaries Merge Strategy, 10% Gap/Peak ratio for Peak RT Splitting, 3 points for *m/z* Smoothing, Ascent-based Peak Detection with 3 points Isolation Threshold, Local Maximum Centre Computation and Maximum Curvature Boundary Determination, 7/ Chromatogram Isotope Clustering with 0.1 min RT Tolerance and 20 ppm *m/z* Tolerance, the Peptide Isotope Shaping method with Protonation Ionisation, Minimum Charge of 2 and Maximum Charge of 10, Maximum Log-Ratio Distance of 0.8, and Variable Charge Dependency for Cluster Size Restriction, 8/ Singleton Filter, 9/ Metadata Import, 10/ Save Snapshot, and 11/ Export Analyst of the Clusters using the Integrated Maximum Intensity.

LC-MS processed quantitative data and metadata (sample description, LMA measurements, sample preparation technical steps, LC-MS sequence, instrument maintenance, etc…) were exported into Genedata Analyst (version 13, Genedata AG, Basel, Switzerland) for normalisation purposes (Supplementary Figure S1D). Data file normalisation with three consecutive steps was reported (41). In brief, first the quantities were normalised using the flour weights (1% accuracy) to account for sample preparation variation, second the IS cluster was used to normalise peptide abundances in order to take into consideration post-digestion technical variation, and third QCs and injection order were taken into account to correct instrument variation over time. The normalised quantitative data was exported as a CSV file for further processing. The CSV file contained 44,444 rows (peptide clusters) and 3,990 columns (wheat samples).

The effects of technical biases on the LC-MS spectra were quantified using ANOVA simultaneous component analysis (ASCA), a generalisation of ANOVA which quantifies the variation induced by fixed experimental factors on complex multivariate datasets (43). Firstly, the normalised data were imported into R where clusters containing 100% missing values were removed (n = 12,108), leaving 32,336 peptide clusters. The resulting dataset was a 3,990 x 32,336 matrix with each row being an individual sample, and each column an LC-MS cluster. All remaining missing values were then imputed to a value zero. A separate metadata matrix (3990 x 4) which contained information on the technical conditions in the LC-MS run for each sample was compiled. These metadata were 1/ LC separation column – Categorical variable with 4 levels, 2/ Mass Calibration – Categorical variable with 6 levels, and 3/ Source heated capillary – categorical variable with 2 levels. A total of 3,090 samples had complete data (LC-MS spectra and corresponding metadata). This complete dataset was then analysed using ASCA in MatLab v.R2017b (Mathworks, Natick, WA, USA) utilising the PLS Toolbox v. 8.5.2 (Eigenvector Research Inc., Manson, WA, USA) to see which, if any, of the fixed experimental effects had a significant impact on the LC-MS cluster data. The statistical significance of the impact of each fixed experimental effect was estimated by calculating a p-value from permutation testing with 100 iterations.

The impact of experimental factors with a significant effect on LC-MS cluster data was then accounted for by correcting the data using multiple linear regression in R (44) as described in (45). The linear model was fitted as follows:

Y ijkl = u + Columni + MassCalj + Capk + eijkl

Where y is the signal intensity of a given cluster, u is the overall mean, Column is the i^th^ LC column (4 levels), MassCal is the j^th^ Mass calibration (6 levels), Cap is k^th^ Source heated capillary (2 levels), and eijkl is the random error term. The “corrected data” was a matrix of the residuals of the above model, which was run iteratively for each of the 32,336 peptide clusters. PCA plots were produced using R (44) and the gg2plot package.

### 2.7. Protein identification

The 63 RAW LC-MS2 files were processed in the Refiner MS module of Genedata Expressionist® 13.0 using a stepwise workflow similar the one described for LC-MS1 data, with the exception of additional activities pertaining to protein database search(Supplementary Figure S2A-C).

RAW files were searched using Mascot program (version: 2.6.1, Matrix Science Ltd, London, UK) within Genedata Refiner. The wheat database searched was retrieved from three independent sources. The first source was UniProtKB (https://www.uniprot.org/uniprot/?query=triticum%20aestivum&fil=organism%3A%22Tritic um+aestivum+%28Wheat%29+%5B4565%5D%22&sort=score) with 142,969 *T. aestivum* protein sequences (accessed on 26 February 2020, (41)). The second source was the EnsemblPlants repository hosting the *T. aestivum* genome initially sequenced by the International Wheat Genome Sequencing Consortium (IWGSC (9)) and containing 143,241 Traes AA sequences (http://ftp.ensemblgenomes.org/pub/plants/release-52/fasta/triticum_aestivum/pep/). A contaminant database was also retrieved (common Repository of Adventitious Proteins (cRAP); ftp://ftp.thegpm.org/fasta/cRAP). All the FASTA files were combined and redundant sequences removed by following the GalaxyP tutorial “Protein FASTA Database Handling” (https://training.galaxyproject.org/training-material/topics/proteomics/tutorials/database-handling/tutorial.html) (46, 47). The decoy database was created by reversing all the sequences and appending them using the GalaxyP tool “DecoyDatabase” (https://github.com/galaxyproteomics). Our Galaxy workflow is available in Supplementary File SF1. The final FASTA file was imported and indexed in Mascot. It contained 286,482 protein sequences and 1,647,476,761 AA residues; its longest sequence bore 5,359 residues.

All MS2 files were searched in one batch using Mascot Daemon (version 2.6.1,Matrix Science Ltd, London, UK) and the following parameters: MS/MS ions search, Mascot generic data format, ESI-TRAP instrument, trypsin enzyme, 9 maximum missed cleavages, carbamidomethyl (C) as fixed modification, guanidyl (K) and oxidation (M) as variable modifications, quantitation none, monoisotopic mass, 2+, 3+ and 4+ peptide charge, 10 ppm peptide tolerance, 0.5 Da MS/MS tolerance, and error tolerant search (Supplementary Figure S2D). Results were exported as .csv files into Excel.

The 32,336 peptide clusters from the corrected dataset produced by the LC-MS analyses were matched in R (44) (version 4.1.0-foss-2021a) to the 29,908 peptide clusters generated by the LC-MS/MS analyses using their respective RT, *m/z* and mass values with ±0.1 accuracy, and then linked to the Mascot identification results. The identification results of the peptide clusters whose RT shifted by more than 1 min were not included.

### 2.8. Statistical analyses of proteomics data

Out of the 4,061 grains samples processed in this work, 3,990 yielded reproducible LC-MS data for 32,336 peptide clusters. The full quantitative data is available from the MassIVE repository ((https://massive.ucsd.edu/ProteoSAFe/static/massive.jsp, MSV000090572). The corrected dataset with Mascot identification results were imported into Genedata Analyst (version 13, Genedata AG, Basel, Switzerland). LMA measurements were obtained on 3,773 (out of 3,990) wheat samples. Whilst LMA trait characterised the wheat samples, we also wanted to analyse it along with the peptides to facilitate biomarker discovery. To this end, we used the inverse function to normally distribute the LMA values (Inv(LMA)) and transposed them as a row to incorporate them into the LC-MS dataset under the label “Cluster_AAA” along with all the other 32,336 peptides, thus bringing the total number of clusters to 32,337. This “Cluster_AAA” row was used in the subsequence statistical analyses to isolate peptides displaying profiles similar to that of LMA.

#### 2.8.1. Principal Component Analysis (PCA)

A PCA was performed on the full dataset (3,990 samples x 32,336 peptides) in R using the prcomp() function of the stats package. The eigenvalues were plotted using the screeplot() function.

#### 2.8.2. Checking the distribution of LC-MS1 data

To redistribute data normally, the corrected dataset rows (peptides and Cluster_AAA) were z-transformed and plotted as a histogram in R. The hist() function was used to plot the corrected and z-transformed dataset as histograms in R (version 4.1.0-foss-2021a). One-sample Kolmogorov-Smirnov tests were applied to check the normality of the distribution of both corrected and z-transformed datasets using the ks.test() function and “pnorm” argument in R. All the subsequent statistical analyses were performed on the z-transformed dataset.

#### 2.8.3. Subsampling wheat samples to eliminate the bias towards low LMA values

LMA values spanned 0 to 8 u/g with the vast majority (95%) below 0.2 u/g (which corresponds to FN 300 s (18)); therefore, the LMA distribution was greatly skewed towards low LMA values. To eliminate this bias, a subset of wheat samples was selected as follows: all the samples bearing a LMA ≥ 0.17 were selected (467 samples in total) and an equivalent number of samples (467) with LMA < 0.17 were randomly selected among the 3,306 remaining wheat samples. This subset of 934 wheat samples was no longer skewed towards low LMA values and is referred as “unbiased samples” hereafter.

#### 2.8.4. Partial Least Squares (PLS) to subset LMA-responding peptides

In Genedata Analyst, a PLS 2-D plot was created using the 934 unbiased samples and all the 32,346 peptides resolved in this study. The parameters were: LMA as a response, 3 latent factors, 10% valid values, and row mean imputation. Both score and loading plots were exported along with the variable importance in projection (VIP) scores. The higher the score, the greater the contribution of the peptide to the PLS and the closer to LMA response. These VIP scores were used to select meaningful subsets of peptides for the subsequent statistical analyses.

#### 2.8.5. Univariate Partial Least Square (PLS) Regression to impute LMA missing values

The missing LMA values were predicted using a univariate PLS regression model in Genedata Analyst. First a model was developed using the 934 unbiased samples and 2,996 peptides with PLS high VIP scores (> 1.5). Second, among the 934 wheat samples, 179 were randomly chosen so that LMA evenly spanned 0 to 5 and those LMA values were erased. Several PLSR models were tested to accurately predict erased LMA values. The most accurate model applied the following parameters: LMA as a response, 20% valid values, and 20 latent factors. The model was then applied to the 217 missing LMA values against the 934 unbiased wheat samples.

#### 2.8.6. Self-Organising Maps (SOM) Clustering

In Genedata Analyst, a SOM was created using the 934 unbiased samples and 7,254 peptides with VIP scores above 1 (including Cluster_AAA) and the following parameters: 6 rows, 8 columns, positive correlation distance, 50 maximum iterations, and 10% valid values.

#### 2.8.7. K-Means

In Genedata Analyst, a k-means was performed using the 934 unbiased samples and 7,254 peptides with VIP scores above 1 (including Cluster_AAA) and the following parameters: k=20, positive correlation distance, mean centroid calculation, 10% valid values, and 50 maximum iterations.

#### 2.8.8. Divisive Hierarchical Clustering Analysis (HCA) and agglomerative HCA

An HCA was produced in Genedata Analyst a divisive HCA using the 934 unbiased samples and 7,254 peptides with VIP scores above 1 (including Cluster_AAA) and the following parameters: clustering peptides, tree with tile plot, positive correlation distance, Ward linkage, 10% valid values, k-means cluster profile, and split by size. The outcome of this analysis enabled us to sort the peptides based on their accumulation patterns in wheat samples.

Still in Genedata Analyst, we also performed an agglomerative HCA using the all the 934 unbiased samples and 532 LMA-related biomarkers (including Cluster_AAA) and the following parameters: clustering samples, tree, positive correlation distance, Ward linkage, 50% valid values. The outcome of this analysis allowed us to sort the grain samples according to their LC-MS molecular similarity which was then exploited in a heat map.

#### 2.8.9. Correlation

An annotation correlation was performed in Genedata Analyst using the full dataset including Cluster_AAA (3,990 samples x 32,337 peptides) against standardised LMA values. This produced R squared (R2) values.

#### 2.8.10. Simple linear mixed regression

The full dataset including Cluster_AAA (3,990 samples x 32,337 peptides) was used to run a linear regression in Genedata Analyst with one explanatoryvariable usingthe followingmodel: y = Inv(LMA) + ε, in which Inv(LMA) is the normal inverse function of LMA measurements. The false discovery rates were computed according to the Benjamini-Hochberg estimates as q-values.

#### 2.8.11. Peptide expression profiles along 2 or 8 LMA bins

Our data matrix of 3,990 columns by 32,337 rows contained 129,024,630 quantities which posed representation challenges. We adopted a data reduction strategy involving binning the samples into 8 or 2 arbitrary bins based on their LMA values in order to produce simpler more legible graphs for individual peptide profiling.

In the first instance, we sorted all 3,990 wheat samples based on an increasing order of LMA values, and then split them into 8 arbitrary bins of 499 samples each. The last bin (0.17132<LMA<7.95442) contained all the 266 unsound grains (LMA > 0.2).

In the second instance and using the 934 unbiased wheat samples, we created 2 bins based on LMA value threshold of 0.17. The bin containing 467 samples with LMA < 0.17 only comprised sound grains. All the 266 unsound grains (LMA > 0.2) were comprised in the bin containing 467 samples with LMA ≥ 0.17.

The peptide quantities were then averaged per bin to produce mean expression profiles along 2 or 8 bins.

#### 2.8.12. T test with effect size and volcano plot

Using the unbiased biomarker dataset (934 samples x 532 peptides including Cluster_AAA), a t test was performed with the LMA threshold of 0.17 as a factor (see sections 2.8.3 and 2.9.1) and the following parameters: boostrap with 10 repeats and balanced permutations, effect size based on group means, and 90% valid values. This produced a volcano plot.

### 2.9. Proteomics data mining

The LC-MS2 experiments followed by Mascot search produced identification results for 5,414 peptide clusters which matched 8,044 protein accessions. These identification results were mined using the databases and tools described below. Resulting outputs were consigned to Supplementary Tables S3.

#### 2.9.1. UniProt database and Gene Ontology (GO)

The list of 8,044 UniProt accessions identified in this study was uploaded in the Retrieve/ID mapping tool of UniProt (https://www.uniprot.org/uploadlists/ accessed on May 2022) (48) to retrieve protein descriptions, FASTA sequences, GO terms, and TRAES accession IDs. Out of the 8,044 UniProt accessions, 5,960 UniProt accessions corresponded to 6,622 TRAES accessions. TRAES accessions were needed to interrogate ShinyGO and BreadwheatCyc databases (described below).

#### 2.9.2. Kyoto Encyclopedia of Genes and Genomes (KEGG) database and pathway maps

The 8,044 FASTA sequences were uploaded into the Assign KO tool (https://www.kegg.jp/kegg/mapper/assign_ko.html accessed on May 2022) (49) by specifying the Poaceae family to retrieve KEGG ORTHOLOGY (KO) identifiers. KO identifiers were then mapped using the KEGG Mapper Reconstruct tool (https://www.genome.jp/kegg/mapper/reconstruct.html accessed on May 2022) to list pathways, Brites and modules involving identified proteins.

#### 2.9.3. ShinyGO, Functional Category enrichment and chromosomal positions

The list of 6,622 TRAES accessions was uploaded into ShinyGO (http://bioinf ormatics.sdstate.edu/go/) (50) to generate Functional Category enrichments, dot plots, tree, networks, as well as retrieve chromosomal positions. Positions were obtained for 4,571 TRAES accessions which were used in Circos plots (detailed below).

#### 2.9.4. Pathway Tools, BreadwheatCyc and perturbed pathways

The list of 6,622 TRAES accessions along with quantitative data along 8 bins was uploaded into the Pathway Tools software (51) and run online via the BreadwheatCyc database (https://pmn.plantcyc.org/organism-summary?object=BREADWHEAT accessed on June 2022) via the Plant Metabolic Network server (52) using the Omics Dashboard (https://pmn.plantcyc.org/dashboard/dashboard-intro.shtml accessed on June 2022), the Cellular Overview tools (https://pmn.plantcyc.org/overviewsWeb/celOv.shtml?orgid=BREADWHEAT accessed on June 2022) to generate Pathway Perturbation Scores (PPS).

The Chrome extension Veed.io was used to create a film capturing the Cellular Overview animation.

#### 2.9.5. Circos and chromosomal position

The 4,571 TRAES accessions whose chromosomal positions were known from ShinyGO were charted along a Circos plot invented by Krzywinski and colleagues (53) and recently wrapped in the Galaxy platform by Rasche and colleagues (https://usegalaxy.eu/?tool_id=circos) (46, 54, 55). The details of the various layers are indicated in the figure’s legend.

#### 2.9.6. R and Power BI Desktop

Most identified peptide matched several UniProt accessions which corresponded to several TRAES IDs, and GO terms. This produced wide tables. In R (version 4.1.0-foss-2021a) (44), wide tables were converted to longtables usingthe pivot_longer() function from tidyr package. Long tables were merged using the merge() function of the R base package using peptide Cluster IDs as unique references.

Wheat sample metadata, peptide metadata and quantitative dataset and identities for the biomarkers were imported into Microsoft Power BI Desktop (Version: 2.106.883.0 64-bit June 2022) and linked via the Clusters names to produce dashboards using multiple visuals (word clouds, tree maps, histograms, scatterplots, waterfall plots, pie charts, violin plots and ribbon charts).

## 3. Results and Discussion

### 3.1. Resources for scientific studies on wheat

#### 3.1.1. Wheat resources

A total 858 wheat genotypes, sourced from all over the world, grown over 8 years since 2012 and stored in optimal conditions amounting to 4,061 grain samples were analysed in this work (Supplementary Table S1). Because LMA measurements occurred simultaneously to the proteomics analyses in 2019, we did not consider storage time for the statistics. We also did not statistically test for varietal differences which was outside the focus of this study.

#### 3.1.2. High-throughput proteomics workflow to efficiently process and analyse thousands of samples

We have developed a high-throughput proteomics LC-MS method (41) that was applied to 4,061 wheatgrain samples followingthe workflow described in Figure 1. The technical aspects pertaining to sample preparation/tracking and data acquisition steps that ensured a high-throughput workflow are available in Supplementary File SF1. Overall, the LC-MS continuous run lasted for 143 days (20.4 weeks or 4.5 months) and included regular system maintenance (mass calibration, source cleaning, HPLC column swapping). A total of 4,370 RAW files were acquired. A Gantt chart illustrates the timeline of the workflow steps along with data accumulation (Figure 2).

**Figure 2.**
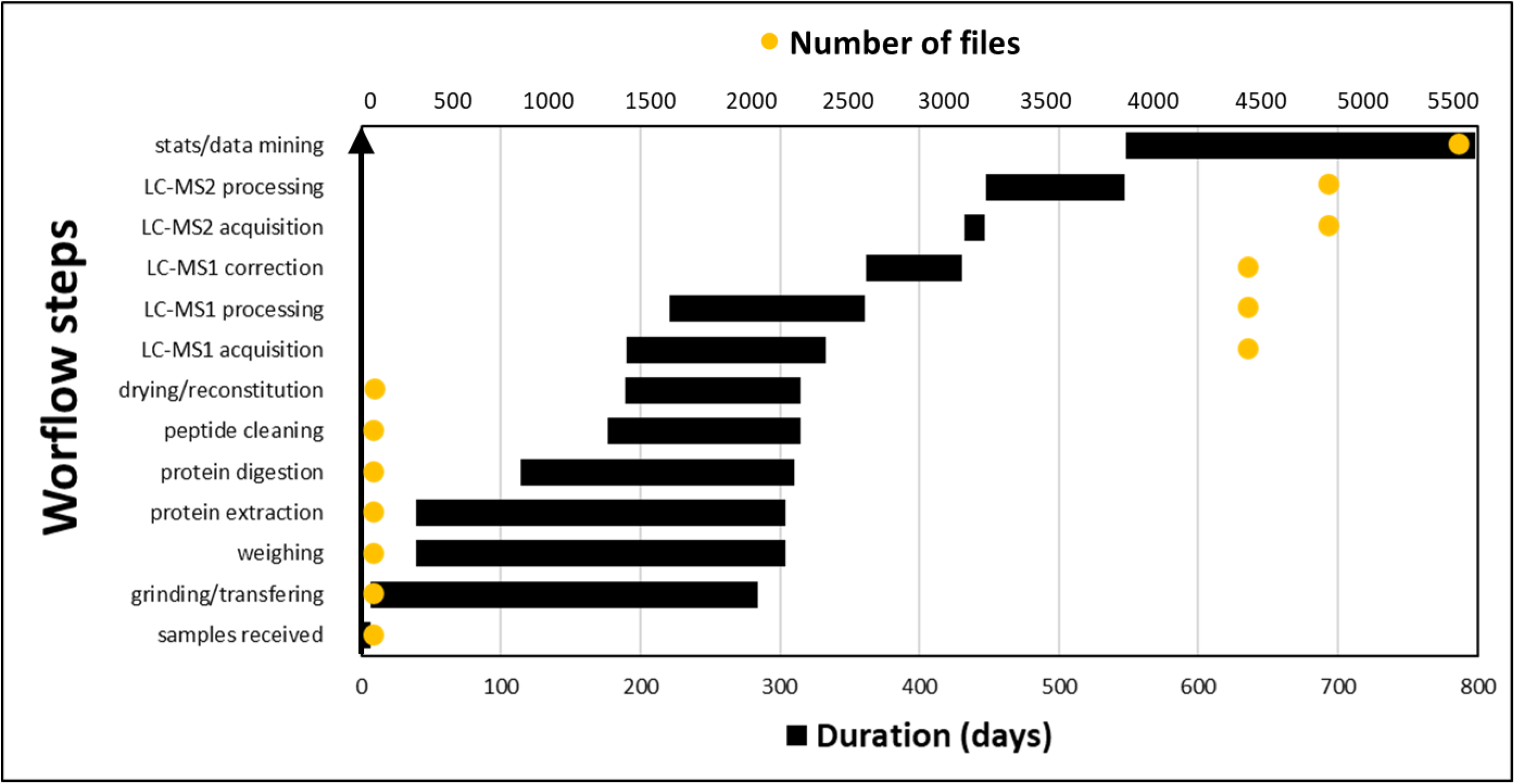
Gantt chart capturing the timeline for each step of the proteomics workflow and file accumulation.

The wet experiment bottlenecks were resolved where possible as explained in (41). Most time was spent grinding, transferring, weighing and extracting the samples as there was no option to greatly up-scale those steps (Figure 2). The workflow became much faster when 96-well plates were introduced (from digestion step onward) allowing for high throughput multipipetting and multidispensing activities, as well as minimising the footprint of sample freezer storage. Although steps were sequential, they could overlap with two experimenters operating in a staggered fashion from one lab workstation to the next.

LC-MS1 acquisition started when enough plates were ready to ensure continuous instrument run while samples processing was still happening. Data acquisition was completed 18 days after the last wheat sample was fully processed, demonstrating minimum time loss (Figure 2). The Genedata Refiner workflow used to process LC-MS1 data was previously optimised (41) (Supplementary Figure S1 described in section 3.1.2); its first step was applied to batches of ∼200 LC-MS1 files. The time limiting factor was the server computing ability.

Overall, all 4,061 wheat samples were processed and analysed (from receiving the samples to processing the LC-MS1 data) in 334 days (∼11 months). Purchasing all required consumable ahead, keeping track of the samples, good logistics by setting up working stations for each wet lab step, as well as overlapping activities across experimenters guaranteed efficient time management. Stowing samples in the freezer in-between steps allowed to safely interrupt the sample preparation procedure to accommodate equipment/experimenter downtime without compromising the quality of the samples processed so far.

The subsequent steps had to follow one another. LC-MS2 acquisition necessitated LC-MS1 data processing to be finished in order to produce parent mass lists and consequently had to be performed post-hoc. Whilst LC-MS2 acquisition was rapid (2 weeks), its processing took longer (3 months) because it required another Genedata Refiner workflow (Supplementary Figure S2 described in section 3.1.3), a more recent non redundant database with decoy sequences, testing several Mascot parameters (data not shown), and linking LC-MS2 clusters to LC-MS1 clusters (data not shown).

The final bottleneck in the workflow pertained to statistical analyses and data mining (8 months) which necessitated trying different statistical methods with multiple trial and error stages working out optimal parameters, testing and using different data mining tools which required training and a lot of strategising on how best to present big data. Running such large datasets proved computationally taxing, necessitated extensive dwell times; it often ran out of memory and triggered server crashes.

One way to increase the throughput and therefore shrink the timeline would be to use an automated sample preparation station. A robot (Bravo Automated Liquid Handling Platform from Agilent) was used to automate peptide clean-up and phosphopeptide enrichment from wheat and maize vegetative samples (56). We could notfind any other high throughputmethod in wheat or cereals.

#### 3.1.2. LC-MS1 quantitative data processing, normalisation, correction and standardisation to remove technical biases

The Genedata Refiner workflow described in (41) was applied to 4,147 LC-MS1 files (4,061 wheat + 86 QCs; Supplementary Figure S1). Step 1 covered noise subtraction nodes that could be run on individual data file. It was performed throughout LC-MS1 acquisition activity on weekly batches (∼230 files) to optimise server dwell time. Step 1 helped assess data reproducibility and non-reproducible files (71 samples) were omitted from the remainder of the processing, leaving 3,990 wheat and 86 QC data files. Step 2 encapsulated all alignment, peak detection and quantitation, as well as isotope clustering and singleton filtering activities. This step had to be performed on all 4,076 reproducible data files simultaneously and therefore could only be attempted when the LC-MS1 run was finalised. The experiment metadata captured in Excel was associated to the quantitative data and exported to Genedata Analyst for data normalisation purpose.

The data was normalised as described in (41) following three steps: using flour weights, IS cluster and QC replicates along with LC-MS injection order (Figure 3).

**Figure 3:**
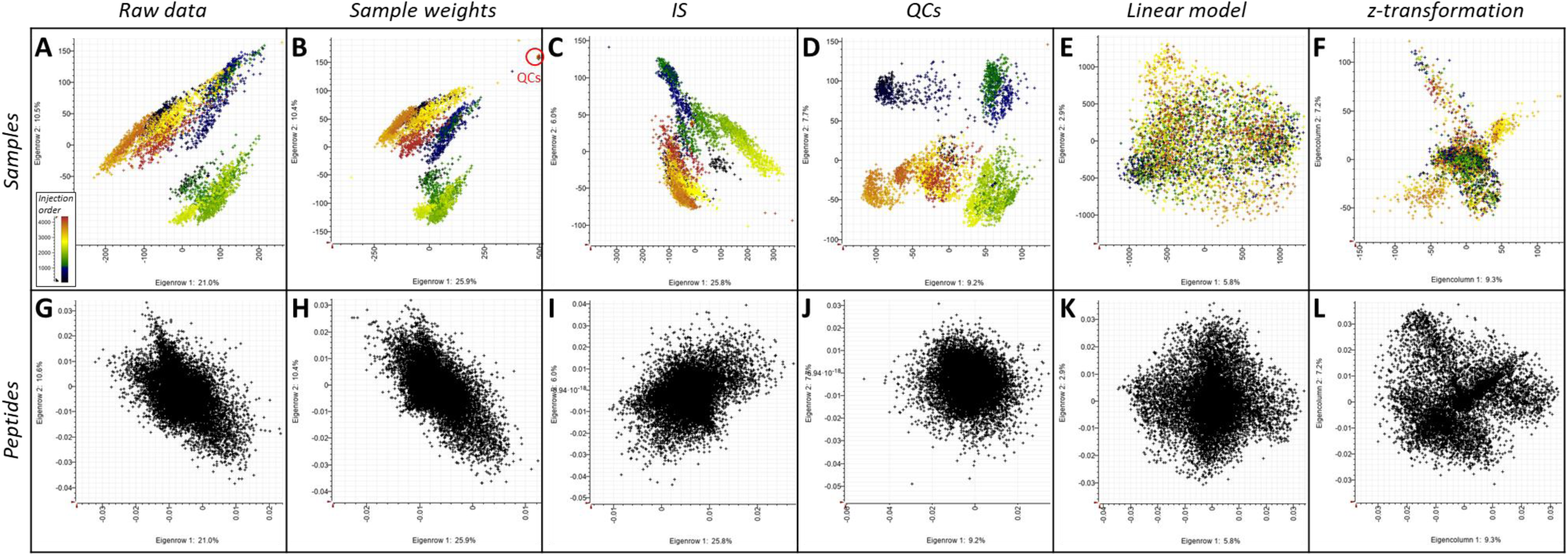
Normalisation, correction and standardisation of the raw data visualised using PCA projection plots of the samples (A-F) and loading plots of the peptides (F-K). Samples are coloured accordingly to LC-MS injection order from blue-green to yellow-orange-red. (A,G) PC1 vs. PC2 plot based on unnormalised LC-MS1 quantitative data; (B,H) PC1 vs. PC2 plot based on data from panels A,G normalised using the sample weights; QCs are all condensed in a tight group (C,I) PC1 vs. PC2 plot based on data from panels B,H normalised using the IS cluster; (D,J) PC1 vs. PC2 plot based using data from panels C,I normalised using the injection order and the ‘intensity drift’ algorithm; (E,K) PC1 vs. PC2 plot using normalised data from panels D,J corrected using a linear model and keeping the residuals; (F,L) PC1 vs. PC2 plot using corrected data from panels E,L and z-transformed per row (peptides).

Raw data displayed a clear sample grouping based on injection order during the LC-MS1 run (Figure 3A) and mirrored the instrument maintenance events (mass calibration, etc…). Two large groups appeared that could not be explained by any experimental steps. Normalising using flour weight accuracy of 1% helped creating tighter wheat sample groups with four outliers, and isolated QCs (Figure 3B). The two larger groups of samples were less distinct. This first normalisation step did not significantly impact the peptide distribution as can be seen on the PCA loading plots (Figure S3G,H). Normalisingagainstthe IS shifted the sample groups around but did not combine or homogenise them (Figure 3C). The two larger sample groups observed in panels A-B became indistinguishable in panel C. This normalisation step also affected peptide distribution assuming a more oval shape on the loading plot (Figure S3I). The final normalisation step further scattered the samples more widely across the PCA plot and accentuated the technical variation gradually expanding overtime during the instrument run (Figure 3D). Yet at the peptide level, this last normalisation activity further shrunk the grouping assuming a more circular distribution with less outliers (Figure S3J). The benefits of normalisation were discussed before (41) with respect to precise sample weights mandated in metabolomics (57), spiking IS post-digestion to alleviate for sample to sample variations (58, 59), and QCs to account for batch differences over time and minimise cross run effects (59–61). In their ground-breaking study to assess and ameliorate the reproducibility of large-scale proteomics experiments, Poulos and colleagues have highlighted the decrease over time in mass analyser sensitivity in-between cleaning events and how technical replicates, such as QCs, help remove unwanted variation (62). Despite all the normalisation steps applied to our data, not all technical biases could be removed, thus necessitating further data correction.

The fully normalised dataset of 3,990 wheat samples and 32,336 reproducible peptides was exported as a CSV file and imported into R to run a linear model fitting the technical factors that bore the greatest variance and were associated with LC-MS maintenance. The experimental variation was successfully eradicated as illustrated by PCA (Figure 3E,K). The results showed that while instrument mass calibration had a much bigger effect, all three technical factors had a significant effect (P < 0.05 based on permutation testing with 100 iterations) on the spectraldata (data notshown). This correctionmethod was initially developed in a metabolomics study to account for uncontrollable environmental effects (45). Quantitative geneticists routinely exploit linear models to measure the influence of systematic environmental effects (fixed effects) which impact phenotypic variation and unscramble genetic from non-genetic factors (63). To our knowledge, this is the first time such correction method was applied to proteomics data.

The final data transformation step involved a z-transformation (scaling and centring) to level out extreme quantities and facilitate the comparison and clustering of peptide profiles during statistical analyses. Finding linear combinations of predictors based on how much variation they explain is achieved by centring to a mean of 0 and scaling to a standard deviation of 1 (64). Such mathematical transformation is common practice in post-genomics expression studies, and MS-proteomics is no exception (65, 66). In our study, z-transformation radically modified the data froman homogenous plot to defined groups stretching in four main directions (Figure 3F,L), which could not be attributed to any of our metadata. Peptide quantities that originally ranged from 0 to 1 x 10^7^ ultimately spanned a mere -22 to 63 scale.

#### 3.1.3. A non-redundant wheat database to annotate LC-MS2 results and identify post-translational modifications (PTMs)

A *T. aestivum* database was created by combining all the protein sequences publicly available from UniProt and IWGSC EnsemblPlants repositories. Because protein annotations from the IWGSC (hereafter called TRAES sequences) referred to UniProt, we used the latter as a template to eliminate AA sequence redundancy. This completely removed all IWGSC TRAES sequences (data not shown) from our merged data file indicating they were all included in the UniProt repository. The database was reversed to create a decoy database which was then concatenated to the latter. This way, not only a single file has to be interrogated in Mascot system, but also false positives are only recorded when a match from the decoy sequences exceeds any match from the target sequences (67, 68). All LC-MS2 files were processed in Genedata Refiner and searched using the Mascot algorithm with an error tolerant search to maximise PTM discovery. The search outputs were merged into a single file and exported to Excel (Supplementary Figure S2E).

Our strategy to quickly identify as many peptides as possible was to multiply the number of data-dependent LC-MS2 methods rather than multiplying the number of samples analysed. We thus pooled 10% of the wheatsamples randomly choseninto one tube and subjected this pooled sample to 11 methods (passes) with replicates, varied ITMS parameters and 10 unique parent lists of 2,000 ions each. Each method had a drastic impact of the selection of precursor ion, with some areas being thoroughly samples whilst others were ignored (Supplementary Figure S3).

A total of 63 LC-MS2 files were thus obtained. The LC-MS2 methods varied in their efficiencies, identifying as few as 104 peptides (pass 7) up to 11,662 peptides (pass 8), irrespective of the number of MS2 events (Supplementary Figure S4).

Passes 8-10 yielded by far the largest identity counts across all 10 parent lists, even though they did not feature the highest MS2 event counts (Supplementary Figure S4). Key MS parameters to maximise peptide identifications were the inclusion of the parent lists into the data-dependent settings (passes 8-11) albeit not the at the global level (pass 7) as well as allowing for wider mass tolerance window during precursor selection. The widest tolerance (2 m/z) achieved the greatest counts (pass 8, Supplementary Figure S4). Overall, a total of 315,934 peptides were identified, comprising only 6,550 unique peptides which matched 10,437 unique wheat proteins, 277 decoy accessions, and 3 contaminant proteins. The huge peptide redundancy was explained by the fact that a single pooled sample (from 400 individual samples) was repeatedly analysed using various LC-MS2 methods. Pooling digests erased sample-to-sample variation. More protein identities could have been realised with a diverse sample set subject to all the methods developed here but that would have extended the data acquisition, analysis and mining by many more months. A greater proteome coverage was achieved in our method optimisation study yielding 13,165 identified peptides even though far less samples were analysed because two extraction protocols and three orthogonal digestions were applied which produced more diverse LC-MS profiles (41). An array of strategies can be employed to increase the proteome coverage of plant seeds, including depletion and pre - fractionation strategies as well as exploring different organs, developmental stages, and cell cultures (69, 70). However, these additional experimental steps are time-consuming, labour-intensive, as well as costly thus unsuitable for large-scale high-throughput experiments like ours. Our strategy was first to rapidly and reproducibly quantify digested peptides from thousands of wheat samples using a label-free LC-MS approach and apply robust statistical analyses to detect potential trait-related biomarkers, and second to quickly identify as many peptides as possible using LC-MS2. Large-scale proteomics studies have been applied to human (71); to our knowledge, this is the largest plant proteomics study carried out to date.

In this study, we opted for an error-tolerant search which accrued a plethora of modifications (Supplementary Table S2). A total of 21,486 carbamidomethylations of Cys residues were identified as fixed modifications. This was expected to occur during our denaturing protein extraction procedure. The most prevalent dynamic modifications were non-specific cleavages (5, 480), followed by N-terminal ammonia losses (907), and conversion from N-terminal Gln to pyroGlu (815). Duringthe digestion process involvingtrypsin, proteomics studies haveoften reported the formation of semi-tryptic and non-specific peptides besides cleavages after Arg or Lys residues (72). Therefore, some of our non-specific peptides could have resulted from the digestion step, but we cannot rule out that non-tryptic peptides were naturally present on our stored grains, resulting from residual enzymatic activities. Ammonia losses are neutral losses commonly triggered by CID upon creating b and y ions, and can be detected by high resolution mass analysers such as FTMS instruments (73). C-terminal Arg or Lys of tryptic peptides often leads to abundant y ions with ammonia loss (74) and as well as b ions specific enough to detect the presence of Gln, Asn, His, Lys, and Arg residues (73). PyroGlu formation is a common cyclization side reaction of Glu and/or Gln residues in peptides and proteins that occurs when those residue are located at the N-terminus and under slightly acidic conditions (75), such as our experimental conditions therefore this PTM could also be a process artifact. Other frequent PTMs in our study were N-terminal ethylation (265 occurences), deamidation (147 occurrences), guanidylation (141 occurrences), the latter of which could have been triggered during protein resuspension in Guanidine-HCl solution as discussed in (41), as well as oxidation of Met (100 occurrences) (Supplementary Table S2). Numerous PTMs have been identified in plants (69) and cereals in particular (76), including barley (77), and wheat (2, 78, 79). Deamidations of glutamine residues in glutenins have been reported (5), along with C-terminal loss of tyrosine potentially facilitating protein sorting during seed maturation (2). Starch content and starch-related proteins are prominent in wheat grain; PTMs involved in starch quality have been reviewed (80). Our study lists numerous potential PTMs; this warrants more experiments to validate them and decipher their role in LMA response. Future proteomics experiments should endeavour to explore the relationship between structure and functionality of gluten proteoforms arising from key PTMs in response to LMA phenotype.

#### 3.1.4. Linking LC-MS1 and LC-MS2 data to annotate quantities with identities

LC-MS1 files resolved 32,336 reproducible clusters which had to be matched to 29,908 clusters from LC-MS2 data files. Using tolerances of 20 ppm for m/z and mass and 1 min for retention times, 16,874 (52%) peptide clusters were matched across both datasets, of which 5,414 bore peptide identification results. These identified peptides matched 8,044 *T. aestivum* protein accessions. Our experimental results are summarised in Table 1; number of identified peptide numbers aside, they compared well with our previous findings during method optimisation (41).

**Table 1:**
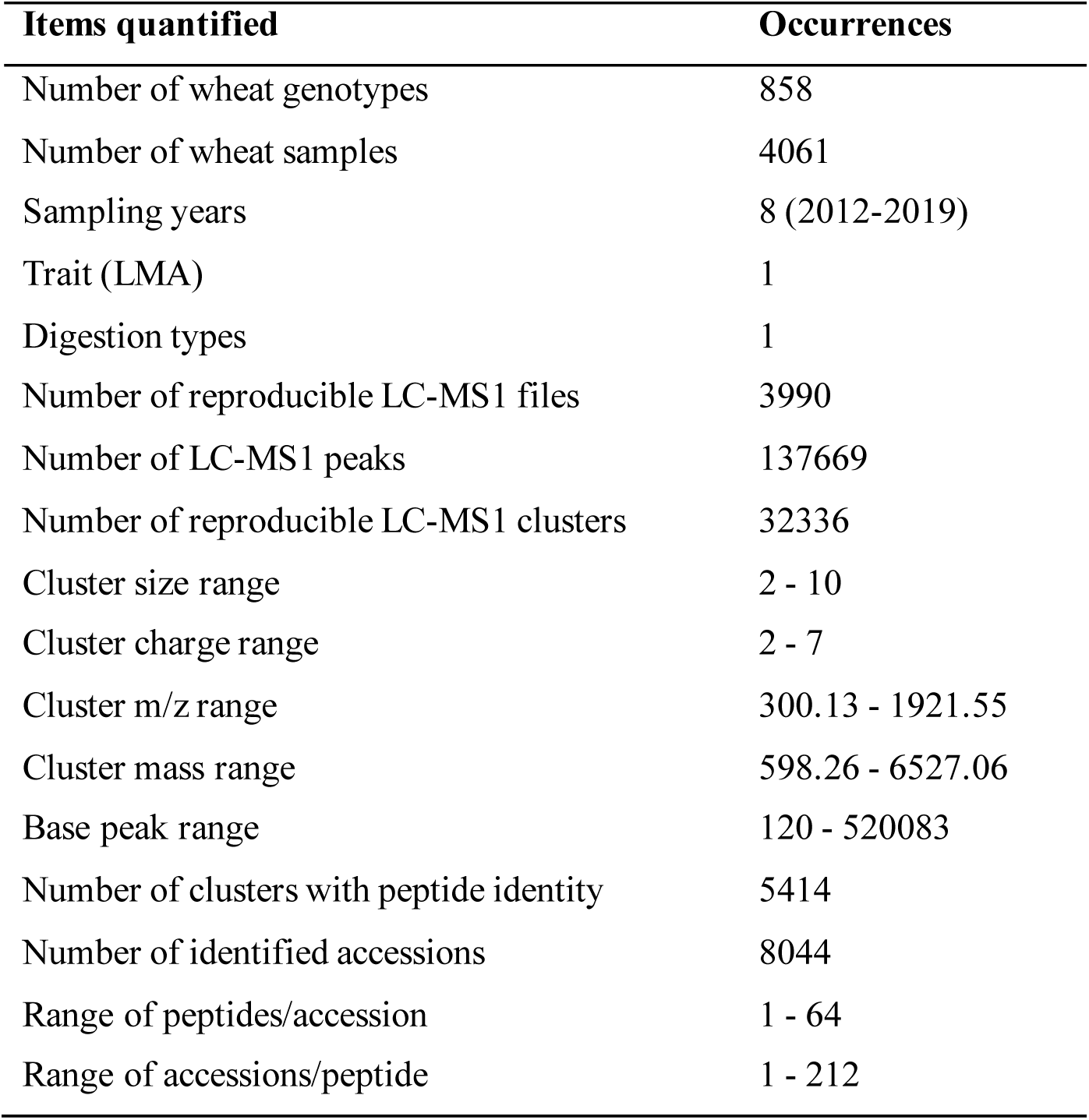
Experiment summary.

Our strategy was to consider all 8,044 protein hits identified fromthe 5,414 sequencedpeptides irrespective of their homology. We thus turned the 5,414 x 212 wide table into a long table containing 32,347 rows of peptides and replicated the quantitative data accordingly for statistical analysis purposes. The list of all identities is captured in Supplementary Table S3. Up to 64 unique peptides matched a particular protein with an average of 4 peptides per hit (Supplementary Figure S5A-B).

A given peptide matched to up to 212 protein accessions with an average of 6 hits per peptide (Cluster_29452, VLQQLNPCK, Supplementary Figure S5C-D). This mirrored the high frequency of homoeologous proteins in the hexaploid wheat samples expressed from three similar subgenomes, A, B and D (81). Another compounding factor was that wheat protein accessions were created from genomic sequences, resulting in multiple accessions bearing identical sequences but arising from different gene accessions (2). This created on one hand protein accessions labelled as “fragments” despite havinga complete codingregion and, on the other hand, other accessions lacking this tag despite having an incomplete coding region (Supplementary Table S3). Finally, the vast number of PTMs identified here also contributed to boosting hits against a particular peptide AA sequence. The most dominant wheat grain proteins are storage proteins such as gliadins and glutenins, which featured prominently in our proteome (Supplementary Figure S5E, Supplementary Table S3), despite the fact that their low Lys/Arg content makes them less prone to trypsin digestion (2). Other major proteins comprised histones, beta-D-glucosidases, and ubiquitin. This list of identified proteins compared well with our previous methodological work (41).Other recent studies on mature wheat seed proteome using gel-based or gel-free technologies also published comparable list of identities (82–84).

### 3.2. Application to a wheat industry problem: Late maturity alpha-amylase (LMA)

Wheatmarketingfor milling grades dictates that below a certain FN value, grains are no longer suitable for human diet and must then be discounted causing significant financial losses to the suppliers (17). FN assesses starch degradation resulting from LMA activity which can be assayed in flour samples using the Ceralpha method (37) for instance. Even though LMA trait is a genetic defect, it persists in wheat germplasm implying that it is either not selected against or alternatively imparts unbeknown beneficial attributes to LMA-prone varieties (24). By unravelling the genetic, biochemical, and physiological mechanisms that lead to LMA expression, scientists strive to understand and eliminate LMA from wheat breeding programs (39). Surprisingly, post-genomics is not one of the strategies adopted by researchers to close the biological knowledge gap, with only one transcriptomics study registered so far (22). Our study constitutes the first proteomics experiment performed to decipher the mechanisms involved. Machine learning was performed on the complete dataset to distinguish LMA-susceptible from non-susceptible wheat genotypes without success (data not shown). Results from statistics and data mining are described and discussed below.

#### 3.2.1. Getting the quantitative data ready for statistical analyses

##### 3.2.1.1. Assessing the normality of LC-MS1 datasets

To assess whether our LC-MS1 datasets following the correction and z-transformation steps was normally distributed, we plotted the data as histogram and boxplot. We further performed the nonparametric one-sample Kolmogorov-Smirnov (K-S) test (85) well suited to analysing big data (86). Both histogram and boxplot of the corrected data were asymmetrical with most values being on the low range (Supplementary Figure 6A-B), which revealed that this dataset was not normally distributed. This was confirmed by the high K-S statistics (D) of 0.41 and a very low p-value (< 2.2 e^16^).

Using the z-transformed data, the histogram and boxplot were more symmetrical (Supplementary Figure 6C-D). Whilst the K-S statistics (D) was reduced to 0.27, it was still too high to conclude to normality. Even though we did not achieve a gaussian distribution by standardising the data, we managed to make it more even which improved statistical analyses for biomarker discovery.

3.2.1.2. PLS of unbiased samples to select a meaningful set of LMA-responsive peptides Analysing such a large dataset (3,990 columns x 32,337 rows) was computationally taxing, necessitating extensive dwell times to finalise statistical analyses, and often triggering Genedata sever crashes due to out-of-memory failures despite recent upgrades. Consequently, we devised a strategy to select a subset of relevant peptides via the supervised cluster method PLS. Using the 934 unbiased samples and all 32,337 peptides (including Cluster_AAA), we executed a PLS analysis with LMA trait as a response. The score plot of the first two components showed that the PLS successfully pulled out the grain samples exhibiting high LMA activities (Supplementary Figure S7A).

The corresponding loading plot allowed us to categorise peptides according to th eir contribution to the PLS model via their Variable Importance in Projection (VIP) scores. The most-contributing peptides (i.e. exhibiting the highest VIP score) were located in the plot area equivalent to that of high LMA samples (Supplementary Figure S7B).

VIP scores indicated the importance of each variable (peptide) in the projection used in the PLS model. Peptide VIP scores were calculated as weighted sums of the squared correlations between the PLS components and the original peptides; weights were inferred from the percentage variation explained by the PLS component in the model (87). VIP scores greater than 0.5, 1.0, and 1.5 segregated 14,440 (45%), 7,252 (22%), and 2,996 (9%) peptides, respectively. By setting up three VIP score thresholds of increasing stringency, we thus created three subsets of peptides of decreasing sizes that could be used in more computationally demanding processes.

##### 3.2.1.3. Wheat subsampling to create an unbiased dataset and transforming LMA trait profile to achieve normal distribution

In the 3,990 reproducible wheat samples, 3,773 featured LMA measurements that ranged from 0.04 to 7.95 u/g (Supplementary Table S1), albeit mostly on the low scale with 88% of the values recording less than 0.2 u/g (Figure 4A), which corresponds to the receival threshold of FN 300 s (18, 23).

**Figure 4:**
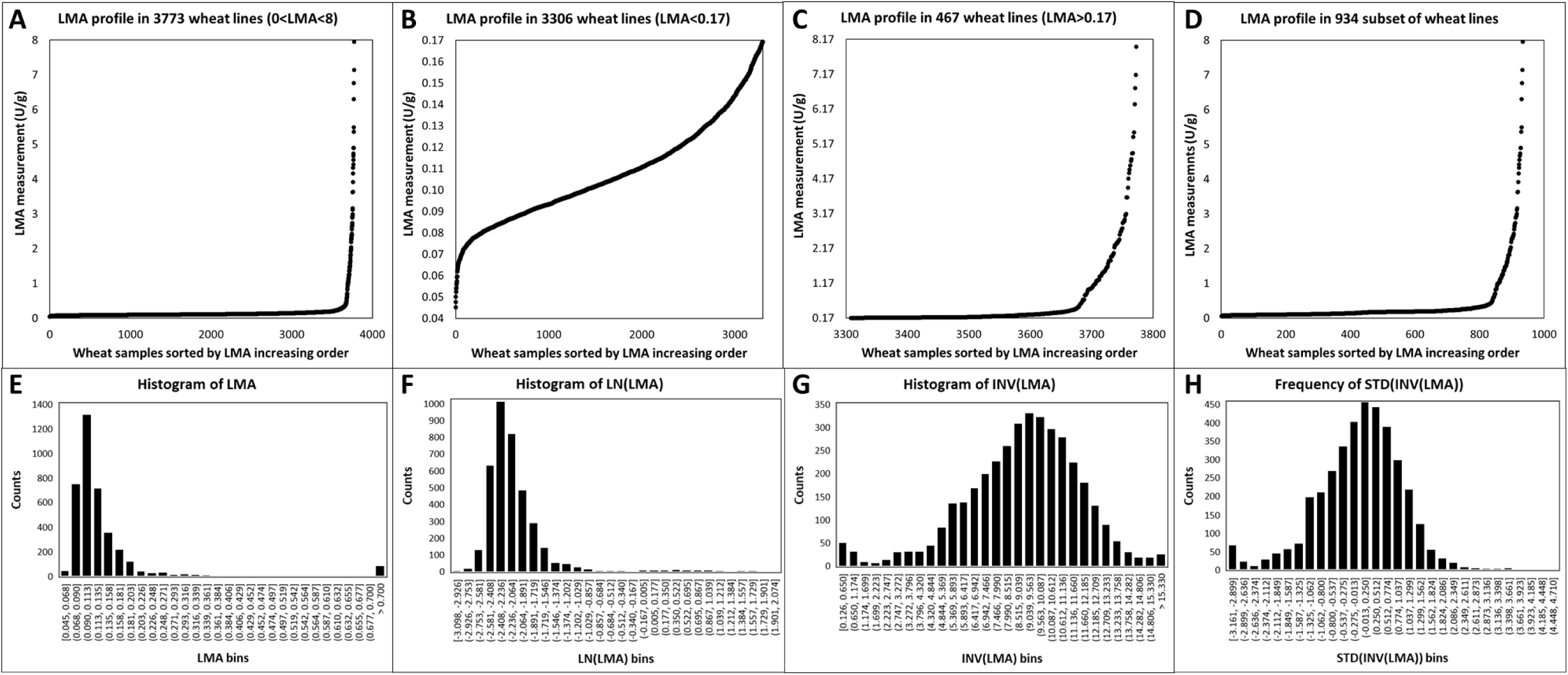
Profiles of LMA measurements for each wheat sample sorted by increasing values illustrated as scatterplots (A-D) and histograms (E-H). (A) Scatterplot of LMA values assayed in 3,773 wheat samples; (B) Scatterplot of LMA values less than 0.17 U/g in 3,306 wheat samples; (C) Scatterplot of LMA values equal to or greater than 0.17 U/g in 467 wheat samples; (D) Scatterplot of LMA values in unbiased set containing 934 samples (see Section 2.8.2 for explanation); (E) Histogram of LMA values assayed in 3,773 wheat samples along 30 bins; (F) Histogram of LMA values assayed in 3773 wheat samples and transformed using a natural logarithm (LN) function along 30 bins; (G) Histogram of LMA values assayed in 3,773 wheatsamples and transformedusingan inverse function(1/LMA=INV(LMA)) along 30 bins; (H) Histogram of LMA values assayed in 3,773 wheat samples and transformed standardising the inversion function (STD(INV(LMA))) from panel G along 30 bins.

Our range far exceeded those reported earlier, spanning either 0.08 to 0.67 u/g across 33 spring wheat cultivars grown across 18 field sites (88), 0.023 to 1.417 u/g over 39 varieties grown under controlled and triggering LMA-conditions (23), or 0.002 to 1.977 u/g among 196 genotypes from three experimental locations (19). We chose a threshold of 0.17 as a tipping point to delineate between grain samples displaying either low (3,306 samples) or high (467 samples) alpha-amylase activity. The LMA profiles below and above this arbitrary value showed a slow gradual increase of enzyme activity up to 3.2 units where datapoints became more scattered (Figure 4B-C). Because the LMA distribution was significantly skewed towards low values and to restore balance to the trait profile, we retained all the wheat samples with an LMA above 0.17 (467 samples) and randomly selected 467 samples (out of 3,306) for which LMA fell below this threshold. The LMA profile of this unbiased subset of 934 samples (Figure 4D) was very similar to the complete distribution (Figure 4A).

When LMA measurements were plotted as a histogram, it confirmed the skewness towards low activities and highlighted that most values fell between 0.068 and 0.203 u/g (Figure 4E). A natural logarithm transformation did not make the data gaussian (Figure 4F); nor did other logarithmic bases (data not shown). A binary logarithm function was used to transform LMA data to ascertain the significant negative correlation with Falling Numbers (FN) (19, 23). FNs inferior to 300 sec, which is the commercialtrade cut-off manifestingsignificant alpha-amylase activity, corresponded to log2 LMA value of -3 (23). In our work, an inverse function normally distributed LMA values, albeit as a slightly asymmetrical bell curve (Figure 4G). This INV(LMA) data was further standardised (centred around zero and scaled down to comparable variance) when it was incorporated at the peptide level which did not compromise its gaussian distribution (Figure 4H).

##### 3.2.1.4. Predicting LMA missing values

Out of the 3,990 reproducibly processed grain samples, 217 were not measured for LMA. We employed a univariate PLS regression strategy to impute them. Using our 2,996 peptide set with the highest VIP scores (see section 3.2.1.2), we tested various PLS regression models (data not shown) using a random selection of 179 samples out of the 934 unbiased sample set which ranged from 0.5 to 4.9. This testing set was analysed against the remainder of the unbiased set (755 samples). The best regression model utilised 20% of the valid values and 20 latent factors; it predicted the 179 tested values with 93% accuracy (Supplementary Figure S8A).

This model was not accurate for small LMA values with a R^2^ of 6%, even imputing negative values (Supplementary Figure S8B). Yet, it was 98% accurate for LMA measurements greater than 0.17 u/g(Supplementary Figure S8C). Itwas more critical to faithfullyestimate high LMA values given that it was the criteria for grain soundness; our PLS regression (PLSR) model fulfilled this. We applied the model’s parameters to predict the 217 LMA missing values against the unbiased set of 934 samples; the imputations ranged from -0.29 to 0.63 u/g (Supplementary Figure S8D). The negative values were converted to zeros. LMA predictions are reported in Supplementary Table S1.

The simplest method for imputing missing data relied on single value imputation, such as the mean (89), whist more complex methods were based on regression (90) or K-Nearest Neighbours (KNN) which estimates a missing data point using distances calculated from its most similar neighbours (91). Invented in 1966 (92), PLS regression has become very popular notably in the fields of bioinformatics (93) and spectroscopy (94). Nengsih and colleagues demonstrated that while computation times increased with the proportion of missing data, up to 30% missing values could be imputed using PLSR (95). In our study, LMA was the single trait provided to analyse LC-MS1 data. Not imputing missing LMA measurements meant that 5.4% (217/3,990 samples) of our dataset would have been useless, therefore it was a worthwhile effort. Alongwith PLSR, we have also tested multivariate linear regression (MLR), univariate polynomial regression and KNN imputation by varyingseveralparameters including valid value percentage, number of latent factors, number of parameters (for MLR), as well as distance computation and number of K (for KNN), albeit without success (data not shown).

##### 3.2.1.5. Incorporating LMA trait at the peptide level for biomarker discovery

Because we only had a single trait to make biological sense of our big data, we introduced all 3,990 LMA values (including the predicted values) which characterised wheat samples at the peptide level by transposing it and renaming “Cluster_AAA”. This added one extra row to our dataset of 32,336 peptides to make a final matrix of 3,990 columns (wheat samples) and 32,337 rows. This way, we could apply statistical analyses that would group peptides that behaved similarly or conversely to our LMA trait thereby facilitating biomarker discovery. To permit the comparison between LMA and grain peptides, we first needed to normalise and standardise LMA values, as detailed above in section 3.2.1.3, prior to their transposition.

Having LMA incorporated with wheat grain peptides (as Cluster_AAA) further helped us assess the relevance of the statistical tests carried out by validating anticipated results. For instance, when performing a correlation analysis with LMA, as expected Cluster_AAA achieved a positive correlation of 1. In another instance, when executing a one factor linear model with LMA as a covariate, Cluster_AAA was confirmed to yield a q-value of 0. Finally, when performingmultivariate clusteringanalyses (HCA, SOM, k-means), this strategy assisted us in finding peptides with profiles similar to that of Cluster_AAA.

#### 3.2.2. Statistical analyses to discover LMA-responsive biomarkers

Big data produced by gene expression studies are too large to analyse by mere sorting in spreadsheets or plotting on few charts. Multivariate data analyses such as clustering and correlating methods are required to make sense of the data (96, 97). Yet, as helpful these multivariate analyses are, they are not as statistically robust as uni- or bivariate analyses (96) to test the relationship between peptides and LMA. We thus performed a few uni-, bi- and multivariate analyses to explore our large dataset against our single LMA trait.

##### 3.2.2.1. Unsupervised multivariate clustering analyses (SOM, k-means, HCA) for pattern recognition and peptide profiling of LMA phenotype

As multivariate analyses handle integral datasets and iteratively impute many statistics, they incur heavy computational costs. Suffering multiple Genedata server crashes, we could only apply such methods to a subset of our data. Using the unbiased set of 934 wheat samples and the list of 7,254 peptides with LMA-responsive VIP scores above 1 (see section 3.2.1.2), we have performed three unsupervised clustering analyses, SOM, k-means and divisive HCA. Because we had incorporated the LMA trait at the peptide level as Cluster_AAA, we could look for groups resulting from these analyses which assembled peptides behaving similarly to Cluster_AAA. Clustering or cluster analysis corresponds to a set of learning methods grouping observations that share similar characteristics. Within a set of related values of the variables analysed, these methods find feature patterns which generate clusters that group similar observations (98). Unsupervised clustering analyses are commonly employed in gene expression studies (97).

In our experiment, the SOM model yielded 48 groups comprising 8 to 555 peptides with mean distances from 0.09 to 0.80. The group including Cluster_AAA (4, 3) contained 26 biomarker peptides; its distance from the group centre ranged from 0.00-0.83 with a mean of 0.38 and a SD of 0.31 (Supplementary Table S4). Cluster_AAA stood 0.70 from the group centre. While SOM has been widely used in exploratory data analyses in diverse fields (99), it has only been applied to proteomics in the context of animal cell culture (100), GPI anchor prediction (101), transmembrane helix predictor (102) protein conformation (103) or protein-protein interaction (104), never in plant grains.

We tested different number of neighbours (k) and observed that the larger k the greater the variance explained by the k-means model (data not shown). Applying the biggest k possible (20) produced a model that overall explained 71.1% of the variance. Group 14 with a variance of 35% contained 93 biomarker peptides spanning a distance of 0.12 to 0.94, including Cluster_AAA whose distance was 0.79 (Supplementary Table S4). K-means clustering was well adopted by the proteomics community to group gene products of similar profiles, notably in plants such as bamboo (105), nightshade (106), or grape (107), but to our knowledge not in wheat. In developing corn grains, coordinated protein expression associated with different functional categories was revealed by a k-means clustering analysis (108).

We successfully applied an agglomerative 2-D HCA to cluster both samples and peptides (data not shown) but failed to select individual groups to retain the one hosting Cluster_AAA. Instead, we performed a divisive HCA which ordered the peptides into clusters that could then be chosen individually. Cluster_AAA belonged to a group of 33 biomarker peptides (order 1915-1947, Supplementary Table S4). We could not find in the literature any proteomics study which resorted to divisive HCA; conversely, classic (agglomerative) HCA created in 1998 (109) and its extension 2-D HCA (110) are widely used by the community, including wheat scientists (111–115). Using agglomerative HCA on 2-DE-resolved proteins, Tasleem-Tahir distinguished nine expression profiles throughout wheat grain growth, from anthesis to maturity (115). In their gel-free iTRAQ analysis of early developing wheat endosperms (from 7-28 days post-anthesis (DPA)), Ma and colleagues employed HCA to delineate starch processes (113). Similarly, five major protein expression patterns across developmental stages 4-12 DPA were outlined using HCA (116). HCA was also employed to explore the change in expression of embryo andendorsperm proteomes duringwheatseed germination (117). In their comprehensive proteomics andproteogenomics study of keydevelopmentalstages of 24 wheat organs and tissues, Duncan and colleagues showed that HCA faithfully assigned samples to three main clusters corresponding to first photosynthetic tissues (leaves, bracts and other green organs), second non-photosynthetic, developmental and reproductive organs (pollen, stem, anther, coleoptiles, roots, immature spike), and third grain (developmental series, embryo, pericarp, endosperm) (111). More recently, Cao and colleagues discriminated differentially expressed proteins in two wheat lines using HCA (82). All these reports demonstrate that genotype-, sample- and tissue-specificity of protein profiles can be highlighted using unsupervised clustering tools.

##### 3.2.2.2. Bivariate analyses (correlation and linear regression) to consider each individual peptide against LMA

As bivariate analyses handle only two variables at a time, they are not computationally taxing. We were thus able to apply such methods on our complete dataset comprising 3,990 samples and 32,337 peptides (including Cluster_AAA). Due to the quantitative nature of LMA trait, we could not perform an analysis of variance (ANOVA). We have thus carried out two bivariates analyses, a correlation and a linear model. Because we had incorporated the LMA trait at the peptide levelas Cluster_AAA, we could assess the validity of our analyses based on the outputs produced by the latter.

In our experiment, correlation coefficients ranged from -0.07 to 0.3, except for Cluster_AAA which as expected attained absolute positive correlation with a R^2^ of 1 (Supplementary Table S4). Our coefficients do not show a strong relationship between peptide profiles and LMA. We arbitrarily chose an absolute value of 0.15 to retain any LMA-associated peptide which excluded all negatively-correlated features but included 28 positively-correlated biomarkers. Correlation analyses are frequently employed in proteomics to unravel proteins underpinning particular sample types, conditions or traits (118), and wheat is no exception (119–127). Concordance of transcript and protein profiles in wheat grain were assessed via correlation coefficients, which increased with seed maturity (120, 126). Grain yield and grain protein content were observed to be negatively correlated, yet both also positively correlated to nitrogen availability in a wheat genotype-specific manner (128).

The q-value for the linear regression slope indicates whether changes in the explanatory variable are significantly linked with changes in the outcome. In our work, we looked for significantrelationships betweenthe 32,337 peptides (including Cluster_AAA) and the inverse function of LMA which assumed normality as a covariate factor. Q-values ranged from 6 x 10^-8^ to 1, with the exception of Cluster_AAA which exhibited a q-value of 0 as expected (Supplementary Table S4). We arbitrarily applied a 5% q-value threshold to consider 494 biomarker peptides whose change in expression profiles were significantly linked to variation in LMA measurements. Linear mixed models are regularly employed by the proteomics community for biomarker discovery approaches (129–132), but as far as we know not on wheat grains.

##### 3.2.2.3. Compiling all statistical analyses to generate a list of candidate peptides and binning LMA values for biomarker profiling and t test

In this study, LMA-responsive biomarkers were selected based on the statistical analyses presented above and had to fulfill at least one of the following criterium: belong to SOM group (4, 3), be included in k-means group 14, bear a divisive HCA order from 1915 to 194, exhibit a correlation R^2^ greater than 15%, or display a q-value inferior to 5%. This created a list of 531 biomarkers, most of which fulfilled several statistical criteria and all of them exhibiting a VIP score for the LMA-responsive PLS greater than 1 (Supplementary Table S4).

When attempting to chart the biomarker profiles, we were faced with the challenge of plotting 3,990 datapoints per gene product which ruled out typical line graphs, scatter plots, histograms or utilising oversized illegible heat maps to represent all data points simultaneously (data not shown). We consequently adopted a data reduction strategy involving binning the samples into 8 or 2 arbitrary bins based on their LMA values.

The 8-bin profiling comprised all 3,990 samples sorted by increasing LMA measurements and partitioned into 8 groups of equal sample size (∼499 samples/bin, Supplementary Table S1). Plotting the average of each bin as a line chart faithfully maintained the pattern of LMA measurement observed in Figure 4A with a flat profile for the first 7 bins followed by a steep increase in the last bin (Supplementary Figure S9A).

This profilingstrategy was notused for statistical purpose butproved usefulduringdata mining of all identified 5,514 peptides upon using tools that offered quantitative charting such as Pathway Tools and Circos (see below).

The 2-bin profiling only featured the 934 unbiased samples separated according to an arbitrary 0.17 u/g threshold (Supplementary Table S1). Plotting the average of each bin as a histogram clearly displayed a marked quantitative increased from bin 1 to bin 2 (Supplementary Figure S9B). This simple representation tool allowed us to categorise the 531 biomarkers as being either up-regulated when bin 2 was taller than bin 1 denoting an accumulation in samples with LMA>0.17 u/g or down-regulated when bin 1 was taller than bin 2 denoting an accumulation in samples with LMA<0.17 u/g.

This oversimplified binning scheme allowed us to perform one last statistical analysis on the 532 (including Cluster_AAA) biomarkers using the unbiased set of 934 samples, namely a Student’s t test with an effect size. We generated a volcano plot based on the p-values and the directed effectsize (i.e. fold change) which clearly delineated the biomarkers according to their accumulation in bin 1 or 2 (Figure 5A).

**Figure 5:**
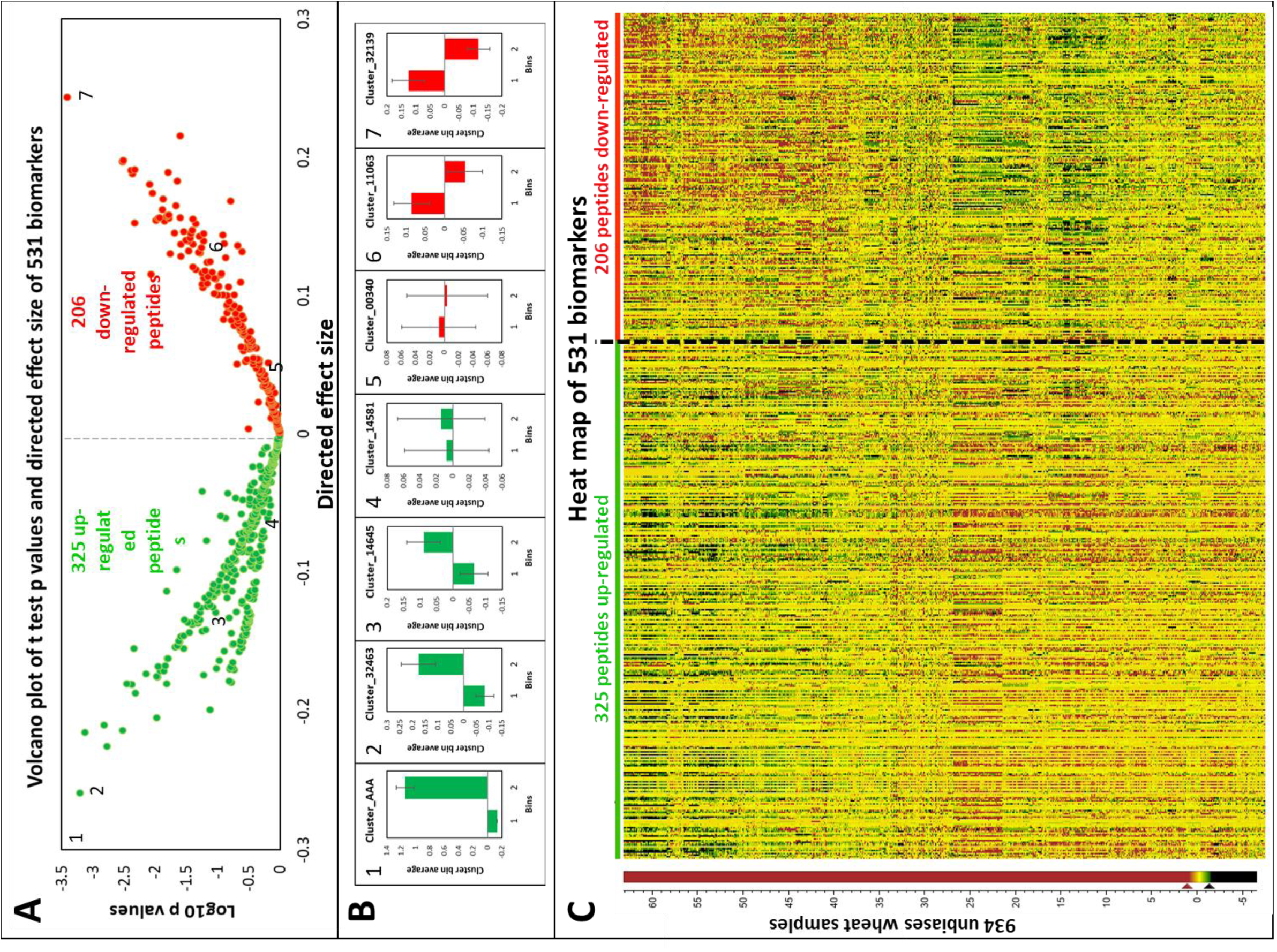
Volcano plot from t test and heat map of up- and down-regulated 531 biomarkers using the unbiased set of 934 wheat samples. (A) Volcano plot of the 325 up-regulated and 206 down-regulated biomarkers. Numbers position exemplary peptides plotted in panel B. Cluster_AAA with coordinates (−1.2, -23.5) is an outlier in the upper left corner and is not featured for display purpose;(B) Mean histograms along 2 bins of clusters illustrating up- and down-regulation patterns and located with numbers on panel A. Standard errors are depicted with the vertical bars. Bin 1 corresponds to 467 samples with LMA < 0.17 u/g and bin 2 corresponds to 467 samples with LMA > 0.17 u/g; (C) Heat map corresponding to the Volcano plot in panel A with peptides sorted according to directed effect size and samples sorted based on HCA cluster order.

More LMA-related biomarkers were up-regulated (325) than down-regulated (206) according to our 2-bin profiling. This was explained by the fact that all our statistical analyses, bar the PLS and linear model, favoured peptides behaving similarly to Cluster_AAA a proxy to LMA actualmeasurements. Some exemplarypatterns are displayed as histograms with error bars and compared to thatof Cluster_AAA to expose the assortmentof up- and downregulation profiles (Figure 5B). Because the 2-bin representation was very reductive, we also present a heat map of all the intensities of the 532 biomarkers (including Cluster_AAA) sorted by directed effect size (i.e. fold change) in each of 934 unbiased wheat samples organised by HCA cluster order (Figure 5C). No strong differential expression trend appeared apart from a horizontal gradient of colours from left to right denoting the change from up- to down-regulation of the biomarkers and a swap in colour vertically suggesting that samples were efficiently classified by the HCA. Despite merely featuring a small subset (934×532) of our global dataset (3,990×32,337), the heat map looked noisy and remained very hard to interpret due to an excessive number of data points (469,888 quantities) and the lack of visually striking pattern. This further reinforced the need to devise simple representations tools such as a Volcano plot when reporting results on big data.

To our knowledge, volcano plots have not been widely adopted by the proteomics community, let alone wheat grain scientists with only one report so far (84), unlike heat maps which are frequently reported in proteomics publications (133). In our work, we sorted the 531 biomarker peptides according to their 2-bin fold changes and wheat sample based on their LC-MS molecular similarity (Figure 5C). Zang and colleagues have adopted heat maps to profile the proteins underpinning seed tissue organogenesis (134).

#### 3.2.3. Mining biomarkers to make biological sense of the data

Among the 531 biomarkers that exhibited significance levels in response to LMA measurements, 390 were identified by LC-MS2 and matched 3,798 protein accessions (Supplementary Table S5). This list included the most abundant and homoeologous proteins such as the prominent storage and starch-related proteins, gliadins, glutenins, avenins, and starch synthases as well as constitutive proteins such as histones, protein disulfide isomerases, and tubulin, or else stress-related proteins such as heat shock and 14-3-3 proteins. We did not identify any peptides belonging to LMA in this study, likely because we did not target high LMA samples. To visualise our peptides of interest in a biological context, we have undertaken a series of data mining steps. We have also made use of our 8- or 2-bins profiling strategy when using quantitative mapping tools. The 2-bin profiling is hereafter referred to it as up- or down-regulated gene products. The data mining tools presented below suited wheat proteins. Many other in silico tools are freely available online which we encourage the community to employ; however, we would not recommend using String or PlantReactome which in our hands yielded very little results.

##### 3.2.3.1 Protein descriptions and GO terms from UniProtKB

Out of the 8,044 identities, 7,939 could be mapped in UniProtKB which flagged 6,457 GOMF terms, 3,769 GOCC terms, 3,991 GOBP terms, as well as 1,385 unique protein names (Supplementary Table S3). Power BI proved very useful to mine identified peptides and simultaneously plot some of their features as histogram, scatterplot, pie chart, violin plot, tree map and word cloud into a single dashboard (Supplementary Figure S10A) and then drill down on some aspects, for instance inhibitor (Supplementary Figure S10B) or deamidation (Supplementary Figure S10C).

The protein names were turned into word clouds and the most frequent GO terms for each category were presented as tree maps. Standing out from the cloud were the words “protein”, “containing”, “domain”, “subunit”, “glutenin”, “LMW”, “molecular”, and “weight”, confirmingthe preponderance of LMWglutenin subunits and domain-containing proteins such as AAI domain-containing protein homoeologous to alpha-amylase inhibitors (Supplementary Figure S11B-D). Also predominant among identified proteins were the words “alpha” and “gliadin”. Word cloud is a text processing method that offers an efficient and compact visualization of the most frequent terms in a text (135), yet it seldom appears in the scientific literature. It has been cleverly used to categorise moonlighting proteins (136) or depict the history of GOMF terms (137), but not in the wheat proteome. Representing our 390 identified LMA-responsive biomarkers as word clouds revealed that up-regulated peptides belonged predominantly to alpha-gliadins whereas down-regulated peptides mostly matched LMW glutenins (Figure 6A,F).

**Figure 6:**
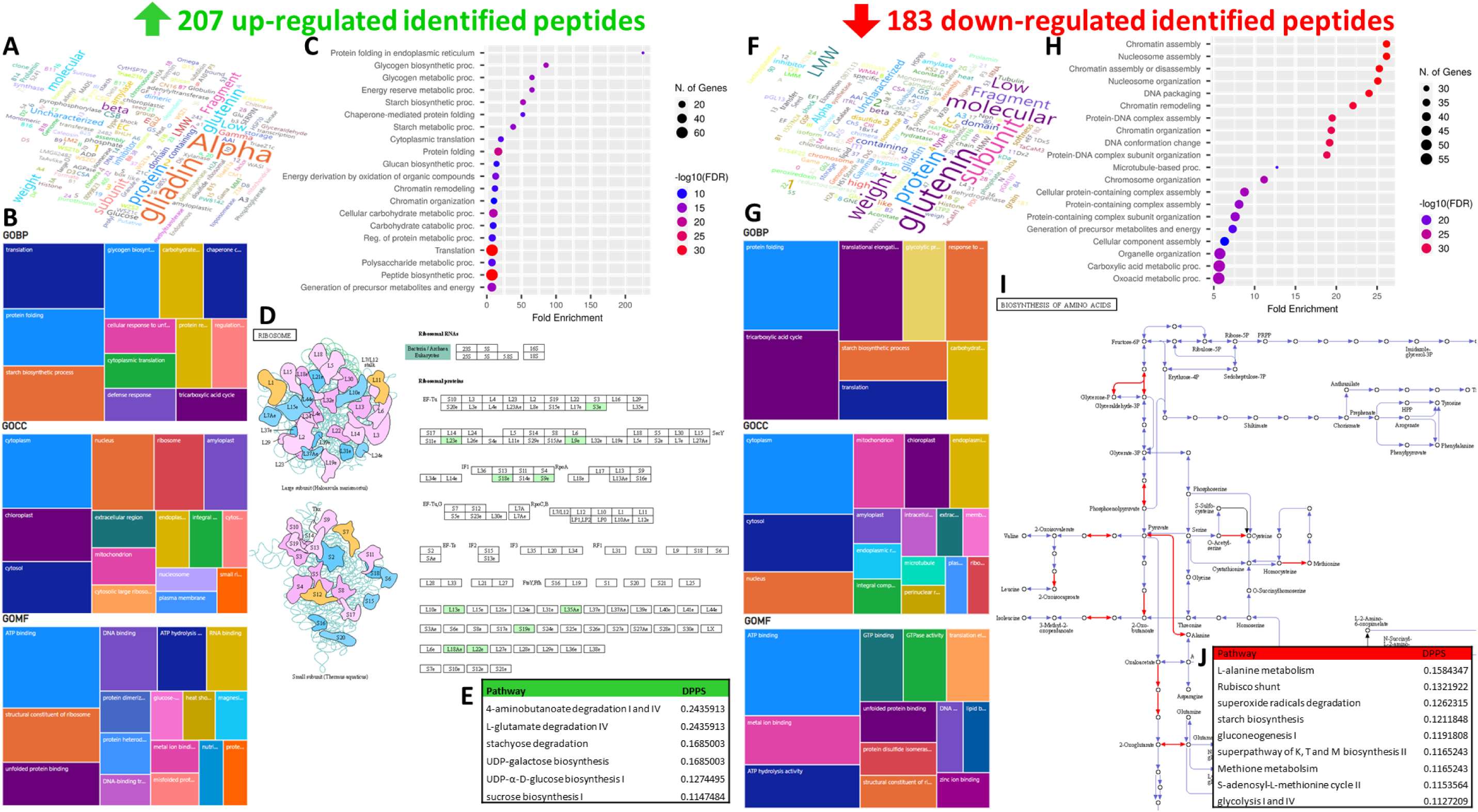
Data mining of up- and down-regulated biomarkers. (A, F) word cloud of protein names; (B, G) tree maps of GO terms for BP, CC and MF categories; (C, H) dot plots from ShinyGO;(D, I) most significant KEGG pathways, ribosomes for up-regulated biomarkers and AA biosynthesis for down-regulated biomarkers; (E, J) differentially perturbed pathways (DPPS) from Pathway Tools.

Rather than adopting a pie chart or histogram to plot the GO terms of all identified proteins as commonly reported, we opted for tree maps which were initially implemented for microarray data (138, 139) and later integrated into the web server REVIGO (140) used during our wheat method optimisation (41). For all 8,044 identified proteins in the present study, we generated the tree maps for all three GO classes using Power BI as it afforded more display options than REVIGO. The most frequent biological processes (GOBP) were “polysaccharide catabolic process” (5, 643), “starch biosynthetic process” (3, 688), “nucleosome assembly” (3, 626), “protein folding” (2,950) and “protein refolding” (2,499) (Supplementary Figure S11E). “Cytoplasm” (11,888), “extracellular region” (9,964), and nucleus” (7,478) were the most common cellular components (GOCC); recording 3,687 entries, the amyloplast was listed in 6^th^ position (Supplementary Figure S11F). With 37,308 occurrences, the “nutrient reservoir activity” was by far the most recurrent molecular function (GOMF), followed by “ATP binding” (7,012) and “serine-type endopeptidase inhibitor activity” (5,811) (Supplementary Figure S11G). The list of dominant proteins and associated GO terms in this work pointed to a storage organ such as the wheatseed and confirmedwhathas previously been reported in wheat grain (41, 126, 134, 141-143). All GO terms against the 390 identified LMA-related biomarkers are listed in Supplementary Table S5. The 207 up-regulated biomarkers came mostly from cytoplasmic and chloroplastic proteins involved in protein translation and folding, with ATP binding activities (Figure 6B). The 183 down-regulated peptides predominantly belonged to cytoplasmic and cytosolic proteins acting in protein folding and TCA cycle and bearing ATP binding activity (Figure 6G).

##### 3.2.3.2. KEGG to retrieve Pathway, Brite and Module names

From the 8,044 fasta sequences, 677 unique KEGG Orthologs (KOs) could be retrieved which mapped to 327 KEGG pathways, 41 brites and 117 modules and annotated 11,888 peptides (Supplementary Table S3). Identified proteins belonged to 179 (26%) KEGG metabolic pathways with 109 (16%) KOs involved in the biosynthesis of secondary metabolites Supplementary Figure S12A), including sugar-related enzymes such as amylases, sucrose synthases, hexokinases, fructokinase sand beta-glucosidases.

Half of KOs pointed to enzymes (336), then exosomes (71, 10%), ribosomes (62, 9%), and chromosome-associated proteins (60, 9%) (Supplementary Figure S12B). Primary metabolisms such as glycolysis, TCA cycle and gluconeogenesis were prominent KEGG modules (Supplementary Figure S12C). Unexpectedly, 62 KOs (exclusively ribosomal proteins) were associated with “Coronavirus disease – COVID 19” pathway. Similarly, many proteins were linked with other human-related afflictions (e.g. sclerosis, neurodegeneration, Parkinson, Huntington, Alzheimer and prion diseases; Supplementary Figure S12A). This demonstrated the limitations of using generalist databases like KEGG that are mostly relevant to human research to map plant proteins. While KEGG plant interface exists (https://www.genome.jp/kegg/genome/plant.html) (144), plant-related datasets are dispersed throughout the whole KEGG server so that one cannot exclusively mine plant-specific entries. There is a need for future KEGG iterations to restrict searches to relevant taxa. Notwithstanding non-plant hits, pathways symptomatic of grains were accurately captured in this experiment such as the carbon metabolism (42, 6%), glycolysis/gluconeogenesis (25, 4%), as well as the starch and sucrose metabolism (18, 3%) (Supplementary Figure S12D-F). Despite the constraint raised above, KEGG remains a database widely employed to explore plant proteomes, including wheat grain proteins (41, 145–147). Mapping our 390 LMA-associated biomarkers (Supplementary Table S5) highlighted that many up-regulated peptides came from ribosomal proteins (Figure 6D) while several down-regulated peptides belonged to enzymes acting in the biosynthesis of AAs (Figure 6I).

##### 3.2.3.3. ShinyGO to retrieve enriched functional categories and chromosomal positions

Multiple online tools exist to efficiently mine GO terms, however only a few cater for non-model species, let alone plants (148–150). When looking for relevant mining tools during our method development stage, we resorted to AgriGO online program which specifically focused on agricultural species and offered valuable illustrations to display enrichment sets (41). Unfortunately, AgriGO server is no longer available. We have found instead ShinyGO (50), recently developed, which surpassed AgriGO not only in terms of enrichment visualisations butalso provided wheatprotein chromosomalpositions,desirable for Circos plots. A downside of ShinyGO was that it did not perform well with UniProt accession IDs, hence the prerequisite to retrieve TRAES IDs from UniProtKB. A total of 6,622 TRAES accessions corresponding to the 8,044 UniProt proteins were thus retrieved, of which 4,571 could be mapped by ShinyGO (Supplementary Table S6). An enrichment analysis ensued and could be visualised as a chart, tree, network and chromosomal map; density plots and histograms were also produced (Supplementary Figure S13).

The most enriched category was the TCA cycle with a fold enrichment in excess of 12.5 and the most significant GO classes were translation and peptide biosynthesis with an FDR inferior to e^-160^ (Supplementary Figure S13A,E). Protein folding and ribonucleoprotein complex biogenesis stood out as well among the proteins identified in this study (Supplementary Figure S13B). Identities covered the whole genome with lower density around centromeres (Supplementary Figure S13F). ShinyGO and other online data mining algorithms were employed to predict genetic components systems implicated in the plant model species *Arabidopsis* in response to high light from transcriptomics datasets publicly available (151). Our results exemplify the relevance of ShinyGO for non-model plant species; we could not find other cereal reports making use of it, probably due to its recent emergence (50). A fold enrichmentexceeding 200 was found amongthe 207 up-regulated peptides from gene products involved in protein folding in endoplasmic reticulum (Figure 6C), followed by glycogen metabolism, energy reserve and starch biosynthesis. ShinyGO enrichment analysis produced very different results for our 183 down-regulated peptides, mostly invoking chromatin assembly and remodeling, nucleosome assembly and organisation, DNA packaging and conformation change, as well as protein-DNA complex assembly and organisation(Figure 6H).

##### 3.2.3.4. Pathway Tools to retrieve differentially perturbed pathways based on 8-bin profiling

As useful as the program described above are, they yet do not accommodate quantitative data, unlike Pathway Tools (51) made available online by the Plant Metabolic Network server and curating the PlantCyc databases encapsulating 126 plant and algae species (https://plantcyc.org/), including BreadwheatCyc (52). We could thus display protein expression data on pathway diagrams in a dynamic and interactive way. Using the 6,622 TRAES accessions corresponding to the proteins identified in this study and the quantitative data averaged along 8 bins, we mapped 1,432 proteins in the *T. aestivum* Pathway Tools website (Supplementary Figure S14A).

The change in expression profiles along the 8 bins was recorded and showed that all peptide quantities varied across sample groups with multiple trends throughout the whole cellular overview (Supplementary Video SV1). As previously reported (41), the primary and secondary metabolisms were well covered. Overall quantities of homoeologous wheat proteins involved in TCA and glyoxylate cycles declined along 8 bin expression profiles (Supplementary Figure S14B).

Also featured was plant hormone biosynthesis (Supplementary Figure S14C) which was lacking in the other exploratory tools, thus demonstrating the superiority of *T. aestivum* Pathway Tools over other databases (41). The 8 bin-profiling hinted an accumulation of proteins related to auxin, cytokinin and gibberellin biosynthesis and a reduction of enzymes participating in 5-deoxystrigol, brassinosteroid, andjasmonate synthesis in LMA-rich samples. Hormonal response was flagged as one of the biochemical mechanisms of LMA expression, in particular gibberellin and ABA signalling (22, 25, 152). Focussing on the ent-kaurene biosynthesis, expression patterns accumulated in low LMA samples at the initial step of the pathway and diminished in high LMA samples at the last step (Supplementary Figure S14D-E). The first biosynthetic step is controlled by ent-copalyl disphosphate synthase (TaCSP) which was reported to be associated with LMA via a major locus on wheat chromosome 7B accordingly renamed as LMA-1 (153). TaCSP (Cluster_22809 in Supplementary Figure S13F) was one of our biomarkers. Even though databases such as Pathway Tools mapped TaCSP to the gibberellin metabolism, its function with this phytohormone was recently contested and it was suggested that high pI alpha-amylase synthesis in the aleurone of developing wheat grains would be independent of gibberellins during LMA response (40). Other biomarkers matching phytohormone-associated proteins included a cytokinin dehydrogenase whose decreasing pattern picked up in the bin containing all the wheat sample registering high LMA (Cluster_24683 in Supplementary Figure S14F), and a Responsive to ABA (Rab) protein whose expression profile closely resembled that of Cluster_AAA (Cluster_36748 in Supplementary Figure S14F). Interestingly, Cluster_24621 with an increasing expression profile belonged to an uncharacterised protein annotated with GO terms “Response to Auxin” and “Response to ethylene” (Supplementary Figure S14F).

Because Pathway Tools handles quantitative data, it produced lists of differentially perturbed pathways (DPPS) for each set of up- and down-regulated biomarkers. Pathways characterising wheat grains with high LMA measurements were degradations of aminobutanoate, glutamate, and stachyose, as well as biosynthesis of UDP-galactose, UDP-glucose and sucrose (Figure 6E). DPPS differentiating samples with low LMA activities were AA metabolisms (A, K, T, and M) rubsico shunt, superoxide radical degradation, starch biosynthesis, gluconeogenesis, S-adenosyl-M cycle and glycolysis (Figure 6J). Our method study aside (41), we could not find any other wheat gene expression study utilising this impressive PlantCyc database. However, work on other plant species have amply demonstrated its value (154–159).

##### 3.2.3.5. Circos plot to visualise chromosomal positions, expression profile and statistics of identified proteins and biomarkers

Invented over a decade ago (53), Circos plots have proven so valuable to efficiently represent qualitative and quantitative information that a multitude of emulations have since arisen, including its packaging within the Galaxy server (55) which we took advantage of here. When the IWGSC released *T. aestivum* genome and published their findings, the genomic features were elegantly and succinctly captured in a circular plot which highlighted homeologous genes and translocated chromosomal regions (9). Being infinitely flexible, Circos plots can chart any data as multiple concentric circular layers provided the correct file format is applied. We opted to chart proteins encoded by genes we could locate on the genome (chromosomal positions retrieved from ShinyGO analysis) and overlay their expression profiles, along with some statistics of candidate LMA-responsive biomarkers (Figure 7).

**Figure 7:**
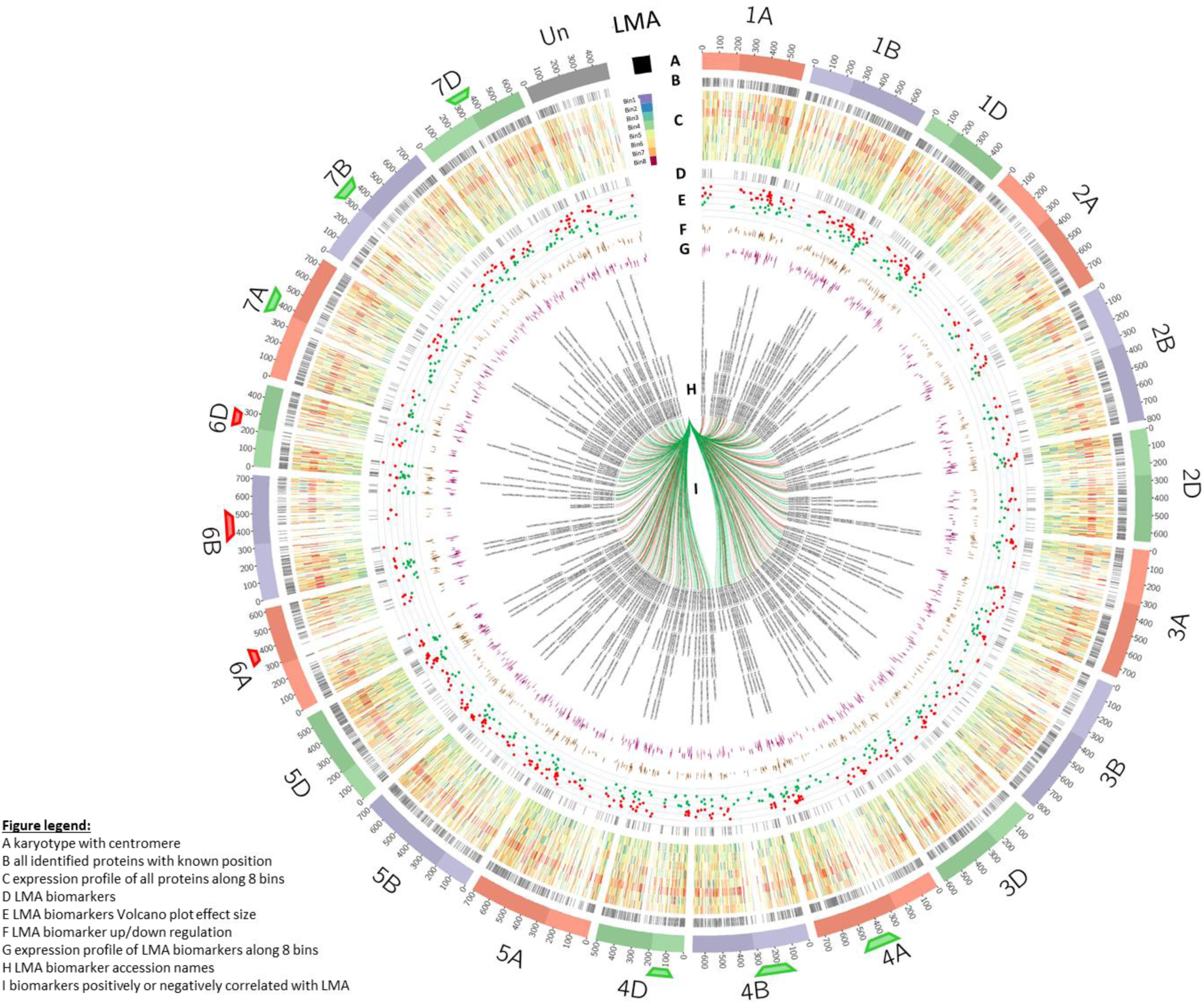
Circos plot of identified proteins and LMA-responsive biomarkers with expression patterns and statistics. (A) *T. aestivum* karyotype with chromosome length marked each 10^6^ cM and centromeres indicated by the change in shade. LMA is displayed as a chromosome to portray the trait’s 8-bin colour pattern in trace C; (B) chromosomal positions of all identified proteins as highlights; (C) profiling of all identified proteins along 8 bins as heatmaps. LMA pattern is provided as a reference; (D) chromosomal positions of all identified LMA-responsive biomarkers as highlights; (E) Volcano plot effect size of biomarkers as scatterplot. Red denotes down-regulation and green denotes up-regulation; (F) profiling of biomarkers along 2 bins as stacked histogram; (G) profiling of biomarkers along 8 bins as stacked histogram;(H) biomarker accession IDs as text labels; (I) positive (green) and negative (red) correlation with LMA as links. Green and red tags under chromosomes 4ABD, 6ABD, and 7ABD denote genomic regions exclusive to biomarkers up- and down-regulated, respectively.

Proteins identified in this experiment aligned with the full genome, densely covering each chromosomealbeitless so around centromeric regions (Figure 7B). Overall, expression profiles along 8-bin accumulated in bins 1-6 corresponding to wheat samples with low LMA and decreased in bins 7-8 characterised by high LMA samples (Figure 7C). LMA-related biomarkers were evenly dispersed on all chromosomes (Figure 7D). Plotting their effect size (fold changes, Figure 7E) outlined thatmostgenome areas hosted both up- and down-regulated biomarkers bar a few exceptions on chromosomes 4, 6 and 7 for all 3 genomes A, B, and D. Only up-regulated biomarkers could be seen on chromosome 4A region 300-500 x 10^6^ cM and chromosome 7A region 300-480 x 10^6^ cM (replicated on genomes B and D). They matched three uncharacterised proteins, a 60S ribosomal protein L18a, a glucose-1-phosphate adenyltransferase, a polyadenylate-binding protein, a 14-3-3 protein and a protein disulfide isomerase (Supplementary Table S5). Conversely, chromosome 6A region 300-410 x 10^6^ cM (replicated on genomes B and D) exclusively located down-regulated biomarkers matching a glyceraldehyde-3-phosphate dehydrogenase, a glutathione peroxidase, a tripeptidyl-peptidase II and an uncharacterised protein. Charting biomarker correlation values with LMA as links failed to isolate stretches of genomic areas specific to LMA-responding proteins (Figure 7I). This could be explained by the fact that LMA expression in our experiment elicited a complex metabolic responseinvolvingmany gene products independentof their genomic position. LMA is indeed a multigenic trait; associated quantitative trait loci (QTLs) have been located across all three genomes and would contribute to the LMA phenotype in an independently effective and additive fashion (39).

## Concluding remarks

For the first time, LMA phenotypewas exploredvia proteomics. All the differentially regulated biological processes highlighted in this study by the various data mining means have been condensed into one summarising diagram and organised into broad functional categories (Figure 8).

**Figure 8:**
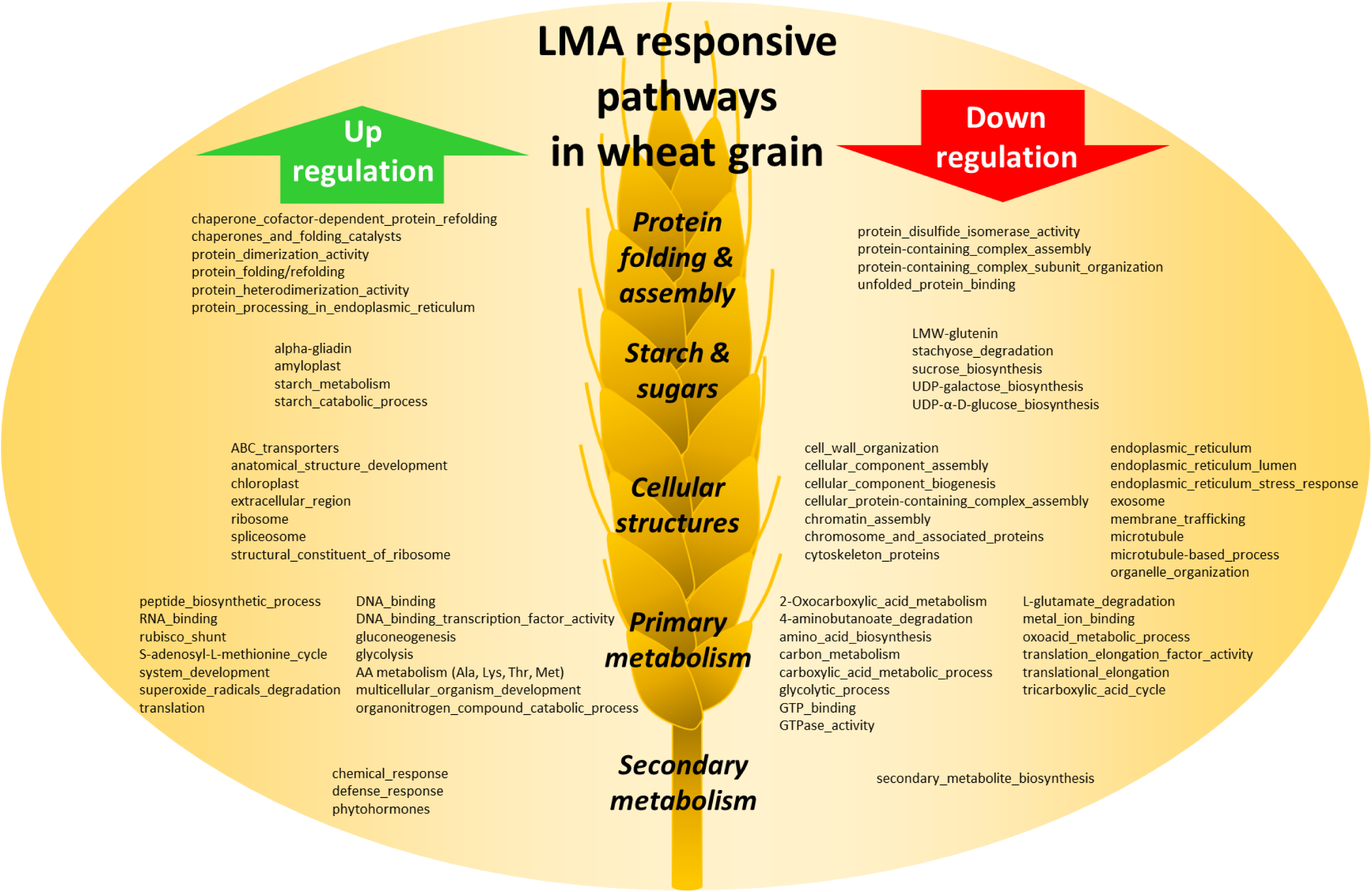
Synopsis of mechanisms involved in LMA response.

In this work, stored LMA-affected grains activated their primary metabolisms such as glycolysis and gluconeogenesis, TCA cycle. It also including DNA- and RNA binding mechanisms, as well as protein translation. This logically transitioned to protein folding activities driven by chaperones and protein disulfide isomerase, as well as protein assembly via dimerisation and complexing. The secondary metabolism was also flagged notably with the up-regulation of phytohormones, chemical and defense responses. LMA further invoked cellular structures among which ribosomes, microtubules, and chromatin. Finally, and unsurprisingly, LMA expression greatly impacted grain starch and other carbohydrates with the up-regulation of alpha-gliadins and starch metabolism, while LMW glutenin, stachyose, sucrose, UDP-galactose and UDP-glucose were down-regulated. This work demonstrates that, whilst we did not find the LMA needle in the proteome haystack, proteomics deserves to be part of the wheat LMA molecular toolkit and should be adopted by LMA scientists and breeders in the future.

## Abbreviations

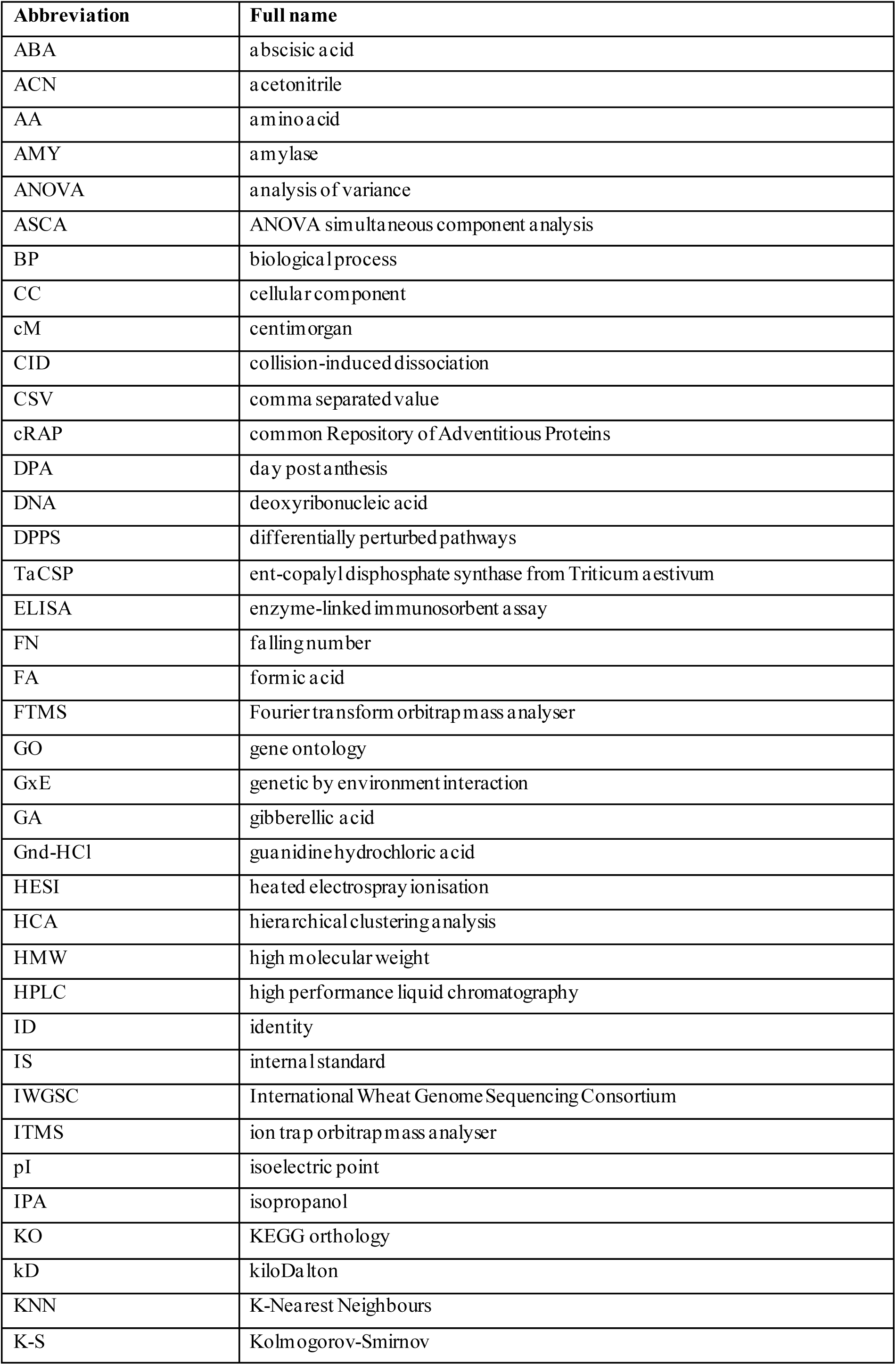
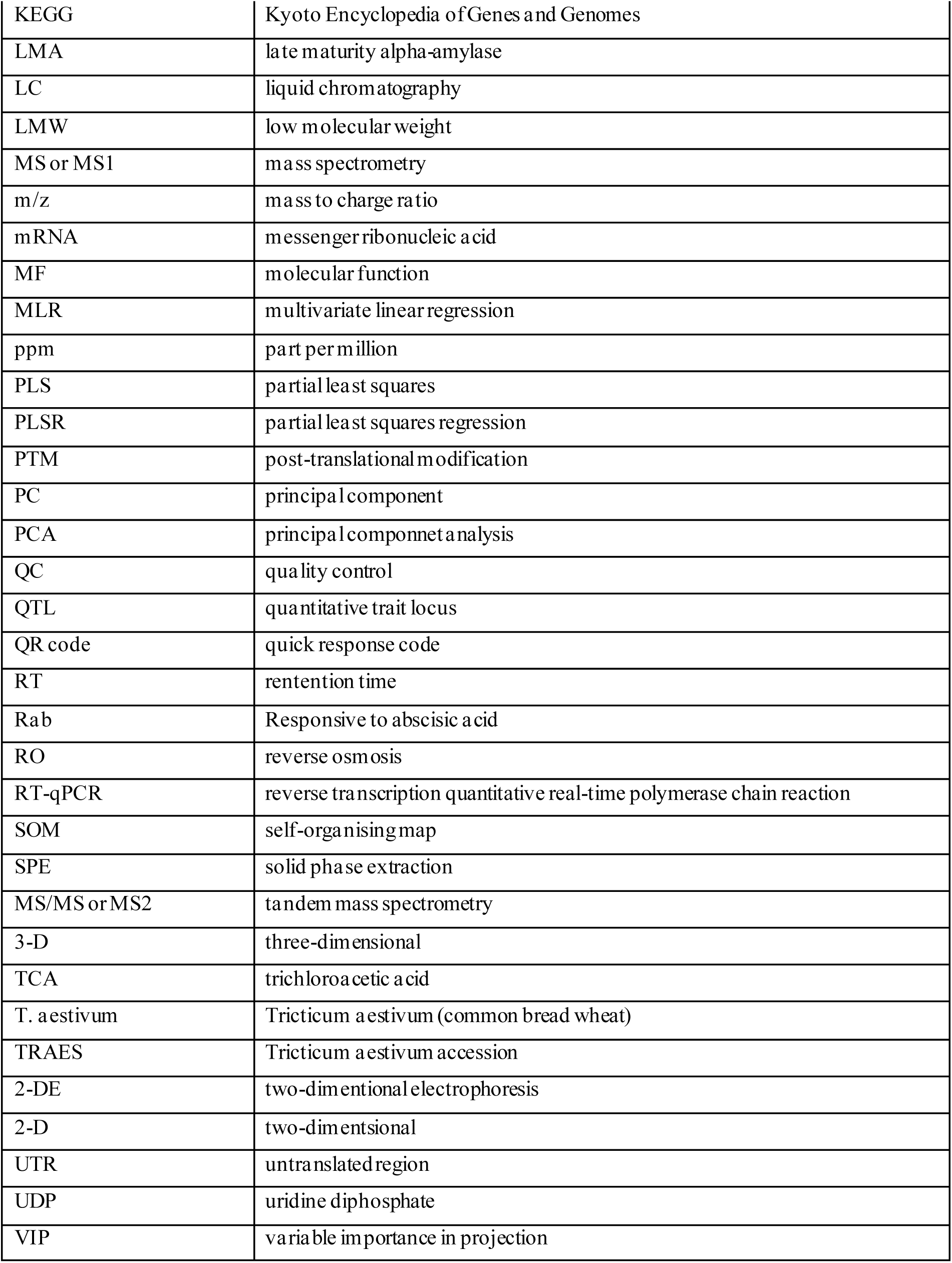

| Abbreviation | Full name |
| --- | --- |
| ABA | abscisic acid |
| ACN | acetonitrile |
| AA | amino acid |
| AMY | amylase |
| ANOVA | analysis of variance |
| ASCA | ANOVA simultaneous component analysis |
| BP | biological process |
| CC | cellular component |
| cM | centimorgan |
| CID | collision-induced dissociation |
| CSV | comma separated value |
| cRAP | common Repository of Adventitious Proteins |
| DPA | day post anthesis |
| DNA | deoxyribonucleic acid |
| DPPS | differentially perturbed pathways |
| TaCSP | ent-copalyl diphosphate synthase from *Triticum aestivum* |
| ELISA | enzyme-linked immunosorbent assay |
| FN | falling number |
| FA | formic acid |
| FTMS | Fourier transform orbitrap mass analyser |
| GO | gene ontology |
| GxE | genetic by environment interaction |
| GA | gibberellic acid |
| Gnd-HCl | guanidine hydrochloric acid |
| HESI | heated electrospray ionisation |
| HCA | hierarchical clustering analysis |
| HMW | high molecular weight |
| HPLC | high performance liquid chromatography |
| ID | identity |
| IS | internal standard |
| IWGSC | International Wheat Genome Sequencing Consortium |
| ITMS | ion trap orbitrap mass analyser |
| pI | isoelectric point |
| IPA | isopropanol |
| KO | KEGG orthology |
| kD | kiloDalton |
| KNN | K-Nearest Neighbours |
| K-S | Kolmogorov-Smirnov |
| KEGG | Kyoto Encyclopedia of Genes and Genomes |
| LMA | late maturity alpha-amylase |
| LC | liquid chromatography |
| LMW | low molecular weight |
| MS or MS1 | mass spectrometry |
| m/z | mass to charge ratio |
| mRNA | messenger ribonucleic acid |
| MF | molecular function |
| MLR | multivariate linear regression |
| ppm | part per million |
| PLS | partial least squares |
| PLSR | partial least squares regression |
| PTM | post-translational modification |
| PC | principal component |
| PCA | principal component analysis |
| QC | quality control |
| QTL | quantitative trait locus |
| QR code | quick response code |
| RT | retention time |
| Rab | Responsive to abscisic acid |
| RO | reverse osmosis |
| RT-qPCR | reverse transcription quantitative real-time polymerase chain reaction |
| SOM | self-organising map |
| SPE | solid phase extraction |
| MS/MS or MS2 | tandem mass spectrometry |
| 3-D | three-dimensional |
| TCA | trichloroacetic acid |
| T. aestivum | Triticum aestivum (common bread wheat) |
| TRAES | Triticum aestivum accession |
| 2-DE | two-dimensional electrophoresis |
| 2-D | two-dimensional |
| UTR | untranslated region |
| UDP | uridine diphosphate |
| VIP | variable importance in projection |

## Declarations

### Ethics approval and consent to participate

Not applicable.

### Consent for publication

Not applicable.

### Availability of data and materials

The LC-MS1 dataset and raw LC-MS2 data generated and analysed during the current study are available in the MassIVE repository, f tp://massive.ucsd.edu/MSV000090572. All data generated or analysed during this study are included in this published article and its supplementary information files.

### Competing interests

The authors declare that they have no competing interests.

### Funding

This research was funded by the Grains Research and Development Corporation (GRDC), Project DJP2001-008RTX.

### Authors’ contributions

Conceptualisation, M.H., H.D., J.P., D.V.; plant materials: J.P., LMA assays: N.R.; grain grinding: D.V., A.B., D.R.; sample processing, D.V., A.B. ; LC-MS maintenance: D.V. and V.E.; LC-MS data acquisition: D.V., and A.B.; LC-MS and LC-MS/MS data acquisition and analysis: D.V.; LC-MS matching with LC-MS/MS in R: S.S.; technical bias removal: T.L.; statistical analyses: D.V. and S.R.; data mining and figures, D.V.; investigation, D.V.; resources, S.R.; data curation, D.V.; writing—original draft preparation, D.V.; review and editing, D.V., T.L., J.P., S.R., and H.D.; visualization, D.V.; logistics: D.V.; supervision, S.R.; project administration, D.V., S.R., H.D., and M.H; funding acquisition, M.H. and H.D. All authors have read and agreed to the published version of the manuscript.

## Acknowledgements

Acknowledgements

We thank Mr Pankaj Maharjan for retrieving all wheat samples from storage. We are grateful for advice on MS/MS targeted methods from Drs Aaron Elkins, Priyanka Reddy from AVR, and Dr Enzo Huang from Thermo Scientific. We are grateful to Carl Thomas and Piotr Malicki from AVR for upgrading Genedata and Mascot servers, as well as maintaining the Bioinformatics Advanced Scientific Computing cluster. We thank Dr Gabriel Keeble-Gagnere from AVR for his critical review of the manuscript.

## Supplementary Figure legends

**Supplementary Figure S1: Genedata Refiner workflow to process all wheat, IS and QC LCMS1 RAW files and export them to Genedata Analyst.** A. Refiner Step 1; B. Refiner Repetition node from Step 1; C. Refiner Step 2; D. Analyst setup. See Materials and Methods for description.

**Supplementary Figure S2: Genedata Refiner workflow to process all wheat LCMS2 RAW files and export them to Excel.** A. Step 1; B. Repetition node from Step 1; C. Step 2; D. Mascot parameters; E. Excel output. See Materials and Methods for description. **Supplementary Figure S3: LC-MS2 RAW maps for each tandem pass.** X-axis delineates 300-2000 m/z. Y-axis delineates 1-35 min Retention Time. White dots represent MS2 events. (A) LC-MS1 map of pooled sample; (B) LC-MS2 map of Pass 1 replicate 1 with 3000 threshold; (C) LC-MS2 map of Pass 2 replicate 1 with exclusion list of 2000 ions fragmented in Pass 1; (D) LC-MS2 map of Pass 3 replicate 1 with exclusion list of 2000 ions fragmented in Pass 2; (E) LC-MS2 map of Pass 4 replicate 1 with exclusion list of 2000 ions fragmented in Pass 3; (F) LC-MS2 map of Pass 5 replicate 1 (same as Pass 1 but with 500 threshold); (G) LC-MS2 map of Pass 6 replicate 1 with inclusion list of 2000 most abundant ions from Pass 1; 1. (H) LC-MS2 map of Pass 7 with inclusion list 1 loaded Global mass tab and 2 m/z tolerance; (I) LC-MS2 map of Pass 8 with inclusion list 1 loaded in data-dependent settings and 2 m/z tolerance; (J) LC-MS2 map of Pass 9 with inclusion list 1 loaded in data-dependent settings and 1 m/z tolerance;(K) LC-MS2 map of Pass 10 with inclusion list 1 loaded in data-dependent settings and 0.5 m/z tolerance; (L) LC-MS2 map of Pass 11 with inclusion list 1 loaded in data-dependent settings and 0.2 m/z tolerance. Maps from other replicates in Passes 1-6 or with inclusion lists 2-10 for Passes 7-11 are not shown.

**Supplementary Figure S4: Histogram of the number of peptides identified using Mascot algorithm and number of MS2 events in each of the LC-MS2 file.** Black bars represent peptide counts (y axis on the left) and orange dots depict MS/MS event counts (y axis on the right).

**Supplementary Figure S5: Histograms (A, C, E) and box plots (B, D) of the number of peptides per accession (A-B, E) and number of accessions per peptides (C-D).** The orange line in panels A and C represents cumulated counts in percent. Panel E displays the peptides with the highest hit counts belonging either to low molecular weight glutenin subunit (LMW-GS), alpha-gliadin (GLIA), or gamma-gliadin (GLIG).

**Supplementary Figure S6: Distribution of LC-MS1 data across 3,990 wheat samples and 32,336 quantified peptides.** (A) Histogram of the corrected dataset using a linear model and keeping the residuals; (B) Boxplot of corrected dataset log10 transformed for display purpose; (C) Histogram of the corrected dataset z-transformed per row of peptides; (D) Boxplot of z-transformed dataset log10 transformed for display purpose. Insets in panels A-B indicate one-sample Kolmogorov-Smirnov (K-S) test results where D is the value of the K-S statistics. **Supplementary Figure S7: Partial Least Square (PLS) using LMA as a response on the unbiased samples and the unbiased samples and all the quantified peptides.** (A) Score plot of Component 1 vs Component 2 of the 934 unbiased samples coloured based on LMA measurements; samples with high LMA are circled; (B) Loading plot of Component 1 vs Component 2 of the 32,337 peptides coloured based on PLS VIP scores; peptides with high LMA are circled; Cluster_AAA resolves in the top right corner and contributes the most to the PLS with a VIP score of 38.84.

**Supplementary Figure S8: Partial least square regression (PLSR) to impute LMA missing values.** (A) Full scatterplot of the measured vs. predicted LMA values of the testing set containing 179 samples; (B) same as panel A but limiting LMA predicted values inferior to 0.17 u/g; (C) same as panel A but limiting LMA predicted values superior to 0.17 u/g; (D) Line chart of the 217 LMA missing values and predicted by our PLSR model and sorted based on increasing LMA.

**Supplementary Figure S9: Binning strategies of wheat samples based on LMA measurements.** (A) all 3990 wheat samples were sorted by increasing order of LMA values and then split into 8 arbitrary bins of 499 samples each; the line chart displays bin averages; (B) the 934 unbiased wheat samples were sorted by increasing order of LMA values and then split into 2 arbitrary bins of 467 samples each based on a LMA value threshold of 0.17 u/g; the histogram displays bin averages. Bins are listed in Supplementary Table S1.

**Supplementary Figure S10: Mining identified proteins using Power BI.** (A) all identified peptides plotted as peptide mass against Mascot peptide scores (dot histogram), peptide missed cleavages (pie chart), peptide PTMs (tree map), peptide lengths (violin plot), peptide charges (vertical bar plot), protein score against sequence coverage (scatterplot) and protein description (word cloud); (B) same charts but drilled down on the term “inhibitor” in the word cloud of protein descriptions; (C) same charts but drilled down on “deamidated” peptides in the tree map of PTMs.

**Supplementary Figure S11: Retrieval of protein descriptions and Gene Ontology (GO) terms for Molecular Function (MF), Cellular Component (CC), and Biological Process (BP) from UniProtKB using all 8,044 protein identities.** (A) UniprotKB output viewed by GO; (B) word cloud of all protein names; (C) word cloud of protein names filtered as “glutenin”; (D) word cloud of protein names filtered as “domain-containing”; (E) tree map of the most abundant terms for GOBP category; (F) tree map of the most abundant terms for GOCC category; (E) tree map of the most abundant terms for GOMF category.

**Supplementary Figure S12: KEGG output using all 8,044 identified proteins matching 677 KOs.** (A) Histogram of the most frequent pathways; (B) Histogram of the most frequent brite terms; (C) Histogram of the most frequent modules; (D) Carbon metabolism map; (E) Glycolysis/gluconeogenesis map; (F) Starch and sucrose metabolism map. Proteins identified in this study are highlighted in green in panels D-F.

**Supplementary Figure S13: ShinyGO outputs using all 6,622 TRAES accessions corresponding to the 8,044 UniProt proteins.** (A) dot plot of the GO categories sorted by fold enrichment; (B) network of nodes representing enriched GO terms. Related GO terms are connected by a line, whose thickness reflects percent of overlapping genes. Node size represents the number of genes;(C-D) statistical analysis on the genomic features. Chi-squared and Student’s t-tests are run to compare the user’s genes to the T. aestivum genome. Results on number of exons, transcript isoforms, GC content, untranslated region (UTR) length, and types of genes (coding, non-coding, pseudogenes) are displayed as density scatterplots or histograms; (E) hierarchical clustering tree of significant enriched pathways. Pathways that share many genes are clustered together and dot size indicates q-values significance; (F) Plot of the chromosomal positions of the genes encoding our identified proteins.

**Supplementary Figure S14: Pathway Tools output using 6622 TRAES accessions and quantitative data averaged along 8 bins.** (A) OMICS dashboard general view; (B) cellular view zoomed in on TCA cycle II and glyoxylate cycle. Each expression profile points to a unique TRAES accession, most of them being homologous. The whole cellular view is available in Supplementary Video SV1; (C) OMICS dashboard zoomed in on hormone biosynthesis; (D) OMICS dashboard zoomed in on gibberellin and gibberellin precursor biosynthesis; (E) Pathway view of ent-kaurene biosynthesis from the Gibberellin biosynthesis pathway further illustrating high homology of wheat proteins; (F) 8-bin profiles of peptide biomarkers belonging to proteins involved in phytohormone biosynthesis; Cluster_AAA is displayed for comparison purpose.

## Graphical abstract

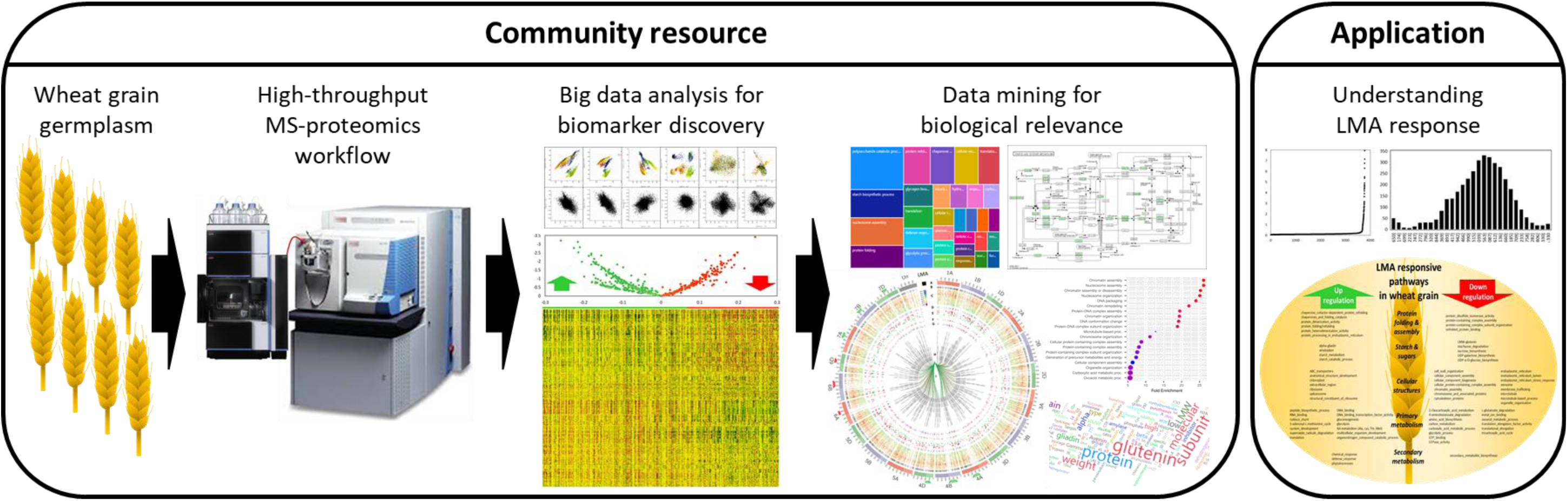

## Notes

### Competing Interest Statement

The authors have declared no competing interest.

