## Supplementary figures for "Finding the LMA needle in the wheat proteome haystack"

Vincent D. et al.

### Suppl. Figures

**Vincent et al – Supplementary Figure S1: Genedata Refiner workflow to process all wheat, IS and QC LCMS1 RAW files and export them to Genedata Analyst.** A. Step 1; B. Repetition node from Step 1; C. Step 2; D. Analyst setup. See Materials and Methods for description.

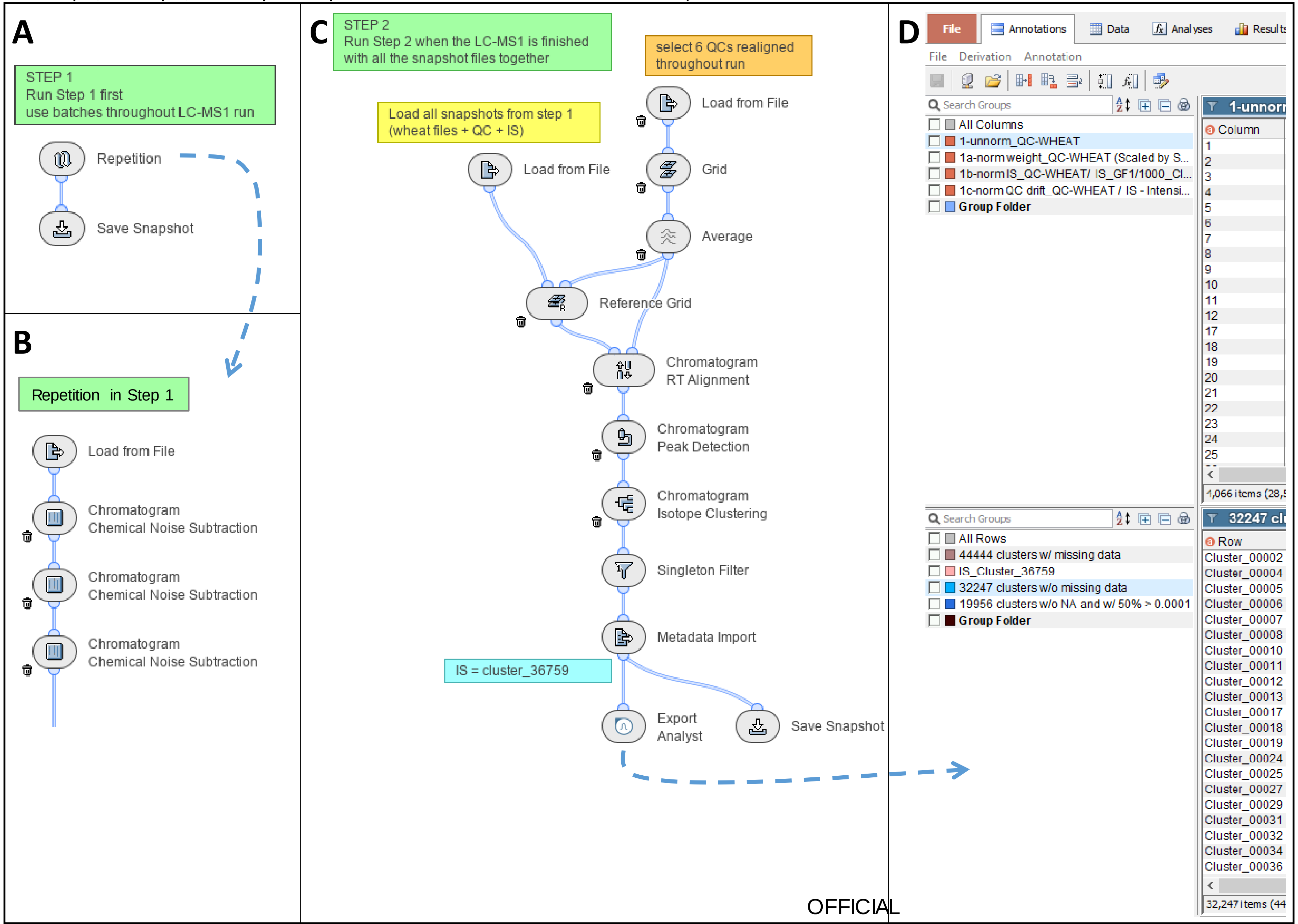


OFFICIAL

**Vincent et al – Supplementary Figure S2: Genedata Refiner workflow to process all wheat LCMS2 RAW files and export them to Excel.** A. Step 1; B. Repetition node from Step 1; C. Step 2; D. Mascot parameters; E. Excel output. See Materials and Methods for description.

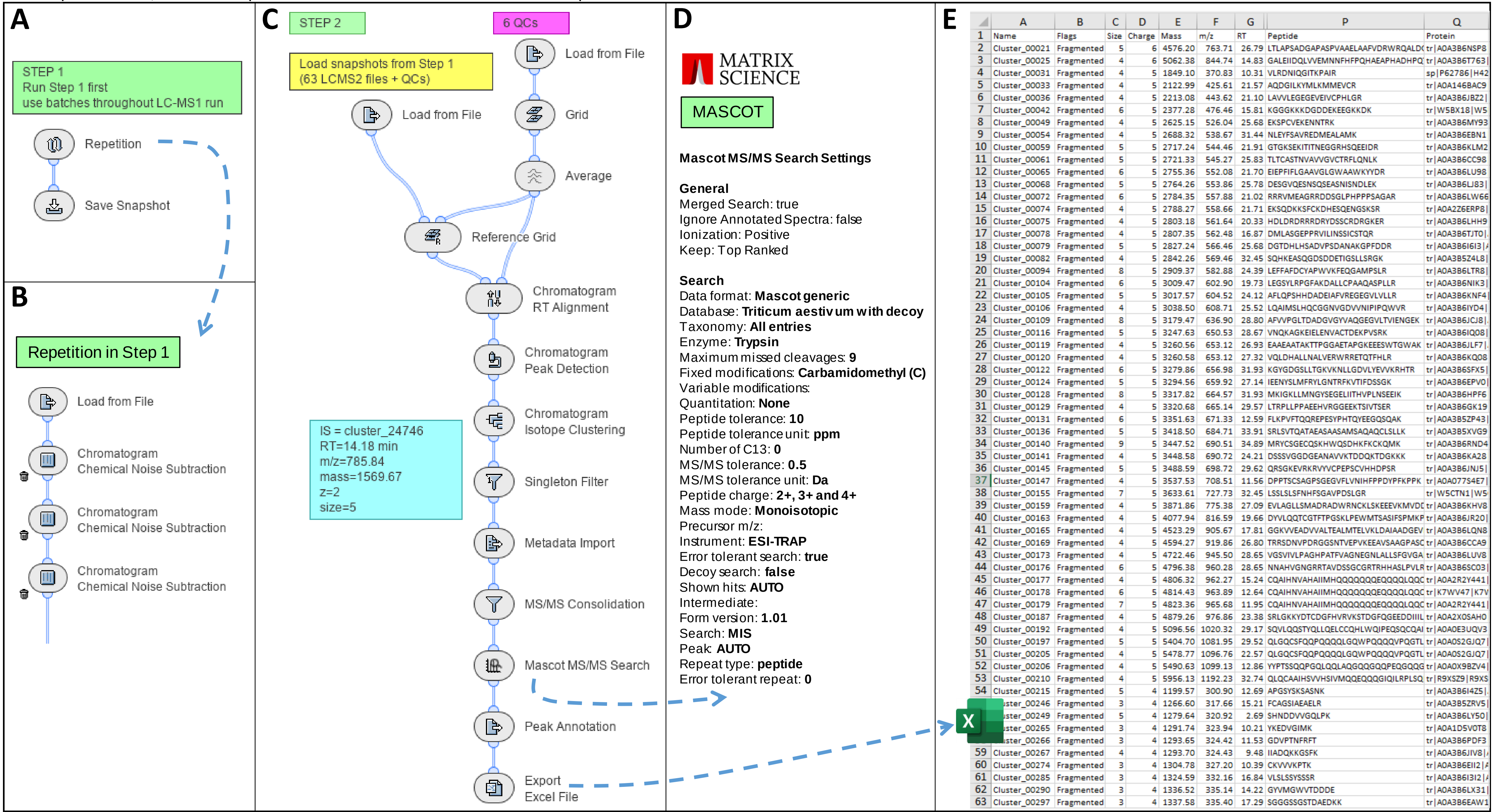


| | A | B | C | D | E | F | G | P | Q |
|---|---|---|---|---|---|---|---|---|---|
| 1 | Name | Flags | Size | Charge | Mass | m/z | RT | Peptide | Protein |
| 2 | Cluster_00021 | Fragmented | 5 | 6 | 4576.20 | 763.71 | 26.79 | LTLAPSADGAPASPVAAELAAFVDRWRQALD | tr\|A0A3B6NSP8 |
| 3 | Cluster_00025 | Fragmented | 4 | 6 | 5062.38 | 844.74 | 14.83 | GALEIIDQLVVEMNNFHFPQHAEAPHADHPQ | tr\|A0A3B6T763\| |
| 4 | Cluster_00031 | Fragmented | 4 | 5 | 1849.10 | 370.83 | 10.31 | VLRDNIQGITKPAIR | sp\|P62786\|H42 |
| 5 | Cluster_00033 | Fragmented | 4 | 5 | 2122.99 | 425.61 | 21.57 | AQDGILKYMLKMMEVCR | tr\|A0A146BAC9 |
| 6 | Cluster_00036 | Fragmented | 4 | 5 | 2213.08 | 443.62 | 21.10 | LAVVLEGEGEVEIVCPHLGR | tr\|A0A3B6JBZ2\| |
| 7 | Cluster_00042 | Fragmented | 6 | 5 | 2377.28 | 476.46 | 15.81 | KGGGKKKDGDDEKEEGKKDK | tr\|W5BX18\|W5 |
| 8 | Cluster_00049 | Fragmented | 4 | 5 | 2625.15 | 526.04 | 25.68 | EKSPCVEKENNTRK | tr\|A0A3B6MY93 |
| 9 | Cluster_00054 | Fragmented | 4 | 5 | 2688.32 | 538.67 | 31.44 | NLEYFSAVREDMEALAMK | tr\|A0A3B6EBN1 |
| 10 | Cluster_00059 | Fragmented | 5 | 5 | 2717.24 | 544.46 | 21.91 | GTGKSEKITITNEGGRHSQEEIDR | tr\|A0A3B6KLM2 |
| 11 | Cluster_00061 | Fragmented | 5 | 5 | 2721.33 | 545.27 | 25.83 | TLTCASTNVAVVGVCTRFLQNLK | tr\|A0A3B6CC98 |
| 12 | Cluster_00065 | Fragmented | 6 | 5 | 2755.36 | 552.08 | 21.70 | EIEPFIFLGAAVGLGWAAWKYYDR | tr\|A0A3B6LU98 |
| 13 | Cluster_00068 | Fragmented | 5 | 5 | 2764.26 | 553.86 | 25.78 | DESGVQESNSQSEASNISNDLEK | tr\|A0A3B6LJ83\| |
| 14 | Cluster_00072 | Fragmented | 6 | 5 | 2784.35 | 557.88 | 21.02 | RRRVMEAGRRDDSGLPHPPPSAGAR | tr\|A0A3B6LW66 |
| 15 | Cluster_00074 | Fragmented | 4 | 5 | 2788.27 | 558.66 | 21.71 | EKSQDKKSFCKDHESQENGSKSR | tr\|A0A2Z6ERP8\| |
| 16 | Cluster_00075 | Fragmented | 4 | 5 | 2803.18 | 561.64 | 20.33 | HDLDRDRRRDRYDSSCRDRGKER | tr\|A0A3B6LHH9 |
| 17 | Cluster_00078 | Fragmented | 4 | 5 | 2807.35 | 562.48 | 16.87 | DMLASGEPPRVILINSSICSTQR | tr\|A0A3B6TJT0\| |
| 18 | Cluster_00079 | Fragmented | 5 | 5 | 2827.24 | 566.46 | 25.68 | DGTDHLHSADVPSDANAKGPFDDR | tr\|A0A3B6I6I3\| |
| 19 | Cluster_00082 | Fragmented | 4 | 5 | 2842.26 | 569.46 | 32.45 | SQHKEASQGDSDDETIGSLLSRGK | tr\|A0A3B5Z4L8\| |
| 20 | Cluster_00094 | Fragmented | 8 | 5 | 2909.37 | 582.88 | 24.39 | LEFFAFDCYAPWVKFEQGAMPSLR | tr\|A0A3B6LTR8\| |
| 21 | Cluster_00104 | Fragmented | 6 | 5 | 3009.47 | 602.90 | 19.73 | LEGSYLRPGFAKDALLCPAAQASPLLR | tr\|A0A3B6NIK3\| |
| 22 | Cluster_00105 | Fragmented | 5 | 5 | 3017.57 | 604.52 | 24.12 | AFLQPSHHDADEIAFVREGEGVLVLLR | tr\|A0A3B6KNF4\| |
| 23 | Cluster_00106 | Fragmented | 4 | 5 | 3038.50 | 608.71 | 25.52 | LQAIMSLHQCGGNVGDVVNIPIPQWVR | tr\|A0A3B6IYD4\| |
| 24 | Cluster_00109 | Fragmented | 8 | 5 | 3179.47 | 636.90 | 28.80 | AFVVPGLTDADGVGYVAQGEGVLTVIENGEK | tr\|A0A3B6JCJ8\| |
| 25 | Cluster_00116 | Fragmented | 5 | 5 | 3247.63 | 650.53 | 28.67 | VNQKAGKEIELENVACTDEKPVSRK | tr\|A0A3B6IQ08\| |
| 26 | Cluster_00119 | Fragmented | 4 | 5 | 3260.56 | 653.12 | 26.93 | EAAEAATAKTTPGGAETAPGKEEESWTGWAK | tr\|A0A3B6JLF7\| |
| 27 | Cluster_00120 | Fragmented | 4 | 5 | 3260.58 | 653.12 | 27.32 | VQLDHALLNALVERWRRETQTFHLR | tr\|A0A3B6KQ08 |
| 28 | Cluster_00122 | Fragmented | 6 | 5 | 3279.86 | 656.98 | 31.93 | KGYGDGSLLTGKVKNLLGDVLYEVVKRHTR | tr\|A0A3B6SFX5\| |
| 29 | Cluster_00124 | Fragmented | 5 | 5 | 3294.56 | 659.92 | 27.14 | IEENYSLMFRYLGNTRFKVTIFDSSGK | tr\|A0A3B6EPV0\| |
| 30 | Cluster_00128 | Fragmented | 8 | 5 | 3317.82 | 664.57 | 31.93 | MKIGKLLMNGYSEGELIITHVPLNSEEIK | tr\|A0A3B6HPF6 |
| 31 | Cluster_00129 | Fragmented | 4 | 5 | 3320.68 | 665.14 | 29.57 | LTRPLLPPAEEHVRGGEEKTSIVTSER | tr\|A0A3B6GK19 |
| 32 | Cluster_00131 | Fragmented | 6 | 5 | 3351.63 | 671.33 | 12.59 | FLKPVFTQQREPESYPHTQYEEGQSQAK | tr\|A0A3B5ZP43\| |
| 33 | Cluster_00136 | Fragmented | 5 | 5 | 3418.50 | 684.71 | 33.91 | SRLSVTQATAEASAASAMSAQAQCLSLLK | tr\|A0A3B5XVG9 |
| 34 | Cluster_00140 | Fragmented | 9 | 5 | 3447.52 | 690.51 | 34.89 | MRYCSGECQSKHWQSDHKFKCKQMK | tr\|A0A3B6RND4 |
| 35 | Cluster_00141 | Fragmented | 4 | 5 | 3448.58 | 690.72 | 24.21 | DSSSVGGDGEANAVVKTDDQKTDGKKK | tr\|A0A3B6KA28 |
| 36 | Cluster_00145 | Fragmented | 5 | 5 | 3488.59 | 698.72 | 29.62 | QRSGKEVRKRVYVCPEPSCVHHDPSR | tr\|A0A3B6JNJ5\| |
| 37 | Cluster_00147 | Fragmented | 4 | 5 | 3537.53 | 708.51 | 11.56 | DPPTSCSAGPSGEGVFLVNIHFPPDYPFKPPK | tr\|A0A077S4E7\| |
| 38 | Cluster_00155 | Fragmented | 7 | 5 | 3633.61 | 727.73 | 32.45 | LSSLSLSFNHFSGAVPDSLGR | tr\|W5CTN1\|W5 |
| 39 | Cluster_00159 | Fragmented | 4 | 5 | 3871.86 | 775.38 | 27.09 | EVLAGLLSMADRADWRNCKLSKEEEVKMVD | tr\|A0A3B6KHV8 |
| 40 | Cluster_00163 | Fragmented | 4 | 5 | 4077.94 | 816.59 | 19.66 | DYVLQQTCGTFTPGSKLPEWMTSASIFSPMKP | tr\|A0A3B6JR20\| |
| 41 | Cluster_00165 | Fragmented | 4 | 5 | 4523.29 | 905.67 | 17.81 | GGKVVEADVVALTEALMTELVKLDAIAADGEV | tr\|A0A3B6LQN8 |
| 42 | Cluster_00169 | Fragmented | 4 | 5 | 4594.27 | 919.86 | 26.80 | TRRSDNVPDRGGSNTVEPVKEEAVSAAGPAS | tr\|A0A3B6CCA9 |
| 43 | Cluster_00173 | Fragmented | 4 | 5 | 4722.46 | 945.50 | 28.65 | VGSVIVLPAGHPATFVAGNEGNLALLSFGVGA | tr\|A0A3B6LUV8 |
| 44 | Cluster_00176 | Fragmented | 6 | 5 | 4796.38 | 960.28 | 28.65 | NNAHVGNGRRTAVDSSGCGRTRHHASLPVLR | tr\|A0A3B6SC03\| |
| 45 | Cluster_00177 | Fragmented | 4 | 5 | 4806.32 | 962.27 | 15.24 | CQAIHNVAHAIIMHQQQQQQEQQQLQQ | tr\|A0A2R2Y441\| |
| 46 | Cluster_00178 | Fragmented | 6 | 5 | 4814.43 | 963.89 | 12.64 | CQAIHNVAHAIIMHQQQQQQEQQQLQQ | tr\|K7WV47\|K7V |
| 47 | Cluster_00179 | Fragmented | 7 | 5 | 4823.36 | 965.68 | 11.95 | CQAIHNVAHAIIMHQQQQQQEQQQLQQ | tr\|A0A2R2Y441\| |
| 48 | Cluster_00187 | Fragmented | 4 | 5 | 4879.26 | 976.86 | 23.38 | SRLGKKYDTCDGFHVRVKSTDGFQGEEDDIIIL | tr\|A0A2X0SAH0 |
| 49 | Cluster_00192 | Fragmented | 4 | 5 | 5096.56 | 1020.32 | 29.17 | SQVLQQSTYQLLQELCCQHLWQIPEQSQCQAI | tr\|A0A0E3UQV3 |
| 50 | Cluster_00197 | Fragmented | 5 | 5 | 5404.70 | 1081.95 | 29.52 | QLGQCSFQQPQQQQLGQWPQQQQVPQGTL | tr\|A0A0S2GJQ7\| |
| 51 | Cluster_00205 | Fragmented | 4 | 5 | 5478.77 | 1096.76 | 22.57 | QLGQCSFQQPQQQQLGQWPQQQQVPQGTL | tr\|A0A0S2GJQ7\| |
| 52 | Cluster_00206 | Fragmented | 4 | 5 | 5490.63 | 1099.13 | 12.86 | YYPTSSQQPGQLQQLAQGQQGQQPEQGQQG | tr\|A0A0X9BZV4\| |
| 53 | Cluster_00210 | Fragmented | 4 | 5 | 5956.13 | 1192.23 | 32.74 | QLQCAAIHSVVHSIVMQQEQQQGIQILRPLSQ | tr\|R9XSZ9\|R9XS |
| 54 | Cluster_00215 | Fragmented | 5 | 4 | 1199.57 | 300.90 | 12.69 | APGSYSKSASNK | tr\|A0A3B6I4Z5\| |
| | ster_00246 | Fragmented | 3 | 4 | 1266.60 | 317.66 | 15.21 | FCAGSIAEAELR | tr\|A0A3B5ZRV5\| |
| | ster_00249 | Fragmented | 5 | 4 | 1279.64 | 320.92 | 2.69 | SHNDDVVGQLPK | tr\|A0A3B6LY50\| |
| | ster_00265 | Fragmented | 3 | 4 | 1291.74 | 323.94 | 10.21 | YKEDVGIMK | tr\|A0A1D5V0T8 |
| | ster_00266 | Fragmented | 3 | 4 | 1293.65 | 324.42 | 11.53 | GDVPTNFRFT | tr\|A0A3B6PDF3 |
| 59 | Cluster_00267 | Fragmented | 4 | 4 | 1293.70 | 324.43 | 9.48 | IIADQKKGSFK | tr\|A0A3B6JIV8\| |
| 60 | Cluster_00274 | Fragmented | 3 | 4 | 1304.78 | 327.20 | 10.39 | CKVVVKPTK | tr\|A0A3B6EII2\| |
| 61 | Cluster_00285 | Fragmented | 3 | 4 | 1324.59 | 332.16 | 16.84 | VLSLSSYSSSR | tr\|A0A3B6I3I2\| |
| 62 | Cluster_00290 | Fragmented | 3 | 4 | 1336.52 | 335.14 | 14.22 | GYVMGWVTDDDE | tr\|A0A3B6LX31\| |
| 63 | Cluster_00297 | Fragmented | 3 | 4 | 1337.58 | 335.40 | 17.29 | SGGGSSGSTDAEDKK | tr\|A0A3B6EAW1 |

**Vincent et al – Supplementary Figure S3: LC-MS2 RAW maps for each tandem pass.** X-axis depicts 300-2000 m/z. Y-axis depicts 1-35 min Retention Time. White dots represent MS2 events. (A) LC-MS1 map of pooled sample; (B) LC-MS2 map of Pass 1 replicate 1 with 3000 threshold; (C) LC-MS2 map of Pass 2 replicate 1 with exclusion list of 2000 ions fragmented in Pass 1; (D) LC-MS2 map of Pass 3 replicate 1 with exclusion list of 2000 ions fragmented in Pass 2; (E) LC-MS2 map of Pass 4 replicate 1 with exclusion list of 2000 ions fragmented in Pass 3; (F) LC-MS2 map of Pass 5 replicate 1 (same as Pass 1 but with 500 threshold); (G) LC-MS2 map of Pass 6 replicate 1 with inclusion list of 2000 most abundant ions from Pass 1; (H) LC-MS2 map of Pass 7 with inclusion list 1 loaded Global mass tab and 2 *m/z* tolerance; (I) LC-MS2 map of Pass 8 with inclusion list 1 loaded in data-dependent settings and 2 *m/z* tolerance; (J) LC-MS2 map of Pass 9 with inclusion list 1 loaded in data-dependent settings and 1 *m/z* tolerance; (K) LC-MS2 map of Pass 10 with inclusion list 1 loaded in data-dependent settings and 0.5 *m/z* tolerance; (L) LC-MS2 map of Pass 11 with inclusion list 1 loaded in data-dependent settings and 0.2 *m/z* tolerance. Maps with inclusion lists 2-10 for Passes 7-11 are not shown.

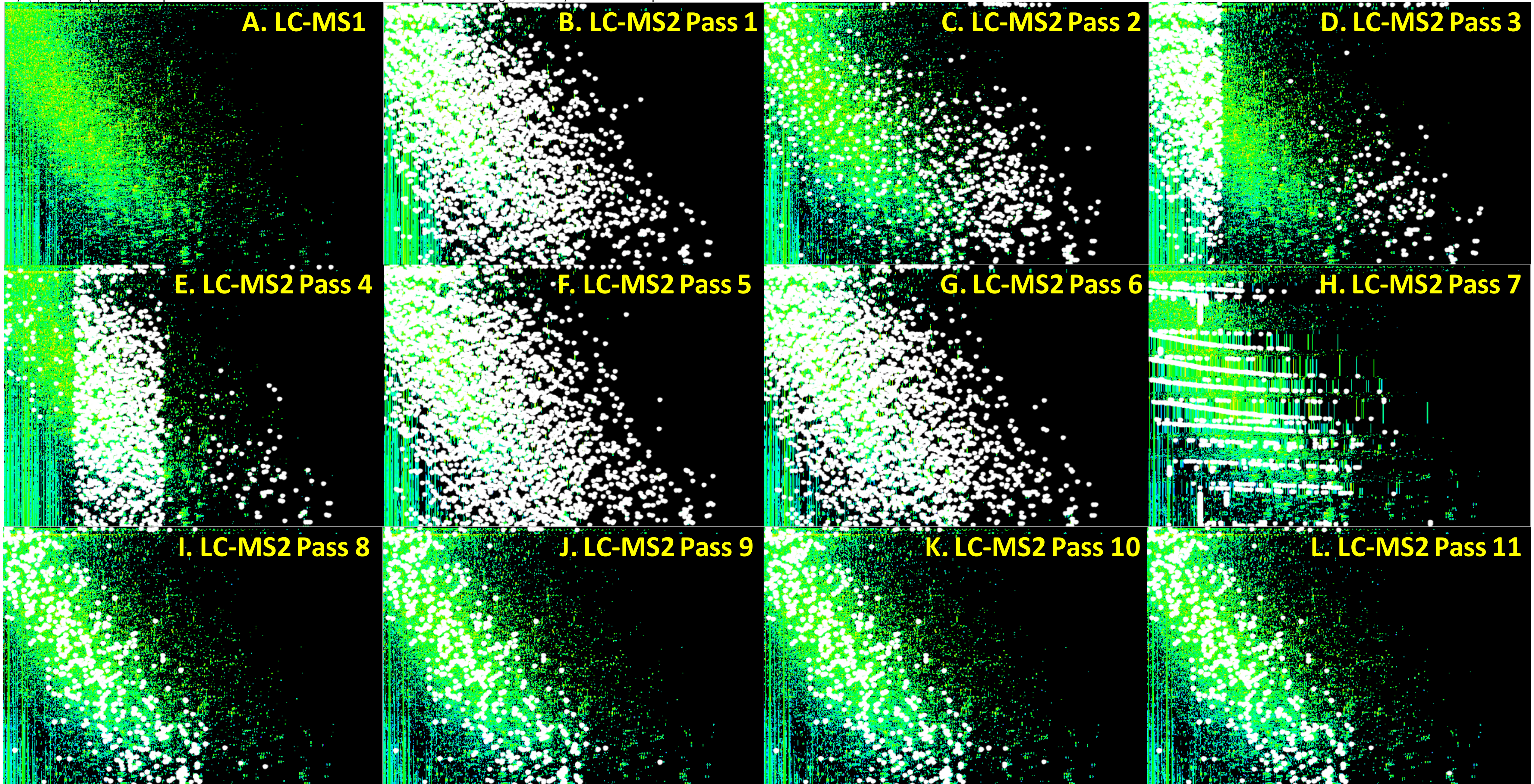

**Vincent et al – Supplementary Figure S4: Histogram of the number of peptides identified using Mascot algorithm and number of MS2 events in each of the LC-MS2 file.** Black bars represent peptide counts (y axis on the left) and orange dots depict MS/MS event counts (y axis on the right).

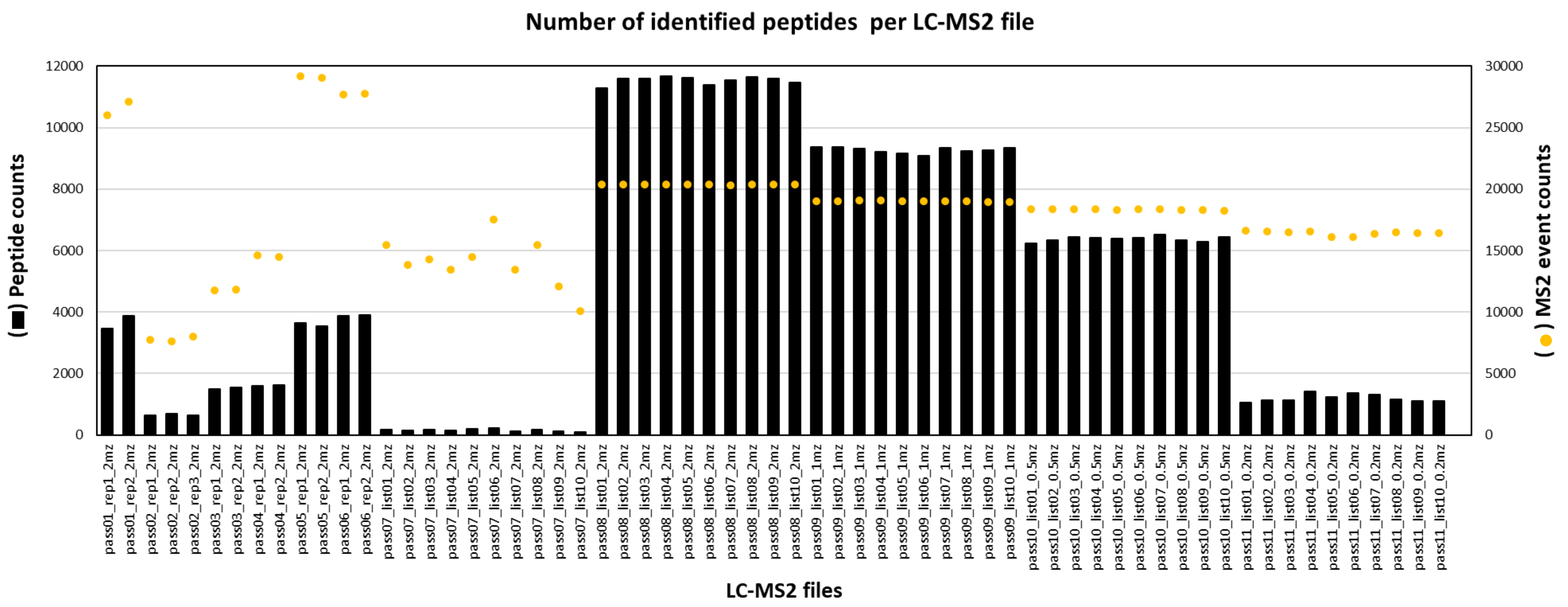

**Vincent et al – Supplementary Figure S5: Histograms (A, C, E) and box plots (B, D) of the number of peptides per accession (A-B, E) and number of accessions per peptides (C-D).** The orange line in panels A and C represents cumulated counts in percent. Panel E displays the peptides with the highest hit counts belonging either to low molecular weight glutenin subunit (LMW-GS) or alpha-gliadin (GLIA).

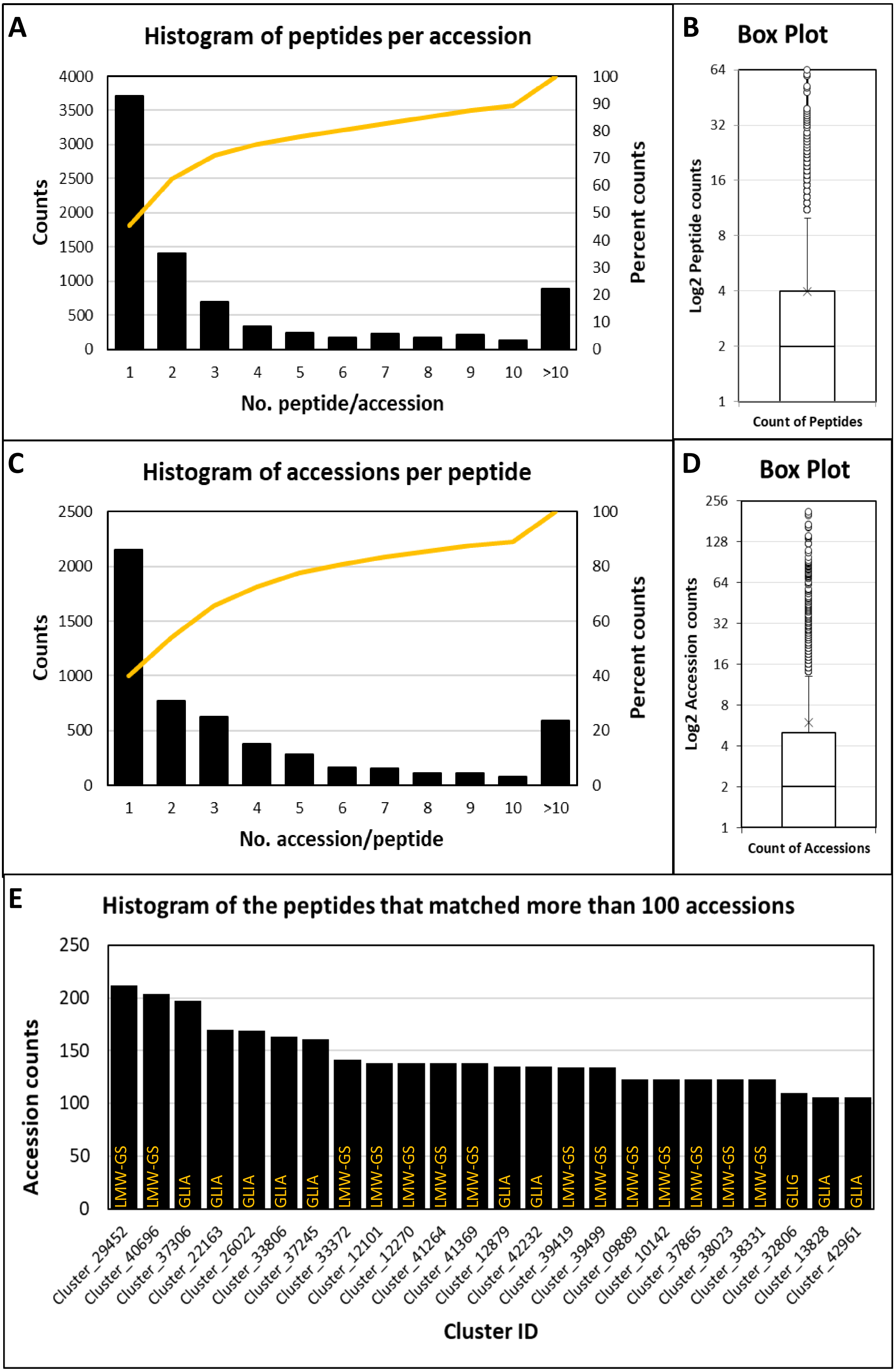

**Vincent et al – Supplementary Figure S6: Distribution of LC-MS1 data across 3,990 wheat samples and 32,336 quantified peptides.** (A) Histogram of the corrected dataset using a linear model and keeping the residuals; (B) Boxplot of corrected dataset log10 transformed for display purpose; (C) Histogram of the corre cted dataset z-transformed per row of peptides; (D) Boxplot of z-transformed dataset log10 transformed for display purpose. Insets in panels A-B indicate One-sample Kolmogorov-Smirnov (K-S) test results where D is the value of the K-S statistics.

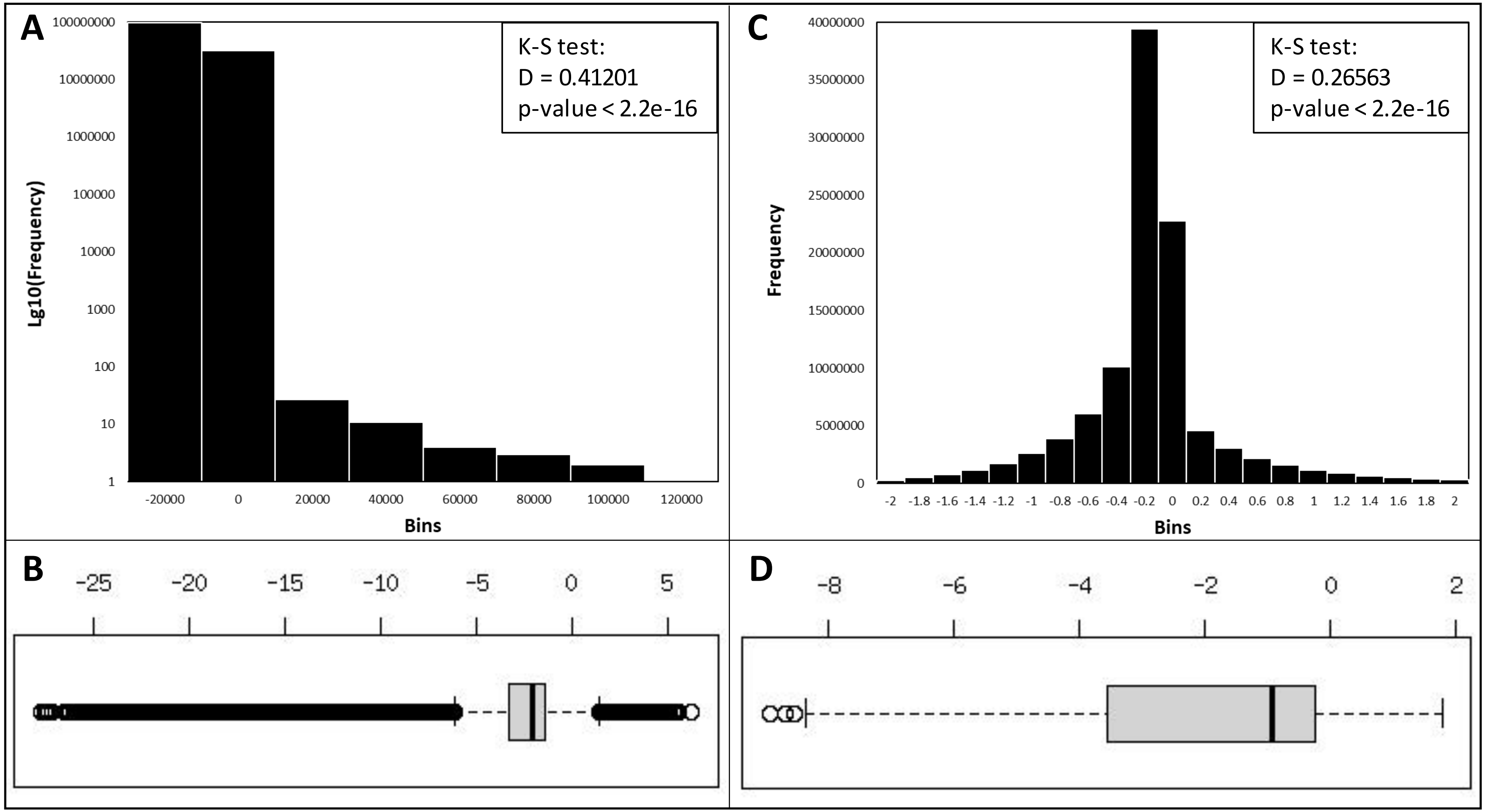

**Vincent et al – Supplementary Figure S7: Partial Least Square (PLS) using LMA as a response on the unbiased samples and all the quantified peptides.** (A) Score plot of PC1 vs PC2 of the 934 unbiased samples coloured based on LMA measurements; (B) Loading plot of PC1 vs PC2 of the 32,337 peptides coloured based on PLS VIP scores; Cluster_AAA resolves in the top right corner and contributes the most to the PLS with a VIP score of 38.84.

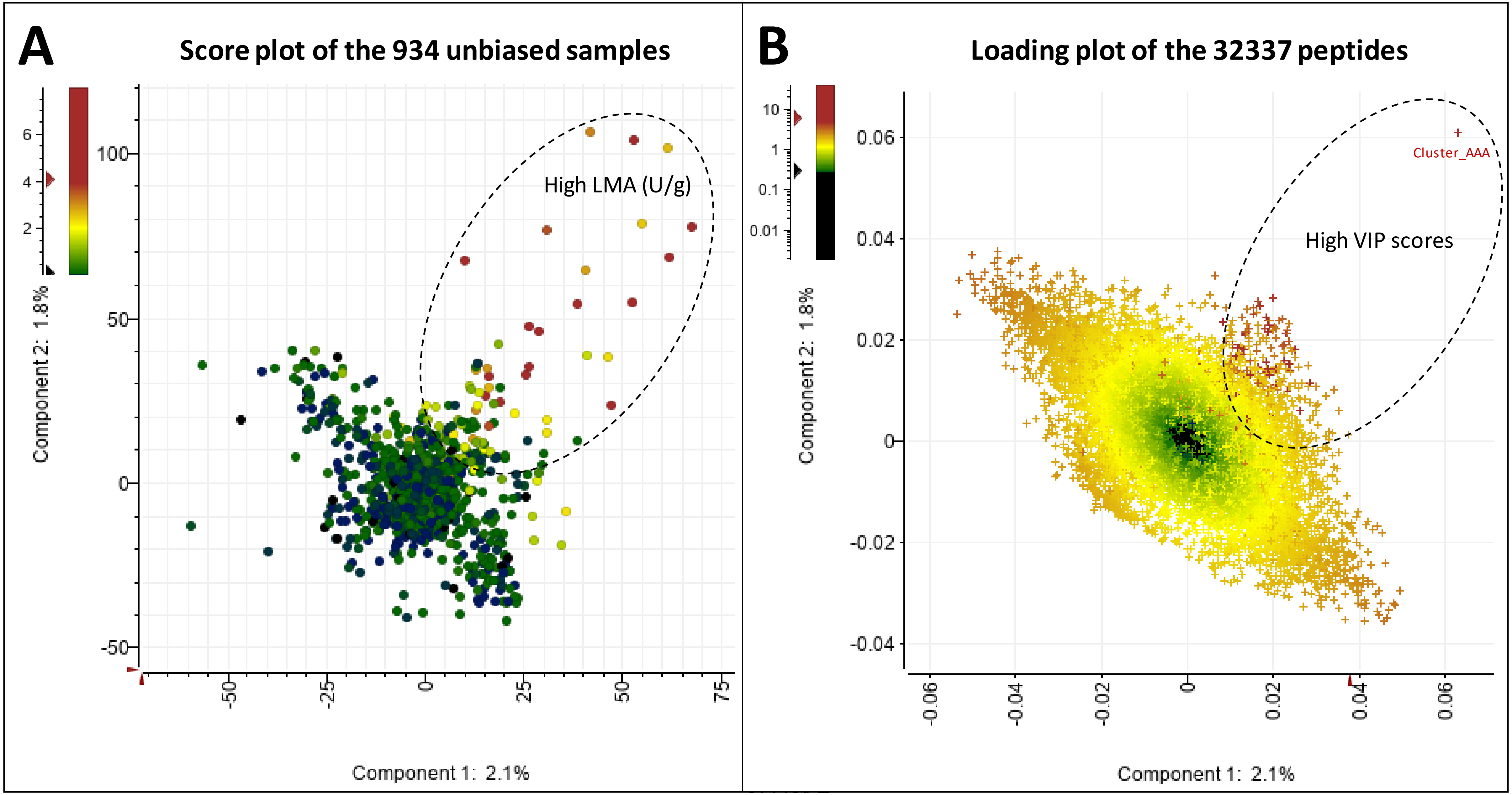

**Vincent et al – Supplementary Figure S8: Partial least square regression (PLSR) to impute LMA missing values.** (A) Full scatterplot of the measured vs. predicted LMA values of the testing set containing 174 samples; (B) same as panel A but limiting LMA predicted values inferior to 0.2; (C) same as panel A but limiting LMA predicted values superior to 0.2; (D) Line chart of the 217 LMA missing values and predicted by our PLSR model.

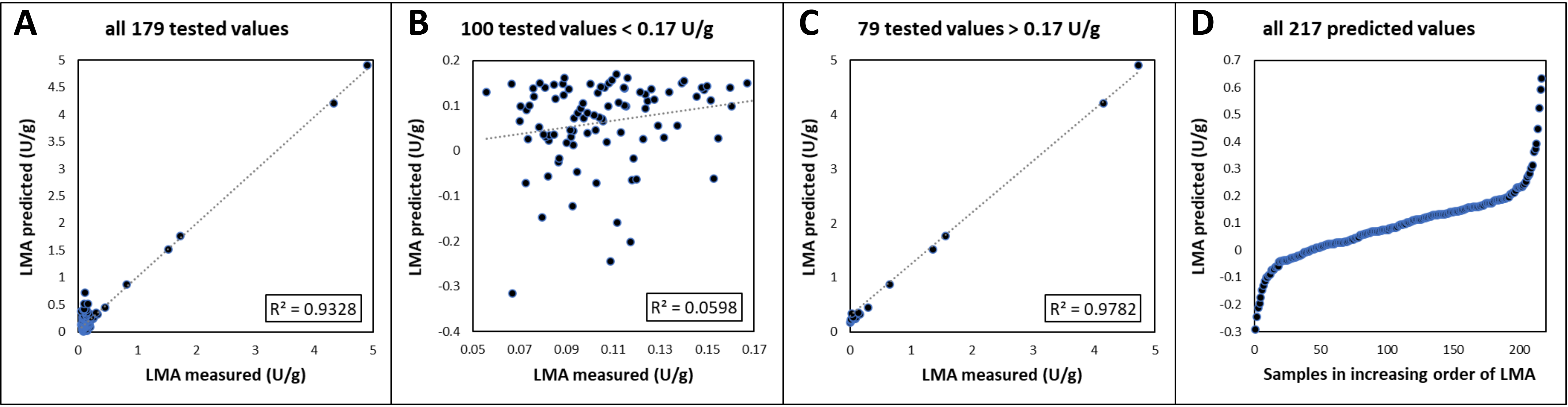

**Vincent et al – Supplementary Figure S9: Binning strategies of wheat samples based on LMA measurements.** (A) all 3990 wheat samples were sorted by increasing order of LMA values and then split into 8 arbitrary bins of 499 samples each; the line chart displays bin averages; (B) the 934 unbiased wheat sample s were sorted by increasing order of LMA values and then split into 2 arbitrary bins of 467 samples each based on a LMA value threshold of 0.17; the histogram displays bin averages.

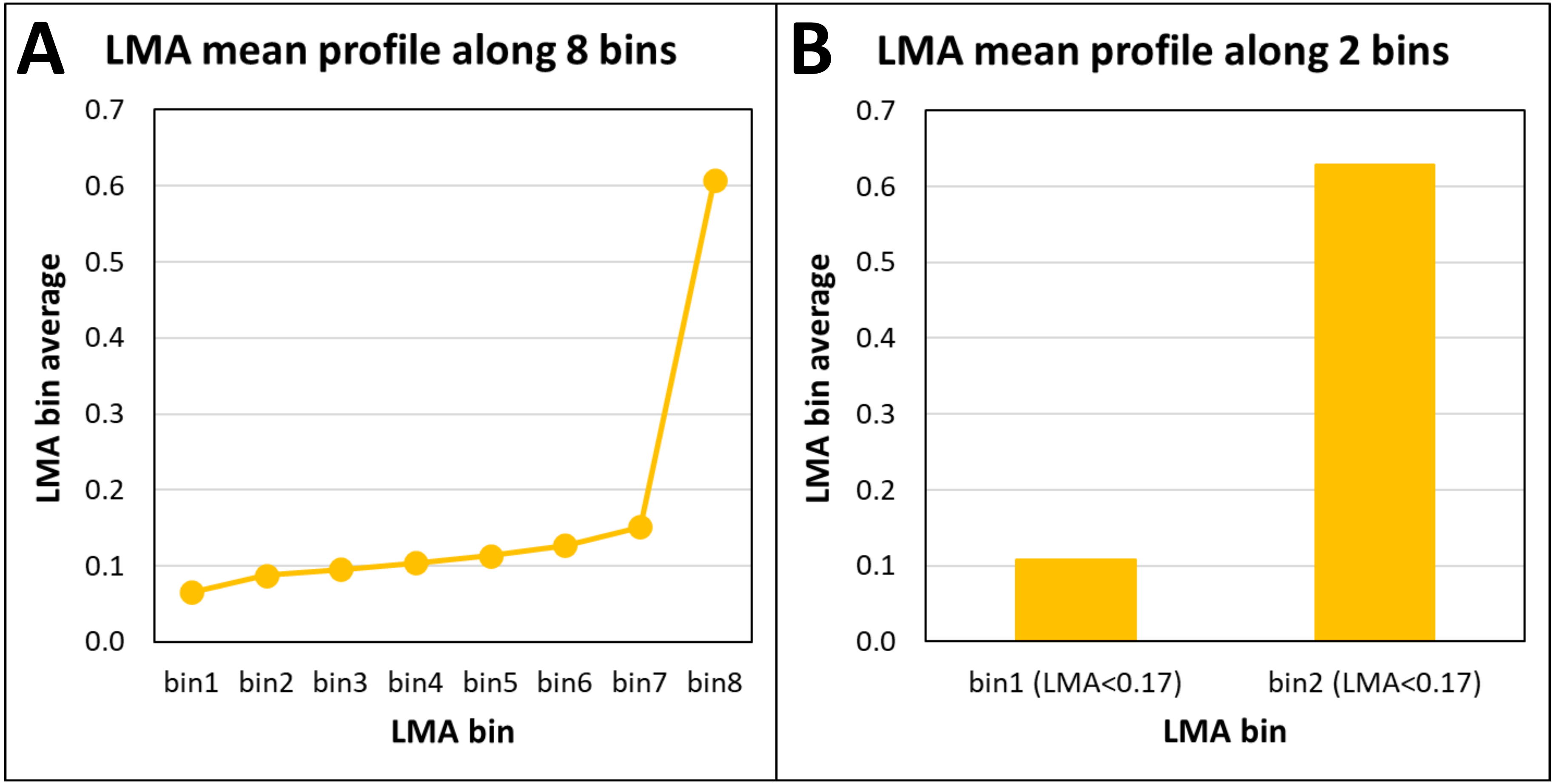

**Vincent et al – Supplementary Figure S10: Mining identified proteins using Power BI** (A) all identified peptides plotted as peptide mass against Mascot peptide scores (dot histogram), peptide missed cleavages (pie chart), peptide PTMs (tree map), peptide lengths (violin plot), peptide charges (vertical bar plot), protein score against sequence coverage (scatterplot) and protein description (word cloud); (B) same charts but drilled down on the term "inhibitor" in the word cloud of protein descriptions; (C) same charts but drilled down on "deamidated" peptides in the tree map of PTMs.

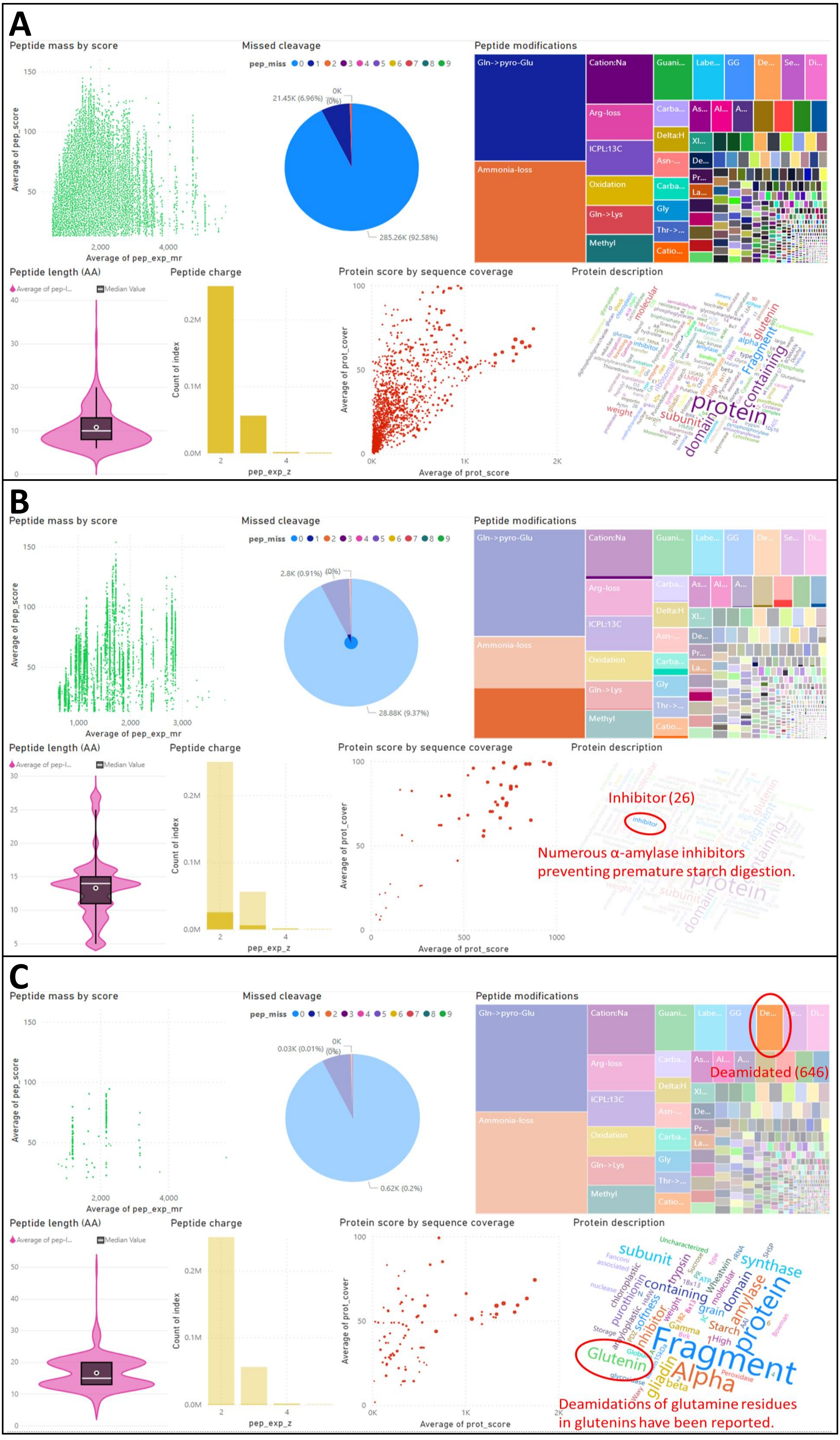

**Vincent et al – Supplementary Figure S11: Retrieval of protein descriptions and Gene Ontology (GO) terms for Molecular Function (MF), Cellular Component (CC), and Biological Process (BP) from UniProtKB using all 8,044 protein identities.** (A) UniprotKB output viewed by GO; (B) word cloud of all protein names; (C) word cloud of protein names filtered as "glutenin"; (D) word cloud of protein names filtered as "domain-containing"; (E) tree map of the most abundant terms for GOBP category; (F) tree map of the most abundant terms for GOCC category; (E) tree map of the most abundant terms for GOMF category.

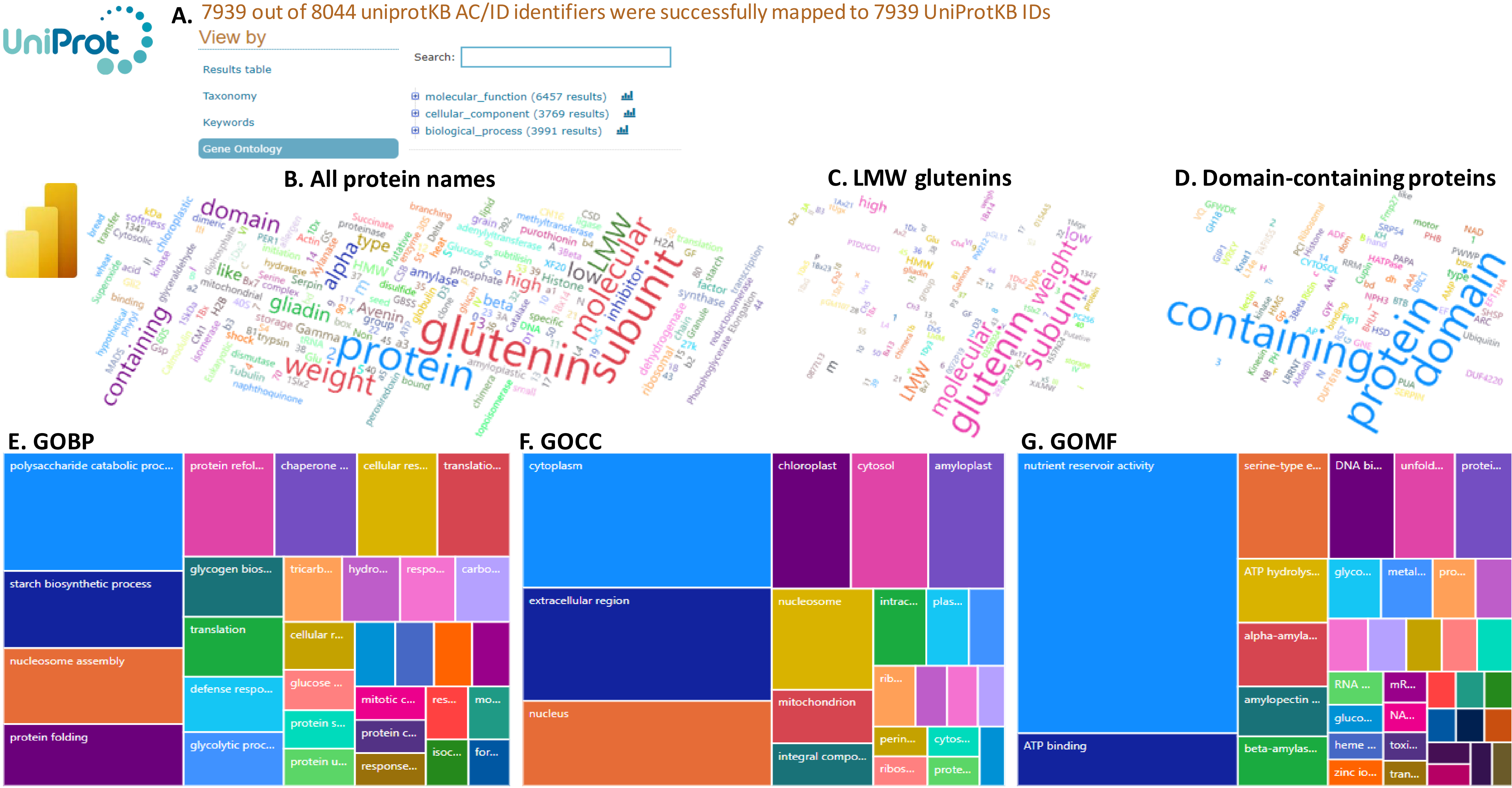

**Vincent et al – Supplementary Figure S12: KEGG output using all 8,044 identified proteins matching 677 KOs.** (A) Histogram of the most frequent pathways; (B) Histogram of the most frequent brite terms; (C) Histogram of the most frequent modules; (D) Carbon metabolism map; (E) Glycolysis/gluconeogenesis map; (F) Starch and sucrose metabolism map. Proteins identified in this study are highlighted in green in panels D-F.

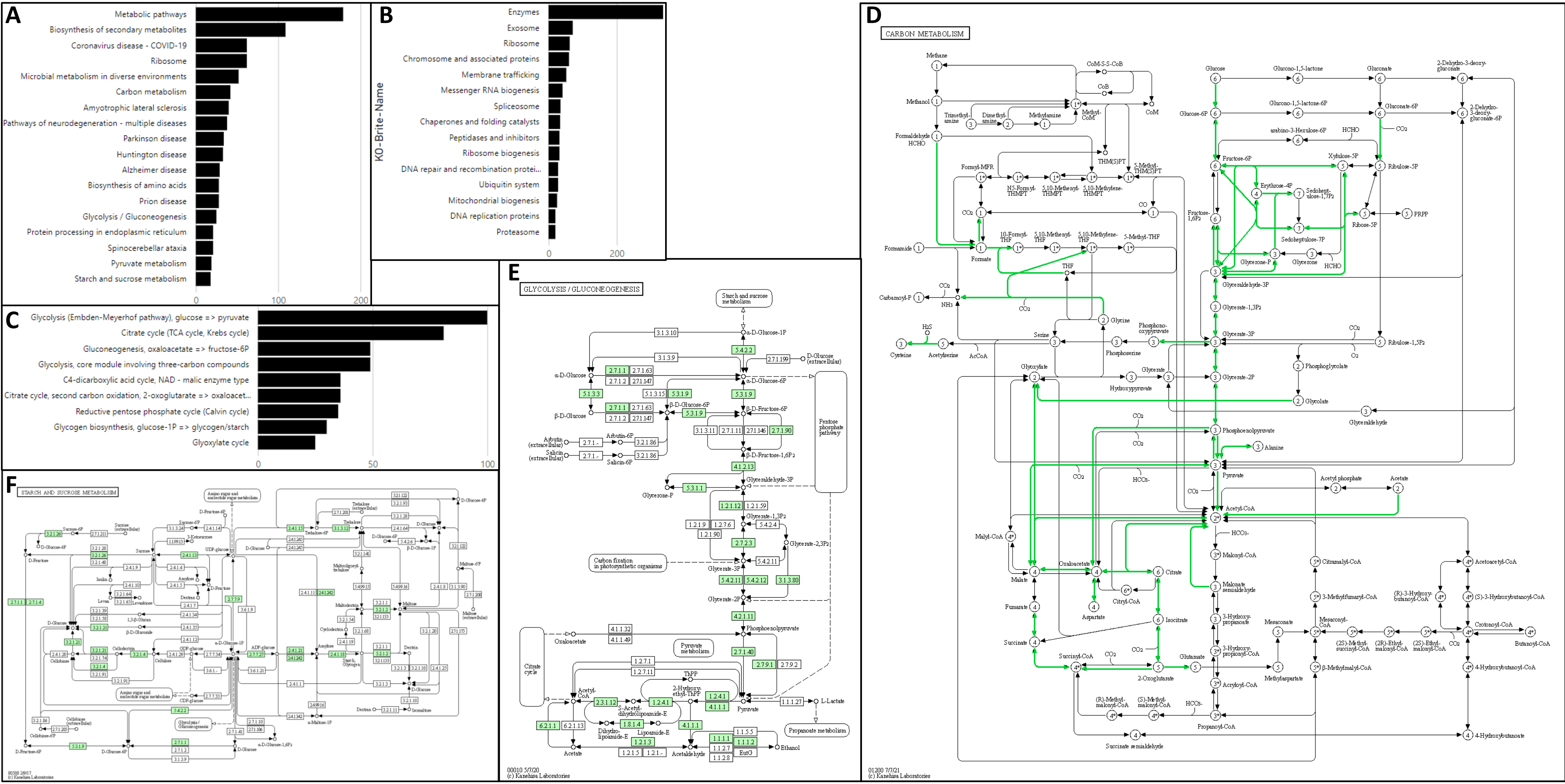

**Vincent et al – Supplementary Figure S13: ShinyGO outputs using all 6,622 TRAES accessions corresponding to the 8,044 UniProt proteins.** (A) dot plot of the GO categories sorted by fold enrichment; (B) network of nodes representing enriched GO terms. Related GO terms are connected by a line, whose thickness reflects percent of overlapping genes. Node size represents the number of genes; (C-D) statistical analysis on the genomic features. Chi-squared and Student's t-tests are run to compare the user's genes to the T. aestivum genome. Results on number of exons, transcript isoforms, GC content, UTR length, and types of genes (coding, non-coding, pseudogenes) are displayed as density scatterplots or histograms; (E) hierarchical clustering tree of significant enriched pathways. Pathways that share many genes are clustered together and dot size indicates P-values significance; (F) Plot of the chromosomal positions of the genes encoding our identified proteins.

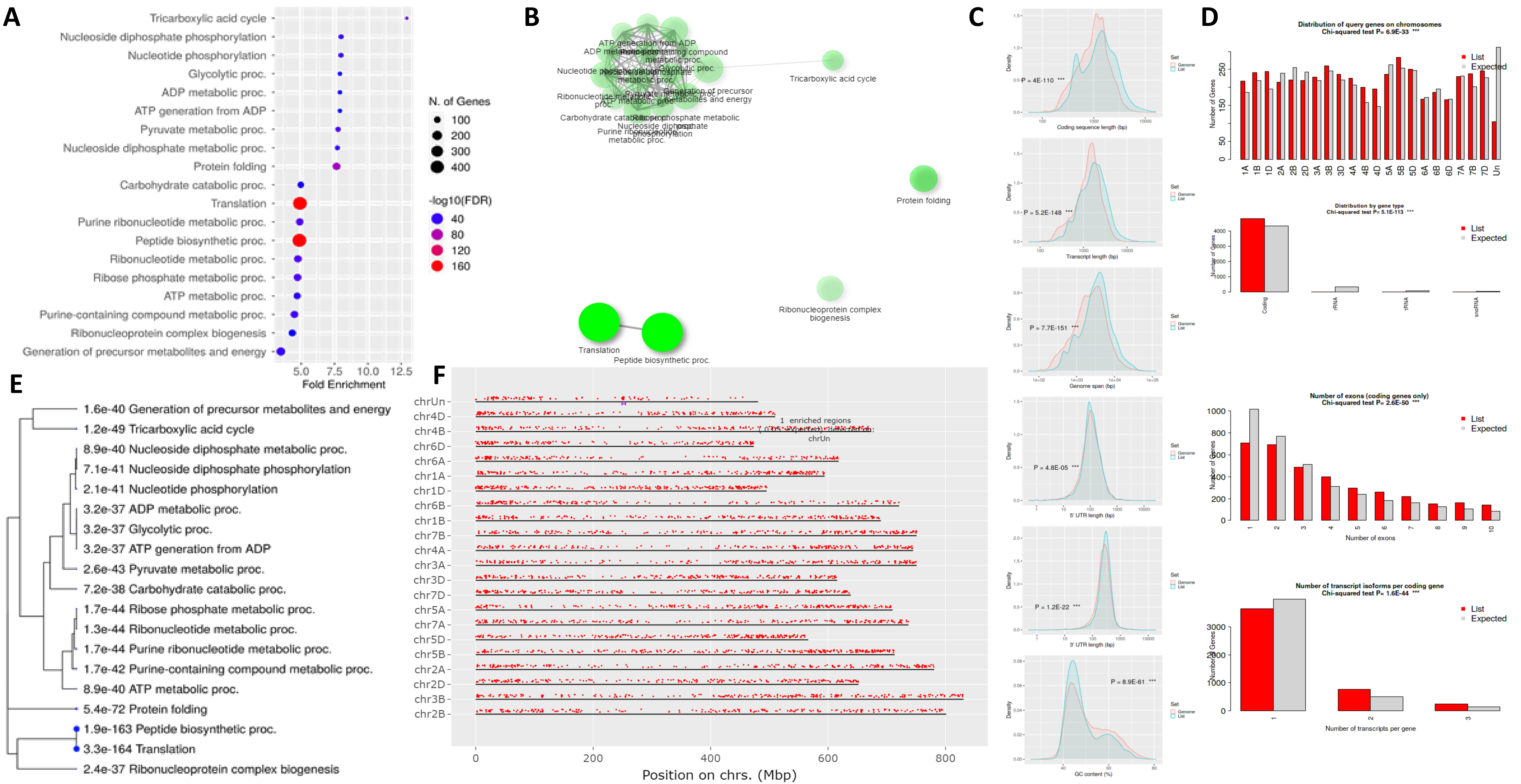

**Vincent et al – Supplementary Figure S14: Pathway Tools output using 6622 TRAES accessions and quantitative data averaged along 8 bins.** (A) OMICS dashboard general view; (B) cellular view zoomed in on TCA cycle II and glyoxylate cycle; (C) OMICS dashboard zoomed in on hormone biosynthesis; (D) OMICS dashboard zoomed in on gibberellin and gibberellin precursor biosynthesis; (E) Pathway view of ent-kaurene biosynthesis; (F) 8-bin profiles of peptide biomarkers belonging to proteins involved in phytohormone biosynthesis.

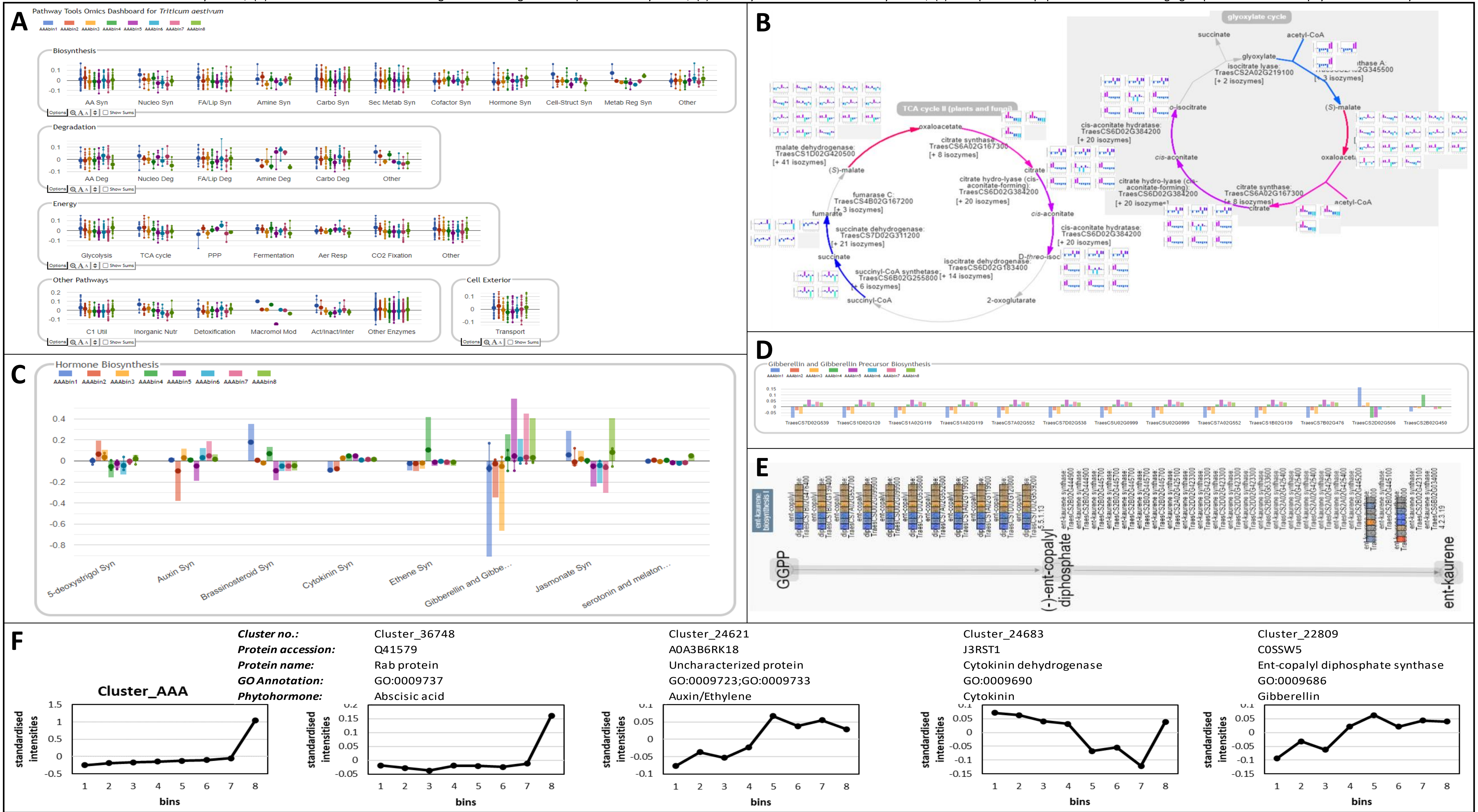
