## Supplementary methods for "Finding the LMA needle in the wheat proteome haystack"

### Vincent et al

### Proteomics of Wheat 4000 lines

### Supplementary File SF1

### 1. High-throughput proteomics workflow to efficiently process and analyse thousands of samples

Samples were packaged in small plastic and randomised by mixing them all up in a big box. The QR codes were scanned to populate an Excel spreadsheet assigning a unique number to each individual sample against sample ID. The Excel spreadsheet was readily accessible and populated on the fly by all involved using Microsoft Sharepoint from any laptop in the lab along with the SOP and other relevant documentation. Every workflow step was captured in that single Excel spreadsheet with the date, experimenter, parameters and comments where applicable. It contained 4,400 rows and 110 columns and was used as sample metadata during the statistical analyses and data correction methods.

The full content of a package was poured into a numbered 15 mL grinding jar with metal balls. Jars were arranged into two cardboard boxes to grind 200 samples per day. The samples were ground in batches of 36 samples simultaneously using a Genogrinder. Finely ground flour was transferred from the grinding jar into a microtube prelabelled with printed labels displaying the unique sample number for tracking purpose and a QR code capturing it along with the unique sample ID. Microtubes containing flour samples were stored in cardboard boxes fitting 81 tubes at -80°C until weighing and extraction. The grinding jars and metal balls were thoroughly washed to be re-used.

The thawed flour was individually and manually weighed with 1% accuracy into a prelabelled microtube. The flour was resuspended into an extraction buffer using a sonicating probe, a vortex and heating blocks in batches of 48 samples per day. Solubilised flours were spun 24 tubes at a time using two benchtop centrifuges; supernatants transferred into prelabelled microtubes. Microtubes containing protein extracts were stored in cardboard boxes fitting 81 tubes at -80°C until protein digestion.

A set of 192 frozen protein extracts were thawed overnight at 4°C and aliquoted (10 µL) into two 96-well plates for Trypsin/Lys-C digestion overnight. The IS was introduced at the end of the digestion step. Each digestion plate had a unique plate number that was tracked against all individual samples it contained. The plates were stored at -80°C until SPE.

The peptide digests were SPE-cleaned using a plate manifold, in batches of 4 plates per day (384 digests). Cleaned digests were eluted into 96-well plates bearing the same unique plate number for tracking purpose and stored at -80°C until evaporation and reconstitution.

Samples were completely evaporated using a SpeedVac fitted with a rotor accommodating 4 frozen plates at a time (384 samples/day). Dried samples were reconstituted and homogenised using a multiplate vortex shaker holding four 96-well plates. Plates with reconstituted samples were stored at -80°C until LC-MS analyses.

The Vanquish UPLC accommodated four 96-well plates simultaneously. Each sample was injected once (unless the raw data was unusable; then a duplicate was obtained) and LC-MS analysed for 43 min so that 33 samples were run daily. Enough QC, IS and blanks were prepared to last the full LC-MS run. They were injected once every 48 wheat samples, amounting at the end of the total run to 86 replicates. Data files from the blank sample were inspected throughout the run to check for carry-over peaks (data not shown). Data files from the IS sample were used throughout the run to check for retention time drift, loss of peak resolution and signal intensity, as well as low mass accuracy (data not shown). They were also used during data file processing to isolate the LC-MS peaks belonging to the spiked IS among the wheat peaks. Data files from the QC sample were reviewed throughout the run to also assess loss of peak resolution and signal intensity, as well as batch reproducibility (data not shown). Instrument maintenance such as source and heated capillary cleaning, mass calibration and LC column swapping occurred to address retention time drift, as well as loss of peak resolution, signal intensity, and mass accuracy.

### 2. Background information on statistical analyses for big data

#### 2.1 Unsupervised multivariate clustering analyses (PCA, SOM, k-means, HCA)

Principal component analysis (PCA) dates from 1901 (Pearson 1901) which reduces the dimensionality of a dataset by linearly transforming the data into a new coordinate system where the variation in the data can be described with fewer dimensions than the initial data. The first two principal components are commonly used to chart the data as a scatterplot and to visually detect clusters of closely related data points. Biological studies such as transcriptomics and proteomics generating large datasets typically resort to PCA to identify trends in the multidimensional data while preserving the maximum amount of information (Jolliffe and Cadima 2016).

Invented in the 80's Kohonen (Kohonen 1982), a self-organising map (SOM) is an unsupervised machine learning technique that generates a low-dimensional representation of a higher dimensional dataset while preserving the topological structure of the data; a two-dimensional organization of the model effectively approximates similarities of high-dimensional data features (Kohonen 2013).

Invented in the 50's, K-means is another unsupervised machine learning method and its ease of implementation, simplicity, computational efficiency, and empirical success make it one of the most popular clustering methods (Jain 2010). In this clustering algorithm, n observations are segmented into k clusters in which each observation belongs to the cluster with the nearest mean, thereby minimising within-cluster variances. K is a key parameter that need to be prespecified. The algorithm then iteratively alternates between assigning each observation to its nearest centroid and shifting the location of the centroid to the mean of all data points until the classification remains unchanged (Bi et al. 2019, Cresta Morgado et al. 2022).

Divisive hierarchical clustering analysis (HCA) was introduced in 1999 to compare microarray data from different cell lines and tissues in order to discover colon tumour biomarkers (Alon et al. 1999). In divisive HCA, two vectors are randomly initialized and assigned each gene product using a probability function; the vectors are iteratively recalculated to form the centroids of two clusters. The same splitting process applies to each successive cluster, until each cluster consists of a single profile, and the binary tree is built from the history of the data splitting (Sherlock 2000). Two-dimensional agglomerative clustering visualised with coloured heat maps was also devised by Alon and colleagues in which not only the rows (gene products such transcripts, peptides) but also the columns (arrays, samples, stages, conditions, etc…) are rearranged (Alon et al. 1999).

#### 2.2. Bivariate analyses (correlation and linear regression)

Formulated in 1895 (Pearson 1895), correlation analysis aims at measuring an association or linear relationship between two continuous variables, by which the change in the magnitude of one variable is associated with a change in the magnitude of the other variable, either in the same (positive correlation) or in the opposite (negative correlation) direction (Schober, Boer, and Schwarte 2018). Correlation coefficients (R2 or r) are scaled from -1 to +1, where 0 denotes no association and the relationship gets stronger towards an absolute value of 1.

Simple linear regression dates from 1805 (Legendre 1805) and determines the relationship between a quantitative outcome and a single quantitative explanatory variable (Seltman 2018), such as LMA.

#### 2.3. Volcano plot and heat map

A volcano plot is a type of scatterplot that shows statistical significance (q-value) versus magnitude of change (effect size or fold change). It is often utilised for biomarker discovery in differential expression studies, in particular transcriptomics (Li 2012). It enables quick visual identification of gene products with large fold changes that are not only statistically significant but potentially biologically significant as well.

Heat maps offer an efficient method of visualizing complex quantitative data sets organized as matrices (Key 2012). A typical matrix is created by arranging the data such that each column contains the amount (intensity) from a single sample and each row corresponds to a single feature such as peptide. The intensities of the dependent variables in each cell are then converted into colours to create a heat map. Heat maps are flexible in that both the rows and columns can be arranged to explore a particular experimental hypothesis thus enabling a fast overview of data with a third quantitative dimension captured as coloured gradients (Pleil et al. 2011).

#### 3. MS/MS method parameters

Below are the parameters of the tandem methods tested.

---

Pass 1 (wheat-mixed-digests_MS2_pass01_1.raw):

Creator: Orbi_30393
Last modified: 10/12/2021 by Orbi_30393
MS Run Time (min): 43.00
Sequence override of method parameters not enabled.
Divert Valve: not used during run
Contact Closure: not used during run
Syringe Pump: not used during run
MS Detector Settings:
Real-time modifications to method not enabled
Stepped collision energy not enabled
Additional Microscans:

| | | |
|---|---|---|
| MS2 | 0 | 0 |
| MS3 | 0 | 0 |
| MS4 | 0 | 0 |
| MS5 | 0 | 0 |
| MS6 | 0 | 0 |
| MS7 | 0 | 0 |
| MS8 | 0 | 0 |
| MS9 | 0 | 0 |
| MS10 | 0 | 0 |

Experiment Type: Nth Order Double Play
Tune Method: Orbitrap-tune-file_2020-03-13_HESI
Scan Event Details:
1: FTMS + p norm o(300.0-2000.0)
  CV = 0.0V
2: ITMS + c norm Dep MS/MS Most intense ion from (1)
  Activation Type: CID
  Min. Signal Required: 3000.0
  Isolation Width: 2.00
  Normalized Coll. Energy: 35.0
  Default Charge State: 2
  Activation Q: 0.250
  Activation Time: 10.000
  CV = 0.0V
Scan Event 2 repeated for top 10 peaks.
Lock Masses:
  Pos List Name: N/A
    Source: API Source
  Mass List: (none)
  Neg List Name: N/A
    Source: API Source
  Mass List: (none)
Data Dependent Settings:
  Use separate polarity settings disabled
  Parent Mass List: (none)
  Reject Mass List: (none)

Neutral Loss Mass List: (none)
Product Mass List: (none)
Neutral loss in top: 3
Product in top: 3
Most intense if no parent masses found not enabled
Add/subtract mass not enabled
FT master scan preview mode enabled
Charge state screening enabled
Charge state dependent ETD time not enabled
Monoisotopic precursor selection enabled
Charge state rejection enabled
Unassigned charge states : rejected
Charge state 1 : rejected
Charge state 2 : not rejected
Charge state 3 : not rejected
Charge states 4+ : not rejected
Chromatography mode is disabled
Global Data Dependent Settings:
Predict ion injection time enabled
Use global parent and reject mass lists not enabled
Exclude parent mass from data dependent selection not enabled
Exclusion mass width by mass
Exclusion mass width low: 1.50000
Exclusion mass width high: 1.50000
Parent mass width by mass
Parent mass width low: 0.50000
Parent mass width high: 0.50000
Reject mass width by mass
Reject mass width low: 0.50000
Reject mass width high: 0.50000
Zoom/UltraZoom scan mass width by mass
Zoom/UltraZoom scan mass width low: 5.00
Zoom/UltraZoom scan mass width high: 5.00
FT SIM scan mass width low: 5.00
FT SIM scan mass width high: 5.00
Neutral Loss candidates processed by decreasing intensity
Neutral Loss mass width by mass
Neutral Loss mass width low: 0.50000
Neutral Loss mass width high: 0.50000
Product candidates processed by decreasing intensity
Product mass width by mass
Product mass width low: 0.50000
Product mass width high: 0.50000
MS mass range: 300.00-2000.00
MSn mass range by mass
MSn mass range: 0.00-1000000.00
Use m/z values as masses not enabled
Analog UV data dep. not enabled
Dynamic exclusion enabled
Repeat Count: 1
Repeat Duration: 30.00

Exclusion List Size: 500
Exclusion Duration: 180.00
Exclusion mass width by mass
Exclusion mass width low: 1.50000
Exclusion mass width high: 1.50000
Expiration: disabled
Isotopic data dependence not enabled
Custom Data Dependent Settings:
Not enabled

---

Pass 2 (wheat-mixed-digests_MS2_pass02_1.raw):

Creator: Orbi_30393
Last modified: 10/12/2021 by Orbi_30393
MS Run Time (min): 43.00
Sequence override of method parameters not enabled.
Divert Valve: not used during run
Contact Closure: not used during run
Syringe Pump: not used during run
MS Detector Settings:
Real-time modifications to method not enabled
Stepped collision energy not enabled
Additional Microscans:
MS2 0 0
MS3 0 0
MS4 0 0
MS5 0 0
MS6 0 0
MS7 0 0
MS8 0 0
MS9 0 0
MS10 0 0
Experiment Type: Nth Order Double Play
Tune Method: Orbitrap-tune-file_2020-03-13_HESI
Scan Event Details:
1: FTMS + p norm o(300.0-2000.0)
CV = 0.0V
2: ITMS + c norm Dep MS/MS Most intense ion from (1)
Activation Type: CID
Min. Signal Required: 3000.0
Isolation Width: 2.00
Normalized Coll. Energy: 35.0
Default Charge State: 2
Activation Q: 0.250
Activation Time: 10.000
CV = 0.0V
Scan Event 2 repeated for top 10 peaks.
Lock Masses:
Pos List Name: N/A
Source: API Source
Mass List: (none)

Neg List Name: N/A
Source: API Source
Mass List: (none)

Data Dependent Settings:

Use separate polarity settings disabled

Parent Mass List: (none)

Reject Mass List:

| MS Mass | Name |
|---|---|
| 300.14000 | exclusion0001 |
| 300.53000 | exclusion0002 |
| 301.15000 | exclusion0003 |
| 301.67000 | exclusion0004 |
| 302.58000 | exclusion0005 |
| 302.80000 | exclusion0006 |
| 303.19000 | exclusion0007 |
| 303.84000 | exclusion0008 |
| 304.11000 | exclusion0009 |
| 304.61000 | exclusion0010 |
| 305.47000 | exclusion0011 |
| 305.68000 | exclusion0012 |
| 305.85000 | exclusion0013 |
| 306.84000 | exclusion0014 |
| 307.71000 | exclusion0015 |
| 308.14000 | exclusion0016 |
| 309.18000 | exclusion0017 |
| 309.20000 | exclusion0018 |
| 309.48000 | exclusion0019 |
| 310.50000 | exclusion0020 |
| 311.67000 | exclusion0021 |
| 312.17000 | exclusion0022 |
| 312.50000 | exclusion0023 |
| 313.72000 | exclusion0024 |
| 314.70000 | exclusion0025 |
| 314.86000 | exclusion0026 |
| 316.21000 | exclusion0027 |
| 316.66000 | exclusion0028 |
| 317.16000 | exclusion0029 |
| 317.51000 | exclusion0030 |
| 318.19000 | exclusion0031 |
| 318.85000 | exclusion0032 |
| 319.17000 | exclusion0033 |
| 320.14000 | exclusion0034 |
| 320.18000 | exclusion0035 |
| 320.53000 | exclusion0036 |
| 320.72000 | exclusion0037 |
| 322.70000 | exclusion0038 |
| 323.15000 | exclusion0039 |
| 324.15000 | exclusion0040 |
| 324.71000 | exclusion0041 |
| 325.52000 | exclusion0042 |
| 326.18000 | exclusion0043 |

| | |
|---|---|
| 327.19000 | exclusion0044 |
| 328.16000 | exclusion0045 |
| 329.19000 | exclusion0046 |
| 329.72000 | exclusion0047 |
| 330.20000 | exclusion0048 |
| 330.21000 | exclusion0049 |
| 331.48000 | exclusion0050 |
| 332.65000 | exclusion0051 |
| 333.40000 | exclusion0052 |
| 333.52000 | exclusion0053 |
| 334.17000 | exclusion0054 |
| 334.65000 | exclusion0055 |
| 335.17000 | exclusion0056 |
| 336.19000 | exclusion0057 |
| 336.74000 | exclusion0058 |
| 337.13000 | exclusion0059 |
| 337.51000 | exclusion0060 |
| 337.84000 | exclusion0061 |
| 339.68000 | exclusion0062 |
| 339.81000 | exclusion0063 |
| 339.94000 | exclusion0064 |
| 340.18000 | exclusion0065 |
| 341.66000 | exclusion0066 |
| 341.87000 | exclusion0067 |
| 341.91000 | exclusion0068 |
| 342.22000 | exclusion0069 |
| 343.49000 | exclusion0070 |
| 343.73000 | exclusion0071 |
| 344.50000 | exclusion0072 |
| 345.70000 | exclusion0073 |
| 346.19000 | exclusion0074 |
| 346.21000 | exclusion0075 |
| 347.68000 | exclusion0076 |
| 347.85000 | exclusion0077 |
| 348.21000 | exclusion0078 |
| 348.21000 | exclusion0079 |
| 349.32000 | exclusion0080 |
| 349.73000 | exclusion0081 |
| 350.16000 | exclusion0082 |
| 350.21000 | exclusion0083 |
| 350.67000 | exclusion0084 |
| 351.68000 | exclusion0085 |
| 352.17000 | exclusion0086 |
| 352.50000 | exclusion0087 |
| 352.66000 | exclusion0088 |
| 353.80000 | exclusion0089 |
| 354.19000 | exclusion0090 |
| 355.19000 | exclusion0091 |
| 356.14000 | exclusion0092 |
| 357.75000 | exclusion0093 |
| 358.21000 | exclusion0094 |

| | |
|---|---|
| 358.55000 | exclusion0095 |
| 359.20000 | exclusion0096 |
| 359.31000 | exclusion0097 |
| 359.72000 | exclusion0098 |
| 359.86000 | exclusion0099 |
| 360.70000 | exclusion0100 |
| 361.69000 | exclusion0101 |
| 362.19000 | exclusion0102 |
| 363.18000 | exclusion0103 |
| 363.20000 | exclusion0104 |
| 363.31000 | exclusion0105 |
| 363.70000 | exclusion0106 |
| 364.12000 | exclusion0107 |
| 364.65000 | exclusion0108 |
| 365.17000 | exclusion0109 |
| 365.23000 | exclusion0110 |
| 365.75000 | exclusion0111 |
| 366.52000 | exclusion0112 |
| 367.21000 | exclusion0113 |
| 367.22000 | exclusion0114 |
| 367.50000 | exclusion0115 |
| 367.70000 | exclusion0116 |
| 369.72000 | exclusion0117 |
| 369.86000 | exclusion0118 |
| 370.77000 | exclusion0119 |
| 371.55000 | exclusion0120 |
| 371.68000 | exclusion0121 |
| 372.74000 | exclusion0122 |
| 373.20000 | exclusion0123 |
| 373.71000 | exclusion0124 |
| 374.20000 | exclusion0125 |
| 374.87000 | exclusion0126 |
| 375.21000 | exclusion0127 |
| 375.71000 | exclusion0128 |
| 376.19000 | exclusion0129 |
| 377.27000 | exclusion0130 |
| 377.49000 | exclusion0131 |
| 377.69000 | exclusion0132 |
| 379.20000 | exclusion0133 |
| 379.23000 | exclusion0134 |
| 379.52000 | exclusion0135 |
| 379.72000 | exclusion0136 |
| 380.18000 | exclusion0137 |
| 381.24000 | exclusion0138 |
| 381.73000 | exclusion0139 |
| 382.19000 | exclusion0140 |
| 382.67000 | exclusion0141 |
| 383.33000 | exclusion0142 |
| 384.17000 | exclusion0143 |
| 384.19000 | exclusion0144 |
| 385.17000 | exclusion0145 |

| | |
|---|---|
| 385.54000 | exclusion0146 |
| 385.71000 | exclusion0147 |
| 386.58000 | exclusion0148 |
| 386.69000 | exclusion0149 |
| 386.73000 | exclusion0150 |
| 387.21000 | exclusion0151 |
| 387.73000 | exclusion0152 |
| 388.22000 | exclusion0153 |
| 388.73000 | exclusion0154 |
| 388.93000 | exclusion0155 |
| 390.94000 | exclusion0156 |
| 390.95000 | exclusion0157 |
| 391.22000 | exclusion0158 |
| 391.71000 | exclusion0159 |
| 393.17000 | exclusion0160 |
| 393.57000 | exclusion0161 |
| 393.70000 | exclusion0162 |
| 394.18000 | exclusion0163 |
| 394.89000 | exclusion0164 |
| 395.20000 | exclusion0165 |
| 395.55000 | exclusion0166 |
| 396.70000 | exclusion0167 |
| 396.85000 | exclusion0168 |
| 397.21000 | exclusion0169 |
| 397.21000 | exclusion0170 |
| 397.58000 | exclusion0171 |
| 397.98000 | exclusion0172 |
| 398.53000 | exclusion0173 |
| 398.74000 | exclusion0174 |
| 399.22000 | exclusion0175 |
| 399.23000 | exclusion0176 |
| 399.53000 | exclusion0177 |
| 400.52000 | exclusion0178 |
| 400.75000 | exclusion0179 |
| 400.89000 | exclusion0180 |
| 401.22000 | exclusion0181 |
| 401.34000 | exclusion0182 |
| 401.72000 | exclusion0183 |
| 403.23000 | exclusion0184 |
| 403.24000 | exclusion0185 |
| 404.20000 | exclusion0186 |
| 404.54000 | exclusion0187 |
| 405.21000 | exclusion0188 |
| 405.21000 | exclusion0189 |
| 405.52000 | exclusion0190 |
| 406.22000 | exclusion0191 |
| 406.75000 | exclusion0192 |
| 407.19000 | exclusion0193 |
| 407.25000 | exclusion0194 |
| 408.69000 | exclusion0195 |
| 409.73000 | exclusion0196 |

| | |
|---|---|
| 409.86000 | exclusion0197 |
| 410.56000 | exclusion0198 |
| 410.73000 | exclusion0199 |
| 411.21000 | exclusion0200 |
| 413.73000 | exclusion0201 |
| 413.93000 | exclusion0202 |
| 414.28000 | exclusion0203 |
| 414.76000 | exclusion0204 |
| 415.04000 | exclusion0205 |
| 415.18000 | exclusion0206 |
| 415.57000 | exclusion0207 |
| 416.19000 | exclusion0208 |
| 417.22000 | exclusion0209 |
| 417.24000 | exclusion0210 |
| 417.67000 | exclusion0211 |
| 417.75000 | exclusion0212 |
| 417.86000 | exclusion0213 |
| 418.60000 | exclusion0214 |
| 419.20000 | exclusion0215 |
| 419.21000 | exclusion0216 |
| 419.72000 | exclusion0217 |
| 420.96000 | exclusion0218 |
| 421.18000 | exclusion0219 |
| 421.67000 | exclusion0220 |
| 422.19000 | exclusion0221 |
| 422.73000 | exclusion0222 |
| 423.73000 | exclusion0223 |
| 423.87000 | exclusion0224 |
| 424.24000 | exclusion0225 |
| 424.90000 | exclusion0226 |
| 425.26000 | exclusion0227 |
| 425.72000 | exclusion0228 |
| 426.55000 | exclusion0229 |
| 426.73000 | exclusion0230 |
| 427.66000 | exclusion0231 |
| 428.54000 | exclusion0232 |
| 428.77000 | exclusion0233 |
| 429.37000 | exclusion0234 |
| 430.21000 | exclusion0235 |
| 430.75000 | exclusion0236 |
| 431.90000 | exclusion0237 |
| 432.22000 | exclusion0238 |
| 432.56000 | exclusion0239 |
| 432.69000 | exclusion0240 |
| 433.25000 | exclusion0241 |
| 433.72000 | exclusion0242 |
| 433.86000 | exclusion0243 |
| 433.98000 | exclusion0244 |
| 434.22000 | exclusion0245 |
| 435.23000 | exclusion0246 |
| 436.29000 | exclusion0247 |

| | |
|---|---|
| 437.53000 | exclusion0248 |
| 438.76000 | exclusion0249 |
| 439.20000 | exclusion0250 |
| 439.24000 | exclusion0251 |
| 439.70000 | exclusion0252 |
| 440.89000 | exclusion0253 |
| 441.18000 | exclusion0254 |
| 441.20000 | exclusion0255 |
| 442.27000 | exclusion0256 |
| 442.69000 | exclusion0257 |
| 443.22000 | exclusion0258 |
| 443.89000 | exclusion0259 |
| 444.22000 | exclusion0260 |
| 444.25000 | exclusion0261 |
| 445.72000 | exclusion0262 |
| 446.21000 | exclusion0263 |
| 446.71000 | exclusion0264 |
| 447.26000 | exclusion0265 |
| 447.77000 | exclusion0266 |
| 447.89000 | exclusion0267 |
| 448.21000 | exclusion0268 |
| 448.88000 | exclusion0269 |
| 449.29000 | exclusion0270 |
| 450.18000 | exclusion0271 |
| 450.20000 | exclusion0272 |
| 450.22000 | exclusion0273 |
| 450.55000 | exclusion0274 |
| 450.73000 | exclusion0275 |
| 451.26000 | exclusion0276 |
| 451.72000 | exclusion0277 |
| 451.90000 | exclusion0278 |
| 452.91000 | exclusion0279 |
| 452.97000 | exclusion0280 |
| 453.55000 | exclusion0281 |
| 453.97000 | exclusion0282 |
| 454.24000 | exclusion0283 |
| 454.55000 | exclusion0284 |
| 455.48000 | exclusion0285 |
| 455.72000 | exclusion0286 |
| 455.93000 | exclusion0287 |
| 456.21000 | exclusion0288 |
| 456.74000 | exclusion0289 |
| 457.20000 | exclusion0290 |
| 457.23000 | exclusion0291 |
| 457.49000 | exclusion0292 |
| 458.53000 | exclusion0293 |
| 458.91000 | exclusion0294 |
| 459.21000 | exclusion0295 |
| 459.22000 | exclusion0296 |
| 459.76000 | exclusion0297 |
| 460.24000 | exclusion0298 |

| | |
|---|---|
| 460.27000 | exclusion0299 |
| 460.57000 | exclusion0300 |
| 460.73000 | exclusion0301 |
| 461.26000 | exclusion0302 |
| 461.72000 | exclusion0303 |
| 462.24000 | exclusion0304 |
| 462.62000 | exclusion0305 |
| 462.71000 | exclusion0306 |
| 463.23000 | exclusion0307 |
| 463.91000 | exclusion0308 |
| 464.24000 | exclusion0309 |
| 464.76000 | exclusion0310 |
| 465.19000 | exclusion0311 |
| 465.24000 | exclusion0312 |
| 465.96000 | exclusion0313 |
| 466.11000 | exclusion0314 |
| 466.27000 | exclusion0315 |
| 466.73000 | exclusion0316 |
| 466.73000 | exclusion0317 |
| 467.74000 | exclusion0318 |
| 467.88000 | exclusion0319 |
| 468.11000 | exclusion0320 |
| 468.22000 | exclusion0321 |
| 468.61000 | exclusion0322 |
| 468.92000 | exclusion0323 |
| 469.23000 | exclusion0324 |
| 469.76000 | exclusion0325 |
| 469.76000 | exclusion0326 |
| 470.26000 | exclusion0327 |
| 470.27000 | exclusion0328 |
| 471.54000 | exclusion0329 |
| 471.78000 | exclusion0330 |
| 472.26000 | exclusion0331 |
| 473.07000 | exclusion0332 |
| 473.58000 | exclusion0333 |
| 473.77000 | exclusion0334 |
| 474.25000 | exclusion0335 |
| 475.57000 | exclusion0336 |
| 476.25000 | exclusion0337 |
| 476.70000 | exclusion0338 |
| 477.22000 | exclusion0339 |
| 477.56000 | exclusion0340 |
| 477.91000 | exclusion0341 |
| 477.99000 | exclusion0342 |
| 478.28000 | exclusion0343 |
| 479.59000 | exclusion0344 |
| 479.76000 | exclusion0345 |
| 479.76000 | exclusion0346 |
| 479.90000 | exclusion0347 |
| 480.22000 | exclusion0348 |
| 480.80000 | exclusion0349 |

| | |
|---|---|
| 481.24000 | exclusion0350 |
| 481.55000 | exclusion0351 |
| 481.78000 | exclusion0352 |
| 481.90000 | exclusion0353 |
| 482.30000 | exclusion0354 |
| 482.89000 | exclusion0355 |
| 483.24000 | exclusion0356 |
| 483.56000 | exclusion0357 |
| 484.25000 | exclusion0358 |
| 484.74000 | exclusion0359 |
| 485.24000 | exclusion0360 |
| 485.73000 | exclusion0361 |
| 486.25000 | exclusion0362 |
| 486.31000 | exclusion0363 |
| 486.74000 | exclusion0364 |
| 486.92000 | exclusion0365 |
| 487.24000 | exclusion0366 |
| 487.75000 | exclusion0367 |
| 488.22000 | exclusion0368 |
| 488.79000 | exclusion0369 |
| 489.25000 | exclusion0370 |
| 489.61000 | exclusion0371 |
| 489.77000 | exclusion0372 |
| 489.91000 | exclusion0373 |
| 490.26000 | exclusion0374 |
| 490.72000 | exclusion0375 |
| 490.94000 | exclusion0376 |
| 491.74000 | exclusion0377 |
| 491.91000 | exclusion0378 |
| 492.25000 | exclusion0379 |
| 492.26000 | exclusion0380 |
| 492.59000 | exclusion0381 |
| 493.23000 | exclusion0382 |
| 493.76000 | exclusion0383 |
| 494.28000 | exclusion0384 |
| 494.62000 | exclusion0385 |
| 495.24000 | exclusion0386 |
| 495.27000 | exclusion0387 |
| 495.72000 | exclusion0388 |
| 496.19000 | exclusion0389 |
| 496.23000 | exclusion0390 |
| 497.26000 | exclusion0391 |
| 497.77000 | exclusion0392 |
| 498.23000 | exclusion0393 |
| 498.27000 | exclusion0394 |
| 498.61000 | exclusion0395 |
| 498.74000 | exclusion0396 |
| 499.58000 | exclusion0397 |
| 499.76000 | exclusion0398 |
| 499.96000 | exclusion0399 |
| 500.24000 | exclusion0400 |

| | |
|---|---|
| 500.29000 | exclusion0401 |
| 500.74000 | exclusion0402 |
| 500.92000 | exclusion0403 |
| 501.27000 | exclusion0404 |
| 501.28000 | exclusion0405 |
| 501.60000 | exclusion0406 |
| 501.91000 | exclusion0407 |
| 502.78000 | exclusion0408 |
| 502.93000 | exclusion0409 |
| 503.11000 | exclusion0410 |
| 503.20000 | exclusion0411 |
| 503.30000 | exclusion0412 |
| 503.75000 | exclusion0413 |
| 504.11000 | exclusion0414 |
| 504.22000 | exclusion0415 |
| 504.73000 | exclusion0416 |
| 504.88000 | exclusion0417 |
| 505.27000 | exclusion0418 |
| 505.92000 | exclusion0419 |
| 506.26000 | exclusion0420 |
| 506.80000 | exclusion0421 |
| 506.94000 | exclusion0422 |
| 507.26000 | exclusion0423 |
| 507.27000 | exclusion0424 |
| 508.62000 | exclusion0425 |
| 508.70000 | exclusion0426 |
| 509.00000 | exclusion0427 |
| 509.26000 | exclusion0428 |
| 510.29000 | exclusion0429 |
| 510.34000 | exclusion0430 |
| 510.59000 | exclusion0431 |
| 510.95000 | exclusion0432 |
| 511.20000 | exclusion0433 |
| 511.76000 | exclusion0434 |
| 511.92000 | exclusion0435 |
| 512.27000 | exclusion0436 |
| 512.75000 | exclusion0437 |
| 512.94000 | exclusion0438 |
| 513.27000 | exclusion0439 |
| 513.59000 | exclusion0440 |
| 513.76000 | exclusion0441 |
| 514.27000 | exclusion0442 |
| 514.77000 | exclusion0443 |
| 514.89000 | exclusion0444 |
| 515.29000 | exclusion0445 |
| 515.73000 | exclusion0446 |
| 515.82000 | exclusion0447 |
| 516.80000 | exclusion0448 |
| 516.96000 | exclusion0449 |
| 517.24000 | exclusion0450 |
| 517.26000 | exclusion0451 |

| | |
|---|---|
| 517.62000 | exclusion0452 |
| 517.77000 | exclusion0453 |
| 518.29000 | exclusion0454 |
| 519.14000 | exclusion0455 |
| 519.56000 | exclusion0456 |
| 519.78000 | exclusion0457 |
| 519.79000 | exclusion0458 |
| 519.92000 | exclusion0459 |
| 520.26000 | exclusion0460 |
| 520.62000 | exclusion0461 |
| 520.78000 | exclusion0462 |
| 521.53000 | exclusion0463 |
| 522.22000 | exclusion0464 |
| 522.28000 | exclusion0465 |
| 522.77000 | exclusion0466 |
| 523.27000 | exclusion0467 |
| 523.82000 | exclusion0468 |
| 523.99000 | exclusion0469 |
| 524.22000 | exclusion0470 |
| 524.59000 | exclusion0471 |
| 524.77000 | exclusion0472 |
| 524.93000 | exclusion0473 |
| 525.26000 | exclusion0474 |
| 525.62000 | exclusion0475 |
| 525.96000 | exclusion0476 |
| 526.52000 | exclusion0477 |
| 526.60000 | exclusion0478 |
| 526.92000 | exclusion0479 |
| 527.27000 | exclusion0480 |
| 527.81000 | exclusion0481 |
| 528.23000 | exclusion0482 |
| 528.73000 | exclusion0483 |
| 529.26000 | exclusion0484 |
| 530.25000 | exclusion0485 |
| 530.74000 | exclusion0486 |
| 531.29000 | exclusion0487 |
| 531.59000 | exclusion0488 |
| 532.25000 | exclusion0489 |
| 532.25000 | exclusion0490 |
| 533.28000 | exclusion0491 |
| 533.35000 | exclusion0492 |
| 533.53000 | exclusion0493 |
| 533.53000 | exclusion0494 |
| 533.82000 | exclusion0495 |
| 534.23000 | exclusion0496 |
| 534.27000 | exclusion0497 |
| 534.94000 | exclusion0498 |
| 535.82000 | exclusion0499 |
| 535.92000 | exclusion0500 |
| 536.09000 | exclusion0501 |
| 536.26000 | exclusion0502 |

| | |
|---|---|
| 536.28000 | exclusion0503 |
| 537.24000 | exclusion0504 |
| 537.79000 | exclusion0505 |
| 537.97000 | exclusion0506 |
| 538.29000 | exclusion0507 |
| 538.59000 | exclusion0508 |
| 538.70000 | exclusion0509 |
| 538.77000 | exclusion0510 |
| 538.80000 | exclusion0511 |
| 540.27000 | exclusion0512 |
| 540.79000 | exclusion0513 |
| 541.23000 | exclusion0514 |
| 541.26000 | exclusion0515 |
| 541.62000 | exclusion0516 |
| 541.92000 | exclusion0517 |
| 542.31000 | exclusion0518 |
| 542.58000 | exclusion0519 |
| 543.27000 | exclusion0520 |
| 543.28000 | exclusion0521 |
| 543.76000 | exclusion0522 |
| 544.01000 | exclusion0523 |
| 544.22000 | exclusion0524 |
| 545.28000 | exclusion0525 |
| 545.67000 | exclusion0526 |
| 545.77000 | exclusion0527 |
| 546.24000 | exclusion0528 |
| 546.96000 | exclusion0529 |
| 547.28000 | exclusion0530 |
| 547.77000 | exclusion0531 |
| 548.77000 | exclusion0532 |
| 548.79000 | exclusion0533 |
| 549.04000 | exclusion0534 |
| 549.28000 | exclusion0535 |
| 549.49000 | exclusion0536 |
| 550.31000 | exclusion0537 |
| 550.33000 | exclusion0538 |
| 550.78000 | exclusion0539 |
| 551.28000 | exclusion0540 |
| 551.29000 | exclusion0541 |
| 551.60000 | exclusion0542 |
| 551.89000 | exclusion0543 |
| 552.04000 | exclusion0544 |
| 552.29000 | exclusion0545 |
| 552.30000 | exclusion0546 |
| 552.74000 | exclusion0547 |
| 553.28000 | exclusion0548 |
| 553.77000 | exclusion0549 |
| 554.37000 | exclusion0550 |
| 554.77000 | exclusion0551 |
| 555.28000 | exclusion0552 |
| 555.72000 | exclusion0553 |

| | |
|---|---|
| 556.30000 | exclusion0554 |
| 556.94000 | exclusion0555 |
| 557.26000 | exclusion0556 |
| 557.30000 | exclusion0557 |
| 557.83000 | exclusion0558 |
| 558.07000 | exclusion0559 |
| 558.28000 | exclusion0560 |
| 558.77000 | exclusion0561 |
| 559.27000 | exclusion0562 |
| 559.80000 | exclusion0563 |
| 560.28000 | exclusion0564 |
| 560.49000 | exclusion0565 |
| 560.66000 | exclusion0566 |
| 560.77000 | exclusion0567 |
| 561.30000 | exclusion0568 |
| 562.27000 | exclusion0569 |
| 562.31000 | exclusion0570 |
| 562.81000 | exclusion0571 |
| 563.08000 | exclusion0572 |
| 563.29000 | exclusion0573 |
| 563.95000 | exclusion0574 |
| 564.78000 | exclusion0575 |
| 565.31000 | exclusion0576 |
| 565.34000 | exclusion0577 |
| 566.27000 | exclusion0578 |
| 566.77000 | exclusion0579 |
| 567.03000 | exclusion0580 |
| 567.29000 | exclusion0581 |
| 567.59000 | exclusion0582 |
| 567.83000 | exclusion0583 |
| 567.93000 | exclusion0584 |
| 568.28000 | exclusion0585 |
| 569.24000 | exclusion0586 |
| 569.30000 | exclusion0587 |
| 569.81000 | exclusion0588 |
| 570.26000 | exclusion0589 |
| 570.32000 | exclusion0590 |
| 571.33000 | exclusion0591 |
| 571.62000 | exclusion0592 |
| 571.76000 | exclusion0593 |
| 572.26000 | exclusion0594 |
| 572.50000 | exclusion0595 |
| 572.67000 | exclusion0596 |
| 573.25000 | exclusion0597 |
| 573.30000 | exclusion0598 |
| 573.81000 | exclusion0599 |
| 573.97000 | exclusion0600 |
| 574.28000 | exclusion0601 |
| 574.79000 | exclusion0602 |
| 574.80000 | exclusion0603 |
| 575.18000 | exclusion0604 |

| | |
|---|---|
| 575.30000 | exclusion0605 |
| 575.66000 | exclusion0606 |
| 575.77000 | exclusion0607 |
| 576.41000 | exclusion0608 |
| 576.66000 | exclusion0609 |
| 576.71000 | exclusion0610 |
| 577.30000 | exclusion0611 |
| 577.81000 | exclusion0612 |
| 578.29000 | exclusion0613 |
| 578.31000 | exclusion0614 |
| 578.77000 | exclusion0615 |
| 578.96000 | exclusion0616 |
| 579.11000 | exclusion0617 |
| 579.29000 | exclusion0618 |
| 579.59000 | exclusion0619 |
| 579.75000 | exclusion0620 |
| 579.84000 | exclusion0621 |
| 580.34000 | exclusion0622 |
| 580.79000 | exclusion0623 |
| 581.25000 | exclusion0624 |
| 581.36000 | exclusion0625 |
| 581.79000 | exclusion0626 |
| 582.31000 | exclusion0627 |
| 582.74000 | exclusion0628 |
| 583.10000 | exclusion0629 |
| 583.79000 | exclusion0630 |
| 583.95000 | exclusion0631 |
| 584.38000 | exclusion0632 |
| 584.55000 | exclusion0633 |
| 584.68000 | exclusion0634 |
| 584.76000 | exclusion0635 |
| 585.83000 | exclusion0636 |
| 586.06000 | exclusion0637 |
| 586.28000 | exclusion0638 |
| 586.30000 | exclusion0639 |
| 586.81000 | exclusion0640 |
| 587.81000 | exclusion0641 |
| 588.29000 | exclusion0642 |
| 588.30000 | exclusion0643 |
| 588.80000 | exclusion0644 |
| 589.31000 | exclusion0645 |
| 590.25000 | exclusion0646 |
| 590.51000 | exclusion0647 |
| 590.76000 | exclusion0648 |
| 591.09000 | exclusion0649 |
| 591.29000 | exclusion0650 |
| 591.79000 | exclusion0651 |
| 592.64000 | exclusion0652 |
| 592.76000 | exclusion0653 |
| 593.16000 | exclusion0654 |
| 593.32000 | exclusion0655 |

| | |
|---|---|
| 593.36000 | exclusion0656 |
| 593.81000 | exclusion0657 |
| 594.76000 | exclusion0658 |
| 594.92000 | exclusion0659 |
| 595.33000 | exclusion0660 |
| 595.92000 | exclusion0661 |
| 596.32000 | exclusion0662 |
| 596.77000 | exclusion0663 |
| 597.28000 | exclusion0664 |
| 597.31000 | exclusion0665 |
| 597.56000 | exclusion0666 |
| 597.85000 | exclusion0667 |
| 598.28000 | exclusion0668 |
| 598.40000 | exclusion0669 |
| 598.97000 | exclusion0670 |
| 599.66000 | exclusion0671 |
| 599.78000 | exclusion0672 |
| 600.26000 | exclusion0673 |
| 601.32000 | exclusion0674 |
| 601.62000 | exclusion0675 |
| 601.77000 | exclusion0676 |
| 601.84000 | exclusion0677 |
| 603.29000 | exclusion0678 |
| 603.29000 | exclusion0679 |
| 603.65000 | exclusion0680 |
| 603.79000 | exclusion0681 |
| 604.28000 | exclusion0682 |
| 605.29000 | exclusion0683 |
| 605.78000 | exclusion0684 |
| 606.28000 | exclusion0685 |
| 606.29000 | exclusion0686 |
| 606.31000 | exclusion0687 |
| 606.62000 | exclusion0688 |
| 607.81000 | exclusion0689 |
| 608.29000 | exclusion0690 |
| 608.31000 | exclusion0691 |
| 609.31000 | exclusion0692 |
| 609.33000 | exclusion0693 |
| 610.36000 | exclusion0694 |
| 610.78000 | exclusion0695 |
| 611.00000 | exclusion0696 |
| 611.27000 | exclusion0697 |
| 611.30000 | exclusion0698 |
| 611.31000 | exclusion0699 |
| 611.81000 | exclusion0700 |
| 612.95000 | exclusion0701 |
| 613.31000 | exclusion0702 |
| 613.32000 | exclusion0703 |
| 613.79000 | exclusion0704 |
| 614.09000 | exclusion0705 |
| 614.48000 | exclusion0706 |

| | |
|---|---|
| 614.82000 | exclusion0707 |
| 614.96000 | exclusion0708 |
| 615.29000 | exclusion0709 |
| 615.83000 | exclusion0710 |
| 616.30000 | exclusion0711 |
| 616.66000 | exclusion0712 |
| 616.79000 | exclusion0713 |
| 616.99000 | exclusion0714 |
| 617.34000 | exclusion0715 |
| 617.80000 | exclusion0716 |
| 617.80000 | exclusion0717 |
| 618.29000 | exclusion0718 |
| 618.93000 | exclusion0719 |
| 619.13000 | exclusion0720 |
| 619.30000 | exclusion0721 |
| 619.63000 | exclusion0722 |
| 619.85000 | exclusion0723 |
| 620.02000 | exclusion0724 |
| 620.30000 | exclusion0725 |
| 620.83000 | exclusion0726 |
| 621.33000 | exclusion0727 |
| 621.79000 | exclusion0728 |
| 621.98000 | exclusion0729 |
| 622.34000 | exclusion0730 |
| 622.66000 | exclusion0731 |
| 622.79000 | exclusion0732 |
| 623.24000 | exclusion0733 |
| 623.33000 | exclusion0734 |
| 623.78000 | exclusion0735 |
| 624.34000 | exclusion0736 |
| 624.77000 | exclusion0737 |
| 625.04000 | exclusion0738 |
| 625.33000 | exclusion0739 |
| 625.33000 | exclusion0740 |
| 625.93000 | exclusion0741 |
| 626.28000 | exclusion0742 |
| 626.81000 | exclusion0743 |
| 627.01000 | exclusion0744 |
| 627.30000 | exclusion0745 |
| 628.29000 | exclusion0746 |
| 628.58000 | exclusion0747 |
| 628.80000 | exclusion0748 |
| 628.96000 | exclusion0749 |
| 629.03000 | exclusion0750 |
| 629.82000 | exclusion0751 |
| 630.30000 | exclusion0752 |
| 630.77000 | exclusion0753 |
| 631.31000 | exclusion0754 |
| 631.83000 | exclusion0755 |
| 632.29000 | exclusion0756 |
| 632.83000 | exclusion0757 |

| | |
|---|---|
| 633.28000 | exclusion0758 |
| 633.31000 | exclusion0759 |
| 633.83000 | exclusion0760 |
| 634.33000 | exclusion0761 |
| 634.79000 | exclusion0762 |
| 635.86000 | exclusion0763 |
| 636.32000 | exclusion0764 |
| 636.83000 | exclusion0765 |
| 637.64000 | exclusion0766 |
| 637.78000 | exclusion0767 |
| 637.85000 | exclusion0768 |
| 638.23000 | exclusion0769 |
| 638.26000 | exclusion0770 |
| 638.85000 | exclusion0771 |
| 639.35000 | exclusion0772 |
| 639.59000 | exclusion0773 |
| 639.61000 | exclusion0774 |
| 639.78000 | exclusion0775 |
| 639.92000 | exclusion0776 |
| 640.30000 | exclusion0777 |
| 640.61000 | exclusion0778 |
| 641.35000 | exclusion0779 |
| 641.44000 | exclusion0780 |
| 641.84000 | exclusion0781 |
| 642.86000 | exclusion0782 |
| 643.29000 | exclusion0783 |
| 643.36000 | exclusion0784 |
| 643.82000 | exclusion0785 |
| 644.02000 | exclusion0786 |
| 644.87000 | exclusion0787 |
| 645.02000 | exclusion0788 |
| 645.35000 | exclusion0789 |
| 645.51000 | exclusion0790 |
| 645.70000 | exclusion0791 |
| 645.98000 | exclusion0792 |
| 647.31000 | exclusion0793 |
| 647.35000 | exclusion0794 |
| 647.87000 | exclusion0795 |
| 648.00000 | exclusion0796 |
| 648.29000 | exclusion0797 |
| 648.30000 | exclusion0798 |
| 648.31000 | exclusion0799 |
| 648.82000 | exclusion0800 |
| 649.28000 | exclusion0801 |
| 649.82000 | exclusion0802 |
| 650.29000 | exclusion0803 |
| 650.82000 | exclusion0804 |
| 651.30000 | exclusion0805 |
| 651.32000 | exclusion0806 |
| 651.78000 | exclusion0807 |
| 652.84000 | exclusion0808 |

| | |
|---|---|
| 653.24000 | exclusion0809 |
| 653.36000 | exclusion0810 |
| 653.63000 | exclusion0811 |
| 654.30000 | exclusion0812 |
| 654.35000 | exclusion0813 |
| 654.85000 | exclusion0814 |
| 655.09000 | exclusion0815 |
| 655.82000 | exclusion0816 |
| 656.32000 | exclusion0817 |
| 656.54000 | exclusion0818 |
| 657.03000 | exclusion0819 |
| 657.35000 | exclusion0820 |
| 657.38000 | exclusion0821 |
| 657.83000 | exclusion0822 |
| 658.03000 | exclusion0823 |
| 658.32000 | exclusion0824 |
| 658.35000 | exclusion0825 |
| 659.28000 | exclusion0826 |
| 659.34000 | exclusion0827 |
| 659.78000 | exclusion0828 |
| 660.30000 | exclusion0829 |
| 660.35000 | exclusion0830 |
| 660.66000 | exclusion0831 |
| 661.30000 | exclusion0832 |
| 661.81000 | exclusion0833 |
| 662.32000 | exclusion0834 |
| 662.83000 | exclusion0835 |
| 663.32000 | exclusion0836 |
| 663.75000 | exclusion0837 |
| 664.11000 | exclusion0838 |
| 664.37000 | exclusion0839 |
| 664.66000 | exclusion0840 |
| 665.32000 | exclusion0841 |
| 665.34000 | exclusion0842 |
| 665.64000 | exclusion0843 |
| 665.87000 | exclusion0844 |
| 666.34000 | exclusion0845 |
| 666.86000 | exclusion0846 |
| 667.03000 | exclusion0847 |
| 667.36000 | exclusion0848 |
| 667.52000 | exclusion0849 |
| 668.66000 | exclusion0850 |
| 668.83000 | exclusion0851 |
| 669.34000 | exclusion0852 |
| 669.51000 | exclusion0853 |
| 669.85000 | exclusion0854 |
| 669.90000 | exclusion0855 |
| 670.29000 | exclusion0856 |
| 670.80000 | exclusion0857 |
| 671.83000 | exclusion0858 |
| 672.32000 | exclusion0859 |

| | |
|---|---|
| 672.82000 | exclusion0860 |
| 673.11000 | exclusion0861 |
| 673.33000 | exclusion0862 |
| 673.34000 | exclusion0863 |
| 674.31000 | exclusion0864 |
| 674.33000 | exclusion0865 |
| 674.58000 | exclusion0866 |
| 674.83000 | exclusion0867 |
| 675.19000 | exclusion0868 |
| 676.05000 | exclusion0869 |
| 676.85000 | exclusion0870 |
| 677.00000 | exclusion0871 |
| 677.28000 | exclusion0872 |
| 677.34000 | exclusion0873 |
| 677.85000 | exclusion0874 |
| 677.88000 | exclusion0875 |
| 678.36000 | exclusion0876 |
| 679.04000 | exclusion0877 |
| 679.31000 | exclusion0878 |
| 679.85000 | exclusion0879 |
| 680.33000 | exclusion0880 |
| 680.78000 | exclusion0881 |
| 681.32000 | exclusion0882 |
| 682.37000 | exclusion0883 |
| 682.67000 | exclusion0884 |
| 682.82000 | exclusion0885 |
| 683.32000 | exclusion0886 |
| 683.37000 | exclusion0887 |
| 683.97000 | exclusion0888 |
| 684.29000 | exclusion0889 |
| 684.78000 | exclusion0890 |
| 685.67000 | exclusion0891 |
| 685.83000 | exclusion0892 |
| 686.31000 | exclusion0893 |
| 686.34000 | exclusion0894 |
| 686.85000 | exclusion0895 |
| 687.30000 | exclusion0896 |
| 687.33000 | exclusion0897 |
| 687.68000 | exclusion0898 |
| 688.78000 | exclusion0899 |
| 689.00000 | exclusion0900 |
| 689.34000 | exclusion0901 |
| 689.37000 | exclusion0902 |
| 689.67000 | exclusion0903 |
| 690.35000 | exclusion0904 |
| 690.72000 | exclusion0905 |
| 691.32000 | exclusion0906 |
| 691.36000 | exclusion0907 |
| 692.02000 | exclusion0908 |
| 692.26000 | exclusion0909 |
| 692.70000 | exclusion0910 |

| | |
|---|---|
| 693.10000 | exclusion0911 |
| 693.56000 | exclusion0912 |
| 693.88000 | exclusion0913 |
| 694.31000 | exclusion0914 |
| 695.33000 | exclusion0915 |
| 695.39000 | exclusion0916 |
| 695.85000 | exclusion0917 |
| 696.33000 | exclusion0918 |
| 696.35000 | exclusion0919 |
| 697.01000 | exclusion0920 |
| 697.36000 | exclusion0921 |
| 697.83000 | exclusion0922 |
| 698.37000 | exclusion0923 |
| 698.89000 | exclusion0924 |
| 699.12000 | exclusion0925 |
| 699.37000 | exclusion0926 |
| 699.38000 | exclusion0927 |
| 699.68000 | exclusion0928 |
| 700.37000 | exclusion0929 |
| 700.90000 | exclusion0930 |
| 701.33000 | exclusion0931 |
| 702.31000 | exclusion0932 |
| 702.33000 | exclusion0933 |
| 702.77000 | exclusion0934 |
| 702.87000 | exclusion0935 |
| 703.38000 | exclusion0936 |
| 703.71000 | exclusion0937 |
| 704.31000 | exclusion0938 |
| 705.27000 | exclusion0939 |
| 705.37000 | exclusion0940 |
| 705.82000 | exclusion0941 |
| 706.81000 | exclusion0942 |
| 707.02000 | exclusion0943 |
| 707.32000 | exclusion0944 |
| 708.04000 | exclusion0945 |
| 708.33000 | exclusion0946 |
| 708.38000 | exclusion0947 |
| 708.85000 | exclusion0948 |
| 709.35000 | exclusion0949 |
| 709.35000 | exclusion0950 |
| 709.90000 | exclusion0951 |
| 710.33000 | exclusion0952 |
| 710.36000 | exclusion0953 |
| 710.81000 | exclusion0954 |
| 712.05000 | exclusion0955 |
| 712.34000 | exclusion0956 |
| 712.35000 | exclusion0957 |
| 712.59000 | exclusion0958 |
| 712.70000 | exclusion0959 |
| 712.80000 | exclusion0960 |
| 713.09000 | exclusion0961 |

| | |
|---|---|
| 713.34000 | exclusion0962 |
| 713.68000 | exclusion0963 |
| 713.89000 | exclusion0964 |
| 714.35000 | exclusion0965 |
| 714.40000 | exclusion0966 |
| 715.01000 | exclusion0967 |
| 715.36000 | exclusion0968 |
| 715.84000 | exclusion0969 |
| 716.31000 | exclusion0970 |
| 716.87000 | exclusion0971 |
| 717.83000 | exclusion0972 |
| 718.31000 | exclusion0973 |
| 718.34000 | exclusion0974 |
| 719.35000 | exclusion0975 |
| 719.71000 | exclusion0976 |
| 719.81000 | exclusion0977 |
| 720.32000 | exclusion0978 |
| 720.90000 | exclusion0979 |
| 721.37000 | exclusion0980 |
| 721.41000 | exclusion0981 |
| 721.73000 | exclusion0982 |
| 721.79000 | exclusion0983 |
| 722.59000 | exclusion0984 |
| 722.87000 | exclusion0985 |
| 723.07000 | exclusion0986 |
| 723.33000 | exclusion0987 |
| 723.40000 | exclusion0988 |
| 724.88000 | exclusion0989 |
| 725.04000 | exclusion0990 |
| 725.38000 | exclusion0991 |
| 725.82000 | exclusion0992 |
| 726.05000 | exclusion0993 |
| 726.72000 | exclusion0994 |
| 726.81000 | exclusion0995 |
| 727.37000 | exclusion0996 |
| 727.50000 | exclusion0997 |
| 728.02000 | exclusion0998 |
| 728.32000 | exclusion0999 |
| 728.88000 | exclusion1000 |
| 729.32000 | exclusion1001 |
| 729.86000 | exclusion1002 |
| 730.36000 | exclusion1003 |
| 730.36000 | exclusion1004 |
| 730.64000 | exclusion1005 |
| 730.87000 | exclusion1006 |
| 731.01000 | exclusion1007 |
| 731.88000 | exclusion1008 |
| 732.35000 | exclusion1009 |
| 732.84000 | exclusion1010 |
| 733.37000 | exclusion1011 |
| 733.86000 | exclusion1012 |

| | |
|---|---|
| 734.32000 | exclusion1013 |
| 734.36000 | exclusion1014 |
| 735.33000 | exclusion1015 |
| 735.90000 | exclusion1016 |
| 736.37000 | exclusion1017 |
| 736.38000 | exclusion1018 |
| 736.86000 | exclusion1019 |
| 737.36000 | exclusion1020 |
| 737.88000 | exclusion1021 |
| 738.04000 | exclusion1022 |
| 738.38000 | exclusion1023 |
| 739.35000 | exclusion1024 |
| 739.86000 | exclusion1025 |
| 740.28000 | exclusion1026 |
| 740.87000 | exclusion1027 |
| 741.37000 | exclusion1028 |
| 742.37000 | exclusion1029 |
| 742.83000 | exclusion1030 |
| 743.06000 | exclusion1031 |
| 743.38000 | exclusion1032 |
| 743.85000 | exclusion1033 |
| 744.25000 | exclusion1034 |
| 744.38000 | exclusion1035 |
| 744.86000 | exclusion1036 |
| 746.28000 | exclusion1037 |
| 746.72000 | exclusion1038 |
| 747.03000 | exclusion1039 |
| 747.73000 | exclusion1040 |
| 747.83000 | exclusion1041 |
| 748.07000 | exclusion1042 |
| 748.32000 | exclusion1043 |
| 748.85000 | exclusion1044 |
| 749.04000 | exclusion1045 |
| 749.38000 | exclusion1046 |
| 749.61000 | exclusion1047 |
| 750.36000 | exclusion1048 |
| 750.37000 | exclusion1049 |
| 750.87000 | exclusion1050 |
| 751.37000 | exclusion1051 |
| 751.38000 | exclusion1052 |
| 751.86000 | exclusion1053 |
| 752.11000 | exclusion1054 |
| 752.36000 | exclusion1055 |
| 753.10000 | exclusion1056 |
| 753.37000 | exclusion1057 |
| 753.38000 | exclusion1058 |
| 754.32000 | exclusion1059 |
| 754.83000 | exclusion1060 |
| 755.34000 | exclusion1061 |
| 755.73000 | exclusion1062 |
| 755.86000 | exclusion1063 |

| | |
|---|---|
| 756.08000 | exclusion1064 |
| 756.32000 | exclusion1065 |
| 756.71000 | exclusion1066 |
| 757.36000 | exclusion1067 |
| 757.40000 | exclusion1068 |
| 757.70000 | exclusion1069 |
| 757.88000 | exclusion1070 |
| 758.84000 | exclusion1071 |
| 758.89000 | exclusion1072 |
| 759.37000 | exclusion1073 |
| 759.39000 | exclusion1074 |
| 759.68000 | exclusion1075 |
| 759.86000 | exclusion1076 |
| 760.36000 | exclusion1077 |
| 760.41000 | exclusion1078 |
| 760.73000 | exclusion1079 |
| 760.91000 | exclusion1080 |
| 761.34000 | exclusion1081 |
| 761.90000 | exclusion1082 |
| 762.40000 | exclusion1083 |
| 762.40000 | exclusion1084 |
| 762.88000 | exclusion1085 |
| 763.34000 | exclusion1086 |
| 763.90000 | exclusion1087 |
| 764.21000 | exclusion1088 |
| 764.36000 | exclusion1089 |
| 764.73000 | exclusion1090 |
| 765.86000 | exclusion1091 |
| 766.05000 | exclusion1092 |
| 766.35000 | exclusion1093 |
| 766.39000 | exclusion1094 |
| 767.87000 | exclusion1095 |
| 768.35000 | exclusion1096 |
| 768.66000 | exclusion1097 |
| 768.90000 | exclusion1098 |
| 769.31000 | exclusion1099 |
| 769.90000 | exclusion1100 |
| 770.37000 | exclusion1101 |
| 770.86000 | exclusion1102 |
| 771.02000 | exclusion1103 |
| 771.38000 | exclusion1104 |
| 771.88000 | exclusion1105 |
| 773.36000 | exclusion1106 |
| 773.37000 | exclusion1107 |
| 773.87000 | exclusion1108 |
| 774.30000 | exclusion1109 |
| 774.61000 | exclusion1110 |
| 775.34000 | exclusion1111 |
| 775.39000 | exclusion1112 |
| 775.83000 | exclusion1113 |
| 776.37000 | exclusion1114 |

| | |
|---|---|
| 776.58000 | exclusion1115 |
| 776.86000 | exclusion1116 |
| 777.39000 | exclusion1117 |
| 777.40000 | exclusion1118 |
| 777.83000 | exclusion1119 |
| 778.40000 | exclusion1120 |
| 778.91000 | exclusion1121 |
| 779.11000 | exclusion1122 |
| 779.34000 | exclusion1123 |
| 779.37000 | exclusion1124 |
| 779.86000 | exclusion1125 |
| 780.75000 | exclusion1126 |
| 780.81000 | exclusion1127 |
| 781.08000 | exclusion1128 |
| 781.16000 | exclusion1129 |
| 781.39000 | exclusion1130 |
| 782.28000 | exclusion1131 |
| 783.30000 | exclusion1132 |
| 783.41000 | exclusion1133 |
| 784.35000 | exclusion1134 |
| 784.87000 | exclusion1135 |
| 785.30000 | exclusion1136 |
| 785.65000 | exclusion1137 |
| 786.39000 | exclusion1138 |
| 786.88000 | exclusion1139 |
| 787.10000 | exclusion1140 |
| 787.36000 | exclusion1141 |
| 787.90000 | exclusion1142 |
| 788.71000 | exclusion1143 |
| 789.04000 | exclusion1144 |
| 789.37000 | exclusion1145 |
| 790.36000 | exclusion1146 |
| 790.38000 | exclusion1147 |
| 791.06000 | exclusion1148 |
| 791.37000 | exclusion1149 |
| 792.29000 | exclusion1150 |
| 792.38000 | exclusion1151 |
| 792.91000 | exclusion1152 |
| 792.91000 | exclusion1153 |
| 793.38000 | exclusion1154 |
| 793.87000 | exclusion1155 |
| 793.94000 | exclusion1156 |
| 794.38000 | exclusion1157 |
| 794.40000 | exclusion1158 |
| 795.36000 | exclusion1159 |
| 795.42000 | exclusion1160 |
| 795.90000 | exclusion1161 |
| 796.12000 | exclusion1162 |
| 796.37000 | exclusion1163 |
| 796.40000 | exclusion1164 |
| 796.88000 | exclusion1165 |

| | |
|---|---|
| 797.43000 | exclusion1166 |
| 797.87000 | exclusion1167 |
| 798.35000 | exclusion1168 |
| 798.41000 | exclusion1169 |
| 799.13000 | exclusion1170 |
| 799.87000 | exclusion1171 |
| 800.35000 | exclusion1172 |
| 800.38000 | exclusion1173 |
| 800.68000 | exclusion1174 |
| 801.38000 | exclusion1175 |
| 801.38000 | exclusion1176 |
| 801.95000 | exclusion1177 |
| 802.27000 | exclusion1178 |
| 802.39000 | exclusion1179 |
| 802.69000 | exclusion1180 |
| 802.82000 | exclusion1181 |
| 803.92000 | exclusion1182 |
| 804.40000 | exclusion1183 |
| 804.41000 | exclusion1184 |
| 804.74000 | exclusion1185 |
| 805.00000 | exclusion1186 |
| 805.40000 | exclusion1187 |
| 806.39000 | exclusion1188 |
| 806.43000 | exclusion1189 |
| 806.88000 | exclusion1190 |
| 807.02000 | exclusion1191 |
| 808.10000 | exclusion1192 |
| 808.36000 | exclusion1193 |
| 808.36000 | exclusion1194 |
| 808.94000 | exclusion1195 |
| 809.40000 | exclusion1196 |
| 809.91000 | exclusion1197 |
| 810.39000 | exclusion1198 |
| 810.39000 | exclusion1199 |
| 810.75000 | exclusion1200 |
| 811.36000 | exclusion1201 |
| 812.38000 | exclusion1202 |
| 812.73000 | exclusion1203 |
| 813.37000 | exclusion1204 |
| 813.43000 | exclusion1205 |
| 815.36000 | exclusion1206 |
| 815.41000 | exclusion1207 |
| 815.93000 | exclusion1208 |
| 817.05000 | exclusion1209 |
| 817.33000 | exclusion1210 |
| 817.75000 | exclusion1211 |
| 817.86000 | exclusion1212 |
| 818.43000 | exclusion1213 |
| 818.55000 | exclusion1214 |
| 818.87000 | exclusion1215 |
| 819.04000 | exclusion1216 |

| | |
|---|---|
| 819.42000 | exclusion1217 |
| 820.37000 | exclusion1218 |
| 821.15000 | exclusion1219 |
| 821.39000 | exclusion1220 |
| 822.38000 | exclusion1221 |
| 822.39000 | exclusion1222 |
| 822.91000 | exclusion1223 |
| 822.92000 | exclusion1224 |
| 823.57000 | exclusion1225 |
| 824.40000 | exclusion1226 |
| 824.81000 | exclusion1227 |
| 825.40000 | exclusion1228 |
| 825.85000 | exclusion1229 |
| 826.22000 | exclusion1230 |
| 826.38000 | exclusion1231 |
| 826.44000 | exclusion1232 |
| 827.42000 | exclusion1233 |
| 827.42000 | exclusion1234 |
| 827.89000 | exclusion1235 |
| 828.41000 | exclusion1236 |
| 828.44000 | exclusion1237 |
| 829.41000 | exclusion1238 |
| 829.43000 | exclusion1239 |
| 830.43000 | exclusion1240 |
| 830.92000 | exclusion1241 |
| 830.96000 | exclusion1242 |
| 831.13000 | exclusion1243 |
| 831.32000 | exclusion1244 |
| 831.94000 | exclusion1245 |
| 832.41000 | exclusion1246 |
| 832.44000 | exclusion1247 |
| 832.88000 | exclusion1248 |
| 833.15000 | exclusion1249 |
| 833.40000 | exclusion1250 |
| 833.78000 | exclusion1251 |
| 834.16000 | exclusion1252 |
| 834.33000 | exclusion1253 |
| 834.43000 | exclusion1254 |
| 834.92000 | exclusion1255 |
| 835.04000 | exclusion1256 |
| 835.13000 | exclusion1257 |
| 836.41000 | exclusion1258 |
| 836.42000 | exclusion1259 |
| 837.42000 | exclusion1260 |
| 837.44000 | exclusion1261 |
| 837.76000 | exclusion1262 |
| 837.93000 | exclusion1263 |
| 838.42000 | exclusion1264 |
| 839.16000 | exclusion1265 |
| 839.25000 | exclusion1266 |
| 839.43000 | exclusion1267 |

| | |
|---|---|
| 839.91000 | exclusion1268 |
| 840.42000 | exclusion1269 |
| 840.67000 | exclusion1270 |
| 840.73000 | exclusion1271 |
| 841.44000 | exclusion1272 |
| 841.45000 | exclusion1273 |
| 842.91000 | exclusion1274 |
| 842.92000 | exclusion1275 |
| 843.40000 | exclusion1276 |
| 843.64000 | exclusion1277 |
| 843.91000 | exclusion1278 |
| 843.94000 | exclusion1279 |
| 845.41000 | exclusion1280 |
| 845.87000 | exclusion1281 |
| 846.16000 | exclusion1282 |
| 846.36000 | exclusion1283 |
| 847.37000 | exclusion1284 |
| 847.39000 | exclusion1285 |
| 847.88000 | exclusion1286 |
| 848.36000 | exclusion1287 |
| 849.41000 | exclusion1288 |
| 849.44000 | exclusion1289 |
| 849.89000 | exclusion1290 |
| 849.90000 | exclusion1291 |
| 850.42000 | exclusion1292 |
| 850.91000 | exclusion1293 |
| 851.17000 | exclusion1294 |
| 851.42000 | exclusion1295 |
| 851.45000 | exclusion1296 |
| 851.78000 | exclusion1297 |
| 852.38000 | exclusion1298 |
| 852.39000 | exclusion1299 |
| 852.88000 | exclusion1300 |
| 853.23000 | exclusion1301 |
| 853.35000 | exclusion1302 |
| 853.66000 | exclusion1303 |
| 854.38000 | exclusion1304 |
| 855.37000 | exclusion1305 |
| 855.40000 | exclusion1306 |
| 855.88000 | exclusion1307 |
| 856.15000 | exclusion1308 |
| 856.40000 | exclusion1309 |
| 857.06000 | exclusion1310 |
| 857.45000 | exclusion1311 |
| 857.89000 | exclusion1312 |
| 858.50000 | exclusion1313 |
| 858.74000 | exclusion1314 |
| 859.47000 | exclusion1315 |
| 859.64000 | exclusion1316 |
| 860.08000 | exclusion1317 |
| 860.39000 | exclusion1318 |

| | |
|---|---|
| 860.88000 | exclusion1319 |
| 861.44000 | exclusion1320 |
| 861.90000 | exclusion1321 |
| 862.46000 | exclusion1322 |
| 862.49000 | exclusion1323 |
| 862.90000 | exclusion1324 |
| 863.39000 | exclusion1325 |
| 863.41000 | exclusion1326 |
| 864.44000 | exclusion1327 |
| 864.45000 | exclusion1328 |
| 864.88000 | exclusion1329 |
| 865.91000 | exclusion1330 |
| 866.41000 | exclusion1331 |
| 866.42000 | exclusion1332 |
| 867.45000 | exclusion1333 |
| 867.75000 | exclusion1334 |
| 868.13000 | exclusion1335 |
| 868.40000 | exclusion1336 |
| 868.40000 | exclusion1337 |
| 868.90000 | exclusion1338 |
| 868.94000 | exclusion1339 |
| 869.22000 | exclusion1340 |
| 869.73000 | exclusion1341 |
| 870.18000 | exclusion1342 |
| 870.42000 | exclusion1343 |
| 870.43000 | exclusion1344 |
| 871.35000 | exclusion1345 |
| 872.38000 | exclusion1346 |
| 872.43000 | exclusion1347 |
| 872.70000 | exclusion1348 |
| 872.92000 | exclusion1349 |
| 873.45000 | exclusion1350 |
| 874.05000 | exclusion1351 |
| 874.42000 | exclusion1352 |
| 874.95000 | exclusion1353 |
| 875.39000 | exclusion1354 |
| 875.90000 | exclusion1355 |
| 876.01000 | exclusion1356 |
| 876.40000 | exclusion1357 |
| 876.40000 | exclusion1358 |
| 877.36000 | exclusion1359 |
| 877.40000 | exclusion1360 |
| 878.05000 | exclusion1361 |
| 878.48000 | exclusion1362 |
| 879.05000 | exclusion1363 |
| 879.41000 | exclusion1364 |
| 879.93000 | exclusion1365 |
| 880.40000 | exclusion1366 |
| 880.41000 | exclusion1367 |
| 880.73000 | exclusion1368 |
| 881.43000 | exclusion1369 |

| | |
|---|---|
| 882.44000 | exclusion1370 |
| 882.46000 | exclusion1371 |
| 882.91000 | exclusion1372 |
| 883.44000 | exclusion1373 |
| 884.41000 | exclusion1374 |
| 884.47000 | exclusion1375 |
| 884.89000 | exclusion1376 |
| 885.42000 | exclusion1377 |
| 885.45000 | exclusion1378 |
| 885.93000 | exclusion1379 |
| 886.40000 | exclusion1380 |
| 886.95000 | exclusion1381 |
| 887.48000 | exclusion1382 |
| 887.96000 | exclusion1383 |
| 888.40000 | exclusion1384 |
| 888.42000 | exclusion1385 |
| 888.74000 | exclusion1386 |
| 889.47000 | exclusion1387 |
| 890.41000 | exclusion1388 |
| 890.66000 | exclusion1389 |
| 890.88000 | exclusion1390 |
| 891.41000 | exclusion1391 |
| 891.90000 | exclusion1392 |
| 892.24000 | exclusion1393 |
| 892.45000 | exclusion1394 |
| 893.44000 | exclusion1395 |
| 893.89000 | exclusion1396 |
| 894.18000 | exclusion1397 |
| 894.44000 | exclusion1398 |
| 895.42000 | exclusion1399 |
| 895.48000 | exclusion1400 |
| 895.89000 | exclusion1401 |
| 896.16000 | exclusion1402 |
| 896.43000 | exclusion1403 |
| 896.45000 | exclusion1404 |
| 897.41000 | exclusion1405 |
| 898.10000 | exclusion1406 |
| 898.42000 | exclusion1407 |
| 898.82000 | exclusion1408 |
| 899.44000 | exclusion1409 |
| 899.45000 | exclusion1410 |
| 900.42000 | exclusion1411 |
| 900.43000 | exclusion1412 |
| 901.14000 | exclusion1413 |
| 901.45000 | exclusion1414 |
| 902.42000 | exclusion1415 |
| 902.70000 | exclusion1416 |
| 903.13000 | exclusion1417 |
| 903.41000 | exclusion1418 |
| 904.79000 | exclusion1419 |
| 905.13000 | exclusion1420 |

| | |
|---|---|
| 905.40000 | exclusion1421 |
| 905.96000 | exclusion1422 |
| 906.41000 | exclusion1423 |
| 907.42000 | exclusion1424 |
| 907.45000 | exclusion1425 |
| 907.96000 | exclusion1426 |
| 908.40000 | exclusion1427 |
| 909.44000 | exclusion1428 |
| 909.44000 | exclusion1429 |
| 909.94000 | exclusion1430 |
| 911.13000 | exclusion1431 |
| 911.46000 | exclusion1432 |
| 911.69000 | exclusion1433 |
| 911.94000 | exclusion1434 |
| 911.97000 | exclusion1435 |
| 912.41000 | exclusion1436 |
| 913.66000 | exclusion1437 |
| 913.70000 | exclusion1438 |
| 913.92000 | exclusion1439 |
| 914.43000 | exclusion1440 |
| 914.47000 | exclusion1441 |
| 915.12000 | exclusion1442 |
| 915.45000 | exclusion1443 |
| 915.95000 | exclusion1444 |
| 916.46000 | exclusion1445 |
| 916.49000 | exclusion1446 |
| 916.75000 | exclusion1447 |
| 917.08000 | exclusion1448 |
| 917.47000 | exclusion1449 |
| 917.93000 | exclusion1450 |
| 918.11000 | exclusion1451 |
| 918.46000 | exclusion1452 |
| 918.48000 | exclusion1453 |
| 918.92000 | exclusion1454 |
| 919.43000 | exclusion1455 |
| 919.54000 | exclusion1456 |
| 920.42000 | exclusion1457 |
| 920.49000 | exclusion1458 |
| 920.87000 | exclusion1459 |
| 921.40000 | exclusion1460 |
| 921.47000 | exclusion1461 |
| 922.30000 | exclusion1462 |
| 922.47000 | exclusion1463 |
| 922.90000 | exclusion1464 |
| 923.46000 | exclusion1465 |
| 923.91000 | exclusion1466 |
| 924.13000 | exclusion1467 |
| 924.43000 | exclusion1468 |
| 924.94000 | exclusion1469 |
| 925.49000 | exclusion1470 |
| 926.46000 | exclusion1471 |

| | |
|---|---|
| 926.79000 | exclusion1472 |
| 927.82000 | exclusion1473 |
| 927.88000 | exclusion1474 |
| 928.47000 | exclusion1475 |
| 928.93000 | exclusion1476 |
| 929.45000 | exclusion1477 |
| 929.77000 | exclusion1478 |
| 930.46000 | exclusion1479 |
| 930.47000 | exclusion1480 |
| 930.68000 | exclusion1481 |
| 931.11000 | exclusion1482 |
| 931.40000 | exclusion1483 |
| 931.41000 | exclusion1484 |
| 931.81000 | exclusion1485 |
| 932.43000 | exclusion1486 |
| 933.47000 | exclusion1487 |
| 933.79000 | exclusion1488 |
| 934.39000 | exclusion1489 |
| 934.97000 | exclusion1490 |
| 935.16000 | exclusion1491 |
| 935.47000 | exclusion1492 |
| 935.91000 | exclusion1493 |
| 936.11000 | exclusion1494 |
| 937.49000 | exclusion1495 |
| 937.52000 | exclusion1496 |
| 937.79000 | exclusion1497 |
| 937.99000 | exclusion1498 |
| 938.49000 | exclusion1499 |
| 939.21000 | exclusion1500 |
| 939.49000 | exclusion1501 |
| 939.93000 | exclusion1502 |
| 941.46000 | exclusion1503 |
| 942.47000 | exclusion1504 |
| 942.98000 | exclusion1505 |
| 943.10000 | exclusion1506 |
| 943.47000 | exclusion1507 |
| 943.73000 | exclusion1508 |
| 944.47000 | exclusion1509 |
| 944.74000 | exclusion1510 |
| 944.92000 | exclusion1511 |
| 945.48000 | exclusion1512 |
| 945.78000 | exclusion1513 |
| 945.98000 | exclusion1514 |
| 946.95000 | exclusion1515 |
| 947.49000 | exclusion1516 |
| 947.96000 | exclusion1517 |
| 948.46000 | exclusion1518 |
| 949.02000 | exclusion1519 |
| 949.46000 | exclusion1520 |
| 949.80000 | exclusion1521 |
| 950.01000 | exclusion1522 |

| | |
|---|---|
| 950.70000 | exclusion1523 |
| 951.13000 | exclusion1524 |
| 951.41000 | exclusion1525 |
| 951.47000 | exclusion1526 |
| 951.82000 | exclusion1527 |
| 952.14000 | exclusion1528 |
| 952.77000 | exclusion1529 |
| 952.96000 | exclusion1530 |
| 953.01000 | exclusion1531 |
| 953.48000 | exclusion1532 |
| 954.44000 | exclusion1533 |
| 954.94000 | exclusion1534 |
| 955.35000 | exclusion1535 |
| 955.57000 | exclusion1536 |
| 956.47000 | exclusion1537 |
| 957.43000 | exclusion1538 |
| 957.74000 | exclusion1539 |
| 958.44000 | exclusion1540 |
| 958.81000 | exclusion1541 |
| 959.41000 | exclusion1542 |
| 960.45000 | exclusion1543 |
| 960.75000 | exclusion1544 |
| 961.00000 | exclusion1545 |
| 961.35000 | exclusion1546 |
| 962.49000 | exclusion1547 |
| 962.87000 | exclusion1548 |
| 963.48000 | exclusion1549 |
| 963.94000 | exclusion1550 |
| 964.44000 | exclusion1551 |
| 964.74000 | exclusion1552 |
| 964.98000 | exclusion1553 |
| 965.02000 | exclusion1554 |
| 965.48000 | exclusion1555 |
| 966.26000 | exclusion1556 |
| 966.48000 | exclusion1557 |
| 967.21000 | exclusion1558 |
| 967.48000 | exclusion1559 |
| 967.48000 | exclusion1560 |
| 967.90000 | exclusion1561 |
| 968.47000 | exclusion1562 |
| 969.50000 | exclusion1563 |
| 969.98000 | exclusion1564 |
| 970.11000 | exclusion1565 |
| 970.45000 | exclusion1566 |
| 970.52000 | exclusion1567 |
| 971.48000 | exclusion1568 |
| 971.96000 | exclusion1569 |
| 972.15000 | exclusion1570 |
| 972.44000 | exclusion1571 |
| 972.48000 | exclusion1572 |
| 972.54000 | exclusion1573 |

| | |
|---|---|
| 973.00000 | exclusion1574 |
| 973.80000 | exclusion1575 |
| 974.47000 | exclusion1576 |
| 975.47000 | exclusion1577 |
| 975.49000 | exclusion1578 |
| 975.82000 | exclusion1579 |
| 976.68000 | exclusion1580 |
| 977.14000 | exclusion1581 |
| 977.49000 | exclusion1582 |
| 977.78000 | exclusion1583 |
| 978.46000 | exclusion1584 |
| 978.51000 | exclusion1585 |
| 979.42000 | exclusion1586 |
| 980.13000 | exclusion1587 |
| 980.51000 | exclusion1588 |
| 980.97000 | exclusion1589 |
| 981.21000 | exclusion1590 |
| 981.30000 | exclusion1591 |
| 981.54000 | exclusion1592 |
| 982.46000 | exclusion1593 |
| 983.45000 | exclusion1594 |
| 983.49000 | exclusion1595 |
| 984.48000 | exclusion1596 |
| 984.50000 | exclusion1597 |
| 984.94000 | exclusion1598 |
| 985.87000 | exclusion1599 |
| 986.00000 | exclusion1600 |
| 986.47000 | exclusion1601 |
| 986.83000 | exclusion1602 |
| 987.01000 | exclusion1603 |
| 987.51000 | exclusion1604 |
| 987.97000 | exclusion1605 |
| 988.18000 | exclusion1606 |
| 988.49000 | exclusion1607 |
| 989.01000 | exclusion1608 |
| 990.02000 | exclusion1609 |
| 990.14000 | exclusion1610 |
| 990.47000 | exclusion1611 |
| 990.50000 | exclusion1612 |
| 991.16000 | exclusion1613 |
| 991.20000 | exclusion1614 |
| 991.48000 | exclusion1615 |
| 991.83000 | exclusion1616 |
| 991.99000 | exclusion1617 |
| 993.52000 | exclusion1618 |
| 993.96000 | exclusion1619 |
| 995.49000 | exclusion1620 |
| 996.48000 | exclusion1621 |
| 997.50000 | exclusion1622 |
| 998.48000 | exclusion1623 |
| 998.81000 | exclusion1624 |

999.39000 exclusion1625
999.82000 exclusion1626
1000.46000 exclusion1627
1000.84000 exclusion1628
1001.50000 exclusion1629
1002.48000 exclusion1630
1003.01000 exclusion1631
1003.53000 exclusion1632
1004.48000 exclusion1633
1004.94000 exclusion1634
1005.45000 exclusion1635
1006.52000 exclusion1636
1006.98000 exclusion1637
1007.95000 exclusion1638
1008.80000 exclusion1639
1009.06000 exclusion1640
1009.92000 exclusion1641
1010.38000 exclusion1642
1011.02000 exclusion1643
1011.77000 exclusion1644
1012.48000 exclusion1645
1012.94000 exclusion1646
1014.47000 exclusion1647
1015.42000 exclusion1648
1016.01000 exclusion1649
1016.49000 exclusion1650
1017.02000 exclusion1651
1017.53000 exclusion1652
1017.91000 exclusion1653
1018.95000 exclusion1654
1019.21000 exclusion1655
1020.05000 exclusion1656
1020.49000 exclusion1657
1021.00000 exclusion1658
1021.53000 exclusion1659
1022.46000 exclusion1660
1022.82000 exclusion1661
1023.45000 exclusion1662
1024.18000 exclusion1663
1024.51000 exclusion1664
1025.42000 exclusion1665
1025.86000 exclusion1666
1026.79000 exclusion1667
1027.46000 exclusion1668
1028.02000 exclusion1669
1028.98000 exclusion1670
1030.46000 exclusion1671
1031.47000 exclusion1672
1032.46000 exclusion1673
1032.86000 exclusion1674
1033.15000 exclusion1675

| | |
|---|---|
| 1033.52000 | exclusion1676 |
| 1034.02000 | exclusion1677 |
| 1035.55000 | exclusion1678 |
| 1036.20000 | exclusion1679 |
| 1036.53000 | exclusion1680 |
| 1036.77000 | exclusion1681 |
| 1037.45000 | exclusion1682 |
| 1038.98000 | exclusion1683 |
| 1039.29000 | exclusion1684 |
| 1039.50000 | exclusion1685 |
| 1040.04000 | exclusion1686 |
| 1040.51000 | exclusion1687 |
| 1041.47000 | exclusion1688 |
| 1041.77000 | exclusion1689 |
| 1042.02000 | exclusion1690 |
| 1042.52000 | exclusion1691 |
| 1043.14000 | exclusion1692 |
| 1044.17000 | exclusion1693 |
| 1044.52000 | exclusion1694 |
| 1044.97000 | exclusion1695 |
| 1045.46000 | exclusion1696 |
| 1046.50000 | exclusion1697 |
| 1047.82000 | exclusion1698 |
| 1048.05000 | exclusion1699 |
| 1048.84000 | exclusion1700 |
| 1049.47000 | exclusion1701 |
| 1049.87000 | exclusion1702 |
| 1050.51000 | exclusion1703 |
| 1050.77000 | exclusion1704 |
| 1051.47000 | exclusion1705 |
| 1052.53000 | exclusion1706 |
| 1053.06000 | exclusion1707 |
| 1053.53000 | exclusion1708 |
| 1053.84000 | exclusion1709 |
| 1054.50000 | exclusion1710 |
| 1055.04000 | exclusion1711 |
| 1055.48000 | exclusion1712 |
| 1056.04000 | exclusion1713 |
| 1056.52000 | exclusion1714 |
| 1057.56000 | exclusion1715 |
| 1057.84000 | exclusion1716 |
| 1058.07000 | exclusion1717 |
| 1058.48000 | exclusion1718 |
| 1059.27000 | exclusion1719 |
| 1059.48000 | exclusion1720 |
| 1060.03000 | exclusion1721 |
| 1060.53000 | exclusion1722 |
| 1061.02000 | exclusion1723 |
| 1061.83000 | exclusion1724 |
| 1062.24000 | exclusion1725 |
| 1062.54000 | exclusion1726 |

| | |
|---|---|
| 1063.01000 | exclusion1727 |
| 1063.48000 | exclusion1728 |
| 1064.48000 | exclusion1729 |
| 1065.47000 | exclusion1730 |
| 1067.04000 | exclusion1731 |
| 1067.49000 | exclusion1732 |
| 1068.04000 | exclusion1733 |
| 1068.47000 | exclusion1734 |
| 1070.06000 | exclusion1735 |
| 1070.52000 | exclusion1736 |
| 1071.15000 | exclusion1737 |
| 1071.54000 | exclusion1738 |
| 1072.87000 | exclusion1739 |
| 1073.53000 | exclusion1740 |
| 1074.04000 | exclusion1741 |
| 1075.53000 | exclusion1742 |
| 1076.06000 | exclusion1743 |
| 1077.04000 | exclusion1744 |
| 1077.54000 | exclusion1745 |
| 1078.99000 | exclusion1746 |
| 1079.51000 | exclusion1747 |
| 1080.50000 | exclusion1748 |
| 1080.85000 | exclusion1749 |
| 1081.53000 | exclusion1750 |
| 1082.19000 | exclusion1751 |
| 1082.53000 | exclusion1752 |
| 1083.58000 | exclusion1753 |
| 1084.06000 | exclusion1754 |
| 1084.55000 | exclusion1755 |
| 1085.04000 | exclusion1756 |
| 1085.53000 | exclusion1757 |
| 1086.05000 | exclusion1758 |
| 1086.49000 | exclusion1759 |
| 1087.55000 | exclusion1760 |
| 1087.88000 | exclusion1761 |
| 1088.24000 | exclusion1762 |
| 1088.49000 | exclusion1763 |
| 1090.06000 | exclusion1764 |
| 1090.53000 | exclusion1765 |
| 1091.06000 | exclusion1766 |
| 1091.89000 | exclusion1767 |
| 1093.53000 | exclusion1768 |
| 1093.87000 | exclusion1769 |
| 1094.55000 | exclusion1770 |
| 1095.50000 | exclusion1771 |
| 1095.86000 | exclusion1772 |
| 1096.12000 | exclusion1773 |
| 1097.29000 | exclusion1774 |
| 1097.54000 | exclusion1775 |
| 1098.20000 | exclusion1776 |
| 1099.53000 | exclusion1777 |

| | |
|---|---|
| 1101.09000 | exclusion1778 |
| 1102.22000 | exclusion1779 |
| 1102.51000 | exclusion1780 |
| 1102.88000 | exclusion1781 |
| 1103.99000 | exclusion1782 |
| 1104.22000 | exclusion1783 |
| 1104.58000 | exclusion1784 |
| 1105.54000 | exclusion1785 |
| 1105.77000 | exclusion1786 |
| 1106.53000 | exclusion1787 |
| 1107.54000 | exclusion1788 |
| 1108.58000 | exclusion1789 |
| 1109.52000 | exclusion1790 |
| 1109.88000 | exclusion1791 |
| 1110.23000 | exclusion1792 |
| 1111.56000 | exclusion1793 |
| 1111.87000 | exclusion1794 |
| 1112.08000 | exclusion1795 |
| 1112.80000 | exclusion1796 |
| 1113.00000 | exclusion1797 |
| 1113.52000 | exclusion1798 |
| 1114.94000 | exclusion1799 |
| 1115.55000 | exclusion1800 |
| 1116.08000 | exclusion1801 |
| 1116.88000 | exclusion1802 |
| 1117.52000 | exclusion1803 |
| 1118.09000 | exclusion1804 |
| 1118.58000 | exclusion1805 |
| 1118.86000 | exclusion1806 |
| 1119.89000 | exclusion1807 |
| 1120.52000 | exclusion1808 |
| 1121.05000 | exclusion1809 |
| 1122.19000 | exclusion1810 |
| 1122.51000 | exclusion1811 |
| 1123.21000 | exclusion1812 |
| 1123.56000 | exclusion1813 |
| 1124.57000 | exclusion1814 |
| 1125.02000 | exclusion1815 |
| 1125.54000 | exclusion1816 |
| 1126.04000 | exclusion1817 |
| 1127.55000 | exclusion1818 |
| 1127.87000 | exclusion1819 |
| 1128.09000 | exclusion1820 |
| 1128.53000 | exclusion1821 |
| 1130.21000 | exclusion1822 |
| 1130.58000 | exclusion1823 |
| 1131.03000 | exclusion1824 |
| 1131.57000 | exclusion1825 |
| 1132.20000 | exclusion1826 |
| 1132.87000 | exclusion1827 |
| 1133.08000 | exclusion1828 |

| | |
|---|---|
| 1133.62000 | exclusion1829 |
| 1133.89000 | exclusion1830 |
| 1134.54000 | exclusion1831 |
| 1135.11000 | exclusion1832 |
| 1136.04000 | exclusion1833 |
| 1136.53000 | exclusion1834 |
| 1137.44000 | exclusion1835 |
| 1138.55000 | exclusion1836 |
| 1139.18000 | exclusion1837 |
| 1139.54000 | exclusion1838 |
| 1141.52000 | exclusion1839 |
| 1141.90000 | exclusion1840 |
| 1142.51000 | exclusion1841 |
| 1143.05000 | exclusion1842 |
| 1143.56000 | exclusion1843 |
| 1145.04000 | exclusion1844 |
| 1145.53000 | exclusion1845 |
| 1146.56000 | exclusion1846 |
| 1147.00000 | exclusion1847 |
| 1147.53000 | exclusion1848 |
| 1148.07000 | exclusion1849 |
| 1148.56000 | exclusion1850 |
| 1149.06000 | exclusion1851 |
| 1149.56000 | exclusion1852 |
| 1150.56000 | exclusion1853 |
| 1151.05000 | exclusion1854 |
| 1151.60000 | exclusion1855 |
| 1152.12000 | exclusion1856 |
| 1152.55000 | exclusion1857 |
| 1153.54000 | exclusion1858 |
| 1154.20000 | exclusion1859 |
| 1154.55000 | exclusion1860 |
| 1155.08000 | exclusion1861 |
| 1156.07000 | exclusion1862 |
| 1156.52000 | exclusion1863 |
| 1157.56000 | exclusion1864 |
| 1158.23000 | exclusion1865 |
| 1158.51000 | exclusion1866 |
| 1159.44000 | exclusion1867 |
| 1159.64000 | exclusion1868 |
| 1159.91000 | exclusion1869 |
| 1161.57000 | exclusion1870 |
| 1162.04000 | exclusion1871 |
| 1162.58000 | exclusion1872 |
| 1163.03000 | exclusion1873 |
| 1164.56000 | exclusion1874 |
| 1165.57000 | exclusion1875 |
| 1167.20000 | exclusion1876 |
| 1167.52000 | exclusion1877 |
| 1168.20000 | exclusion1878 |
| 1168.55000 | exclusion1879 |

| | |
|---|---|
| 1169.58000 | exclusion1880 |
| 1171.09000 | exclusion1881 |
| 1171.61000 | exclusion1882 |
| 1173.27000 | exclusion1883 |
| 1173.56000 | exclusion1884 |
| 1174.54000 | exclusion1885 |
| 1175.48000 | exclusion1886 |
| 1176.59000 | exclusion1887 |
| 1177.58000 | exclusion1888 |
| 1178.09000 | exclusion1889 |
| 1178.50000 | exclusion1890 |
| 1179.58000 | exclusion1891 |
| 1180.14000 | exclusion1892 |
| 1181.56000 | exclusion1893 |
| 1182.54000 | exclusion1894 |
| 1182.93000 | exclusion1895 |
| 1184.10000 | exclusion1896 |
| 1184.55000 | exclusion1897 |
| 1185.54000 | exclusion1898 |
| 1187.10000 | exclusion1899 |
| 1187.58000 | exclusion1900 |
| 1188.57000 | exclusion1901 |
| 1189.54000 | exclusion1902 |
| 1189.91000 | exclusion1903 |
| 1192.62000 | exclusion1904 |
| 1192.89000 | exclusion1905 |
| 1193.59000 | exclusion1906 |
| 1195.03000 | exclusion1907 |
| 1195.25000 | exclusion1908 |
| 1196.25000 | exclusion1909 |
| 1196.54000 | exclusion1910 |
| 1197.55000 | exclusion1911 |
| 1198.55000 | exclusion1912 |
| 1200.23000 | exclusion1913 |
| 1200.56000 | exclusion1914 |
| 1200.84000 | exclusion1915 |
| 1201.11000 | exclusion1916 |
| 1202.57000 | exclusion1917 |
| 1203.50000 | exclusion1918 |
| 1204.54000 | exclusion1919 |
| 1204.91000 | exclusion1920 |
| 1205.63000 | exclusion1921 |
| 1206.03000 | exclusion1922 |
| 1206.59000 | exclusion1923 |
| 1207.88000 | exclusion1924 |
| 1208.87000 | exclusion1925 |
| 1209.91000 | exclusion1926 |
| 1210.64000 | exclusion1927 |
| 1211.01000 | exclusion1928 |
| 1211.60000 | exclusion1929 |
| 1212.25000 | exclusion1930 |

| | |
|---|---|
| 1213.61000 | exclusion1931 |
| 1214.58000 | exclusion1932 |
| 1215.65000 | exclusion1933 |
| 1216.12000 | exclusion1934 |
| 1217.86000 | exclusion1935 |
| 1218.54000 | exclusion1936 |
| 1219.58000 | exclusion1937 |
| 1221.12000 | exclusion1938 |
| 1223.05000 | exclusion1939 |
| 1223.57000 | exclusion1940 |
| 1224.08000 | exclusion1941 |
| 1224.59000 | exclusion1942 |
| 1225.92000 | exclusion1943 |
| 1226.62000 | exclusion1944 |
| 1227.13000 | exclusion1945 |
| 1228.28000 | exclusion1946 |
| 1228.82000 | exclusion1947 |
| 1230.06000 | exclusion1948 |
| 1232.05000 | exclusion1949 |
| 1233.30000 | exclusion1950 |
| 1234.12000 | exclusion1951 |
| 1235.07000 | exclusion1952 |
| 1235.58000 | exclusion1953 |
| 1237.12000 | exclusion1954 |
| 1241.05000 | exclusion1955 |
| 1241.64000 | exclusion1956 |
| 1241.87000 | exclusion1957 |
| 1245.11000 | exclusion1958 |
| 1245.92000 | exclusion1959 |
| 1246.97000 | exclusion1960 |
| 1247.36000 | exclusion1961 |
| 1247.59000 | exclusion1962 |
| 1248.57000 | exclusion1963 |
| 1249.04000 | exclusion1964 |
| 1250.60000 | exclusion1965 |
| 1252.19000 | exclusion1966 |
| 1252.59000 | exclusion1967 |
| 1253.07000 | exclusion1968 |
| 1253.59000 | exclusion1969 |
| 1254.63000 | exclusion1970 |
| 1256.79000 | exclusion1971 |
| 1257.09000 | exclusion1972 |
| 1258.57000 | exclusion1973 |
| 1259.21000 | exclusion1974 |
| 1260.63000 | exclusion1975 |
| 1264.16000 | exclusion1976 |
| 1265.08000 | exclusion1977 |
| 1265.59000 | exclusion1978 |
| 1265.97000 | exclusion1979 |
| 1270.27000 | exclusion1980 |
| 1271.18000 | exclusion1981 |

1272.68000 exclusion1982
1275.65000 exclusion1983
1276.64000 exclusion1984
1279.02000 exclusion1985
1279.61000 exclusion1986
1283.66000 exclusion1987
1284.64000 exclusion1988
1286.66000 exclusion1989
1287.08000 exclusion1990
1288.64000 exclusion1991
1289.45000 exclusion1992
1291.08000 exclusion1993
1292.39000 exclusion1994
1293.18000 exclusion1995
1294.61000 exclusion1996
1295.33000 exclusion1997
1298.08000 exclusion1998
1300.07000 exclusion1999
1301.06000 exclusion2000
Neutral Loss Mass List: (none)
Product Mass List: (none)
Neutral loss in top: 3
Product in top: 3
Most intense if no parent masses found not enabled
Add/subtract mass not enabled
FT master scan preview mode enabled
Charge state screening enabled
Charge state dependent ETD time not enabled
Monoisotopic precursor selection enabled
Charge state rejection enabled
Unassigned charge states : rejected
Charge state 1 : rejected
Charge state 2 : not rejected
Charge state 3 : not rejected
Charge states 4+ : not rejected
Chromatography mode is disabled
Global Data Dependent Settings:
Predict ion injection time enabled
Use global parent and reject mass lists not enabled
Exclude parent mass from data dependent selection not enabled
Exclusion mass width by mass
Exclusion mass width low: 1.50000
Exclusion mass width high: 1.50000
Parent mass width by mass
Parent mass width low: 0.50000
Parent mass width high: 0.50000
Reject mass width by mass
Reject mass width low: 0.50000
Reject mass width high: 0.50000
Zoom/UltraZoom scan mass width by mass
Zoom/UltraZoom scan mass width low: 5.00

Zoom/UltraZoom scan mass width high: 5.00
FT SIM scan mass width low: 5.00
FT SIM scan mass width high: 5.00
Neutral Loss candidates processed by decreasing intensity
Neutral Loss mass width by mass
Neutral Loss mass width low: 0.50000
Neutral Loss mass width high: 0.50000
Product candidates processed by decreasing intensity
Product mass width by mass
Product mass width low: 0.50000
Product mass width high: 0.50000
MS mass range: 300.00-2000.00
MSn mass range by mass
MSn mass range: 0.00-1000000.00
Use m/z values as masses not enabled
Analog UV data dep. not enabled
Dynamic exclusion enabled
Repeat Count: 1
Repeat Duration: 30.00
Exclusion List Size: 500
Exclusion Duration: 180.00
Exclusion mass width by mass
Exclusion mass width low: 1.50000
Exclusion mass width high: 1.50000
Expiration: disabled
Isotopic data dependence not enabled
Custom Data Dependent Settings:
Not enabled

---

Pass 3 (wheat-mixed-digests_MS2_pass03_1.raw):

Creator: Orbi_30393
Last modified: 10/12/2021 by Orbi_30393
MS Run Time (min): 43.00
Sequence override of method parameters not enabled.
Divert Valve: not used during run
Contact Closure: not used during run
Syringe Pump: not used during run
MS Detector Settings:
Real-time modifications to method not enabled
Stepped collision energy not enabled
Additional Microscans:
MS2 0 0
MS3 0 0
MS4 0 0
MS5 0 0
MS6 0 0
MS7 0 0
MS8 0 0
MS9 0 0
MS10 0 0

Experiment Type: Nth Order Double Play
Tune Method: Orbitrap-tune-file_2020-03-13_HESI
Scan Event Details:
1: FTMS + p norm o(300.0-2000.0)
CV = 0.0V
2: ITMS + c norm Dep MS/MS Most intense ion from (1)
Activation Type: CID
Min. Signal Required: 3000.0
Isolation Width: 2.00
Normalized Coll. Energy: 35.0
Default Charge State: 2
Activation Q: 0.250
Activation Time: 10.000
CV = 0.0V
Scan Event 2 repeated for top 10 peaks.
Lock Masses:
Pos List Name: N/A
Source: API Source
Mass List: (none)
Neg List Name: N/A
Source: API Source
Mass List: (none)
Data Dependent Settings:
Use separate polarity settings disabled
Parent Mass List: (none)
Reject Mass List:

| MS Mass | Name |
|---|---|
| 308.66000 | exclusion0001 |
| 311.10000 | exclusion0002 |
| 313.17000 | exclusion0003 |
| 315.47000 | exclusion0004 |
| 315.70000 | exclusion0005 |
| 321.24000 | exclusion0006 |
| 321.66000 | exclusion0007 |
| 321.68000 | exclusion0008 |
| 328.69000 | exclusion0009 |
| 330.83000 | exclusion0010 |
| 335.68000 | exclusion0011 |
| 338.67000 | exclusion0012 |
| 338.69000 | exclusion0013 |
| 339.17000 | exclusion0014 |
| 340.69000 | exclusion0015 |
| 340.83000 | exclusion0016 |
| 342.84000 | exclusion0017 |
| 345.13000 | exclusion0018 |
| 345.15000 | exclusion0019 |
| 346.71000 | exclusion0020 |
| 346.85000 | exclusion0021 |
| 346.93000 | exclusion0022 |
| 347.17000 | exclusion0023 |
| 353.18000 | exclusion0024 |

| | |
|---|---|
| 353.19000 | exclusion0025 |
| 356.69000 | exclusion0026 |
| 356.87000 | exclusion0027 |
| 357.19000 | exclusion0028 |
| 357.23000 | exclusion0029 |
| 368.22000 | exclusion0030 |
| 368.86000 | exclusion0031 |
| 369.16000 | exclusion0032 |
| 369.26000 | exclusion0033 |
| 372.22000 | exclusion0034 |
| 372.23000 | exclusion0035 |
| 376.69000 | exclusion0036 |
| 377.68000 | exclusion0037 |
| 378.22000 | exclusion0038 |
| 378.22000 | exclusion0039 |
| 378.52000 | exclusion0040 |
| 379.20000 | exclusion0041 |
| 380.70000 | exclusion0042 |
| 389.89000 | exclusion0043 |
| 390.19000 | exclusion0044 |
| 392.25000 | exclusion0045 |
| 396.18000 | exclusion0046 |
| 396.20000 | exclusion0047 |
| 402.23000 | exclusion0048 |
| 402.44000 | exclusion0049 |
| 402.53000 | exclusion0050 |
| 403.22000 | exclusion0051 |
| 407.76000 | exclusion0052 |
| 408.19000 | exclusion0053 |
| 409.20000 | exclusion0054 |
| 409.23000 | exclusion0055 |
| 412.75000 | exclusion0056 |
| 413.23000 | exclusion0057 |
| 416.71000 | exclusion0058 |
| 416.71000 | exclusion0059 |
| 420.22000 | exclusion0060 |
| 431.27000 | exclusion0061 |
| 434.72000 | exclusion0062 |
| 435.73000 | exclusion0063 |
| 438.20000 | exclusion0064 |
| 438.24000 | exclusion0065 |
| 439.24000 | exclusion0066 |
| 440.21000 | exclusion0067 |
| 440.26000 | exclusion0068 |
| 441.21000 | exclusion0069 |
| 442.25000 | exclusion0070 |
| 444.76000 | exclusion0071 |
| 444.87000 | exclusion0072 |
| 445.24000 | exclusion0073 |
| 446.23000 | exclusion0074 |
| 467.23000 | exclusion0075 |

| | |
|---|---|
| 467.23000 | exclusion0076 |
| 470.78000 | exclusion0077 |
| 470.91000 | exclusion0078 |
| 474.76000 | exclusion0079 |
| 475.22000 | exclusion0080 |
| 478.78000 | exclusion0081 |
| 478.90000 | exclusion0082 |
| 496.74000 | exclusion0083 |
| 497.49000 | exclusion0084 |
| 507.80000 | exclusion0085 |
| 507.88000 | exclusion0086 |
| 507.99000 | exclusion0087 |
| 508.26000 | exclusion0088 |
| 509.76000 | exclusion0089 |
| 510.28000 | exclusion0090 |
| 510.76000 | exclusion0091 |
| 532.77000 | exclusion0092 |
| 539.30000 | exclusion0093 |
| 539.72000 | exclusion0094 |
| 540.27000 | exclusion0095 |
| 544.76000 | exclusion0096 |
| 545.25000 | exclusion0097 |
| 548.77000 | exclusion0098 |
| 585.26000 | exclusion0099 |
| 585.28000 | exclusion0100 |
| 585.28000 | exclusion0101 |
| 585.65000 | exclusion0102 |
| 600.78000 | exclusion0103 |
| 602.84000 | exclusion0104 |
| 603.28000 | exclusion0105 |
| 604.79000 | exclusion0106 |
| 607.28000 | exclusion0107 |
| 607.30000 | exclusion0108 |
| 607.30000 | exclusion0109 |
| 608.32000 | exclusion0110 |
| 609.85000 | exclusion0111 |
| 612.32000 | exclusion0112 |
| 612.33000 | exclusion0113 |
| 612.35000 | exclusion0114 |
| 613.32000 | exclusion0115 |
| 625.93000 | exclusion0606 |
| 626.28000 | exclusion0607 |
| 626.81000 | exclusion0608 |
| 627.01000 | exclusion0609 |
| 627.30000 | exclusion0610 |
| 628.29000 | exclusion0611 |
| 628.58000 | exclusion0612 |
| 628.80000 | exclusion0613 |
| 628.96000 | exclusion0614 |
| 629.03000 | exclusion0615 |
| 629.82000 | exclusion0616 |

| | |
|---|---|
| 630.30000 | exclusion0617 |
| 630.77000 | exclusion0618 |
| 631.31000 | exclusion0619 |
| 631.83000 | exclusion0620 |
| 632.29000 | exclusion0621 |
| 632.83000 | exclusion0622 |
| 633.28000 | exclusion0623 |
| 633.31000 | exclusion0624 |
| 633.83000 | exclusion0625 |
| 634.33000 | exclusion0626 |
| 634.79000 | exclusion0627 |
| 635.29000 | exclusion0116 |
| 635.32000 | exclusion0117 |
| 635.86000 | exclusion0118 |
| 635.86000 | exclusion0628 |
| 636.32000 | exclusion0629 |
| 636.83000 | exclusion0630 |
| 637.64000 | exclusion0631 |
| 637.78000 | exclusion0632 |
| 637.85000 | exclusion0633 |
| 638.23000 | exclusion0634 |
| 638.26000 | exclusion0635 |
| 638.85000 | exclusion0636 |
| 639.35000 | exclusion0637 |
| 639.59000 | exclusion0638 |
| 639.61000 | exclusion0639 |
| 639.78000 | exclusion0640 |
| 639.92000 | exclusion0641 |
| 640.30000 | exclusion0642 |
| 640.61000 | exclusion0643 |
| 641.35000 | exclusion0644 |
| 641.44000 | exclusion0645 |
| 641.84000 | exclusion0646 |
| 642.35000 | exclusion0119 |
| 642.86000 | exclusion0647 |
| 643.29000 | exclusion0648 |
| 643.36000 | exclusion0649 |
| 643.82000 | exclusion0650 |
| 644.02000 | exclusion0651 |
| 644.87000 | exclusion0652 |
| 645.02000 | exclusion0653 |
| 645.35000 | exclusion0654 |
| 645.51000 | exclusion0655 |
| 645.70000 | exclusion0656 |
| 645.98000 | exclusion0657 |
| 646.80000 | exclusion0120 |
| 647.03000 | exclusion0121 |
| 647.31000 | exclusion0658 |
| 647.35000 | exclusion0659 |
| 647.87000 | exclusion0660 |
| 648.00000 | exclusion0661 |

| | |
|---|---|
| 648.29000 | exclusion0662 |
| 648.30000 | exclusion0663 |
| 648.31000 | exclusion0664 |
| 648.82000 | exclusion0665 |
| 649.28000 | exclusion0666 |
| 649.82000 | exclusion0667 |
| 650.29000 | exclusion0668 |
| 650.82000 | exclusion0669 |
| 651.30000 | exclusion0670 |
| 651.32000 | exclusion0671 |
| 651.78000 | exclusion0672 |
| 652.30000 | exclusion0122 |
| 652.32000 | exclusion0123 |
| 652.84000 | exclusion0673 |
| 653.24000 | exclusion0674 |
| 653.32000 | exclusion0124 |
| 653.35000 | exclusion0125 |
| 653.36000 | exclusion0675 |
| 653.63000 | exclusion0676 |
| 654.30000 | exclusion0677 |
| 654.35000 | exclusion0678 |
| 654.85000 | exclusion0679 |
| 655.09000 | exclusion0680 |
| 655.82000 | exclusion0681 |
| 656.32000 | exclusion0682 |
| 656.54000 | exclusion0683 |
| 657.03000 | exclusion0684 |
| 657.35000 | exclusion0685 |
| 657.38000 | exclusion0686 |
| 657.83000 | exclusion0687 |
| 658.03000 | exclusion0688 |
| 658.32000 | exclusion0689 |
| 658.35000 | exclusion0690 |
| 659.28000 | exclusion0691 |
| 659.34000 | exclusion0692 |
| 659.78000 | exclusion0693 |
| 660.30000 | exclusion0694 |
| 660.35000 | exclusion0695 |
| 660.66000 | exclusion0696 |
| 661.30000 | exclusion0697 |
| 661.81000 | exclusion0698 |
| 662.32000 | exclusion0699 |
| 662.83000 | exclusion0700 |
| 663.32000 | exclusion0701 |
| 663.75000 | exclusion0702 |
| 664.11000 | exclusion0703 |
| 664.37000 | exclusion0704 |
| 664.66000 | exclusion0705 |
| 665.32000 | exclusion0706 |
| 665.34000 | exclusion0707 |
| 665.64000 | exclusion0708 |

| | |
|---|---|
| 665.87000 | exclusion0709 |
| 666.34000 | exclusion0710 |
| 666.86000 | exclusion0711 |
| 667.03000 | exclusion0712 |
| 667.36000 | exclusion0713 |
| 667.52000 | exclusion0714 |
| 668.66000 | exclusion0715 |
| 668.83000 | exclusion0716 |
| 669.34000 | exclusion0717 |
| 669.51000 | exclusion0718 |
| 669.85000 | exclusion0719 |
| 669.90000 | exclusion0720 |
| 670.29000 | exclusion0721 |
| 670.80000 | exclusion0722 |
| 671.30000 | exclusion0126 |
| 671.53000 | exclusion0127 |
| 671.83000 | exclusion0128 |
| 671.83000 | exclusion0723 |
| 672.32000 | exclusion0724 |
| 672.82000 | exclusion0725 |
| 673.11000 | exclusion0726 |
| 673.33000 | exclusion0727 |
| 673.34000 | exclusion0728 |
| 674.31000 | exclusion0729 |
| 674.33000 | exclusion0730 |
| 674.58000 | exclusion0731 |
| 674.83000 | exclusion0732 |
| 675.19000 | exclusion0733 |
| 676.05000 | exclusion0734 |
| 676.85000 | exclusion0735 |
| 677.00000 | exclusion0736 |
| 677.28000 | exclusion0737 |
| 677.34000 | exclusion0738 |
| 677.85000 | exclusion0739 |
| 677.88000 | exclusion0740 |
| 678.36000 | exclusion0741 |
| 679.04000 | exclusion0742 |
| 679.31000 | exclusion0743 |
| 679.85000 | exclusion0744 |
| 680.33000 | exclusion0745 |
| 680.78000 | exclusion0746 |
| 681.32000 | exclusion0747 |
| 681.83000 | exclusion0129 |
| 681.84000 | exclusion0130 |
| 682.37000 | exclusion0748 |
| 682.67000 | exclusion0749 |
| 682.82000 | exclusion0750 |
| 682.85000 | exclusion0131 |
| 683.32000 | exclusion0751 |
| 683.37000 | exclusion0752 |
| 683.97000 | exclusion0753 |

| | |
|---|---|
| 684.29000 | exclusion0754 |
| 684.78000 | exclusion0755 |
| 685.67000 | exclusion0756 |
| 685.83000 | exclusion0757 |
| 686.31000 | exclusion0758 |
| 686.34000 | exclusion0759 |
| 686.85000 | exclusion0760 |
| 687.30000 | exclusion0761 |
| 687.33000 | exclusion0762 |
| 687.68000 | exclusion0763 |
| 688.27000 | exclusion0132 |
| 688.78000 | exclusion0764 |
| 689.00000 | exclusion0765 |
| 689.34000 | exclusion0766 |
| 689.37000 | exclusion0767 |
| 689.67000 | exclusion0768 |
| 690.35000 | exclusion0769 |
| 690.72000 | exclusion0770 |
| 691.32000 | exclusion0771 |
| 691.36000 | exclusion0772 |
| 692.02000 | exclusion0773 |
| 692.26000 | exclusion0774 |
| 692.70000 | exclusion0775 |
| 693.10000 | exclusion0776 |
| 693.56000 | exclusion0777 |
| 693.88000 | exclusion0778 |
| 694.31000 | exclusion0779 |
| 694.81000 | exclusion0133 |
| 695.33000 | exclusion0780 |
| 695.39000 | exclusion0781 |
| 695.85000 | exclusion0782 |
| 696.33000 | exclusion0783 |
| 696.35000 | exclusion0784 |
| 697.01000 | exclusion0785 |
| 697.36000 | exclusion0786 |
| 697.83000 | exclusion0787 |
| 698.37000 | exclusion0788 |
| 698.89000 | exclusion0789 |
| 699.12000 | exclusion0790 |
| 699.37000 | exclusion0791 |
| 699.38000 | exclusion0792 |
| 699.68000 | exclusion0793 |
| 700.37000 | exclusion0794 |
| 700.90000 | exclusion0795 |
| 701.33000 | exclusion0796 |
| 702.31000 | exclusion0797 |
| 702.33000 | exclusion0798 |
| 702.77000 | exclusion0799 |
| 702.87000 | exclusion0800 |
| 703.38000 | exclusion0801 |
| 703.71000 | exclusion0802 |

| | |
|---|---|
| 704.31000 | exclusion0803 |
| 705.27000 | exclusion0804 |
| 705.37000 | exclusion0805 |
| 705.82000 | exclusion0806 |
| 706.81000 | exclusion0807 |
| 707.02000 | exclusion0808 |
| 707.32000 | exclusion0809 |
| 708.04000 | exclusion0810 |
| 708.33000 | exclusion0811 |
| 708.38000 | exclusion0812 |
| 708.85000 | exclusion0813 |
| 709.35000 | exclusion0814 |
| 709.35000 | exclusion0815 |
| 709.90000 | exclusion0816 |
| 710.33000 | exclusion0817 |
| 710.36000 | exclusion0818 |
| 710.81000 | exclusion0819 |
| 711.35000 | exclusion0134 |
| 711.69000 | exclusion0135 |
| 711.89000 | exclusion0136 |
| 712.05000 | exclusion0820 |
| 712.34000 | exclusion0821 |
| 712.35000 | exclusion0822 |
| 712.38000 | exclusion0137 |
| 712.39000 | exclusion0138 |
| 712.59000 | exclusion0823 |
| 712.70000 | exclusion0824 |
| 712.80000 | exclusion0825 |
| 713.09000 | exclusion0826 |
| 713.34000 | exclusion0827 |
| 713.68000 | exclusion0828 |
| 713.89000 | exclusion0829 |
| 714.35000 | exclusion0830 |
| 714.40000 | exclusion0831 |
| 715.01000 | exclusion0832 |
| 715.36000 | exclusion0833 |
| 715.84000 | exclusion0834 |
| 716.31000 | exclusion0835 |
| 716.87000 | exclusion0836 |
| 717.83000 | exclusion0837 |
| 718.31000 | exclusion0838 |
| 718.34000 | exclusion0839 |
| 718.85000 | exclusion0139 |
| 718.85000 | exclusion0140 |
| 719.35000 | exclusion0840 |
| 719.71000 | exclusion0841 |
| 719.81000 | exclusion0842 |
| 720.32000 | exclusion0843 |
| 720.90000 | exclusion0844 |
| 721.37000 | exclusion0845 |
| 721.41000 | exclusion0846 |

| | |
|---|---|
| 721.73000 | exclusion0847 |
| 721.79000 | exclusion0848 |
| 722.59000 | exclusion0849 |
| 722.87000 | exclusion0850 |
| 723.07000 | exclusion0851 |
| 723.33000 | exclusion0852 |
| 723.40000 | exclusion0853 |
| 723.93000 | exclusion0141 |
| 724.34000 | exclusion0142 |
| 724.37000 | exclusion0143 |
| 724.68000 | exclusion0144 |
| 724.88000 | exclusion0854 |
| 725.04000 | exclusion0855 |
| 725.37000 | exclusion0145 |
| 725.38000 | exclusion0856 |
| 725.82000 | exclusion0857 |
| 726.05000 | exclusion0858 |
| 726.72000 | exclusion0859 |
| 726.81000 | exclusion0860 |
| 727.37000 | exclusion0861 |
| 727.50000 | exclusion0862 |
| 728.02000 | exclusion0863 |
| 728.32000 | exclusion0864 |
| 728.88000 | exclusion0865 |
| 729.32000 | exclusion0866 |
| 729.86000 | exclusion0867 |
| 730.36000 | exclusion0868 |
| 730.36000 | exclusion0869 |
| 730.64000 | exclusion0870 |
| 730.87000 | exclusion0871 |
| 731.01000 | exclusion0872 |
| 731.88000 | exclusion0873 |
| 732.35000 | exclusion0874 |
| 732.84000 | exclusion0875 |
| 733.37000 | exclusion0876 |
| 733.86000 | exclusion0877 |
| 734.32000 | exclusion0878 |
| 734.36000 | exclusion0879 |
| 735.33000 | exclusion0880 |
| 735.90000 | exclusion0881 |
| 736.37000 | exclusion0882 |
| 736.38000 | exclusion0883 |
| 736.86000 | exclusion0884 |
| 737.36000 | exclusion0885 |
| 737.88000 | exclusion0886 |
| 738.04000 | exclusion0887 |
| 738.38000 | exclusion0888 |
| 739.35000 | exclusion0889 |
| 739.86000 | exclusion0890 |
| 740.28000 | exclusion0891 |
| 740.87000 | exclusion0892 |

| | |
|---|---|
| 741.37000 | exclusion0893 |
| 742.37000 | exclusion0894 |
| 742.83000 | exclusion0895 |
| 743.06000 | exclusion0896 |
| 743.38000 | exclusion0897 |
| 743.85000 | exclusion0898 |
| 744.25000 | exclusion0899 |
| 744.38000 | exclusion0900 |
| 744.86000 | exclusion0901 |
| 745.36000 | exclusion0146 |
| 745.90000 | exclusion0147 |
| 746.28000 | exclusion0902 |
| 746.72000 | exclusion0903 |
| 747.03000 | exclusion0904 |
| 747.73000 | exclusion0905 |
| 747.83000 | exclusion0906 |
| 748.07000 | exclusion0907 |
| 748.32000 | exclusion0908 |
| 748.85000 | exclusion0909 |
| 749.04000 | exclusion0910 |
| 749.38000 | exclusion0911 |
| 749.61000 | exclusion0912 |
| 750.36000 | exclusion0913 |
| 750.37000 | exclusion0914 |
| 750.87000 | exclusion0915 |
| 751.37000 | exclusion0916 |
| 751.38000 | exclusion0917 |
| 751.86000 | exclusion0918 |
| 752.11000 | exclusion0919 |
| 752.36000 | exclusion0920 |
| 753.10000 | exclusion0921 |
| 753.37000 | exclusion0922 |
| 753.38000 | exclusion0923 |
| 754.32000 | exclusion0924 |
| 754.83000 | exclusion0925 |
| 755.34000 | exclusion0926 |
| 755.73000 | exclusion0927 |
| 755.86000 | exclusion0928 |
| 756.08000 | exclusion0929 |
| 756.32000 | exclusion0930 |
| 756.71000 | exclusion0931 |
| 757.36000 | exclusion0932 |
| 757.40000 | exclusion0933 |
| 757.70000 | exclusion0934 |
| 757.88000 | exclusion0935 |
| 758.84000 | exclusion0936 |
| 758.89000 | exclusion0937 |
| 759.37000 | exclusion0938 |
| 759.39000 | exclusion0939 |
| 759.68000 | exclusion0940 |
| 759.86000 | exclusion0941 |

| | |
|---|---|
| 760.36000 | exclusion0942 |
| 760.41000 | exclusion0943 |
| 760.73000 | exclusion0944 |
| 760.91000 | exclusion0945 |
| 761.34000 | exclusion0946 |
| 761.90000 | exclusion0947 |
| 762.40000 | exclusion0948 |
| 762.40000 | exclusion0949 |
| 762.88000 | exclusion0950 |
| 763.34000 | exclusion0951 |
| 763.90000 | exclusion0952 |
| 764.21000 | exclusion0953 |
| 764.36000 | exclusion0954 |
| 764.73000 | exclusion0955 |
| 765.34000 | exclusion0148 |
| 765.36000 | exclusion0149 |
| 765.86000 | exclusion0956 |
| 766.05000 | exclusion0957 |
| 766.35000 | exclusion0958 |
| 766.36000 | exclusion0150 |
| 766.39000 | exclusion0959 |
| 767.16000 | exclusion0151 |
| 767.37000 | exclusion0152 |
| 767.37000 | exclusion0153 |
| 767.86000 | exclusion0154 |
| 767.87000 | exclusion0960 |
| 768.35000 | exclusion0961 |
| 768.66000 | exclusion0962 |
| 768.90000 | exclusion0963 |
| 769.31000 | exclusion0964 |
| 769.90000 | exclusion0965 |
| 770.37000 | exclusion0966 |
| 770.86000 | exclusion0967 |
| 771.02000 | exclusion0968 |
| 771.38000 | exclusion0969 |
| 771.88000 | exclusion0970 |
| 772.39000 | exclusion0155 |
| 772.39000 | exclusion0156 |
| 772.85000 | exclusion0157 |
| 773.36000 | exclusion0971 |
| 773.37000 | exclusion0972 |
| 773.38000 | exclusion0158 |
| 773.87000 | exclusion0973 |
| 774.30000 | exclusion0974 |
| 774.61000 | exclusion0975 |
| 775.34000 | exclusion0976 |
| 775.39000 | exclusion0977 |
| 775.83000 | exclusion0978 |
| 776.37000 | exclusion0979 |
| 776.58000 | exclusion0980 |
| 776.86000 | exclusion0981 |

| | |
|---|---|
| 777.39000 | exclusion0982 |
| 777.40000 | exclusion0983 |
| 777.83000 | exclusion0984 |
| 778.40000 | exclusion0985 |
| 778.91000 | exclusion0986 |
| 779.11000 | exclusion0987 |
| 779.34000 | exclusion0988 |
| 779.37000 | exclusion0989 |
| 779.86000 | exclusion0990 |
| 780.75000 | exclusion0991 |
| 780.81000 | exclusion0992 |
| 781.08000 | exclusion0993 |
| 781.16000 | exclusion0994 |
| 781.39000 | exclusion0995 |
| 782.28000 | exclusion0996 |
| 782.79000 | exclusion0159 |
| 783.30000 | exclusion0997 |
| 783.41000 | exclusion0998 |
| 784.35000 | exclusion0999 |
| 784.87000 | exclusion1000 |
| 785.30000 | exclusion1001 |
| 785.65000 | exclusion1002 |
| 786.39000 | exclusion1003 |
| 786.88000 | exclusion1004 |
| 787.10000 | exclusion1005 |
| 787.36000 | exclusion1006 |
| 787.90000 | exclusion1007 |
| 788.71000 | exclusion1008 |
| 789.04000 | exclusion1009 |
| 789.37000 | exclusion1010 |
| 790.36000 | exclusion1011 |
| 790.38000 | exclusion1012 |
| 791.06000 | exclusion1013 |
| 791.37000 | exclusion1014 |
| 792.29000 | exclusion1015 |
| 792.38000 | exclusion1016 |
| 792.91000 | exclusion1017 |
| 792.91000 | exclusion1018 |
| 793.38000 | exclusion1019 |
| 793.87000 | exclusion1020 |
| 793.94000 | exclusion1021 |
| 794.38000 | exclusion1022 |
| 794.40000 | exclusion1023 |
| 795.36000 | exclusion1024 |
| 795.42000 | exclusion1025 |
| 795.90000 | exclusion1026 |
| 796.12000 | exclusion1027 |
| 796.37000 | exclusion1028 |
| 796.40000 | exclusion1029 |
| 796.88000 | exclusion1030 |
| 797.43000 | exclusion1031 |

| | |
|---|---|
| 797.87000 | exclusion1032 |
| 798.35000 | exclusion1033 |
| 798.41000 | exclusion1034 |
| 799.13000 | exclusion1035 |
| 799.87000 | exclusion1036 |
| 800.35000 | exclusion1037 |
| 800.38000 | exclusion1038 |
| 800.68000 | exclusion1039 |
| 801.38000 | exclusion1040 |
| 801.38000 | exclusion1041 |
| 801.95000 | exclusion1042 |
| 802.27000 | exclusion1043 |
| 802.39000 | exclusion1044 |
| 802.69000 | exclusion1045 |
| 802.82000 | exclusion1046 |
| 803.37000 | exclusion0160 |
| 803.92000 | exclusion1047 |
| 804.40000 | exclusion1048 |
| 804.41000 | exclusion1049 |
| 804.74000 | exclusion1050 |
| 805.00000 | exclusion1051 |
| 805.40000 | exclusion1052 |
| 806.39000 | exclusion1053 |
| 806.43000 | exclusion1054 |
| 806.88000 | exclusion1055 |
| 807.02000 | exclusion1056 |
| 808.10000 | exclusion1057 |
| 808.36000 | exclusion1058 |
| 808.36000 | exclusion1059 |
| 808.94000 | exclusion1060 |
| 809.40000 | exclusion1061 |
| 809.91000 | exclusion1062 |
| 810.39000 | exclusion1063 |
| 810.39000 | exclusion1064 |
| 810.75000 | exclusion1065 |
| 811.36000 | exclusion1066 |
| 812.36000 | exclusion0161 |
| 812.38000 | exclusion1067 |
| 812.73000 | exclusion1068 |
| 812.86000 | exclusion0162 |
| 813.37000 | exclusion1069 |
| 813.43000 | exclusion1070 |
| 814.36000 | exclusion0163 |
| 814.41000 | exclusion0164 |
| 814.74000 | exclusion0165 |
| 815.36000 | exclusion1071 |
| 815.41000 | exclusion1072 |
| 815.93000 | exclusion1073 |
| 816.44000 | exclusion0166 |
| 817.05000 | exclusion1074 |
| 817.33000 | exclusion1075 |

| | |
|---|---|
| 817.75000 | exclusion1076 |
| 817.86000 | exclusion1077 |
| 818.43000 | exclusion1078 |
| 818.55000 | exclusion1079 |
| 818.87000 | exclusion1080 |
| 819.04000 | exclusion1081 |
| 819.42000 | exclusion1082 |
| 820.37000 | exclusion1083 |
| 821.15000 | exclusion1084 |
| 821.39000 | exclusion1085 |
| 822.38000 | exclusion1086 |
| 822.39000 | exclusion1087 |
| 822.91000 | exclusion1088 |
| 822.92000 | exclusion1089 |
| 823.57000 | exclusion1090 |
| 824.40000 | exclusion1091 |
| 824.81000 | exclusion1092 |
| 825.40000 | exclusion1093 |
| 825.85000 | exclusion1094 |
| 826.22000 | exclusion1095 |
| 826.38000 | exclusion1096 |
| 826.44000 | exclusion1097 |
| 827.42000 | exclusion1098 |
| 827.42000 | exclusion1099 |
| 827.89000 | exclusion1100 |
| 828.41000 | exclusion1101 |
| 828.44000 | exclusion1102 |
| 829.41000 | exclusion1103 |
| 829.43000 | exclusion1104 |
| 830.43000 | exclusion1105 |
| 830.92000 | exclusion1106 |
| 830.96000 | exclusion1107 |
| 831.13000 | exclusion1108 |
| 831.32000 | exclusion1109 |
| 831.94000 | exclusion1110 |
| 832.41000 | exclusion1111 |
| 832.44000 | exclusion1112 |
| 832.88000 | exclusion1113 |
| 833.15000 | exclusion1114 |
| 833.40000 | exclusion1115 |
| 833.78000 | exclusion1116 |
| 834.16000 | exclusion1117 |
| 834.33000 | exclusion1118 |
| 834.43000 | exclusion1119 |
| 834.92000 | exclusion1120 |
| 835.04000 | exclusion1121 |
| 835.13000 | exclusion1122 |
| 835.72000 | exclusion0167 |
| 836.06000 | exclusion0168 |
| 836.40000 | exclusion0169 |
| 836.41000 | exclusion1123 |

| | |
|---|---|
| 836.42000 | exclusion1124 |
| 837.42000 | exclusion1125 |
| 837.44000 | exclusion1126 |
| 837.76000 | exclusion1127 |
| 837.93000 | exclusion1128 |
| 838.42000 | exclusion1129 |
| 839.16000 | exclusion1130 |
| 839.25000 | exclusion1131 |
| 839.43000 | exclusion1132 |
| 839.91000 | exclusion1133 |
| 840.42000 | exclusion1134 |
| 840.67000 | exclusion1135 |
| 840.73000 | exclusion1136 |
| 841.44000 | exclusion1137 |
| 841.45000 | exclusion1138 |
| 842.11000 | exclusion0170 |
| 842.14000 | exclusion0171 |
| 842.38000 | exclusion0172 |
| 842.39000 | exclusion0173 |
| 842.91000 | exclusion1139 |
| 842.92000 | exclusion1140 |
| 843.40000 | exclusion1141 |
| 843.64000 | exclusion1142 |
| 843.91000 | exclusion1143 |
| 843.94000 | exclusion1144 |
| 844.44000 | exclusion0174 |
| 844.58000 | exclusion0175 |
| 844.89000 | exclusion0176 |
| 845.37000 | exclusion0177 |
| 845.41000 | exclusion0178 |
| 845.41000 | exclusion1145 |
| 845.87000 | exclusion1146 |
| 846.16000 | exclusion1147 |
| 846.36000 | exclusion1148 |
| 847.37000 | exclusion1149 |
| 847.39000 | exclusion1150 |
| 847.88000 | exclusion1151 |
| 848.36000 | exclusion1152 |
| 848.91000 | exclusion0179 |
| 849.37000 | exclusion0180 |
| 849.41000 | exclusion1153 |
| 849.44000 | exclusion1154 |
| 849.89000 | exclusion1155 |
| 849.90000 | exclusion1156 |
| 850.42000 | exclusion1157 |
| 850.91000 | exclusion1158 |
| 851.17000 | exclusion1159 |
| 851.42000 | exclusion1160 |
| 851.45000 | exclusion1161 |
| 851.78000 | exclusion1162 |
| 852.38000 | exclusion1163 |

| | |
|---|---|
| 852.39000 | exclusion1164 |
| 852.88000 | exclusion1165 |
| 853.23000 | exclusion1166 |
| 853.35000 | exclusion1167 |
| 853.66000 | exclusion1168 |
| 854.38000 | exclusion1169 |
| 855.37000 | exclusion1170 |
| 855.40000 | exclusion1171 |
| 855.88000 | exclusion1172 |
| 856.15000 | exclusion1173 |
| 856.40000 | exclusion1174 |
| 857.06000 | exclusion1175 |
| 857.45000 | exclusion1176 |
| 857.89000 | exclusion1177 |
| 858.50000 | exclusion1178 |
| 858.74000 | exclusion1179 |
| 859.47000 | exclusion1180 |
| 859.64000 | exclusion1181 |
| 860.08000 | exclusion1182 |
| 860.39000 | exclusion1183 |
| 860.88000 | exclusion1184 |
| 861.44000 | exclusion1185 |
| 861.90000 | exclusion1186 |
| 862.46000 | exclusion1187 |
| 862.49000 | exclusion1188 |
| 862.90000 | exclusion1189 |
| 863.39000 | exclusion1190 |
| 863.41000 | exclusion1191 |
| 863.92000 | exclusion0181 |
| 864.42000 | exclusion0182 |
| 864.44000 | exclusion1192 |
| 864.45000 | exclusion1193 |
| 864.88000 | exclusion1194 |
| 865.40000 | exclusion0183 |
| 865.91000 | exclusion1195 |
| 866.41000 | exclusion0184 |
| 866.41000 | exclusion1196 |
| 866.42000 | exclusion1197 |
| 867.43000 | exclusion0185 |
| 867.45000 | exclusion1198 |
| 867.75000 | exclusion1199 |
| 868.13000 | exclusion1200 |
| 868.40000 | exclusion1201 |
| 868.40000 | exclusion1202 |
| 868.90000 | exclusion1203 |
| 868.94000 | exclusion1204 |
| 869.22000 | exclusion1205 |
| 869.73000 | exclusion1206 |
| 870.18000 | exclusion1207 |
| 870.42000 | exclusion1208 |
| 870.43000 | exclusion1209 |

| | |
|---|---|
| 871.35000 | exclusion1210 |
| 872.38000 | exclusion1211 |
| 872.43000 | exclusion1212 |
| 872.70000 | exclusion1213 |
| 872.92000 | exclusion1214 |
| 873.45000 | exclusion1215 |
| 874.05000 | exclusion1216 |
| 874.42000 | exclusion1217 |
| 874.95000 | exclusion1218 |
| 875.39000 | exclusion1219 |
| 875.90000 | exclusion1220 |
| 876.01000 | exclusion1221 |
| 876.40000 | exclusion1222 |
| 876.40000 | exclusion1223 |
| 877.36000 | exclusion1224 |
| 877.40000 | exclusion1225 |
| 878.05000 | exclusion1226 |
| 878.48000 | exclusion1227 |
| 879.05000 | exclusion1228 |
| 879.41000 | exclusion1229 |
| 879.93000 | exclusion1230 |
| 880.40000 | exclusion1231 |
| 880.41000 | exclusion1232 |
| 880.73000 | exclusion1233 |
| 881.43000 | exclusion1234 |
| 881.94000 | exclusion0186 |
| 882.44000 | exclusion1235 |
| 882.46000 | exclusion1236 |
| 882.91000 | exclusion1237 |
| 883.44000 | exclusion1238 |
| 884.41000 | exclusion1239 |
| 884.47000 | exclusion1240 |
| 884.89000 | exclusion1241 |
| 885.42000 | exclusion1242 |
| 885.45000 | exclusion1243 |
| 885.93000 | exclusion1244 |
| 886.40000 | exclusion1245 |
| 886.95000 | exclusion1246 |
| 887.48000 | exclusion1247 |
| 887.96000 | exclusion1248 |
| 888.40000 | exclusion1249 |
| 888.42000 | exclusion1250 |
| 888.74000 | exclusion1251 |
| 889.47000 | exclusion1252 |
| 890.41000 | exclusion1253 |
| 890.66000 | exclusion1254 |
| 890.88000 | exclusion1255 |
| 891.41000 | exclusion1256 |
| 891.90000 | exclusion1257 |
| 892.24000 | exclusion1258 |
| 892.45000 | exclusion1259 |

| | |
|---|---|
| 893.44000 | exclusion1260 |
| 893.89000 | exclusion1261 |
| 894.18000 | exclusion1262 |
| 894.44000 | exclusion1263 |
| 895.42000 | exclusion1264 |
| 895.48000 | exclusion1265 |
| 895.89000 | exclusion1266 |
| 896.16000 | exclusion1267 |
| 896.43000 | exclusion1268 |
| 896.45000 | exclusion1269 |
| 897.41000 | exclusion1270 |
| 898.10000 | exclusion1271 |
| 898.42000 | exclusion1272 |
| 898.82000 | exclusion1273 |
| 899.44000 | exclusion1274 |
| 899.45000 | exclusion1275 |
| 900.42000 | exclusion1276 |
| 900.43000 | exclusion1277 |
| 901.14000 | exclusion1278 |
| 901.45000 | exclusion1279 |
| 902.42000 | exclusion1280 |
| 902.70000 | exclusion1281 |
| 903.13000 | exclusion1282 |
| 903.41000 | exclusion1283 |
| 904.43000 | exclusion0187 |
| 904.44000 | exclusion0188 |
| 904.79000 | exclusion1284 |
| 905.13000 | exclusion1285 |
| 905.40000 | exclusion1286 |
| 905.96000 | exclusion1287 |
| 906.41000 | exclusion1288 |
| 907.42000 | exclusion1289 |
| 907.45000 | exclusion1290 |
| 907.96000 | exclusion1291 |
| 908.40000 | exclusion1292 |
| 908.93000 | exclusion0189 |
| 909.43000 | exclusion0190 |
| 909.44000 | exclusion1293 |
| 909.44000 | exclusion1294 |
| 909.94000 | exclusion1295 |
| 910.44000 | exclusion0191 |
| 910.44000 | exclusion0192 |
| 911.13000 | exclusion1296 |
| 911.46000 | exclusion1297 |
| 911.69000 | exclusion1298 |
| 911.94000 | exclusion1299 |
| 911.97000 | exclusion1300 |
| 912.41000 | exclusion1301 |
| 912.91000 | exclusion0193 |
| 912.99000 | exclusion0194 |
| 913.45000 | exclusion0195 |

| | |
|---|---|
| 913.66000 | exclusion1302 |
| 913.70000 | exclusion1303 |
| 913.92000 | exclusion1304 |
| 913.93000 | exclusion0196 |
| 914.43000 | exclusion1305 |
| 914.47000 | exclusion1306 |
| 915.12000 | exclusion1307 |
| 915.45000 | exclusion1308 |
| 915.95000 | exclusion1309 |
| 916.46000 | exclusion1310 |
| 916.49000 | exclusion1311 |
| 916.75000 | exclusion1312 |
| 917.08000 | exclusion1313 |
| 917.47000 | exclusion1314 |
| 917.93000 | exclusion1315 |
| 918.11000 | exclusion1316 |
| 918.46000 | exclusion1317 |
| 918.48000 | exclusion1318 |
| 918.92000 | exclusion1319 |
| 919.43000 | exclusion1320 |
| 919.54000 | exclusion1321 |
| 920.42000 | exclusion1322 |
| 920.49000 | exclusion1323 |
| 920.87000 | exclusion1324 |
| 921.40000 | exclusion1325 |
| 921.47000 | exclusion1326 |
| 922.30000 | exclusion1327 |
| 922.47000 | exclusion1328 |
| 922.90000 | exclusion1329 |
| 923.46000 | exclusion1330 |
| 923.91000 | exclusion1331 |
| 924.13000 | exclusion1332 |
| 924.43000 | exclusion1333 |
| 924.94000 | exclusion1334 |
| 925.49000 | exclusion1335 |
| 926.46000 | exclusion1336 |
| 926.79000 | exclusion1337 |
| 927.48000 | exclusion0197 |
| 927.82000 | exclusion1338 |
| 927.88000 | exclusion1339 |
| 928.47000 | exclusion1340 |
| 928.93000 | exclusion1341 |
| 929.45000 | exclusion1342 |
| 929.77000 | exclusion1343 |
| 930.46000 | exclusion1344 |
| 930.47000 | exclusion1345 |
| 930.68000 | exclusion1346 |
| 931.11000 | exclusion1347 |
| 931.40000 | exclusion1348 |
| 931.41000 | exclusion1349 |
| 931.81000 | exclusion1350 |

| | |
|---|---|
| 932.43000 | exclusion1351 |
| 932.94000 | exclusion0198 |
| 933.14000 | exclusion0199 |
| 933.47000 | exclusion0200 |
| 933.47000 | exclusion1352 |
| 933.79000 | exclusion1353 |
| 934.39000 | exclusion1354 |
| 934.97000 | exclusion1355 |
| 935.16000 | exclusion1356 |
| 935.47000 | exclusion1357 |
| 935.91000 | exclusion1358 |
| 936.11000 | exclusion1359 |
| 937.12000 | exclusion0201 |
| 937.46000 | exclusion0202 |
| 937.49000 | exclusion1360 |
| 937.52000 | exclusion1361 |
| 937.79000 | exclusion1362 |
| 937.99000 | exclusion1363 |
| 938.49000 | exclusion1364 |
| 939.21000 | exclusion1365 |
| 939.49000 | exclusion1366 |
| 939.93000 | exclusion1367 |
| 940.44000 | exclusion0203 |
| 940.48000 | exclusion0204 |
| 941.09000 | exclusion0205 |
| 941.46000 | exclusion1368 |
| 942.47000 | exclusion1369 |
| 942.48000 | exclusion0206 |
| 942.98000 | exclusion1370 |
| 943.10000 | exclusion1371 |
| 943.47000 | exclusion1372 |
| 943.73000 | exclusion1373 |
| 944.47000 | exclusion1374 |
| 944.74000 | exclusion1375 |
| 944.92000 | exclusion1376 |
| 945.48000 | exclusion1377 |
| 945.78000 | exclusion1378 |
| 945.98000 | exclusion1379 |
| 946.95000 | exclusion1380 |
| 947.49000 | exclusion1381 |
| 947.96000 | exclusion1382 |
| 948.46000 | exclusion1383 |
| 949.02000 | exclusion1384 |
| 949.46000 | exclusion1385 |
| 949.80000 | exclusion1386 |
| 950.01000 | exclusion1387 |
| 950.70000 | exclusion1388 |
| 951.13000 | exclusion1389 |
| 951.41000 | exclusion1390 |
| 951.47000 | exclusion1391 |
| 951.82000 | exclusion1392 |

| | |
|---|---|
| 952.14000 | exclusion1393 |
| 952.77000 | exclusion1394 |
| 952.96000 | exclusion1395 |
| 953.01000 | exclusion1396 |
| 953.48000 | exclusion1397 |
| 954.44000 | exclusion1398 |
| 954.94000 | exclusion1399 |
| 955.35000 | exclusion1400 |
| 955.57000 | exclusion1401 |
| 956.47000 | exclusion1402 |
| 957.43000 | exclusion1403 |
| 957.74000 | exclusion1404 |
| 958.44000 | exclusion1405 |
| 958.81000 | exclusion1406 |
| 959.41000 | exclusion1407 |
| 960.41000 | exclusion0207 |
| 960.45000 | exclusion1408 |
| 960.75000 | exclusion1409 |
| 961.00000 | exclusion1410 |
| 961.35000 | exclusion1411 |
| 961.98000 | exclusion0208 |
| 962.49000 | exclusion1412 |
| 962.87000 | exclusion1413 |
| 963.48000 | exclusion1414 |
| 963.94000 | exclusion1415 |
| 964.44000 | exclusion1416 |
| 964.74000 | exclusion1417 |
| 964.98000 | exclusion1418 |
| 965.02000 | exclusion1419 |
| 965.48000 | exclusion1420 |
| 966.26000 | exclusion1421 |
| 966.48000 | exclusion1422 |
| 967.21000 | exclusion1423 |
| 967.48000 | exclusion1424 |
| 967.48000 | exclusion1425 |
| 967.90000 | exclusion1426 |
| 968.47000 | exclusion1427 |
| 968.97000 | exclusion0209 |
| 969.50000 | exclusion1428 |
| 969.98000 | exclusion1429 |
| 970.11000 | exclusion1430 |
| 970.45000 | exclusion1431 |
| 970.52000 | exclusion1432 |
| 971.48000 | exclusion1433 |
| 971.96000 | exclusion1434 |
| 972.15000 | exclusion1435 |
| 972.44000 | exclusion1436 |
| 972.48000 | exclusion1437 |
| 972.54000 | exclusion1438 |
| 973.00000 | exclusion1439 |
| 973.80000 | exclusion1440 |

| | |
|---|---|
| 974.47000 | exclusion1441 |
| 975.47000 | exclusion1442 |
| 975.49000 | exclusion1443 |
| 975.82000 | exclusion1444 |
| 976.68000 | exclusion1445 |
| 977.14000 | exclusion1446 |
| 977.49000 | exclusion1447 |
| 977.78000 | exclusion1448 |
| 978.46000 | exclusion1449 |
| 978.51000 | exclusion1450 |
| 979.42000 | exclusion1451 |
| 980.13000 | exclusion1452 |
| 980.51000 | exclusion1453 |
| 980.97000 | exclusion1454 |
| 981.21000 | exclusion1455 |
| 981.30000 | exclusion1456 |
| 981.54000 | exclusion1457 |
| 982.46000 | exclusion1458 |
| 983.45000 | exclusion1459 |
| 983.49000 | exclusion1460 |
| 984.48000 | exclusion1461 |
| 984.50000 | exclusion1462 |
| 984.94000 | exclusion1463 |
| 985.87000 | exclusion1464 |
| 986.00000 | exclusion1465 |
| 986.47000 | exclusion1466 |
| 986.83000 | exclusion1467 |
| 987.01000 | exclusion1468 |
| 987.51000 | exclusion1469 |
| 987.97000 | exclusion1470 |
| 988.18000 | exclusion1471 |
| 988.49000 | exclusion1472 |
| 989.01000 | exclusion1473 |
| 989.52000 | exclusion0210 |
| 990.02000 | exclusion1474 |
| 990.14000 | exclusion1475 |
| 990.47000 | exclusion1476 |
| 990.50000 | exclusion1477 |
| 991.16000 | exclusion1478 |
| 991.20000 | exclusion1479 |
| 991.48000 | exclusion1480 |
| 991.83000 | exclusion1481 |
| 991.99000 | exclusion1482 |
| 992.50000 | exclusion0211 |
| 993.02000 | exclusion0212 |
| 993.50000 | exclusion0213 |
| 993.52000 | exclusion1483 |
| 993.96000 | exclusion1484 |
| 994.47000 | exclusion0214 |
| 994.98000 | exclusion0215 |
| 995.46000 | exclusion0216 |

| | |
|---|---|
| 995.49000 | exclusion1485 |
| 996.48000 | exclusion1486 |
| 996.98000 | exclusion0217 |
| 997.50000 | exclusion1487 |
| 998.48000 | exclusion1488 |
| 998.81000 | exclusion1489 |
| 999.39000 | exclusion1490 |
| 999.82000 | exclusion1491 |
| 1000.46000 | exclusion1492 |
| 1000.84000 | exclusion1493 |
| 1001.50000 | exclusion1494 |
| 1002.48000 | exclusion1495 |
| 1003.01000 | exclusion1496 |
| 1003.53000 | exclusion1497 |
| 1004.48000 | exclusion1498 |
| 1004.94000 | exclusion1499 |
| 1005.45000 | exclusion1500 |
| 1005.98000 | exclusion0218 |
| 1006.52000 | exclusion1501 |
| 1006.98000 | exclusion1502 |
| 1007.95000 | exclusion1503 |
| 1008.80000 | exclusion1504 |
| 1009.06000 | exclusion1505 |
| 1009.92000 | exclusion1506 |
| 1010.38000 | exclusion1507 |
| 1011.02000 | exclusion1508 |
| 1011.77000 | exclusion1509 |
| 1012.48000 | exclusion1510 |
| 1012.94000 | exclusion1511 |
| 1013.49000 | exclusion0219 |
| 1013.79000 | exclusion0220 |
| 1014.47000 | exclusion0221 |
| 1014.47000 | exclusion1512 |
| 1015.42000 | exclusion1513 |
| 1016.01000 | exclusion1514 |
| 1016.49000 | exclusion1515 |
| 1017.02000 | exclusion1516 |
| 1017.53000 | exclusion1517 |
| 1017.91000 | exclusion1518 |
| 1018.41000 | exclusion0222 |
| 1018.95000 | exclusion1519 |
| 1019.21000 | exclusion1520 |
| 1020.05000 | exclusion1521 |
| 1020.49000 | exclusion1522 |
| 1021.00000 | exclusion1523 |
| 1021.53000 | exclusion1524 |
| 1022.46000 | exclusion1525 |
| 1022.82000 | exclusion1526 |
| 1023.45000 | exclusion1527 |
| 1024.18000 | exclusion1528 |
| 1024.51000 | exclusion1529 |

| | |
|---|---|
| 1025.42000 | exclusion1530 |
| 1025.86000 | exclusion1531 |
| 1026.79000 | exclusion1532 |
| 1027.46000 | exclusion1533 |
| 1028.02000 | exclusion1534 |
| 1028.98000 | exclusion1535 |
| 1029.51000 | exclusion0223 |
| 1030.46000 | exclusion1536 |
| 1030.53000 | exclusion0224 |
| 1031.47000 | exclusion1537 |
| 1032.46000 | exclusion1538 |
| 1032.86000 | exclusion1539 |
| 1033.15000 | exclusion1540 |
| 1033.52000 | exclusion1541 |
| 1034.02000 | exclusion1542 |
| 1034.80000 | exclusion0225 |
| 1035.54000 | exclusion0226 |
| 1035.55000 | exclusion1543 |
| 1036.20000 | exclusion1544 |
| 1036.53000 | exclusion1545 |
| 1036.77000 | exclusion1546 |
| 1037.45000 | exclusion1547 |
| 1038.03000 | exclusion0227 |
| 1038.26000 | exclusion0228 |
| 1038.46000 | exclusion0229 |
| 1038.98000 | exclusion1548 |
| 1039.29000 | exclusion1549 |
| 1039.50000 | exclusion1550 |
| 1040.04000 | exclusion1551 |
| 1040.51000 | exclusion1552 |
| 1041.47000 | exclusion1553 |
| 1041.77000 | exclusion1554 |
| 1042.02000 | exclusion1555 |
| 1042.52000 | exclusion1556 |
| 1043.14000 | exclusion1557 |
| 1044.17000 | exclusion1558 |
| 1044.52000 | exclusion1559 |
| 1044.97000 | exclusion1560 |
| 1045.46000 | exclusion1561 |
| 1046.50000 | exclusion0230 |
| 1046.50000 | exclusion1562 |
| 1047.50000 | exclusion0231 |
| 1047.82000 | exclusion1563 |
| 1048.05000 | exclusion1564 |
| 1048.84000 | exclusion1565 |
| 1049.47000 | exclusion1566 |
| 1049.87000 | exclusion1567 |
| 1050.51000 | exclusion1568 |
| 1050.77000 | exclusion1569 |
| 1051.47000 | exclusion1570 |
| 1052.00000 | exclusion0232 |

| | |
|---|---|
| 1052.53000 | exclusion1571 |
| 1053.06000 | exclusion1572 |
| 1053.53000 | exclusion1573 |
| 1053.84000 | exclusion1574 |
| 1054.50000 | exclusion1575 |
| 1055.04000 | exclusion1576 |
| 1055.48000 | exclusion1577 |
| 1056.04000 | exclusion1578 |
| 1056.52000 | exclusion1579 |
| 1057.55000 | exclusion0233 |
| 1057.56000 | exclusion1580 |
| 1057.84000 | exclusion1581 |
| 1058.07000 | exclusion1582 |
| 1058.48000 | exclusion1583 |
| 1059.27000 | exclusion1584 |
| 1059.48000 | exclusion1585 |
| 1060.03000 | exclusion1586 |
| 1060.53000 | exclusion1587 |
| 1061.02000 | exclusion1588 |
| 1061.83000 | exclusion1589 |
| 1062.24000 | exclusion1590 |
| 1062.54000 | exclusion1591 |
| 1063.01000 | exclusion1592 |
| 1063.48000 | exclusion1593 |
| 1064.48000 | exclusion1594 |
| 1065.47000 | exclusion1595 |
| 1066.11000 | exclusion0234 |
| 1066.54000 | exclusion0235 |
| 1067.04000 | exclusion1596 |
| 1067.49000 | exclusion1597 |
| 1067.53000 | exclusion0236 |
| 1068.04000 | exclusion1598 |
| 1068.47000 | exclusion1599 |
| 1069.03000 | exclusion0237 |
| 1069.45000 | exclusion0238 |
| 1070.06000 | exclusion1600 |
| 1070.52000 | exclusion1601 |
| 1071.15000 | exclusion1602 |
| 1071.54000 | exclusion1603 |
| 1072.07000 | exclusion0239 |
| 1072.54000 | exclusion0240 |
| 1072.84000 | exclusion0241 |
| 1072.87000 | exclusion1604 |
| 1073.53000 | exclusion1605 |
| 1074.04000 | exclusion1606 |
| 1074.55000 | exclusion0242 |
| 1074.85000 | exclusion0243 |
| 1075.13000 | exclusion0244 |
| 1075.52000 | exclusion0245 |
| 1075.53000 | exclusion1607 |
| 1076.06000 | exclusion1608 |

| | |
|---|---|
| 1077.04000 | exclusion1609 |
| 1077.54000 | exclusion1610 |
| 1078.99000 | exclusion1611 |
| 1079.51000 | exclusion1612 |
| 1080.50000 | exclusion1613 |
| 1080.85000 | exclusion1614 |
| 1081.53000 | exclusion1615 |
| 1082.19000 | exclusion1616 |
| 1082.53000 | exclusion1617 |
| 1083.55000 | exclusion0246 |
| 1083.58000 | exclusion1618 |
| 1084.06000 | exclusion1619 |
| 1084.55000 | exclusion1620 |
| 1085.04000 | exclusion1621 |
| 1085.53000 | exclusion1622 |
| 1086.05000 | exclusion1623 |
| 1086.49000 | exclusion1624 |
| 1086.99000 | exclusion0247 |
| 1087.55000 | exclusion0248 |
| 1087.55000 | exclusion1625 |
| 1087.88000 | exclusion1626 |
| 1088.24000 | exclusion1627 |
| 1088.49000 | exclusion1628 |
| 1089.00000 | exclusion0249 |
| 1089.53000 | exclusion0250 |
| 1090.05000 | exclusion0251 |
| 1090.06000 | exclusion1629 |
| 1090.53000 | exclusion1630 |
| 1091.06000 | exclusion1631 |
| 1091.89000 | exclusion1632 |
| 1092.50000 | exclusion0252 |
| 1093.03000 | exclusion0253 |
| 1093.53000 | exclusion1633 |
| 1093.87000 | exclusion1634 |
| 1094.55000 | exclusion1635 |
| 1095.50000 | exclusion1636 |
| 1095.86000 | exclusion1637 |
| 1096.12000 | exclusion1638 |
| 1097.16000 | exclusion0254 |
| 1097.29000 | exclusion1639 |
| 1097.54000 | exclusion1640 |
| 1098.20000 | exclusion1641 |
| 1099.29000 | exclusion0255 |
| 1099.53000 | exclusion1642 |
| 1100.03000 | exclusion0256 |
| 1100.56000 | exclusion0257 |
| 1100.81000 | exclusion0258 |
| 1101.09000 | exclusion0259 |
| 1101.09000 | exclusion1643 |
| 1102.22000 | exclusion1644 |
| 1102.51000 | exclusion1645 |

| | |
|---|---|
| 1102.88000 | exclusion1646 |
| 1103.99000 | exclusion1647 |
| 1104.22000 | exclusion1648 |
| 1104.58000 | exclusion1649 |
| 1105.54000 | exclusion1650 |
| 1105.77000 | exclusion1651 |
| 1106.53000 | exclusion1652 |
| 1107.54000 | exclusion1653 |
| 1108.05000 | exclusion0260 |
| 1108.58000 | exclusion1654 |
| 1109.52000 | exclusion1655 |
| 1109.88000 | exclusion1656 |
| 1110.23000 | exclusion1657 |
| 1110.91000 | exclusion0261 |
| 1111.54000 | exclusion0262 |
| 1111.56000 | exclusion1658 |
| 1111.87000 | exclusion1659 |
| 1112.08000 | exclusion1660 |
| 1112.80000 | exclusion1661 |
| 1113.00000 | exclusion1662 |
| 1113.52000 | exclusion1663 |
| 1114.04000 | exclusion0263 |
| 1114.85000 | exclusion0264 |
| 1114.94000 | exclusion1664 |
| 1115.55000 | exclusion1665 |
| 1116.08000 | exclusion1666 |
| 1116.88000 | exclusion1667 |
| 1117.52000 | exclusion1668 |
| 1118.09000 | exclusion1669 |
| 1118.58000 | exclusion1670 |
| 1118.86000 | exclusion1671 |
| 1119.89000 | exclusion1672 |
| 1120.52000 | exclusion1673 |
| 1121.05000 | exclusion1674 |
| 1121.56000 | exclusion0265 |
| 1121.88000 | exclusion0266 |
| 1122.19000 | exclusion1675 |
| 1122.51000 | exclusion1676 |
| 1123.21000 | exclusion1677 |
| 1123.56000 | exclusion1678 |
| 1124.57000 | exclusion1679 |
| 1125.02000 | exclusion1680 |
| 1125.54000 | exclusion1681 |
| 1126.04000 | exclusion1682 |
| 1126.54000 | exclusion0267 |
| 1127.00000 | exclusion0268 |
| 1127.21000 | exclusion0269 |
| 1127.55000 | exclusion1683 |
| 1127.87000 | exclusion1684 |
| 1128.09000 | exclusion1685 |
| 1128.53000 | exclusion1686 |

| | |
|---|---|
| 1129.58000 | exclusion0270 |
| 1130.12000 | exclusion0271 |
| 1130.21000 | exclusion1687 |
| 1130.54000 | exclusion0272 |
| 1130.58000 | exclusion1688 |
| 1131.03000 | exclusion1689 |
| 1131.57000 | exclusion1690 |
| 1132.20000 | exclusion1691 |
| 1132.87000 | exclusion1692 |
| 1133.08000 | exclusion1693 |
| 1133.62000 | exclusion1694 |
| 1133.89000 | exclusion1695 |
| 1134.54000 | exclusion1696 |
| 1135.11000 | exclusion1697 |
| 1136.04000 | exclusion1698 |
| 1136.53000 | exclusion1699 |
| 1137.44000 | exclusion1700 |
| 1138.55000 | exclusion0273 |
| 1138.55000 | exclusion1701 |
| 1139.18000 | exclusion1702 |
| 1139.54000 | exclusion1703 |
| 1140.20000 | exclusion0274 |
| 1140.58000 | exclusion0275 |
| 1140.83000 | exclusion0276 |
| 1141.52000 | exclusion1704 |
| 1141.90000 | exclusion1705 |
| 1142.51000 | exclusion1706 |
| 1143.05000 | exclusion1707 |
| 1143.56000 | exclusion1708 |
| 1144.09000 | exclusion0277 |
| 1144.54000 | exclusion0278 |
| 1145.04000 | exclusion1709 |
| 1145.51000 | exclusion0279 |
| 1145.53000 | exclusion1710 |
| 1146.56000 | exclusion1711 |
| 1147.00000 | exclusion1712 |
| 1147.53000 | exclusion1713 |
| 1148.07000 | exclusion1714 |
| 1148.56000 | exclusion1715 |
| 1149.06000 | exclusion1716 |
| 1149.56000 | exclusion1717 |
| 1150.56000 | exclusion0280 |
| 1150.56000 | exclusion1718 |
| 1151.05000 | exclusion1719 |
| 1151.60000 | exclusion1720 |
| 1152.12000 | exclusion1721 |
| 1152.55000 | exclusion1722 |
| 1153.54000 | exclusion1723 |
| 1154.20000 | exclusion1724 |
| 1154.55000 | exclusion1725 |
| 1155.08000 | exclusion1726 |

| | |
|---|---|
| 1156.07000 | exclusion1727 |
| 1156.52000 | exclusion1728 |
| 1157.56000 | exclusion1729 |
| 1158.23000 | exclusion1730 |
| 1158.51000 | exclusion1731 |
| 1159.44000 | exclusion1732 |
| 1159.64000 | exclusion1733 |
| 1159.91000 | exclusion1734 |
| 1161.55000 | exclusion0281 |
| 1161.57000 | exclusion1735 |
| 1162.04000 | exclusion1736 |
| 1162.58000 | exclusion1737 |
| 1163.03000 | exclusion1738 |
| 1163.58000 | exclusion0282 |
| 1163.89000 | exclusion0283 |
| 1164.56000 | exclusion1739 |
| 1165.57000 | exclusion1740 |
| 1166.27000 | exclusion0284 |
| 1166.53000 | exclusion0285 |
| 1166.90000 | exclusion0286 |
| 1167.20000 | exclusion1741 |
| 1167.21000 | exclusion0287 |
| 1167.52000 | exclusion1742 |
| 1168.20000 | exclusion1743 |
| 1168.55000 | exclusion1744 |
| 1169.58000 | exclusion1745 |
| 1170.08000 | exclusion0288 |
| 1170.56000 | exclusion0289 |
| 1171.09000 | exclusion1746 |
| 1171.61000 | exclusion1747 |
| 1172.54000 | exclusion0290 |
| 1173.26000 | exclusion0291 |
| 1173.27000 | exclusion1748 |
| 1173.56000 | exclusion1749 |
| 1174.54000 | exclusion1750 |
| 1175.48000 | exclusion1751 |
| 1176.00000 | exclusion0292 |
| 1176.58000 | exclusion0293 |
| 1176.59000 | exclusion1752 |
| 1177.58000 | exclusion1753 |
| 1178.09000 | exclusion1754 |
| 1178.50000 | exclusion1755 |
| 1179.08000 | exclusion0294 |
| 1179.58000 | exclusion1756 |
| 1180.14000 | exclusion1757 |
| 1180.94000 | exclusion0295 |
| 1181.56000 | exclusion1758 |
| 1181.91000 | exclusion0296 |
| 1182.54000 | exclusion1759 |
| 1182.93000 | exclusion1760 |
| 1183.53000 | exclusion0297 |

| | |
|---|---|
| 1184.10000 | exclusion1761 |
| 1184.55000 | exclusion1762 |
| 1185.54000 | exclusion1763 |
| 1186.47000 | exclusion0298 |
| 1187.05000 | exclusion0299 |
| 1187.10000 | exclusion1764 |
| 1187.58000 | exclusion1765 |
| 1188.57000 | exclusion1766 |
| 1189.54000 | exclusion1767 |
| 1189.91000 | exclusion1768 |
| 1190.58000 | exclusion0300 |
| 1191.08000 | exclusion0301 |
| 1192.06000 | exclusion0302 |
| 1192.62000 | exclusion0303 |
| 1192.62000 | exclusion1769 |
| 1192.89000 | exclusion1770 |
| 1193.59000 | exclusion1771 |
| 1194.23000 | exclusion0304 |
| 1195.03000 | exclusion0305 |
| 1195.03000 | exclusion1772 |
| 1195.25000 | exclusion1773 |
| 1196.25000 | exclusion1774 |
| 1196.54000 | exclusion1775 |
| 1197.55000 | exclusion0306 |
| 1197.55000 | exclusion1776 |
| 1198.55000 | exclusion1777 |
| 1199.06000 | exclusion0307 |
| 1200.23000 | exclusion1778 |
| 1200.56000 | exclusion1779 |
| 1200.57000 | exclusion0308 |
| 1200.84000 | exclusion1780 |
| 1201.11000 | exclusion1781 |
| 1202.05000 | exclusion0309 |
| 1202.57000 | exclusion0310 |
| 1202.57000 | exclusion1782 |
| 1203.50000 | exclusion1783 |
| 1204.53000 | exclusion0311 |
| 1204.54000 | exclusion1784 |
| 1204.91000 | exclusion1785 |
| 1205.63000 | exclusion1786 |
| 1206.03000 | exclusion1787 |
| 1206.59000 | exclusion1788 |
| 1207.23000 | exclusion0312 |
| 1207.88000 | exclusion1789 |
| 1208.87000 | exclusion1790 |
| 1209.91000 | exclusion1791 |
| 1210.64000 | exclusion1792 |
| 1211.01000 | exclusion1793 |
| 1211.60000 | exclusion1794 |
| 1212.25000 | exclusion1795 |
| 1212.76000 | exclusion0313 |

| | |
|---|---|
| 1213.52000 | exclusion0314 |
| 1213.61000 | exclusion1796 |
| 1214.58000 | exclusion1797 |
| 1215.65000 | exclusion1798 |
| 1216.12000 | exclusion1799 |
| 1217.14000 | exclusion0315 |
| 1217.61000 | exclusion0316 |
| 1217.86000 | exclusion1800 |
| 1218.54000 | exclusion1801 |
| 1219.58000 | exclusion1802 |
| 1220.22000 | exclusion0317 |
| 1220.58000 | exclusion0318 |
| 1221.12000 | exclusion1803 |
| 1222.07000 | exclusion0319 |
| 1223.05000 | exclusion0320 |
| 1223.05000 | exclusion1804 |
| 1223.57000 | exclusion1805 |
| 1224.08000 | exclusion1806 |
| 1224.59000 | exclusion1807 |
| 1225.09000 | exclusion0321 |
| 1225.60000 | exclusion0322 |
| 1225.92000 | exclusion1808 |
| 1226.62000 | exclusion1809 |
| 1227.13000 | exclusion1810 |
| 1228.28000 | exclusion1811 |
| 1228.82000 | exclusion1812 |
| 1229.35000 | exclusion0323 |
| 1229.56000 | exclusion0324 |
| 1230.06000 | exclusion1813 |
| 1230.66000 | exclusion0325 |
| 1231.11000 | exclusion0326 |
| 1232.05000 | exclusion1814 |
| 1232.59000 | exclusion0327 |
| 1233.10000 | exclusion0328 |
| 1233.30000 | exclusion1815 |
| 1234.12000 | exclusion1816 |
| 1235.07000 | exclusion1817 |
| 1235.58000 | exclusion1818 |
| 1237.06000 | exclusion0329 |
| 1237.12000 | exclusion1819 |
| 1237.95000 | exclusion0330 |
| 1240.06000 | exclusion0331 |
| 1241.05000 | exclusion1820 |
| 1241.64000 | exclusion1821 |
| 1241.87000 | exclusion1822 |
| 1243.30000 | exclusion0332 |
| 1244.09000 | exclusion0333 |
| 1245.11000 | exclusion1823 |
| 1245.92000 | exclusion1824 |
| 1246.97000 | exclusion1825 |
| 1247.36000 | exclusion1826 |

| | |
|---|---|
| 1247.59000 | exclusion1827 |
| 1248.57000 | exclusion1828 |
| 1249.04000 | exclusion1829 |
| 1249.65000 | exclusion0334 |
| 1250.60000 | exclusion0335 |
| 1250.60000 | exclusion1830 |
| 1252.19000 | exclusion1831 |
| 1252.20000 | exclusion0336 |
| 1252.59000 | exclusion1832 |
| 1253.07000 | exclusion1833 |
| 1253.59000 | exclusion1834 |
| 1254.63000 | exclusion1835 |
| 1255.14000 | exclusion0337 |
| 1256.59000 | exclusion0338 |
| 1256.79000 | exclusion1836 |
| 1257.09000 | exclusion0339 |
| 1257.09000 | exclusion1837 |
| 1257.64000 | exclusion0340 |
| 1258.57000 | exclusion1838 |
| 1259.21000 | exclusion1839 |
| 1260.13000 | exclusion0341 |
| 1260.63000 | exclusion1840 |
| 1260.64000 | exclusion0342 |
| 1261.60000 | exclusion0343 |
| 1263.27000 | exclusion0344 |
| 1264.07000 | exclusion0345 |
| 1264.16000 | exclusion1841 |
| 1265.08000 | exclusion1842 |
| 1265.59000 | exclusion1843 |
| 1265.97000 | exclusion1844 |
| 1266.61000 | exclusion0346 |
| 1267.25000 | exclusion0347 |
| 1268.19000 | exclusion0348 |
| 1268.81000 | exclusion0349 |
| 1269.15000 | exclusion0350 |
| 1270.26000 | exclusion0351 |
| 1270.27000 | exclusion1845 |
| 1271.18000 | exclusion1846 |
| 1272.11000 | exclusion0352 |
| 1272.67000 | exclusion0353 |
| 1272.68000 | exclusion1847 |
| 1274.07000 | exclusion0354 |
| 1275.16000 | exclusion0355 |
| 1275.62000 | exclusion0356 |
| 1275.65000 | exclusion1848 |
| 1276.64000 | exclusion1849 |
| 1277.14000 | exclusion0357 |
| 1277.67000 | exclusion0358 |
| 1278.46000 | exclusion0359 |
| 1279.01000 | exclusion0360 |
| 1279.02000 | exclusion1850 |

| | |
|---|---|
| 1279.61000 | exclusion1851 |
| 1283.66000 | exclusion0361 |
| 1283.66000 | exclusion1852 |
| 1284.64000 | exclusion1853 |
| 1286.65000 | exclusion0362 |
| 1286.66000 | exclusion1854 |
| 1287.08000 | exclusion1855 |
| 1287.66000 | exclusion0363 |
| 1288.63000 | exclusion0364 |
| 1288.64000 | exclusion1856 |
| 1289.45000 | exclusion1857 |
| 1291.08000 | exclusion1858 |
| 1292.39000 | exclusion1859 |
| 1293.18000 | exclusion1860 |
| 1294.10000 | exclusion0365 |
| 1294.61000 | exclusion1861 |
| 1295.33000 | exclusion1862 |
| 1296.64000 | exclusion0366 |
| 1298.08000 | exclusion1863 |
| 1298.58000 | exclusion0367 |
| 1299.13000 | exclusion0368 |
| 1300.07000 | exclusion1864 |
| 1301.06000 | exclusion1865 |
| 1301.57000 | exclusion0369 |
| 1302.09000 | exclusion1866 |
| 1302.10000 | exclusion0370 |
| 1304.08000 | exclusion0371 |
| 1304.09000 | exclusion1867 |
| 1304.65000 | exclusion0372 |
| 1305.69000 | exclusion0373 |
| 1306.06000 | exclusion0374 |
| 1307.12000 | exclusion1868 |
| 1308.00000 | exclusion0375 |
| 1309.07000 | exclusion0376 |
| 1310.67000 | exclusion0377 |
| 1311.63000 | exclusion0378 |
| 1312.61000 | exclusion1869 |
| 1312.67000 | exclusion0379 |
| 1313.62000 | exclusion0380 |
| 1313.62000 | exclusion1870 |
| 1314.61000 | exclusion0381 |
| 1315.13000 | exclusion0382 |
| 1315.59000 | exclusion1871 |
| 1315.69000 | exclusion0383 |
| 1316.64000 | exclusion0384 |
| 1318.03000 | exclusion0385 |
| 1318.61000 | exclusion0386 |
| 1319.74000 | exclusion1872 |
| 1320.09000 | exclusion0387 |
| 1320.66000 | exclusion1873 |
| 1321.12000 | exclusion0388 |

| | |
|---|---|
| 1321.12000 | exclusion1874 |
| 1322.61000 | exclusion0389 |
| 1323.60000 | exclusion0390 |
| 1324.18000 | exclusion1875 |
| 1324.61000 | exclusion1876 |
| 1325.11000 | exclusion0391 |
| 1326.10000 | exclusion0392 |
| 1328.19000 | exclusion0393 |
| 1328.25000 | exclusion1877 |
| 1330.61000 | exclusion0394 |
| 1331.19000 | exclusion0395 |
| 1331.20000 | exclusion1878 |
| 1331.82000 | exclusion0396 |
| 1332.12000 | exclusion0397 |
| 1332.19000 | exclusion1879 |
| 1332.65000 | exclusion0398 |
| 1333.65000 | exclusion0399 |
| 1333.65000 | exclusion1880 |
| 1334.06000 | exclusion0400 |
| 1335.19000 | exclusion0401 |
| 1335.67000 | exclusion0402 |
| 1335.68000 | exclusion1881 |
| 1336.17000 | exclusion0403 |
| 1337.18000 | exclusion1882 |
| 1337.19000 | exclusion0404 |
| 1339.66000 | exclusion0405 |
| 1339.68000 | exclusion1883 |
| 1341.16000 | exclusion0406 |
| 1341.71000 | exclusion1884 |
| 1342.13000 | exclusion0407 |
| 1342.13000 | exclusion1885 |
| 1343.68000 | exclusion0408 |
| 1344.18000 | exclusion1886 |
| 1344.63000 | exclusion0409 |
| 1346.20000 | exclusion1887 |
| 1346.61000 | exclusion1888 |
| 1346.62000 | exclusion0410 |
| 1347.12000 | exclusion0411 |
| 1347.63000 | exclusion1889 |
| 1348.36000 | exclusion0412 |
| 1348.36000 | exclusion1890 |
| 1348.67000 | exclusion0413 |
| 1349.10000 | exclusion0414 |
| 1349.63000 | exclusion0415 |
| 1351.64000 | exclusion0416 |
| 1352.66000 | exclusion0417 |
| 1352.66000 | exclusion1891 |
| 1354.16000 | exclusion0418 |
| 1354.19000 | exclusion1892 |
| 1354.68000 | exclusion0419 |
| 1355.21000 | exclusion0420 |

| | |
|---|---|
| 1355.61000 | exclusion0421 |
| 1355.65000 | exclusion1893 |
| 1356.17000 | exclusion0422 |
| 1356.66000 | exclusion0423 |
| 1356.72000 | exclusion1894 |
| 1357.19000 | exclusion1895 |
| 1357.61000 | exclusion0424 |
| 1359.64000 | exclusion0425 |
| 1360.23000 | exclusion0426 |
| 1361.19000 | exclusion1896 |
| 1361.70000 | exclusion0427 |
| 1363.66000 | exclusion0428 |
| 1364.21000 | exclusion0429 |
| 1365.18000 | exclusion1897 |
| 1365.19000 | exclusion0430 |
| 1365.69000 | exclusion0431 |
| 1366.64000 | exclusion0432 |
| 1367.70000 | exclusion0433 |
| 1368.17000 | exclusion0434 |
| 1368.17000 | exclusion1898 |
| 1368.69000 | exclusion1899 |
| 1368.74000 | exclusion0435 |
| 1371.20000 | exclusion0436 |
| 1372.19000 | exclusion0437 |
| 1373.15000 | exclusion0438 |
| 1374.18000 | exclusion0439 |
| 1374.38000 | exclusion1900 |
| 1374.68000 | exclusion1901 |
| 1375.70000 | exclusion0440 |
| 1376.17000 | exclusion0441 |
| 1376.21000 | exclusion1902 |
| 1376.68000 | exclusion0442 |
| 1377.18000 | exclusion1903 |
| 1377.19000 | exclusion0443 |
| 1380.13000 | exclusion0444 |
| 1380.13000 | exclusion1904 |
| 1381.15000 | exclusion0445 |
| 1381.15000 | exclusion1905 |
| 1382.04000 | exclusion0446 |
| 1382.71000 | exclusion1906 |
| 1382.72000 | exclusion0447 |
| 1383.62000 | exclusion0448 |
| 1384.69000 | exclusion0449 |
| 1384.69000 | exclusion1907 |
| 1385.11000 | exclusion1908 |
| 1385.12000 | exclusion0450 |
| 1387.20000 | exclusion0451 |
| 1387.20000 | exclusion1909 |
| 1388.70000 | exclusion0452 |
| 1389.68000 | exclusion0453 |
| 1390.19000 | exclusion1910 |

| | |
|---|---|
| 1392.67000 | exclusion0454 |
| 1392.69000 | exclusion1911 |
| 1395.15000 | exclusion0455 |
| 1395.16000 | exclusion1912 |
| 1395.68000 | exclusion0456 |
| 1395.68000 | exclusion1913 |
| 1396.70000 | exclusion0457 |
| 1397.11000 | exclusion1914 |
| 1397.12000 | exclusion0458 |
| 1397.73000 | exclusion0459 |
| 1398.71000 | exclusion1915 |
| 1399.21000 | exclusion0460 |
| 1399.69000 | exclusion1916 |
| 1399.70000 | exclusion0461 |
| 1402.16000 | exclusion0462 |
| 1402.16000 | exclusion1917 |
| 1402.79000 | exclusion0463 |
| 1403.67000 | exclusion0464 |
| 1404.17000 | exclusion1918 |
| 1404.44000 | exclusion0465 |
| 1405.23000 | exclusion0466 |
| 1405.23000 | exclusion1919 |
| 1406.31000 | exclusion1920 |
| 1406.33000 | exclusion0467 |
| 1406.66000 | exclusion0468 |
| 1407.11000 | exclusion0469 |
| 1407.12000 | exclusion1921 |
| 1408.36000 | exclusion0470 |
| 1413.19000 | exclusion1922 |
| 1413.22000 | exclusion0471 |
| 1413.64000 | exclusion0472 |
| 1413.71000 | exclusion1923 |
| 1415.02000 | exclusion0473 |
| 1415.68000 | exclusion0474 |
| 1415.69000 | exclusion1924 |
| 1417.66000 | exclusion0475 |
| 1417.68000 | exclusion1925 |
| 1418.03000 | exclusion0476 |
| 1419.37000 | exclusion0477 |
| 1419.38000 | exclusion1926 |
| 1420.34000 | exclusion0478 |
| 1420.93000 | exclusion1927 |
| 1421.12000 | exclusion0479 |
| 1422.22000 | exclusion1928 |
| 1422.70000 | exclusion0480 |
| 1423.71000 | exclusion0481 |
| 1424.20000 | exclusion0482 |
| 1425.33000 | exclusion0483 |
| 1425.68000 | exclusion0484 |
| 1427.97000 | exclusion0485 |
| 1428.67000 | exclusion0486 |

| | |
|---|---|
| 1429.32000 | exclusion1929 |
| 1430.20000 | exclusion0487 |
| 1431.14000 | exclusion1930 |
| 1432.34000 | exclusion0488 |
| 1432.69000 | exclusion0489 |
| 1433.21000 | exclusion0490 |
| 1433.71000 | exclusion1931 |
| 1435.72000 | exclusion0491 |
| 1436.21000 | exclusion0492 |
| 1437.23000 | exclusion0493 |
| 1440.25000 | exclusion0494 |
| 1440.73000 | exclusion1932 |
| 1440.79000 | exclusion0495 |
| 1441.23000 | exclusion0496 |
| 1442.71000 | exclusion0497 |
| 1442.72000 | exclusion1933 |
| 1443.66000 | exclusion0498 |
| 1444.21000 | exclusion0499 |
| 1445.74000 | exclusion0500 |
| 1445.74000 | exclusion1934 |
| 1446.19000 | exclusion0501 |
| 1448.17000 | exclusion0502 |
| 1449.06000 | exclusion0503 |
| 1450.25000 | exclusion1935 |
| 1450.72000 | exclusion0504 |
| 1451.24000 | exclusion1936 |
| 1453.70000 | exclusion0505 |
| 1454.23000 | exclusion1937 |
| 1456.72000 | exclusion0506 |
| 1456.74000 | exclusion1938 |
| 1457.24000 | exclusion1939 |
| 1458.26000 | exclusion0507 |
| 1458.26000 | exclusion1940 |
| 1459.21000 | exclusion0508 |
| 1460.09000 | exclusion0509 |
| 1460.42000 | exclusion1941 |
| 1460.75000 | exclusion1942 |
| 1461.23000 | exclusion0510 |
| 1461.70000 | exclusion0511 |
| 1461.71000 | exclusion1943 |
| 1463.72000 | exclusion0512 |
| 1465.73000 | exclusion1944 |
| 1465.74000 | exclusion0513 |
| 1467.73000 | exclusion1945 |
| 1468.23000 | exclusion0514 |
| 1471.31000 | exclusion1946 |
| 1471.73000 | exclusion0515 |
| 1473.73000 | exclusion0516 |
| 1473.74000 | exclusion1947 |
| 1474.23000 | exclusion0517 |
| 1479.23000 | exclusion0518 |

| | |
|---|---|
| 1479.23000 | exclusion1948 |
| 1479.71000 | exclusion0519 |
| 1479.72000 | exclusion1949 |
| 1480.24000 | exclusion0520 |
| 1481.77000 | exclusion0521 |
| 1482.27000 | exclusion1950 |
| 1485.74000 | exclusion0522 |
| 1486.23000 | exclusion1951 |
| 1487.33000 | exclusion0523 |
| 1489.72000 | exclusion0524 |
| 1489.73000 | exclusion1952 |
| 1490.72000 | exclusion1953 |
| 1490.73000 | exclusion0525 |
| 1492.77000 | exclusion0526 |
| 1492.77000 | exclusion1954 |
| 1493.77000 | exclusion0527 |
| 1497.22000 | exclusion1955 |
| 1497.73000 | exclusion0528 |
| 1498.24000 | exclusion1956 |
| 1498.73000 | exclusion0529 |
| 1504.74000 | exclusion0530 |
| 1507.24000 | exclusion0531 |
| 1507.24000 | exclusion1957 |
| 1507.74000 | exclusion0532 |
| 1509.24000 | exclusion0533 |
| 1510.28000 | exclusion1958 |
| 1510.29000 | exclusion0534 |
| 1512.19000 | exclusion0535 |
| 1512.69000 | exclusion1959 |
| 1512.70000 | exclusion0536 |
| 1514.74000 | exclusion1960 |
| 1515.24000 | exclusion0537 |
| 1518.25000 | exclusion0538 |
| 1519.22000 | exclusion0539 |
| 1520.79000 | exclusion0540 |
| 1522.20000 | exclusion0541 |
| 1524.25000 | exclusion0542 |
| 1524.25000 | exclusion1961 |
| 1528.30000 | exclusion0543 |
| 1528.32000 | exclusion1962 |
| 1532.74000 | exclusion0544 |
| 1532.76000 | exclusion1963 |
| 1535.75000 | exclusion0545 |
| 1537.22000 | exclusion0546 |
| 1537.29000 | exclusion1964 |
| 1537.73000 | exclusion0547 |
| 1543.74000 | exclusion0548 |
| 1544.29000 | exclusion1965 |
| 1553.26000 | exclusion0549 |
| 1553.78000 | exclusion0550 |
| 1553.80000 | exclusion1966 |

| | |
|---|---|
| 1555.25000 | exclusion0551 |
| 1559.76000 | exclusion0552 |
| 1561.27000 | exclusion0553 |
| 1561.75000 | exclusion1967 |
| 1563.30000 | exclusion1968 |
| 1564.79000 | exclusion0554 |
| 1565.79000 | exclusion0555 |
| 1568.27000 | exclusion0556 |
| 1568.27000 | exclusion1969 |
| 1569.27000 | exclusion0557 |
| 1570.29000 | exclusion0558 |
| 1572.26000 | exclusion0559 |
| 1572.26000 | exclusion1970 |
| 1574.30000 | exclusion1971 |
| 1575.32000 | exclusion1972 |
| 1575.80000 | exclusion0560 |
| 1576.43000 | exclusion0561 |
| 1579.25000 | exclusion0562 |
| 1579.26000 | exclusion1973 |
| 1580.48000 | exclusion0563 |
| 1580.51000 | exclusion1974 |
| 1583.23000 | exclusion0564 |
| 1584.31000 | exclusion0565 |
| 1584.32000 | exclusion1975 |
| 1589.77000 | exclusion0566 |
| 1589.78000 | exclusion1976 |
| 1592.33000 | exclusion0567 |
| 1596.28000 | exclusion0568 |
| 1599.72000 | exclusion0569 |
| 1635.31000 | exclusion1977 |
| 1635.33000 | exclusion0570 |
| 1643.75000 | exclusion0571 |
| 1648.81000 | exclusion0572 |
| 1654.77000 | exclusion0573 |
| 1656.31000 | exclusion0574 |
| 1656.32000 | exclusion1978 |
| 1662.32000 | exclusion0575 |
| 1662.32000 | exclusion1979 |
| 1663.84000 | exclusion0576 |
| 1664.33000 | exclusion0577 |
| 1667.28000 | exclusion1980 |
| 1667.33000 | exclusion0578 |
| 1668.30000 | exclusion0579 |
| 1671.31000 | exclusion0580 |
| 1671.84000 | exclusion1981 |
| 1673.32000 | exclusion1982 |
| 1673.35000 | exclusion0581 |
| 1675.31000 | exclusion1983 |
| 1675.33000 | exclusion0582 |
| 1677.31000 | exclusion0583 |
| 1677.81000 | exclusion0584 |

1678.32000 exclusion1984
1679.40000 exclusion0585
1679.80000 exclusion1985
1682.80000 exclusion0586
1683.30000 exclusion0587
1684.32000 exclusion0588
1684.32000 exclusion1986
1684.85000 exclusion1987
1686.39000 exclusion0589
1687.38000 exclusion1988
1687.81000 exclusion0590
1688.81000 exclusion0591
1690.39000 exclusion0592
1690.84000 exclusion1989
1698.33000 exclusion0593
1698.34000 exclusion1990
1700.37000 exclusion0594
1701.33000 exclusion1991
1701.81000 exclusion0595
1707.86000 exclusion0596
1708.36000 exclusion1992
1760.89000 exclusion0597
1760.89000 exclusion1993
1764.34000 exclusion0598
1773.89000 exclusion0599
1773.89000 exclusion1994
1775.34000 exclusion0600
1780.39000 exclusion0601
1780.89000 exclusion1995
1784.88000 exclusion1996
1784.89000 exclusion0602
1791.88000 exclusion0603
1791.88000 exclusion1997
1802.39000 exclusion0604
1809.40000 exclusion0605
Neutral Loss Mass List: (none)
Product Mass List: (none)
Neutral loss in top: 3
Product in top: 3
Most intense if no parent masses found not enabled
Add/subtract mass not enabled
FT master scan preview mode enabled
Charge state screening enabled
Charge state dependent ETD time not enabled
Monoisotopic precursor selection enabled
Charge state rejection enabled
Unassigned charge states : rejected
Charge state 1 : rejected
Charge state 2 : not rejected
Charge state 3 : not rejected
Charge states 4+ : not rejected

Chromatography mode is disabled
Global Data Dependent Settings:
Predict ion injection time enabled
Use global parent and reject mass lists not enabled
Exclude parent mass from data dependent selection not enabled
Exclusion mass width by mass
Exclusion mass width low: 1.50000
Exclusion mass width high: 1.50000
Parent mass width by mass
Parent mass width low: 0.50000
Parent mass width high: 0.50000
Reject mass width by mass
Reject mass width low: 0.50000
Reject mass width high: 0.50000
Zoom/UltraZoom scan mass width by mass
Zoom/UltraZoom scan mass width low: 5.00
Zoom/UltraZoom scan mass width high: 5.00
FT SIM scan mass width low: 5.00
FT SIM scan mass width high: 5.00
Neutral Loss candidates processed by decreasing intensity
Neutral Loss mass width by mass
Neutral Loss mass width low: 0.50000
Neutral Loss mass width high: 0.50000
Product candidates processed by decreasing intensity
Product mass width by mass
Product mass width low: 0.50000
Product mass width high: 0.50000
MS mass range: 300.00-2000.00
MSn mass range by mass
MSn mass range: 0.00-1000000.00
Use m/z values as masses not enabled
Analog UV data dep. not enabled
Dynamic exclusion enabled
Repeat Count: 1
Repeat Duration: 30.00
Exclusion List Size: 500
Exclusion Duration: 180.00
Exclusion mass width by mass
Exclusion mass width low: 1.50000
Exclusion mass width high: 1.50000
Expiration: disabled
Isotopic data dependence not enabled
Custom Data Dependent Settings:
Not enabled

---

Pass 4 (wheat-mixed-digests_MS2_pass04_1.raw):

Creator: Orbi_30393
Last modified: 10/12/2021 by Orbi_30393
MS Run Time (min): 43.00
Sequence override of method parameters not enabled.

Divert Valve: not used during run
Contact Closure: not used during run
Syringe Pump: not used during run
MS Detector Settings:
Real-time modifications to method not enabled
Stepped collision energy not enabled
Additional Microscans:
MS2 0 0
MS3 0 0
MS4 0 0
MS5 0 0
MS6 0 0
MS7 0 0
MS8 0 0
MS9 0 0
MS10 0 0
Experiment Type: Nth Order Double Play
Tune Method: Orbitrap-tune-file_2020-03-13_HESI
Scan Event Details:
1: FTMS + p norm o(300.0-2000.0)
CV = 0.0V
2: ITMS + c norm Dep MS/MS Most intense ion from (1)
Activation Type: CID
Min. Signal Required: 3000.0
Isolation Width: 2.00
Normalized Coll. Energy: 35.0
Default Charge State: 2
Activation Q: 0.250
Activation Time: 10.000
CV = 0.0V
Scan Event 2 repeated for top 10 peaks.
Lock Masses:
Pos List Name: N/A
Source: API Source
Mass List: (none)
Neg List Name: N/A
Source: API Source
Mass List: (none)
Data Dependent Settings:
Use separate polarity settings disabled
Parent Mass List: (none)
Reject Mass List:

| MS Mass | Name |
|---|---|
| 300.14000 | exclusion0001 |
| 300.19000 | exclusion0002 |
| 300.71000 | exclusion0003 |
| 301.67000 | exclusion0004 |
| 302.19000 | exclusion0005 |
| 302.20000 | exclusion0006 |
| 302.59000 | exclusion0007 |
| 302.80000 | exclusion0008 |

| | |
|---|---|
| 303.84000 | exclusion0009 |
| 304.11000 | exclusion0010 |
| 304.16000 | exclusion0011 |
| 304.39000 | exclusion0012 |
| 304.66000 | exclusion0013 |
| 305.81000 | exclusion0014 |
| 306.18000 | exclusion0015 |
| 306.50000 | exclusion0016 |
| 306.84000 | exclusion0017 |
| 308.14000 | exclusion0018 |
| 308.66000 | exclusion0925 |
| 308.70000 | exclusion0019 |
| 309.18000 | exclusion0020 |
| 309.48000 | exclusion0021 |
| 309.67000 | exclusion0022 |
| 310.46000 | exclusion0023 |
| 311.10000 | exclusion0926 |
| 311.67000 | exclusion0024 |
| 312.17000 | exclusion0025 |
| 312.50000 | exclusion0026 |
| 313.17000 | exclusion0927 |
| 313.72000 | exclusion0027 |
| 314.70000 | exclusion0028 |
| 314.86000 | exclusion0029 |
| 315.47000 | exclusion0928 |
| 315.70000 | exclusion0929 |
| 316.87000 | exclusion0030 |
| 317.16000 | exclusion0031 |
| 317.64000 | exclusion0032 |
| 318.16000 | exclusion0033 |
| 318.49000 | exclusion0034 |
| 318.69000 | exclusion0035 |
| 319.14000 | exclusion0036 |
| 320.14000 | exclusion0037 |
| 320.49000 | exclusion0038 |
| 320.66000 | exclusion0039 |
| 320.71000 | exclusion0040 |
| 321.24000 | exclusion0930 |
| 321.66000 | exclusion0931 |
| 321.68000 | exclusion0932 |
| 322.20000 | exclusion0041 |
| 322.57000 | exclusion0042 |
| 324.15000 | exclusion0043 |
| 324.18000 | exclusion0044 |
| 325.70000 | exclusion0045 |
| 325.85000 | exclusion0046 |
| 326.18000 | exclusion0047 |
| 327.01000 | exclusion0048 |
| 327.67000 | exclusion0049 |
| 328.11000 | exclusion0050 |
| 328.69000 | exclusion0933 |

| | |
|---|---|
| 329.71000 | exclusion0051 |
| 330.19000 | exclusion0052 |
| 330.19000 | exclusion0053 |
| 330.83000 | exclusion0934 |
| 332.14000 | exclusion0054 |
| 332.20000 | exclusion0055 |
| 332.69000 | exclusion0056 |
| 333.15000 | exclusion0057 |
| 333.42000 | exclusion0058 |
| 333.83000 | exclusion0059 |
| 334.52000 | exclusion0060 |
| 335.12000 | exclusion0061 |
| 335.18000 | exclusion0062 |
| 335.68000 | exclusion0935 |
| 336.19000 | exclusion0063 |
| 336.73000 | exclusion0064 |
| 337.13000 | exclusion0065 |
| 337.51000 | exclusion0066 |
| 337.84000 | exclusion0067 |
| 338.67000 | exclusion0936 |
| 338.69000 | exclusion0937 |
| 339.17000 | exclusion0938 |
| 339.68000 | exclusion0068 |
| 340.18000 | exclusion0069 |
| 340.69000 | exclusion0939 |
| 340.83000 | exclusion0940 |
| 341.53000 | exclusion0070 |
| 341.66000 | exclusion0071 |
| 341.84000 | exclusion0072 |
| 341.91000 | exclusion0073 |
| 342.84000 | exclusion0941 |
| 343.49000 | exclusion0074 |
| 343.73000 | exclusion0075 |
| 344.23000 | exclusion0076 |
| 344.50000 | exclusion0077 |
| 345.13000 | exclusion0942 |
| 345.15000 | exclusion0943 |
| 345.70000 | exclusion0078 |
| 346.01000 | exclusion0079 |
| 346.19000 | exclusion0080 |
| 346.22000 | exclusion0081 |
| 346.71000 | exclusion0944 |
| 346.85000 | exclusion0945 |
| 346.93000 | exclusion0946 |
| 347.17000 | exclusion0947 |
| 347.68000 | exclusion0082 |
| 347.85000 | exclusion0083 |
| 348.21000 | exclusion0084 |
| 348.60000 | exclusion0085 |
| 348.65000 | exclusion0086 |
| 349.23000 | exclusion0087 |

| | |
|---|---|
| 349.68000 | exclusion0088 |
| 350.42000 | exclusion0089 |
| 350.67000 | exclusion0090 |
| 351.19000 | exclusion0091 |
| 351.40000 | exclusion0092 |
| 351.71000 | exclusion0093 |
| 352.17000 | exclusion0094 |
| 352.21000 | exclusion0095 |
| 352.65000 | exclusion0096 |
| 353.18000 | exclusion0948 |
| 353.19000 | exclusion0949 |
| 353.70000 | exclusion0097 |
| 353.80000 | exclusion0098 |
| 353.92000 | exclusion0099 |
| 354.21000 | exclusion0100 |
| 354.52000 | exclusion0101 |
| 354.64000 | exclusion0102 |
| 355.55000 | exclusion0103 |
| 355.86000 | exclusion0104 |
| 356.14000 | exclusion0105 |
| 356.69000 | exclusion0950 |
| 356.87000 | exclusion0951 |
| 357.19000 | exclusion0952 |
| 357.23000 | exclusion0953 |
| 358.22000 | exclusion0106 |
| 358.50000 | exclusion0107 |
| 358.71000 | exclusion0108 |
| 358.84000 | exclusion0109 |
| 359.83000 | exclusion0110 |
| 360.36000 | exclusion0111 |
| 360.70000 | exclusion0112 |
| 361.42000 | exclusion0113 |
| 361.69000 | exclusion0114 |
| 362.12000 | exclusion0115 |
| 363.20000 | exclusion0116 |
| 363.66000 | exclusion0117 |
| 363.83000 | exclusion0118 |
| 364.13000 | exclusion0119 |
| 364.71000 | exclusion0120 |
| 364.86000 | exclusion0121 |
| 365.23000 | exclusion0122 |
| 365.56000 | exclusion0123 |
| 365.80000 | exclusion0124 |
| 366.42000 | exclusion0125 |
| 366.70000 | exclusion0126 |
| 367.21000 | exclusion0127 |
| 367.22000 | exclusion0128 |
| 367.49000 | exclusion0129 |
| 367.70000 | exclusion0130 |
| 368.22000 | exclusion0954 |
| 368.86000 | exclusion0955 |

| | |
|---|---|
| 369.16000 | exclusion0956 |
| 369.26000 | exclusion0957 |
| 369.86000 | exclusion0131 |
| 370.18000 | exclusion0132 |
| 370.77000 | exclusion0133 |
| 371.17000 | exclusion0134 |
| 371.54000 | exclusion0135 |
| 371.67000 | exclusion0136 |
| 372.22000 | exclusion0958 |
| 372.23000 | exclusion0959 |
| 372.83000 | exclusion0137 |
| 373.19000 | exclusion0138 |
| 373.19000 | exclusion0139 |
| 374.23000 | exclusion0140 |
| 374.67000 | exclusion0141 |
| 375.21000 | exclusion0142 |
| 375.71000 | exclusion0143 |
| 376.19000 | exclusion0144 |
| 376.69000 | exclusion0960 |
| 377.68000 | exclusion0961 |
| 378.22000 | exclusion0962 |
| 378.22000 | exclusion0963 |
| 378.52000 | exclusion0964 |
| 379.20000 | exclusion0965 |
| 379.71000 | exclusion0145 |
| 379.87000 | exclusion0146 |
| 380.19000 | exclusion0147 |
| 380.70000 | exclusion0966 |
| 380.80000 | exclusion0148 |
| 381.24000 | exclusion0149 |
| 382.17000 | exclusion0150 |
| 382.19000 | exclusion0151 |
| 382.85000 | exclusion0152 |
| 383.20000 | exclusion0153 |
| 383.33000 | exclusion0154 |
| 383.49000 | exclusion0155 |
| 384.17000 | exclusion0156 |
| 384.52000 | exclusion0157 |
| 385.17000 | exclusion0158 |
| 385.54000 | exclusion0159 |
| 385.70000 | exclusion0160 |
| 386.19000 | exclusion0161 |
| 386.25000 | exclusion0162 |
| 386.58000 | exclusion0163 |
| 387.21000 | exclusion0164 |
| 387.72000 | exclusion0165 |
| 388.20000 | exclusion0166 |
| 388.23000 | exclusion0167 |
| 388.50000 | exclusion0168 |
| 388.72000 | exclusion0169 |
| 388.93000 | exclusion0170 |

| | |
|---|---|
| 389.89000 | exclusion0967 |
| 390.19000 | exclusion0968 |
| 390.94000 | exclusion0171 |
| 391.21000 | exclusion0172 |
| 391.22000 | exclusion0173 |
| 392.25000 | exclusion0969 |
| 392.87000 | exclusion0174 |
| 393.51000 | exclusion0175 |
| 393.73000 | exclusion0176 |
| 394.75000 | exclusion0177 |
| 394.89000 | exclusion0178 |
| 395.20000 | exclusion0179 |
| 395.24000 | exclusion0180 |
| 395.55000 | exclusion0181 |
| 395.70000 | exclusion0182 |
| 396.18000 | exclusion0970 |
| 396.20000 | exclusion0971 |
| 397.13000 | exclusion0183 |
| 397.21000 | exclusion0184 |
| 397.39000 | exclusion0185 |
| 397.68000 | exclusion0186 |
| 397.71000 | exclusion0187 |
| 398.54000 | exclusion0188 |
| 399.18000 | exclusion0189 |
| 399.22000 | exclusion0190 |
| 399.53000 | exclusion0191 |
| 400.76000 | exclusion0192 |
| 400.90000 | exclusion0193 |
| 401.23000 | exclusion0194 |
| 401.34000 | exclusion0195 |
| 401.41000 | exclusion0196 |
| 401.66000 | exclusion0197 |
| 402.23000 | exclusion0972 |
| 402.25000 | exclusion0198 |
| 402.44000 | exclusion0973 |
| 402.53000 | exclusion0974 |
| 403.22000 | exclusion0975 |
| 403.72000 | exclusion0199 |
| 404.19000 | exclusion0200 |
| 404.54000 | exclusion0201 |
| 405.52000 | exclusion0202 |
| 405.73000 | exclusion0203 |
| 406.22000 | exclusion0204 |
| 407.19000 | exclusion0205 |
| 407.19000 | exclusion0206 |
| 407.19000 | exclusion0207 |
| 407.76000 | exclusion0976 |
| 408.19000 | exclusion0977 |
| 409.20000 | exclusion0978 |
| 409.23000 | exclusion0979 |
| 409.86000 | exclusion0208 |

| | |
|---|---|
| 410.21000 | exclusion0209 |
| 410.72000 | exclusion0210 |
| 411.21000 | exclusion0211 |
| 411.36000 | exclusion0212 |
| 412.75000 | exclusion0980 |
| 413.23000 | exclusion0981 |
| 413.77000 | exclusion0213 |
| 414.22000 | exclusion0214 |
| 414.52000 | exclusion0215 |
| 414.73000 | exclusion0216 |
| 414.87000 | exclusion0217 |
| 415.76000 | exclusion0218 |
| 416.71000 | exclusion0982 |
| 416.71000 | exclusion0983 |
| 417.22000 | exclusion0219 |
| 417.22000 | exclusion0220 |
| 417.70000 | exclusion0221 |
| 417.95000 | exclusion0222 |
| 418.28000 | exclusion0223 |
| 418.57000 | exclusion0224 |
| 418.73000 | exclusion0225 |
| 419.20000 | exclusion0226 |
| 419.21000 | exclusion0227 |
| 420.22000 | exclusion0984 |
| 420.93000 | exclusion0228 |
| 421.26000 | exclusion0229 |
| 421.72000 | exclusion0230 |
| 421.85000 | exclusion0231 |
| 422.25000 | exclusion0232 |
| 423.21000 | exclusion0233 |
| 423.44000 | exclusion0234 |
| 423.70000 | exclusion0235 |
| 423.73000 | exclusion0236 |
| 423.96000 | exclusion0237 |
| 424.24000 | exclusion0238 |
| 424.69000 | exclusion0239 |
| 425.72000 | exclusion0240 |
| 425.96000 | exclusion0241 |
| 426.24000 | exclusion0242 |
| 426.73000 | exclusion0243 |
| 427.77000 | exclusion0244 |
| 427.96000 | exclusion0245 |
| 428.54000 | exclusion0246 |
| 428.71000 | exclusion0247 |
| 429.96000 | exclusion0248 |
| 430.25000 | exclusion0249 |
| 430.54000 | exclusion0250 |
| 430.75000 | exclusion0251 |
| 431.27000 | exclusion0985 |
| 432.46000 | exclusion0252 |
| 432.69000 | exclusion0253 |

| | |
|---|---|
| 433.01000 | exclusion0254 |
| 433.23000 | exclusion0255 |
| 433.24000 | exclusion0256 |
| 433.74000 | exclusion0257 |
| 433.86000 | exclusion0258 |
| 434.72000 | exclusion0986 |
| 435.73000 | exclusion0987 |
| 436.29000 | exclusion0259 |
| 436.43000 | exclusion0260 |
| 436.53000 | exclusion0261 |
| 436.74000 | exclusion0262 |
| 437.23000 | exclusion0263 |
| 438.20000 | exclusion0988 |
| 438.24000 | exclusion0989 |
| 439.24000 | exclusion0990 |
| 440.21000 | exclusion0991 |
| 440.26000 | exclusion0992 |
| 441.21000 | exclusion0993 |
| 441.71000 | exclusion0264 |
| 442.25000 | exclusion0265 |
| 442.25000 | exclusion0994 |
| 442.90000 | exclusion0266 |
| 443.23000 | exclusion0267 |
| 443.26000 | exclusion0268 |
| 443.71000 | exclusion0269 |
| 443.92000 | exclusion0270 |
| 444.23000 | exclusion0271 |
| 444.26000 | exclusion0272 |
| 444.76000 | exclusion0995 |
| 444.87000 | exclusion0996 |
| 445.24000 | exclusion0997 |
| 446.23000 | exclusion0998 |
| 446.75000 | exclusion0273 |
| 447.24000 | exclusion0274 |
| 448.20000 | exclusion0275 |
| 448.63000 | exclusion0276 |
| 448.97000 | exclusion0277 |
| 449.19000 | exclusion0278 |
| 449.27000 | exclusion0279 |
| 449.58000 | exclusion0280 |
| 450.24000 | exclusion0281 |
| 450.45000 | exclusion0282 |
| 450.55000 | exclusion0283 |
| 450.73000 | exclusion0284 |
| 451.19000 | exclusion0285 |
| 451.22000 | exclusion0286 |
| 451.88000 | exclusion0287 |
| 452.24000 | exclusion0288 |
| 452.72000 | exclusion0289 |
| 452.92000 | exclusion0290 |
| 453.22000 | exclusion0291 |

| | |
|---|---|
| 453.47000 | exclusion0292 |
| 454.26000 | exclusion0293 |
| 454.71000 | exclusion0294 |
| 455.22000 | exclusion0295 |
| 455.23000 | exclusion0296 |
| 455.24000 | exclusion0297 |
| 455.72000 | exclusion0298 |
| 456.25000 | exclusion0299 |
| 456.74000 | exclusion0300 |
| 457.73000 | exclusion0301 |
| 457.89000 | exclusion0302 |
| 458.23000 | exclusion0303 |
| 458.26000 | exclusion0304 |
| 459.02000 | exclusion0305 |
| 459.22000 | exclusion0306 |
| 459.57000 | exclusion0307 |
| 459.74000 | exclusion0308 |
| 460.76000 | exclusion0309 |
| 461.25000 | exclusion0310 |
| 461.62000 | exclusion0311 |
| 461.72000 | exclusion0312 |
| 462.20000 | exclusion0313 |
| 462.91000 | exclusion0314 |
| 463.23000 | exclusion0315 |
| 463.28000 | exclusion0316 |
| 463.73000 | exclusion0317 |
| 464.29000 | exclusion0318 |
| 464.52000 | exclusion0319 |
| 464.73000 | exclusion0320 |
| 464.76000 | exclusion0321 |
| 465.19000 | exclusion0322 |
| 466.24000 | exclusion0323 |
| 466.72000 | exclusion0324 |
| 467.23000 | exclusion0999 |
| 467.23000 | exclusion1000 |
| 467.76000 | exclusion0325 |
| 467.88000 | exclusion0326 |
| 468.22000 | exclusion0327 |
| 468.74000 | exclusion0328 |
| 469.24000 | exclusion0329 |
| 469.73000 | exclusion0330 |
| 469.78000 | exclusion0331 |
| 469.87000 | exclusion0332 |
| 470.23000 | exclusion0333 |
| 470.26000 | exclusion0334 |
| 470.78000 | exclusion1001 |
| 470.91000 | exclusion1002 |
| 471.54000 | exclusion0335 |
| 471.78000 | exclusion0336 |
| 472.22000 | exclusion0337 |
| 472.71000 | exclusion0338 |

| | |
|---|---|
| 473.27000 | exclusion0339 |
| 473.58000 | exclusion0340 |
| 474.21000 | exclusion0341 |
| 474.21000 | exclusion0342 |
| 474.76000 | exclusion1003 |
| 475.22000 | exclusion1004 |
| 475.75000 | exclusion0343 |
| 475.95000 | exclusion0344 |
| 476.23000 | exclusion0345 |
| 476.70000 | exclusion0346 |
| 476.73000 | exclusion0347 |
| 477.22000 | exclusion0348 |
| 477.26000 | exclusion0349 |
| 477.92000 | exclusion0350 |
| 478.21000 | exclusion0351 |
| 478.78000 | exclusion1005 |
| 478.90000 | exclusion1006 |
| 479.48000 | exclusion0352 |
| 479.77000 | exclusion0353 |
| 479.91000 | exclusion0354 |
| 480.23000 | exclusion0355 |
| 480.24000 | exclusion0356 |
| 480.58000 | exclusion0357 |
| 481.22000 | exclusion0358 |
| 481.31000 | exclusion0359 |
| 481.48000 | exclusion0360 |
| 481.75000 | exclusion0361 |
| 482.27000 | exclusion0362 |
| 483.24000 | exclusion0363 |
| 483.25000 | exclusion0364 |
| 483.75000 | exclusion0365 |
| 484.25000 | exclusion0366 |
| 484.29000 | exclusion0367 |
| 484.57000 | exclusion0368 |
| 484.91000 | exclusion0369 |
| 485.20000 | exclusion0370 |
| 485.28000 | exclusion0371 |
| 486.25000 | exclusion0372 |
| 486.62000 | exclusion0373 |
| 486.77000 | exclusion0374 |
| 486.88000 | exclusion0375 |
| 487.24000 | exclusion0376 |
| 487.75000 | exclusion0377 |
| 488.21000 | exclusion0378 |
| 488.22000 | exclusion0379 |
| 488.74000 | exclusion0380 |
| 489.24000 | exclusion0381 |
| 489.25000 | exclusion0382 |
| 489.55000 | exclusion0383 |
| 489.95000 | exclusion0384 |
| 490.23000 | exclusion0385 |

| | |
|---|---|
| 490.26000 | exclusion0386 |
| 490.78000 | exclusion0387 |
| 491.05000 | exclusion0388 |
| 491.25000 | exclusion0389 |
| 491.26000 | exclusion0390 |
| 491.74000 | exclusion0391 |
| 492.25000 | exclusion0392 |
| 492.50000 | exclusion0393 |
| 493.23000 | exclusion0394 |
| 493.25000 | exclusion0395 |
| 493.76000 | exclusion0396 |
| 494.26000 | exclusion0397 |
| 494.77000 | exclusion0398 |
| 494.92000 | exclusion0399 |
| 495.27000 | exclusion0400 |
| 495.72000 | exclusion0401 |
| 495.76000 | exclusion0402 |
| 496.19000 | exclusion0403 |
| 496.74000 | exclusion1007 |
| 497.49000 | exclusion1008 |
| 498.24000 | exclusion0404 |
| 498.27000 | exclusion0405 |
| 498.76000 | exclusion0406 |
| 499.26000 | exclusion0407 |
| 499.55000 | exclusion0408 |
| 499.80000 | exclusion0409 |
| 499.95000 | exclusion0410 |
| 500.56000 | exclusion0411 |
| 500.70000 | exclusion0412 |
| 500.79000 | exclusion0413 |
| 501.21000 | exclusion0414 |
| 501.26000 | exclusion0415 |
| 501.57000 | exclusion0416 |
| 501.78000 | exclusion0417 |
| 502.24000 | exclusion0418 |
| 502.76000 | exclusion0419 |
| 503.11000 | exclusion0420 |
| 503.25000 | exclusion0421 |
| 503.59000 | exclusion0422 |
| 503.71000 | exclusion0423 |
| 504.22000 | exclusion0424 |
| 504.23000 | exclusion0425 |
| 505.23000 | exclusion0426 |
| 505.25000 | exclusion0427 |
| 505.58000 | exclusion0428 |
| 505.94000 | exclusion0429 |
| 506.26000 | exclusion0430 |
| 506.27000 | exclusion0431 |
| 506.81000 | exclusion0432 |
| 506.91000 | exclusion0433 |
| 507.25000 | exclusion0434 |

| | |
|---|---|
| 507.27000 | exclusion0435 |
| 507.80000 | exclusion1009 |
| 507.88000 | exclusion1010 |
| 507.99000 | exclusion1011 |
| 508.26000 | exclusion1012 |
| 508.78000 | exclusion0436 |
| 509.24000 | exclusion0437 |
| 509.26000 | exclusion0438 |
| 509.29000 | exclusion0439 |
| 509.76000 | exclusion1013 |
| 510.28000 | exclusion1014 |
| 510.76000 | exclusion1015 |
| 511.26000 | exclusion0440 |
| 511.26000 | exclusion0441 |
| 511.76000 | exclusion0442 |
| 512.23000 | exclusion0443 |
| 512.28000 | exclusion0444 |
| 512.56000 | exclusion0445 |
| 512.75000 | exclusion0446 |
| 513.24000 | exclusion0447 |
| 513.63000 | exclusion0448 |
| 513.77000 | exclusion0449 |
| 514.57000 | exclusion0450 |
| 514.81000 | exclusion0451 |
| 515.28000 | exclusion0452 |
| 515.78000 | exclusion0453 |
| 515.92000 | exclusion0454 |
| 516.23000 | exclusion0455 |
| 516.80000 | exclusion0456 |
| 517.24000 | exclusion0457 |
| 517.25000 | exclusion0458 |
| 517.73000 | exclusion0459 |
| 518.24000 | exclusion0460 |
| 518.65000 | exclusion0461 |
| 518.78000 | exclusion0462 |
| 519.14000 | exclusion0463 |
| 519.73000 | exclusion0464 |
| 519.93000 | exclusion0465 |
| 520.31000 | exclusion0466 |
| 520.92000 | exclusion0467 |
| 521.02000 | exclusion0468 |
| 521.24000 | exclusion0469 |
| 521.53000 | exclusion0470 |
| 521.76000 | exclusion0471 |
| 522.24000 | exclusion0472 |
| 522.60000 | exclusion0473 |
| 522.77000 | exclusion0474 |
| 523.29000 | exclusion0475 |
| 523.52000 | exclusion0476 |
| 524.22000 | exclusion0477 |
| 524.59000 | exclusion0478 |

| | |
|---|---|
| 524.77000 | exclusion0479 |
| 524.81000 | exclusion0480 |
| 524.89000 | exclusion0481 |
| 525.24000 | exclusion0482 |
| 526.21000 | exclusion0483 |
| 526.44000 | exclusion0484 |
| 526.59000 | exclusion0485 |
| 526.77000 | exclusion0486 |
| 527.76000 | exclusion0487 |
| 528.24000 | exclusion0488 |
| 529.23000 | exclusion0489 |
| 529.79000 | exclusion0490 |
| 530.26000 | exclusion0491 |
| 530.30000 | exclusion0492 |
| 530.74000 | exclusion0493 |
| 531.29000 | exclusion0494 |
| 531.75000 | exclusion0495 |
| 532.25000 | exclusion0496 |
| 532.58000 | exclusion0497 |
| 532.77000 | exclusion1016 |
| 533.28000 | exclusion0498 |
| 533.63000 | exclusion0499 |
| 533.76000 | exclusion0500 |
| 533.90000 | exclusion0501 |
| 534.28000 | exclusion0502 |
| 534.71000 | exclusion0503 |
| 535.27000 | exclusion0504 |
| 535.59000 | exclusion0505 |
| 535.76000 | exclusion0506 |
| 536.09000 | exclusion0507 |
| 536.28000 | exclusion0508 |
| 536.33000 | exclusion0509 |
| 536.51000 | exclusion0510 |
| 537.79000 | exclusion0511 |
| 537.80000 | exclusion0512 |
| 537.95000 | exclusion0513 |
| 538.29000 | exclusion0514 |
| 538.77000 | exclusion0515 |
| 538.78000 | exclusion0516 |
| 539.30000 | exclusion1017 |
| 539.72000 | exclusion1018 |
| 540.27000 | exclusion1019 |
| 540.78000 | exclusion0517 |
| 540.94000 | exclusion0518 |
| 541.26000 | exclusion0519 |
| 541.76000 | exclusion0520 |
| 542.31000 | exclusion0521 |
| 542.78000 | exclusion0522 |
| 542.92000 | exclusion0523 |
| 543.27000 | exclusion0524 |
| 543.32000 | exclusion0525 |

| | |
|---|---|
| 543.76000 | exclusion0526 |
| 543.95000 | exclusion0527 |
| 544.22000 | exclusion0528 |
| 544.30000 | exclusion0529 |
| 544.51000 | exclusion0530 |
| 544.76000 | exclusion1020 |
| 545.25000 | exclusion1021 |
| 545.76000 | exclusion0531 |
| 546.02000 | exclusion0532 |
| 546.28000 | exclusion0533 |
| 546.61000 | exclusion0534 |
| 546.75000 | exclusion0535 |
| 547.28000 | exclusion0536 |
| 547.31000 | exclusion0537 |
| 547.61000 | exclusion0538 |
| 547.77000 | exclusion0539 |
| 548.24000 | exclusion0540 |
| 548.77000 | exclusion1022 |
| 548.79000 | exclusion0541 |
| 549.27000 | exclusion0542 |
| 549.75000 | exclusion0543 |
| 549.92000 | exclusion0544 |
| 549.98000 | exclusion0545 |
| 550.26000 | exclusion0546 |
| 550.78000 | exclusion0547 |
| 551.28000 | exclusion0548 |
| 551.78000 | exclusion0549 |
| 551.95000 | exclusion0550 |
| 552.04000 | exclusion0551 |
| 552.29000 | exclusion0552 |
| 552.74000 | exclusion0553 |
| 553.28000 | exclusion0554 |
| 553.30000 | exclusion0555 |
| 553.59000 | exclusion0556 |
| 553.77000 | exclusion0557 |
| 553.82000 | exclusion0558 |
| 554.37000 | exclusion0559 |
| 554.75000 | exclusion0560 |
| 555.29000 | exclusion0561 |
| 555.31000 | exclusion0562 |
| 555.61000 | exclusion0563 |
| 555.64000 | exclusion0564 |
| 555.75000 | exclusion0565 |
| 556.90000 | exclusion0566 |
| 557.26000 | exclusion0567 |
| 557.30000 | exclusion0568 |
| 558.28000 | exclusion0569 |
| 558.77000 | exclusion0570 |
| 558.78000 | exclusion0571 |
| 559.06000 | exclusion0572 |
| 559.27000 | exclusion0573 |

| | |
|---|---|
| 559.56000 | exclusion0574 |
| 559.80000 | exclusion0575 |
| 560.78000 | exclusion0576 |
| 561.29000 | exclusion0577 |
| 561.30000 | exclusion0578 |
| 561.57000 | exclusion0579 |
| 561.79000 | exclusion0580 |
| 562.30000 | exclusion0581 |
| 562.61000 | exclusion0582 |
| 562.78000 | exclusion0583 |
| 563.30000 | exclusion0584 |
| 563.78000 | exclusion0585 |
| 564.27000 | exclusion0586 |
| 564.29000 | exclusion0587 |
| 564.58000 | exclusion0588 |
| 564.82000 | exclusion0589 |
| 565.42000 | exclusion0590 |
| 565.59000 | exclusion0591 |
| 566.22000 | exclusion0592 |
| 566.58000 | exclusion0593 |
| 566.76000 | exclusion0594 |
| 566.86000 | exclusion0595 |
| 567.29000 | exclusion0596 |
| 567.59000 | exclusion0597 |
| 567.81000 | exclusion0598 |
| 568.29000 | exclusion0599 |
| 568.31000 | exclusion0600 |
| 568.75000 | exclusion0601 |
| 569.31000 | exclusion0602 |
| 569.81000 | exclusion0603 |
| 570.08000 | exclusion0604 |
| 570.30000 | exclusion0605 |
| 570.75000 | exclusion0606 |
| 571.27000 | exclusion0607 |
| 571.77000 | exclusion0608 |
| 571.81000 | exclusion0609 |
| 572.30000 | exclusion0610 |
| 572.31000 | exclusion0611 |
| 572.67000 | exclusion0612 |
| 573.29000 | exclusion0613 |
| 573.30000 | exclusion0614 |
| 573.81000 | exclusion0615 |
| 573.96000 | exclusion0616 |
| 574.28000 | exclusion0617 |
| 574.31000 | exclusion0618 |
| 574.63000 | exclusion0619 |
| 574.79000 | exclusion0620 |
| 575.28000 | exclusion0621 |
| 575.80000 | exclusion0622 |
| 575.99000 | exclusion0623 |
| 576.21000 | exclusion0624 |

| | |
|---|---|
| 576.25000 | exclusion0625 |
| 576.71000 | exclusion0626 |
| 577.67000 | exclusion0627 |
| 577.83000 | exclusion0628 |
| 577.98000 | exclusion0629 |
| 578.25000 | exclusion0630 |
| 578.26000 | exclusion0631 |
| 578.65000 | exclusion0632 |
| 578.97000 | exclusion0633 |
| 579.34000 | exclusion0634 |
| 579.78000 | exclusion0635 |
| 580.03000 | exclusion0636 |
| 580.30000 | exclusion0637 |
| 580.31000 | exclusion0638 |
| 581.00000 | exclusion0639 |
| 581.29000 | exclusion0640 |
| 581.83000 | exclusion0641 |
| 581.96000 | exclusion0642 |
| 582.10000 | exclusion0643 |
| 582.25000 | exclusion0644 |
| 582.31000 | exclusion0645 |
| 582.74000 | exclusion0646 |
| 583.03000 | exclusion0647 |
| 583.79000 | exclusion0648 |
| 583.95000 | exclusion0649 |
| 584.27000 | exclusion0650 |
| 584.65000 | exclusion0651 |
| 584.76000 | exclusion0652 |
| 585.26000 | exclusion1023 |
| 585.28000 | exclusion1024 |
| 585.28000 | exclusion1025 |
| 585.65000 | exclusion1026 |
| 586.24000 | exclusion0653 |
| 586.31000 | exclusion0654 |
| 586.75000 | exclusion0655 |
| 587.31000 | exclusion0656 |
| 587.33000 | exclusion0657 |
| 588.00000 | exclusion0658 |
| 588.28000 | exclusion0659 |
| 588.30000 | exclusion0660 |
| 588.80000 | exclusion0661 |
| 589.30000 | exclusion0662 |
| 590.25000 | exclusion0663 |
| 590.29000 | exclusion0664 |
| 590.80000 | exclusion0665 |
| 590.84000 | exclusion0666 |
| 590.97000 | exclusion0667 |
| 591.34000 | exclusion0668 |
| 591.65000 | exclusion0669 |
| 591.79000 | exclusion0670 |
| 592.31000 | exclusion0671 |

| | |
|---|---|
| 592.85000 | exclusion0672 |
| 592.98000 | exclusion0673 |
| 593.16000 | exclusion0674 |
| 593.59000 | exclusion0675 |
| 593.61000 | exclusion0676 |
| 593.81000 | exclusion0677 |
| 594.58000 | exclusion0678 |
| 594.76000 | exclusion0679 |
| 594.80000 | exclusion0680 |
| 595.31000 | exclusion0681 |
| 595.63000 | exclusion0682 |
| 595.76000 | exclusion0683 |
| 595.92000 | exclusion0684 |
| 596.77000 | exclusion0685 |
| 597.28000 | exclusion0686 |
| 597.29000 | exclusion0687 |
| 597.78000 | exclusion0688 |
| 598.31000 | exclusion0689 |
| 598.36000 | exclusion0690 |
| 598.81000 | exclusion0691 |
| 599.35000 | exclusion0692 |
| 599.67000 | exclusion0693 |
| 599.78000 | exclusion0694 |
| 600.00000 | exclusion0695 |
| 600.26000 | exclusion0696 |
| 600.26000 | exclusion0697 |
| 600.78000 | exclusion1027 |
| 601.30000 | exclusion0698 |
| 601.31000 | exclusion0699 |
| 601.81000 | exclusion0700 |
| 602.26000 | exclusion0701 |
| 602.30000 | exclusion0702 |
| 602.31000 | exclusion0703 |
| 602.84000 | exclusion1028 |
| 603.28000 | exclusion1029 |
| 603.80000 | exclusion0704 |
| 603.98000 | exclusion0705 |
| 604.28000 | exclusion0706 |
| 604.32000 | exclusion0707 |
| 604.32000 | exclusion0708 |
| 604.79000 | exclusion1030 |
| 605.78000 | exclusion0709 |
| 606.00000 | exclusion0710 |
| 606.28000 | exclusion0711 |
| 606.28000 | exclusion0712 |
| 606.29000 | exclusion0713 |
| 606.98000 | exclusion0714 |
| 607.28000 | exclusion1031 |
| 607.30000 | exclusion1032 |
| 607.30000 | exclusion1033 |
| 607.81000 | exclusion0715 |

| | |
|---|---|
| 608.32000 | exclusion1034 |
| 608.85000 | exclusion0716 |
| 608.95000 | exclusion0717 |
| 609.31000 | exclusion0718 |
| 609.85000 | exclusion1035 |
| 610.32000 | exclusion0719 |
| 611.05000 | exclusion0720 |
| 611.27000 | exclusion0721 |
| 611.29000 | exclusion0722 |
| 611.77000 | exclusion0723 |
| 612.32000 | exclusion1036 |
| 612.33000 | exclusion0724 |
| 612.33000 | exclusion1037 |
| 612.35000 | exclusion1038 |
| 613.32000 | exclusion1039 |
| 614.09000 | exclusion0725 |
| 614.14000 | exclusion0726 |
| 614.29000 | exclusion0727 |
| 614.30000 | exclusion0728 |
| 614.77000 | exclusion0729 |
| 615.30000 | exclusion0730 |
| 615.82000 | exclusion0731 |
| 616.28000 | exclusion0732 |
| 616.32000 | exclusion0733 |
| 616.83000 | exclusion0734 |
| 617.31000 | exclusion0735 |
| 617.34000 | exclusion0736 |
| 617.62000 | exclusion0737 |
| 617.80000 | exclusion0738 |
| 617.93000 | exclusion0739 |
| 619.26000 | exclusion0740 |
| 619.30000 | exclusion0741 |
| 619.63000 | exclusion0742 |
| 620.59000 | exclusion0743 |
| 620.81000 | exclusion0744 |
| 621.29000 | exclusion0745 |
| 621.29000 | exclusion0746 |
| 621.66000 | exclusion0747 |
| 621.71000 | exclusion0748 |
| 621.98000 | exclusion0749 |
| 623.32000 | exclusion0750 |
| 623.32000 | exclusion0751 |
| 623.82000 | exclusion0752 |
| 624.01000 | exclusion0753 |
| 624.83000 | exclusion0754 |
| 625.01000 | exclusion0755 |
| 625.30000 | exclusion0756 |
| 625.33000 | exclusion0757 |
| 625.87000 | exclusion0758 |
| 635.29000 | exclusion1040 |
| 635.32000 | exclusion1041 |

| | |
|---|---|
| 635.86000 | exclusion1042 |
| 642.35000 | exclusion1043 |
| 646.80000 | exclusion1044 |
| 647.03000 | exclusion1045 |
| 652.30000 | exclusion1046 |
| 652.32000 | exclusion1047 |
| 653.32000 | exclusion1048 |
| 653.35000 | exclusion1049 |
| 671.30000 | exclusion1050 |
| 671.53000 | exclusion1051 |
| 671.83000 | exclusion1052 |
| 681.83000 | exclusion1053 |
| 681.84000 | exclusion1054 |
| 682.85000 | exclusion1055 |
| 688.27000 | exclusion1056 |
| 694.81000 | exclusion1057 |
| 711.35000 | exclusion1058 |
| 711.69000 | exclusion1059 |
| 711.89000 | exclusion1060 |
| 712.38000 | exclusion1061 |
| 712.39000 | exclusion1062 |
| 718.85000 | exclusion1063 |
| 718.85000 | exclusion1064 |
| 723.93000 | exclusion1065 |
| 724.34000 | exclusion1066 |
| 724.37000 | exclusion1067 |
| 724.68000 | exclusion1068 |
| 725.37000 | exclusion1069 |
| 745.36000 | exclusion1070 |
| 745.90000 | exclusion1071 |
| 765.34000 | exclusion1072 |
| 765.36000 | exclusion1073 |
| 766.36000 | exclusion1074 |
| 767.16000 | exclusion1075 |
| 767.37000 | exclusion1076 |
| 767.37000 | exclusion1077 |
| 767.86000 | exclusion1078 |
| 772.39000 | exclusion1079 |
| 772.39000 | exclusion1080 |
| 772.85000 | exclusion1081 |
| 773.38000 | exclusion1082 |
| 782.79000 | exclusion1083 |
| 803.37000 | exclusion1084 |
| 812.36000 | exclusion1085 |
| 812.86000 | exclusion1086 |
| 814.36000 | exclusion1087 |
| 814.41000 | exclusion1088 |
| 814.74000 | exclusion1089 |
| 816.44000 | exclusion1090 |
| 835.72000 | exclusion1091 |
| 836.06000 | exclusion1092 |

| | |
|---|---|
| 836.40000 | exclusion1093 |
| 842.11000 | exclusion1094 |
| 842.14000 | exclusion1095 |
| 842.38000 | exclusion1096 |
| 842.39000 | exclusion1097 |
| 844.44000 | exclusion1098 |
| 844.58000 | exclusion1099 |
| 844.89000 | exclusion1100 |
| 845.37000 | exclusion1101 |
| 845.41000 | exclusion1102 |
| 848.91000 | exclusion1103 |
| 849.37000 | exclusion1104 |
| 863.92000 | exclusion1105 |
| 864.42000 | exclusion1106 |
| 865.40000 | exclusion1107 |
| 866.41000 | exclusion1108 |
| 867.43000 | exclusion1109 |
| 867.44000 | exclusion0759 |
| 881.94000 | exclusion1110 |
| 903.92000 | exclusion0760 |
| 904.43000 | exclusion1111 |
| 904.44000 | exclusion1112 |
| 904.45000 | exclusion0761 |
| 906.91000 | exclusion0762 |
| 908.93000 | exclusion1113 |
| 909.43000 | exclusion1114 |
| 910.44000 | exclusion1115 |
| 910.44000 | exclusion1116 |
| 912.91000 | exclusion1117 |
| 912.99000 | exclusion1118 |
| 913.45000 | exclusion1119 |
| 913.93000 | exclusion1120 |
| 927.48000 | exclusion1121 |
| 932.94000 | exclusion1122 |
| 933.14000 | exclusion1123 |
| 933.47000 | exclusion1124 |
| 937.12000 | exclusion1125 |
| 937.46000 | exclusion1126 |
| 940.44000 | exclusion1127 |
| 940.48000 | exclusion1128 |
| 941.09000 | exclusion1129 |
| 941.96000 | exclusion0763 |
| 942.48000 | exclusion1130 |
| 960.41000 | exclusion1131 |
| 961.98000 | exclusion1132 |
| 968.97000 | exclusion1133 |
| 989.52000 | exclusion1134 |
| 992.50000 | exclusion1135 |
| 993.02000 | exclusion1136 |
| 993.50000 | exclusion1137 |
| 994.47000 | exclusion1138 |

| | |
|---|---|
| 994.98000 | exclusion1139 |
| 995.46000 | exclusion1140 |
| 996.98000 | exclusion1141 |
| 1005.98000 | exclusion1142 |
| 1013.49000 | exclusion1143 |
| 1013.79000 | exclusion1144 |
| 1014.47000 | exclusion1145 |
| 1018.41000 | exclusion1146 |
| 1024.51000 | exclusion1530 |
| 1025.42000 | exclusion1531 |
| 1025.86000 | exclusion1532 |
| 1026.79000 | exclusion1533 |
| 1027.46000 | exclusion1534 |
| 1028.02000 | exclusion1535 |
| 1028.98000 | exclusion1536 |
| 1029.51000 | exclusion1147 |
| 1030.46000 | exclusion1537 |
| 1030.53000 | exclusion1148 |
| 1031.47000 | exclusion1538 |
| 1032.46000 | exclusion1539 |
| 1032.86000 | exclusion1540 |
| 1033.15000 | exclusion1541 |
| 1033.52000 | exclusion1542 |
| 1034.02000 | exclusion1543 |
| 1034.80000 | exclusion1149 |
| 1035.54000 | exclusion1150 |
| 1035.55000 | exclusion1544 |
| 1036.20000 | exclusion1545 |
| 1036.53000 | exclusion1546 |
| 1036.77000 | exclusion1547 |
| 1037.45000 | exclusion1548 |
| 1038.03000 | exclusion1151 |
| 1038.26000 | exclusion1152 |
| 1038.46000 | exclusion1153 |
| 1038.98000 | exclusion1549 |
| 1039.29000 | exclusion1550 |
| 1039.50000 | exclusion1551 |
| 1040.04000 | exclusion1552 |
| 1040.51000 | exclusion1553 |
| 1041.47000 | exclusion1554 |
| 1041.77000 | exclusion1555 |
| 1042.02000 | exclusion1556 |
| 1042.52000 | exclusion1557 |
| 1043.14000 | exclusion1558 |
| 1044.17000 | exclusion1559 |
| 1044.52000 | exclusion1560 |
| 1044.97000 | exclusion1561 |
| 1045.46000 | exclusion1562 |
| 1046.50000 | exclusion0764 |
| 1046.50000 | exclusion1154 |
| 1046.50000 | exclusion1563 |

| | |
|---|---|
| 1047.50000 | exclusion1155 |
| 1047.82000 | exclusion1564 |
| 1048.05000 | exclusion1565 |
| 1048.84000 | exclusion1566 |
| 1049.47000 | exclusion1567 |
| 1049.87000 | exclusion1568 |
| 1050.51000 | exclusion1569 |
| 1050.77000 | exclusion1570 |
| 1051.47000 | exclusion1571 |
| 1052.00000 | exclusion1156 |
| 1052.53000 | exclusion1572 |
| 1053.06000 | exclusion1573 |
| 1053.53000 | exclusion1574 |
| 1053.84000 | exclusion1575 |
| 1054.50000 | exclusion1576 |
| 1055.04000 | exclusion1577 |
| 1055.48000 | exclusion1578 |
| 1056.04000 | exclusion1579 |
| 1056.52000 | exclusion1580 |
| 1057.55000 | exclusion0765 |
| 1057.55000 | exclusion1157 |
| 1057.56000 | exclusion1581 |
| 1057.84000 | exclusion1582 |
| 1058.07000 | exclusion1583 |
| 1058.48000 | exclusion1584 |
| 1059.27000 | exclusion1585 |
| 1059.48000 | exclusion1586 |
| 1060.03000 | exclusion1587 |
| 1060.53000 | exclusion1588 |
| 1061.02000 | exclusion1589 |
| 1061.83000 | exclusion1590 |
| 1062.24000 | exclusion1591 |
| 1062.54000 | exclusion1592 |
| 1063.01000 | exclusion1593 |
| 1063.48000 | exclusion1594 |
| 1064.48000 | exclusion1595 |
| 1065.47000 | exclusion1596 |
| 1066.11000 | exclusion1158 |
| 1066.54000 | exclusion1159 |
| 1067.04000 | exclusion1597 |
| 1067.49000 | exclusion1598 |
| 1067.53000 | exclusion1160 |
| 1068.04000 | exclusion1599 |
| 1068.47000 | exclusion1600 |
| 1069.03000 | exclusion1161 |
| 1069.45000 | exclusion1162 |
| 1070.06000 | exclusion1601 |
| 1070.52000 | exclusion1602 |
| 1071.15000 | exclusion1603 |
| 1071.54000 | exclusion1604 |
| 1072.07000 | exclusion1163 |

| | |
|---|---|
| 1072.54000 | exclusion1164 |
| 1072.84000 | exclusion1165 |
| 1072.87000 | exclusion1605 |
| 1073.53000 | exclusion1606 |
| 1074.04000 | exclusion1607 |
| 1074.55000 | exclusion1166 |
| 1074.85000 | exclusion1167 |
| 1075.13000 | exclusion1168 |
| 1075.52000 | exclusion1169 |
| 1075.53000 | exclusion1608 |
| 1076.06000 | exclusion1609 |
| 1077.04000 | exclusion1610 |
| 1077.54000 | exclusion1611 |
| 1078.07000 | exclusion0766 |
| 1078.56000 | exclusion0767 |
| 1078.99000 | exclusion1612 |
| 1079.51000 | exclusion1613 |
| 1080.50000 | exclusion1614 |
| 1080.85000 | exclusion1615 |
| 1081.53000 | exclusion1616 |
| 1082.19000 | exclusion1617 |
| 1082.53000 | exclusion1618 |
| 1083.55000 | exclusion1170 |
| 1083.58000 | exclusion1619 |
| 1084.06000 | exclusion1620 |
| 1084.55000 | exclusion1621 |
| 1085.04000 | exclusion1622 |
| 1085.53000 | exclusion1623 |
| 1086.05000 | exclusion1624 |
| 1086.49000 | exclusion1625 |
| 1086.99000 | exclusion1171 |
| 1087.55000 | exclusion1172 |
| 1087.55000 | exclusion1626 |
| 1087.88000 | exclusion1627 |
| 1088.24000 | exclusion1628 |
| 1088.49000 | exclusion1629 |
| 1089.00000 | exclusion1173 |
| 1089.53000 | exclusion1174 |
| 1090.05000 | exclusion1175 |
| 1090.06000 | exclusion1630 |
| 1090.53000 | exclusion1631 |
| 1091.06000 | exclusion1632 |
| 1091.89000 | exclusion1633 |
| 1092.50000 | exclusion1176 |
| 1093.03000 | exclusion1177 |
| 1093.53000 | exclusion1634 |
| 1093.87000 | exclusion1635 |
| 1094.55000 | exclusion1636 |
| 1095.50000 | exclusion1637 |
| 1095.86000 | exclusion1638 |
| 1096.12000 | exclusion1639 |

| | |
|---|---|
| 1097.16000 | exclusion1178 |
| 1097.29000 | exclusion1640 |
| 1097.54000 | exclusion1641 |
| 1098.20000 | exclusion1642 |
| 1099.29000 | exclusion1179 |
| 1099.53000 | exclusion1643 |
| 1099.54000 | exclusion0768 |
| 1100.03000 | exclusion1180 |
| 1100.56000 | exclusion1181 |
| 1100.81000 | exclusion1182 |
| 1101.09000 | exclusion1183 |
| 1101.09000 | exclusion1644 |
| 1102.22000 | exclusion1645 |
| 1102.51000 | exclusion1646 |
| 1102.88000 | exclusion1647 |
| 1103.99000 | exclusion1648 |
| 1104.22000 | exclusion1649 |
| 1104.58000 | exclusion1650 |
| 1105.54000 | exclusion1651 |
| 1105.77000 | exclusion1652 |
| 1106.53000 | exclusion1653 |
| 1107.54000 | exclusion1654 |
| 1108.05000 | exclusion1184 |
| 1108.58000 | exclusion1655 |
| 1109.52000 | exclusion1656 |
| 1109.88000 | exclusion1657 |
| 1110.23000 | exclusion1658 |
| 1110.91000 | exclusion1185 |
| 1111.54000 | exclusion1186 |
| 1111.56000 | exclusion1659 |
| 1111.87000 | exclusion1660 |
| 1112.08000 | exclusion1661 |
| 1112.80000 | exclusion1662 |
| 1113.00000 | exclusion1663 |
| 1113.52000 | exclusion1664 |
| 1114.04000 | exclusion1187 |
| 1114.85000 | exclusion1188 |
| 1114.94000 | exclusion1665 |
| 1115.55000 | exclusion1666 |
| 1116.08000 | exclusion1667 |
| 1116.88000 | exclusion1668 |
| 1117.52000 | exclusion1669 |
| 1118.09000 | exclusion1670 |
| 1118.58000 | exclusion1671 |
| 1118.86000 | exclusion1672 |
| 1119.89000 | exclusion1673 |
| 1120.52000 | exclusion1674 |
| 1121.05000 | exclusion1675 |
| 1121.56000 | exclusion1189 |
| 1121.88000 | exclusion1190 |
| 1122.19000 | exclusion1676 |

| | |
|---|---|
| 1122.51000 | exclusion1677 |
| 1123.21000 | exclusion1678 |
| 1123.56000 | exclusion1679 |
| 1124.57000 | exclusion1680 |
| 1125.02000 | exclusion1681 |
| 1125.54000 | exclusion1682 |
| 1126.04000 | exclusion1683 |
| 1126.54000 | exclusion1191 |
| 1127.00000 | exclusion1192 |
| 1127.21000 | exclusion1193 |
| 1127.55000 | exclusion1684 |
| 1127.87000 | exclusion1685 |
| 1128.09000 | exclusion1686 |
| 1128.53000 | exclusion1687 |
| 1129.58000 | exclusion0769 |
| 1129.58000 | exclusion1194 |
| 1130.12000 | exclusion1195 |
| 1130.21000 | exclusion1688 |
| 1130.54000 | exclusion1196 |
| 1130.58000 | exclusion1689 |
| 1131.03000 | exclusion1690 |
| 1131.57000 | exclusion1691 |
| 1132.20000 | exclusion1692 |
| 1132.87000 | exclusion1693 |
| 1133.08000 | exclusion1694 |
| 1133.62000 | exclusion1695 |
| 1133.89000 | exclusion1696 |
| 1134.54000 | exclusion1697 |
| 1135.11000 | exclusion1698 |
| 1136.04000 | exclusion1699 |
| 1136.53000 | exclusion1700 |
| 1137.44000 | exclusion1701 |
| 1138.55000 | exclusion0770 |
| 1138.55000 | exclusion1197 |
| 1138.55000 | exclusion1702 |
| 1139.18000 | exclusion1703 |
| 1139.54000 | exclusion1704 |
| 1140.20000 | exclusion1198 |
| 1140.58000 | exclusion1199 |
| 1140.83000 | exclusion1200 |
| 1141.52000 | exclusion1705 |
| 1141.90000 | exclusion1706 |
| 1142.51000 | exclusion1707 |
| 1143.05000 | exclusion1708 |
| 1143.56000 | exclusion1709 |
| 1144.09000 | exclusion1201 |
| 1144.54000 | exclusion1202 |
| 1145.04000 | exclusion1710 |
| 1145.51000 | exclusion1203 |
| 1145.53000 | exclusion1711 |
| 1146.56000 | exclusion1712 |

| | |
|---|---|
| 1147.00000 | exclusion1713 |
| 1147.53000 | exclusion1714 |
| 1148.07000 | exclusion1715 |
| 1148.56000 | exclusion1716 |
| 1149.06000 | exclusion1717 |
| 1149.56000 | exclusion1718 |
| 1150.56000 | exclusion1204 |
| 1150.56000 | exclusion1719 |
| 1151.05000 | exclusion1720 |
| 1151.60000 | exclusion1721 |
| 1152.12000 | exclusion1722 |
| 1152.55000 | exclusion1723 |
| 1153.54000 | exclusion1724 |
| 1154.20000 | exclusion1725 |
| 1154.55000 | exclusion1726 |
| 1155.08000 | exclusion1727 |
| 1156.07000 | exclusion1728 |
| 1156.52000 | exclusion1729 |
| 1157.56000 | exclusion1730 |
| 1158.23000 | exclusion1731 |
| 1158.51000 | exclusion1732 |
| 1159.44000 | exclusion1733 |
| 1159.64000 | exclusion1734 |
| 1159.91000 | exclusion1735 |
| 1160.55000 | exclusion0771 |
| 1161.05000 | exclusion0772 |
| 1161.55000 | exclusion0773 |
| 1161.55000 | exclusion1205 |
| 1161.57000 | exclusion1736 |
| 1162.04000 | exclusion1737 |
| 1162.58000 | exclusion1738 |
| 1163.03000 | exclusion1739 |
| 1163.58000 | exclusion1206 |
| 1163.89000 | exclusion1207 |
| 1164.56000 | exclusion1740 |
| 1165.57000 | exclusion0774 |
| 1165.57000 | exclusion1741 |
| 1166.27000 | exclusion1208 |
| 1166.53000 | exclusion1209 |
| 1166.90000 | exclusion1210 |
| 1167.20000 | exclusion1742 |
| 1167.21000 | exclusion1211 |
| 1167.52000 | exclusion1743 |
| 1168.20000 | exclusion1744 |
| 1168.55000 | exclusion1745 |
| 1169.07000 | exclusion0775 |
| 1169.58000 | exclusion1746 |
| 1170.08000 | exclusion1212 |
| 1170.56000 | exclusion1213 |
| 1171.09000 | exclusion1747 |
| 1171.61000 | exclusion1748 |

| | |
|---|---|
| 1172.54000 | exclusion1214 |
| 1173.26000 | exclusion1215 |
| 1173.27000 | exclusion1749 |
| 1173.56000 | exclusion1750 |
| 1174.54000 | exclusion1751 |
| 1175.48000 | exclusion1752 |
| 1176.00000 | exclusion1216 |
| 1176.58000 | exclusion1217 |
| 1176.59000 | exclusion1753 |
| 1177.58000 | exclusion1754 |
| 1178.09000 | exclusion1755 |
| 1178.50000 | exclusion1756 |
| 1179.08000 | exclusion1218 |
| 1179.58000 | exclusion1757 |
| 1180.14000 | exclusion1758 |
| 1180.94000 | exclusion1219 |
| 1181.56000 | exclusion1759 |
| 1181.91000 | exclusion1220 |
| 1182.54000 | exclusion1760 |
| 1182.93000 | exclusion1761 |
| 1183.53000 | exclusion1221 |
| 1184.10000 | exclusion1762 |
| 1184.55000 | exclusion1763 |
| 1185.54000 | exclusion1764 |
| 1186.47000 | exclusion1222 |
| 1187.05000 | exclusion1223 |
| 1187.10000 | exclusion1765 |
| 1187.58000 | exclusion1766 |
| 1188.57000 | exclusion1767 |
| 1189.54000 | exclusion1768 |
| 1189.91000 | exclusion1769 |
| 1190.58000 | exclusion1224 |
| 1191.08000 | exclusion1225 |
| 1192.06000 | exclusion1226 |
| 1192.62000 | exclusion1227 |
| 1192.62000 | exclusion1770 |
| 1192.89000 | exclusion1771 |
| 1193.59000 | exclusion1772 |
| 1194.23000 | exclusion1228 |
| 1195.03000 | exclusion1229 |
| 1195.03000 | exclusion1773 |
| 1195.25000 | exclusion1774 |
| 1196.25000 | exclusion1775 |
| 1196.54000 | exclusion1776 |
| 1197.05000 | exclusion0776 |
| 1197.55000 | exclusion1230 |
| 1197.55000 | exclusion1777 |
| 1198.55000 | exclusion1778 |
| 1199.06000 | exclusion1231 |
| 1199.57000 | exclusion0777 |
| 1200.23000 | exclusion0778 |

| | |
|---|---|
| 1200.23000 | exclusion1779 |
| 1200.56000 | exclusion0779 |
| 1200.56000 | exclusion1780 |
| 1200.57000 | exclusion1232 |
| 1200.84000 | exclusion1781 |
| 1201.11000 | exclusion1782 |
| 1202.05000 | exclusion1233 |
| 1202.57000 | exclusion1234 |
| 1202.57000 | exclusion1783 |
| 1203.50000 | exclusion1784 |
| 1204.53000 | exclusion1235 |
| 1204.54000 | exclusion1785 |
| 1204.91000 | exclusion1786 |
| 1205.63000 | exclusion1787 |
| 1206.03000 | exclusion1788 |
| 1206.59000 | exclusion1789 |
| 1207.23000 | exclusion1236 |
| 1207.88000 | exclusion1790 |
| 1208.87000 | exclusion1791 |
| 1209.91000 | exclusion1792 |
| 1210.64000 | exclusion1793 |
| 1211.01000 | exclusion1794 |
| 1211.60000 | exclusion1795 |
| 1212.25000 | exclusion1796 |
| 1212.76000 | exclusion1237 |
| 1213.52000 | exclusion1238 |
| 1213.61000 | exclusion1797 |
| 1214.58000 | exclusion1798 |
| 1215.10000 | exclusion0780 |
| 1215.65000 | exclusion1799 |
| 1216.12000 | exclusion1800 |
| 1217.14000 | exclusion1239 |
| 1217.61000 | exclusion1240 |
| 1217.86000 | exclusion1801 |
| 1218.54000 | exclusion1802 |
| 1219.58000 | exclusion1803 |
| 1220.22000 | exclusion1241 |
| 1220.58000 | exclusion1242 |
| 1221.12000 | exclusion1804 |
| 1222.07000 | exclusion1243 |
| 1223.05000 | exclusion1244 |
| 1223.05000 | exclusion1805 |
| 1223.57000 | exclusion1806 |
| 1224.08000 | exclusion1807 |
| 1224.59000 | exclusion1808 |
| 1225.09000 | exclusion1245 |
| 1225.60000 | exclusion1246 |
| 1225.92000 | exclusion1809 |
| 1226.62000 | exclusion1810 |
| 1227.13000 | exclusion1811 |
| 1227.66000 | exclusion0781 |

| | |
|---|---|
| 1228.28000 | exclusion1812 |
| 1228.82000 | exclusion1813 |
| 1229.35000 | exclusion1247 |
| 1229.56000 | exclusion1248 |
| 1230.06000 | exclusion1814 |
| 1230.66000 | exclusion1249 |
| 1231.11000 | exclusion1250 |
| 1232.05000 | exclusion1815 |
| 1232.59000 | exclusion1251 |
| 1233.10000 | exclusion1252 |
| 1233.30000 | exclusion1816 |
| 1234.12000 | exclusion1817 |
| 1235.07000 | exclusion1818 |
| 1235.58000 | exclusion1819 |
| 1236.08000 | exclusion0782 |
| 1237.06000 | exclusion1253 |
| 1237.12000 | exclusion1820 |
| 1237.95000 | exclusion1254 |
| 1240.06000 | exclusion0783 |
| 1240.06000 | exclusion1255 |
| 1241.05000 | exclusion1821 |
| 1241.64000 | exclusion1822 |
| 1241.87000 | exclusion1823 |
| 1242.58000 | exclusion0784 |
| 1243.30000 | exclusion1256 |
| 1244.09000 | exclusion1257 |
| 1244.59000 | exclusion0785 |
| 1245.11000 | exclusion1824 |
| 1245.92000 | exclusion1825 |
| 1246.97000 | exclusion1826 |
| 1247.36000 | exclusion1827 |
| 1247.59000 | exclusion1828 |
| 1248.57000 | exclusion1829 |
| 1249.04000 | exclusion1830 |
| 1249.65000 | exclusion1258 |
| 1250.60000 | exclusion1259 |
| 1250.60000 | exclusion1831 |
| 1252.18000 | exclusion0786 |
| 1252.19000 | exclusion1832 |
| 1252.20000 | exclusion1260 |
| 1252.59000 | exclusion1833 |
| 1253.07000 | exclusion1834 |
| 1253.59000 | exclusion1835 |
| 1254.63000 | exclusion1836 |
| 1255.13000 | exclusion0787 |
| 1255.14000 | exclusion1261 |
| 1256.59000 | exclusion1262 |
| 1256.79000 | exclusion1837 |
| 1257.09000 | exclusion1263 |
| 1257.09000 | exclusion1838 |
| 1257.64000 | exclusion1264 |

| | |
|---|---|
| 1258.57000 | exclusion1839 |
| 1259.21000 | exclusion1840 |
| 1260.13000 | exclusion1265 |
| 1260.63000 | exclusion1841 |
| 1260.64000 | exclusion1266 |
| 1261.60000 | exclusion1267 |
| 1262.37000 | exclusion0788 |
| 1262.95000 | exclusion0789 |
| 1263.27000 | exclusion1268 |
| 1264.07000 | exclusion1269 |
| 1264.16000 | exclusion1842 |
| 1265.08000 | exclusion1843 |
| 1265.59000 | exclusion1844 |
| 1265.97000 | exclusion1845 |
| 1266.61000 | exclusion1270 |
| 1267.25000 | exclusion1271 |
| 1268.19000 | exclusion1272 |
| 1268.81000 | exclusion1273 |
| 1269.15000 | exclusion1274 |
| 1269.66000 | exclusion0790 |
| 1270.26000 | exclusion1275 |
| 1270.27000 | exclusion1846 |
| 1271.18000 | exclusion1847 |
| 1272.11000 | exclusion1276 |
| 1272.67000 | exclusion1277 |
| 1272.68000 | exclusion1848 |
| 1274.07000 | exclusion0791 |
| 1274.07000 | exclusion1278 |
| 1274.65000 | exclusion0792 |
| 1275.16000 | exclusion1279 |
| 1275.62000 | exclusion1280 |
| 1275.63000 | exclusion0793 |
| 1275.65000 | exclusion1849 |
| 1276.64000 | exclusion1850 |
| 1277.14000 | exclusion1281 |
| 1277.67000 | exclusion1282 |
| 1278.46000 | exclusion1283 |
| 1279.01000 | exclusion1284 |
| 1279.02000 | exclusion1851 |
| 1279.61000 | exclusion1852 |
| 1280.64000 | exclusion0794 |
| 1281.38000 | exclusion0795 |
| 1283.14000 | exclusion0796 |
| 1283.66000 | exclusion1285 |
| 1283.66000 | exclusion1853 |
| 1284.64000 | exclusion1854 |
| 1285.14000 | exclusion0797 |
| 1286.65000 | exclusion0798 |
| 1286.65000 | exclusion1286 |
| 1286.66000 | exclusion1855 |
| 1287.08000 | exclusion1856 |

| | |
|---|---|
| 1287.66000 | exclusion1287 |
| 1288.63000 | exclusion1288 |
| 1288.64000 | exclusion1857 |
| 1289.45000 | exclusion1858 |
| 1291.08000 | exclusion1859 |
| 1292.39000 | exclusion1860 |
| 1293.18000 | exclusion1861 |
| 1294.10000 | exclusion1289 |
| 1294.61000 | exclusion1862 |
| 1295.33000 | exclusion1863 |
| 1296.63000 | exclusion0799 |
| 1296.64000 | exclusion1290 |
| 1298.01000 | exclusion0800 |
| 1298.08000 | exclusion1864 |
| 1298.58000 | exclusion1291 |
| 1299.13000 | exclusion1292 |
| 1300.07000 | exclusion1865 |
| 1301.06000 | exclusion1866 |
| 1301.57000 | exclusion1293 |
| 1302.09000 | exclusion1867 |
| 1302.10000 | exclusion1294 |
| 1302.70000 | exclusion0801 |
| 1303.06000 | exclusion0802 |
| 1304.08000 | exclusion1295 |
| 1304.09000 | exclusion1868 |
| 1304.65000 | exclusion1296 |
| 1305.19000 | exclusion0803 |
| 1305.69000 | exclusion1297 |
| 1306.06000 | exclusion1298 |
| 1306.58000 | exclusion0804 |
| 1307.12000 | exclusion1869 |
| 1308.00000 | exclusion1299 |
| 1309.06000 | exclusion0805 |
| 1309.07000 | exclusion1300 |
| 1310.67000 | exclusion1301 |
| 1311.63000 | exclusion1302 |
| 1312.61000 | exclusion1870 |
| 1312.67000 | exclusion1303 |
| 1313.62000 | exclusion1304 |
| 1313.62000 | exclusion1871 |
| 1314.61000 | exclusion1305 |
| 1315.13000 | exclusion1306 |
| 1315.59000 | exclusion1872 |
| 1315.69000 | exclusion1307 |
| 1316.64000 | exclusion1308 |
| 1318.03000 | exclusion1309 |
| 1318.61000 | exclusion1310 |
| 1319.61000 | exclusion0806 |
| 1319.74000 | exclusion1873 |
| 1320.09000 | exclusion1311 |
| 1320.66000 | exclusion1874 |

| | |
|---|---|
| 1321.12000 | exclusion1312 |
| 1321.12000 | exclusion1875 |
| 1321.67000 | exclusion0807 |
| 1322.10000 | exclusion0808 |
| 1322.61000 | exclusion1313 |
| 1323.60000 | exclusion1314 |
| 1324.18000 | exclusion1876 |
| 1324.61000 | exclusion1877 |
| 1325.11000 | exclusion1315 |
| 1326.10000 | exclusion1316 |
| 1328.19000 | exclusion0809 |
| 1328.19000 | exclusion1317 |
| 1328.25000 | exclusion1878 |
| 1330.61000 | exclusion1318 |
| 1331.19000 | exclusion1319 |
| 1331.20000 | exclusion1879 |
| 1331.82000 | exclusion1320 |
| 1332.12000 | exclusion1321 |
| 1332.19000 | exclusion1880 |
| 1332.65000 | exclusion1322 |
| 1333.65000 | exclusion1323 |
| 1333.65000 | exclusion1881 |
| 1334.06000 | exclusion1324 |
| 1335.19000 | exclusion0810 |
| 1335.19000 | exclusion1325 |
| 1335.67000 | exclusion1326 |
| 1335.68000 | exclusion1882 |
| 1336.17000 | exclusion1327 |
| 1337.17000 | exclusion0811 |
| 1337.18000 | exclusion1883 |
| 1337.19000 | exclusion1328 |
| 1339.66000 | exclusion1329 |
| 1339.68000 | exclusion1884 |
| 1340.19000 | exclusion0812 |
| 1341.16000 | exclusion1330 |
| 1341.71000 | exclusion1885 |
| 1342.13000 | exclusion1331 |
| 1342.13000 | exclusion1886 |
| 1343.18000 | exclusion0813 |
| 1343.68000 | exclusion1332 |
| 1344.18000 | exclusion1887 |
| 1344.63000 | exclusion1333 |
| 1345.24000 | exclusion0814 |
| 1345.68000 | exclusion0815 |
| 1346.20000 | exclusion1888 |
| 1346.61000 | exclusion1889 |
| 1346.62000 | exclusion1334 |
| 1347.12000 | exclusion1335 |
| 1347.63000 | exclusion1890 |
| 1348.36000 | exclusion1336 |
| 1348.36000 | exclusion1891 |

| | |
|---|---|
| 1348.67000 | exclusion1337 |
| 1349.10000 | exclusion1338 |
| 1349.63000 | exclusion1339 |
| 1350.14000 | exclusion0816 |
| 1350.34000 | exclusion0817 |
| 1351.64000 | exclusion1340 |
| 1352.65000 | exclusion0818 |
| 1352.66000 | exclusion1341 |
| 1352.66000 | exclusion1892 |
| 1353.92000 | exclusion0819 |
| 1354.16000 | exclusion1342 |
| 1354.19000 | exclusion1893 |
| 1354.68000 | exclusion1343 |
| 1355.21000 | exclusion1344 |
| 1355.61000 | exclusion1345 |
| 1355.65000 | exclusion1894 |
| 1356.17000 | exclusion1346 |
| 1356.66000 | exclusion1347 |
| 1356.72000 | exclusion1895 |
| 1357.19000 | exclusion1896 |
| 1357.61000 | exclusion1348 |
| 1359.64000 | exclusion0820 |
| 1359.64000 | exclusion1349 |
| 1360.23000 | exclusion1350 |
| 1361.19000 | exclusion1897 |
| 1361.70000 | exclusion1351 |
| 1362.20000 | exclusion0821 |
| 1363.64000 | exclusion0822 |
| 1363.66000 | exclusion1352 |
| 1364.21000 | exclusion1353 |
| 1365.18000 | exclusion1898 |
| 1365.19000 | exclusion1354 |
| 1365.69000 | exclusion1355 |
| 1366.64000 | exclusion1356 |
| 1367.67000 | exclusion0823 |
| 1367.70000 | exclusion1357 |
| 1368.17000 | exclusion1358 |
| 1368.17000 | exclusion1899 |
| 1368.69000 | exclusion1900 |
| 1368.74000 | exclusion1359 |
| 1370.19000 | exclusion0824 |
| 1370.64000 | exclusion0825 |
| 1371.20000 | exclusion1360 |
| 1372.19000 | exclusion1361 |
| 1373.15000 | exclusion1362 |
| 1374.17000 | exclusion0826 |
| 1374.18000 | exclusion1363 |
| 1374.38000 | exclusion1901 |
| 1374.68000 | exclusion1902 |
| 1375.70000 | exclusion1364 |
| 1376.17000 | exclusion1365 |

| | |
|---|---|
| 1376.21000 | exclusion0827 |
| 1376.21000 | exclusion1903 |
| 1376.68000 | exclusion1366 |
| 1377.18000 | exclusion1904 |
| 1377.19000 | exclusion1367 |
| 1380.13000 | exclusion1368 |
| 1380.13000 | exclusion1905 |
| 1380.14000 | exclusion0828 |
| 1381.15000 | exclusion1369 |
| 1381.15000 | exclusion1906 |
| 1382.04000 | exclusion1370 |
| 1382.71000 | exclusion1907 |
| 1382.72000 | exclusion1371 |
| 1383.62000 | exclusion1372 |
| 1384.69000 | exclusion0829 |
| 1384.69000 | exclusion1373 |
| 1384.69000 | exclusion1908 |
| 1385.11000 | exclusion1909 |
| 1385.12000 | exclusion1374 |
| 1385.69000 | exclusion0830 |
| 1386.71000 | exclusion0831 |
| 1387.20000 | exclusion1375 |
| 1387.20000 | exclusion1910 |
| 1388.70000 | exclusion1376 |
| 1389.68000 | exclusion1377 |
| 1390.19000 | exclusion1911 |
| 1392.67000 | exclusion1378 |
| 1392.69000 | exclusion1912 |
| 1394.63000 | exclusion0832 |
| 1395.15000 | exclusion1379 |
| 1395.16000 | exclusion1913 |
| 1395.68000 | exclusion1380 |
| 1395.68000 | exclusion1914 |
| 1396.19000 | exclusion0833 |
| 1396.70000 | exclusion1381 |
| 1397.11000 | exclusion1915 |
| 1397.12000 | exclusion1382 |
| 1397.73000 | exclusion1383 |
| 1398.71000 | exclusion1916 |
| 1399.21000 | exclusion1384 |
| 1399.69000 | exclusion1917 |
| 1399.70000 | exclusion1385 |
| 1400.20000 | exclusion0834 |
| 1402.15000 | exclusion0835 |
| 1402.16000 | exclusion1386 |
| 1402.16000 | exclusion1918 |
| 1402.79000 | exclusion1387 |
| 1403.67000 | exclusion1388 |
| 1404.17000 | exclusion1919 |
| 1404.44000 | exclusion1389 |
| 1405.23000 | exclusion1390 |

| | |
|---|---|
| 1405.23000 | exclusion1920 |
| 1406.31000 | exclusion1921 |
| 1406.33000 | exclusion1391 |
| 1406.66000 | exclusion1392 |
| 1407.11000 | exclusion1393 |
| 1407.12000 | exclusion1922 |
| 1408.02000 | exclusion0836 |
| 1408.36000 | exclusion1394 |
| 1412.69000 | exclusion0837 |
| 1413.19000 | exclusion1923 |
| 1413.20000 | exclusion0838 |
| 1413.22000 | exclusion1395 |
| 1413.64000 | exclusion1396 |
| 1413.71000 | exclusion1924 |
| 1415.02000 | exclusion1397 |
| 1415.68000 | exclusion1398 |
| 1415.69000 | exclusion1925 |
| 1417.65000 | exclusion0839 |
| 1417.66000 | exclusion1399 |
| 1417.68000 | exclusion1926 |
| 1418.03000 | exclusion1400 |
| 1419.37000 | exclusion1401 |
| 1419.38000 | exclusion1927 |
| 1420.34000 | exclusion1402 |
| 1420.93000 | exclusion1928 |
| 1421.12000 | exclusion1403 |
| 1422.19000 | exclusion0840 |
| 1422.22000 | exclusion1929 |
| 1422.70000 | exclusion1404 |
| 1423.71000 | exclusion1405 |
| 1424.20000 | exclusion1406 |
| 1424.71000 | exclusion0841 |
| 1425.33000 | exclusion1407 |
| 1425.68000 | exclusion1408 |
| 1427.97000 | exclusion1409 |
| 1428.67000 | exclusion1410 |
| 1429.32000 | exclusion1930 |
| 1430.20000 | exclusion1411 |
| 1431.14000 | exclusion1931 |
| 1432.24000 | exclusion0842 |
| 1432.34000 | exclusion1412 |
| 1432.69000 | exclusion1413 |
| 1433.21000 | exclusion1414 |
| 1433.71000 | exclusion1932 |
| 1434.25000 | exclusion0843 |
| 1435.72000 | exclusion1415 |
| 1436.21000 | exclusion1416 |
| 1437.21000 | exclusion0844 |
| 1437.23000 | exclusion1417 |
| 1440.24000 | exclusion0845 |
| 1440.25000 | exclusion1418 |

| | |
|---|---|
| 1440.73000 | exclusion1933 |
| 1440.79000 | exclusion1419 |
| 1441.23000 | exclusion1420 |
| 1442.71000 | exclusion1421 |
| 1442.72000 | exclusion1934 |
| 1443.66000 | exclusion1422 |
| 1444.21000 | exclusion1423 |
| 1445.70000 | exclusion0846 |
| 1445.74000 | exclusion1424 |
| 1445.74000 | exclusion1935 |
| 1446.19000 | exclusion1425 |
| 1446.71000 | exclusion0847 |
| 1448.16000 | exclusion0848 |
| 1448.17000 | exclusion1426 |
| 1449.06000 | exclusion1427 |
| 1450.25000 | exclusion1936 |
| 1450.28000 | exclusion0849 |
| 1450.72000 | exclusion1428 |
| 1451.24000 | exclusion1937 |
| 1453.68000 | exclusion0850 |
| 1453.70000 | exclusion1429 |
| 1454.23000 | exclusion1938 |
| 1456.72000 | exclusion1430 |
| 1456.74000 | exclusion1939 |
| 1457.22000 | exclusion0851 |
| 1457.24000 | exclusion1940 |
| 1458.26000 | exclusion1431 |
| 1458.26000 | exclusion1941 |
| 1458.75000 | exclusion0852 |
| 1459.21000 | exclusion1432 |
| 1460.09000 | exclusion1433 |
| 1460.42000 | exclusion1942 |
| 1460.75000 | exclusion1943 |
| 1461.23000 | exclusion1434 |
| 1461.70000 | exclusion1435 |
| 1461.71000 | exclusion1944 |
| 1462.73000 | exclusion0853 |
| 1463.72000 | exclusion1436 |
| 1464.70000 | exclusion0854 |
| 1465.43000 | exclusion0855 |
| 1465.73000 | exclusion1945 |
| 1465.74000 | exclusion1437 |
| 1467.71000 | exclusion0856 |
| 1467.73000 | exclusion1946 |
| 1468.23000 | exclusion1438 |
| 1469.73000 | exclusion0857 |
| 1471.30000 | exclusion0858 |
| 1471.31000 | exclusion1947 |
| 1471.73000 | exclusion0859 |
| 1471.73000 | exclusion1439 |
| 1473.73000 | exclusion1440 |

| | |
|---|---|
| 1473.74000 | exclusion1948 |
| 1474.23000 | exclusion1441 |
| 1475.23000 | exclusion0860 |
| 1478.29000 | exclusion0861 |
| 1479.22000 | exclusion0862 |
| 1479.23000 | exclusion1442 |
| 1479.23000 | exclusion1949 |
| 1479.71000 | exclusion1443 |
| 1479.72000 | exclusion1950 |
| 1480.24000 | exclusion1444 |
| 1481.77000 | exclusion1445 |
| 1482.27000 | exclusion0863 |
| 1482.27000 | exclusion1951 |
| 1485.24000 | exclusion0864 |
| 1485.74000 | exclusion1446 |
| 1486.23000 | exclusion1952 |
| 1487.33000 | exclusion1447 |
| 1489.72000 | exclusion0865 |
| 1489.72000 | exclusion1448 |
| 1489.73000 | exclusion1953 |
| 1490.72000 | exclusion1954 |
| 1490.73000 | exclusion1449 |
| 1492.77000 | exclusion1450 |
| 1492.77000 | exclusion1955 |
| 1493.26000 | exclusion0866 |
| 1493.77000 | exclusion1451 |
| 1497.22000 | exclusion0867 |
| 1497.22000 | exclusion1956 |
| 1497.73000 | exclusion1452 |
| 1498.24000 | exclusion1957 |
| 1498.73000 | exclusion1453 |
| 1504.74000 | exclusion0868 |
| 1504.74000 | exclusion1454 |
| 1507.24000 | exclusion0869 |
| 1507.24000 | exclusion1455 |
| 1507.24000 | exclusion1958 |
| 1507.74000 | exclusion1456 |
| 1507.75000 | exclusion0870 |
| 1508.24000 | exclusion0871 |
| 1509.23000 | exclusion0872 |
| 1509.24000 | exclusion1457 |
| 1509.75000 | exclusion0873 |
| 1510.28000 | exclusion1959 |
| 1510.29000 | exclusion1458 |
| 1510.78000 | exclusion0874 |
| 1512.19000 | exclusion1459 |
| 1512.69000 | exclusion1960 |
| 1512.70000 | exclusion1460 |
| 1514.74000 | exclusion1961 |
| 1515.24000 | exclusion1461 |
| 1518.25000 | exclusion1462 |

| | |
|---|---|
| 1518.73000 | exclusion0875 |
| 1519.22000 | exclusion1463 |
| 1520.79000 | exclusion1464 |
| 1522.20000 | exclusion1465 |
| 1524.25000 | exclusion1466 |
| 1524.25000 | exclusion1962 |
| 1524.75000 | exclusion0876 |
| 1528.30000 | exclusion1467 |
| 1528.32000 | exclusion1963 |
| 1530.83000 | exclusion0877 |
| 1532.25000 | exclusion0878 |
| 1532.74000 | exclusion1468 |
| 1532.76000 | exclusion1964 |
| 1535.73000 | exclusion0879 |
| 1535.75000 | exclusion1469 |
| 1537.22000 | exclusion1470 |
| 1537.29000 | exclusion1965 |
| 1537.30000 | exclusion0880 |
| 1537.73000 | exclusion1471 |
| 1538.72000 | exclusion0881 |
| 1543.22000 | exclusion0882 |
| 1543.74000 | exclusion1472 |
| 1544.29000 | exclusion1966 |
| 1546.30000 | exclusion0883 |
| 1552.75000 | exclusion0884 |
| 1553.26000 | exclusion1473 |
| 1553.78000 | exclusion1474 |
| 1553.80000 | exclusion1967 |
| 1555.25000 | exclusion1475 |
| 1559.76000 | exclusion1476 |
| 1561.27000 | exclusion1477 |
| 1561.75000 | exclusion1968 |
| 1561.76000 | exclusion0885 |
| 1563.30000 | exclusion1969 |
| 1564.79000 | exclusion1478 |
| 1565.79000 | exclusion1479 |
| 1567.76000 | exclusion0886 |
| 1568.27000 | exclusion1480 |
| 1568.27000 | exclusion1970 |
| 1569.27000 | exclusion1481 |
| 1570.29000 | exclusion0887 |
| 1570.29000 | exclusion1482 |
| 1572.25000 | exclusion0888 |
| 1572.26000 | exclusion1483 |
| 1572.26000 | exclusion1971 |
| 1574.30000 | exclusion1972 |
| 1574.79000 | exclusion0889 |
| 1575.32000 | exclusion1973 |
| 1575.80000 | exclusion1484 |
| 1576.43000 | exclusion1485 |
| 1578.75000 | exclusion0890 |

| | |
|---|---|
| 1579.25000 | exclusion1486 |
| 1579.26000 | exclusion1974 |
| 1580.48000 | exclusion1487 |
| 1580.51000 | exclusion1975 |
| 1580.81000 | exclusion0891 |
| 1581.78000 | exclusion0892 |
| 1583.23000 | exclusion1488 |
| 1584.31000 | exclusion1489 |
| 1584.32000 | exclusion1976 |
| 1584.81000 | exclusion0893 |
| 1586.30000 | exclusion0894 |
| 1589.77000 | exclusion1490 |
| 1589.78000 | exclusion1977 |
| 1590.28000 | exclusion0895 |
| 1591.81000 | exclusion0896 |
| 1592.33000 | exclusion1491 |
| 1596.28000 | exclusion1492 |
| 1598.84000 | exclusion0897 |
| 1599.72000 | exclusion1493 |
| 1635.31000 | exclusion1978 |
| 1635.32000 | exclusion0898 |
| 1635.33000 | exclusion1494 |
| 1635.90000 | exclusion0899 |
| 1643.27000 | exclusion0900 |
| 1643.75000 | exclusion1495 |
| 1646.30000 | exclusion0901 |
| 1648.81000 | exclusion1496 |
| 1649.31000 | exclusion0902 |
| 1651.37000 | exclusion0903 |
| 1654.77000 | exclusion1497 |
| 1655.83000 | exclusion0904 |
| 1656.31000 | exclusion1498 |
| 1656.32000 | exclusion1979 |
| 1662.32000 | exclusion1499 |
| 1662.32000 | exclusion1980 |
| 1663.84000 | exclusion1500 |
| 1664.33000 | exclusion1501 |
| 1667.28000 | exclusion1981 |
| 1667.32000 | exclusion0905 |
| 1667.33000 | exclusion1502 |
| 1668.30000 | exclusion1503 |
| 1669.87000 | exclusion0906 |
| 1670.37000 | exclusion0907 |
| 1671.31000 | exclusion1504 |
| 1671.84000 | exclusion1982 |
| 1673.31000 | exclusion0908 |
| 1673.32000 | exclusion1983 |
| 1673.35000 | exclusion1505 |
| 1675.30000 | exclusion0909 |
| 1675.31000 | exclusion1984 |
| 1675.33000 | exclusion1506 |

| | |
|---|---|
| 1677.31000 | exclusion1507 |
| 1677.81000 | exclusion1508 |
| 1678.32000 | exclusion1985 |
| 1679.40000 | exclusion1509 |
| 1679.80000 | exclusion1986 |
| 1679.85000 | exclusion0910 |
| 1682.33000 | exclusion0911 |
| 1682.80000 | exclusion1510 |
| 1683.30000 | exclusion1511 |
| 1683.82000 | exclusion0912 |
| 1684.32000 | exclusion1512 |
| 1684.32000 | exclusion1987 |
| 1684.85000 | exclusion1988 |
| 1686.39000 | exclusion1513 |
| 1687.38000 | exclusion1989 |
| 1687.81000 | exclusion1514 |
| 1688.81000 | exclusion1515 |
| 1690.34000 | exclusion0913 |
| 1690.39000 | exclusion1516 |
| 1690.84000 | exclusion1990 |
| 1698.33000 | exclusion1517 |
| 1698.34000 | exclusion1991 |
| 1699.86000 | exclusion0914 |
| 1700.37000 | exclusion1518 |
| 1701.33000 | exclusion1992 |
| 1701.81000 | exclusion1519 |
| 1703.85000 | exclusion0915 |
| 1707.86000 | exclusion0916 |
| 1707.86000 | exclusion1520 |
| 1708.36000 | exclusion1993 |
| 1722.42000 | exclusion0917 |
| 1760.89000 | exclusion0918 |
| 1760.89000 | exclusion1521 |
| 1760.89000 | exclusion1994 |
| 1764.34000 | exclusion1522 |
| 1773.39000 | exclusion0919 |
| 1773.89000 | exclusion1523 |
| 1773.89000 | exclusion1995 |
| 1775.34000 | exclusion1524 |
| 1780.39000 | exclusion1525 |
| 1780.89000 | exclusion0920 |
| 1780.89000 | exclusion1996 |
| 1784.88000 | exclusion0921 |
| 1784.88000 | exclusion1997 |
| 1784.89000 | exclusion1526 |
| 1791.37000 | exclusion0922 |
| 1791.88000 | exclusion1527 |
| 1791.88000 | exclusion1998 |
| 1801.89000 | exclusion0923 |
| 1802.39000 | exclusion1528 |
| 1808.89000 | exclusion0924 |

1809.40000 exclusion1529
Neutral Loss Mass List: (none)
Product Mass List: (none)
Neutral loss in top: 3
Product in top: 3
Most intense if no parent masses found not enabled
Add/subtract mass not enabled
FT master scan preview mode enabled
Charge state screening enabled
Charge state dependent ETD time not enabled
Monoisotopic precursor selection enabled
Charge state rejection enabled
Unassigned charge states : rejected
Charge state 1 : rejected
Charge state 2 : not rejected
Charge state 3 : not rejected
Charge states 4+ : not rejected
Chromatography mode is disabled
Global Data Dependent Settings:
Predict ion injection time enabled
Use global parent and reject mass lists not enabled
Exclude parent mass from data dependent selection not enabled
Exclusion mass width by mass
Exclusion mass width low: 1.50000
Exclusion mass width high: 1.50000
Parent mass width by mass
Parent mass width low: 0.50000
Parent mass width high: 0.50000
Reject mass width by mass
Reject mass width low: 0.50000
Reject mass width high: 0.50000
Zoom/UltraZoom scan mass width by mass
Zoom/UltraZoom scan mass width low: 5.00
Zoom/UltraZoom scan mass width high: 5.00
FT SIM scan mass width low: 5.00
FT SIM scan mass width high: 5.00
Neutral Loss candidates processed by decreasing intensity
Neutral Loss mass width by mass
Neutral Loss mass width low: 0.50000
Neutral Loss mass width high: 0.50000
Product candidates processed by decreasing intensity
Product mass width by mass
Product mass width low: 0.50000
Product mass width high: 0.50000
MS mass range: 300.00-2000.00
MSn mass range by mass
MSn mass range: 0.00-1000000.00
Use m/z values as masses not enabled
Analog UV data dep. not enabled
Dynamic exclusion enabled
Repeat Count: 1

Repeat Duration: 30.00
Exclusion List Size: 500
Exclusion Duration: 180.00
Exclusion mass width by mass
Exclusion mass width low: 1.50000
Exclusion mass width high: 1.50000
Expiration: disabled
Isotopic data dependence not enabled
Custom Data Dependent Settings:
Not enabled

---

Pass 5 (wheat-mixed-digests_MS2_pass05_1.raw):

Creator: Orbi_30393
Last modified: 10/13/2021 by Orbi_30393
MS Run Time (min): 43.00
Sequence override of method parameters not enabled.
Divert Valve: not used during run
Contact Closure: not used during run
Syringe Pump: not used during run
MS Detector Settings:
Real-time modifications to method not enabled
Stepped collision energy not enabled
Additional Microscans:
MS2 0 0
MS3 0 0
MS4 0 0
MS5 0 0
MS6 0 0
MS7 0 0
MS8 0 0
MS9 0 0
MS10 0 0
Experiment Type: Nth Order Double Play
Tune Method: Orbitrap-tune-file_2020-03-13_HESI
Scan Event Details:
1: FTMS + p norm o(300.0-2000.0)
CV = 0.0V
2: ITMS + c norm Dep MS/MS Most intense ion from (1)
Activation Type: CID
Min. Signal Required: 500.0
Isolation Width: 2.00
Normalized Coll. Energy: 35.0
Default Charge State: 2
Activation Q: 0.250
Activation Time: 10.000
CV = 0.0V
Scan Event 2 repeated for top 10 peaks.
Lock Masses:
Pos List Name: N/A
Source: API Source

Mass List: (none)
Neg List Name: N/A
Source: API Source
Mass List: (none)
Data Dependent Settings:
Use separate polarity settings disabled
Parent Mass List: (none)
Reject Mass List: (none)
Neutral Loss Mass List: (none)
Product Mass List: (none)
Neutral loss in top: 3
Product in top: 3
Most intense if no parent masses found not enabled
Add/subtract mass not enabled
FT master scan preview mode enabled
Charge state screening enabled
Charge state dependent ETD time not enabled
Monoisotopic precursor selection enabled
Charge state rejection enabled
Unassigned charge states : rejected
Charge state 1 : rejected
Charge state 2 : not rejected
Charge state 3 : not rejected
Charge states 4+ : not rejected
Chromatography mode is disabled
Global Data Dependent Settings:
Predict ion injection time enabled
Use global parent and reject mass lists not enabled
Exclude parent mass from data dependent selection not enabled
Exclusion mass width by mass
Exclusion mass width low: 1.50000
Exclusion mass width high: 1.50000
Parent mass width by mass
Parent mass width low: 0.50000
Parent mass width high: 0.50000
Reject mass width by mass
Reject mass width low: 0.50000
Reject mass width high: 0.50000
Zoom/UltraZoom scan mass width by mass
Zoom/UltraZoom scan mass width low: 5.00
Zoom/UltraZoom scan mass width high: 5.00
FT SIM scan mass width low: 5.00
FT SIM scan mass width high: 5.00
Neutral Loss candidates processed by decreasing intensity
Neutral Loss mass width by mass
Neutral Loss mass width low: 0.50000
Neutral Loss mass width high: 0.50000
Product candidates processed by decreasing intensity
Product mass width by mass
Product mass width low: 0.50000
Product mass width high: 0.50000

MS mass range: 300.00-2000.00
MSn mass range by mass
MSn mass range: 0.00-1000000.00
Use m/z values as masses not enabled
Analog UV data dep. not enabled
Dynamic exclusion enabled
Repeat Count: 1
Repeat Duration: 30.00
Exclusion List Size: 500
Exclusion Duration: 180.00
Exclusion mass width by mass
Exclusion mass width low: 1.50000
Exclusion mass width high: 1.50000
Expiration: disabled
Isotopic data dependence not enabled
Custom Data Dependent Settings:
Not enabled

---

Pass 6 (wheat-mixed-digests_MS2_pass06_1.raw):

Creator: Orbi_30393
Last modified: 10/13/2021 by Orbi_30393
MS Run Time (min): 43.00
Sequence override of method parameters not enabled.
Divert Valve: not used during run
Contact Closure: not used during run
Syringe Pump: not used during run
MS Detector Settings:
Real-time modifications to method not enabled
Stepped collision energy not enabled
Additional Microscans:
MS2 0 0
MS3 0 0
MS4 0 0
MS5 0 0
MS6 0 0
MS7 0 0
MS8 0 0
MS9 0 0
MS10 0 0
Experiment Type: Nth Order Double Play
Tune Method: Orbitrap-tune-file_2020-03-13_HESI
Scan Event Details:
1: FTMS + p norm o(300.0-2000.0)
CV = 0.0V
2: ITMS + c norm Dep MS/MS Most intense ion from (1)
Activation Type: CID
Min. Signal Required: 3000.0
Isolation Width: 2.00
Normalized Coll. Energy: 35.0
Default Charge State: 2

Activation Q: 0.250
Activation Time: 10.000
CV = 0.0V
Scan Event 2 repeated for top 10 peaks.
Lock Masses:
Pos List Name: N/A
Source: API Source
Mass List: (none)
Neg List Name: N/A
Source: API Source
Mass List: (none)
Data Dependent Settings:
Use separate polarity settings disabled
Parent Mass List:

| MS Mass | MS FAIMS CV | MS Normalized Collision Energy | MS2 Mass | MS2 Normalized Collision Energy | Name |
|---|---|---|---|---|---|
| 300.18139 | | 35.0 | | | Cluster_18939 |
| 302.17822 | | 35.0 | | | Cluster_18998 |
| 304.10723 | | 35.0 | | | Cluster_19042 |
| 305.50412 | | 35.0 | | | Cluster_19081 |
| 307.12252 | | 35.0 | | | Cluster_19119 |
| 307.17583 | | 35.0 | | | Cluster_19131 |
| 307.17623 | | 35.0 | | | Cluster_19118 |
| 307.18423 | | 35.0 | | | Cluster_19126 |
| 307.50589 | | 35.0 | | | Cluster_06882 |
| 307.71572 | | 35.0 | | | Cluster_19142 |
| 307.84574 | | 35.0 | | | Cluster_06883 |
| 308.14269 | | 35.0 | | | Cluster_06884 |
| 308.14312 | | 35.0 | | | Cluster_06885 |
| 308.14934 | | 35.0 | | | Cluster_19160 |
| 308.64632 | | 35.0 | | | Cluster_19177 |
| 309.10936 | | 35.0 | | | Cluster_19188 |
| 309.67276 | | 35.0 | | | Cluster_19196 |
| 310.66456 | | 35.0 | | | Cluster_19221 |
| 310.86504 | | 35.0 | | | Cluster_19230 |
| 311.10568 | | 35.0 | | | Cluster_19240 |
| 312.17580 | | 35.0 | | | Cluster_19273 |
| 312.29236 | | 35.0 | | | Cluster_19281 |
| 312.52450 | | 35.0 | | | Cluster_19284 |
| 313.17441 | | 35.0 | | | Cluster_19302 |
| 313.50880 | | 35.0 | | | Cluster_19312 |
| 313.94655 | | 35.0 | | | Cluster_06933 |
| 314.11405 | | 35.0 | | | Cluster_19325 |
| 314.52501 | | 35.0 | | | Cluster_06938 |
| 314.85813 | | 35.0 | | | Cluster_06945 |
| 314.90987 | | 35.0 | | | Cluster_19353 |
| 315.17966 | | 35.0 | | | Cluster_19370 |
| 315.18040 | | 35.0 | | | Cluster_19367 |
| 315.18043 | | 35.0 | | | Cluster_19368 |

| | | |
|---|---|---|
| 315.71422 | 35.0 | Cluster_19400 |
| 318.85827 | 35.0 | Cluster_06976 |
| 319.21104 | 35.0 | Cluster_19555 |
| 319.68517 | 35.0 | Cluster_19563 |
| 319.68541 | 35.0 | Cluster_00847 |
| 319.68602 | 35.0 | Cluster_00844 |
| 319.68635 | 35.0 | Cluster_00845 |
| 320.91801 | 35.0 | Cluster_19585 |
| 320.91883 | 35.0 | Cluster_00861 |
| 320.91896 | 35.0 | Cluster_00856 |
| 320.91918 | 35.0 | Cluster_00852 |
| 321.15308 | 35.0 | Cluster_06995 |
| 321.61792 | 35.0 | Cluster_19601 |
| 321.72018 | 35.0 | Cluster_19607 |
| 322.53614 | 35.0 | Cluster_19633 |
| 323.19190 | 35.0 | Cluster_19656 |
| 323.65030 | 35.0 | Cluster_19672 |
| 323.68024 | 35.0 | Cluster_19667 |
| 324.13156 | 35.0 | Cluster_19688 |
| 324.42069 | 35.0 | Cluster_00878 |
| 324.60316 | 35.0 | Cluster_19711 |
| 324.66388 | 35.0 | Cluster_19720 |
| 324.66471 | 35.0 | Cluster_19719 |
| 325.15298 | 35.0 | Cluster_19731 |
| 325.19669 | 35.0 | Cluster_19741 |
| 326.61613 | 35.0 | Cluster_19767 |
| 327.17150 | 35.0 | Cluster_19785 |
| 327.44838 | 35.0 | Cluster_19797 |
| 328.20600 | 35.0 | Cluster_19817 |
| 329.00657 | 35.0 | Cluster_19842 |
| 329.82193 | 35.0 | Cluster_07063 |
| 330.10925 | 35.0 | Cluster_19877 |
| 330.19915 | 35.0 | Cluster_19884 |
| 330.90022 | 35.0 | Cluster_19916 |
| 331.20771 | 35.0 | Cluster_19930 |
| 331.22467 | 35.0 | Cluster_19925 |
| 331.91322 | 35.0 | Cluster_19943 |
| 331.92626 | 35.0 | Cluster_00898 |
| 333.14933 | 35.0 | Cluster_19981 |
| 333.86488 | 35.0 | Cluster_07093 |
| 333.87455 | 35.0 | Cluster_07090 |
| 334.51482 | 35.0 | Cluster_20027 |
| 335.39009 | 35.0 | Cluster_00911 |
| 335.42425 | 35.0 | Cluster_20059 |
| 335.90453 | 35.0 | Cluster_20079 |
| 336.18771 | 35.0 | Cluster_07109 |
| 336.21532 | 35.0 | Cluster_20094 |
| 336.46579 | 35.0 | Cluster_20097 |
| 336.70926 | 35.0 | Cluster_20102 |
| 337.19049 | 35.0 | Cluster_20132 |
| 338.68639 | 35.0 | Cluster_20181 |

| | | |
|---|---|---|
| 338.93601 | 35.0 | Cluster_20187 |
| 339.28949 | 35.0 | Cluster_20198 |
| 339.49537 | 35.0 | Cluster_07140 |
| 340.12729 | 35.0 | Cluster_07154 |
| 341.18643 | 35.0 | Cluster_20258 |
| 342.20057 | 35.0 | Cluster_07171 |
| 342.22830 | 35.0 | Cluster_20300 |
| 343.66746 | 35.0 | Cluster_20336 |
| 344.15675 | 35.0 | Cluster_20368 |
| 344.72229 | 35.0 | Cluster_20384 |
| 345.15297 | 35.0 | Cluster_07192 |
| 345.91798 | 35.0 | Cluster_20415 |
| 346.71073 | 35.0 | Cluster_20448 |
| 348.04600 | 35.0 | Cluster_20480 |
| 348.64686 | 35.0 | Cluster_00973 |
| 348.69149 | 35.0 | Cluster_20506 |
| 349.24306 | 35.0 | Cluster_07239 |
| 349.63790 | 35.0 | Cluster_20543 |
| 350.11421 | 35.0 | Cluster_20551 |
| 350.15978 | 35.0 | Cluster_20553 |
| 351.23268 | 35.0 | Cluster_20627 |
| 351.44817 | 35.0 | Cluster_07249 |
| 351.70979 | 35.0 | Cluster_20644 |
| 352.17409 | 35.0 | Cluster_07256 |
| 352.65949 | 35.0 | Cluster_20669 |
| 352.68040 | 35.0 | Cluster_20674 |
| 354.69171 | 35.0 | Cluster_20769 |
| 356.18535 | 35.0 | Cluster_20822 |
| 356.21147 | 35.0 | Cluster_20827 |
| 356.21666 | 35.0 | Cluster_20830 |
| 356.54633 | 35.0 | Cluster_07307 |
| 356.64829 | 35.0 | Cluster_20835 |
| 356.86733 | 35.0 | Cluster_07310 |
| 357.15015 | 35.0 | Cluster_07312 |
| 357.63894 | 35.0 | Cluster_20878 |
| 357.81560 | 35.0 | Cluster_07316 |
| 357.83203 | 35.0 | Cluster_20894 |
| 359.46536 | 35.0 | Cluster_01042 |
| 359.87192 | 35.0 | Cluster_07329 |
| 360.20680 | 35.0 | Cluster_20980 |
| 360.53233 | 35.0 | Cluster_07338 |
| 360.91407 | 35.0 | Cluster_21012 |
| 361.42083 | 35.0 | Cluster_01055 |
| 361.90202 | 35.0 | Cluster_07352 |
| 362.22641 | 35.0 | Cluster_21071 |
| 362.69593 | 35.0 | Cluster_21078 |
| 363.10528 | 35.0 | Cluster_21089 |
| 363.17084 | 35.0 | Cluster_21090 |
| 363.18642 | 35.0 | Cluster_21093 |
| 363.24378 | 35.0 | Cluster_21098 |
| 364.15928 | 35.0 | Cluster_21135 |

| | | |
|---|---|---|
| 364.53487 | 35.0 | Cluster_07386 |
| 365.21954 | 35.0 | Cluster_21200 |
| 365.74467 | 35.0 | Cluster_21233 |
| 365.86270 | 35.0 | Cluster_07398 |
| 366.66719 | 35.0 | Cluster_21282 |
| 367.19893 | 35.0 | Cluster_21312 |
| 367.23906 | 35.0 | Cluster_07411 |
| 367.54820 | 35.0 | Cluster_21323 |
| 367.67913 | 35.0 | Cluster_21329 |
| 368.18275 | 35.0 | Cluster_21363 |
| 368.19973 | 35.0 | Cluster_21366 |
| 368.20426 | 35.0 | Cluster_21358 |
| 368.20533 | 35.0 | Cluster_21369 |
| 368.35175 | 35.0 | Cluster_21372 |
| 368.66636 | 35.0 | Cluster_21380 |
| 368.68342 | 35.0 | Cluster_21381 |
| 368.69841 | 35.0 | Cluster_21379 |
| 369.18125 | 35.0 | Cluster_21406 |
| 369.18336 | 35.0 | Cluster_21407 |
| 369.68028 | 35.0 | Cluster_21424 |
| 369.68403 | 35.0 | Cluster_21443 |
| 370.08367 | 35.0 | Cluster_21453 |
| 370.19785 | 35.0 | Cluster_21468 |
| 370.25193 | 35.0 | Cluster_21469 |
| 370.54002 | 35.0 | Cluster_21470 |
| 370.73721 | 35.0 | Cluster_21476 |
| 371.17419 | 35.0 | Cluster_21493 |
| 371.27007 | 35.0 | Cluster_21509 |
| 371.45807 | 35.0 | Cluster_21511 |
| 371.60677 | 35.0 | Cluster_21514 |
| 372.22058 | 35.0 | Cluster_21545 |
| 372.67345 | 35.0 | Cluster_21575 |
| 372.70722 | 35.0 | Cluster_21577 |
| 373.21517 | 35.0 | Cluster_21597 |
| 373.21690 | 35.0 | Cluster_21590 |
| 374.70173 | 35.0 | Cluster_21660 |
| 374.72356 | 35.0 | Cluster_21659 |
| 375.17267 | 35.0 | Cluster_21676 |
| 376.18825 | 35.0 | Cluster_07499 |
| 376.19398 | 35.0 | Cluster_21713 |
| 376.24488 | 35.0 | Cluster_21722 |
| 377.20229 | 35.0 | Cluster_21745 |
| 377.23336 | 35.0 | Cluster_21758 |
| 377.69338 | 35.0 | Cluster_21776 |
| 377.71834 | 35.0 | Cluster_21774 |
| 378.18100 | 35.0 | Cluster_21788 |
| 380.45221 | 35.0 | Cluster_21886 |
| 380.72076 | 35.0 | Cluster_21888 |
| 381.23578 | 35.0 | Cluster_21923 |
| 381.72765 | 35.0 | Cluster_21933 |
| 382.19501 | 35.0 | Cluster_07559 |

| | | |
|---|---|---|
| 382.19970 | 35.0 | Cluster_21959 |
| 382.73846 | 35.0 | Cluster_21985 |
| 383.16199 | 35.0 | Cluster_07572 |
| 383.70026 | 35.0 | Cluster_22026 |
| 383.89718 | 35.0 | Cluster_07580 |
| 384.20618 | 35.0 | Cluster_22055 |
| 384.20882 | 35.0 | Cluster_22063 |
| 384.22734 | 35.0 | Cluster_22048 |
| 384.23699 | 35.0 | Cluster_22058 |
| 384.23964 | 35.0 | Cluster_22056 |
| 384.95294 | 35.0 | Cluster_22080 |
| 386.22103 | 35.0 | Cluster_22144 |
| 386.36180 | 35.0 | Cluster_22148 |
| 387.15735 | 35.0 | Cluster_22178 |
| 387.92865 | 35.0 | Cluster_22226 |
| 388.21127 | 35.0 | Cluster_22239 |
| 390.07893 | 35.0 | Cluster_22287 |
| 391.83529 | 35.0 | Cluster_07662 |
| 391.88959 | 35.0 | Cluster_07660 |
| 392.19049 | 35.0 | Cluster_01189 |
| 393.16093 | 35.0 | Cluster_22393 |
| 393.35115 | 35.0 | Cluster_22422 |
| 393.35150 | 35.0 | Cluster_22418 |
| 394.12425 | 35.0 | Cluster_22482 |
| 394.23632 | 35.0 | Cluster_22487 |
| 395.18490 | 35.0 | Cluster_07695 |
| 395.22912 | 35.0 | Cluster_22537 |
| 395.23140 | 35.0 | Cluster_22535 |
| 396.19889 | 35.0 | Cluster_07708 |
| 396.67333 | 35.0 | Cluster_22609 |
| 396.88504 | 35.0 | Cluster_22610 |
| 396.90372 | 35.0 | Cluster_01219 |
| 397.21106 | 35.0 | Cluster_07728 |
| 397.21183 | 35.0 | Cluster_07721 |
| 397.21517 | 35.0 | Cluster_07729 |
| 397.21518 | 35.0 | Cluster_07727 |
| 398.54590 | 35.0 | Cluster_07741 |
| 400.68518 | 35.0 | Cluster_22777 |
| 400.73920 | 35.0 | Cluster_22787 |
| 400.76739 | 35.0 | Cluster_22792 |
| 401.12569 | 35.0 | Cluster_22796 |
| 402.47822 | 35.0 | Cluster_22868 |
| 403.21431 | 35.0 | Cluster_22897 |
| 403.22681 | 35.0 | Cluster_22898 |
| 403.47621 | 35.0 | Cluster_22910 |
| 403.50556 | 35.0 | Cluster_01262 |
| 403.54121 | 35.0 | Cluster_07797 |
| 403.71289 | 35.0 | Cluster_22924 |
| 405.20527 | 35.0 | Cluster_22990 |
| 405.95533 | 35.0 | Cluster_01279 |
| 406.19169 | 35.0 | Cluster_23017 |

| | | |
|---|---|---|
| 406.25032 | 35.0 | Cluster_23028 |
| 406.25211 | 35.0 | Cluster_23025 |
| 407.19440 | 35.0 | Cluster_01299 |
| 407.23778 | 35.0 | Cluster_23063 |
| 407.69970 | 35.0 | Cluster_23073 |
| 408.23545 | 35.0 | Cluster_07858 |
| 408.54132 | 35.0 | Cluster_07863 |
| 409.12920 | 35.0 | Cluster_23141 |
| 409.13151 | 35.0 | Cluster_23142 |
| 409.21581 | 35.0 | Cluster_07870 |
| 409.22568 | 35.0 | Cluster_23153 |
| 409.43498 | 35.0 | Cluster_01310 |
| 409.57292 | 35.0 | Cluster_23167 |
| 409.73324 | 35.0 | Cluster_23180 |
| 410.21306 | 35.0 | Cluster_23192 |
| 410.43506 | 35.0 | Cluster_23203 |
| 410.69545 | 35.0 | Cluster_23208 |
| 410.88152 | 35.0 | Cluster_23217 |
| 411.29757 | 35.0 | Cluster_23239 |
| 411.46011 | 35.0 | Cluster_01319 |
| 411.46179 | 35.0 | Cluster_23246 |
| 411.87723 | 35.0 | Cluster_07897 |
| 412.22721 | 35.0 | Cluster_23277 |
| 412.36399 | 35.0 | Cluster_07901 |
| 412.44074 | 35.0 | Cluster_23284 |
| 412.99218 | 35.0 | Cluster_23308 |
| 413.16510 | 35.0 | Cluster_01330 |
| 413.42901 | 35.0 | Cluster_23332 |
| 413.70849 | 35.0 | Cluster_01337 |
| 413.72640 | 35.0 | Cluster_23344 |
| 413.76861 | 35.0 | Cluster_23346 |
| 414.75386 | 35.0 | Cluster_23381 |
| 415.19319 | 35.0 | Cluster_23407 |
| 416.76401 | 35.0 | Cluster_23476 |
| 416.96144 | 35.0 | Cluster_23484 |
| 417.24118 | 35.0 | Cluster_23495 |
| 417.70707 | 35.0 | Cluster_23504 |
| 417.93206 | 35.0 | Cluster_07965 |
| 418.27534 | 35.0 | Cluster_23531 |
| 418.68587 | 35.0 | Cluster_23550 |
| 418.73063 | 35.0 | Cluster_23554 |
| 419.16928 | 35.0 | Cluster_23561 |
| 420.19295 | 35.0 | Cluster_23617 |
| 420.48611 | 35.0 | Cluster_23629 |
| 420.99622 | 35.0 | Cluster_23649 |
| 421.17949 | 35.0 | Cluster_07996 |
| 421.25549 | 35.0 | Cluster_23656 |
| 421.74177 | 35.0 | Cluster_23691 |
| 422.24188 | 35.0 | Cluster_08011 |
| 422.75222 | 35.0 | Cluster_23720 |
| 423.24596 | 35.0 | Cluster_23732 |

| | | |
|---|---|---|
| 423.25021 | 35.0 | Cluster_23734 |
| 423.26452 | 35.0 | Cluster_23744 |
| 423.71445 | 35.0 | Cluster_23753 |
| 423.71797 | 35.0 | Cluster_23757 |
| 423.86317 | 35.0 | Cluster_08030 |
| 424.21310 | 35.0 | Cluster_23791 |
| 424.24130 | 35.0 | Cluster_08033 |
| 424.71578 | 35.0 | Cluster_23830 |
| 425.21353 | 35.0 | Cluster_23847 |
| 425.22563 | 35.0 | Cluster_23845 |
| 425.23397 | 35.0 | Cluster_08046 |
| 425.75390 | 35.0 | Cluster_23890 |
| 425.75537 | 35.0 | Cluster_23891 |
| 425.86680 | 35.0 | Cluster_08056 |
| 426.21748 | 35.0 | Cluster_23928 |
| 426.25210 | 35.0 | Cluster_23935 |
| 426.85610 | 35.0 | Cluster_23956 |
| 426.96516 | 35.0 | Cluster_23959 |
| 427.45165 | 35.0 | Cluster_01418 |
| 427.73954 | 35.0 | Cluster_23990 |
| 427.77055 | 35.0 | Cluster_23996 |
| 427.77629 | 35.0 | Cluster_23986 |
| 428.20390 | 35.0 | Cluster_01423 |
| 428.26325 | 35.0 | Cluster_24023 |
| 428.27627 | 35.0 | Cluster_24030 |
| 428.73429 | 35.0 | Cluster_24052 |
| 429.17967 | 35.0 | Cluster_24066 |
| 429.56584 | 35.0 | Cluster_08120 |
| 430.22949 | 35.0 | Cluster_08134 |
| 431.27472 | 35.0 | Cluster_24169 |
| 431.55978 | 35.0 | Cluster_08150 |
| 432.46095 | 35.0 | Cluster_24230 |
| 433.16955 | 35.0 | Cluster_24257 |
| 433.24369 | 35.0 | Cluster_24263 |
| 433.28448 | 35.0 | Cluster_24266 |
| 434.76872 | 35.0 | Cluster_24329 |
| 435.22149 | 35.0 | Cluster_24369 |
| 435.22203 | 35.0 | Cluster_24359 |
| 435.90332 | 35.0 | Cluster_08208 |
| 436.27565 | 35.0 | Cluster_24432 |
| 436.47569 | 35.0 | Cluster_01491 |
| 436.56693 | 35.0 | Cluster_24436 |
| 436.56866 | 35.0 | Cluster_01492 |
| 436.72309 | 35.0 | Cluster_24448 |
| 436.77680 | 35.0 | Cluster_24452 |
| 436.78365 | 35.0 | Cluster_24453 |
| 437.24974 | 35.0 | Cluster_08221 |
| 438.52285 | 35.0 | Cluster_24544 |
| 438.73222 | 35.0 | Cluster_24563 |
| 438.90950 | 35.0 | Cluster_24583 |
| 439.25435 | 35.0 | Cluster_24589 |

| | | |
|---|---|---|
| 439.25668 | 35.0 | Cluster_24587 |
| 439.76705 | 35.0 | Cluster_24627 |
| 439.90281 | 35.0 | Cluster_08267 |
| 440.21796 | 35.0 | Cluster_24645 |
| 440.73012 | 35.0 | Cluster_24657 |
| 440.95090 | 35.0 | Cluster_24671 |
| 441.06270 | 35.0 | Cluster_24673 |
| 441.14804 | 35.0 | Cluster_08284 |
| 441.19838 | 35.0 | Cluster_08283 |
| 441.65909 | 35.0 | Cluster_24701 |
| 442.22077 | 35.0 | Cluster_08299 |
| 442.48368 | 35.0 | Cluster_24747 |
| 442.73721 | 35.0 | Cluster_24752 |
| 442.89667 | 35.0 | Cluster_24764 |
| 443.22990 | 35.0 | Cluster_01524 |
| 443.29019 | 35.0 | Cluster_24795 |
| 443.72248 | 35.0 | Cluster_01529 |
| 444.56992 | 35.0 | Cluster_24864 |
| 444.62609 | 35.0 | Cluster_24865 |
| 444.80902 | 35.0 | Cluster_24880 |
| 444.98546 | 35.0 | Cluster_24883 |
| 444.98560 | 35.0 | Cluster_24882 |
| 444.98721 | 35.0 | Cluster_01544 |
| 445.55386 | 35.0 | Cluster_24900 |
| 445.66341 | 35.0 | Cluster_01549 |
| 445.70794 | 35.0 | Cluster_24905 |
| 445.72605 | 35.0 | Cluster_24909 |
| 446.30454 | 35.0 | Cluster_24942 |
| 446.55756 | 35.0 | Cluster_24946 |
| 446.74957 | 35.0 | Cluster_01557 |
| 447.19331 | 35.0 | Cluster_24975 |
| 447.29041 | 35.0 | Cluster_24988 |
| 447.29684 | 35.0 | Cluster_24980 |
| 447.47895 | 35.0 | Cluster_24992 |
| 447.76941 | 35.0 | Cluster_25002 |
| 448.21671 | 35.0 | Cluster_08397 |
| 448.23477 | 35.0 | Cluster_25021 |
| 448.57791 | 35.0 | Cluster_08400 |
| 449.27084 | 35.0 | Cluster_25066 |
| 449.95854 | 35.0 | Cluster_25103 |
| 450.27739 | 35.0 | Cluster_25114 |
| 450.56658 | 35.0 | Cluster_25130 |
| 451.77488 | 35.0 | Cluster_25180 |
| 452.01254 | 35.0 | Cluster_25192 |
| 452.75396 | 35.0 | Cluster_25235 |
| 453.70221 | 35.0 | Cluster_01604 |
| 454.19131 | 35.0 | Cluster_08500 |
| 454.58119 | 35.0 | Cluster_08511 |
| 454.70208 | 35.0 | Cluster_25319 |
| 455.20057 | 35.0 | Cluster_08518 |
| 455.47836 | 35.0 | Cluster_01621 |

| | | |
|---|---|---|
| 455.81190 | 35.0 | Cluster_25364 |
| 455.92600 | 35.0 | Cluster_08527 |
| 456.79120 | 35.0 | Cluster_25411 |
| 457.23466 | 35.0 | Cluster_25428 |
| 457.23627 | 35.0 | Cluster_25426 |
| 457.26552 | 35.0 | Cluster_25430 |
| 457.95445 | 35.0 | Cluster_08555 |
| 458.78823 | 35.0 | Cluster_25504 |
| 458.79020 | 35.0 | Cluster_25505 |
| 459.99743 | 35.0 | Cluster_25551 |
| 461.16305 | 35.0 | Cluster_08588 |
| 461.24082 | 35.0 | Cluster_08590 |
| 461.26795 | 35.0 | Cluster_25609 |
| 461.26968 | 35.0 | Cluster_25607 |
| 461.45638 | 35.0 | Cluster_25615 |
| 461.55248 | 35.0 | Cluster_25617 |
| 461.57128 | 35.0 | Cluster_08594 |
| 461.67636 | 35.0 | Cluster_25619 |
| 461.72794 | 35.0 | Cluster_25620 |
| 461.92659 | 35.0 | Cluster_08602 |
| 462.24726 | 35.0 | Cluster_25635 |
| 462.52195 | 35.0 | Cluster_08614 |
| 462.71081 | 35.0 | Cluster_25647 |
| 462.71773 | 35.0 | Cluster_25648 |
| 463.22978 | 35.0 | Cluster_25667 |
| 463.28631 | 35.0 | Cluster_25669 |
| 463.51536 | 35.0 | Cluster_01676 |
| 463.74334 | 35.0 | Cluster_25680 |
| 463.98229 | 35.0 | Cluster_25697 |
| 464.22084 | 35.0 | Cluster_08632 |
| 464.97394 | 35.0 | Cluster_25759 |
| 464.97417 | 35.0 | Cluster_25764 |
| 464.97567 | 35.0 | Cluster_25761 |
| 465.24645 | 35.0 | Cluster_25784 |
| 465.27096 | 35.0 | Cluster_25773 |
| 465.27897 | 35.0 | Cluster_25781 |
| 466.75077 | 35.0 | Cluster_25838 |
| 466.90772 | 35.0 | Cluster_25848 |
| 467.57948 | 35.0 | Cluster_08687 |
| 468.74197 | 35.0 | Cluster_25932 |
| 468.75878 | 35.0 | Cluster_25925 |
| 468.93506 | 35.0 | Cluster_08710 |
| 469.20920 | 35.0 | Cluster_08717 |
| 469.24421 | 35.0 | Cluster_25946 |
| 469.25453 | 35.0 | Cluster_08719 |
| 469.25778 | 35.0 | Cluster_08720 |
| 469.48106 | 35.0 | Cluster_01715 |
| 469.78252 | 35.0 | Cluster_25970 |
| 470.23518 | 35.0 | Cluster_25990 |
| 470.23714 | 35.0 | Cluster_25992 |
| 470.24168 | 35.0 | Cluster_08729 |

| | | |
|---|---|---|
| 470.31187 | 35.0 | Cluster_25999 |
| 470.73989 | 35.0 | Cluster_01726 |
| 470.95744 | 35.0 | Cluster_26025 |
| 471.25377 | 35.0 | Cluster_01737 |
| 471.78022 | 35.0 | Cluster_26056 |
| 471.85656 | 35.0 | Cluster_26069 |
| 472.28933 | 35.0 | Cluster_26090 |
| 473.00613 | 35.0 | Cluster_26120 |
| 473.25522 | 35.0 | Cluster_26141 |
| 473.26784 | 35.0 | Cluster_26146 |
| 473.50841 | 35.0 | Cluster_26151 |
| 473.61584 | 35.0 | Cluster_08782 |
| 474.25392 | 35.0 | Cluster_01752 |
| 474.74844 | 35.0 | Cluster_26188 |
| 474.82517 | 35.0 | Cluster_26201 |
| 475.20902 | 35.0 | Cluster_26205 |
| 475.25075 | 35.0 | Cluster_26223 |
| 475.91606 | 35.0 | Cluster_08825 |
| 476.77033 | 35.0 | Cluster_26267 |
| 477.91699 | 35.0 | Cluster_08866 |
| 478.21889 | 35.0 | Cluster_26309 |
| 478.50171 | 35.0 | Cluster_26327 |
| 478.51894 | 35.0 | Cluster_08877 |
| 478.79293 | 35.0 | Cluster_26336 |
| 479.23947 | 35.0 | Cluster_26383 |
| 479.24655 | 35.0 | Cluster_26380 |
| 479.24822 | 35.0 | Cluster_26381 |
| 479.32696 | 35.0 | Cluster_26391 |
| 479.48057 | 35.0 | Cluster_26410 |
| 479.98975 | 35.0 | Cluster_26444 |
| 480.19347 | 35.0 | Cluster_26456 |
| 480.23128 | 35.0 | Cluster_26461 |
| 480.87318 | 35.0 | Cluster_08912 |
| 481.21585 | 35.0 | Cluster_26502 |
| 481.22448 | 35.0 | Cluster_26493 |
| 481.55886 | 35.0 | Cluster_08922 |
| 482.56792 | 35.0 | Cluster_26551 |
| 482.75063 | 35.0 | Cluster_26575 |
| 482.75333 | 35.0 | Cluster_26573 |
| 483.26706 | 35.0 | Cluster_01822 |
| 483.28102 | 35.0 | Cluster_26592 |
| 483.51623 | 35.0 | Cluster_26609 |
| 483.77330 | 35.0 | Cluster_26614 |
| 484.26988 | 35.0 | Cluster_26656 |
| 484.56376 | 35.0 | Cluster_26659 |
| 484.74531 | 35.0 | Cluster_26674 |
| 486.05408 | 35.0 | Cluster_26738 |
| 486.30822 | 35.0 | Cluster_26750 |
| 486.77412 | 35.0 | Cluster_26773 |
| 487.78232 | 35.0 | Cluster_26821 |
| 488.49865 | 35.0 | Cluster_01848 |

| | | |
|---|---|---|
| 488.59040 | 35.0 | Cluster_09031 |
| 488.59139 | 35.0 | Cluster_09030 |
| 488.99668 | 35.0 | Cluster_26865 |
| 488.99712 | 35.0 | Cluster_26866 |
| 489.27346 | 35.0 | Cluster_09043 |
| 489.31443 | 35.0 | Cluster_26892 |
| 489.62012 | 35.0 | Cluster_09044 |
| 489.95299 | 35.0 | Cluster_09049 |
| 490.92353 | 35.0 | Cluster_26947 |
| 491.21912 | 35.0 | Cluster_26962 |
| 491.24758 | 35.0 | Cluster_26961 |
| 491.29815 | 35.0 | Cluster_09076 |
| 491.79438 | 35.0 | Cluster_26978 |
| 491.79691 | 35.0 | Cluster_26976 |
| 491.95912 | 35.0 | Cluster_09085 |
| 492.23402 | 35.0 | Cluster_26996 |
| 492.25131 | 35.0 | Cluster_01878 |
| 492.26447 | 35.0 | Cluster_27001 |
| 492.56112 | 35.0 | Cluster_27005 |
| 492.62984 | 35.0 | Cluster_00178 |
| 493.20484 | 35.0 | Cluster_27036 |
| 493.70557 | 35.0 | Cluster_27072 |
| 493.77046 | 35.0 | Cluster_27075 |
| 493.97846 | 35.0 | Cluster_27098 |
| 494.24345 | 35.0 | Cluster_09119 |
| 494.26012 | 35.0 | Cluster_27100 |
| 494.30737 | 35.0 | Cluster_27115 |
| 494.58050 | 35.0 | Cluster_09124 |
| 495.18779 | 35.0 | Cluster_27149 |
| 495.24938 | 35.0 | Cluster_27147 |
| 495.25575 | 35.0 | Cluster_09135 |
| 495.49799 | 35.0 | Cluster_27166 |
| 495.90563 | 35.0 | Cluster_09146 |
| 496.76962 | 35.0 | Cluster_27229 |
| 497.24142 | 35.0 | Cluster_27241 |
| 497.56231 | 35.0 | Cluster_09183 |
| 497.74856 | 35.0 | Cluster_01911 |
| 497.75069 | 35.0 | Cluster_01921 |
| 497.78784 | 35.0 | Cluster_27268 |
| 498.25766 | 35.0 | Cluster_09195 |
| 498.26429 | 35.0 | Cluster_27284 |
| 498.28088 | 35.0 | Cluster_27285 |
| 498.50412 | 35.0 | Cluster_27299 |
| 498.60981 | 35.0 | Cluster_09200 |
| 498.77805 | 35.0 | Cluster_27303 |
| 498.78258 | 35.0 | Cluster_27312 |
| 499.29594 | 35.0 | Cluster_27338 |
| 499.50829 | 35.0 | Cluster_01942 |
| 499.70562 | 35.0 | Cluster_27342 |
| 500.74414 | 35.0 | Cluster_01949 |
| 501.76450 | 35.0 | Cluster_27459 |

| | | |
|---|---|---|
| 502.78120 | 35.0 | Cluster_27488 |
| 502.81600 | 35.0 | Cluster_27498 |
| 502.81913 | 35.0 | Cluster_27495 |
| 503.24700 | 35.0 | Cluster_27512 |
| 503.25273 | 35.0 | Cluster_27520 |
| 503.50051 | 35.0 | Cluster_27526 |
| 504.28071 | 35.0 | Cluster_27541 |
| 504.29921 | 35.0 | Cluster_27556 |
| 505.25105 | 35.0 | Cluster_09295 |
| 505.28005 | 35.0 | Cluster_27584 |
| 505.28920 | 35.0 | Cluster_27589 |
| 505.29389 | 35.0 | Cluster_27576 |
| 505.71880 | 35.0 | Cluster_27594 |
| 505.72781 | 35.0 | Cluster_01992 |
| 505.77895 | 35.0 | Cluster_27595 |
| 506.20698 | 35.0 | Cluster_09317 |
| 506.23976 | 35.0 | Cluster_09316 |
| 506.25029 | 35.0 | Cluster_09318 |
| 506.25580 | 35.0 | Cluster_09321 |
| 506.72657 | 35.0 | Cluster_27611 |
| 507.27444 | 35.0 | Cluster_27625 |
| 507.32513 | 35.0 | Cluster_09337 |
| 507.55637 | 35.0 | Cluster_27647 |
| 507.56368 | 35.0 | Cluster_09340 |
| 507.73798 | 35.0 | Cluster_02005 |
| 507.80290 | 35.0 | Cluster_27650 |
| 507.80497 | 35.0 | Cluster_27658 |
| 508.29129 | 35.0 | Cluster_27676 |
| 508.75919 | 35.0 | Cluster_27690 |
| 509.57744 | 35.0 | Cluster_09372 |
| 510.02497 | 35.0 | Cluster_02027 |
| 510.24236 | 35.0 | Cluster_27767 |
| 510.26739 | 35.0 | Cluster_27773 |
| 510.60055 | 35.0 | Cluster_09393 |
| 510.62285 | 35.0 | Cluster_09390 |
| 510.80102 | 35.0 | Cluster_27809 |
| 511.00698 | 35.0 | Cluster_27816 |
| 511.20283 | 35.0 | Cluster_27818 |
| 511.23726 | 35.0 | Cluster_09403 |
| 511.67620 | 35.0 | Cluster_27837 |
| 512.25496 | 35.0 | Cluster_27867 |
| 512.27120 | 35.0 | Cluster_27865 |
| 512.27809 | 35.0 | Cluster_27872 |
| 513.21850 | 35.0 | Cluster_27907 |
| 513.28698 | 35.0 | Cluster_09437 |
| 513.30292 | 35.0 | Cluster_27916 |
| 513.82143 | 35.0 | Cluster_27928 |
| 514.23162 | 35.0 | Cluster_27944 |
| 514.29844 | 35.0 | Cluster_27956 |
| 514.72678 | 35.0 | Cluster_27962 |
| 515.03858 | 35.0 | Cluster_02052 |

| | | |
|---|---|---|
| 515.42879 | 35.0 | Cluster_28005 |
| 515.90242 | 35.0 | Cluster_09478 |
| 516.23238 | 35.0 | Cluster_28027 |
| 516.24464 | 35.0 | Cluster_28041 |
| 516.78373 | 35.0 | Cluster_28064 |
| 516.78653 | 35.0 | Cluster_28060 |
| 518.59799 | 35.0 | Cluster_09517 |
| 518.63157 | 35.0 | Cluster_09519 |
| 518.68185 | 35.0 | Cluster_28149 |
| 518.75080 | 35.0 | Cluster_28148 |
| 519.29071 | 35.0 | Cluster_28177 |
| 519.44164 | 35.0 | Cluster_28179 |
| 519.52561 | 35.0 | Cluster_28182 |
| 520.72991 | 35.0 | Cluster_28238 |
| 521.03117 | 35.0 | Cluster_02096 |
| 521.72792 | 35.0 | Cluster_28263 |
| 522.21350 | 35.0 | Cluster_02103 |
| 522.25391 | 35.0 | Cluster_28293 |
| 522.28500 | 35.0 | Cluster_09575 |
| 523.26552 | 35.0 | Cluster_28334 |
| 523.29131 | 35.0 | Cluster_09594 |
| 523.83465 | 35.0 | Cluster_28370 |
| 523.99562 | 35.0 | Cluster_09605 |
| 524.75967 | 35.0 | Cluster_28399 |
| 524.76802 | 35.0 | Cluster_28395 |
| 524.82869 | 35.0 | Cluster_28404 |
| 524.96550 | 35.0 | Cluster_09628 |
| 525.01312 | 35.0 | Cluster_28407 |
| 525.24753 | 35.0 | Cluster_28422 |
| 525.26257 | 35.0 | Cluster_28419 |
| 525.31479 | 35.0 | Cluster_28430 |
| 525.74833 | 35.0 | Cluster_02122 |
| 525.78681 | 35.0 | Cluster_28443 |
| 526.21218 | 35.0 | Cluster_09659 |
| 526.28131 | 35.0 | Cluster_28458 |
| 526.30647 | 35.0 | Cluster_09666 |
| 526.55381 | 35.0 | Cluster_28469 |
| 526.77481 | 35.0 | Cluster_28471 |
| 526.92737 | 35.0 | Cluster_09676 |
| 527.31121 | 35.0 | Cluster_28488 |
| 527.40923 | 35.0 | Cluster_28492 |
| 527.73779 | 35.0 | Cluster_28503 |
| 527.80726 | 35.0 | Cluster_28514 |
| 527.95744 | 35.0 | Cluster_09698 |
| 527.96172 | 35.0 | Cluster_09696 |
| 527.97250 | 35.0 | Cluster_09700 |
| 528.26935 | 35.0 | Cluster_00184 |
| 528.30280 | 35.0 | Cluster_09702 |
| 528.30501 | 35.0 | Cluster_28550 |
| 528.32878 | 35.0 | Cluster_28540 |
| 528.46671 | 35.0 | Cluster_28558 |

| | | |
|---|---|---|
| 528.49276 | 35.0 | Cluster_28556 |
| 529.27669 | 35.0 | Cluster_28578 |
| 529.46801 | 35.0 | Cluster_02143 |
| 530.02188 | 35.0 | Cluster_28624 |
| 530.29494 | 35.0 | Cluster_28649 |
| 530.30380 | 35.0 | Cluster_28641 |
| 530.78294 | 35.0 | Cluster_28675 |
| 531.19942 | 35.0 | Cluster_09748 |
| 531.25743 | 35.0 | Cluster_28699 |
| 531.26425 | 35.0 | Cluster_28687 |
| 532.96674 | 35.0 | Cluster_09773 |
| 533.26423 | 35.0 | Cluster_28757 |
| 533.26639 | 35.0 | Cluster_28756 |
| 533.28262 | 35.0 | Cluster_28768 |
| 533.34161 | 35.0 | Cluster_28774 |
| 533.75392 | 35.0 | Cluster_28791 |
| 533.77168 | 35.0 | Cluster_28786 |
| 533.91745 | 35.0 | Cluster_09785 |
| 533.92278 | 35.0 | Cluster_09791 |
| 534.53000 | 35.0 | Cluster_28822 |
| 534.75817 | 35.0 | Cluster_28839 |
| 535.32665 | 35.0 | Cluster_09813 |
| 537.26984 | 35.0 | Cluster_28925 |
| 537.29559 | 35.0 | Cluster_28936 |
| 537.30451 | 35.0 | Cluster_02195 |
| 537.52975 | 35.0 | Cluster_02196 |
| 537.79197 | 35.0 | Cluster_28941 |
| 537.91326 | 35.0 | Cluster_09869 |
| 538.52689 | 35.0 | Cluster_28962 |
| 538.79448 | 35.0 | Cluster_28978 |
| 539.73333 | 35.0 | Cluster_02212 |
| 539.87213 | 35.0 | Cluster_29005 |
| 539.97028 | 35.0 | Cluster_09906 |
| 539.98204 | 35.0 | Cluster_09905 |
| 540.02231 | 35.0 | Cluster_09899 |
| 540.26271 | 35.0 | Cluster_29033 |
| 540.58133 | 35.0 | Cluster_09919 |
| 540.93901 | 35.0 | Cluster_09927 |
| 541.04334 | 35.0 | Cluster_29059 |
| 541.12109 | 35.0 | Cluster_29068 |
| 541.26220 | 35.0 | Cluster_29080 |
| 541.79336 | 35.0 | Cluster_29114 |
| 542.33816 | 35.0 | Cluster_29137 |
| 542.52109 | 35.0 | Cluster_02234 |
| 542.52124 | 35.0 | Cluster_02233 |
| 542.67811 | 35.0 | Cluster_29147 |
| 542.80738 | 35.0 | Cluster_29176 |
| 542.81002 | 35.0 | Cluster_29175 |
| 543.23404 | 35.0 | Cluster_02240 |
| 543.26414 | 35.0 | Cluster_09960 |
| 543.27696 | 35.0 | Cluster_29192 |

| | | |
|---|---|---|
| 543.48293 | 35.0 | Cluster_02239 |
| 543.97154 | 35.0 | Cluster_09968 |
| 544.61757 | 35.0 | Cluster_09977 |
| 544.75729 | 35.0 | Cluster_29233 |
| 545.24926 | 35.0 | Cluster_29272 |
| 545.53016 | 35.0 | Cluster_02260 |
| 545.53044 | 35.0 | Cluster_29276 |
| 545.76869 | 35.0 | Cluster_29280 |
| 545.77989 | 35.0 | Cluster_29284 |
| 545.82018 | 35.0 | Cluster_29288 |
| 546.78186 | 35.0 | Cluster_29325 |
| 546.78372 | 35.0 | Cluster_29324 |
| 546.83805 | 35.0 | Cluster_29334 |
| 547.62967 | 35.0 | Cluster_10018 |
| 548.62278 | 35.0 | Cluster_10038 |
| 548.63524 | 35.0 | Cluster_10037 |
| 548.98839 | 35.0 | Cluster_29415 |
| 549.26773 | 35.0 | Cluster_29422 |
| 549.98566 | 35.0 | Cluster_10058 |
| 550.04160 | 35.0 | Cluster_29449 |
| 550.28949 | 35.0 | Cluster_10064 |
| 550.79143 | 35.0 | Cluster_29466 |
| 551.79855 | 35.0 | Cluster_29517 |
| 552.72294 | 35.0 | Cluster_29549 |
| 552.79997 | 35.0 | Cluster_29559 |
| 552.81760 | 35.0 | Cluster_29561 |
| 553.30563 | 35.0 | Cluster_10111 |
| 553.51882 | 35.0 | Cluster_02298 |
| 553.54594 | 35.0 | Cluster_29579 |
| 554.25070 | 35.0 | Cluster_02300 |
| 554.75239 | 35.0 | Cluster_29631 |
| 555.28796 | 35.0 | Cluster_10147 |
| 555.29621 | 35.0 | Cluster_29653 |
| 555.65390 | 35.0 | Cluster_10160 |
| 555.65807 | 35.0 | Cluster_10161 |
| 556.27635 | 35.0 | Cluster_29690 |
| 557.01747 | 35.0 | Cluster_29723 |
| 557.30706 | 35.0 | Cluster_29740 |
| 557.31245 | 35.0 | Cluster_10191 |
| 557.83386 | 35.0 | Cluster_29760 |
| 559.54108 | 35.0 | Cluster_02335 |
| 559.55624 | 35.0 | Cluster_29814 |
| 559.70118 | 35.0 | Cluster_29812 |
| 559.79273 | 35.0 | Cluster_29826 |
| 560.23774 | 35.0 | Cluster_29837 |
| 560.25365 | 35.0 | Cluster_10240 |
| 560.34293 | 35.0 | Cluster_29841 |
| 560.61957 | 35.0 | Cluster_10242 |
| 561.03156 | 35.0 | Cluster_29867 |
| 561.05796 | 35.0 | Cluster_29871 |
| 561.28991 | 35.0 | Cluster_29882 |

| | | |
|---|---|---|
| 561.30986 | 35.0 | Cluster_29873 |
| 561.55399 | 35.0 | Cluster_29893 |
| 561.79208 | 35.0 | Cluster_29903 |
| 562.19999 | 35.0 | Cluster_29911 |
| 562.80554 | 35.0 | Cluster_02348 |
| 562.81178 | 35.0 | Cluster_29927 |
| 563.05489 | 35.0 | Cluster_29945 |
| 563.06345 | 35.0 | Cluster_29939 |
| 563.06864 | 35.0 | Cluster_29942 |
| 563.79657 | 35.0 | Cluster_02357 |
| 565.58903 | 35.0 | Cluster_10320 |
| 565.81858 | 35.0 | Cluster_30054 |
| 566.29562 | 35.0 | Cluster_30075 |
| 566.29597 | 35.0 | Cluster_30076 |
| 566.90411 | 35.0 | Cluster_30092 |
| 567.32070 | 35.0 | Cluster_10347 |
| 568.29705 | 35.0 | Cluster_10359 |
| 568.75120 | 35.0 | Cluster_30170 |
| 568.79681 | 35.0 | Cluster_30176 |
| 569.53416 | 35.0 | Cluster_30199 |
| 571.81791 | 35.0 | Cluster_30271 |
| 571.87618 | 35.0 | Cluster_10410 |
| 571.98268 | 35.0 | Cluster_10418 |
| 572.32261 | 35.0 | Cluster_30285 |
| 574.26044 | 35.0 | Cluster_10456 |
| 574.28417 | 35.0 | Cluster_10460 |
| 574.47512 | 35.0 | Cluster_00219 |
| 576.51907 | 35.0 | Cluster_30411 |
| 576.53992 | 35.0 | Cluster_30412 |
| 577.47477 | 35.0 | Cluster_00221 |
| 577.74768 | 35.0 | Cluster_30445 |
| 577.78800 | 35.0 | Cluster_30435 |
| 578.81497 | 35.0 | Cluster_30477 |
| 579.78136 | 35.0 | Cluster_30511 |
| 580.28827 | 35.0 | Cluster_30554 |
| 580.52520 | 35.0 | Cluster_02465 |
| 580.85968 | 35.0 | Cluster_02466 |
| 581.79816 | 35.0 | Cluster_02475 |
| 581.89041 | 35.0 | Cluster_10562 |
| 582.06200 | 35.0 | Cluster_30614 |
| 582.30425 | 35.0 | Cluster_30634 |
| 582.35907 | 35.0 | Cluster_30651 |
| 584.75963 | 35.0 | Cluster_30741 |
| 584.99621 | 35.0 | Cluster_10603 |
| 586.03148 | 35.0 | Cluster_30798 |
| 586.05382 | 35.0 | Cluster_02506 |
| 586.06372 | 35.0 | Cluster_30797 |
| 586.34272 | 35.0 | Cluster_30803 |
| 586.65752 | 35.0 | Cluster_10632 |
| 586.66077 | 35.0 | Cluster_10633 |
| 587.81633 | 35.0 | Cluster_30865 |

| | | |
|---|---|---|
| 587.99851 | 35.0 | Cluster_10651 |
| 588.27868 | 35.0 | Cluster_30879 |
| 588.29050 | 35.0 | Cluster_30880 |
| 588.31813 | 35.0 | Cluster_10661 |
| 588.32128 | 35.0 | Cluster_10664 |
| 588.79784 | 35.0 | Cluster_30889 |
| 588.80006 | 35.0 | Cluster_30887 |
| 589.09090 | 35.0 | Cluster_02521 |
| 589.31162 | 35.0 | Cluster_30920 |
| 589.31328 | 35.0 | Cluster_30912 |
| 589.34271 | 35.0 | Cluster_30916 |
| 590.28217 | 35.0 | Cluster_30964 |
| 590.29951 | 35.0 | Cluster_30953 |
| 590.49205 | 35.0 | Cluster_02529 |
| 590.87542 | 35.0 | Cluster_30993 |
| 590.87542 | 35.0 | Cluster_30994 |
| 590.99826 | 35.0 | Cluster_30999 |
| 591.64593 | 35.0 | Cluster_10702 |
| 591.77316 | 35.0 | Cluster_31010 |
| 591.77316 | 35.0 | Cluster_31015 |
| 592.03205 | 35.0 | Cluster_31028 |
| 592.37681 | 35.0 | Cluster_31036 |
| 592.90589 | 35.0 | Cluster_31063 |
| 592.97018 | 35.0 | Cluster_10720 |
| 593.06175 | 35.0 | Cluster_02547 |
| 593.29794 | 35.0 | Cluster_31068 |
| 593.99147 | 35.0 | Cluster_10739 |
| 594.81425 | 35.0 | Cluster_31119 |
| 595.07225 | 35.0 | Cluster_02556 |
| 595.32870 | 35.0 | Cluster_10766 |
| 595.57788 | 35.0 | Cluster_31139 |
| 597.33731 | 35.0 | Cluster_31195 |
| 597.35703 | 35.0 | Cluster_31199 |
| 598.32380 | 35.0 | Cluster_02578 |
| 598.62702 | 35.0 | Cluster_10823 |
| 599.81361 | 35.0 | Cluster_02581 |
| 599.84927 | 35.0 | Cluster_31286 |
| 601.76195 | 35.0 | Cluster_02594 |
| 601.85727 | 35.0 | Cluster_31366 |
| 601.86414 | 35.0 | Cluster_31365 |
| 602.31203 | 35.0 | Cluster_31391 |
| 603.53345 | 35.0 | Cluster_31419 |
| 603.60560 | 35.0 | Cluster_31422 |
| 603.94866 | 35.0 | Cluster_31432 |
| 604.36908 | 35.0 | Cluster_31449 |
| 605.67389 | 35.0 | Cluster_10910 |
| 606.36332 | 35.0 | Cluster_31521 |
| 606.62627 | 35.0 | Cluster_10929 |
| 606.63827 | 35.0 | Cluster_10932 |
| 606.99602 | 35.0 | Cluster_10938 |
| 607.81374 | 35.0 | Cluster_31552 |

| | | |
|---|---|---|
| 607.86967 | 35.0 | Cluster_31561 |
| 608.00218 | 35.0 | Cluster_10951 |
| 608.32379 | 35.0 | Cluster_31587 |
| 608.37907 | 35.0 | Cluster_31579 |
| 609.53174 | 35.0 | Cluster_31620 |
| 609.54399 | 35.0 | Cluster_31621 |
| 610.58909 | 35.0 | Cluster_31662 |
| 610.81214 | 35.0 | Cluster_31672 |
| 611.77453 | 35.0 | Cluster_31712 |
| 611.95982 | 35.0 | Cluster_11001 |
| 612.30404 | 35.0 | Cluster_31723 |
| 613.33866 | 35.0 | Cluster_31764 |
| 613.84388 | 35.0 | Cluster_31788 |
| 614.30736 | 35.0 | Cluster_31801 |
| 614.32829 | 35.0 | Cluster_31804 |
| 614.58419 | 35.0 | Cluster_31817 |
| 615.31368 | 35.0 | Cluster_31835 |
| 615.36301 | 35.0 | Cluster_31840 |
| 615.65857 | 35.0 | Cluster_11061 |
| 616.34935 | 35.0 | Cluster_31871 |
| 616.54862 | 35.0 | Cluster_31873 |
| 616.81547 | 35.0 | Cluster_31888 |
| 617.81621 | 35.0 | Cluster_31926 |
| 617.99735 | 35.0 | Cluster_11092 |
| 619.23737 | 35.0 | Cluster_31977 |
| 619.23825 | 35.0 | Cluster_31978 |
| 619.33872 | 35.0 | Cluster_31982 |
| 619.63456 | 35.0 | Cluster_11118 |
| 619.65655 | 35.0 | Cluster_11120 |
| 619.78725 | 35.0 | Cluster_31996 |
| 620.32652 | 35.0 | Cluster_32018 |
| 620.38097 | 35.0 | Cluster_32024 |
| 620.83282 | 35.0 | Cluster_32031 |
| 621.29918 | 35.0 | Cluster_32049 |
| 621.39040 | 35.0 | Cluster_32052 |
| 622.79607 | 35.0 | Cluster_32093 |
| 623.30190 | 35.0 | Cluster_32111 |
| 623.64928 | 35.0 | Cluster_11191 |
| 623.83853 | 35.0 | Cluster_32135 |
| 624.98651 | 35.0 | Cluster_11209 |
| 625.85548 | 35.0 | Cluster_32203 |
| 625.97156 | 35.0 | Cluster_11220 |
| 626.01878 | 35.0 | Cluster_02740 |
| 626.31307 | 35.0 | Cluster_11232 |
| 626.38225 | 35.0 | Cluster_32224 |
| 627.11200 | 35.0 | Cluster_02743 |
| 627.38196 | 35.0 | Cluster_32271 |
| 627.62311 | 35.0 | Cluster_11256 |
| 628.00286 | 35.0 | Cluster_11263 |
| 628.33999 | 35.0 | Cluster_11261 |
| 628.55578 | 35.0 | Cluster_32312 |

| | | |
|---|---|---|
| 628.84969 | 35.0 | Cluster_32316 |
| 630.09806 | 35.0 | Cluster_02750 |
| 630.86016 | 35.0 | Cluster_32381 |
| 632.72081 | 35.0 | Cluster_32443 |
| 632.80122 | 35.0 | Cluster_32448 |
| 634.29435 | 35.0 | Cluster_11358 |
| 634.31779 | 35.0 | Cluster_32515 |
| 634.35247 | 35.0 | Cluster_11350 |
| 635.82948 | 35.0 | Cluster_32562 |
| 636.99289 | 35.0 | Cluster_11392 |
| 637.33236 | 35.0 | Cluster_32612 |
| 637.62019 | 35.0 | Cluster_11403 |
| 638.31773 | 35.0 | Cluster_11412 |
| 638.34363 | 35.0 | Cluster_32654 |
| 639.31559 | 35.0 | Cluster_32685 |
| 639.68269 | 35.0 | Cluster_11437 |
| 639.80812 | 35.0 | Cluster_32706 |
| 640.82008 | 35.0 | Cluster_32751 |
| 640.83128 | 35.0 | Cluster_32746 |
| 641.04752 | 35.0 | Cluster_32764 |
| 642.81785 | 35.0 | Cluster_32830 |
| 643.08841 | 35.0 | Cluster_32841 |
| 645.09355 | 35.0 | Cluster_02837 |
| 645.31567 | 35.0 | Cluster_00252 |
| 645.84840 | 35.0 | Cluster_32926 |
| 646.05937 | 35.0 | Cluster_02847 |
| 646.37710 | 35.0 | Cluster_32949 |
| 646.95918 | 35.0 | Cluster_11539 |
| 648.39547 | 35.0 | Cluster_33012 |
| 649.07347 | 35.0 | Cluster_33032 |
| 649.25206 | 35.0 | Cluster_33036 |
| 649.56122 | 35.0 | Cluster_02868 |
| 649.76473 | 35.0 | Cluster_33053 |
| 649.86734 | 35.0 | Cluster_33068 |
| 651.06179 | 35.0 | Cluster_33115 |
| 651.66012 | 35.0 | Cluster_11603 |
| 654.75497 | 35.0 | Cluster_33236 |
| 655.81688 | 35.0 | Cluster_33275 |
| 658.66615 | 35.0 | Cluster_33366 |
| 658.79165 | 35.0 | Cluster_33375 |
| 659.02086 | 35.0 | Cluster_11706 |
| 659.37949 | 35.0 | Cluster_11709 |
| 659.54833 | 35.0 | Cluster_33395 |
| 659.62589 | 35.0 | Cluster_02907 |
| 661.33894 | 35.0 | Cluster_11734 |
| 661.79061 | 35.0 | Cluster_33460 |
| 662.11757 | 35.0 | Cluster_02928 |
| 662.69475 | 35.0 | Cluster_11762 |
| 662.82543 | 35.0 | Cluster_02934 |
| 662.94506 | 35.0 | Cluster_00261 |
| 663.08145 | 35.0 | Cluster_33501 |

| | | |
|---|---|---|
| 663.25841 | 35.0 | Cluster_33503 |
| 663.38606 | 35.0 | Cluster_33507 |
| 663.60178 | 35.0 | Cluster_02940 |
| 664.37085 | 35.0 | Cluster_33540 |
| 664.81843 | 35.0 | Cluster_33548 |
| 665.35530 | 35.0 | Cluster_33569 |
| 665.82233 | 35.0 | Cluster_02959 |
| 665.84730 | 35.0 | Cluster_33575 |
| 666.34744 | 35.0 | Cluster_02964 |
| 666.37680 | 35.0 | Cluster_33591 |
| 667.81938 | 35.0 | Cluster_33644 |
| 667.84739 | 35.0 | Cluster_33646 |
| 667.89684 | 35.0 | Cluster_33648 |
| 667.97367 | 35.0 | Cluster_11845 |
| 667.98825 | 35.0 | Cluster_11843 |
| 668.32263 | 35.0 | Cluster_11851 |
| 668.33838 | 35.0 | Cluster_33655 |
| 668.65012 | 35.0 | Cluster_11854 |
| 669.09668 | 35.0 | Cluster_02977 |
| 669.11109 | 35.0 | Cluster_02976 |
| 669.16846 | 35.0 | Cluster_00267 |
| 669.32694 | 35.0 | Cluster_33680 |
| 669.69725 | 35.0 | Cluster_00270 |
| 669.85195 | 35.0 | Cluster_02980 |
| 670.38121 | 35.0 | Cluster_33722 |
| 670.38953 | 35.0 | Cluster_33710 |
| 670.38953 | 35.0 | Cluster_33726 |
| 670.81349 | 35.0 | Cluster_33737 |
| 671.10547 | 35.0 | Cluster_02987 |
| 671.36374 | 35.0 | Cluster_33756 |
| 671.86108 | 35.0 | Cluster_33771 |
| 672.38383 | 35.0 | Cluster_11909 |
| 673.69262 | 35.0 | Cluster_11928 |
| 673.84601 | 35.0 | Cluster_33828 |
| 674.37802 | 35.0 | Cluster_33841 |
| 675.28713 | 35.0 | Cluster_11947 |
| 675.63966 | 35.0 | Cluster_03025 |
| 675.86022 | 35.0 | Cluster_33867 |
| 676.04126 | 35.0 | Cluster_33882 |
| 676.06023 | 35.0 | Cluster_33875 |
| 676.82567 | 35.0 | Cluster_33894 |
| 677.38049 | 35.0 | Cluster_33924 |
| 677.66310 | 35.0 | Cluster_11983 |
| 678.05380 | 35.0 | Cluster_11985 |
| 678.07682 | 35.0 | Cluster_03041 |
| 678.10788 | 35.0 | Cluster_33950 |
| 678.61132 | 35.0 | Cluster_03048 |
| 679.35854 | 35.0 | Cluster_12002 |
| 680.39313 | 35.0 | Cluster_12012 |
| 680.68664 | 35.0 | Cluster_12015 |
| 680.79621 | 35.0 | Cluster_34046 |

| | | |
|---|---|---|
| 680.80240 | 35.0 | Cluster_34051 |
| 681.68356 | 35.0 | Cluster_12028 |
| 683.17868 | 35.0 | Cluster_34121 |
| 683.50325 | 35.0 | Cluster_00281 |
| 683.61484 | 35.0 | Cluster_12059 |
| 684.33882 | 35.0 | Cluster_34147 |
| 684.61364 | 35.0 | Cluster_12066 |
| 684.78469 | 35.0 | Cluster_34165 |
| 684.81441 | 35.0 | Cluster_34164 |
| 684.82466 | 35.0 | Cluster_34171 |
| 685.08807 | 35.0 | Cluster_03103 |
| 685.08939 | 35.0 | Cluster_03095 |
| 685.08962 | 35.0 | Cluster_34186 |
| 685.33558 | 35.0 | Cluster_34202 |
| 685.34195 | 35.0 | Cluster_12078 |
| 685.84087 | 35.0 | Cluster_34213 |
| 687.37344 | 35.0 | Cluster_34250 |
| 687.56541 | 35.0 | Cluster_03119 |
| 687.79947 | 35.0 | Cluster_34262 |
| 687.80190 | 35.0 | Cluster_34258 |
| 687.88397 | 35.0 | Cluster_03131 |
| 687.88623 | 35.0 | Cluster_03124 |
| 688.01588 | 35.0 | Cluster_12113 |
| 688.27721 | 35.0 | Cluster_34272 |
| 688.34085 | 35.0 | Cluster_34279 |
| 688.38593 | 35.0 | Cluster_34282 |
| 688.84881 | 35.0 | Cluster_34292 |
| 688.85810 | 35.0 | Cluster_03136 |
| 688.85987 | 35.0 | Cluster_34290 |
| 690.32779 | 35.0 | Cluster_03148 |
| 690.82663 | 35.0 | Cluster_34322 |
| 691.07637 | 35.0 | Cluster_34336 |
| 691.10745 | 35.0 | Cluster_34337 |
| 691.81429 | 35.0 | Cluster_03161 |
| 692.58945 | 35.0 | Cluster_03180 |
| 692.59083 | 35.0 | Cluster_03181 |
| 692.59314 | 35.0 | Cluster_03176 |
| 692.59958 | 35.0 | Cluster_03182 |
| 693.01929 | 35.0 | Cluster_12161 |
| 693.09427 | 35.0 | Cluster_34407 |
| 693.99699 | 35.0 | Cluster_12168 |
| 694.06755 | 35.0 | Cluster_34421 |
| 694.67508 | 35.0 | Cluster_12172 |
| 695.04142 | 35.0 | Cluster_12177 |
| 695.33781 | 35.0 | Cluster_03206 |
| 695.61408 | 35.0 | Cluster_34466 |
| 696.02156 | 35.0 | Cluster_12198 |
| 696.03203 | 35.0 | Cluster_12194 |
| 696.31580 | 35.0 | Cluster_34489 |
| 696.36747 | 35.0 | Cluster_34481 |
| 696.99121 | 35.0 | Cluster_12209 |

| | | |
|---|---|---|
| 697.02744 | 35.0 | Cluster_12212 |
| 697.40157 | 35.0 | Cluster_34514 |
| 698.56565 | 35.0 | Cluster_03224 |
| 698.56567 | 35.0 | Cluster_03225 |
| 698.69069 | 35.0 | Cluster_12232 |
| 699.09398 | 35.0 | Cluster_34549 |
| 699.34183 | 35.0 | Cluster_12236 |
| 699.70254 | 35.0 | Cluster_12244 |
| 699.90969 | 35.0 | Cluster_34574 |
| 700.06571 | 35.0 | Cluster_03240 |
| 700.31327 | 35.0 | Cluster_34581 |
| 700.87788 | 35.0 | Cluster_34610 |
| 701.80476 | 35.0 | Cluster_34627 |
| 702.28565 | 35.0 | Cluster_34650 |
| 702.29475 | 35.0 | Cluster_34647 |
| 702.62288 | 35.0 | Cluster_12286 |
| 702.93375 | 35.0 | Cluster_34673 |
| 703.31422 | 35.0 | Cluster_34685 |
| 703.60021 | 35.0 | Cluster_34686 |
| 704.21072 | 35.0 | Cluster_34700 |
| 704.32933 | 35.0 | Cluster_34707 |
| 704.35627 | 35.0 | Cluster_12313 |
| 704.78009 | 35.0 | Cluster_34721 |
| 704.85837 | 35.0 | Cluster_34731 |
| 704.88607 | 35.0 | Cluster_34735 |
| 705.02263 | 35.0 | Cluster_12328 |
| 705.34861 | 35.0 | Cluster_12324 |
| 705.41349 | 35.0 | Cluster_34748 |
| 705.70698 | 35.0 | Cluster_12329 |
| 706.56252 | 35.0 | Cluster_00307 |
| 706.62232 | 35.0 | Cluster_03289 |
| 706.62925 | 35.0 | Cluster_03297 |
| 706.63126 | 35.0 | Cluster_03300 |
| 706.76398 | 35.0 | Cluster_00306 |
| 706.86361 | 35.0 | Cluster_03301 |
| 708.60835 | 35.0 | Cluster_34818 |
| 708.87437 | 35.0 | Cluster_34828 |
| 709.28860 | 35.0 | Cluster_34838 |
| 709.34977 | 35.0 | Cluster_12378 |
| 709.66787 | 35.0 | Cluster_34855 |
| 709.81635 | 35.0 | Cluster_34851 |
| 710.60612 | 35.0 | Cluster_34899 |
| 710.90108 | 35.0 | Cluster_34904 |
| 711.83266 | 35.0 | Cluster_03342 |
| 711.83663 | 35.0 | Cluster_03347 |
| 711.86301 | 35.0 | Cluster_34922 |
| 713.27764 | 35.0 | Cluster_34969 |
| 713.29080 | 35.0 | Cluster_03356 |
| 713.36980 | 35.0 | Cluster_34978 |
| 713.86227 | 35.0 | Cluster_03365 |
| 713.91951 | 35.0 | Cluster_34998 |

| | | |
|---|---|---|
| 714.36691 | 35.0 | Cluster_35003 |
| 714.82840 | 35.0 | Cluster_35015 |
| 714.83014 | 35.0 | Cluster_35026 |
| 715.18884 | 35.0 | Cluster_35027 |
| 717.16182 | 35.0 | Cluster_35084 |
| 717.26810 | 35.0 | Cluster_35093 |
| 719.01137 | 35.0 | Cluster_12500 |
| 719.31062 | 35.0 | Cluster_35143 |
| 719.61357 | 35.0 | Cluster_03408 |
| 719.62148 | 35.0 | Cluster_03410 |
| 719.86761 | 35.0 | Cluster_35165 |
| 720.05737 | 35.0 | Cluster_12523 |
| 721.42197 | 35.0 | Cluster_35196 |
| 721.86747 | 35.0 | Cluster_03429 |
| 721.92070 | 35.0 | Cluster_35214 |
| 722.42456 | 35.0 | Cluster_35223 |
| 722.60635 | 35.0 | Cluster_03438 |
| 722.61381 | 35.0 | Cluster_03439 |
| 722.70199 | 35.0 | Cluster_12561 |
| 723.69044 | 35.0 | Cluster_35246 |
| 723.73145 | 35.0 | Cluster_00325 |
| 724.01495 | 35.0 | Cluster_12583 |
| 724.63248 | 35.0 | Cluster_35271 |
| 727.04057 | 35.0 | Cluster_12624 |
| 727.11444 | 35.0 | Cluster_03480 |
| 727.21257 | 35.0 | Cluster_35312 |
| 727.72402 | 35.0 | Cluster_35325 |
| 728.02187 | 35.0 | Cluster_12644 |
| 729.13426 | 35.0 | Cluster_35352 |
| 729.99899 | 35.0 | Cluster_12666 |
| 731.01313 | 35.0 | Cluster_12681 |
| 731.86730 | 35.0 | Cluster_35428 |
| 732.35032 | 35.0 | Cluster_12702 |
| 732.63776 | 35.0 | Cluster_03534 |
| 733.11661 | 35.0 | Cluster_03543 |
| 733.11760 | 35.0 | Cluster_35455 |
| 733.11909 | 35.0 | Cluster_03544 |
| 733.12735 | 35.0 | Cluster_03548 |
| 733.36842 | 35.0 | Cluster_03556 |
| 733.36858 | 35.0 | Cluster_03555 |
| 735.35959 | 35.0 | Cluster_35520 |
| 736.22851 | 35.0 | Cluster_35536 |
| 736.53601 | 35.0 | Cluster_35546 |
| 736.85330 | 35.0 | Cluster_03578 |
| 737.11647 | 35.0 | Cluster_03579 |
| 737.33266 | 35.0 | Cluster_35583 |
| 737.69447 | 35.0 | Cluster_12767 |
| 738.10935 | 35.0 | Cluster_03586 |
| 738.30853 | 35.0 | Cluster_35618 |
| 738.33760 | 35.0 | Cluster_12777 |
| 738.37561 | 35.0 | Cluster_35615 |

| | | |
|---|---|---|
| 738.88226 | 35.0 | Cluster_35625 |
| 739.32896 | 35.0 | Cluster_12785 |
| 740.02039 | 35.0 | Cluster_12796 |
| 740.69260 | 35.0 | Cluster_12808 |
| 742.33310 | 35.0 | Cluster_35695 |
| 742.39069 | 35.0 | Cluster_35694 |
| 742.68914 | 35.0 | Cluster_12829 |
| 743.35586 | 35.0 | Cluster_35729 |
| 744.07867 | 35.0 | Cluster_12846 |
| 744.35590 | 35.0 | Cluster_35745 |
| 744.42886 | 35.0 | Cluster_35754 |
| 744.87374 | 35.0 | Cluster_35760 |
| 745.71379 | 35.0 | Cluster_12858 |
| 745.87885 | 35.0 | Cluster_35786 |
| 746.06574 | 35.0 | Cluster_12867 |
| 746.39046 | 35.0 | Cluster_12876 |
| 748.06008 | 35.0 | Cluster_12907 |
| 748.86103 | 35.0 | Cluster_03654 |
| 749.38170 | 35.0 | Cluster_35861 |
| 749.39351 | 35.0 | Cluster_35865 |
| 749.43777 | 35.0 | Cluster_35870 |
| 750.39152 | 35.0 | Cluster_35881 |
| 751.69588 | 35.0 | Cluster_12965 |
| 751.80275 | 35.0 | Cluster_35915 |
| 753.38453 | 35.0 | Cluster_35960 |
| 753.78417 | 35.0 | Cluster_35982 |
| 754.58916 | 35.0 | Cluster_03681 |
| 754.73608 | 35.0 | Cluster_13017 |
| 754.80036 | 35.0 | Cluster_35999 |
| 755.85015 | 35.0 | Cluster_03685 |
| 755.87117 | 35.0 | Cluster_36038 |
| 756.39066 | 35.0 | Cluster_36047 |
| 756.42287 | 35.0 | Cluster_36046 |
| 756.59788 | 35.0 | Cluster_03689 |
| 757.07652 | 35.0 | Cluster_13067 |
| 757.30855 | 35.0 | Cluster_36071 |
| 758.22619 | 35.0 | Cluster_36095 |
| 758.73364 | 35.0 | Cluster_36107 |
| 759.06890 | 35.0 | Cluster_13093 |
| 759.37381 | 35.0 | Cluster_36124 |
| 761.02441 | 35.0 | Cluster_13116 |
| 761.45335 | 35.0 | Cluster_13117 |
| 761.64099 | 35.0 | Cluster_36196 |
| 762.05458 | 35.0 | Cluster_13131 |
| 762.07253 | 35.0 | Cluster_13132 |
| 762.10790 | 35.0 | Cluster_36203 |
| 762.14821 | 35.0 | Cluster_03718 |
| 762.28681 | 35.0 | Cluster_36205 |
| 762.33840 | 35.0 | Cluster_36213 |
| 762.34180 | 35.0 | Cluster_13133 |
| 762.38228 | 35.0 | Cluster_13130 |

| | | |
|---|---|---|
| 764.06043 | 35.0 | Cluster_13156 |
| 766.08420 | 35.0 | Cluster_03757 |
| 766.32954 | 35.0 | Cluster_13190 |
| 766.87855 | 35.0 | Cluster_36304 |
| 767.16279 | 35.0 | Cluster_03766 |
| 768.06607 | 35.0 | Cluster_13209 |
| 769.12088 | 35.0 | Cluster_36363 |
| 769.40797 | 35.0 | Cluster_36377 |
| 769.46550 | 35.0 | Cluster_36384 |
| 769.72586 | 35.0 | Cluster_13232 |
| 770.69391 | 35.0 | Cluster_13238 |
| 771.07220 | 35.0 | Cluster_13246 |
| 771.56399 | 35.0 | Cluster_00342 |
| 771.88334 | 35.0 | Cluster_03784 |
| 771.93573 | 35.0 | Cluster_36427 |
| 773.13438 | 35.0 | Cluster_13272 |
| 773.87560 | 35.0 | Cluster_03789 |
| 774.04780 | 35.0 | Cluster_13292 |
| 774.17570 | 35.0 | Cluster_36483 |
| 774.37503 | 35.0 | Cluster_13304 |
| 774.85555 | 35.0 | Cluster_36496 |
| 775.66717 | 35.0 | Cluster_13318 |
| 775.87209 | 35.0 | Cluster_36521 |
| 777.70948 | 35.0 | Cluster_13338 |
| 777.83832 | 35.0 | Cluster_36578 |
| 777.84116 | 35.0 | Cluster_36575 |
| 778.37272 | 35.0 | Cluster_36591 |
| 780.30002 | 35.0 | Cluster_36632 |
| 780.74770 | 35.0 | Cluster_13383 |
| 781.40025 | 35.0 | Cluster_36661 |
| 782.03197 | 35.0 | Cluster_13403 |
| 783.12152 | 35.0 | Cluster_03835 |
| 783.88454 | 35.0 | Cluster_36719 |
| 784.06281 | 35.0 | Cluster_13433 |
| 784.14461 | 35.0 | Cluster_03845 |
| 784.88961 | 35.0 | Cluster_36733 |
| 784.95613 | 35.0 | Cluster_36736 |
| 785.02199 | 35.0 | Cluster_13443 |
| 785.09844 | 35.0 | Cluster_36737 |
| 785.41303 | 35.0 | Cluster_36755 |
| 785.61996 | 35.0 | Cluster_03854 |
| 785.87805 | 35.0 | Cluster_03853 |
| 786.60068 | 35.0 | Cluster_03865 |
| 786.82147 | 35.0 | Cluster_36782 |
| 787.05983 | 35.0 | Cluster_13459 |
| 787.40128 | 35.0 | Cluster_03872 |
| 788.36857 | 35.0 | Cluster_36828 |
| 788.38543 | 35.0 | Cluster_36836 |
| 788.41338 | 35.0 | Cluster_13472 |
| 788.70878 | 35.0 | Cluster_13473 |
| 788.86008 | 35.0 | Cluster_36852 |

| | | |
|---|---|---|
| 788.86793 | 35.0 | Cluster_36840 |
| 789.11768 | 35.0 | Cluster_03883 |
| 789.19890 | 35.0 | Cluster_00345 |
| 789.38869 | 35.0 | Cluster_36865 |
| 789.61728 | 35.0 | Cluster_03891 |
| 789.83948 | 35.0 | Cluster_36869 |
| 789.85099 | 35.0 | Cluster_36877 |
| 789.86668 | 35.0 | Cluster_36866 |
| 790.03748 | 35.0 | Cluster_13497 |
| 790.07355 | 35.0 | Cluster_13501 |
| 790.34222 | 35.0 | Cluster_36880 |
| 791.39206 | 35.0 | Cluster_03901 |
| 791.45763 | 35.0 | Cluster_36916 |
| 792.14125 | 35.0 | Cluster_36929 |
| 792.68615 | 35.0 | Cluster_13537 |
| 792.80105 | 35.0 | Cluster_36946 |
| 792.96916 | 35.0 | Cluster_13541 |
| 793.45015 | 35.0 | Cluster_36966 |
| 793.95164 | 35.0 | Cluster_36976 |
| 794.13239 | 35.0 | Cluster_36978 |
| 794.42894 | 35.0 | Cluster_36983 |
| 795.05070 | 35.0 | Cluster_13574 |
| 795.77104 | 35.0 | Cluster_37009 |
| 796.39992 | 35.0 | Cluster_37032 |
| 796.40163 | 35.0 | Cluster_37036 |
| 796.88868 | 35.0 | Cluster_03956 |
| 796.98818 | 35.0 | Cluster_37058 |
| 797.13475 | 35.0 | Cluster_03957 |
| 798.35794 | 35.0 | Cluster_13632 |
| 798.91924 | 35.0 | Cluster_37110 |
| 799.16011 | 35.0 | Cluster_03969 |
| 799.90299 | 35.0 | Cluster_37123 |
| 799.92816 | 35.0 | Cluster_37119 |
| 800.38300 | 35.0 | Cluster_37139 |
| 800.39257 | 35.0 | Cluster_37138 |
| 800.89786 | 35.0 | Cluster_37148 |
| 800.90112 | 35.0 | Cluster_00030 |
| 800.91677 | 35.0 | Cluster_03988 |
| 800.91942 | 35.0 | Cluster_37153 |
| 801.46098 | 35.0 | Cluster_37165 |
| 801.70986 | 35.0 | Cluster_13663 |
| 801.73389 | 35.0 | Cluster_13658 |
| 802.04041 | 35.0 | Cluster_13662 |
| 802.36221 | 35.0 | Cluster_03998 |
| 802.38466 | 35.0 | Cluster_37185 |
| 802.41035 | 35.0 | Cluster_13666 |
| 802.42282 | 35.0 | Cluster_37192 |
| 802.74372 | 35.0 | Cluster_13681 |
| 802.91353 | 35.0 | Cluster_04000 |
| 804.33872 | 35.0 | Cluster_13691 |
| 804.41270 | 35.0 | Cluster_37237 |

| | | |
|---|---|---|
| 804.44409 | 35.0 | Cluster_13695 |
| 804.90323 | 35.0 | Cluster_37264 |
| 804.92086 | 35.0 | Cluster_37265 |
| 805.64630 | 35.0 | Cluster_04018 |
| 806.00995 | 35.0 | Cluster_37294 |
| 806.43863 | 35.0 | Cluster_37309 |
| 806.92500 | 35.0 | Cluster_37317 |
| 807.14878 | 35.0 | Cluster_04029 |
| 807.73159 | 35.0 | Cluster_13735 |
| 807.87638 | 35.0 | Cluster_04031 |
| 807.89009 | 35.0 | Cluster_37340 |
| 807.89822 | 35.0 | Cluster_37345 |
| 808.39507 | 35.0 | Cluster_37359 |
| 808.84749 | 35.0 | Cluster_37370 |
| 808.89580 | 35.0 | Cluster_37373 |
| 808.92878 | 35.0 | Cluster_04036 |
| 809.05025 | 35.0 | Cluster_13760 |
| 809.73734 | 35.0 | Cluster_13764 |
| 809.78549 | 35.0 | Cluster_13767 |
| 811.43019 | 35.0 | Cluster_37431 |
| 811.74034 | 35.0 | Cluster_00036 |
| 811.85919 | 35.0 | Cluster_37446 |
| 812.10995 | 35.0 | Cluster_13797 |
| 812.41776 | 35.0 | Cluster_37457 |
| 812.65606 | 35.0 | Cluster_04058 |
| 812.85184 | 35.0 | Cluster_37467 |
| 813.72689 | 35.0 | Cluster_13823 |
| 813.74559 | 35.0 | Cluster_13819 |
| 814.00285 | 35.0 | Cluster_37491 |
| 814.42062 | 35.0 | Cluster_37495 |
| 814.57404 | 35.0 | Cluster_37500 |
| 814.74378 | 35.0 | Cluster_13828 |
| 814.89596 | 35.0 | Cluster_04074 |
| 815.16227 | 35.0 | Cluster_04078 |
| 815.68634 | 35.0 | Cluster_13842 |
| 815.80355 | 35.0 | Cluster_00353 |
| 815.87690 | 35.0 | Cluster_37524 |
| 815.87690 | 35.0 | Cluster_37536 |
| 815.93641 | 35.0 | Cluster_37532 |
| 816.70520 | 35.0 | Cluster_04090 |
| 818.38020 | 35.0 | Cluster_37581 |
| 818.87672 | 35.0 | Cluster_37589 |
| 819.16928 | 35.0 | Cluster_04097 |
| 820.08348 | 35.0 | Cluster_13916 |
| 820.84883 | 35.0 | Cluster_37627 |
| 821.66013 | 35.0 | Cluster_13928 |
| 822.07133 | 35.0 | Cluster_13931 |
| 823.39219 | 35.0 | Cluster_04123 |
| 826.90824 | 35.0 | Cluster_37758 |
| 827.36936 | 35.0 | Cluster_37773 |
| 827.77245 | 35.0 | Cluster_13990 |

| | | |
|---|---|---|
| 828.79852 | 35.0 | Cluster_14000 |
| 829.43734 | 35.0 | Cluster_37818 |
| 829.90674 | 35.0 | Cluster_04169 |
| 830.07611 | 35.0 | Cluster_14011 |
| 830.40984 | 35.0 | Cluster_37831 |
| 831.04351 | 35.0 | Cluster_14024 |
| 831.16303 | 35.0 | Cluster_04175 |
| 831.37479 | 35.0 | Cluster_37854 |
| 832.06644 | 35.0 | Cluster_14030 |
| 832.16480 | 35.0 | Cluster_04195 |
| 832.16812 | 35.0 | Cluster_04194 |
| 833.89799 | 35.0 | Cluster_37898 |
| 834.49238 | 35.0 | Cluster_37906 |
| 834.79922 | 35.0 | Cluster_14057 |
| 834.92208 | 35.0 | Cluster_37924 |
| 836.67277 | 35.0 | Cluster_04230 |
| 836.75775 | 35.0 | Cluster_14083 |
| 836.87714 | 35.0 | Cluster_04232 |
| 836.96388 | 35.0 | Cluster_37964 |
| 838.42552 | 35.0 | Cluster_37988 |
| 839.42310 | 35.0 | Cluster_38013 |
| 839.42443 | 35.0 | Cluster_38016 |
| 840.38600 | 35.0 | Cluster_38045 |
| 840.42961 | 35.0 | Cluster_38043 |
| 840.65991 | 35.0 | Cluster_04260 |
| 840.93222 | 35.0 | Cluster_38057 |
| 842.15556 | 35.0 | Cluster_04278 |
| 842.42996 | 35.0 | Cluster_38083 |
| 843.17212 | 35.0 | Cluster_38098 |
| 843.91630 | 35.0 | Cluster_38116 |
| 843.92674 | 35.0 | Cluster_04290 |
| 844.40997 | 35.0 | Cluster_04296 |
| 845.41720 | 35.0 | Cluster_38145 |
| 845.69271 | 35.0 | Cluster_04313 |
| 846.16906 | 35.0 | Cluster_04326 |
| 846.64883 | 35.0 | Cluster_38171 |
| 846.91185 | 35.0 | Cluster_38178 |
| 847.87944 | 35.0 | Cluster_38203 |
| 847.97354 | 35.0 | Cluster_38213 |
| 848.09241 | 35.0 | Cluster_14192 |
| 848.42873 | 35.0 | Cluster_14196 |
| 848.44349 | 35.0 | Cluster_14193 |
| 849.05340 | 35.0 | Cluster_14199 |
| 849.66179 | 35.0 | Cluster_04366 |
| 849.89533 | 35.0 | Cluster_04373 |
| 850.70279 | 35.0 | Cluster_38261 |
| 850.93488 | 35.0 | Cluster_04378 |
| 851.45581 | 35.0 | Cluster_38292 |
| 851.93495 | 35.0 | Cluster_38300 |
| 852.36811 | 35.0 | Cluster_38309 |
| 852.39296 | 35.0 | Cluster_04398 |

| | | |
|---|---|---|
| 852.87918 | 35.0 | Cluster_38320 |
| 853.91600 | 35.0 | Cluster_38358 |
| 853.91840 | 35.0 | Cluster_38362 |
| 854.86483 | 35.0 | Cluster_38381 |
| 854.89483 | 35.0 | Cluster_38383 |
| 855.42613 | 35.0 | Cluster_38394 |
| 856.19549 | 35.0 | Cluster_38404 |
| 856.42831 | 35.0 | Cluster_14249 |
| 857.47441 | 35.0 | Cluster_38452 |
| 858.90187 | 35.0 | Cluster_04463 |
| 859.48300 | 35.0 | Cluster_38507 |
| 859.90011 | 35.0 | Cluster_38518 |
| 859.90579 | 35.0 | Cluster_04476 |
| 859.94338 | 35.0 | Cluster_38512 |
| 860.40068 | 35.0 | Cluster_38529 |
| 860.43330 | 35.0 | Cluster_38536 |
| 860.86555 | 35.0 | Cluster_00368 |
| 861.37862 | 35.0 | Cluster_38563 |
| 862.39101 | 35.0 | Cluster_04505 |
| 862.89909 | 35.0 | Cluster_04512 |
| 862.94643 | 35.0 | Cluster_04515 |
| 863.41282 | 35.0 | Cluster_38604 |
| 863.91763 | 35.0 | Cluster_38614 |
| 863.93693 | 35.0 | Cluster_38615 |
| 864.91738 | 35.0 | Cluster_38635 |
| 865.65650 | 35.0 | Cluster_04549 |
| 865.94266 | 35.0 | Cluster_38656 |
| 866.50122 | 35.0 | Cluster_14332 |
| 866.64805 | 35.0 | Cluster_04564 |
| 866.90274 | 35.0 | Cluster_38675 |
| 867.15163 | 35.0 | Cluster_04567 |
| 867.42398 | 35.0 | Cluster_14342 |
| 867.94135 | 35.0 | Cluster_38687 |
| 868.95585 | 35.0 | Cluster_38713 |
| 869.39554 | 35.0 | Cluster_38720 |
| 869.40591 | 35.0 | Cluster_04599 |
| 869.65175 | 35.0 | Cluster_04607 |
| 871.15725 | 35.0 | Cluster_04624 |
| 871.38291 | 35.0 | Cluster_04626 |
| 871.44686 | 35.0 | Cluster_38770 |
| 871.48385 | 35.0 | Cluster_14389 |
| 871.93492 | 35.0 | Cluster_38790 |
| 871.99104 | 35.0 | Cluster_38783 |
| 872.21335 | 35.0 | Cluster_38793 |
| 872.42942 | 35.0 | Cluster_38804 |
| 872.93760 | 35.0 | Cluster_38810 |
| 873.02168 | 35.0 | Cluster_38812 |
| 873.92713 | 35.0 | Cluster_38826 |
| 874.05293 | 35.0 | Cluster_38841 |
| 874.91378 | 35.0 | Cluster_04666 |
| 874.98369 | 35.0 | Cluster_38855 |

| | | |
|---|---|---|
| 875.77964 | 35.0 | Cluster_14433 |
| 876.13511 | 35.0 | Cluster_04697 |
| 876.75188 | 35.0 | Cluster_14443 |
| 876.75497 | 35.0 | Cluster_14439 |
| 876.75613 | 35.0 | Cluster_14442 |
| 876.80258 | 35.0 | Cluster_14444 |
| 877.39658 | 35.0 | Cluster_04709 |
| 877.44878 | 35.0 | Cluster_14450 |
| 877.88623 | 35.0 | Cluster_04717 |
| 877.91326 | 35.0 | Cluster_04713 |
| 878.14699 | 35.0 | Cluster_04718 |
| 879.16542 | 35.0 | Cluster_14474 |
| 879.16633 | 35.0 | Cluster_14473 |
| 879.82195 | 35.0 | Cluster_14482 |
| 879.82480 | 35.0 | Cluster_14476 |
| 879.90195 | 35.0 | Cluster_04734 |
| 881.79493 | 35.0 | Cluster_14500 |
| 881.97445 | 35.0 | Cluster_38983 |
| 882.02414 | 35.0 | Cluster_39002 |
| 882.08171 | 35.0 | Cluster_14501 |
| 882.66243 | 35.0 | Cluster_04748 |
| 882.91454 | 35.0 | Cluster_39015 |
| 883.40859 | 35.0 | Cluster_04754 |
| 883.43072 | 35.0 | Cluster_14515 |
| 883.77664 | 35.0 | Cluster_14526 |
| 884.44669 | 35.0 | Cluster_04756 |
| 885.13034 | 35.0 | Cluster_14552 |
| 885.68403 | 35.0 | Cluster_04765 |
| 885.69766 | 35.0 | Cluster_04767 |
| 885.97171 | 35.0 | Cluster_39076 |
| 885.99322 | 35.0 | Cluster_39073 |
| 886.12034 | 35.0 | Cluster_14560 |
| 887.13762 | 35.0 | Cluster_14569 |
| 887.51949 | 35.0 | Cluster_39107 |
| 888.09975 | 35.0 | Cluster_14576 |
| 888.48577 | 35.0 | Cluster_39122 |
| 889.43944 | 35.0 | Cluster_39147 |
| 889.95457 | 35.0 | Cluster_39156 |
| 892.45399 | 35.0 | Cluster_39210 |
| 893.42606 | 35.0 | Cluster_39231 |
| 894.93787 | 35.0 | Cluster_39259 |
| 894.99352 | 35.0 | Cluster_04854 |
| 895.80879 | 35.0 | Cluster_14625 |
| 896.76001 | 35.0 | Cluster_14640 |
| 896.77212 | 35.0 | Cluster_14637 |
| 897.39803 | 35.0 | Cluster_39296 |
| 897.48380 | 35.0 | Cluster_14652 |
| 897.94487 | 35.0 | Cluster_39306 |
| 897.95625 | 35.0 | Cluster_39315 |
| 899.10766 | 35.0 | Cluster_14671 |
| 899.12716 | 35.0 | Cluster_14674 |

| | | |
|---|---|---|
| 899.46375 | 35.0 | Cluster_39340 |
| 899.92997 | 35.0 | Cluster_39346 |
| 900.44745 | 35.0 | Cluster_04893 |
| 900.93632 | 35.0 | Cluster_39368 |
| 900.96315 | 35.0 | Cluster_39369 |
| 901.00269 | 35.0 | Cluster_39384 |
| 902.56355 | 35.0 | Cluster_39414 |
| 902.56355 | 35.0 | Cluster_39415 |
| 902.80635 | 35.0 | Cluster_14691 |
| 902.83968 | 35.0 | Cluster_14697 |
| 903.09960 | 35.0 | Cluster_14690 |
| 903.43465 | 35.0 | Cluster_39426 |
| 903.95125 | 35.0 | Cluster_04920 |
| 904.40367 | 35.0 | Cluster_39437 |
| 904.45901 | 35.0 | Cluster_39443 |
| 904.93200 | 35.0 | Cluster_39455 |
| 906.97714 | 35.0 | Cluster_39478 |
| 907.38470 | 35.0 | Cluster_39484 |
| 907.49299 | 35.0 | Cluster_39483 |
| 907.96227 | 35.0 | Cluster_39490 |
| 907.99865 | 35.0 | Cluster_39496 |
| 908.68258 | 35.0 | Cluster_04964 |
| 908.85062 | 35.0 | Cluster_00383 |
| 908.94704 | 35.0 | Cluster_39504 |
| 909.97831 | 35.0 | Cluster_39519 |
| 910.12391 | 35.0 | Cluster_39518 |
| 910.12524 | 35.0 | Cluster_14768 |
| 911.98028 | 35.0 | Cluster_39560 |
| 912.42574 | 35.0 | Cluster_14806 |
| 912.80213 | 35.0 | Cluster_14805 |
| 912.92380 | 35.0 | Cluster_39580 |
| 912.92797 | 35.0 | Cluster_39581 |
| 912.95093 | 35.0 | Cluster_39585 |
| 914.26396 | 35.0 | Cluster_00389 |
| 914.45343 | 35.0 | Cluster_14816 |
| 915.11024 | 35.0 | Cluster_14822 |
| 916.09860 | 35.0 | Cluster_14835 |
| 916.44537 | 35.0 | Cluster_39639 |
| 916.94625 | 35.0 | Cluster_39657 |
| 917.97343 | 35.0 | Cluster_39674 |
| 918.44980 | 35.0 | Cluster_39693 |
| 918.80128 | 35.0 | Cluster_14871 |
| 919.26354 | 35.0 | Cluster_39707 |
| 919.47261 | 35.0 | Cluster_14874 |
| 920.10752 | 35.0 | Cluster_14882 |
| 920.28060 | 35.0 | Cluster_39719 |
| 920.62551 | 35.0 | Cluster_00053 |
| 920.77045 | 35.0 | Cluster_14885 |
| 922.44243 | 35.0 | Cluster_14905 |
| 922.96183 | 35.0 | Cluster_39769 |
| 923.09415 | 35.0 | Cluster_14931 |

| | | |
|---|---|---|
| 923.94294 | 35.0 | Cluster_39783 |
| 923.98960 | 35.0 | Cluster_39784 |
| 924.43536 | 35.0 | Cluster_14953 |
| 924.92883 | 35.0 | Cluster_39798 |
| 924.96027 | 35.0 | Cluster_39797 |
| 926.01701 | 35.0 | Cluster_39810 |
| 926.13223 | 35.0 | Cluster_14978 |
| 926.45313 | 35.0 | Cluster_14969 |
| 926.46599 | 35.0 | Cluster_14973 |
| 926.49297 | 35.0 | Cluster_39814 |
| 926.49478 | 35.0 | Cluster_14979 |
| 927.43778 | 35.0 | Cluster_39836 |
| 927.92586 | 35.0 | Cluster_39851 |
| 928.44165 | 35.0 | Cluster_39864 |
| 928.47669 | 35.0 | Cluster_39862 |
| 930.81778 | 35.0 | Cluster_15028 |
| 930.92703 | 35.0 | Cluster_05035 |
| 930.97959 | 35.0 | Cluster_39916 |
| 930.98681 | 35.0 | Cluster_39912 |
| 931.41413 | 35.0 | Cluster_39913 |
| 931.75879 | 35.0 | Cluster_15046 |
| 931.80684 | 35.0 | Cluster_15048 |
| 931.89565 | 35.0 | Cluster_39924 |
| 932.63768 | 35.0 | Cluster_15062 |
| 932.75835 | 35.0 | Cluster_15059 |
| 933.92763 | 35.0 | Cluster_39962 |
| 934.94244 | 35.0 | Cluster_39977 |
| 936.16095 | 35.0 | Cluster_15109 |
| 936.48194 | 35.0 | Cluster_39994 |
| 936.97952 | 35.0 | Cluster_40008 |
| 937.48268 | 35.0 | Cluster_15142 |
| 937.71270 | 35.0 | Cluster_40015 |
| 937.73608 | 35.0 | Cluster_40014 |
| 937.94932 | 35.0 | Cluster_05054 |
| 938.37752 | 35.0 | Cluster_40030 |
| 938.37914 | 35.0 | Cluster_40031 |
| 938.80176 | 35.0 | Cluster_15159 |
| 938.94420 | 35.0 | Cluster_40033 |
| 938.96404 | 35.0 | Cluster_05057 |
| 938.96620 | 35.0 | Cluster_40042 |
| 939.15327 | 35.0 | Cluster_15165 |
| 939.68688 | 35.0 | Cluster_05058 |
| 941.11983 | 35.0 | Cluster_15193 |
| 941.98683 | 35.0 | Cluster_40092 |
| 942.14657 | 35.0 | Cluster_15209 |
| 942.48402 | 35.0 | Cluster_40107 |
| 943.92893 | 35.0 | Cluster_40133 |
| 943.96400 | 35.0 | Cluster_40118 |
| 944.17376 | 35.0 | Cluster_05071 |
| 945.38704 | 35.0 | Cluster_15242 |
| 945.69833 | 35.0 | Cluster_00409 |

| | | |
|---|---|---|
| 946.02003 | 35.0 | Cluster_05076 |
| 946.96633 | 35.0 | Cluster_05081 |
| 946.97997 | 35.0 | Cluster_40166 |
| 947.13555 | 35.0 | Cluster_15263 |
| 947.51320 | 35.0 | Cluster_15268 |
| 948.12920 | 35.0 | Cluster_15278 |
| 949.44613 | 35.0 | Cluster_40193 |
| 949.47546 | 35.0 | Cluster_15306 |
| 949.48378 | 35.0 | Cluster_15297 |
| 949.79869 | 35.0 | Cluster_40191 |
| 951.98931 | 35.0 | Cluster_40222 |
| 952.14699 | 35.0 | Cluster_15339 |
| 952.88349 | 35.0 | Cluster_15346 |
| 952.89465 | 35.0 | Cluster_00415 |
| 954.41272 | 35.0 | Cluster_15371 |
| 954.78930 | 35.0 | Cluster_15383 |
| 954.80409 | 35.0 | Cluster_15390 |
| 954.80629 | 35.0 | Cluster_15384 |
| 954.82575 | 35.0 | Cluster_15380 |
| 955.47325 | 35.0 | Cluster_15404 |
| 955.49162 | 35.0 | Cluster_40268 |
| 956.15976 | 35.0 | Cluster_15421 |
| 956.44311 | 35.0 | Cluster_15422 |
| 956.47325 | 35.0 | Cluster_40279 |
| 956.47921 | 35.0 | Cluster_15426 |
| 956.80746 | 35.0 | Cluster_15424 |
| 956.81074 | 35.0 | Cluster_15425 |
| 957.42854 | 35.0 | Cluster_40288 |
| 958.70795 | 35.0 | Cluster_05141 |
| 960.16356 | 35.0 | Cluster_15469 |
| 960.83092 | 35.0 | Cluster_15476 |
| 961.21712 | 35.0 | Cluster_40324 |
| 961.71471 | 35.0 | Cluster_05152 |
| 962.78316 | 35.0 | Cluster_15504 |
| 963.14215 | 35.0 | Cluster_15508 |
| 964.47449 | 35.0 | Cluster_40369 |
| 964.71819 | 35.0 | Cluster_40371 |
| 964.98137 | 35.0 | Cluster_05166 |
| 965.80056 | 35.0 | Cluster_15544 |
| 965.82093 | 35.0 | Cluster_15540 |
| 966.92472 | 35.0 | Cluster_40391 |
| 968.00687 | 35.0 | Cluster_05185 |
| 968.51104 | 35.0 | Cluster_15580 |
| 971.14245 | 35.0 | Cluster_15612 |
| 971.59682 | 35.0 | Cluster_40462 |
| 971.99294 | 35.0 | Cluster_40453 |
| 972.00212 | 35.0 | Cluster_40459 |
| 972.26876 | 35.0 | Cluster_05201 |
| 972.45860 | 35.0 | Cluster_15627 |
| 972.48455 | 35.0 | Cluster_15634 |
| 973.26231 | 35.0 | Cluster_40484 |

| | | |
|---|---|---|
| 975.16800 | 35.0 | Cluster_15670 |
| 977.10216 | 35.0 | Cluster_15692 |
| 977.15613 | 35.0 | Cluster_15689 |
| 978.75156 | 35.0 | Cluster_05224 |
| 979.02301 | 35.0 | Cluster_40541 |
| 979.49673 | 35.0 | Cluster_15713 |
| 980.13256 | 35.0 | Cluster_15723 |
| 981.41397 | 35.0 | Cluster_40580 |
| 981.44889 | 35.0 | Cluster_15750 |
| 981.45712 | 35.0 | Cluster_15741 |
| 981.45924 | 35.0 | Cluster_15747 |
| 982.13387 | 35.0 | Cluster_15752 |
| 983.47704 | 35.0 | Cluster_40610 |
| 984.15760 | 35.0 | Cluster_15784 |
| 985.95900 | 35.0 | Cluster_40633 |
| 986.82341 | 35.0 | Cluster_15809 |
| 987.47846 | 35.0 | Cluster_40647 |
| 988.49825 | 35.0 | Cluster_15833 |
| 988.94776 | 35.0 | Cluster_40676 |
| 990.42795 | 35.0 | Cluster_40698 |
| 990.48099 | 35.0 | Cluster_15858 |
| 991.15972 | 35.0 | Cluster_15864 |
| 991.47567 | 35.0 | Cluster_15867 |
| 991.97924 | 35.0 | Cluster_05270 |
| 992.48241 | 35.0 | Cluster_40726 |
| 992.74985 | 35.0 | Cluster_05272 |
| 992.82102 | 35.0 | Cluster_15885 |
| 993.01345 | 35.0 | Cluster_40733 |
| 994.14517 | 35.0 | Cluster_15902 |
| 996.00812 | 35.0 | Cluster_40774 |
| 996.70093 | 35.0 | Cluster_40783 |
| 996.86309 | 35.0 | Cluster_15945 |
| 997.76759 | 35.0 | Cluster_05282 |
| 997.96271 | 35.0 | Cluster_40814 |
| 998.87343 | 35.0 | Cluster_15967 |
| 998.87590 | 35.0 | Cluster_15968 |
| 999.99081 | 35.0 | Cluster_40861 |
| 1001.02474 | 35.0 | Cluster_40870 |
| 1003.12198 | 35.0 | Cluster_16013 |
| 1003.14396 | 35.0 | Cluster_16019 |
| 1003.15815 | 35.0 | Cluster_16017 |
| 1003.53342 | 35.0 | Cluster_40926 |
| 1006.19509 | 35.0 | Cluster_16058 |
| 1007.01591 | 35.0 | Cluster_40963 |
| 1007.54887 | 35.0 | Cluster_40979 |
| 1007.80918 | 35.0 | Cluster_16072 |
| 1007.96099 | 35.0 | Cluster_40980 |
| 1008.19150 | 35.0 | Cluster_16075 |
| 1009.53477 | 35.0 | Cluster_41016 |
| 1009.82372 | 35.0 | Cluster_16100 |
| 1012.00753 | 35.0 | Cluster_05327 |

| | | |
|---|---|---|
| 1013.18893 | 35.0 | Cluster_16139 |
| 1013.46270 | 35.0 | Cluster_16144 |
| 1014.19342 | 35.0 | Cluster_16147 |
| 1014.50348 | 35.0 | Cluster_41066 |
| 1015.75807 | 35.0 | Cluster_05350 |
| 1015.79694 | 35.0 | Cluster_16185 |
| 1015.91838 | 35.0 | Cluster_41088 |
| 1016.17395 | 35.0 | Cluster_16182 |
| 1016.48392 | 35.0 | Cluster_16187 |
| 1016.78489 | 35.0 | Cluster_05351 |
| 1017.91335 | 35.0 | Cluster_41107 |
| 1018.01928 | 35.0 | Cluster_41119 |
| 1018.03014 | 35.0 | Cluster_41116 |
| 1018.15185 | 35.0 | Cluster_16199 |
| 1018.27963 | 35.0 | Cluster_41121 |
| 1022.17658 | 35.0 | Cluster_16227 |
| 1022.18316 | 35.0 | Cluster_16229 |
| 1022.49717 | 35.0 | Cluster_16231 |
| 1023.01121 | 35.0 | Cluster_41172 |
| 1023.18114 | 35.0 | Cluster_16239 |
| 1023.19682 | 35.0 | Cluster_16242 |
| 1024.50414 | 35.0 | Cluster_16256 |
| 1024.92184 | 35.0 | Cluster_41190 |
| 1026.15295 | 35.0 | Cluster_16265 |
| 1027.88031 | 35.0 | Cluster_16291 |
| 1028.00760 | 35.0 | Cluster_05398 |
| 1028.51822 | 35.0 | Cluster_16304 |
| 1028.55896 | 35.0 | Cluster_41243 |
| 1028.82739 | 35.0 | Cluster_16303 |
| 1030.05344 | 35.0 | Cluster_41261 |
| 1030.51311 | 35.0 | Cluster_16316 |
| 1030.83122 | 35.0 | Cluster_16315 |
| 1031.52220 | 35.0 | Cluster_41282 |
| 1031.95890 | 35.0 | Cluster_41279 |
| 1032.06454 | 35.0 | Cluster_41290 |
| 1034.50686 | 35.0 | Cluster_41326 |
| 1035.51947 | 35.0 | Cluster_16366 |
| 1035.52766 | 35.0 | Cluster_16368 |
| 1035.88293 | 35.0 | Cluster_16378 |
| 1038.04978 | 35.0 | Cluster_41356 |
| 1038.52523 | 35.0 | Cluster_41369 |
| 1039.11544 | 35.0 | Cluster_41372 |
| 1039.52126 | 35.0 | Cluster_41374 |
| 1039.55516 | 35.0 | Cluster_41383 |
| 1041.01058 | 35.0 | Cluster_41401 |
| 1041.85584 | 35.0 | Cluster_16451 |
| 1042.51021 | 35.0 | Cluster_41413 |
| 1042.53113 | 35.0 | Cluster_16458 |
| 1045.27743 | 35.0 | Cluster_05445 |
| 1046.18482 | 35.0 | Cluster_16494 |
| 1046.22057 | 35.0 | Cluster_16495 |

| | | |
|---|---|---|
| 1046.52458 | 35.0 | Cluster_16503 |
| 1048.00284 | 35.0 | Cluster_41465 |
| 1048.97425 | 35.0 | Cluster_41478 |
| 1050.51799 | 35.0 | Cluster_16549 |
| 1052.08579 | 35.0 | Cluster_41500 |
| 1053.27199 | 35.0 | Cluster_41517 |
| 1053.47816 | 35.0 | Cluster_41529 |
| 1053.51090 | 35.0 | Cluster_16579 |
| 1053.87018 | 35.0 | Cluster_16587 |
| 1054.86259 | 35.0 | Cluster_16590 |
| 1054.86458 | 35.0 | Cluster_16593 |
| 1055.04277 | 35.0 | Cluster_41552 |
| 1055.21811 | 35.0 | Cluster_16595 |
| 1055.52224 | 35.0 | Cluster_16604 |
| 1056.84213 | 35.0 | Cluster_16627 |
| 1056.84310 | 35.0 | Cluster_16621 |
| 1056.86273 | 35.0 | Cluster_16631 |
| 1056.86682 | 35.0 | Cluster_16628 |
| 1056.98835 | 35.0 | Cluster_41572 |
| 1057.48459 | 35.0 | Cluster_41573 |
| 1057.99718 | 35.0 | Cluster_41576 |
| 1058.07318 | 35.0 | Cluster_41590 |
| 1058.82604 | 35.0 | Cluster_16656 |
| 1059.48627 | 35.0 | Cluster_41600 |
| 1061.05662 | 35.0 | Cluster_05509 |
| 1062.01230 | 35.0 | Cluster_41632 |
| 1062.51278 | 35.0 | Cluster_41641 |
| 1062.53761 | 35.0 | Cluster_41640 |
| 1062.94023 | 35.0 | Cluster_41638 |
| 1063.51637 | 35.0 | Cluster_16710 |
| 1064.19367 | 35.0 | Cluster_16719 |
| 1064.20364 | 35.0 | Cluster_16713 |
| 1064.26120 | 35.0 | Cluster_41669 |
| 1064.52753 | 35.0 | Cluster_41674 |
| 1064.82145 | 35.0 | Cluster_16722 |
| 1065.19208 | 35.0 | Cluster_16729 |
| 1065.54336 | 35.0 | Cluster_16738 |
| 1065.55172 | 35.0 | Cluster_16733 |
| 1065.88246 | 35.0 | Cluster_16740 |
| 1065.95736 | 35.0 | Cluster_00582 |
| 1066.21797 | 35.0 | Cluster_16744 |
| 1066.84581 | 35.0 | Cluster_16751 |
| 1068.74086 | 35.0 | Cluster_41735 |
| 1068.74952 | 35.0 | Cluster_41734 |
| 1069.06383 | 35.0 | Cluster_41739 |
| 1069.87675 | 35.0 | Cluster_16777 |
| 1070.03189 | 35.0 | Cluster_41745 |
| 1070.11798 | 35.0 | Cluster_41751 |
| 1070.86551 | 35.0 | Cluster_16790 |
| 1073.53750 | 35.0 | Cluster_41789 |
| 1074.52560 | 35.0 | Cluster_16830 |

| | | |
|---|---|---|
| 1075.19675 | 35.0 | Cluster_16840 |
| 1077.55035 | 35.0 | Cluster_16862 |
| 1077.55228 | 35.0 | Cluster_16863 |
| 1079.08133 | 35.0 | Cluster_41847 |
| 1079.58373 | 35.0 | Cluster_41846 |
| 1080.99907 | 35.0 | Cluster_41855 |
| 1081.04145 | 35.0 | Cluster_41859 |
| 1083.50260 | 35.0 | Cluster_05592 |
| 1083.60866 | 35.0 | Cluster_41894 |
| 1085.05222 | 35.0 | Cluster_41919 |
| 1086.49351 | 35.0 | Cluster_41934 |
| 1087.03798 | 35.0 | Cluster_41945 |
| 1089.06441 | 35.0 | Cluster_41967 |
| 1093.87964 | 35.0 | Cluster_17034 |
| 1098.22930 | 35.0 | Cluster_17071 |
| 1098.80067 | 35.0 | Cluster_42074 |
| 1098.88169 | 35.0 | Cluster_17084 |
| 1099.13859 | 35.0 | Cluster_00634 |
| 1103.12066 | 35.0 | Cluster_42104 |
| 1103.50753 | 35.0 | Cluster_42106 |
| 1103.54965 | 35.0 | Cluster_00644 |
| 1105.53482 | 35.0 | Cluster_17143 |
| 1105.55729 | 35.0 | Cluster_17144 |
| 1108.17135 | 35.0 | Cluster_17167 |
| 1108.90668 | 35.0 | Cluster_17181 |
| 1111.53648 | 35.0 | Cluster_17220 |
| 1113.59218 | 35.0 | Cluster_17234 |
| 1113.60048 | 35.0 | Cluster_17235 |
| 1114.57480 | 35.0 | Cluster_17251 |
| 1116.59135 | 35.0 | Cluster_42204 |
| 1117.22661 | 35.0 | Cluster_17271 |
| 1118.01598 | 35.0 | Cluster_42221 |
| 1118.08766 | 35.0 | Cluster_42224 |
| 1118.87403 | 35.0 | Cluster_17290 |
| 1121.01097 | 35.0 | Cluster_42253 |
| 1121.03337 | 35.0 | Cluster_42254 |
| 1122.55016 | 35.0 | Cluster_17335 |
| 1122.86660 | 35.0 | Cluster_17332 |
| 1123.53963 | 35.0 | Cluster_17344 |
| 1123.97198 | 35.0 | Cluster_42271 |
| 1124.00796 | 35.0 | Cluster_42274 |
| 1124.51758 | 35.0 | Cluster_42279 |
| 1124.52443 | 35.0 | Cluster_05696 |
| 1124.52521 | 35.0 | Cluster_17361 |
| 1125.56818 | 35.0 | Cluster_42283 |
| 1127.87407 | 35.0 | Cluster_17396 |
| 1129.16094 | 35.0 | Cluster_17421 |
| 1129.57408 | 35.0 | Cluster_42318 |
| 1130.53740 | 35.0 | Cluster_42327 |
| 1132.22675 | 35.0 | Cluster_17466 |
| 1136.05741 | 35.0 | Cluster_42359 |

| | | |
|---|---|---|
| 1137.19776 | 35.0 | Cluster_17532 |
| 1137.21620 | 35.0 | Cluster_17536 |
| 1137.22189 | 35.0 | Cluster_17533 |
| 1137.22952 | 35.0 | Cluster_17529 |
| 1138.87058 | 35.0 | Cluster_17553 |
| 1139.87279 | 35.0 | Cluster_17566 |
| 1139.92314 | 35.0 | Cluster_17563 |
| 1140.20245 | 35.0 | Cluster_17575 |
| 1141.54454 | 35.0 | Cluster_17592 |
| 1141.77200 | 35.0 | Cluster_00673 |
| 1141.89056 | 35.0 | Cluster_17593 |
| 1143.03536 | 35.0 | Cluster_42405 |
| 1143.25402 | 35.0 | Cluster_17604 |
| 1143.54127 | 35.0 | Cluster_17610 |
| 1144.55115 | 35.0 | Cluster_17624 |
| 1144.55511 | 35.0 | Cluster_17632 |
| 1144.86522 | 35.0 | Cluster_17636 |
| 1145.57649 | 35.0 | Cluster_05763 |
| 1147.10296 | 35.0 | Cluster_42440 |
| 1147.85107 | 35.0 | Cluster_17678 |
| 1150.20248 | 35.0 | Cluster_17718 |
| 1151.04837 | 35.0 | Cluster_42474 |
| 1151.06782 | 35.0 | Cluster_42470 |
| 1152.20245 | 35.0 | Cluster_17734 |
| 1152.54176 | 35.0 | Cluster_17735 |
| 1152.57782 | 35.0 | Cluster_17744 |
| 1153.03400 | 35.0 | Cluster_05790 |
| 1153.87912 | 35.0 | Cluster_17753 |
| 1153.88976 | 35.0 | Cluster_17763 |
| 1155.08158 | 35.0 | Cluster_42492 |
| 1155.24348 | 35.0 | Cluster_17775 |
| 1155.59409 | 35.0 | Cluster_17779 |
| 1155.89824 | 35.0 | Cluster_17801 |
| 1156.88520 | 35.0 | Cluster_17793 |
| 1161.58506 | 35.0 | Cluster_42549 |
| 1163.55909 | 35.0 | Cluster_17884 |
| 1166.21448 | 35.0 | Cluster_17894 |
| 1168.08892 | 35.0 | Cluster_42606 |
| 1168.58026 | 35.0 | Cluster_05832 |
| 1172.61481 | 35.0 | Cluster_17968 |
| 1173.06914 | 35.0 | Cluster_42645 |
| 1176.56192 | 35.0 | Cluster_42661 |
| 1177.58775 | 35.0 | Cluster_42668 |
| 1177.59766 | 35.0 | Cluster_42670 |
| 1178.92015 | 35.0 | Cluster_18022 |
| 1182.12534 | 35.0 | Cluster_42707 |
| 1182.58890 | 35.0 | Cluster_18037 |
| 1183.23440 | 35.0 | Cluster_18040 |
| 1183.54983 | 35.0 | Cluster_18043 |
| 1184.10037 | 35.0 | Cluster_42721 |
| 1187.11937 | 35.0 | Cluster_05925 |

| | | |
|---|---|---|
| 1187.49288 | 35.0 | Cluster_42749 |
| 1187.51857 | 35.0 | Cluster_42746 |
| 1187.59216 | 35.0 | Cluster_18075 |
| 1187.92731 | 35.0 | Cluster_18077 |
| 1187.92845 | 35.0 | Cluster_18076 |
| 1190.56370 | 35.0 | Cluster_18091 |
| 1192.59067 | 35.0 | Cluster_18107 |
| 1195.88339 | 35.0 | Cluster_18133 |
| 1198.06720 | 35.0 | Cluster_42794 |
| 1198.63811 | 35.0 | Cluster_42803 |
| 1200.25888 | 35.0 | Cluster_18168 |
| 1200.52867 | 35.0 | Cluster_18173 |
| 1200.56319 | 35.0 | Cluster_18177 |
| 1202.23828 | 35.0 | Cluster_18187 |
| 1202.23877 | 35.0 | Cluster_18194 |
| 1202.24327 | 35.0 | Cluster_18188 |
| 1204.59157 | 35.0 | Cluster_18204 |
| 1205.52768 | 35.0 | Cluster_42840 |
| 1206.89327 | 35.0 | Cluster_18228 |
| 1208.38799 | 35.0 | Cluster_42881 |
| 1208.56000 | 35.0 | Cluster_18237 |
| 1209.25140 | 35.0 | Cluster_18244 |
| 1214.21516 | 35.0 | Cluster_18272 |
| 1219.90840 | 35.0 | Cluster_18297 |
| 1221.61327 | 35.0 | Cluster_42961 |
| 1222.91304 | 35.0 | Cluster_18319 |
| 1224.59870 | 35.0 | Cluster_42983 |
| 1229.56319 | 35.0 | Cluster_43017 |
| 1230.62662 | 35.0 | Cluster_43026 |
| 1232.60076 | 35.0 | Cluster_43040 |
| 1233.11912 | 35.0 | Cluster_43048 |
| 1235.08055 | 35.0 | Cluster_43061 |
| 1235.57065 | 35.0 | Cluster_06173 |
| 1242.32292 | 35.0 | Cluster_18386 |
| 1242.56480 | 35.0 | Cluster_18385 |
| 1244.94841 | 35.0 | Cluster_18392 |
| 1248.07817 | 35.0 | Cluster_43132 |
| 1248.07817 | 35.0 | Cluster_43134 |
| 1249.82369 | 35.0 | Cluster_06249 |
| 1256.58258 | 35.0 | Cluster_06280 |
| 1256.84647 | 35.0 | Cluster_43180 |
| 1265.60966 | 35.0 | Cluster_43219 |
| 1267.20414 | 35.0 | Cluster_43224 |
| 1281.84230 | 35.0 | Cluster_06351 |
| 1287.13293 | 35.0 | Cluster_43302 |
| 1297.34487 | 35.0 | Cluster_18563 |
| 1297.57738 | 35.0 | Cluster_43344 |
| 1303.91742 | 35.0 | Cluster_06419 |
| 1309.16699 | 35.0 | Cluster_06433 |
| 1323.96017 | 35.0 | Cluster_06474 |
| 1324.59971 | 35.0 | Cluster_43461 |

| | | |
|---|---|---|
| 1333.16281 | 35.0 | Cluster_43496 |
| 1343.21645 | 35.0 | Cluster_43538 |
| 1353.42347 | 35.0 | Cluster_06570 |
| 1356.69696 | 35.0 | Cluster_43600 |
| 1356.70119 | 35.0 | Cluster_43599 |
| 1358.65593 | 35.0 | Cluster_06587 |
| 1358.66525 | 35.0 | Cluster_43610 |
| 1371.69886 | 35.0 | Cluster_43665 |
| 1372.75349 | 35.0 | Cluster_43667 |
| 1375.22316 | 35.0 | Cluster_43678 |
| 1382.71602 | 35.0 | Cluster_43695 |
| 1383.64043 | 35.0 | Cluster_43699 |
| 1404.23437 | 35.0 | Cluster_43777 |
| 1429.02179 | 35.0 | Cluster_18795 |
| 1442.73652 | 35.0 | Cluster_43921 |
| 1445.24833 | 35.0 | Cluster_43930 |
| 1464.40330 | 35.0 | Cluster_18820 |
| 1470.73553 | 35.0 | Cluster_43998 |
| 1473.26210 | 35.0 | Cluster_44004 |
| 1477.30369 | 35.0 | Cluster_44015 |
| 1478.73704 | 35.0 | Cluster_44019 |
| 1530.32978 | 35.0 | Cluster_44125 |
| 1564.32207 | 35.0 | Cluster_44184 |
| 1571.75675 | 35.0 | Cluster_44194 |
| 1616.85500 | 35.0 | Cluster_44250 |
| 1655.34019 | 35.0 | Cluster_44282 |
| 1689.84084 | 35.0 | Cluster_44356 |
| 1700.84338 | 35.0 | Cluster_44377 |
| 1739.94490 | 35.0 | Cluster_44403 |

Reject Mass List: (none)
Neutral Loss Mass List: (none)
Product Mass List: (none)
Neutral loss in top: 3
Product in top: 3
Most intense if no parent masses found not enabled
Add/subtract mass not enabled
FT master scan preview mode enabled
Charge state screening enabled
Charge state dependent ETD time not enabled
Monoisotopic precursor selection enabled
Charge state rejection enabled
Unassigned charge states : rejected
Charge state 1 : rejected
Charge state 2 : not rejected
Charge state 3 : not rejected
Charge states 4+ : not rejected
Chromatography mode is disabled

Global Data Dependent Settings:
Predict ion injection time enabled
Use global parent and reject mass lists not enabled
Exclude parent mass from data dependent selection not enabled

Exclusion mass width by mass
Exclusion mass width low: 1.50000
Exclusion mass width high: 1.50000
Parent mass width by mass
Parent mass width low: 0.50000
Parent mass width high: 0.50000
Reject mass width by mass
Reject mass width low: 0.50000
Reject mass width high: 0.50000
Zoom/UltraZoom scan mass width by mass
Zoom/UltraZoom scan mass width low: 5.00
Zoom/UltraZoom scan mass width high: 5.00
FT SIM scan mass width low: 5.00
FT SIM scan mass width high: 5.00
Neutral Loss candidates processed by decreasing intensity
Neutral Loss mass width by mass
Neutral Loss mass width low: 0.50000
Neutral Loss mass width high: 0.50000
Product candidates processed by decreasing intensity
Product mass width by mass
Product mass width low: 0.50000
Product mass width high: 0.50000
MS mass range: 300.00-2000.00
MSn mass range by mass
MSn mass range: 0.00-1000000.00
Use m/z values as masses not enabled
Analog UV data dep. not enabled
Dynamic exclusion enabled
Repeat Count: 1
Repeat Duration: 30.00
Exclusion List Size: 500
Exclusion Duration: 180.00
Exclusion mass width by mass
Exclusion mass width low: 1.50000
Exclusion mass width high: 1.50000
Expiration: disabled
Isotopic data dependence not enabled
Custom Data Dependent Settings:
Not enabled

---

Pass 7 (wheat-mixed-digests_FTMS1-ITMS2-only_inclusion-RT-01.raw):

Creator: Orbi_30393
Last modified: 10/20/2021 by Orbi_30393
MS Run Time (min): 43.00
Sequence override of method parameters not enabled.
Divert Valve: not used during run
Contact Closure: not used during run
Syringe Pump: not used during run
MS Detector Settings:
Real-time modifications to method not enabled

Stepped collision energy not enabled
Additional Microscans:
MS2 0 0
MS3 0 0
MS4 0 0
MS5 0 0
MS6 0 0
MS7 0 0
MS8 0 0
MS9 0 0
MS10 0 0
Segment 1 Information
Duration (min): 43.00
Number of Scan Events: 2
Tune Method: Orbitrap-tune-file_2020-03-13_HESI
Scan Event Details:
1: FTMS + p norm res=15000 o(300.0-2000.0)
CV = 0.0V
2: ITMS + c norm ·(list)->o(auto range)
Using global MS/MS mass lists
MS/MS: AT CID CE 35.0% Q 0.250 Time 10.000 IsoW 1.0
CV = 0.0V
Lock Masses:
Pos List Name: N/A
Source: API Source
Mass List: (none)
Neg List Name: N/A
Source: API Source
Mass List: (none)
Global Non-Data Dependent Settings:
(+) Global MS/MS Masses:

| MS Mass | Start (min) | End (min) | MS FAIMS CV | MS Normalized Collision Energy | Last Mass | Name |
|---|---|---|---|---|---|---|
| 300.13000 | 27.890 | 28.060 | | 35.0 | | Cluster_06844 |
| 300.14000 | 1.590 | 2.170 | | 35.0 | | Cluster_18928 |
| 300.71000 | 7.520 | 8.290 | | 35.0 | | Cluster_18955 |
| 301.50000 | 1.580 | 2.120 | | 35.0 | | Cluster_06851 |
| 301.65000 | 11.650 | 12.150 | | 35.0 | | Cluster_18984 |
| 302.64000 | 1.240 | 1.560 | | 35.0 | | Cluster_19016 |
| 302.65000 | 1.560 | 2.040 | | 35.0 | | Cluster_19017 |
| 303.99000 | 1.650 | 2.110 | | 35.0 | | Cluster_19040 |
| 304.11000 | 17.480 | 17.640 | | 35.0 | | Cluster_19042 |
| 304.14000 | 1.280 | 1.580 | | 35.0 | | Cluster_19049 |
| 304.16000 | 1.560 | 2.030 | | 35.0 | | Cluster_19050 |
| 304.20000 | 7.520 | 8.260 | | 35.0 | | Cluster_19052 |
| 304.49000 | 7.390 | 7.980 | | 35.0 | | Cluster_06863 |
| 304.64000 | 1.270 | 1.570 | | 35.0 | | Cluster_19062 |
| 306.13000 | 7.630 | 8.420 | | 35.0 | | Cluster_06873 |
| 306.67000 | 11.740 | 12.080 | | 35.0 | | Cluster_19109 |

| | | | | |
|---|---|---|---|---|
| 307.12000 | 19.890 | 20.520 | 35.0 | Cluster_19119 |
| 307.13000 | 7.640 | 8.390 | 35.0 | Cluster_06880 |
| 308.65000 | 1.480 | 1.910 | 35.0 | Cluster_19177 |
| 308.66000 | 7.530 | 8.280 | 35.0 | Cluster_19179 |
| 309.53000 | 7.580 | 8.060 | 35.0 | Cluster_06901 |
| 310.58000 | 17.390 | 18.040 | 35.0 | Cluster_19217 |
| 310.65000 | 7.630 | 7.900 | 35.0 | Cluster_19222 |
| 310.66000 | 1.580 | 2.020 | 35.0 | Cluster_19221 |
| 311.17000 | 7.570 | 8.330 | 35.0 | Cluster_19246 |
| 311.68000 | 7.770 | 8.930 | 35.0 | Cluster_19257 |
| 313.65000 | 1.240 | 1.630 | 35.0 | Cluster_19315 |
| 314.82000 | 1.460 | 1.880 | 35.0 | Cluster_06941 |
| 315.65000 | 1.540 | 2.000 | 35.0 | Cluster_19384 |
| 316.69000 | 11.690 | 12.070 | 35.0 | Cluster_19442 |
| 317.06000 | 11.640 | 11.860 | 35.0 | Cluster_19459 |
| 317.16000 | 1.470 | 2.010 | 35.0 | Cluster_19466 |
| 317.64000 | 14.650 | 15.060 | 35.0 | Cluster_19490 |
| 318.64000 | 1.530 | 1.960 | 35.0 | Cluster_19532 |
| 318.85000 | 7.830 | 9.360 | 35.0 | Cluster_06973 |
| 319.17000 | 1.260 | 1.540 | 35.0 | Cluster_19552 |
| 319.67000 | 1.530 | 2.040 | 35.0 | Cluster_19560 |
| 319.94000 | 7.730 | 8.090 | 35.0 | Cluster_00850 |
| 320.49000 | 1.510 | 2.130 | 35.0 | Cluster_06993 |
| 320.80000 | 11.560 | 12.030 | 35.0 | Cluster_06994 |
| 321.15000 | 1.560 | 1.880 | 35.0 | Cluster_06995 |
| 321.17000 | 14.840 | 15.780 | 35.0 | Cluster_00869 |
| 321.52000 | 7.600 | 8.110 | 35.0 | Cluster_19594 |
| 322.15000 | 19.880 | 20.020 | 35.0 | Cluster_19619 |
| 322.17000 | 1.490 | 1.910 | 35.0 | Cluster_19617 |
| 322.50000 | 1.590 | 2.130 | 35.0 | Cluster_06998 |
| 323.65000 | 1.230 | 1.590 | 35.0 | Cluster_19672 |
| 323.69000 | 19.910 | 20.420 | 35.0 | Cluster_19677 |
| 323.83000 | 7.450 | 7.800 | 35.0 | Cluster_19683 |
| 324.42000 | 11.820 | 12.680 | 35.0 | Cluster_00878 |
| 324.60000 | 11.610 | 11.830 | 35.0 | Cluster_19711 |
| 324.65000 | 1.530 | 1.960 | 35.0 | Cluster_19713 |
| 324.83000 | 11.690 | 12.010 | 35.0 | Cluster_07017 |
| 325.14000 | 7.400 | 7.970 | 35.0 | Cluster_19730 |
| 325.82000 | 1.570 | 2.110 | 35.0 | Cluster_07023 |
| 325.85000 | 11.630 | 12.670 | 35.0 | Cluster_07028 |
| 326.17000 | 7.780 | 8.310 | 35.0 | Cluster_19753 |
| 326.40000 | 1.640 | 2.050 | 35.0 | Cluster_00887 |
| 326.90000 | 7.400 | 7.730 | 35.0 | Cluster_00888 |
| 327.21000 | 14.670 | 15.080 | 35.0 | Cluster_19791 |
| 327.96000 | 7.700 | 8.060 | 35.0 | Cluster_07043 |
| 328.48000 | 1.240 | 1.570 | 35.0 | Cluster_07051 |
| 329.18000 | 1.520 | 2.030 | 35.0 | Cluster_19855 |
| 329.82000 | 11.770 | 12.160 | 35.0 | Cluster_07063 |
| 329.93000 | 1.230 | 1.630 | 35.0 | Cluster_19873 |
| 330.18000 | 14.850 | 15.020 | 35.0 | Cluster_19886 |
| 330.90000 | 1.650 | 2.040 | 35.0 | Cluster_19916 |

| | | | | |
|---|---|---|---|---|
| 331.17000 | 1.560 | 2.070 | 35.0 | Cluster_19926 |
| 331.20000 | 7.320 | 7.700 | 35.0 | Cluster_19928 |
| 331.21000 | 7.550 | 7.760 | 35.0 | Cluster_19930 |
| 331.22000 | 17.390 | 17.790 | 35.0 | Cluster_19925 |
| 331.66000 | 1.240 | 1.580 | 35.0 | Cluster_19936 |
| 332.14000 | 1.450 | 1.820 | 35.0 | Cluster_19948 |
| 333.47000 | 1.650 | 2.010 | 35.0 | Cluster_07081 |
| 333.82000 | 17.380 | 17.810 | 35.0 | Cluster_07087 |
| 333.87000 | 7.640 | 8.170 | 35.0 | Cluster_07090 |
| 335.12000 | 7.740 | 8.570 | 35.0 | Cluster_20034 |
| 335.83000 | 1.230 | 1.640 | 35.0 | Cluster_07104 |
| 336.19000 | 7.470 | 7.920 | 35.0 | Cluster_07109 |
| 336.23000 | 11.790 | 12.070 | 35.0 | Cluster_20093 |
| 336.91000 | 7.390 | 7.630 | 35.0 | Cluster_00920 |
| 337.19000 | 19.960 | 20.100 | 35.0 | Cluster_20132 |
| 337.21000 | 11.720 | 12.080 | 35.0 | Cluster_20126 |
| 337.83000 | 11.850 | 12.060 | 35.0 | Cluster_07126 |
| 338.11000 | 7.370 | 7.610 | 35.0 | Cluster_20150 |
| 338.67000 | 7.620 | 8.030 | 35.0 | Cluster_20172 |
| 339.16000 | 1.450 | 1.910 | 35.0 | Cluster_20192 |
| 339.47000 | 1.440 | 2.030 | 35.0 | Cluster_07139 |
| 339.81000 | 14.710 | 15.090 | 35.0 | Cluster_07150 |
| 339.94000 | 7.660 | 8.070 | 35.0 | Cluster_00929 |
| 340.18000 | 19.890 | 20.250 | 35.0 | Cluster_20223 |
| 340.26000 | 22.630 | 22.970 | 35.0 | Cluster_20229 |
| 341.15000 | 1.270 | 1.550 | 35.0 | Cluster_20257 |
| 341.16000 | 1.650 | 2.020 | 35.0 | Cluster_20261 |
| 342.16000 | 1.620 | 2.020 | 35.0 | Cluster_07167 |
| 343.11000 | 7.800 | 8.560 | 35.0 | Cluster_20316 |
| 343.18000 | 7.750 | 8.420 | 35.0 | Cluster_20318 |
| 343.66000 | 1.250 | 1.550 | 35.0 | Cluster_20344 |
| 345.19000 | 14.850 | 15.320 | 35.0 | Cluster_20390 |
| 345.67000 | 1.450 | 1.780 | 35.0 | Cluster_20402 |
| 346.15000 | 17.420 | 18.290 | 35.0 | Cluster_07199 |
| 346.67000 | 7.760 | 8.250 | 35.0 | Cluster_20444 |
| 347.20000 | 7.310 | 7.800 | 35.0 | Cluster_20457 |
| 347.71000 | 7.760 | 8.410 | 35.0 | Cluster_20471 |
| 348.44000 | 19.970 | 20.260 | 35.0 | Cluster_20495 |
| 348.87000 | 7.420 | 7.900 | 35.0 | Cluster_07234 |
| 348.92000 | 14.640 | 14.870 | 35.0 | Cluster_00975 |
| 348.93000 | 14.860 | 15.120 | 35.0 | Cluster_00976 |
| 349.20000 | 7.770 | 8.090 | 35.0 | Cluster_20517 |
| 349.21000 | 11.780 | 12.250 | 35.0 | Cluster_20520 |
| 349.64000 | 11.600 | 11.900 | 35.0 | Cluster_20543 |
| 349.73000 | 11.610 | 11.990 | 35.0 | Cluster_20549 |
| 350.42000 | 14.810 | 15.540 | 35.0 | Cluster_00982 |
| 351.18000 | 1.240 | 1.560 | 35.0 | Cluster_20611 |
| 351.19000 | 17.520 | 17.990 | 35.0 | Cluster_20622 |
| 352.66000 | 11.800 | 12.220 | 35.0 | Cluster_20669 |
| 353.17000 | 14.630 | 15.020 | 35.0 | Cluster_01002 |
| 353.21000 | 7.750 | 8.140 | 35.0 | Cluster_20705 |

| | | | | |
|---|---|---|---|---|
| 353.66000 | 1.540 | 2.210 | 35.0 | Cluster_01003 |
| 353.80000 | 19.900 | 20.390 | 35.0 | Cluster_07278 |
| 353.87000 | 7.510 | 7.740 | 35.0 | Cluster_20720 |
| 353.91000 | 1.610 | 2.100 | 35.0 | Cluster_01004 |
| 354.16000 | 1.610 | 2.040 | 35.0 | Cluster_20732 |
| 354.20000 | 11.790 | 12.210 | 35.0 | Cluster_20739 |
| 354.40000 | 7.680 | 8.720 | 35.0 | Cluster_01009 |
| 354.42000 | 11.860 | 12.480 | 35.0 | Cluster_01011 |
| 355.14000 | 1.230 | 1.620 | 35.0 | Cluster_20781 |
| 355.16000 | 1.450 | 1.850 | 35.0 | Cluster_20782 |
| 355.22000 | 7.500 | 7.920 | 35.0 | Cluster_20787 |
| 355.36000 | 1.330 | 1.600 | 35.0 | Cluster_20789 |
| 355.70000 | 1.440 | 2.070 | 35.0 | Cluster_20796 |
| 356.24000 | 14.870 | 15.320 | 35.0 | Cluster_20824 |
| 356.55000 | 14.880 | 16.740 | 35.0 | Cluster_07307 |
| 358.42000 | 1.590 | 2.110 | 35.0 | Cluster_20933 |
| 359.13000 | 19.930 | 20.340 | 35.0 | Cluster_07326 |
| 359.17000 | 11.670 | 12.360 | 35.0 | Cluster_00118 |
| 359.37000 | 11.560 | 12.150 | 35.0 | Cluster_20960 |
| 360.16000 | 1.260 | 1.450 | 35.0 | Cluster_20986 |
| 360.42000 | 11.840 | 12.330 | 35.0 | Cluster_01047 |
| 360.83000 | 7.800 | 8.000 | 35.0 | Cluster_21008 |
| 361.18000 | 11.800 | 12.350 | 35.0 | Cluster_21024 |
| 361.42000 | 7.620 | 8.490 | 35.0 | Cluster_01055 |
| 361.66000 | 1.430 | 2.060 | 35.0 | Cluster_21038 |
| 361.67000 | 11.800 | 12.310 | 35.0 | Cluster_21036 |
| 361.69000 | 7.390 | 7.940 | 35.0 | Cluster_21039 |
| 361.84000 | 7.750 | 8.230 | 35.0 | Cluster_07347 |
| 362.10000 | 1.550 | 1.980 | 35.0 | Cluster_21048 |
| 362.42000 | 17.390 | 17.620 | 35.0 | Cluster_01059 |
| 362.43000 | 7.800 | 8.230 | 35.0 | Cluster_21075 |
| 362.60000 | 1.520 | 2.040 | 35.0 | Cluster_21077 |
| 362.65000 | 1.280 | 1.550 | 35.0 | Cluster_21076 |
| 363.11000 | 1.580 | 2.030 | 35.0 | Cluster_21089 |
| 363.19000 | 11.880 | 12.080 | 35.0 | Cluster_21093 |
| 363.20000 | 7.390 | 8.120 | 35.0 | Cluster_21094 |
| 363.77000 | 17.520 | 17.890 | 35.0 | Cluster_21121 |
| 364.16000 | 11.580 | 12.110 | 35.0 | Cluster_21135 |
| 364.50000 | 1.550 | 1.900 | 35.0 | Cluster_21161 |
| 364.65000 | 19.910 | 20.370 | 35.0 | Cluster_21162 |
| 365.24000 | 17.500 | 17.690 | 35.0 | Cluster_21221 |
| 365.79000 | 11.730 | 12.200 | 35.0 | Cluster_07396 |
| 365.90000 | 1.310 | 1.630 | 35.0 | Cluster_21241 |
| 366.27000 | 7.710 | 8.040 | 35.0 | Cluster_21267 |
| 366.74000 | 11.800 | 12.060 | 35.0 | Cluster_21285 |
| 367.22000 | 14.610 | 15.000 | 35.0 | Cluster_21306 |
| 367.50000 | 11.840 | 12.340 | 35.0 | Cluster_07416 |
| 367.53000 | 11.850 | 12.350 | 35.0 | Cluster_21324 |
| 368.16000 | 11.830 | 12.240 | 35.0 | Cluster_21368 |
| 368.20000 | 7.600 | 8.110 | 35.0 | Cluster_21366 |
| 368.47000 | 7.520 | 8.860 | 35.0 | Cluster_07428 |

| | | | | |
|---|---|---|---|---|
| 369.67000 | 1.240 | 1.570 | 35.0 | Cluster_21437 |
| 370.08000 | 1.590 | 1.940 | 35.0 | Cluster_21453 |
| 370.50000 | 11.840 | 12.340 | 35.0 | Cluster_07445 |
| 370.74000 | 22.670 | 23.120 | 35.0 | Cluster_21476 |
| 370.93000 | 14.750 | 14.960 | 35.0 | Cluster_21486 |
| 371.18000 | 1.570 | 1.930 | 35.0 | Cluster_07449 |
| 371.24000 | 11.560 | 11.900 | 35.0 | Cluster_21501 |
| 371.71000 | 17.530 | 17.830 | 35.0 | Cluster_21517 |
| 371.89000 | 1.270 | 1.470 | 35.0 | Cluster_21531 |
| 372.17000 | 7.580 | 7.970 | 35.0 | Cluster_21537 |
| 372.23000 | 11.560 | 12.260 | 35.0 | Cluster_07464 |
| 372.67000 | 20.000 | 20.220 | 35.0 | Cluster_21575 |
| 372.69000 | 1.240 | 1.640 | 35.0 | Cluster_21572 |
| 373.12000 | 1.600 | 1.800 | 35.0 | Cluster_21589 |
| 373.14000 | 20.040 | 20.290 | 35.0 | Cluster_21585 |
| 373.16000 | 14.870 | 15.050 | 35.0 | Cluster_07470 |
| 373.19000 | 7.420 | 8.060 | 35.0 | Cluster_07471 |
| 373.22000 | 22.670 | 23.130 | 35.0 | Cluster_21597 |
| 373.90000 | 14.600 | 14.750 | 35.0 | Cluster_21628 |
| 374.19000 | 7.480 | 8.090 | 35.0 | Cluster_07480 |
| 374.40000 | 1.250 | 1.670 | 35.0 | Cluster_21650 |
| 374.68000 | 14.750 | 15.180 | 35.0 | Cluster_21656 |
| 375.15000 | 1.650 | 2.000 | 35.0 | Cluster_21668 |
| 375.89000 | 17.460 | 17.840 | 35.0 | Cluster_07496 |
| 377.48000 | 14.870 | 15.440 | 35.0 | Cluster_07512 |
| 377.77000 | 14.780 | 14.930 | 35.0 | Cluster_21780 |
| 378.17000 | 11.710 | 12.170 | 35.0 | Cluster_01122 |
| 378.18000 | 1.230 | 1.600 | 35.0 | Cluster_21788 |
| 378.20000 | 11.840 | 12.330 | 35.0 | Cluster_07523 |
| 378.67000 | 14.750 | 14.950 | 35.0 | Cluster_01125 |
| 379.26000 | 14.750 | 15.010 | 35.0 | Cluster_21841 |
| 380.45000 | 7.770 | 8.430 | 35.0 | Cluster_21886 |
| 380.68000 | 14.810 | 15.050 | 35.0 | Cluster_21891 |
| 381.22000 | 17.270 | 17.840 | 35.0 | Cluster_07550 |
| 381.70000 | 7.800 | 8.490 | 35.0 | Cluster_01141 |
| 382.18000 | 1.540 | 1.730 | 35.0 | Cluster_21953 |
| 382.66000 | 14.630 | 15.040 | 35.0 | Cluster_01144 |
| 383.63000 | 19.940 | 20.250 | 35.0 | Cluster_22024 |
| 383.68000 | 7.470 | 8.030 | 35.0 | Cluster_22025 |
| 384.16000 | 14.730 | 15.060 | 35.0 | Cluster_01146 |
| 384.21000 | 22.650 | 22.920 | 35.0 | Cluster_22055 |
| 384.22000 | 7.530 | 7.780 | 35.0 | Cluster_22065 |
| 385.52000 | 22.480 | 22.910 | 35.0 | Cluster_07597 |
| 385.54000 | 31.410 | 31.800 | 35.0 | Cluster_07596 |
| 385.86000 | 27.960 | 28.270 | 35.0 | Cluster_22123 |
| 386.66000 | 20.010 | 20.720 | 35.0 | Cluster_22155 |
| 386.68000 | 14.620 | 14.820 | 35.0 | Cluster_22158 |
| 386.90000 | 14.610 | 14.880 | 35.0 | Cluster_22172 |
| 387.11000 | 11.820 | 12.000 | 35.0 | Cluster_22177 |
| 387.63000 | 17.350 | 17.770 | 35.0 | Cluster_22217 |
| 387.93000 | 7.700 | 8.280 | 35.0 | Cluster_22226 |

| | | | | |
|---|---|---|---|---|
| 388.23000 | 11.660 | 12.060 | 35.0 | Cluster_22233 |
| 388.50000 | 11.630 | 12.080 | 35.0 | Cluster_07621 |
| 388.64000 | 14.810 | 15.110 | 35.0 | Cluster_22244 |
| 388.67000 | 19.930 | 20.200 | 35.0 | Cluster_22245 |
| 388.83000 | 22.620 | 22.930 | 35.0 | Cluster_07623 |
| 389.25000 | 17.510 | 18.000 | 35.0 | Cluster_22269 |
| 389.51000 | 14.810 | 15.110 | 35.0 | Cluster_07627 |
| 389.91000 | 14.830 | 14.940 | 35.0 | Cluster_22284 |
| 390.71000 | 14.680 | 15.090 | 35.0 | Cluster_22316 |
| 390.87000 | 31.450 | 31.800 | 35.0 | Cluster_07651 |
| 390.88000 | 7.800 | 9.140 | 35.0 | Cluster_07646 |
| 391.50000 | 1.450 | 2.120 | 35.0 | Cluster_07654 |
| 391.65000 | 19.880 | 20.230 | 35.0 | Cluster_22334 |
| 391.75000 | 11.860 | 12.080 | 35.0 | Cluster_22344 |
| 391.82000 | 7.470 | 7.980 | 35.0 | Cluster_07658 |
| 392.22000 | 14.780 | 15.190 | 35.0 | Cluster_07665 |
| 392.25000 | 7.750 | 8.230 | 35.0 | Cluster_22358 |
| 392.73000 | 7.380 | 7.780 | 35.0 | Cluster_22381 |
| 392.77000 | 11.560 | 11.910 | 35.0 | Cluster_22377 |
| 393.19000 | 14.820 | 15.160 | 35.0 | Cluster_01195 |
| 393.53000 | 11.840 | 12.050 | 35.0 | Cluster_07674 |
| 393.69000 | 14.830 | 15.060 | 35.0 | Cluster_01197 |
| 393.71000 | 1.590 | 2.000 | 35.0 | Cluster_22428 |
| 393.98000 | 17.500 | 17.660 | 35.0 | Cluster_22479 |
| 394.21000 | 25.240 | 25.400 | 35.0 | Cluster_22490 |
| 394.71000 | 14.870 | 15.050 | 35.0 | Cluster_22522 |
| 394.72000 | 11.850 | 11.980 | 35.0 | Cluster_22521 |
| 395.89000 | 11.840 | 12.350 | 35.0 | Cluster_07703 |
| 396.23000 | 19.970 | 20.450 | 35.0 | Cluster_22589 |
| 396.30000 | 22.510 | 22.690 | 35.0 | Cluster_22595 |
| 397.39000 | 25.090 | 26.350 | 35.0 | Cluster_22628 |
| 397.67000 | 1.530 | 1.910 | 35.0 | Cluster_22634 |
| 397.72000 | 7.580 | 8.070 | 35.0 | Cluster_22641 |
| 397.90000 | 14.840 | 15.260 | 35.0 | Cluster_07736 |
| 398.18000 | 17.310 | 17.770 | 35.0 | Cluster_01226 |
| 398.44000 | 17.280 | 17.770 | 35.0 | Cluster_01230 |
| 400.24000 | 14.690 | 14.900 | 35.0 | Cluster_22747 |
| 400.68000 | 22.560 | 22.770 | 35.0 | Cluster_22780 |
| 400.90000 | 20.030 | 20.230 | 35.0 | Cluster_07771 |
| 401.70000 | 7.350 | 7.800 | 35.0 | Cluster_22833 |
| 402.32000 | 7.400 | 7.580 | 35.0 | Cluster_22866 |
| 402.48000 | 11.860 | 12.560 | 35.0 | Cluster_22868 |
| 402.72000 | 11.830 | 11.990 | 35.0 | Cluster_22878 |
| 403.44000 | 1.560 | 1.910 | 35.0 | Cluster_01260 |
| 403.54000 | 28.070 | 28.440 | 35.0 | Cluster_07797 |
| 404.47000 | 14.760 | 15.050 | 35.0 | Cluster_01272 |
| 405.88000 | 25.070 | 25.350 | 35.0 | Cluster_07831 |
| 406.22000 | 22.600 | 23.040 | 35.0 | Cluster_23023 |
| 406.68000 | 1.270 | 1.540 | 35.0 | Cluster_01285 |
| 406.73000 | 20.010 | 20.210 | 35.0 | Cluster_23046 |
| 406.95000 | 1.450 | 2.070 | 35.0 | Cluster_01293 |

| | | | | |
|---|---|---|---|---|
| 407.17000 | 7.360 | 7.710 | 35.0 | Cluster_23059 |
| 407.25000 | 11.740 | 12.110 | 35.0 | Cluster_23066 |
| 407.67000 | 11.640 | 12.000 | 35.0 | Cluster_01302 |
| 407.75000 | 14.850 | 15.150 | 35.0 | Cluster_23083 |
| 407.93000 | 1.570 | 2.110 | 35.0 | Cluster_01303 |
| 408.22000 | 14.810 | 14.990 | 35.0 | Cluster_23104 |
| 408.54000 | 11.800 | 12.160 | 35.0 | Cluster_07863 |
| 408.66000 | 17.290 | 17.790 | 35.0 | Cluster_01305 |
| 408.91000 | 7.750 | 8.310 | 35.0 | Cluster_07864 |
| 408.92000 | 20.020 | 20.430 | 35.0 | Cluster_01307 |
| 409.19000 | 1.560 | 1.970 | 35.0 | Cluster_07867 |
| 409.25000 | 11.580 | 11.910 | 35.0 | Cluster_23160 |
| 409.69000 | 1.240 | 1.530 | 35.0 | Cluster_23170 |
| 410.53000 | 17.390 | 17.810 | 35.0 | Cluster_07882 |
| 411.72000 | 11.630 | 12.000 | 35.0 | Cluster_23258 |
| 412.67000 | 1.440 | 1.970 | 35.0 | Cluster_23299 |
| 412.68000 | 11.850 | 12.330 | 35.0 | Cluster_01326 |
| 413.23000 | 11.660 | 11.900 | 35.0 | Cluster_23320 |
| 413.87000 | 1.590 | 2.080 | 35.0 | Cluster_07913 |
| 413.92000 | 19.920 | 20.450 | 35.0 | Cluster_07916 |
| 413.94000 | 17.500 | 17.750 | 35.0 | Cluster_23350 |
| 413.98000 | 11.740 | 12.090 | 35.0 | Cluster_23354 |
| 414.71000 | 1.530 | 2.160 | 35.0 | Cluster_23379 |
| 414.85000 | 1.610 | 2.210 | 35.0 | Cluster_23397 |
| 414.87000 | 14.710 | 14.970 | 35.0 | Cluster_07924 |
| 415.22000 | 1.230 | 1.480 | 35.0 | Cluster_23422 |
| 415.73000 | 22.500 | 23.100 | 35.0 | Cluster_23434 |
| 416.00000 | 14.750 | 15.180 | 35.0 | Cluster_23448 |
| 416.22000 | 7.630 | 7.950 | 35.0 | Cluster_23463 |
| 416.76000 | 7.470 | 7.800 | 35.0 | Cluster_23476 |
| 416.88000 | 1.230 | 1.600 | 35.0 | Cluster_23482 |
| 416.94000 | 17.320 | 17.820 | 35.0 | Cluster_01353 |
| 418.25000 | 17.280 | 17.520 | 35.0 | Cluster_23534 |
| 418.47000 | 7.650 | 7.880 | 35.0 | Cluster_23549 |
| 419.18000 | 17.330 | 17.880 | 35.0 | Cluster_01360 |
| 419.21000 | 7.510 | 7.930 | 35.0 | Cluster_07978 |
| 419.88000 | 1.270 | 1.430 | 35.0 | Cluster_23600 |
| 420.19000 | 11.580 | 11.880 | 35.0 | Cluster_23617 |
| 420.23000 | 14.640 | 15.400 | 35.0 | Cluster_23612 |
| 420.71000 | 7.310 | 8.340 | 35.0 | Cluster_23632 |
| 421.56000 | 7.310 | 8.090 | 35.0 | Cluster_08005 |
| 422.12000 | 1.580 | 2.070 | 35.0 | Cluster_08008 |
| 422.55000 | 31.510 | 31.860 | 35.0 | Cluster_08014 |
| 422.58000 | 25.320 | 25.840 | 35.0 | Cluster_08015 |
| 423.14000 | 11.570 | 12.000 | 35.0 | Cluster_23727 |
| 423.21000 | 14.870 | 15.370 | 35.0 | Cluster_08022 |
| 423.24000 | 11.740 | 12.070 | 35.0 | Cluster_23740 |
| 423.44000 | 17.530 | 18.010 | 35.0 | Cluster_01387 |
| 423.71000 | 27.930 | 28.180 | 35.0 | Cluster_23753 |
| 423.74000 | 22.510 | 23.020 | 35.0 | Cluster_23766 |
| 424.18000 | 7.300 | 7.920 | 35.0 | Cluster_23787 |

| | | | | |
|---|---|---|---|---|
| 424.51000 | 11.730 | 12.470 | 35.0 | Cluster_23811 |
| 424.68000 | 11.820 | 12.260 | 35.0 | Cluster_23813 |
| 425.18000 | 7.350 | 7.860 | 35.0 | Cluster_23850 |
| 425.46000 | 11.660 | 11.990 | 35.0 | Cluster_23865 |
| 425.87000 | 7.600 | 8.080 | 35.0 | Cluster_08056 |
| 425.88000 | 1.340 | 1.550 | 35.0 | Cluster_23895 |
| 426.14000 | 7.470 | 7.920 | 35.0 | Cluster_23901 |
| 426.24000 | 25.190 | 25.440 | 35.0 | Cluster_23924 |
| 426.44000 | 7.530 | 7.670 | 35.0 | Cluster_23942 |
| 426.88000 | 11.830 | 12.340 | 35.0 | Cluster_08077 |
| 427.51000 | 17.320 | 17.760 | 35.0 | Cluster_08094 |
| 428.56000 | 14.860 | 15.120 | 35.0 | Cluster_08107 |
| 428.72000 | 7.840 | 8.690 | 35.0 | Cluster_24045 |
| 428.79000 | 17.280 | 17.930 | 35.0 | Cluster_24059 |
| 429.24000 | 7.630 | 7.910 | 35.0 | Cluster_24074 |
| 429.57000 | 22.600 | 23.060 | 35.0 | Cluster_08120 |
| 429.72000 | 14.590 | 14.880 | 35.0 | Cluster_24100 |
| 429.76000 | 7.420 | 7.660 | 35.0 | Cluster_24093 |
| 429.85000 | 22.580 | 22.890 | 35.0 | Cluster_08124 |
| 429.95000 | 17.400 | 17.830 | 35.0 | Cluster_01432 |
| 430.73000 | 27.890 | 28.160 | 35.0 | Cluster_24136 |
| 431.21000 | 1.610 | 1.860 | 35.0 | Cluster_24172 |
| 431.41000 | 17.290 | 17.520 | 35.0 | Cluster_24179 |
| 431.74000 | 14.710 | 14.950 | 35.0 | Cluster_24194 |
| 432.16000 | 11.860 | 12.140 | 35.0 | Cluster_24207 |
| 432.21000 | 11.630 | 11.940 | 35.0 | Cluster_24215 |
| 432.87000 | 14.770 | 14.990 | 35.0 | Cluster_08164 |
| 433.96000 | 1.640 | 1.960 | 35.0 | Cluster_24289 |
| 434.79000 | 17.290 | 17.460 | 35.0 | Cluster_24347 |
| 434.97000 | 17.370 | 17.820 | 35.0 | Cluster_24350 |
| 435.17000 | 11.710 | 11.980 | 35.0 | Cluster_24357 |
| 435.24000 | 14.610 | 15.010 | 35.0 | Cluster_08201 |
| 435.37000 | 25.040 | 25.410 | 35.0 | Cluster_24383 |
| 435.70000 | 1.590 | 2.060 | 35.0 | Cluster_01481 |
| 435.97000 | 17.270 | 18.110 | 35.0 | Cluster_01487 |
| 436.48000 | 22.610 | 22.960 | 35.0 | Cluster_01491 |
| 437.37000 | 7.350 | 7.700 | 35.0 | Cluster_24491 |
| 437.46000 | 14.860 | 15.410 | 35.0 | Cluster_01497 |
| 438.22000 | 7.510 | 8.020 | 35.0 | Cluster_24532 |
| 438.30000 | 20.000 | 20.340 | 35.0 | Cluster_24543 |
| 438.96000 | 17.260 | 17.670 | 35.0 | Cluster_01503 |
| 438.97000 | 22.600 | 22.960 | 35.0 | Cluster_01504 |
| 439.82000 | 7.670 | 8.890 | 35.0 | Cluster_24632 |
| 440.21000 | 14.830 | 15.150 | 35.0 | Cluster_01512 |
| 440.56000 | 7.700 | 7.920 | 35.0 | Cluster_24655 |
| 440.74000 | 7.450 | 7.710 | 35.0 | Cluster_24659 |
| 440.86000 | 1.610 | 2.060 | 35.0 | Cluster_08277 |
| 441.21000 | 14.690 | 14.880 | 35.0 | Cluster_24682 |
| 441.24000 | 25.130 | 25.470 | 35.0 | Cluster_01516 |
| 441.73000 | 7.710 | 7.920 | 35.0 | Cluster_24703 |
| 441.87000 | 11.740 | 12.340 | 35.0 | Cluster_08293 |

| | | | | |
|---|---|---|---|---|
| 442.00000 | 14.790 | 14.990 | 35.0 | Cluster_24716 |
| 442.28000 | 14.810 | 15.170 | 35.0 | Cluster_24730 |
| 442.57000 | 14.840 | 15.230 | 35.0 | Cluster_08303 |
| 442.77000 | 11.790 | 12.030 | 35.0 | Cluster_24758 |
| 442.89000 | 25.110 | 25.560 | 35.0 | Cluster_24763 |
| 442.98000 | 11.680 | 12.030 | 35.0 | Cluster_24761 |
| 443.12000 | 1.590 | 2.030 | 35.0 | Cluster_24771 |
| 443.22000 | 17.370 | 17.550 | 35.0 | Cluster_24782 |
| 443.28000 | 22.540 | 22.660 | 35.0 | Cluster_24785 |
| 443.29000 | 22.640 | 22.950 | 35.0 | Cluster_24795 |
| 443.89000 | 7.480 | 8.970 | 35.0 | Cluster_08322 |
| 443.92000 | 11.750 | 12.130 | 35.0 | Cluster_08324 |
| 444.72000 | 1.250 | 1.570 | 35.0 | Cluster_24866 |
| 445.18000 | 14.620 | 14.980 | 35.0 | Cluster_08343 |
| 445.55000 | 11.740 | 12.010 | 35.0 | Cluster_24900 |
| 446.27000 | 14.850 | 15.060 | 35.0 | Cluster_24941 |
| 446.73000 | 11.850 | 12.030 | 35.0 | Cluster_24959 |
| 446.91000 | 19.900 | 20.460 | 35.0 | Cluster_08369 |
| 447.17000 | 7.690 | 8.410 | 35.0 | Cluster_24974 |
| 447.25000 | 22.590 | 23.160 | 35.0 | Cluster_08376 |
| 447.28000 | 7.590 | 7.950 | 35.0 | Cluster_24982 |
| 447.29000 | 7.410 | 7.710 | 35.0 | Cluster_24988 |
| 447.60000 | 11.750 | 12.350 | 35.0 | Cluster_08380 |
| 448.30000 | 11.860 | 12.310 | 35.0 | Cluster_25022 |
| 448.79000 | 7.770 | 8.190 | 35.0 | Cluster_25051 |
| 448.97000 | 11.580 | 12.360 | 35.0 | Cluster_01573 |
| 449.25000 | 17.320 | 18.580 | 35.0 | Cluster_08416 |
| 449.74000 | 14.850 | 15.190 | 35.0 | Cluster_25092 |
| 449.84000 | 1.260 | 1.540 | 35.0 | Cluster_25102 |
| 450.23000 | 14.590 | 14.860 | 35.0 | Cluster_25125 |
| 450.93000 | 17.440 | 18.090 | 35.0 | Cluster_08435 |
| 451.14000 | 7.390 | 7.860 | 35.0 | Cluster_25153 |
| 451.17000 | 1.580 | 1.900 | 35.0 | Cluster_25168 |
| 451.57000 | 20.060 | 20.280 | 35.0 | Cluster_08446 |
| 452.13000 | 1.600 | 1.910 | 35.0 | Cluster_25195 |
| 452.26000 | 14.770 | 15.110 | 35.0 | Cluster_08462 |
| 452.75000 | 22.580 | 23.210 | 35.0 | Cluster_25235 |
| 453.14000 | 1.450 | 1.910 | 35.0 | Cluster_25244 |
| 453.24000 | 11.780 | 12.210 | 35.0 | Cluster_25248 |
| 454.68000 | 28.050 | 28.170 | 35.0 | Cluster_25320 |
| 454.70000 | 1.440 | 1.890 | 35.0 | Cluster_25319 |
| 454.74000 | 11.700 | 12.160 | 35.0 | Cluster_25325 |
| 455.15000 | 7.410 | 7.710 | 35.0 | Cluster_25343 |
| 455.18000 | 17.430 | 17.890 | 35.0 | Cluster_01617 |
| 455.20000 | 25.200 | 25.350 | 35.0 | Cluster_08518 |
| 456.24000 | 7.470 | 7.700 | 35.0 | Cluster_25393 |
| 456.26000 | 20.060 | 20.430 | 35.0 | Cluster_08536 |
| 456.57000 | 11.860 | 12.310 | 35.0 | Cluster_08539 |
| 456.75000 | 14.690 | 14.920 | 35.0 | Cluster_25404 |
| 456.78000 | 11.630 | 12.040 | 35.0 | Cluster_25413 |
| 457.20000 | 22.620 | 22.950 | 35.0 | Cluster_25422 |

| | | | | |
|---|---|---|---|---|
| 457.75000 | 7.570 | 8.250 | 35.0 | Cluster_25448 |
| 457.76000 | 19.950 | 20.130 | 35.0 | Cluster_25454 |
| 458.69000 | 7.650 | 8.440 | 35.0 | Cluster_25493 |
| 458.76000 | 11.740 | 12.070 | 35.0 | Cluster_25500 |
| 459.22000 | 7.770 | 8.510 | 35.0 | Cluster_08566 |
| 459.25000 | 7.430 | 7.750 | 35.0 | Cluster_25514 |
| 459.78000 | 11.640 | 12.160 | 35.0 | Cluster_25540 |
| 459.90000 | 22.680 | 23.130 | 35.0 | Cluster_08580 |
| 460.27000 | 14.580 | 14.840 | 35.0 | Cluster_25572 |
| 460.77000 | 7.570 | 7.800 | 35.0 | Cluster_25581 |
| 460.97000 | 19.890 | 20.150 | 35.0 | Cluster_01663 |
| 461.60000 | 17.530 | 17.830 | 35.0 | Cluster_08592 |
| 461.70000 | 7.800 | 8.280 | 35.0 | Cluster_25618 |
| 461.73000 | 11.600 | 12.100 | 35.0 | Cluster_25620 |
| 462.22000 | 14.740 | 15.170 | 35.0 | Cluster_08608 |
| 462.50000 | 11.680 | 12.120 | 35.0 | Cluster_25643 |
| 463.61000 | 17.260 | 17.520 | 35.0 | Cluster_25677 |
| 463.73000 | 7.690 | 7.930 | 35.0 | Cluster_25692 |
| 464.22000 | 20.010 | 20.270 | 35.0 | Cluster_08632 |
| 464.30000 | 22.630 | 23.030 | 35.0 | Cluster_25714 |
| 464.55000 | 22.620 | 23.550 | 35.0 | Cluster_08640 |
| 465.77000 | 7.690 | 7.940 | 35.0 | Cluster_25807 |
| 465.95000 | 17.270 | 17.670 | 35.0 | Cluster_08659 |
| 465.99000 | 11.720 | 12.350 | 35.0 | Cluster_01692 |
| 466.27000 | 14.870 | 15.280 | 35.0 | Cluster_25822 |
| 466.90000 | 14.760 | 15.170 | 35.0 | Cluster_08673 |
| 467.18000 | 7.360 | 7.770 | 35.0 | Cluster_08677 |
| 467.25000 | 7.800 | 8.810 | 35.0 | Cluster_08678 |
| 467.30000 | 25.180 | 25.510 | 35.0 | Cluster_25858 |
| 467.67000 | 25.110 | 25.470 | 35.0 | Cluster_00144 |
| 467.72000 | 17.470 | 17.920 | 35.0 | Cluster_25877 |
| 468.89000 | 17.410 | 17.840 | 35.0 | Cluster_08711 |
| 469.26000 | 31.370 | 31.710 | 35.0 | Cluster_08720 |
| 469.74000 | 11.690 | 11.980 | 35.0 | Cluster_25967 |
| 469.88000 | 14.730 | 15.000 | 35.0 | Cluster_08724 |
| 470.43000 | 19.910 | 20.270 | 35.0 | Cluster_26002 |
| 470.56000 | 7.620 | 7.940 | 35.0 | Cluster_00010 |
| 470.92000 | 11.610 | 12.050 | 35.0 | Cluster_08745 |
| 471.00000 | 11.820 | 12.340 | 35.0 | Cluster_01731 |
| 471.22000 | 1.510 | 2.030 | 35.0 | Cluster_08748 |
| 471.25000 | 22.510 | 22.710 | 35.0 | Cluster_01737 |
| 471.26000 | 22.690 | 22.990 | 35.0 | Cluster_01738 |
| 471.48000 | 17.440 | 17.680 | 35.0 | Cluster_26044 |
| 471.74000 | 14.730 | 14.930 | 35.0 | Cluster_26063 |
| 471.84000 | 7.370 | 7.650 | 35.0 | Cluster_26052 |
| 472.23000 | 22.680 | 22.910 | 35.0 | Cluster_26081 |
| 472.24000 | 11.690 | 12.530 | 35.0 | Cluster_08764 |
| 472.72000 | 11.740 | 12.060 | 35.0 | Cluster_26107 |
| 472.76000 | 11.600 | 11.750 | 35.0 | Cluster_26106 |
| 472.77000 | 14.810 | 15.350 | 35.0 | Cluster_26113 |
| 473.27000 | 11.610 | 11.880 | 35.0 | Cluster_26146 |

| | | | | |
|---|---|---|---|---|
| 473.77000 | 17.300 | 17.560 | 35.0 | Cluster_26157 |
| 474.54000 | 17.320 | 17.750 | 35.0 | Cluster_08802 |
| 474.58000 | 14.870 | 15.300 | 35.0 | Cluster_08800 |
| 474.85000 | 7.400 | 7.630 | 35.0 | Cluster_08806 |
| 475.58000 | 19.890 | 20.200 | 35.0 | Cluster_08820 |
| 475.78000 | 20.000 | 20.420 | 35.0 | Cluster_26234 |
| 476.21000 | 1.250 | 1.470 | 35.0 | Cluster_26244 |
| 476.23000 | 7.750 | 8.120 | 35.0 | Cluster_08830 |
| 476.28000 | 14.840 | 15.180 | 35.0 | Cluster_08832 |
| 477.07000 | 19.920 | 20.380 | 35.0 | Cluster_26275 |
| 477.20000 | 22.680 | 23.010 | 35.0 | Cluster_08843 |
| 477.27000 | 17.480 | 17.750 | 35.0 | Cluster_08840 |
| 477.55000 | 14.840 | 15.510 | 35.0 | Cluster_08851 |
| 477.61000 | 22.510 | 22.910 | 35.0 | Cluster_08856 |
| 477.93000 | 19.960 | 20.240 | 35.0 | Cluster_08861 |
| 478.22000 | 1.490 | 1.880 | 35.0 | Cluster_26309 |
| 478.25000 | 7.330 | 7.580 | 35.0 | Cluster_26313 |
| 478.76000 | 11.590 | 11.940 | 35.0 | Cluster_26329 |
| 478.77000 | 14.670 | 14.890 | 35.0 | Cluster_26332 |
| 478.93000 | 25.120 | 26.580 | 35.0 | Cluster_08880 |
| 478.99000 | 7.590 | 7.870 | 35.0 | Cluster_01780 |
| 479.57000 | 7.510 | 8.630 | 35.0 | Cluster_08888 |
| 479.71000 | 17.270 | 17.470 | 35.0 | Cluster_26429 |
| 479.76000 | 19.940 | 20.380 | 35.0 | Cluster_26423 |
| 480.22000 | 11.730 | 12.180 | 35.0 | Cluster_08900 |
| 480.70000 | 11.660 | 11.900 | 35.0 | Cluster_26476 |
| 481.22000 | 1.550 | 1.900 | 35.0 | Cluster_26493 |
| 481.27000 | 14.680 | 15.340 | 35.0 | Cluster_26504 |
| 481.28000 | 27.810 | 28.270 | 35.0 | Cluster_08920 |
| 481.56000 | 7.600 | 8.560 | 35.0 | Cluster_08922 |
| 481.58000 | 11.570 | 12.170 | 35.0 | Cluster_08924 |
| 481.72000 | 17.290 | 17.460 | 35.0 | Cluster_26523 |
| 481.73000 | 19.870 | 20.200 | 35.0 | Cluster_01804 |
| 481.79000 | 11.560 | 11.750 | 35.0 | Cluster_26518 |
| 482.20000 | 7.630 | 8.370 | 35.0 | Cluster_01810 |
| 482.28000 | 7.540 | 7.710 | 35.0 | Cluster_26547 |
| 482.49000 | 19.890 | 20.100 | 35.0 | Cluster_26549 |
| 483.24000 | 1.580 | 2.000 | 35.0 | Cluster_26594 |
| 483.30000 | 22.610 | 22.980 | 35.0 | Cluster_26603 |
| 483.76000 | 7.690 | 7.970 | 35.0 | Cluster_26625 |
| 483.84000 | 1.270 | 1.660 | 35.0 | Cluster_26627 |
| 484.09000 | 17.390 | 17.580 | 35.0 | Cluster_26635 |
| 484.73000 | 19.950 | 20.210 | 35.0 | Cluster_26685 |
| 484.76000 | 11.760 | 12.370 | 35.0 | Cluster_26681 |
| 485.06000 | 25.120 | 25.450 | 35.0 | Cluster_00166 |
| 485.54000 | 11.860 | 12.200 | 35.0 | Cluster_08981 |
| 488.29000 | 17.460 | 17.920 | 35.0 | Cluster_26837 |
| 488.61000 | 27.970 | 28.270 | 35.0 | Cluster_09032 |
| 488.79000 | 19.960 | 20.480 | 35.0 | Cluster_26859 |
| 488.95000 | 19.910 | 20.540 | 35.0 | Cluster_09038 |
| 489.74000 | 19.910 | 20.150 | 35.0 | Cluster_26904 |

| | | | | |
|---|---|---|---|---|
| 489.93000 | 11.670 | 12.140 | 35.0 | Cluster_09048 |
| 490.50000 | 11.570 | 12.530 | 35.0 | Cluster_26931 |
| 490.70000 | 1.610 | 2.010 | 35.0 | Cluster_26935 |
| 490.77000 | 14.620 | 14.740 | 35.0 | Cluster_26939 |
| 492.55000 | 14.810 | 14.940 | 35.0 | Cluster_09097 |
| 492.60000 | 14.680 | 14.860 | 35.0 | Cluster_09099 |
| 493.27000 | 17.370 | 17.580 | 35.0 | Cluster_27041 |
| 494.26000 | 7.650 | 7.870 | 35.0 | Cluster_27100 |
| 495.19000 | 19.900 | 20.310 | 35.0 | Cluster_27149 |
| 495.71000 | 1.570 | 1.910 | 35.0 | Cluster_27170 |
| 495.95000 | 22.630 | 22.900 | 35.0 | Cluster_09160 |
| 496.24000 | 19.960 | 20.280 | 35.0 | Cluster_01895 |
| 496.95000 | 31.350 | 31.660 | 35.0 | Cluster_09176 |
| 497.35000 | 1.270 | 1.610 | 35.0 | Cluster_27258 |
| 497.74000 | 7.370 | 7.780 | 35.0 | Cluster_27265 |
| 497.80000 | 17.360 | 17.690 | 35.0 | Cluster_27275 |
| 497.94000 | 22.570 | 23.390 | 35.0 | Cluster_09191 |
| 498.01000 | 25.130 | 25.440 | 35.0 | Cluster_01929 |
| 498.50000 | 11.610 | 11.980 | 35.0 | Cluster_27299 |
| 498.73000 | 7.580 | 7.760 | 35.0 | Cluster_27302 |
| 499.23000 | 14.690 | 15.160 | 35.0 | Cluster_09211 |
| 499.33000 | 22.660 | 23.010 | 35.0 | Cluster_27324 |
| 499.76000 | 22.660 | 22.870 | 35.0 | Cluster_01941 |
| 499.92000 | 19.990 | 20.350 | 35.0 | Cluster_09218 |
| 499.96000 | 27.850 | 28.580 | 35.0 | Cluster_09219 |
| 500.02000 | 1.280 | 1.610 | 35.0 | Cluster_27355 |
| 500.63000 | 22.660 | 23.320 | 35.0 | Cluster_09231 |
| 500.70000 | 7.470 | 8.230 | 35.0 | Cluster_27401 |
| 500.96000 | 1.250 | 1.490 | 35.0 | Cluster_01956 |
| 501.22000 | 14.600 | 15.020 | 35.0 | Cluster_27436 |
| 501.59000 | 19.900 | 20.410 | 35.0 | Cluster_09244 |
| 501.76000 | 25.270 | 25.430 | 35.0 | Cluster_27459 |
| 502.21000 | 14.840 | 15.400 | 35.0 | Cluster_01970 |
| 502.22000 | 19.940 | 20.310 | 35.0 | Cluster_27473 |
| 502.59000 | 11.820 | 12.250 | 35.0 | Cluster_09268 |
| 502.76000 | 7.580 | 8.030 | 35.0 | Cluster_27485 |
| 503.25000 | 7.500 | 7.680 | 35.0 | Cluster_27512 |
| 503.28000 | 27.860 | 28.290 | 35.0 | Cluster_27518 |
| 504.21000 | 17.260 | 17.490 | 35.0 | Cluster_27547 |
| 504.56000 | 14.800 | 15.010 | 35.0 | Cluster_09288 |
| 504.98000 | 25.280 | 25.740 | 35.0 | Cluster_01984 |
| 505.01000 | 17.500 | 18.120 | 35.0 | Cluster_27567 |
| 505.24000 | 19.920 | 20.380 | 35.0 | Cluster_09298 |
| 505.52000 | 11.640 | 11.980 | 35.0 | Cluster_01990 |
| 505.60000 | 14.600 | 14.880 | 35.0 | Cluster_09305 |
| 505.79000 | 14.760 | 15.190 | 35.0 | Cluster_27593 |
| 506.73000 | 1.450 | 1.900 | 35.0 | Cluster_27611 |
| 506.74000 | 22.540 | 22.870 | 35.0 | Cluster_27621 |
| 506.76000 | 11.670 | 11.950 | 35.0 | Cluster_27614 |
| 507.25000 | 11.830 | 11.980 | 35.0 | Cluster_27638 |
| 507.97000 | 14.640 | 15.020 | 35.0 | Cluster_09350 |

| | | | | |
|---|---|---|---|---|
| 508.62000 | 20.060 | 20.640 | 35.0 | Cluster_09366 |
| 508.70000 | 1.600 | 2.010 | 35.0 | Cluster_27686 |
| 508.71000 | 31.670 | 31.910 | 35.0 | Cluster_27692 |
| 509.20000 | 14.750 | 15.200 | 35.0 | Cluster_27715 |
| 509.26000 | 17.360 | 17.720 | 35.0 | Cluster_27718 |
| 509.87000 | 19.870 | 20.150 | 35.0 | Cluster_09377 |
| 509.91000 | 11.720 | 12.580 | 35.0 | Cluster_09383 |
| 511.23000 | 19.900 | 20.500 | 35.0 | Cluster_27820 |
| 511.79000 | 25.080 | 25.960 | 35.0 | Cluster_27852 |
| 511.91000 | 7.750 | 8.370 | 35.0 | Cluster_09420 |
| 512.25000 | 22.680 | 22.900 | 35.0 | Cluster_27867 |
| 512.27000 | 17.380 | 17.700 | 35.0 | Cluster_27865 |
| 513.09000 | 7.390 | 7.620 | 35.0 | Cluster_27898 |
| 513.77000 | 14.700 | 15.030 | 35.0 | Cluster_27929 |
| 514.72000 | 7.380 | 7.820 | 35.0 | Cluster_27964 |
| 514.74000 | 11.660 | 11.800 | 35.0 | Cluster_27971 |
| 515.25000 | 11.820 | 12.340 | 35.0 | Cluster_02053 |
| 515.48000 | 14.690 | 15.030 | 35.0 | Cluster_02061 |
| 515.90000 | 17.500 | 17.700 | 35.0 | Cluster_09478 |
| 516.74000 | 7.410 | 8.010 | 35.0 | Cluster_28062 |
| 517.92000 | 17.270 | 17.770 | 35.0 | Cluster_09503 |
| 518.00000 | 7.400 | 8.010 | 35.0 | Cluster_28115 |
| 518.26000 | 7.480 | 8.140 | 35.0 | Cluster_09508 |
| 518.31000 | 22.490 | 22.790 | 35.0 | Cluster_28130 |
| 518.51000 | 11.730 | 11.950 | 35.0 | Cluster_28138 |
| 518.59000 | 25.120 | 25.590 | 35.0 | Cluster_09521 |
| 518.78000 | 11.860 | 12.260 | 35.0 | Cluster_28152 |
| 519.59000 | 19.910 | 20.490 | 35.0 | Cluster_09531 |
| 519.81000 | 11.710 | 11.900 | 35.0 | Cluster_28200 |
| 520.04000 | 11.820 | 12.330 | 35.0 | Cluster_28202 |
| 520.51000 | 19.930 | 20.220 | 35.0 | Cluster_28225 |
| 520.80000 | 19.990 | 20.360 | 35.0 | Cluster_28240 |
| 520.81000 | 14.720 | 14.890 | 35.0 | Cluster_28232 |
| 521.02000 | 27.920 | 28.760 | 35.0 | Cluster_02095 |
| 521.26000 | 25.120 | 25.760 | 35.0 | Cluster_09553 |
| 521.66000 | 14.750 | 14.990 | 35.0 | Cluster_28271 |
| 522.22000 | 19.900 | 20.320 | 35.0 | Cluster_28286 |
| 522.45000 | 17.530 | 17.930 | 35.0 | Cluster_02102 |
| 522.50000 | 14.850 | 15.550 | 35.0 | Cluster_28303 |
| 522.62000 | 22.580 | 23.260 | 35.0 | Cluster_09582 |
| 522.75000 | 14.580 | 14.890 | 35.0 | Cluster_28315 |
| 523.22000 | 1.540 | 1.970 | 35.0 | Cluster_28350 |
| 523.27000 | 22.580 | 22.760 | 35.0 | Cluster_28334 |
| 523.35000 | 7.710 | 8.070 | 35.0 | Cluster_28353 |
| 523.92000 | 27.830 | 28.280 | 35.0 | Cluster_09603 |
| 523.97000 | 19.890 | 20.330 | 35.0 | Cluster_28374 |
| 524.28000 | 7.440 | 7.750 | 35.0 | Cluster_28386 |
| 526.18000 | 27.810 | 27.980 | 35.0 | Cluster_28467 |
| 526.20000 | 19.930 | 20.490 | 35.0 | Cluster_09656 |
| 526.93000 | 17.300 | 17.490 | 35.0 | Cluster_09676 |
| 526.95000 | 17.420 | 17.830 | 35.0 | Cluster_09677 |

| | | | | |
|---|---|---|---|---|
| 527.76000 | 14.630 | 14.940 | 35.0 | Cluster_28507 |
| 528.23000 | 25.180 | 25.480 | 35.0 | Cluster_28536 |
| 528.27000 | 31.350 | 31.880 | 35.0 | Cluster_00184 |
| 528.32000 | 11.860 | 12.150 | 35.0 | Cluster_28539 |
| 528.36000 | 1.420 | 1.660 | 35.0 | Cluster_28552 |
| 528.97000 | 25.130 | 25.350 | 35.0 | Cluster_09710 |
| 529.25000 | 1.250 | 1.600 | 35.0 | Cluster_09712 |
| 529.79000 | 27.930 | 28.250 | 35.0 | Cluster_28615 |
| 530.20000 | 19.970 | 20.370 | 35.0 | Cluster_28629 |
| 530.26000 | 28.010 | 28.680 | 35.0 | Cluster_02153 |
| 530.51000 | 28.000 | 28.590 | 35.0 | Cluster_28657 |
| 530.81000 | 19.940 | 20.120 | 35.0 | Cluster_28677 |
| 530.94000 | 17.380 | 17.810 | 35.0 | Cluster_09741 |
| 531.75000 | 11.600 | 11.860 | 35.0 | Cluster_28707 |
| 531.77000 | 11.850 | 12.330 | 35.0 | Cluster_28708 |
| 532.24000 | 19.900 | 20.560 | 35.0 | Cluster_09763 |
| 532.27000 | 19.900 | 20.070 | 35.0 | Cluster_09766 |
| 532.62000 | 27.970 | 28.560 | 35.0 | Cluster_09768 |
| 532.97000 | 17.500 | 18.660 | 35.0 | Cluster_09773 |
| 533.15000 | 1.420 | 1.860 | 35.0 | Cluster_28752 |
| 533.16000 | 1.570 | 1.940 | 35.0 | Cluster_28753 |
| 533.19000 | 27.910 | 28.680 | 35.0 | Cluster_09774 |
| 533.32000 | 31.520 | 34.610 | 35.0 | Cluster_09778 |
| 533.70000 | 27.990 | 28.510 | 35.0 | Cluster_28783 |
| 533.82000 | 25.050 | 25.470 | 35.0 | Cluster_28799 |
| 533.91000 | 25.110 | 25.630 | 35.0 | Cluster_09792 |
| 534.28000 | 7.780 | 8.110 | 35.0 | Cluster_28807 |
| 534.59000 | 17.310 | 17.790 | 35.0 | Cluster_09803 |
| 535.32000 | 31.570 | 34.610 | 35.0 | Cluster_09812 |
| 535.80000 | 14.830 | 15.050 | 35.0 | Cluster_28867 |
| 535.94000 | 22.660 | 23.170 | 35.0 | Cluster_09822 |
| 536.06000 | 14.680 | 14.880 | 35.0 | Cluster_28878 |
| 536.26000 | 17.340 | 17.850 | 35.0 | Cluster_09823 |
| 536.27000 | 11.740 | 12.330 | 35.0 | Cluster_28888 |
| 536.48000 | 1.270 | 1.570 | 35.0 | Cluster_28899 |
| 536.60000 | 28.030 | 28.230 | 35.0 | Cluster_09834 |
| 537.23000 | 19.940 | 20.380 | 35.0 | Cluster_09849 |
| 537.56000 | 20.040 | 20.430 | 35.0 | Cluster_09853 |
| 537.69000 | 20.000 | 20.310 | 35.0 | Cluster_28943 |
| 537.91000 | 17.280 | 17.830 | 35.0 | Cluster_09869 |
| 538.84000 | 27.940 | 29.330 | 35.0 | Cluster_28980 |
| 538.97000 | 25.250 | 26.160 | 35.0 | Cluster_09886 |
| 538.98000 | 25.040 | 25.270 | 35.0 | Cluster_09885 |
| 539.01000 | 14.600 | 14.840 | 35.0 | Cluster_28985 |
| 539.28000 | 11.840 | 12.000 | 35.0 | Cluster_28993 |
| 539.52000 | 7.750 | 8.260 | 35.0 | Cluster_29001 |
| 540.02000 | 7.730 | 8.260 | 35.0 | Cluster_09899 |
| 540.27000 | 20.030 | 20.690 | 35.0 | Cluster_29031 |
| 540.79000 | 11.560 | 12.000 | 35.0 | Cluster_29052 |
| 541.34000 | 11.580 | 11.860 | 35.0 | Cluster_29085 |
| 541.44000 | 19.890 | 20.040 | 35.0 | Cluster_29098 |

| | | | | |
|---|---|---|---|---|
| 541.82000 | 11.830 | 12.010 | 35.0 | Cluster_29115 |
| 542.26000 | 31.550 | 31.980 | 35.0 | Cluster_02236 |
| 542.28000 | 17.530 | 17.810 | 35.0 | Cluster_02229 |
| 542.52000 | 19.870 | 20.080 | 35.0 | Cluster_02233 |
| 542.77000 | 11.860 | 12.200 | 35.0 | Cluster_29162 |
| 543.27000 | 22.630 | 23.250 | 35.0 | Cluster_29193 |
| 544.95000 | 11.820 | 12.320 | 35.0 | Cluster_09982 |
| 545.77000 | 7.360 | 7.830 | 35.0 | Cluster_29280 |
| 545.92000 | 17.420 | 18.000 | 35.0 | Cluster_09992 |
| 546.31000 | 14.620 | 14.810 | 35.0 | Cluster_29312 |
| 547.32000 | 17.390 | 17.760 | 35.0 | Cluster_29345 |
| 548.27000 | 14.730 | 15.170 | 35.0 | Cluster_29389 |
| 548.46000 | 25.260 | 25.750 | 35.0 | Cluster_00194 |
| 548.61000 | 1.570 | 2.000 | 35.0 | Cluster_10035 |
| 548.64000 | 22.690 | 23.540 | 35.0 | Cluster_10037 |
| 549.25000 | 27.830 | 28.270 | 35.0 | Cluster_10044 |
| 549.28000 | 7.590 | 7.870 | 35.0 | Cluster_29418 |
| 549.90000 | 11.780 | 12.150 | 35.0 | Cluster_10052 |
| 549.99000 | 25.100 | 25.460 | 35.0 | Cluster_10058 |
| 550.22000 | 7.560 | 7.770 | 35.0 | Cluster_29451 |
| 550.31000 | 22.610 | 23.060 | 35.0 | Cluster_29461 |
| 550.55000 | 17.370 | 17.630 | 35.0 | Cluster_10069 |
| 550.61000 | 14.840 | 15.150 | 35.0 | Cluster_10068 |
| 550.78000 | 7.640 | 7.920 | 35.0 | Cluster_29465 |
| 550.84000 | 17.510 | 17.830 | 35.0 | Cluster_29476 |
| 551.01000 | 25.090 | 25.830 | 35.0 | Cluster_29483 |
| 551.29000 | 11.610 | 11.870 | 35.0 | Cluster_29499 |
| 551.57000 | 7.610 | 8.130 | 35.0 | Cluster_10082 |
| 551.97000 | 17.510 | 17.780 | 35.0 | Cluster_10089 |
| 552.36000 | 19.970 | 20.430 | 35.0 | Cluster_29545 |
| 552.74000 | 7.590 | 7.850 | 35.0 | Cluster_29553 |
| 552.75000 | 1.570 | 2.190 | 35.0 | Cluster_29548 |
| 552.96000 | 25.250 | 25.480 | 35.0 | Cluster_10104 |
| 553.55000 | 19.990 | 20.610 | 35.0 | Cluster_29579 |
| 553.63000 | 11.870 | 12.380 | 35.0 | Cluster_10116 |
| 554.58000 | 27.910 | 28.110 | 35.0 | Cluster_10130 |
| 554.60000 | 17.510 | 17.920 | 35.0 | Cluster_10136 |
| 554.97000 | 22.600 | 23.080 | 35.0 | Cluster_10144 |
| 556.25000 | 14.750 | 15.130 | 35.0 | Cluster_29692 |
| 556.68000 | 27.880 | 28.460 | 35.0 | Cluster_10177 |
| 556.96000 | 14.720 | 14.880 | 35.0 | Cluster_10184 |
| 557.73000 | 14.790 | 15.330 | 35.0 | Cluster_29750 |
| 558.23000 | 17.360 | 17.700 | 35.0 | Cluster_10209 |
| 558.65000 | 14.880 | 15.260 | 35.0 | Cluster_10215 |
| 558.81000 | 22.580 | 23.130 | 35.0 | Cluster_29792 |
| 559.29000 | 7.590 | 7.940 | 35.0 | Cluster_29811 |
| 560.35000 | 28.000 | 28.210 | 35.0 | Cluster_29845 |
| 560.65000 | 19.890 | 20.480 | 35.0 | Cluster_10245 |
| 560.83000 | 25.190 | 25.430 | 35.0 | Cluster_29862 |
| 561.28000 | 28.030 | 28.230 | 35.0 | Cluster_29879 |
| 561.64000 | 31.590 | 32.110 | 35.0 | Cluster_10262 |

| | | | | |
|---|---|---|---|---|
| 561.70000 | 25.100 | 26.430 | 35.0 | Cluster_29892 |
| 561.81000 | 17.510 | 17.830 | 35.0 | Cluster_29901 |
| 561.98000 | 22.680 | 25.390 | 35.0 | Cluster_10266 |
| 562.07000 | 7.430 | 7.710 | 35.0 | Cluster_29906 |
| 562.29000 | 19.900 | 20.370 | 35.0 | Cluster_29915 |
| 563.32000 | 19.870 | 20.460 | 35.0 | Cluster_10288 |
| 563.50000 | 22.690 | 23.220 | 35.0 | Cluster_02355 |
| 563.75000 | 14.750 | 15.200 | 35.0 | Cluster_29975 |
| 563.76000 | 22.530 | 22.950 | 35.0 | Cluster_29965 |
| 564.78000 | 14.710 | 15.270 | 35.0 | Cluster_30003 |
| 564.79000 | 20.010 | 20.400 | 35.0 | Cluster_02366 |
| 565.56000 | 19.960 | 20.140 | 35.0 | Cluster_30039 |
| 565.60000 | 25.100 | 25.530 | 35.0 | Cluster_10318 |
| 566.33000 | 11.630 | 12.010 | 35.0 | Cluster_30080 |
| 566.74000 | 20.060 | 20.350 | 35.0 | Cluster_02376 |
| 566.79000 | 7.390 | 7.780 | 35.0 | Cluster_30088 |
| 566.90000 | 22.620 | 22.840 | 35.0 | Cluster_30092 |
| 566.98000 | 22.640 | 23.320 | 35.0 | Cluster_10333 |
| 567.07000 | 17.470 | 17.720 | 35.0 | Cluster_30105 |
| 567.24000 | 14.640 | 14.920 | 35.0 | Cluster_30110 |
| 567.30000 | 28.010 | 28.530 | 35.0 | Cluster_30123 |
| 567.31000 | 11.850 | 12.010 | 35.0 | Cluster_30114 |
| 568.56000 | 25.160 | 25.510 | 35.0 | Cluster_02384 |
| 568.73000 | 14.840 | 15.260 | 35.0 | Cluster_30166 |
| 568.79000 | 31.530 | 31.820 | 35.0 | Cluster_30177 |
| 568.80000 | 17.460 | 17.930 | 35.0 | Cluster_30176 |
| 568.85000 | 28.010 | 28.520 | 35.0 | Cluster_30178 |
| 569.06000 | 25.130 | 25.340 | 35.0 | Cluster_02386 |
| 569.57000 | 25.040 | 25.390 | 35.0 | Cluster_02390 |
| 569.82000 | 17.530 | 17.910 | 35.0 | Cluster_30204 |
| 570.30000 | 17.480 | 18.020 | 35.0 | Cluster_02396 |
| 570.36000 | 22.500 | 22.780 | 35.0 | Cluster_30219 |
| 570.73000 | 20.060 | 20.310 | 35.0 | Cluster_02397 |
| 570.77000 | 7.460 | 7.770 | 35.0 | Cluster_30236 |
| 571.22000 | 17.470 | 17.700 | 35.0 | Cluster_30252 |
| 571.34000 | 27.880 | 28.210 | 35.0 | Cluster_30256 |
| 571.64000 | 11.820 | 12.560 | 35.0 | Cluster_10409 |
| 571.77000 | 7.480 | 8.050 | 35.0 | Cluster_30264 |
| 572.13000 | 22.500 | 23.000 | 35.0 | Cluster_00213 |
| 572.30000 | 17.340 | 18.190 | 35.0 | Cluster_10420 |
| 572.79000 | 14.660 | 14.850 | 35.0 | Cluster_30301 |
| 573.35000 | 22.660 | 22.940 | 35.0 | Cluster_30317 |
| 574.26000 | 11.830 | 12.020 | 35.0 | Cluster_10456 |
| 574.80000 | 11.580 | 11.830 | 35.0 | Cluster_30355 |
| 575.25000 | 31.530 | 33.580 | 35.0 | Cluster_02433 |
| 575.33000 | 22.510 | 22.760 | 35.0 | Cluster_02435 |
| 576.55000 | 17.430 | 17.830 | 35.0 | Cluster_02439 |
| 576.77000 | 14.720 | 14.850 | 35.0 | Cluster_30417 |
| 576.98000 | 11.830 | 12.410 | 35.0 | Cluster_10494 |
| 577.30000 | 19.960 | 20.490 | 35.0 | Cluster_10500 |
| 577.81000 | 31.450 | 31.790 | 35.0 | Cluster_30446 |

| | | | | |
|---|---|---|---|---|
| 578.32000 | 17.430 | 17.920 | 35.0 | Cluster_30460 |
| 578.55000 | 19.870 | 20.210 | 35.0 | Cluster_02454 |
| 578.86000 | 22.490 | 22.830 | 35.0 | Cluster_30486 |
| 578.98000 | 14.820 | 15.290 | 35.0 | Cluster_10523 |
| 579.33000 | 11.750 | 12.130 | 35.0 | Cluster_30494 |
| 579.55000 | 25.260 | 25.600 | 35.0 | Cluster_02460 |
| 579.61000 | 7.580 | 8.870 | 35.0 | Cluster_10531 |
| 579.63000 | 22.500 | 23.120 | 35.0 | Cluster_10533 |
| 579.77000 | 11.700 | 12.170 | 35.0 | Cluster_30512 |
| 579.81000 | 17.460 | 17.700 | 35.0 | Cluster_30517 |
| 579.83000 | 14.670 | 15.090 | 35.0 | Cluster_30515 |
| 580.36000 | 17.280 | 17.480 | 35.0 | Cluster_30555 |
| 580.86000 | 25.310 | 25.670 | 35.0 | Cluster_02466 |
| 581.28000 | 20.030 | 20.280 | 35.0 | Cluster_02468 |
| 581.31000 | 7.630 | 7.950 | 35.0 | Cluster_30577 |
| 581.96000 | 11.650 | 12.310 | 35.0 | Cluster_10563 |
| 582.31000 | 7.770 | 8.190 | 35.0 | Cluster_30648 |
| 582.54000 | 17.320 | 17.540 | 35.0 | Cluster_02478 |
| 582.65000 | 25.140 | 25.550 | 35.0 | Cluster_10574 |
| 582.70000 | 22.490 | 23.000 | 35.0 | Cluster_00226 |
| 582.74000 | 1.640 | 2.170 | 35.0 | Cluster_30654 |
| 584.05000 | 19.980 | 20.140 | 35.0 | Cluster_30718 |
| 584.31000 | 14.750 | 15.070 | 35.0 | Cluster_30724 |
| 584.54000 | 25.270 | 25.550 | 35.0 | Cluster_30733 |
| 585.02000 | 14.690 | 15.040 | 35.0 | Cluster_30763 |
| 585.29000 | 25.160 | 25.780 | 35.0 | Cluster_10612 |
| 585.35000 | 31.700 | 32.250 | 35.0 | Cluster_02501 |
| 585.80000 | 31.510 | 31.730 | 35.0 | Cluster_30791 |
| 585.84000 | 27.950 | 28.760 | 35.0 | Cluster_30792 |
| 586.62000 | 14.830 | 15.010 | 35.0 | Cluster_10630 |
| 586.75000 | 1.640 | 2.010 | 35.0 | Cluster_30815 |
| 587.30000 | 14.680 | 14.880 | 35.0 | Cluster_10642 |
| 587.33000 | 11.630 | 11.810 | 35.0 | Cluster_30848 |
| 587.34000 | 17.340 | 17.720 | 35.0 | Cluster_30837 |
| 587.65000 | 14.830 | 15.170 | 35.0 | Cluster_10646 |
| 588.69000 | 11.810 | 12.000 | 35.0 | Cluster_30892 |
| 588.82000 | 19.980 | 20.330 | 35.0 | Cluster_30900 |
| 589.55000 | 20.030 | 20.200 | 35.0 | Cluster_30932 |
| 589.74000 | 7.420 | 7.770 | 35.0 | Cluster_30933 |
| 589.96000 | 20.020 | 20.580 | 35.0 | Cluster_10677 |
| 590.22000 | 17.360 | 17.460 | 35.0 | Cluster_30952 |
| 590.30000 | 17.400 | 17.930 | 35.0 | Cluster_30953 |
| 591.04000 | 20.040 | 20.300 | 35.0 | Cluster_30998 |
| 591.44000 | 7.460 | 7.700 | 35.0 | Cluster_31008 |
| 591.52000 | 25.310 | 25.570 | 35.0 | Cluster_02538 |
| 591.66000 | 17.410 | 17.790 | 35.0 | Cluster_10703 |
| 591.81000 | 31.510 | 31.730 | 35.0 | Cluster_31027 |
| 592.01000 | 22.470 | 22.920 | 35.0 | Cluster_10706 |
| 592.79000 | 7.750 | 8.200 | 35.0 | Cluster_31056 |
| 592.82000 | 28.040 | 28.460 | 35.0 | Cluster_31060 |
| 593.06000 | 25.120 | 25.660 | 35.0 | Cluster_02547 |

| | | | | |
|---|---|---|---|---|
| 593.07000 | 25.320 | 25.710 | 35.0 | Cluster_02548 |
| 593.33000 | 22.540 | 22.890 | 35.0 | Cluster_10724 |
| 593.34000 | 11.740 | 12.200 | 35.0 | Cluster_31073 |
| 593.93000 | 17.370 | 17.520 | 35.0 | Cluster_10743 |
| 594.32000 | 27.980 | 28.390 | 35.0 | Cluster_10753 |
| 595.59000 | 17.510 | 17.810 | 35.0 | Cluster_31146 |
| 595.62000 | 17.380 | 17.730 | 35.0 | Cluster_10769 |
| 596.75000 | 7.800 | 8.600 | 35.0 | Cluster_31177 |
| 597.67000 | 31.700 | 32.040 | 35.0 | Cluster_10808 |
| 597.81000 | 22.500 | 22.900 | 35.0 | Cluster_31212 |
| 597.82000 | 17.410 | 17.830 | 35.0 | Cluster_31208 |
| 598.29000 | 14.840 | 15.150 | 35.0 | Cluster_10814 |
| 598.36000 | 27.860 | 28.450 | 35.0 | Cluster_10818 |
| 599.36000 | 19.990 | 20.260 | 35.0 | Cluster_31267 |
| 599.66000 | 31.720 | 32.150 | 35.0 | Cluster_10840 |
| 599.95000 | 19.900 | 20.430 | 35.0 | Cluster_10848 |
| 600.81000 | 7.740 | 8.050 | 35.0 | Cluster_31316 |
| 600.97000 | 11.800 | 12.190 | 35.0 | Cluster_10858 |
| 601.36000 | 22.680 | 23.100 | 35.0 | Cluster_31339 |
| 602.32000 | 27.800 | 28.110 | 35.0 | Cluster_31390 |
| 602.83000 | 20.070 | 20.220 | 35.0 | Cluster_31406 |
| 603.30000 | 11.680 | 12.200 | 35.0 | Cluster_31409 |
| 603.51000 | 7.800 | 7.990 | 35.0 | Cluster_02598 |
| 603.53000 | 19.920 | 20.190 | 35.0 | Cluster_31419 |
| 603.96000 | 11.700 | 12.340 | 35.0 | Cluster_10888 |
| 604.80000 | 17.520 | 17.930 | 35.0 | Cluster_31463 |
| 605.31000 | 11.820 | 12.480 | 35.0 | Cluster_31473 |
| 605.67000 | 25.090 | 25.450 | 35.0 | Cluster_10910 |
| 606.07000 | 31.390 | 31.590 | 35.0 | Cluster_31501 |
| 606.27000 | 7.750 | 7.930 | 35.0 | Cluster_31514 |
| 606.87000 | 25.100 | 25.350 | 35.0 | Cluster_31531 |
| 606.89000 | 25.280 | 25.620 | 35.0 | Cluster_10941 |
| 607.32000 | 14.800 | 15.120 | 35.0 | Cluster_31536 |
| 608.09000 | 22.520 | 22.950 | 35.0 | Cluster_02626 |
| 608.32000 | 31.350 | 31.690 | 35.0 | Cluster_31587 |
| 608.49000 | 19.940 | 20.290 | 35.0 | Cluster_31590 |
| 608.78000 | 14.760 | 15.080 | 35.0 | Cluster_31595 |
| 608.79000 | 1.520 | 1.980 | 35.0 | Cluster_31601 |
| 609.06000 | 17.530 | 17.740 | 35.0 | Cluster_31605 |
| 609.81000 | 7.750 | 8.000 | 35.0 | Cluster_31625 |
| 610.31000 | 31.440 | 31.740 | 35.0 | Cluster_31655 |
| 611.34000 | 22.690 | 22.940 | 35.0 | Cluster_31685 |
| 612.32000 | 14.860 | 14.970 | 35.0 | Cluster_11010 |
| 613.83000 | 27.990 | 28.170 | 35.0 | Cluster_31794 |
| 614.06000 | 14.760 | 14.970 | 35.0 | Cluster_31796 |
| 614.07000 | 17.250 | 17.600 | 35.0 | Cluster_02670 |
| 614.19000 | 1.610 | 1.960 | 35.0 | Cluster_31797 |
| 614.26000 | 17.420 | 18.050 | 35.0 | Cluster_11035 |
| 615.83000 | 31.340 | 31.550 | 35.0 | Cluster_31852 |
| 616.08000 | 31.590 | 31.820 | 35.0 | Cluster_02680 |
| 616.32000 | 28.000 | 28.760 | 35.0 | Cluster_11075 |

| | | | | |
|---|---|---|---|---|
| 616.55000 | 14.620 | 14.950 | 35.0 | Cluster_31873 |
| 616.83000 | 14.850 | 14.940 | 35.0 | Cluster_31882 |
| 617.02000 | 25.040 | 25.250 | 35.0 | Cluster_31892 |
| 617.35000 | 31.420 | 32.000 | 35.0 | Cluster_31911 |
| 617.93000 | 31.430 | 31.700 | 35.0 | Cluster_31935 |
| 618.05000 | 11.870 | 12.540 | 35.0 | Cluster_02693 |
| 618.79000 | 22.510 | 22.750 | 35.0 | Cluster_31952 |
| 619.29000 | 27.910 | 28.450 | 35.0 | Cluster_02701 |
| 619.33000 | 11.880 | 12.150 | 35.0 | Cluster_31985 |
| 619.62000 | 14.870 | 15.340 | 35.0 | Cluster_11116 |
| 620.53000 | 25.290 | 25.590 | 35.0 | Cluster_32028 |
| 620.60000 | 17.420 | 17.850 | 35.0 | Cluster_11128 |
| 621.30000 | 7.510 | 7.700 | 35.0 | Cluster_32049 |
| 621.36000 | 31.580 | 31.760 | 35.0 | Cluster_32053 |
| 621.39000 | 25.140 | 25.350 | 35.0 | Cluster_32052 |
| 621.79000 | 22.560 | 22.900 | 35.0 | Cluster_32057 |
| 621.98000 | 27.860 | 28.620 | 35.0 | Cluster_11154 |
| 622.78000 | 14.670 | 14.880 | 35.0 | Cluster_32095 |
| 622.98000 | 19.870 | 20.320 | 35.0 | Cluster_11175 |
| 623.28000 | 11.660 | 12.090 | 35.0 | Cluster_32112 |
| 623.36000 | 14.850 | 15.070 | 35.0 | Cluster_32126 |
| 623.65000 | 28.070 | 28.400 | 35.0 | Cluster_11191 |
| 623.98000 | 20.050 | 20.290 | 35.0 | Cluster_11189 |
| 624.20000 | 1.600 | 1.870 | 35.0 | Cluster_32153 |
| 624.28000 | 7.840 | 8.040 | 35.0 | Cluster_32154 |
| 624.63000 | 28.070 | 28.390 | 35.0 | Cluster_11202 |
| 625.04000 | 19.980 | 20.630 | 35.0 | Cluster_02732 |
| 625.32000 | 11.570 | 12.160 | 35.0 | Cluster_11203 |
| 625.58000 | 27.880 | 28.190 | 35.0 | Cluster_02735 |
| 626.00000 | 20.030 | 20.250 | 35.0 | Cluster_11225 |
| 626.07000 | 27.840 | 28.080 | 35.0 | Cluster_32208 |
| 626.27000 | 25.120 | 25.590 | 35.0 | Cluster_32218 |
| 626.28000 | 7.550 | 8.420 | 35.0 | Cluster_32212 |
| 626.83000 | 11.680 | 11.830 | 35.0 | Cluster_32244 |
| 627.30000 | 31.710 | 32.020 | 35.0 | Cluster_32266 |
| 627.31000 | 17.250 | 17.550 | 35.0 | Cluster_32263 |
| 627.50000 | 1.390 | 1.660 | 35.0 | Cluster_32275 |
| 627.79000 | 17.480 | 17.610 | 35.0 | Cluster_32283 |
| 628.07000 | 7.800 | 8.180 | 35.0 | Cluster_32292 |
| 628.86000 | 1.450 | 1.770 | 35.0 | Cluster_32321 |
| 629.63000 | 11.830 | 12.310 | 35.0 | Cluster_11289 |
| 629.83000 | 14.580 | 15.190 | 35.0 | Cluster_32344 |
| 629.96000 | 7.690 | 8.530 | 35.0 | Cluster_11292 |
| 630.84000 | 11.630 | 11.990 | 35.0 | Cluster_32375 |
| 631.37000 | 17.410 | 17.660 | 35.0 | Cluster_11312 |
| 631.80000 | 14.840 | 15.180 | 35.0 | Cluster_32410 |
| 632.05000 | 28.070 | 28.630 | 35.0 | Cluster_11324 |
| 632.35000 | 31.350 | 31.640 | 35.0 | Cluster_11329 |
| 632.78000 | 17.370 | 17.650 | 35.0 | Cluster_32445 |
| 632.84000 | 11.640 | 11.850 | 35.0 | Cluster_32450 |
| 633.00000 | 22.660 | 23.130 | 35.0 | Cluster_11339 |

| | | | | |
|---|---|---|---|---|
| 633.32000 | 14.850 | 15.230 | 35.0 | Cluster_32465 |
| 633.36000 | 25.310 | 25.480 | 35.0 | Cluster_32472 |
| 635.07000 | 22.620 | 23.020 | 35.0 | Cluster_02772 |
| 635.82000 | 25.280 | 25.600 | 35.0 | Cluster_02775 |
| 636.32000 | 31.510 | 31.820 | 35.0 | Cluster_32584 |
| 636.84000 | 14.840 | 15.110 | 35.0 | Cluster_32596 |
| 637.62000 | 7.690 | 8.090 | 35.0 | Cluster_11403 |
| 637.83000 | 17.450 | 17.890 | 35.0 | Cluster_32632 |
| 638.35000 | 22.570 | 22.950 | 35.0 | Cluster_32655 |
| 638.36000 | 27.860 | 28.070 | 35.0 | Cluster_11416 |
| 638.39000 | 20.000 | 20.130 | 35.0 | Cluster_32659 |
| 638.79000 | 7.640 | 7.920 | 35.0 | Cluster_32669 |
| 638.85000 | 17.450 | 17.760 | 35.0 | Cluster_32673 |
| 639.85000 | 11.870 | 12.150 | 35.0 | Cluster_32724 |
| 639.99000 | 11.800 | 12.120 | 35.0 | Cluster_11446 |
| 640.32000 | 19.880 | 20.330 | 35.0 | Cluster_32732 |
| 641.64000 | 19.900 | 20.110 | 35.0 | Cluster_11472 |
| 641.84000 | 14.670 | 15.030 | 35.0 | Cluster_32794 |
| 642.02000 | 25.230 | 25.570 | 35.0 | Cluster_11476 |
| 643.07000 | 25.200 | 25.430 | 35.0 | Cluster_32838 |
| 643.83000 | 11.760 | 12.250 | 35.0 | Cluster_32863 |
| 644.00000 | 31.400 | 31.630 | 35.0 | Cluster_11510 |
| 644.30000 | 7.630 | 8.960 | 35.0 | Cluster_02836 |
| 644.36000 | 27.970 | 28.160 | 35.0 | Cluster_32878 |
| 644.37000 | 25.080 | 25.430 | 35.0 | Cluster_11511 |
| 645.00000 | 31.710 | 31.970 | 35.0 | Cluster_11529 |
| 645.85000 | 14.780 | 14.940 | 35.0 | Cluster_32926 |
| 646.30000 | 7.730 | 8.000 | 35.0 | Cluster_32945 |
| 648.34000 | 27.840 | 28.040 | 35.0 | Cluster_33005 |
| 648.67000 | 20.000 | 20.410 | 35.0 | Cluster_11566 |
| 648.88000 | 17.540 | 17.890 | 35.0 | Cluster_33029 |
| 649.01000 | 22.600 | 23.360 | 35.0 | Cluster_11574 |
| 649.77000 | 7.630 | 7.950 | 35.0 | Cluster_33054 |
| 650.31000 | 17.480 | 17.840 | 35.0 | Cluster_33085 |
| 651.01000 | 22.660 | 23.440 | 35.0 | Cluster_11595 |
| 651.08000 | 22.660 | 23.610 | 35.0 | Cluster_33114 |
| 651.32000 | 25.070 | 25.170 | 35.0 | Cluster_33124 |
| 651.36000 | 25.060 | 26.830 | 35.0 | Cluster_11600 |
| 651.86000 | 31.600 | 32.050 | 35.0 | Cluster_33158 |
| 651.99000 | 22.630 | 23.030 | 35.0 | Cluster_11614 |
| 652.06000 | 22.590 | 23.040 | 35.0 | Cluster_33159 |
| 652.35000 | 17.450 | 17.730 | 35.0 | Cluster_33162 |
| 652.86000 | 11.860 | 12.170 | 35.0 | Cluster_33187 |
| 653.33000 | 22.490 | 23.010 | 35.0 | Cluster_33199 |
| 653.57000 | 14.650 | 14.940 | 35.0 | Cluster_33207 |
| 653.60000 | 31.520 | 31.800 | 35.0 | Cluster_02883 |
| 654.05000 | 22.470 | 22.620 | 35.0 | Cluster_33221 |
| 654.68000 | 14.810 | 15.140 | 35.0 | Cluster_11655 |
| 654.78000 | 7.650 | 7.850 | 35.0 | Cluster_33237 |
| 655.99000 | 14.600 | 14.720 | 35.0 | Cluster_11675 |
| 656.63000 | 14.820 | 15.370 | 35.0 | Cluster_11676 |

| | | | | |
|---|---|---|---|---|
| 657.90000 | 22.660 | 22.950 | 35.0 | Cluster_33331 |
| 657.95000 | 22.660 | 22.890 | 35.0 | Cluster_33350 |
| 658.07000 | 17.500 | 17.770 | 35.0 | Cluster_33340 |
| 658.66000 | 14.810 | 15.150 | 35.0 | Cluster_11699 |
| 659.28000 | 7.340 | 7.640 | 35.0 | Cluster_33386 |
| 659.60000 | 14.800 | 15.150 | 35.0 | Cluster_33396 |
| 659.63000 | 25.250 | 25.590 | 35.0 | Cluster_02907 |
| 659.98000 | 7.810 | 8.090 | 35.0 | Cluster_11725 |
| 660.33000 | 17.390 | 17.640 | 35.0 | Cluster_33416 |
| 660.46000 | 17.280 | 17.490 | 35.0 | Cluster_33423 |
| 660.84000 | 14.710 | 14.980 | 35.0 | Cluster_33430 |
| 661.01000 | 27.910 | 28.720 | 35.0 | Cluster_11741 |
| 661.31000 | 7.510 | 8.260 | 35.0 | Cluster_33436 |
| 662.02000 | 22.620 | 23.480 | 35.0 | Cluster_11748 |
| 662.12000 | 25.110 | 25.570 | 35.0 | Cluster_02928 |
| 662.30000 | 11.760 | 12.160 | 35.0 | Cluster_33472 |
| 663.37000 | 22.610 | 23.380 | 35.0 | Cluster_33509 |
| 663.55000 | 14.680 | 15.020 | 35.0 | Cluster_02941 |
| 663.60000 | 31.540 | 33.600 | 35.0 | Cluster_02940 |
| 665.32000 | 11.790 | 12.000 | 35.0 | Cluster_33570 |
| 666.35000 | 25.220 | 25.970 | 35.0 | Cluster_02964 |
| 666.68000 | 31.630 | 31.810 | 35.0 | Cluster_11814 |
| 666.95000 | 22.560 | 22.710 | 35.0 | Cluster_33617 |
| 667.82000 | 11.770 | 12.000 | 35.0 | Cluster_33644 |
| 668.33000 | 14.580 | 14.830 | 35.0 | Cluster_33651 |
| 668.35000 | 22.530 | 22.820 | 35.0 | Cluster_11848 |
| 669.10000 | 31.550 | 31.990 | 35.0 | Cluster_02977 |
| 669.33000 | 7.730 | 7.920 | 35.0 | Cluster_33680 |
| 669.78000 | 11.880 | 12.330 | 35.0 | Cluster_33695 |
| 670.01000 | 20.010 | 20.600 | 35.0 | Cluster_11876 |
| 670.40000 | 11.810 | 12.090 | 35.0 | Cluster_33725 |
| 670.80000 | 1.580 | 2.170 | 35.0 | Cluster_33734 |
| 670.82000 | 14.840 | 15.230 | 35.0 | Cluster_33742 |
| 670.87000 | 31.500 | 31.700 | 35.0 | Cluster_02986 |
| 671.44000 | 22.680 | 23.230 | 35.0 | Cluster_33760 |
| 671.82000 | 14.870 | 15.080 | 35.0 | Cluster_33770 |
| 671.97000 | 19.880 | 20.710 | 35.0 | Cluster_11894 |
| 672.35000 | 17.390 | 17.900 | 35.0 | Cluster_33781 |
| 672.81000 | 22.560 | 22.940 | 35.0 | Cluster_33789 |
| 673.09000 | 31.550 | 31.860 | 35.0 | Cluster_03005 |
| 673.37000 | 17.280 | 17.620 | 35.0 | Cluster_33802 |
| 673.62000 | 31.510 | 31.800 | 35.0 | Cluster_33824 |
| 673.65000 | 11.600 | 11.860 | 35.0 | Cluster_11927 |
| 674.11000 | 31.240 | 31.630 | 35.0 | Cluster_03014 |
| 674.32000 | 14.640 | 15.510 | 35.0 | Cluster_11933 |
| 674.95000 | 31.550 | 32.110 | 35.0 | Cluster_00274 |
| 675.37000 | 22.640 | 23.000 | 35.0 | Cluster_33861 |
| 675.60000 | 31.330 | 31.510 | 35.0 | Cluster_33864 |
| 675.64000 | 25.070 | 25.330 | 35.0 | Cluster_03025 |
| 675.84000 | 25.240 | 25.960 | 35.0 | Cluster_03026 |
| 676.00000 | 27.960 | 28.390 | 35.0 | Cluster_11956 |

| | | | | |
|---|---|---|---|---|
| 676.32000 | 17.510 | 17.840 | 35.0 | Cluster_11959 |
| 677.20000 | 1.550 | 1.800 | 35.0 | Cluster_33905 |
| 677.89000 | 17.290 | 17.370 | 35.0 | Cluster_33946 |
| 678.81000 | 14.610 | 14.810 | 35.0 | Cluster_33981 |
| 678.84000 | 19.900 | 20.130 | 35.0 | Cluster_33980 |
| 678.85000 | 25.320 | 25.840 | 35.0 | Cluster_03050 |
| 679.38000 | 31.570 | 31.860 | 35.0 | Cluster_33996 |
| 679.79000 | 7.500 | 7.750 | 35.0 | Cluster_34007 |
| 679.86000 | 22.690 | 23.500 | 35.0 | Cluster_34018 |
| 680.08000 | 25.280 | 25.680 | 35.0 | Cluster_03060 |
| 680.09000 | 19.920 | 20.480 | 35.0 | Cluster_34023 |
| 680.87000 | 14.720 | 15.220 | 35.0 | Cluster_34054 |
| 681.08000 | 25.320 | 25.720 | 35.0 | Cluster_03066 |
| 681.32000 | 17.470 | 17.820 | 35.0 | Cluster_12020 |
| 681.35000 | 11.710 | 11.930 | 35.0 | Cluster_34074 |
| 682.09000 | 25.270 | 25.680 | 35.0 | Cluster_34089 |
| 682.31000 | 1.380 | 1.610 | 35.0 | Cluster_34106 |
| 682.38000 | 31.630 | 31.920 | 35.0 | Cluster_03074 |
| 683.61000 | 17.540 | 17.710 | 35.0 | Cluster_12059 |
| 683.81000 | 14.650 | 15.140 | 35.0 | Cluster_34134 |
| 683.99000 | 11.820 | 12.300 | 35.0 | Cluster_12062 |
| 684.08000 | 31.210 | 31.860 | 35.0 | Cluster_03088 |
| 684.82000 | 17.510 | 17.670 | 35.0 | Cluster_34171 |
| 684.83000 | 20.060 | 20.260 | 35.0 | Cluster_34174 |
| 685.09000 | 31.700 | 32.020 | 35.0 | Cluster_03095 |
| 685.10000 | 27.910 | 28.210 | 35.0 | Cluster_03102 |
| 685.35000 | 25.210 | 25.420 | 35.0 | Cluster_34201 |
| 686.04000 | 19.890 | 20.200 | 35.0 | Cluster_12086 |
| 686.50000 | 27.930 | 28.200 | 35.0 | Cluster_00285 |
| 686.75000 | 17.440 | 17.750 | 35.0 | Cluster_00286 |
| 687.66000 | 27.960 | 28.390 | 35.0 | Cluster_34255 |
| 689.34000 | 11.570 | 11.990 | 35.0 | Cluster_34298 |
| 689.36000 | 14.790 | 15.320 | 35.0 | Cluster_34295 |
| 689.57000 | 31.350 | 31.700 | 35.0 | Cluster_03142 |
| 690.82000 | 7.740 | 8.350 | 35.0 | Cluster_34324 |
| 690.84000 | 14.610 | 15.040 | 35.0 | Cluster_34327 |
| 691.59000 | 22.500 | 22.700 | 35.0 | Cluster_03162 |
| 691.79000 | 7.360 | 7.590 | 35.0 | Cluster_34357 |
| 691.83000 | 25.240 | 25.490 | 35.0 | Cluster_34369 |
| 692.26000 | 7.370 | 7.970 | 35.0 | Cluster_34373 |
| 692.32000 | 19.900 | 20.660 | 35.0 | Cluster_03167 |
| 692.35000 | 22.680 | 22.990 | 35.0 | Cluster_03168 |
| 692.60000 | 31.670 | 32.670 | 35.0 | Cluster_03182 |
| 692.84000 | 22.550 | 22.680 | 35.0 | Cluster_34402 |
| 693.56000 | 31.300 | 31.830 | 35.0 | Cluster_03192 |
| 693.60000 | 19.950 | 20.630 | 35.0 | Cluster_34413 |
| 693.88000 | 14.790 | 15.050 | 35.0 | Cluster_34418 |
| 694.36000 | 11.580 | 11.990 | 35.0 | Cluster_34424 |
| 694.84000 | 22.670 | 22.890 | 35.0 | Cluster_34436 |
| 694.87000 | 14.830 | 15.400 | 35.0 | Cluster_34443 |
| 695.01000 | 19.930 | 20.930 | 35.0 | Cluster_12183 |

| | | | | |
|---|---|---|---|---|
| 695.36000 | 11.820 | 12.170 | 35.0 | Cluster_34458 |
| 695.70000 | 25.120 | 25.630 | 35.0 | Cluster_12190 |
| 695.87000 | 14.660 | 14.860 | 35.0 | Cluster_34468 |
| 696.32000 | 7.450 | 7.550 | 35.0 | Cluster_34489 |
| 696.34000 | 1.370 | 1.810 | 35.0 | Cluster_03210 |
| 696.39000 | 17.310 | 17.590 | 35.0 | Cluster_34483 |
| 696.58000 | 25.070 | 25.430 | 35.0 | Cluster_03211 |
| 697.31000 | 31.260 | 31.680 | 35.0 | Cluster_03215 |
| 697.37000 | 31.590 | 31.830 | 35.0 | Cluster_34515 |
| 697.85000 | 14.600 | 15.110 | 35.0 | Cluster_34520 |
| 697.87000 | 17.490 | 17.790 | 35.0 | Cluster_34523 |
| 698.82000 | 17.530 | 17.830 | 35.0 | Cluster_34541 |
| 698.83000 | 11.730 | 12.010 | 35.0 | Cluster_34546 |
| 699.70000 | 19.920 | 20.280 | 35.0 | Cluster_12244 |
| 699.85000 | 14.630 | 14.990 | 35.0 | Cluster_03235 |
| 700.26000 | 1.650 | 1.780 | 35.0 | Cluster_34580 |
| 700.36000 | 22.500 | 22.890 | 35.0 | Cluster_34589 |
| 700.65000 | 22.540 | 22.880 | 35.0 | Cluster_34590 |
| 700.85000 | 11.860 | 12.200 | 35.0 | Cluster_34594 |
| 701.36000 | 14.670 | 14.980 | 35.0 | Cluster_34614 |
| 702.34000 | 25.110 | 25.300 | 35.0 | Cluster_34653 |
| 702.62000 | 14.690 | 14.940 | 35.0 | Cluster_12286 |
| 703.04000 | 19.990 | 20.450 | 35.0 | Cluster_12291 |
| 703.05000 | 31.270 | 31.700 | 35.0 | Cluster_03266 |
| 703.33000 | 17.280 | 17.830 | 35.0 | Cluster_34677 |
| 703.69000 | 25.310 | 25.600 | 35.0 | Cluster_12310 |
| 703.91000 | 22.540 | 22.890 | 35.0 | Cluster_03274 |
| 704.17000 | 22.580 | 22.900 | 35.0 | Cluster_34698 |
| 704.21000 | 1.560 | 1.920 | 35.0 | Cluster_34700 |
| 704.31000 | 7.790 | 7.920 | 35.0 | Cluster_34706 |
| 704.81000 | 7.820 | 8.870 | 35.0 | Cluster_34728 |
| 704.85000 | 28.070 | 28.260 | 35.0 | Cluster_34734 |
| 705.04000 | 31.460 | 33.040 | 35.0 | Cluster_12319 |
| 705.37000 | 19.890 | 20.240 | 35.0 | Cluster_34742 |
| 705.83000 | 14.660 | 15.040 | 35.0 | Cluster_34756 |
| 705.88000 | 11.630 | 12.130 | 35.0 | Cluster_34757 |
| 706.86000 | 22.650 | 22.880 | 35.0 | Cluster_03301 |
| 707.02000 | 25.090 | 25.400 | 35.0 | Cluster_12346 |
| 707.80000 | 7.820 | 8.290 | 35.0 | Cluster_34791 |
| 708.51000 | 11.650 | 12.330 | 35.0 | Cluster_00309 |
| 709.28000 | 7.800 | 8.070 | 35.0 | Cluster_34840 |
| 709.39000 | 17.500 | 17.840 | 35.0 | Cluster_34846 |
| 710.88000 | 25.290 | 25.490 | 35.0 | Cluster_03335 |
| 711.06000 | 22.690 | 23.590 | 35.0 | Cluster_12397 |
| 711.85000 | 11.860 | 12.330 | 35.0 | Cluster_34925 |
| 712.39000 | 22.630 | 22.910 | 35.0 | Cluster_34948 |
| 712.84000 | 11.860 | 12.340 | 35.0 | Cluster_34967 |
| 713.08000 | 28.070 | 28.340 | 35.0 | Cluster_03351 |
| 713.29000 | 14.660 | 14.920 | 35.0 | Cluster_03356 |
| 713.38000 | 7.720 | 8.060 | 35.0 | Cluster_12431 |
| 713.72000 | 31.630 | 33.460 | 35.0 | Cluster_12437 |

| | | | | |
|---|---|---|---|---|
| 713.77000 | 7.320 | 7.800 | 35.0 | Cluster_34987 |
| 713.92000 | 14.850 | 15.200 | 35.0 | Cluster_34998 |
| 714.32000 | 17.390 | 17.720 | 35.0 | Cluster_35004 |
| 714.36000 | 19.930 | 20.350 | 35.0 | Cluster_35010 |
| 715.38000 | 19.960 | 20.320 | 35.0 | Cluster_35041 |
| 715.54000 | 11.860 | 12.010 | 35.0 | Cluster_00320 |
| 715.71000 | 14.660 | 14.920 | 35.0 | Cluster_12456 |
| 715.85000 | 17.470 | 17.620 | 35.0 | Cluster_35046 |
| 715.99000 | 31.390 | 31.720 | 35.0 | Cluster_12465 |
| 716.39000 | 19.880 | 20.360 | 35.0 | Cluster_35061 |
| 717.36000 | 31.540 | 32.090 | 35.0 | Cluster_12478 |
| 717.90000 | 25.120 | 25.550 | 35.0 | Cluster_35102 |
| 718.36000 | 19.940 | 20.380 | 35.0 | Cluster_35122 |
| 718.85000 | 7.680 | 8.400 | 35.0 | Cluster_35127 |
| 718.89000 | 17.510 | 17.800 | 35.0 | Cluster_35137 |
| 718.91000 | 27.800 | 28.330 | 35.0 | Cluster_35135 |
| 718.96000 | 28.020 | 28.340 | 35.0 | Cluster_12511 |
| 719.39000 | 19.980 | 20.260 | 35.0 | Cluster_35145 |
| 719.83000 | 11.610 | 12.340 | 35.0 | Cluster_35161 |
| 719.85000 | 25.100 | 25.670 | 35.0 | Cluster_03411 |
| 719.98000 | 19.940 | 20.400 | 35.0 | Cluster_12519 |
| 720.12000 | 25.110 | 25.240 | 35.0 | Cluster_35168 |
| 720.13000 | 28.000 | 28.180 | 35.0 | Cluster_35170 |
| 720.35000 | 17.530 | 17.700 | 35.0 | Cluster_03417 |
| 720.58000 | 17.300 | 17.590 | 35.0 | Cluster_03421 |
| 720.83000 | 7.750 | 8.350 | 35.0 | Cluster_35178 |
| 721.32000 | 19.890 | 20.270 | 35.0 | Cluster_35195 |
| 721.41000 | 17.260 | 17.540 | 35.0 | Cluster_35191 |
| 721.72000 | 22.640 | 22.890 | 35.0 | Cluster_12548 |
| 721.88000 | 22.670 | 23.060 | 35.0 | Cluster_35209 |
| 721.90000 | 25.050 | 25.270 | 35.0 | Cluster_35213 |
| 722.02000 | 31.660 | 32.060 | 35.0 | Cluster_12553 |
| 722.09000 | 28.070 | 28.500 | 35.0 | Cluster_03436 |
| 722.34000 | 25.130 | 26.370 | 35.0 | Cluster_03433 |
| 722.68000 | 31.370 | 32.020 | 35.0 | Cluster_12565 |
| 723.40000 | 19.960 | 20.360 | 35.0 | Cluster_12568 |
| 723.42000 | 22.560 | 22.940 | 35.0 | Cluster_12570 |
| 723.95000 | 27.910 | 28.520 | 35.0 | Cluster_00326 |
| 724.04000 | 31.210 | 31.650 | 35.0 | Cluster_12575 |
| 724.90000 | 22.470 | 22.960 | 35.0 | Cluster_03465 |
| 725.06000 | 27.970 | 28.270 | 35.0 | Cluster_12593 |
| 725.32000 | 7.780 | 8.500 | 35.0 | Cluster_03469 |
| 725.41000 | 17.370 | 17.560 | 35.0 | Cluster_35286 |
| 725.42000 | 31.360 | 31.600 | 35.0 | Cluster_35293 |
| 727.38000 | 25.090 | 25.480 | 35.0 | Cluster_35317 |
| 728.10000 | 25.290 | 25.660 | 35.0 | Cluster_03487 |
| 728.13000 | 22.560 | 22.730 | 35.0 | Cluster_03484 |
| 728.42000 | 25.140 | 25.320 | 35.0 | Cluster_35338 |
| 728.98000 | 31.520 | 31.690 | 35.0 | Cluster_12653 |
| 729.15000 | 31.210 | 31.440 | 35.0 | Cluster_03497 |
| 729.84000 | 17.500 | 17.710 | 35.0 | Cluster_35373 |

| | | | | |
|---|---|---|---|---|
| 729.89000 | 22.500 | 22.710 | 35.0 | Cluster_03507 |
| 730.91000 | 22.680 | 22.870 | 35.0 | Cluster_35408 |
| 731.10000 | 28.060 | 28.490 | 35.0 | Cluster_35410 |
| 731.83000 | 25.260 | 25.650 | 35.0 | Cluster_03518 |
| 732.11000 | 17.410 | 17.790 | 35.0 | Cluster_03522 |
| 732.64000 | 31.490 | 31.690 | 35.0 | Cluster_03534 |
| 733.03000 | 17.410 | 17.820 | 35.0 | Cluster_12708 |
| 733.13000 | 25.140 | 25.560 | 35.0 | Cluster_03548 |
| 733.83000 | 11.880 | 12.390 | 35.0 | Cluster_35471 |
| 733.85000 | 14.650 | 15.150 | 35.0 | Cluster_35474 |
| 734.58000 | 7.790 | 8.040 | 35.0 | Cluster_35493 |
| 735.04000 | 20.040 | 20.350 | 35.0 | Cluster_12729 |
| 735.74000 | 17.500 | 17.780 | 35.0 | Cluster_12741 |
| 736.37000 | 11.800 | 12.200 | 35.0 | Cluster_35541 |
| 737.04000 | 27.800 | 28.310 | 35.0 | Cluster_12754 |
| 737.16000 | 22.680 | 22.850 | 35.0 | Cluster_35560 |
| 737.82000 | 7.630 | 8.050 | 35.0 | Cluster_35593 |
| 738.39000 | 19.880 | 20.480 | 35.0 | Cluster_35611 |
| 738.63000 | 11.820 | 12.200 | 35.0 | Cluster_03590 |
| 739.38000 | 27.950 | 28.260 | 35.0 | Cluster_12791 |
| 739.39000 | 27.810 | 27.990 | 35.0 | Cluster_12789 |
| 739.84000 | 1.420 | 1.740 | 35.0 | Cluster_35645 |
| 740.37000 | 14.720 | 15.210 | 35.0 | Cluster_35657 |
| 740.38000 | 22.620 | 22.940 | 35.0 | Cluster_35661 |
| 740.93000 | 27.860 | 28.230 | 35.0 | Cluster_35676 |
| 741.85000 | 17.470 | 17.680 | 35.0 | Cluster_35683 |
| 742.37000 | 11.870 | 12.390 | 35.0 | Cluster_35697 |
| 742.67000 | 7.750 | 8.360 | 35.0 | Cluster_12831 |
| 742.84000 | 14.750 | 15.520 | 35.0 | Cluster_35707 |
| 744.37000 | 25.210 | 25.740 | 35.0 | Cluster_35750 |
| 745.38000 | 25.200 | 25.420 | 35.0 | Cluster_35777 |
| 745.68000 | 22.680 | 23.580 | 35.0 | Cluster_12857 |
| 746.06000 | 22.680 | 22.970 | 35.0 | Cluster_12863 |
| 746.15000 | 28.030 | 28.180 | 35.0 | Cluster_35792 |
| 746.30000 | 7.780 | 8.080 | 35.0 | Cluster_35797 |
| 747.12000 | 11.800 | 12.230 | 35.0 | Cluster_35817 |
| 747.37000 | 11.710 | 12.180 | 35.0 | Cluster_35818 |
| 748.00000 | 22.610 | 22.950 | 35.0 | Cluster_12903 |
| 748.86000 | 27.800 | 28.150 | 35.0 | Cluster_03654 |
| 748.88000 | 19.970 | 20.640 | 35.0 | Cluster_35857 |
| 749.07000 | 19.960 | 20.200 | 35.0 | Cluster_12920 |
| 749.69000 | 17.250 | 17.550 | 35.0 | Cluster_12929 |
| 749.98000 | 19.860 | 20.090 | 35.0 | Cluster_12936 |
| 750.01000 | 22.600 | 23.500 | 35.0 | Cluster_12941 |
| 750.38000 | 14.680 | 14.960 | 35.0 | Cluster_35883 |
| 750.44000 | 22.600 | 22.990 | 35.0 | Cluster_35886 |
| 750.94000 | 22.590 | 23.000 | 35.0 | Cluster_35903 |
| 751.06000 | 28.050 | 28.290 | 35.0 | Cluster_12961 |
| 751.15000 | 25.250 | 25.670 | 35.0 | Cluster_03662 |
| 751.36000 | 14.640 | 15.100 | 35.0 | Cluster_35908 |
| 752.37000 | 11.840 | 12.380 | 35.0 | Cluster_35932 |

| | | | | |
|---|---|---|---|---|
| 753.05000 | 31.680 | 31.970 | 35.0 | Cluster_12988 |
| 753.06000 | 25.170 | 25.500 | 35.0 | Cluster_12984 |
| 753.39000 | 22.670 | 22.900 | 35.0 | Cluster_35964 |
| 753.91000 | 27.920 | 28.160 | 35.0 | Cluster_35981 |
| 754.06000 | 20.050 | 20.510 | 35.0 | Cluster_13009 |
| 754.90000 | 14.660 | 15.500 | 35.0 | Cluster_36004 |
| 755.06000 | 25.260 | 25.460 | 35.0 | Cluster_13021 |
| 755.30000 | 20.050 | 20.270 | 35.0 | Cluster_13019 |
| 755.33000 | 22.630 | 23.190 | 35.0 | Cluster_13020 |
| 755.42000 | 19.910 | 20.150 | 35.0 | Cluster_36013 |
| 755.75000 | 17.360 | 17.740 | 35.0 | Cluster_13034 |
| 755.92000 | 19.960 | 20.150 | 35.0 | Cluster_36043 |
| 756.13000 | 25.270 | 25.620 | 35.0 | Cluster_03688 |
| 757.31000 | 14.830 | 14.950 | 35.0 | Cluster_36071 |
| 757.73000 | 27.890 | 28.230 | 35.0 | Cluster_13076 |
| 758.41000 | 25.080 | 25.600 | 35.0 | Cluster_36102 |
| 758.70000 | 14.580 | 14.990 | 35.0 | Cluster_13088 |
| 759.63000 | 22.600 | 22.980 | 35.0 | Cluster_03704 |
| 759.90000 | 14.610 | 14.870 | 35.0 | Cluster_36129 |
| 759.91000 | 17.320 | 17.630 | 35.0 | Cluster_36132 |
| 760.35000 | 14.800 | 15.030 | 35.0 | Cluster_36146 |
| 761.05000 | 17.490 | 17.830 | 35.0 | Cluster_00024 |
| 761.41000 | 17.280 | 17.520 | 35.0 | Cluster_36182 |
| 762.38000 | 27.960 | 28.200 | 35.0 | Cluster_13130 |
| 763.03000 | 7.560 | 9.080 | 35.0 | Cluster_13141 |
| 763.08000 | 17.400 | 17.840 | 35.0 | Cluster_13137 |
| 763.35000 | 25.210 | 26.120 | 35.0 | Cluster_03726 |
| 763.71000 | 31.480 | 31.660 | 35.0 | Cluster_13147 |
| 764.09000 | 31.660 | 32.060 | 35.0 | Cluster_13148 |
| 764.36000 | 11.800 | 12.080 | 35.0 | Cluster_36253 |
| 765.15000 | 31.680 | 31.980 | 35.0 | Cluster_03741 |
| 765.79000 | 11.610 | 11.860 | 35.0 | Cluster_36274 |
| 765.80000 | 11.860 | 12.080 | 35.0 | Cluster_36275 |
| 766.37000 | 11.600 | 11.750 | 35.0 | Cluster_36292 |
| 766.55000 | 17.450 | 18.040 | 35.0 | Cluster_00339 |
| 767.70000 | 27.940 | 28.270 | 35.0 | Cluster_13203 |
| 767.85000 | 11.790 | 12.130 | 35.0 | Cluster_36325 |
| 768.07000 | 17.280 | 17.780 | 35.0 | Cluster_13209 |
| 768.08000 | 31.360 | 31.640 | 35.0 | Cluster_13205 |
| 768.38000 | 7.480 | 7.660 | 35.0 | Cluster_36351 |
| 769.33000 | 22.640 | 22.980 | 35.0 | Cluster_36370 |
| 769.59000 | 19.930 | 20.190 | 35.0 | Cluster_36368 |
| 769.82000 | 19.960 | 20.600 | 35.0 | Cluster_36385 |
| 772.41000 | 22.650 | 23.180 | 35.0 | Cluster_36441 |
| 772.74000 | 25.070 | 25.310 | 35.0 | Cluster_13265 |
| 772.80000 | 31.680 | 32.590 | 35.0 | Cluster_13273 |
| 772.88000 | 14.580 | 14.710 | 35.0 | Cluster_36459 |
| 772.92000 | 14.670 | 14.950 | 35.0 | Cluster_36460 |
| 773.08000 | 19.950 | 20.220 | 35.0 | Cluster_03785 |
| 773.09000 | 20.050 | 20.330 | 35.0 | Cluster_03786 |
| 773.13000 | 31.430 | 31.710 | 35.0 | Cluster_13272 |

| | | | | |
|---|---|---|---|---|
| 773.35000 | 14.740 | 15.130 | 35.0 | Cluster_13283 |
| 773.41000 | 27.880 | 28.470 | 35.0 | Cluster_13286 |
| 774.12000 | 20.050 | 20.350 | 35.0 | Cluster_36481 |
| 774.73000 | 17.470 | 18.010 | 35.0 | Cluster_13306 |
| 775.06000 | 11.690 | 12.150 | 35.0 | Cluster_13312 |
| 775.07000 | 19.940 | 20.600 | 35.0 | Cluster_13311 |
| 775.42000 | 27.800 | 28.440 | 35.0 | Cluster_36510 |
| 776.37000 | 22.480 | 22.750 | 35.0 | Cluster_03803 |
| 777.15000 | 27.810 | 28.220 | 35.0 | Cluster_03806 |
| 777.35000 | 11.810 | 12.190 | 35.0 | Cluster_36561 |
| 777.38000 | 25.120 | 25.450 | 35.0 | Cluster_36567 |
| 777.39000 | 14.650 | 14.840 | 35.0 | Cluster_36564 |
| 778.37000 | 17.250 | 17.390 | 35.0 | Cluster_36591 |
| 778.77000 | 25.180 | 25.660 | 35.0 | Cluster_13350 |
| 778.87000 | 17.380 | 17.700 | 35.0 | Cluster_36605 |
| 780.73000 | 27.980 | 28.270 | 35.0 | Cluster_13379 |
| 780.81000 | 19.910 | 20.480 | 35.0 | Cluster_36636 |
| 780.88000 | 31.460 | 32.210 | 35.0 | Cluster_03821 |
| 781.40000 | 31.410 | 31.620 | 35.0 | Cluster_36661 |
| 782.03000 | 11.610 | 12.170 | 35.0 | Cluster_13403 |
| 783.13000 | 22.500 | 22.860 | 35.0 | Cluster_03834 |
| 783.46000 | 25.110 | 25.420 | 35.0 | Cluster_36708 |
| 783.72000 | 19.910 | 20.190 | 35.0 | Cluster_13422 |
| 784.06000 | 27.900 | 28.250 | 35.0 | Cluster_13433 |
| 784.89000 | 1.480 | 1.760 | 35.0 | Cluster_36733 |
| 784.93000 | 25.120 | 25.800 | 35.0 | Cluster_36732 |
| 785.45000 | 19.890 | 20.320 | 35.0 | Cluster_36744 |
| 785.83000 | 22.660 | 22.780 | 35.0 | Cluster_36763 |
| 787.06000 | 28.010 | 28.200 | 35.0 | Cluster_13459 |
| 787.12000 | 22.530 | 22.900 | 35.0 | Cluster_03868 |
| 787.35000 | 19.870 | 20.400 | 35.0 | Cluster_03870 |
| 788.87000 | 22.600 | 22.810 | 35.0 | Cluster_36840 |
| 789.06000 | 31.630 | 32.000 | 35.0 | Cluster_13483 |
| 790.87000 | 31.560 | 32.090 | 35.0 | Cluster_36906 |
| 791.12000 | 22.590 | 22.780 | 35.0 | Cluster_36909 |
| 791.38000 | 25.170 | 25.540 | 35.0 | Cluster_36914 |
| 791.90000 | 27.890 | 28.500 | 35.0 | Cluster_36919 |
| 792.38000 | 31.390 | 31.540 | 35.0 | Cluster_36933 |
| 792.64000 | 17.450 | 17.820 | 35.0 | Cluster_36941 |
| 792.86000 | 22.690 | 23.220 | 35.0 | Cluster_03914 |
| 792.87000 | 31.460 | 31.670 | 35.0 | Cluster_36948 |
| 792.97000 | 14.580 | 14.860 | 35.0 | Cluster_13541 |
| 793.78000 | 31.320 | 31.590 | 35.0 | Cluster_13547 |
| 794.36000 | 31.670 | 32.080 | 35.0 | Cluster_36984 |
| 794.37000 | 19.940 | 20.210 | 35.0 | Cluster_36982 |
| 794.45000 | 1.420 | 1.790 | 35.0 | Cluster_36985 |
| 794.65000 | 17.390 | 17.780 | 35.0 | Cluster_36986 |
| 795.07000 | 25.160 | 25.360 | 35.0 | Cluster_13571 |
| 795.42000 | 22.670 | 22.790 | 35.0 | Cluster_37004 |
| 795.44000 | 1.400 | 1.800 | 35.0 | Cluster_37000 |
| 795.68000 | 14.590 | 14.880 | 35.0 | Cluster_13585 |

| | | | | |
|---|---|---|---|---|
| 795.87000 | 31.560 | 32.170 | 35.0 | Cluster_03947 |
| 795.90000 | 28.060 | 28.270 | 35.0 | Cluster_37017 |
| 795.96000 | 14.760 | 15.070 | 35.0 | Cluster_37020 |
| 796.41000 | 31.450 | 31.680 | 35.0 | Cluster_13605 |
| 796.72000 | 19.940 | 20.100 | 35.0 | Cluster_13597 |
| 796.91000 | 22.500 | 22.720 | 35.0 | Cluster_37044 |
| 797.86000 | 19.930 | 20.440 | 35.0 | Cluster_37077 |
| 797.87000 | 7.430 | 7.730 | 35.0 | Cluster_37073 |
| 797.88000 | 7.310 | 7.440 | 35.0 | Cluster_37072 |
| 797.89000 | 22.500 | 22.690 | 35.0 | Cluster_37068 |
| 799.38000 | 22.500 | 23.460 | 35.0 | Cluster_37115 |
| 799.40000 | 14.770 | 15.530 | 35.0 | Cluster_37112 |
| 799.89000 | 31.620 | 32.180 | 35.0 | Cluster_03975 |
| 800.35000 | 31.530 | 31.630 | 35.0 | Cluster_37146 |
| 800.41000 | 19.890 | 20.120 | 35.0 | Cluster_13643 |
| 800.84000 | 31.290 | 31.550 | 35.0 | Cluster_37149 |
| 800.86000 | 31.650 | 32.090 | 35.0 | Cluster_03991 |
| 801.06000 | 11.650 | 12.070 | 35.0 | Cluster_00029 |
| 801.37000 | 17.480 | 17.810 | 35.0 | Cluster_03995 |
| 801.42000 | 25.180 | 25.660 | 35.0 | Cluster_37163 |
| 801.44000 | 31.270 | 31.660 | 35.0 | Cluster_13660 |
| 802.43000 | 20.050 | 20.550 | 35.0 | Cluster_37188 |
| 802.74000 | 27.900 | 28.080 | 35.0 | Cluster_13681 |
| 803.41000 | 25.130 | 25.600 | 35.0 | Cluster_13684 |
| 803.88000 | 17.370 | 17.770 | 35.0 | Cluster_37226 |
| 804.43000 | 22.650 | 22.870 | 35.0 | Cluster_37243 |
| 804.91000 | 31.690 | 32.140 | 35.0 | Cluster_04015 |
| 805.16000 | 14.600 | 14.940 | 35.0 | Cluster_37269 |
| 805.39000 | 27.930 | 28.290 | 35.0 | Cluster_37283 |
| 805.94000 | 31.440 | 31.630 | 35.0 | Cluster_37284 |
| 806.39000 | 14.640 | 14.860 | 35.0 | Cluster_37304 |
| 807.01000 | 19.980 | 20.440 | 35.0 | Cluster_13723 |
| 807.46000 | 19.920 | 20.080 | 35.0 | Cluster_37330 |
| 807.92000 | 27.870 | 28.030 | 35.0 | Cluster_37352 |
| 808.85000 | 19.960 | 20.450 | 35.0 | Cluster_37370 |
| 808.91000 | 31.470 | 31.730 | 35.0 | Cluster_37377 |
| 809.40000 | 22.660 | 23.070 | 35.0 | Cluster_04037 |
| 809.91000 | 14.850 | 15.310 | 35.0 | Cluster_37401 |
| 810.39000 | 19.920 | 20.390 | 35.0 | Cluster_37407 |
| 810.44000 | 22.630 | 23.110 | 35.0 | Cluster_13774 |
| 811.74000 | 22.480 | 22.860 | 35.0 | Cluster_00036 |
| 811.86000 | 20.040 | 20.560 | 35.0 | Cluster_37446 |
| 811.91000 | 14.870 | 15.290 | 35.0 | Cluster_37441 |
| 812.39000 | 22.630 | 22.830 | 35.0 | Cluster_37458 |
| 812.41000 | 27.930 | 28.400 | 35.0 | Cluster_13796 |
| 813.82000 | 11.790 | 12.190 | 35.0 | Cluster_00352 |
| 813.90000 | 17.360 | 17.730 | 35.0 | Cluster_37483 |
| 814.10000 | 25.250 | 25.470 | 35.0 | Cluster_13827 |
| 814.16000 | 27.890 | 28.190 | 35.0 | Cluster_37492 |
| 814.17000 | 25.120 | 25.320 | 35.0 | Cluster_04067 |
| 814.90000 | 22.680 | 23.030 | 35.0 | Cluster_04074 |

| | | | | |
|---|---|---|---|---|
| 815.06000 | 20.000 | 20.430 | 35.0 | Cluster_13832 |
| 815.79000 | 31.570 | 31.790 | 35.0 | Cluster_13844 |
| 815.92000 | 31.670 | 31.980 | 35.0 | Cluster_37530 |
| 816.38000 | 11.830 | 12.230 | 35.0 | Cluster_37540 |
| 816.43000 | 17.290 | 17.640 | 35.0 | Cluster_04088 |
| 816.72000 | 25.320 | 25.810 | 35.0 | Cluster_13857 |
| 817.34000 | 17.250 | 17.390 | 35.0 | Cluster_37561 |
| 817.88000 | 17.360 | 17.810 | 35.0 | Cluster_37570 |
| 818.42000 | 14.620 | 15.040 | 35.0 | Cluster_13883 |
| 819.05000 | 25.150 | 26.050 | 35.0 | Cluster_13897 |
| 819.42000 | 27.910 | 28.360 | 35.0 | Cluster_13906 |
| 820.17000 | 31.380 | 32.010 | 35.0 | Cluster_04104 |
| 821.23000 | 31.570 | 34.850 | 35.0 | Cluster_04111 |
| 821.38000 | 17.280 | 17.680 | 35.0 | Cluster_37637 |
| 822.06000 | 19.930 | 20.470 | 35.0 | Cluster_13934 |
| 823.42000 | 17.480 | 17.870 | 35.0 | Cluster_37669 |
| 823.91000 | 19.880 | 20.220 | 35.0 | Cluster_37674 |
| 823.94000 | 31.630 | 31.990 | 35.0 | Cluster_37688 |
| 825.32000 | 17.260 | 17.390 | 35.0 | Cluster_37717 |
| 825.39000 | 31.380 | 31.780 | 35.0 | Cluster_04137 |
| 826.38000 | 25.130 | 25.960 | 35.0 | Cluster_13981 |
| 826.86000 | 7.610 | 8.080 | 35.0 | Cluster_37765 |
| 827.04000 | 25.110 | 25.810 | 35.0 | Cluster_13986 |
| 827.06000 | 25.130 | 25.350 | 35.0 | Cluster_13988 |
| 827.09000 | 27.900 | 28.200 | 35.0 | Cluster_13989 |
| 827.88000 | 17.410 | 17.760 | 35.0 | Cluster_37794 |
| 828.80000 | 31.260 | 31.860 | 35.0 | Cluster_14000 |
| 828.98000 | 11.690 | 11.970 | 35.0 | Cluster_14004 |
| 829.16000 | 28.030 | 28.450 | 35.0 | Cluster_04165 |
| 829.36000 | 17.310 | 17.660 | 35.0 | Cluster_37821 |
| 829.43000 | 1.390 | 1.780 | 35.0 | Cluster_37825 |
| 829.91000 | 31.400 | 31.980 | 35.0 | Cluster_04169 |
| 830.74000 | 25.120 | 25.400 | 35.0 | Cluster_14020 |
| 830.89000 | 17.440 | 17.890 | 35.0 | Cluster_00042 |
| 831.45000 | 22.630 | 22.880 | 35.0 | Cluster_37853 |
| 832.41000 | 19.980 | 20.380 | 35.0 | Cluster_37873 |
| 832.68000 | 27.950 | 28.150 | 35.0 | Cluster_37879 |
| 833.49000 | 25.150 | 25.550 | 35.0 | Cluster_37894 |
| 833.99000 | 25.080 | 25.180 | 35.0 | Cluster_37902 |
| 834.10000 | 27.890 | 28.190 | 35.0 | Cluster_14054 |
| 834.94000 | 17.250 | 17.590 | 35.0 | Cluster_37918 |
| 835.05000 | 11.600 | 12.000 | 35.0 | Cluster_14062 |
| 836.39000 | 25.120 | 25.370 | 35.0 | Cluster_14076 |
| 836.42000 | 28.040 | 28.340 | 35.0 | Cluster_14077 |
| 836.76000 | 7.520 | 8.200 | 35.0 | Cluster_14083 |
| 836.86000 | 17.310 | 17.800 | 35.0 | Cluster_37959 |
| 836.94000 | 31.630 | 32.340 | 35.0 | Cluster_04234 |
| 837.41000 | 31.230 | 31.570 | 35.0 | Cluster_37970 |
| 837.95000 | 25.170 | 25.490 | 35.0 | Cluster_37976 |
| 839.10000 | 31.510 | 32.010 | 35.0 | Cluster_14102 |
| 840.66000 | 31.220 | 31.560 | 35.0 | Cluster_04260 |

| | | | | |
|---|---|---|---|---|
| 840.78000 | 25.110 | 25.290 | 35.0 | Cluster_14117 |
| 841.08000 | 22.570 | 22.840 | 35.0 | Cluster_14128 |
| 841.10000 | 19.960 | 20.190 | 35.0 | Cluster_14126 |
| 841.67000 | 25.300 | 25.610 | 35.0 | Cluster_00364 |
| 841.76000 | 27.960 | 28.220 | 35.0 | Cluster_38066 |
| 842.73000 | 25.140 | 25.740 | 35.0 | Cluster_14140 |
| 843.87000 | 17.390 | 17.890 | 35.0 | Cluster_38114 |
| 844.18000 | 31.530 | 32.150 | 35.0 | Cluster_04298 |
| 844.44000 | 25.120 | 25.510 | 35.0 | Cluster_14157 |
| 846.98000 | 1.510 | 1.880 | 35.0 | Cluster_38180 |
| 847.48000 | 1.390 | 1.840 | 35.0 | Cluster_38200 |
| 847.75000 | 19.880 | 20.160 | 35.0 | Cluster_14185 |
| 847.88000 | 11.560 | 11.920 | 35.0 | Cluster_38203 |
| 848.40000 | 27.880 | 28.510 | 35.0 | Cluster_04350 |
| 848.91000 | 31.520 | 32.250 | 35.0 | Cluster_04352 |
| 849.46000 | 25.150 | 25.670 | 35.0 | Cluster_38235 |
| 849.64000 | 27.920 | 28.360 | 35.0 | Cluster_04364 |
| 849.97000 | 22.620 | 23.510 | 35.0 | Cluster_38245 |
| 849.99000 | 25.120 | 25.700 | 35.0 | Cluster_38248 |
| 850.33000 | 14.660 | 14.780 | 35.0 | Cluster_38250 |
| 850.40000 | 19.990 | 20.460 | 35.0 | Cluster_04376 |
| 850.76000 | 19.870 | 20.160 | 35.0 | Cluster_14213 |
| 851.36000 | 11.750 | 12.340 | 35.0 | Cluster_14215 |
| 851.77000 | 31.300 | 31.540 | 35.0 | Cluster_38298 |
| 852.03000 | 22.650 | 23.290 | 35.0 | Cluster_38307 |
| 852.91000 | 22.610 | 23.310 | 35.0 | Cluster_04400 |
| 852.93000 | 31.510 | 32.160 | 35.0 | Cluster_04401 |
| 853.45000 | 20.010 | 20.160 | 35.0 | Cluster_38334 |
| 854.44000 | 20.050 | 20.750 | 35.0 | Cluster_38377 |
| 854.47000 | 22.680 | 23.080 | 35.0 | Cluster_38375 |
| 854.88000 | 22.650 | 22.930 | 35.0 | Cluster_38385 |
| 855.41000 | 27.900 | 28.200 | 35.0 | Cluster_38392 |
| 855.73000 | 17.250 | 17.700 | 35.0 | Cluster_14240 |
| 856.43000 | 25.110 | 25.490 | 35.0 | Cluster_14249 |
| 856.91000 | 31.520 | 32.060 | 35.0 | Cluster_04444 |
| 857.06000 | 17.480 | 17.920 | 35.0 | Cluster_14257 |
| 857.89000 | 14.810 | 15.530 | 35.0 | Cluster_38459 |
| 857.91000 | 27.910 | 28.720 | 35.0 | Cluster_04449 |
| 858.07000 | 11.840 | 12.350 | 35.0 | Cluster_14269 |
| 858.77000 | 25.070 | 25.380 | 35.0 | Cluster_14275 |
| 859.45000 | 17.420 | 18.280 | 35.0 | Cluster_38500 |
| 859.47000 | 7.790 | 8.140 | 35.0 | Cluster_38497 |
| 860.47000 | 31.240 | 31.540 | 35.0 | Cluster_14284 |
| 861.39000 | 27.870 | 28.200 | 35.0 | Cluster_04493 |
| 862.39000 | 22.610 | 23.110 | 35.0 | Cluster_04505 |
| 862.49000 | 25.140 | 25.580 | 35.0 | Cluster_38591 |
| 863.05000 | 14.600 | 15.530 | 35.0 | Cluster_14302 |
| 863.76000 | 22.540 | 23.030 | 35.0 | Cluster_14307 |
| 863.94000 | 25.070 | 25.410 | 35.0 | Cluster_38615 |
| 864.07000 | 14.650 | 14.990 | 35.0 | Cluster_14314 |
| 865.03000 | 17.390 | 17.520 | 35.0 | Cluster_38646 |

| | | | | |
|---|---|---|---|---|
| 865.75000 | 25.320 | 25.950 | 35.0 | Cluster_14326 |
| 865.92000 | 7.380 | 8.040 | 35.0 | Cluster_38659 |
| 865.93000 | 25.050 | 25.440 | 35.0 | Cluster_04555 |
| 866.41000 | 25.100 | 25.470 | 35.0 | Cluster_14334 |
| 869.07000 | 25.170 | 26.260 | 35.0 | Cluster_14353 |
| 869.28000 | 1.610 | 1.970 | 35.0 | Cluster_38719 |
| 869.41000 | 17.420 | 17.720 | 35.0 | Cluster_04599 |
| 869.78000 | 25.260 | 25.530 | 35.0 | Cluster_14363 |
| 871.17000 | 14.840 | 14.960 | 35.0 | Cluster_38758 |
| 871.27000 | 1.620 | 1.910 | 35.0 | Cluster_38759 |
| 871.44000 | 14.800 | 15.060 | 35.0 | Cluster_38769 |
| 871.47000 | 31.540 | 31.790 | 35.0 | Cluster_14388 |
| 872.27000 | 1.620 | 1.840 | 35.0 | Cluster_38802 |
| 874.09000 | 25.150 | 25.530 | 35.0 | Cluster_14414 |
| 874.26000 | 1.450 | 1.800 | 35.0 | Cluster_38844 |
| 874.78000 | 17.480 | 17.850 | 35.0 | Cluster_14424 |
| 875.28000 | 1.610 | 1.910 | 35.0 | Cluster_38860 |
| 875.66000 | 14.760 | 15.120 | 35.0 | Cluster_04698 |
| 876.08000 | 22.620 | 23.800 | 35.0 | Cluster_14437 |
| 876.53000 | 22.590 | 22.860 | 35.0 | Cluster_38895 |
| 877.38000 | 31.440 | 31.940 | 35.0 | Cluster_14453 |
| 877.44000 | 25.130 | 25.990 | 35.0 | Cluster_38909 |
| 878.48000 | 22.630 | 22.810 | 35.0 | Cluster_38928 |
| 880.46000 | 25.120 | 25.320 | 35.0 | Cluster_14483 |
| 881.96000 | 31.500 | 31.750 | 35.0 | Cluster_38993 |
| 882.44000 | 14.860 | 15.200 | 35.0 | Cluster_39008 |
| 882.94000 | 22.620 | 22.950 | 35.0 | Cluster_39018 |
| 884.40000 | 14.610 | 15.200 | 35.0 | Cluster_14528 |
| 884.66000 | 17.330 | 17.790 | 35.0 | Cluster_39047 |
| 885.12000 | 25.300 | 25.650 | 35.0 | Cluster_14544 |
| 885.68000 | 22.680 | 22.970 | 35.0 | Cluster_04765 |
| 885.95000 | 28.020 | 28.260 | 35.0 | Cluster_39078 |
| 886.43000 | 19.990 | 20.220 | 35.0 | Cluster_39095 |
| 886.96000 | 17.420 | 18.080 | 35.0 | Cluster_39099 |
| 887.52000 | 22.650 | 22.880 | 35.0 | Cluster_39107 |
| 887.68000 | 27.940 | 28.230 | 35.0 | Cluster_04781 |
| 887.74000 | 19.860 | 20.230 | 35.0 | Cluster_14571 |
| 888.13000 | 25.110 | 25.970 | 35.0 | Cluster_14577 |
| 888.40000 | 22.640 | 23.100 | 35.0 | Cluster_14578 |
| 888.91000 | 14.830 | 15.120 | 35.0 | Cluster_39135 |
| 888.92000 | 22.520 | 22.680 | 35.0 | Cluster_04786 |
| 889.11000 | 20.050 | 20.460 | 35.0 | Cluster_14581 |
| 889.44000 | 1.430 | 1.770 | 35.0 | Cluster_39147 |
| 890.17000 | 11.610 | 11.980 | 35.0 | Cluster_04796 |
| 890.38000 | 17.520 | 18.070 | 35.0 | Cluster_39161 |
| 890.39000 | 17.270 | 17.520 | 35.0 | Cluster_39160 |
| 890.46000 | 22.600 | 22.960 | 35.0 | Cluster_39173 |
| 890.99000 | 22.610 | 23.040 | 35.0 | Cluster_39180 |
| 891.46000 | 25.300 | 25.500 | 35.0 | Cluster_39186 |
| 891.81000 | 31.560 | 31.880 | 35.0 | Cluster_14598 |
| 892.21000 | 14.840 | 15.360 | 35.0 | Cluster_04824 |

| | | | | |
|---|---|---|---|---|
| 893.43000 | 27.950 | 28.230 | 35.0 | Cluster_39231 |
| 893.78000 | 19.930 | 20.280 | 35.0 | Cluster_14615 |
| 894.12000 | 22.690 | 23.160 | 35.0 | Cluster_14617 |
| 894.71000 | 27.820 | 28.230 | 35.0 | Cluster_04845 |
| 895.21000 | 14.810 | 15.350 | 35.0 | Cluster_04851 |
| 895.45000 | 25.300 | 25.980 | 35.0 | Cluster_14622 |
| 895.81000 | 27.910 | 28.270 | 35.0 | Cluster_14625 |
| 896.15000 | 31.390 | 31.880 | 35.0 | Cluster_14634 |
| 897.04000 | 27.950 | 28.390 | 35.0 | Cluster_39292 |
| 898.77000 | 22.550 | 22.980 | 35.0 | Cluster_14664 |
| 899.43000 | 19.930 | 20.330 | 35.0 | Cluster_39337 |
| 899.51000 | 1.420 | 1.800 | 35.0 | Cluster_39345 |
| 899.96000 | 22.500 | 22.910 | 35.0 | Cluster_04891 |
| 900.94000 | 14.600 | 15.030 | 35.0 | Cluster_39368 |
| 900.97000 | 19.860 | 20.300 | 35.0 | Cluster_39380 |
| 900.99000 | 1.400 | 1.920 | 35.0 | Cluster_39377 |
| 902.46000 | 17.480 | 17.700 | 35.0 | Cluster_39412 |
| 904.46000 | 31.430 | 31.860 | 35.0 | Cluster_39443 |
| 905.43000 | 11.850 | 12.310 | 35.0 | Cluster_39461 |
| 906.77000 | 31.320 | 31.760 | 35.0 | Cluster_14739 |
| 907.80000 | 19.890 | 20.410 | 35.0 | Cluster_14749 |
| 908.46000 | 19.880 | 20.220 | 35.0 | Cluster_39500 |
| 908.48000 | 25.100 | 25.280 | 35.0 | Cluster_14756 |
| 908.94000 | 25.120 | 25.750 | 35.0 | Cluster_04958 |
| 908.95000 | 31.420 | 31.720 | 35.0 | Cluster_39504 |
| 909.44000 | 20.020 | 20.410 | 35.0 | Cluster_14760 |
| 909.50000 | 31.530 | 31.860 | 35.0 | Cluster_14765 |
| 909.89000 | 17.400 | 17.650 | 35.0 | Cluster_39520 |
| 910.11000 | 25.230 | 26.320 | 35.0 | Cluster_14769 |
| 910.19000 | 27.860 | 28.480 | 35.0 | Cluster_04966 |
| 911.96000 | 19.920 | 20.160 | 35.0 | Cluster_39564 |
| 914.56000 | 27.860 | 28.530 | 35.0 | Cluster_39615 |
| 915.13000 | 22.590 | 23.080 | 35.0 | Cluster_14823 |
| 915.93000 | 27.880 | 28.130 | 35.0 | Cluster_39626 |
| 916.44000 | 17.290 | 17.670 | 35.0 | Cluster_39640 |
| 916.49000 | 19.900 | 20.670 | 35.0 | Cluster_39633 |
| 917.45000 | 25.240 | 25.410 | 35.0 | Cluster_14854 |
| 917.97000 | 31.670 | 31.940 | 35.0 | Cluster_39674 |
| 918.66000 | 31.620 | 32.200 | 35.0 | Cluster_39698 |
| 920.45000 | 19.920 | 20.220 | 35.0 | Cluster_39721 |
| 920.76000 | 25.120 | 25.460 | 35.0 | Cluster_14891 |
| 920.79000 | 1.550 | 1.670 | 35.0 | Cluster_39725 |
| 920.88000 | 17.500 | 18.010 | 35.0 | Cluster_39726 |
| 920.95000 | 22.500 | 22.690 | 35.0 | Cluster_39731 |
| 922.42000 | 22.630 | 22.920 | 35.0 | Cluster_14907 |
| 922.98000 | 25.090 | 25.510 | 35.0 | Cluster_39772 |
| 923.30000 | 28.040 | 28.540 | 35.0 | Cluster_14924 |
| 923.45000 | 25.290 | 25.740 | 35.0 | Cluster_14932 |
| 923.92000 | 7.790 | 8.320 | 35.0 | Cluster_39782 |
| 924.49000 | 28.040 | 28.700 | 35.0 | Cluster_39788 |
| 925.11000 | 25.090 | 25.380 | 35.0 | Cluster_14961 |

| | | | | |
|---|---|---|---|---|
| 926.42000 | 31.250 | 31.670 | 35.0 | Cluster_14975 |
| 927.48000 | 31.560 | 31.800 | 35.0 | Cluster_39834 |
| 928.47000 | 19.990 | 20.340 | 35.0 | Cluster_39858 |
| 928.96000 | 28.060 | 28.490 | 35.0 | Cluster_39861 |
| 929.12000 | 31.300 | 31.890 | 35.0 | Cluster_15009 |
| 929.43000 | 25.120 | 25.330 | 35.0 | Cluster_39880 |
| 929.90000 | 17.420 | 17.750 | 35.0 | Cluster_39897 |
| 931.73000 | 31.290 | 31.680 | 35.0 | Cluster_15043 |
| 931.90000 | 14.590 | 15.170 | 35.0 | Cluster_39924 |
| 932.64000 | 28.020 | 30.050 | 35.0 | Cluster_15062 |
| 932.95000 | 20.060 | 20.220 | 35.0 | Cluster_39941 |
| 933.14000 | 31.530 | 32.260 | 35.0 | Cluster_15072 |
| 933.33000 | 25.110 | 25.530 | 35.0 | Cluster_05038 |
| 934.00000 | 1.390 | 1.810 | 35.0 | Cluster_39963 |
| 934.13000 | 25.040 | 25.480 | 35.0 | Cluster_15084 |
| 934.43000 | 22.600 | 23.410 | 35.0 | Cluster_15083 |
| 934.78000 | 27.990 | 28.710 | 35.0 | Cluster_15089 |
| 935.78000 | 31.660 | 32.620 | 35.0 | Cluster_15100 |
| 936.45000 | 20.050 | 20.400 | 35.0 | Cluster_15116 |
| 936.80000 | 19.860 | 20.120 | 35.0 | Cluster_15129 |
| 937.00000 | 17.430 | 17.980 | 35.0 | Cluster_39997 |
| 937.44000 | 27.840 | 28.110 | 35.0 | Cluster_15144 |
| 937.49000 | 31.290 | 31.910 | 35.0 | Cluster_05051 |
| 938.07000 | 14.680 | 15.560 | 35.0 | Cluster_15154 |
| 938.13000 | 28.060 | 28.400 | 35.0 | Cluster_15156 |
| 938.21000 | 22.680 | 22.870 | 35.0 | Cluster_15160 |
| 939.69000 | 20.000 | 20.360 | 35.0 | Cluster_05058 |
| 940.46000 | 31.580 | 32.160 | 35.0 | Cluster_15180 |
| 941.72000 | 17.260 | 17.790 | 35.0 | Cluster_40098 |
| 942.00000 | 22.540 | 23.200 | 35.0 | Cluster_40094 |
| 943.45000 | 31.330 | 31.630 | 35.0 | Cluster_15229 |
| 945.13000 | 27.970 | 28.380 | 35.0 | Cluster_15244 |
| 945.39000 | 14.670 | 14.920 | 35.0 | Cluster_15242 |
| 945.72000 | 22.620 | 23.600 | 35.0 | Cluster_05075 |
| 945.73000 | 19.890 | 20.380 | 35.0 | Cluster_05078 |
| 945.78000 | 31.650 | 32.180 | 35.0 | Cluster_15250 |
| 945.98000 | 25.310 | 25.950 | 35.0 | Cluster_40152 |
| 947.16000 | 25.310 | 25.490 | 35.0 | Cluster_15264 |
| 947.22000 | 31.420 | 31.860 | 35.0 | Cluster_05084 |
| 947.79000 | 31.690 | 31.920 | 35.0 | Cluster_15273 |
| 948.19000 | 27.930 | 28.220 | 35.0 | Cluster_15277 |
| 948.52000 | 31.340 | 31.750 | 35.0 | Cluster_40180 |
| 949.14000 | 31.350 | 31.630 | 35.0 | Cluster_15301 |
| 949.44000 | 20.060 | 20.260 | 35.0 | Cluster_40190 |
| 949.46000 | 31.610 | 31.940 | 35.0 | Cluster_15302 |
| 950.44000 | 28.070 | 28.420 | 35.0 | Cluster_15314 |
| 951.17000 | 22.650 | 22.880 | 35.0 | Cluster_15324 |
| 951.96000 | 7.510 | 8.760 | 35.0 | Cluster_40227 |
| 952.14000 | 28.060 | 28.500 | 35.0 | Cluster_15337 |
| 952.15000 | 31.590 | 31.830 | 35.0 | Cluster_15339 |
| 952.44000 | 31.610 | 32.570 | 35.0 | Cluster_15342 |

| | | | | |
|---|---|---|---|---|
| 952.53000 | 1.380 | 1.840 | 35.0 | Cluster_40233 |
| 953.98000 | 19.900 | 20.100 | 35.0 | Cluster_40255 |
| 954.82000 | 25.130 | 25.390 | 35.0 | Cluster_15379 |
| 954.99000 | 19.910 | 20.370 | 35.0 | Cluster_40262 |
| 955.13000 | 17.260 | 17.690 | 35.0 | Cluster_15394 |
| 955.47000 | 25.300 | 25.650 | 35.0 | Cluster_15404 |
| 956.48000 | 22.670 | 22.990 | 35.0 | Cluster_15426 |
| 957.07000 | 11.840 | 12.000 | 35.0 | Cluster_40290 |
| 957.48000 | 22.550 | 22.750 | 35.0 | Cluster_40291 |
| 960.15000 | 27.830 | 28.270 | 35.0 | Cluster_15462 |
| 960.98000 | 25.150 | 25.570 | 35.0 | Cluster_40317 |
| 961.48000 | 25.120 | 25.340 | 35.0 | Cluster_15478 |
| 962.14000 | 22.620 | 22.830 | 35.0 | Cluster_15497 |
| 962.38000 | 17.250 | 17.410 | 35.0 | Cluster_40344 |
| 962.79000 | 25.120 | 25.790 | 35.0 | Cluster_15505 |
| 964.13000 | 25.130 | 25.570 | 35.0 | Cluster_15523 |
| 964.43000 | 17.440 | 17.700 | 35.0 | Cluster_40365 |
| 965.29000 | 1.640 | 1.850 | 35.0 | Cluster_40377 |
| 966.15000 | 27.940 | 28.380 | 35.0 | Cluster_15549 |
| 966.49000 | 17.440 | 17.820 | 35.0 | Cluster_40387 |
| 966.50000 | 31.570 | 32.070 | 35.0 | Cluster_05172 |
| 966.85000 | 25.130 | 25.470 | 35.0 | Cluster_15552 |
| 967.00000 | 31.630 | 32.160 | 35.0 | Cluster_40392 |
| 967.56000 | 22.680 | 23.030 | 35.0 | Cluster_15565 |
| 968.03000 | 17.400 | 17.670 | 35.0 | Cluster_40402 |
| 968.25000 | 25.220 | 25.360 | 35.0 | Cluster_05187 |
| 971.97000 | 19.900 | 20.000 | 35.0 | Cluster_40451 |
| 972.01000 | 19.890 | 20.440 | 35.0 | Cluster_40467 |
| 972.46000 | 17.480 | 17.700 | 35.0 | Cluster_15627 |
| 972.49000 | 22.660 | 22.870 | 35.0 | Cluster_15626 |
| 973.51000 | 22.670 | 23.170 | 35.0 | Cluster_40476 |
| 974.47000 | 28.050 | 28.200 | 35.0 | Cluster_15651 |
| 974.98000 | 17.380 | 17.710 | 35.0 | Cluster_40495 |
| 975.82000 | 17.380 | 17.850 | 35.0 | Cluster_15672 |
| 975.98000 | 31.650 | 32.000 | 35.0 | Cluster_05210 |
| 976.00000 | 22.650 | 23.000 | 35.0 | Cluster_05212 |
| 976.03000 | 25.160 | 25.450 | 35.0 | Cluster_40504 |
| 977.14000 | 27.990 | 28.200 | 35.0 | Cluster_15696 |
| 977.62000 | 31.460 | 31.650 | 35.0 | Cluster_40522 |
| 979.45000 | 22.690 | 23.050 | 35.0 | Cluster_40546 |
| 979.80000 | 28.060 | 28.190 | 35.0 | Cluster_15720 |
| 979.83000 | 31.360 | 32.110 | 35.0 | Cluster_15721 |
| 979.98000 | 7.510 | 7.660 | 35.0 | Cluster_40568 |
| 980.00000 | 17.300 | 17.710 | 35.0 | Cluster_40562 |
| 980.50000 | 19.910 | 20.120 | 35.0 | Cluster_40565 |
| 980.70000 | 14.830 | 15.050 | 35.0 | Cluster_40578 |
| 981.45000 | 28.050 | 28.230 | 35.0 | Cluster_15750 |
| 983.25000 | 20.060 | 20.280 | 35.0 | Cluster_05237 |
| 983.95000 | 14.870 | 15.210 | 35.0 | Cluster_40617 |
| 985.14000 | 22.650 | 22.870 | 35.0 | Cluster_40628 |
| 986.02000 | 25.190 | 25.490 | 35.0 | Cluster_40630 |

| | | | | |
|---|---|---|---|---|
| 986.52000 | 1.390 | 1.800 | 35.0 | Cluster_40639 |
| 986.83000 | 25.160 | 25.490 | 35.0 | Cluster_15810 |
| 987.24000 | 19.890 | 20.200 | 35.0 | Cluster_40654 |
| 987.80000 | 22.650 | 23.040 | 35.0 | Cluster_15825 |
| 988.44000 | 17.480 | 17.670 | 35.0 | Cluster_15835 |
| 991.93000 | 14.670 | 15.150 | 35.0 | Cluster_40717 |
| 993.50000 | 22.650 | 22.790 | 35.0 | Cluster_15895 |
| 997.19000 | 31.660 | 31.950 | 35.0 | Cluster_15954 |
| 1000.06000 | 25.320 | 25.610 | 35.0 | Cluster_40851 |
| 1001.03000 | 28.010 | 28.330 | 35.0 | Cluster_40878 |
| 1001.16000 | 27.820 | 28.010 | 35.0 | Cluster_15996 |
| 1001.49000 | 22.660 | 22.960 | 35.0 | Cluster_40884 |
| 1003.52000 | 11.730 | 11.910 | 35.0 | Cluster_05294 |
| 1004.00000 | 20.000 | 20.330 | 35.0 | Cluster_40937 |
| 1004.31000 | 22.610 | 22.830 | 35.0 | Cluster_05296 |
| 1004.57000 | 1.420 | 1.800 | 35.0 | Cluster_40940 |
| 1007.81000 | 31.270 | 31.580 | 35.0 | Cluster_16072 |
| 1008.78000 | 27.810 | 28.060 | 35.0 | Cluster_16080 |
| 1012.26000 | 27.990 | 29.000 | 35.0 | Cluster_05329 |
| 1012.27000 | 27.980 | 28.720 | 35.0 | Cluster_41044 |
| 1012.93000 | 22.490 | 22.610 | 35.0 | Cluster_41054 |
| 1013.01000 | 27.790 | 28.710 | 35.0 | Cluster_05341 |
| 1013.47000 | 31.580 | 31.950 | 35.0 | Cluster_16138 |
| 1013.49000 | 31.460 | 31.600 | 35.0 | Cluster_16137 |
| 1013.76000 | 25.100 | 25.600 | 35.0 | Cluster_05344 |
| 1013.90000 | 27.810 | 27.940 | 35.0 | Cluster_41056 |
| 1015.92000 | 31.650 | 31.810 | 35.0 | Cluster_41088 |
| 1016.78000 | 11.730 | 12.300 | 35.0 | Cluster_05351 |
| 1017.27000 | 31.660 | 31.940 | 35.0 | Cluster_41099 |
| 1019.25000 | 25.060 | 25.510 | 35.0 | Cluster_05360 |
| 1020.15000 | 19.920 | 20.310 | 35.0 | Cluster_16210 |
| 1022.50000 | 31.700 | 32.310 | 35.0 | Cluster_16231 |
| 1023.82000 | 1.650 | 1.960 | 35.0 | Cluster_41185 |
| 1024.48000 | 17.530 | 17.820 | 35.0 | Cluster_16254 |
| 1024.50000 | 22.590 | 22.900 | 35.0 | Cluster_16256 |
| 1024.93000 | 17.520 | 17.710 | 35.0 | Cluster_41195 |
| 1025.46000 | 19.880 | 20.050 | 35.0 | Cluster_16264 |
| 1025.81000 | 31.690 | 32.100 | 35.0 | Cluster_41211 |
| 1026.83000 | 31.710 | 32.080 | 35.0 | Cluster_16274 |
| 1027.13000 | 19.980 | 20.230 | 35.0 | Cluster_16285 |
| 1027.14000 | 19.940 | 20.060 | 35.0 | Cluster_16284 |
| 1027.88000 | 28.020 | 28.430 | 35.0 | Cluster_16291 |
| 1028.18000 | 31.580 | 32.300 | 35.0 | Cluster_16295 |
| 1028.84000 | 22.590 | 23.480 | 35.0 | Cluster_16300 |
| 1029.99000 | 25.120 | 25.250 | 35.0 | Cluster_41262 |
| 1031.20000 | 31.260 | 31.520 | 35.0 | Cluster_16323 |
| 1032.82000 | 25.220 | 25.760 | 35.0 | Cluster_16335 |
| 1033.20000 | 31.260 | 31.480 | 35.0 | Cluster_16342 |
| 1036.52000 | 22.500 | 23.060 | 35.0 | Cluster_16382 |
| 1037.50000 | 17.460 | 17.740 | 35.0 | Cluster_16400 |
| 1039.05000 | 1.390 | 1.810 | 35.0 | Cluster_41371 |

| | | | | |
|---|---|---|---|---|
| 1039.52000 | 27.970 | 28.620 | 35.0 | Cluster_41374 |
| 1039.81000 | 27.920 | 28.190 | 35.0 | Cluster_16429 |
| 1040.83000 | 25.120 | 25.310 | 35.0 | Cluster_05437 |
| 1041.01000 | 28.030 | 28.270 | 35.0 | Cluster_41401 |
| 1042.17000 | 31.590 | 31.910 | 35.0 | Cluster_16457 |
| 1044.49000 | 27.920 | 28.280 | 35.0 | Cluster_41432 |
| 1046.13000 | 25.260 | 25.540 | 35.0 | Cluster_16488 |
| 1046.22000 | 17.430 | 17.580 | 35.0 | Cluster_16495 |
| 1047.16000 | 19.910 | 20.550 | 35.0 | Cluster_16507 |
| 1047.83000 | 27.840 | 28.260 | 35.0 | Cluster_41470 |
| 1053.50000 | 31.540 | 32.190 | 35.0 | Cluster_16581 |
| 1053.98000 | 28.040 | 28.260 | 35.0 | Cluster_41534 |
| 1054.99000 | 7.750 | 8.370 | 35.0 | Cluster_05476 |
| 1055.22000 | 31.420 | 31.670 | 35.0 | Cluster_16595 |
| 1055.50000 | 31.640 | 31.900 | 35.0 | Cluster_16606 |
| 1056.52000 | 22.510 | 22.750 | 35.0 | Cluster_16616 |
| 1057.06000 | 31.490 | 32.650 | 35.0 | Cluster_41567 |
| 1057.33000 | 25.040 | 25.360 | 35.0 | Cluster_05486 |
| 1057.34000 | 25.310 | 25.570 | 35.0 | Cluster_05487 |
| 1057.58000 | 1.390 | 1.800 | 35.0 | Cluster_41574 |
| 1057.99000 | 11.640 | 11.940 | 35.0 | Cluster_41578 |
| 1058.02000 | 27.910 | 28.270 | 35.0 | Cluster_41582 |
| 1058.81000 | 31.600 | 31.910 | 35.0 | Cluster_16658 |
| 1058.85000 | 22.580 | 22.790 | 35.0 | Cluster_16649 |
| 1058.96000 | 27.840 | 27.920 | 35.0 | Cluster_41595 |
| 1058.97000 | 28.040 | 28.300 | 35.0 | Cluster_41597 |
| 1059.01000 | 27.920 | 28.050 | 35.0 | Cluster_41596 |
| 1060.32000 | 1.650 | 1.860 | 35.0 | Cluster_41612 |
| 1060.75000 | 7.790 | 8.260 | 35.0 | Cluster_05502 |
| 1060.82000 | 31.630 | 31.910 | 35.0 | Cluster_16679 |
| 1061.26000 | 27.800 | 28.300 | 35.0 | Cluster_05510 |
| 1061.33000 | 25.060 | 25.320 | 35.0 | Cluster_05506 |
| 1061.82000 | 1.640 | 1.910 | 35.0 | Cluster_41625 |
| 1062.83000 | 31.620 | 31.910 | 35.0 | Cluster_16700 |
| 1063.27000 | 27.790 | 28.260 | 35.0 | Cluster_05516 |
| 1063.82000 | 1.630 | 1.910 | 35.0 | Cluster_41655 |
| 1063.99000 | 17.430 | 17.700 | 35.0 | Cluster_41663 |
| 1064.72000 | 7.500 | 7.790 | 35.0 | Cluster_05521 |
| 1064.73000 | 7.770 | 8.180 | 35.0 | Cluster_41675 |
| 1064.99000 | 17.450 | 17.700 | 35.0 | Cluster_41673 |
| 1066.26000 | 27.830 | 28.220 | 35.0 | Cluster_41702 |
| 1067.52000 | 17.470 | 17.670 | 35.0 | Cluster_41715 |
| 1068.05000 | 31.450 | 32.110 | 35.0 | Cluster_41719 |
| 1068.75000 | 25.120 | 25.410 | 35.0 | Cluster_41734 |
| 1069.51000 | 27.810 | 28.180 | 35.0 | Cluster_41748 |
| 1070.16000 | 31.560 | 31.910 | 35.0 | Cluster_16781 |
| 1070.51000 | 27.810 | 28.230 | 35.0 | Cluster_05546 |
| 1072.55000 | 31.660 | 32.210 | 35.0 | Cluster_16804 |
| 1072.76000 | 27.820 | 28.190 | 35.0 | Cluster_05555 |
| 1074.19000 | 25.140 | 25.380 | 35.0 | Cluster_16831 |
| 1074.51000 | 27.900 | 28.260 | 35.0 | Cluster_41810 |

| | | | | |
|---|---|---|---|---|
| 1075.53000 | 31.540 | 31.840 | 35.0 | Cluster_41816 |
| 1076.75000 | 27.800 | 28.170 | 35.0 | Cluster_05574 |
| 1078.01000 | 27.800 | 28.160 | 35.0 | Cluster_05578 |
| 1079.49000 | 27.900 | 28.160 | 35.0 | Cluster_05582 |
| 1080.75000 | 19.910 | 20.820 | 35.0 | Cluster_00593 |
| 1082.00000 | 27.820 | 28.190 | 35.0 | Cluster_05588 |
| 1083.50000 | 27.920 | 28.110 | 35.0 | Cluster_05592 |
| 1086.59000 | 27.920 | 28.290 | 35.0 | Cluster_41940 |
| 1091.58000 | 1.390 | 1.800 | 35.0 | Cluster_41998 |
| 1092.90000 | 31.350 | 32.070 | 35.0 | Cluster_17025 |
| 1093.21000 | 31.350 | 32.010 | 35.0 | Cluster_17031 |
| 1093.55000 | 31.340 | 32.060 | 35.0 | Cluster_17035 |
| 1093.58000 | 19.960 | 20.180 | 35.0 | Cluster_42024 |
| 1093.99000 | 20.020 | 20.190 | 35.0 | Cluster_42029 |
| 1094.92000 | 22.480 | 22.800 | 35.0 | Cluster_00625 |
| 1096.55000 | 22.660 | 23.350 | 35.0 | Cluster_42051 |
| 1100.23000 | 31.350 | 31.820 | 35.0 | Cluster_17091 |
| 1100.56000 | 31.360 | 31.640 | 35.0 | Cluster_17097 |
| 1101.96000 | 31.560 | 32.670 | 35.0 | Cluster_17105 |
| 1102.54000 | 27.940 | 28.260 | 35.0 | Cluster_17111 |
| 1103.12000 | 22.610 | 22.840 | 35.0 | Cluster_42104 |
| 1108.23000 | 27.990 | 28.440 | 35.0 | Cluster_17176 |
| 1108.37000 | 27.990 | 28.540 | 35.0 | Cluster_42148 |
| 1108.88000 | 27.940 | 28.220 | 35.0 | Cluster_17175 |
| 1109.86000 | 28.030 | 28.260 | 35.0 | Cluster_17199 |
| 1113.50000 | 19.920 | 20.940 | 35.0 | Cluster_42184 |
| 1113.53000 | 25.230 | 25.610 | 35.0 | Cluster_17241 |
| 1115.18000 | 20.010 | 20.940 | 35.0 | Cluster_17258 |
| 1115.77000 | 28.030 | 28.340 | 35.0 | Cluster_00660 |
| 1117.22000 | 31.210 | 31.520 | 35.0 | Cluster_17276 |
| 1118.21000 | 25.320 | 25.650 | 35.0 | Cluster_17277 |
| 1118.22000 | 27.860 | 28.580 | 35.0 | Cluster_17280 |
| 1119.87000 | 1.650 | 1.840 | 35.0 | Cluster_42234 |
| 1121.54000 | 25.110 | 25.550 | 35.0 | Cluster_17322 |
| 1123.04000 | 25.040 | 25.380 | 35.0 | Cluster_42270 |
| 1123.97000 | 19.950 | 20.110 | 35.0 | Cluster_42271 |
| 1124.25000 | 31.510 | 32.350 | 35.0 | Cluster_17359 |
| 1125.57000 | 1.430 | 1.760 | 35.0 | Cluster_42283 |
| 1126.07000 | 25.070 | 26.000 | 35.0 | Cluster_42291 |
| 1127.88000 | 31.490 | 31.930 | 35.0 | Cluster_17398 |
| 1128.57000 | 27.990 | 28.730 | 35.0 | Cluster_17409 |
| 1129.21000 | 31.220 | 31.570 | 35.0 | Cluster_17418 |
| 1129.55000 | 31.540 | 31.870 | 35.0 | Cluster_17428 |
| 1129.90000 | 31.220 | 31.560 | 35.0 | Cluster_17433 |
| 1130.23000 | 27.880 | 28.550 | 35.0 | Cluster_17434 |
| 1131.58000 | 31.370 | 32.260 | 35.0 | Cluster_17448 |
| 1133.13000 | 17.390 | 17.710 | 35.0 | Cluster_42337 |
| 1133.20000 | 31.230 | 31.590 | 35.0 | Cluster_17479 |
| 1133.22000 | 31.550 | 31.950 | 35.0 | Cluster_17484 |
| 1134.90000 | 31.510 | 32.360 | 35.0 | Cluster_17503 |
| 1135.23000 | 31.430 | 31.940 | 35.0 | Cluster_17509 |

| | | | | |
|---|---|---|---|---|
| 1135.51000 | 22.610 | 22.830 | 35.0 | Cluster_17513 |
| 1135.54000 | 31.210 | 31.470 | 35.0 | Cluster_17519 |
| 1135.55000 | 27.920 | 28.160 | 35.0 | Cluster_17517 |
| 1136.11000 | 25.210 | 25.510 | 35.0 | Cluster_42363 |
| 1136.88000 | 22.560 | 23.520 | 35.0 | Cluster_17525 |
| 1136.89000 | 31.550 | 32.160 | 35.0 | Cluster_17538 |
| 1138.13000 | 31.240 | 31.470 | 35.0 | Cluster_42377 |
| 1140.22000 | 27.830 | 28.300 | 35.0 | Cluster_17567 |
| 1143.18000 | 22.670 | 23.020 | 35.0 | Cluster_00674 |
| 1143.56000 | 27.910 | 28.730 | 35.0 | Cluster_17607 |
| 1143.60000 | 1.410 | 1.860 | 35.0 | Cluster_42409 |
| 1144.20000 | 22.650 | 23.260 | 35.0 | Cluster_17626 |
| 1144.24000 | 31.500 | 32.060 | 35.0 | Cluster_17620 |
| 1145.21000 | 27.830 | 28.220 | 35.0 | Cluster_17645 |
| 1146.57000 | 22.680 | 23.030 | 35.0 | Cluster_00680 |
| 1149.29000 | 31.270 | 31.440 | 35.0 | Cluster_42461 |
| 1151.05000 | 27.960 | 28.350 | 35.0 | Cluster_42474 |
| 1152.54000 | 28.010 | 28.390 | 35.0 | Cluster_17735 |
| 1152.88000 | 28.010 | 28.180 | 35.0 | Cluster_17750 |
| 1153.03000 | 22.500 | 22.700 | 35.0 | Cluster_05790 |
| 1153.76000 | 19.970 | 20.360 | 35.0 | Cluster_00695 |
| 1154.35000 | 1.610 | 1.810 | 35.0 | Cluster_42485 |
| 1155.34000 | 1.640 | 1.840 | 35.0 | Cluster_42501 |
| 1156.84000 | 1.600 | 1.850 | 35.0 | Cluster_42511 |
| 1158.84000 | 1.650 | 1.860 | 35.0 | Cluster_42519 |
| 1159.61000 | 27.920 | 28.320 | 35.0 | Cluster_42529 |
| 1160.34000 | 1.650 | 1.840 | 35.0 | Cluster_42534 |
| 1162.11000 | 20.010 | 20.380 | 35.0 | Cluster_42551 |
| 1162.64000 | 1.410 | 1.800 | 35.0 | Cluster_42555 |
| 1163.09000 | 25.260 | 26.480 | 35.0 | Cluster_05817 |
| 1166.08000 | 25.300 | 25.730 | 35.0 | Cluster_05827 |
| 1166.14000 | 25.310 | 25.480 | 35.0 | Cluster_42592 |
| 1166.21000 | 20.060 | 20.770 | 35.0 | Cluster_17894 |
| 1167.19000 | 25.060 | 25.610 | 35.0 | Cluster_17908 |
| 1167.57000 | 28.030 | 28.490 | 35.0 | Cluster_17921 |
| 1168.21000 | 17.500 | 17.810 | 35.0 | Cluster_17931 |
| 1168.33000 | 25.190 | 25.800 | 35.0 | Cluster_42608 |
| 1168.58000 | 25.190 | 25.720 | 35.0 | Cluster_05832 |
| 1173.07000 | 27.970 | 28.150 | 35.0 | Cluster_42645 |
| 1180.80000 | 27.990 | 28.420 | 35.0 | Cluster_05888 |
| 1183.23000 | 27.940 | 28.290 | 35.0 | Cluster_18040 |
| 1187.59000 | 31.420 | 31.800 | 35.0 | Cluster_18075 |
| 1190.56000 | 27.950 | 28.230 | 35.0 | Cluster_18091 |
| 1192.81000 | 27.930 | 28.170 | 35.0 | Cluster_42765 |
| 1192.92000 | 25.320 | 26.100 | 35.0 | Cluster_18117 |
| 1195.88000 | 27.950 | 28.200 | 35.0 | Cluster_18133 |
| 1196.63000 | 1.390 | 1.800 | 35.0 | Cluster_42787 |
| 1197.59000 | 25.080 | 25.680 | 35.0 | Cluster_18145 |
| 1197.90000 | 27.950 | 28.180 | 35.0 | Cluster_18140 |
| 1198.64000 | 22.610 | 22.860 | 35.0 | Cluster_42803 |
| 1199.32000 | 22.530 | 22.720 | 35.0 | Cluster_05982 |

| | | | | |
|---|---|---|---|---|
| 1199.57000 | 31.670 | 32.590 | 35.0 | Cluster_18161 |
| 1200.26000 | 25.090 | 25.780 | 35.0 | Cluster_18168 |
| 1200.56000 | 25.320 | 25.570 | 35.0 | Cluster_18177 |
| 1201.89000 | 11.730 | 11.870 | 35.0 | Cluster_18192 |
| 1203.44000 | 1.500 | 1.680 | 35.0 | Cluster_42830 |
| 1206.62000 | 22.520 | 22.840 | 35.0 | Cluster_42853 |
| 1208.56000 | 31.640 | 32.560 | 35.0 | Cluster_18237 |
| 1209.51000 | 19.980 | 20.450 | 35.0 | Cluster_42887 |
| 1213.12000 | 14.580 | 14.970 | 35.0 | Cluster_42910 |
| 1215.15000 | 1.390 | 1.800 | 35.0 | Cluster_42924 |
| 1218.11000 | 27.880 | 28.380 | 35.0 | Cluster_42950 |
| 1219.09000 | 31.570 | 31.720 | 35.0 | Cluster_42956 |
| 1221.25000 | 19.960 | 20.370 | 35.0 | Cluster_18299 |
| 1222.10000 | 19.970 | 20.690 | 35.0 | Cluster_42968 |
| 1224.08000 | 25.090 | 25.700 | 35.0 | Cluster_42977 |
| 1224.60000 | 25.150 | 25.340 | 35.0 | Cluster_42983 |
| 1225.59000 | 25.280 | 25.630 | 35.0 | Cluster_42994 |
| 1228.07000 | 25.130 | 25.960 | 35.0 | Cluster_43008 |
| 1228.11000 | 22.540 | 22.780 | 35.0 | Cluster_43007 |
| 1230.63000 | 1.450 | 1.740 | 35.0 | Cluster_43026 |
| 1232.57000 | 25.160 | 25.320 | 35.0 | Cluster_43045 |
| 1234.58000 | 25.160 | 25.330 | 35.0 | Cluster_43065 |
| 1238.57000 | 25.240 | 25.570 | 35.0 | Cluster_43079 |
| 1245.28000 | 19.970 | 20.630 | 35.0 | Cluster_18390 |
| 1249.16000 | 1.380 | 1.790 | 35.0 | Cluster_43135 |
| 1251.87000 | 1.650 | 1.850 | 35.0 | Cluster_43149 |
| 1253.87000 | 1.650 | 1.850 | 35.0 | Cluster_43162 |
| 1255.86000 | 1.600 | 1.910 | 35.0 | Cluster_43176 |
| 1257.08000 | 22.640 | 22.820 | 35.0 | Cluster_06282 |
| 1264.27000 | 17.430 | 18.580 | 35.0 | Cluster_18463 |
| 1267.20000 | 1.430 | 1.790 | 35.0 | Cluster_43224 |
| 1267.68000 | 1.420 | 1.800 | 35.0 | Cluster_43228 |
| 1274.29000 | 31.230 | 31.620 | 35.0 | Cluster_18502 |
| 1284.15000 | 22.660 | 23.000 | 35.0 | Cluster_43287 |
| 1300.69000 | 31.520 | 31.710 | 35.0 | Cluster_43366 |
| 1300.70000 | 31.390 | 31.530 | 35.0 | Cluster_43360 |
| 1300.73000 | 31.690 | 31.890 | 35.0 | Cluster_43361 |
| 1300.89000 | 1.430 | 1.820 | 35.0 | Cluster_43362 |
| 1301.68000 | 1.410 | 1.690 | 35.0 | Cluster_43365 |
| 1320.21000 | 1.390 | 1.780 | 35.0 | Cluster_43449 |
| 1329.73000 | 25.160 | 25.710 | 35.0 | Cluster_43484 |
| 1330.45000 | 27.860 | 28.160 | 35.0 | Cluster_06499 |
| 1331.69000 | 25.220 | 25.950 | 35.0 | Cluster_43492 |
| 1334.68000 | 31.300 | 31.730 | 35.0 | Cluster_43505 |
| 1349.89000 | 1.650 | 1.840 | 35.0 | Cluster_43562 |
| 1353.71000 | 25.040 | 25.230 | 35.0 | Cluster_43578 |
| 1354.21000 | 25.210 | 25.480 | 35.0 | Cluster_43579 |
| 1354.66000 | 27.960 | 28.180 | 35.0 | Cluster_06571 |
| 1361.03000 | 27.810 | 27.940 | 35.0 | Cluster_18681 |
| 1367.16000 | 31.300 | 31.700 | 35.0 | Cluster_43645 |
| 1372.75000 | 1.410 | 1.790 | 35.0 | Cluster_43667 |

| | | | | |
|---|---|---|---|---|
| 1376.18000 | 19.930 | 20.320 | 35.0 | Cluster_43680 |
| 1376.34000 | 27.820 | 27.970 | 35.0 | Cluster_18707 |
| 1383.67000 | 28.020 | 28.220 | 35.0 | Cluster_18716 |
| 1394.65000 | 27.850 | 28.080 | 35.0 | Cluster_43739 |
| 1396.44000 | 28.030 | 28.270 | 35.0 | Cluster_18732 |
| 1402.12000 | 25.310 | 25.610 | 35.0 | Cluster_18741 |
| 1406.24000 | 1.420 | 1.740 | 35.0 | Cluster_43793 |
| 1407.03000 | 27.830 | 28.260 | 35.0 | Cluster_18752 |
| 1414.35000 | 27.850 | 28.190 | 35.0 | Cluster_18768 |
| 1417.37000 | 27.810 | 28.230 | 35.0 | Cluster_18772 |
| 1420.01000 | 27.880 | 28.160 | 35.0 | Cluster_18783 |
| 1422.96000 | 22.600 | 23.030 | 35.0 | Cluster_43843 |
| 1423.47000 | 22.650 | 23.080 | 35.0 | Cluster_43844 |
| 1424.69000 | 27.830 | 28.170 | 35.0 | Cluster_18789 |
| 1424.79000 | 1.450 | 1.800 | 35.0 | Cluster_43851 |
| 1425.27000 | 1.420 | 1.800 | 35.0 | Cluster_43853 |
| 1427.03000 | 27.920 | 28.150 | 35.0 | Cluster_18794 |
| 1428.82000 | 22.560 | 22.720 | 35.0 | Cluster_43862 |
| 1429.02000 | 27.940 | 28.120 | 35.0 | Cluster_18795 |
| 1430.01000 | 27.860 | 28.130 | 35.0 | Cluster_18796 |
| 1431.68000 | 27.860 | 28.120 | 35.0 | Cluster_18799 |
| 1437.03000 | 27.910 | 28.120 | 35.0 | Cluster_18803 |
| 1439.35000 | 27.920 | 28.100 | 35.0 | Cluster_18807 |
| 1459.27000 | 1.420 | 1.790 | 35.0 | Cluster_43959 |
| 1477.30000 | 1.420 | 1.800 | 35.0 | Cluster_44015 |
| 1489.22000 | 31.260 | 31.580 | 35.0 | Cluster_44043 |
| 1492.94000 | 1.440 | 1.810 | 35.0 | Cluster_44054 |
| 1495.14000 | 1.520 | 1.720 | 35.0 | Cluster_44065 |
| 1511.80000 | 1.420 | 1.800 | 35.0 | Cluster_44102 |
| 1518.72000 | 19.930 | 20.060 | 35.0 | Cluster_44112 |
| 1530.33000 | 1.420 | 1.800 | 35.0 | Cluster_44125 |
| 1560.76000 | 31.560 | 31.910 | 35.0 | Cluster_44175 |
| 1564.32000 | 1.420 | 1.780 | 35.0 | Cluster_44184 |
| 1571.76000 | 31.610 | 31.910 | 35.0 | Cluster_44194 |
| 1582.85000 | 1.420 | 1.730 | 35.0 | Cluster_44218 |
| 1616.85000 | 1.440 | 1.780 | 35.0 | Cluster_44250 |
| 1633.97000 | 1.560 | 1.810 | 35.0 | Cluster_44257 |
| 1635.40000 | 1.430 | 1.720 | 35.0 | Cluster_44260 |
| 1676.82000 | 27.900 | 28.270 | 35.0 | Cluster_44320 |
| 1683.84000 | 27.900 | 28.150 | 35.0 | Cluster_44341 |
| 1687.92000 | 1.440 | 1.800 | 35.0 | Cluster_44353 |
| 1739.94000 | 1.490 | 1.800 | 35.0 | Cluster_44403 |
| 1792.99000 | 1.520 | 1.790 | 35.0 | Cluster_44425 |

(-) Global MS/MS Masses:
(none)
Custom Data Dependent Settings:
Not enabled

---

Pass 8 (wheat-mixed-digests_MS2_TO-USE-inclusion-01_1.raw):

Creator: Orbi_30393

Last modified: 10/19/2021 by Orbi_30393
MS Run Time (min): 43.00
Sequence override of method parameters not enabled.
Divert Valve: not used during run
Contact Closure: not used during run
Syringe Pump: not used during run
MS Detector Settings:
Real-time modifications to method not enabled
Stepped collision energy not enabled
Additional Microscans:

| | | |
|---|---|---|
| MS2 | 0 | 0 |
| MS3 | 0 | 0 |
| MS4 | 0 | 0 |
| MS5 | 0 | 0 |
| MS6 | 0 | 0 |
| MS7 | 0 | 0 |
| MS8 | 0 | 0 |
| MS9 | 0 | 0 |
| MS10 | 0 | 0 |

Segment 1 Information
Duration (min): 43.00
Number of Scan Events: 2
Tune Method: Orbitrap-tune-file_2020-03-13_HESI
Scan Event Details:
1: FTMS + p norm res=15000 o(300.0-2000.0)
CV = 0.0V
2: ITMS + c norm Dep MS/MS Most intense ion from (1)
Activation Type: CID
Min. Signal Required: 500.0
Isolation Width: 2.00
Normalized Coll. Energy: 35.0
Default Charge State: 2
Activation Q: 0.250
Activation Time: 10.000
CV = 0.0V
Lock Masses:
Pos List Name: N/A
Source: API Source
Mass List: (none)
Neg List Name: N/A
Source: API Source
Mass List: (none)
Data Dependent Settings:
Use separate polarity settings disabled
Parent Mass List:

| MS Mass | MS FAIMS CV | MS Normalized Collision Energy | MS2 Mass | MS2 Normalized Collision Energy | Name |
|---|---|---|---|---|---|
| 300.13000 | | 35.0 | | | Cluster_06844 |
| 300.14000 | | 35.0 | | | Cluster_18928 |

| | | |
|---|---|---|
| 300.71000 | 35.0 | Cluster_18955 |
| 301.50000 | 35.0 | Cluster_06851 |
| 301.65000 | 35.0 | Cluster_18984 |
| 302.64000 | 35.0 | Cluster_19016 |
| 302.65000 | 35.0 | Cluster_19017 |
| 303.99000 | 35.0 | Cluster_19040 |
| 304.11000 | 35.0 | Cluster_19042 |
| 304.14000 | 35.0 | Cluster_19049 |
| 304.16000 | 35.0 | Cluster_19050 |
| 304.20000 | 35.0 | Cluster_19052 |
| 304.49000 | 35.0 | Cluster_06863 |
| 304.64000 | 35.0 | Cluster_19062 |
| 306.13000 | 35.0 | Cluster_06873 |
| 306.67000 | 35.0 | Cluster_19109 |
| 307.12000 | 35.0 | Cluster_19119 |
| 307.13000 | 35.0 | Cluster_06880 |
| 308.65000 | 35.0 | Cluster_19177 |
| 308.66000 | 35.0 | Cluster_19179 |
| 309.53000 | 35.0 | Cluster_06901 |
| 310.58000 | 35.0 | Cluster_19217 |
| 310.65000 | 35.0 | Cluster_19222 |
| 310.66000 | 35.0 | Cluster_19221 |
| 311.17000 | 35.0 | Cluster_19246 |
| 311.68000 | 35.0 | Cluster_19257 |
| 313.65000 | 35.0 | Cluster_19315 |
| 314.82000 | 35.0 | Cluster_06941 |
| 315.65000 | 35.0 | Cluster_19384 |
| 316.69000 | 35.0 | Cluster_19442 |
| 317.06000 | 35.0 | Cluster_19459 |
| 317.16000 | 35.0 | Cluster_19466 |
| 317.64000 | 35.0 | Cluster_19490 |
| 318.64000 | 35.0 | Cluster_19532 |
| 318.85000 | 35.0 | Cluster_06973 |
| 319.17000 | 35.0 | Cluster_19552 |
| 319.67000 | 35.0 | Cluster_19560 |
| 319.94000 | 35.0 | Cluster_00850 |
| 320.49000 | 35.0 | Cluster_06993 |
| 320.80000 | 35.0 | Cluster_06994 |
| 321.15000 | 35.0 | Cluster_06995 |
| 321.17000 | 35.0 | Cluster_00869 |
| 321.52000 | 35.0 | Cluster_19594 |
| 322.15000 | 35.0 | Cluster_19619 |
| 322.17000 | 35.0 | Cluster_19617 |
| 322.50000 | 35.0 | Cluster_06998 |
| 323.65000 | 35.0 | Cluster_19672 |
| 323.69000 | 35.0 | Cluster_19677 |
| 323.83000 | 35.0 | Cluster_19683 |
| 324.42000 | 35.0 | Cluster_00878 |
| 324.60000 | 35.0 | Cluster_19711 |
| 324.65000 | 35.0 | Cluster_19713 |
| 324.83000 | 35.0 | Cluster_07017 |

| | | |
|---|---|---|
| 325.14000 | 35.0 | Cluster_19730 |
| 325.82000 | 35.0 | Cluster_07023 |
| 325.85000 | 35.0 | Cluster_07028 |
| 326.17000 | 35.0 | Cluster_19753 |
| 326.40000 | 35.0 | Cluster_00887 |
| 326.90000 | 35.0 | Cluster_00888 |
| 327.21000 | 35.0 | Cluster_19791 |
| 327.96000 | 35.0 | Cluster_07043 |
| 328.48000 | 35.0 | Cluster_07051 |
| 329.18000 | 35.0 | Cluster_19855 |
| 329.82000 | 35.0 | Cluster_07063 |
| 329.93000 | 35.0 | Cluster_19873 |
| 330.18000 | 35.0 | Cluster_19886 |
| 330.90000 | 35.0 | Cluster_19916 |
| 331.17000 | 35.0 | Cluster_19926 |
| 331.20000 | 35.0 | Cluster_19928 |
| 331.21000 | 35.0 | Cluster_19930 |
| 331.22000 | 35.0 | Cluster_19925 |
| 331.66000 | 35.0 | Cluster_19936 |
| 332.14000 | 35.0 | Cluster_19948 |
| 333.47000 | 35.0 | Cluster_07081 |
| 333.82000 | 35.0 | Cluster_07087 |
| 333.87000 | 35.0 | Cluster_07090 |
| 335.12000 | 35.0 | Cluster_20034 |
| 335.83000 | 35.0 | Cluster_07104 |
| 336.19000 | 35.0 | Cluster_07109 |
| 336.23000 | 35.0 | Cluster_20093 |
| 336.91000 | 35.0 | Cluster_00920 |
| 337.19000 | 35.0 | Cluster_20132 |
| 337.21000 | 35.0 | Cluster_20126 |
| 337.83000 | 35.0 | Cluster_07126 |
| 338.11000 | 35.0 | Cluster_20150 |
| 338.67000 | 35.0 | Cluster_20172 |
| 339.16000 | 35.0 | Cluster_20192 |
| 339.47000 | 35.0 | Cluster_07139 |
| 339.81000 | 35.0 | Cluster_07150 |
| 339.94000 | 35.0 | Cluster_00929 |
| 340.18000 | 35.0 | Cluster_20223 |
| 340.26000 | 35.0 | Cluster_20229 |
| 341.15000 | 35.0 | Cluster_20257 |
| 341.16000 | 35.0 | Cluster_20261 |
| 342.16000 | 35.0 | Cluster_07167 |
| 343.11000 | 35.0 | Cluster_20316 |
| 343.18000 | 35.0 | Cluster_20318 |
| 343.66000 | 35.0 | Cluster_20344 |
| 345.19000 | 35.0 | Cluster_20390 |
| 345.67000 | 35.0 | Cluster_20402 |
| 346.15000 | 35.0 | Cluster_07199 |
| 346.67000 | 35.0 | Cluster_20444 |
| 347.20000 | 35.0 | Cluster_20457 |
| 347.71000 | 35.0 | Cluster_20471 |

| | | |
|---|---|---|
| 348.44000 | 35.0 | Cluster_20495 |
| 348.87000 | 35.0 | Cluster_07234 |
| 348.92000 | 35.0 | Cluster_00975 |
| 348.93000 | 35.0 | Cluster_00976 |
| 349.20000 | 35.0 | Cluster_20517 |
| 349.21000 | 35.0 | Cluster_20520 |
| 349.64000 | 35.0 | Cluster_20543 |
| 349.73000 | 35.0 | Cluster_20549 |
| 350.42000 | 35.0 | Cluster_00982 |
| 351.18000 | 35.0 | Cluster_20611 |
| 351.19000 | 35.0 | Cluster_20622 |
| 352.66000 | 35.0 | Cluster_20669 |
| 353.17000 | 35.0 | Cluster_01002 |
| 353.21000 | 35.0 | Cluster_20705 |
| 353.66000 | 35.0 | Cluster_01003 |
| 353.80000 | 35.0 | Cluster_07278 |
| 353.87000 | 35.0 | Cluster_20720 |
| 353.91000 | 35.0 | Cluster_01004 |
| 354.16000 | 35.0 | Cluster_20732 |
| 354.20000 | 35.0 | Cluster_20739 |
| 354.40000 | 35.0 | Cluster_01009 |
| 354.42000 | 35.0 | Cluster_01011 |
| 355.14000 | 35.0 | Cluster_20781 |
| 355.16000 | 35.0 | Cluster_20782 |
| 355.22000 | 35.0 | Cluster_20787 |
| 355.36000 | 35.0 | Cluster_20789 |
| 355.70000 | 35.0 | Cluster_20796 |
| 356.24000 | 35.0 | Cluster_20824 |
| 356.55000 | 35.0 | Cluster_07307 |
| 358.42000 | 35.0 | Cluster_20933 |
| 359.13000 | 35.0 | Cluster_07326 |
| 359.17000 | 35.0 | Cluster_00118 |
| 359.37000 | 35.0 | Cluster_20960 |
| 360.16000 | 35.0 | Cluster_20986 |
| 360.42000 | 35.0 | Cluster_01047 |
| 360.83000 | 35.0 | Cluster_21008 |
| 361.18000 | 35.0 | Cluster_21024 |
| 361.42000 | 35.0 | Cluster_01055 |
| 361.66000 | 35.0 | Cluster_21038 |
| 361.67000 | 35.0 | Cluster_21036 |
| 361.69000 | 35.0 | Cluster_21039 |
| 361.84000 | 35.0 | Cluster_07347 |
| 362.10000 | 35.0 | Cluster_21048 |
| 362.42000 | 35.0 | Cluster_01059 |
| 362.43000 | 35.0 | Cluster_21075 |
| 362.60000 | 35.0 | Cluster_21077 |
| 362.65000 | 35.0 | Cluster_21076 |
| 363.11000 | 35.0 | Cluster_21089 |
| 363.19000 | 35.0 | Cluster_21093 |
| 363.20000 | 35.0 | Cluster_21094 |
| 363.77000 | 35.0 | Cluster_21121 |

| | | |
|---|---|---|
| 364.16000 | 35.0 | Cluster_21135 |
| 364.50000 | 35.0 | Cluster_21161 |
| 364.65000 | 35.0 | Cluster_21162 |
| 365.24000 | 35.0 | Cluster_21221 |
| 365.79000 | 35.0 | Cluster_07396 |
| 365.90000 | 35.0 | Cluster_21241 |
| 366.27000 | 35.0 | Cluster_21267 |
| 366.74000 | 35.0 | Cluster_21285 |
| 367.22000 | 35.0 | Cluster_21306 |
| 367.50000 | 35.0 | Cluster_07416 |
| 367.53000 | 35.0 | Cluster_21324 |
| 368.16000 | 35.0 | Cluster_21368 |
| 368.20000 | 35.0 | Cluster_21366 |
| 368.47000 | 35.0 | Cluster_07428 |
| 369.67000 | 35.0 | Cluster_21437 |
| 370.08000 | 35.0 | Cluster_21453 |
| 370.50000 | 35.0 | Cluster_07445 |
| 370.74000 | 35.0 | Cluster_21476 |
| 370.93000 | 35.0 | Cluster_21486 |
| 371.18000 | 35.0 | Cluster_07449 |
| 371.24000 | 35.0 | Cluster_21501 |
| 371.71000 | 35.0 | Cluster_21517 |
| 371.89000 | 35.0 | Cluster_21531 |
| 372.17000 | 35.0 | Cluster_21537 |
| 372.23000 | 35.0 | Cluster_07464 |
| 372.67000 | 35.0 | Cluster_21575 |
| 372.69000 | 35.0 | Cluster_21572 |
| 373.12000 | 35.0 | Cluster_21589 |
| 373.14000 | 35.0 | Cluster_21585 |
| 373.16000 | 35.0 | Cluster_07470 |
| 373.19000 | 35.0 | Cluster_07471 |
| 373.22000 | 35.0 | Cluster_21597 |
| 373.90000 | 35.0 | Cluster_21628 |
| 374.19000 | 35.0 | Cluster_07480 |
| 374.40000 | 35.0 | Cluster_21650 |
| 374.68000 | 35.0 | Cluster_21656 |
| 375.15000 | 35.0 | Cluster_21668 |
| 375.89000 | 35.0 | Cluster_07496 |
| 377.48000 | 35.0 | Cluster_07512 |
| 377.77000 | 35.0 | Cluster_21780 |
| 378.17000 | 35.0 | Cluster_01122 |
| 378.18000 | 35.0 | Cluster_21788 |
| 378.20000 | 35.0 | Cluster_07523 |
| 378.67000 | 35.0 | Cluster_01125 |
| 379.26000 | 35.0 | Cluster_21841 |
| 380.45000 | 35.0 | Cluster_21886 |
| 380.68000 | 35.0 | Cluster_21891 |
| 381.22000 | 35.0 | Cluster_07550 |
| 381.70000 | 35.0 | Cluster_01141 |
| 382.18000 | 35.0 | Cluster_21953 |
| 382.66000 | 35.0 | Cluster_01144 |

| | | |
|---|---|---|
| 383.63000 | 35.0 | Cluster_22024 |
| 383.68000 | 35.0 | Cluster_22025 |
| 384.16000 | 35.0 | Cluster_01146 |
| 384.21000 | 35.0 | Cluster_22055 |
| 384.22000 | 35.0 | Cluster_22065 |
| 385.52000 | 35.0 | Cluster_07597 |
| 385.54000 | 35.0 | Cluster_07596 |
| 385.86000 | 35.0 | Cluster_22123 |
| 386.66000 | 35.0 | Cluster_22155 |
| 386.68000 | 35.0 | Cluster_22158 |
| 386.90000 | 35.0 | Cluster_22172 |
| 387.11000 | 35.0 | Cluster_22177 |
| 387.63000 | 35.0 | Cluster_22217 |
| 387.93000 | 35.0 | Cluster_22226 |
| 388.23000 | 35.0 | Cluster_22233 |
| 388.50000 | 35.0 | Cluster_07621 |
| 388.64000 | 35.0 | Cluster_22244 |
| 388.67000 | 35.0 | Cluster_22245 |
| 388.83000 | 35.0 | Cluster_07623 |
| 389.25000 | 35.0 | Cluster_22269 |
| 389.51000 | 35.0 | Cluster_07627 |
| 389.91000 | 35.0 | Cluster_22284 |
| 390.71000 | 35.0 | Cluster_22316 |
| 390.87000 | 35.0 | Cluster_07651 |
| 390.88000 | 35.0 | Cluster_07646 |
| 391.50000 | 35.0 | Cluster_07654 |
| 391.65000 | 35.0 | Cluster_22334 |
| 391.75000 | 35.0 | Cluster_22344 |
| 391.82000 | 35.0 | Cluster_07658 |
| 392.22000 | 35.0 | Cluster_07665 |
| 392.25000 | 35.0 | Cluster_22358 |
| 392.73000 | 35.0 | Cluster_22381 |
| 392.77000 | 35.0 | Cluster_22377 |
| 393.19000 | 35.0 | Cluster_01195 |
| 393.53000 | 35.0 | Cluster_07674 |
| 393.69000 | 35.0 | Cluster_01197 |
| 393.71000 | 35.0 | Cluster_22428 |
| 393.98000 | 35.0 | Cluster_22479 |
| 394.21000 | 35.0 | Cluster_22490 |
| 394.71000 | 35.0 | Cluster_22522 |
| 394.72000 | 35.0 | Cluster_22521 |
| 395.89000 | 35.0 | Cluster_07703 |
| 396.23000 | 35.0 | Cluster_22589 |
| 396.30000 | 35.0 | Cluster_22595 |
| 397.39000 | 35.0 | Cluster_22628 |
| 397.67000 | 35.0 | Cluster_22634 |
| 397.72000 | 35.0 | Cluster_22641 |
| 397.90000 | 35.0 | Cluster_07736 |
| 398.18000 | 35.0 | Cluster_01226 |
| 398.44000 | 35.0 | Cluster_01230 |
| 400.24000 | 35.0 | Cluster_22747 |

| | | |
|---|---|---|
| 400.68000 | 35.0 | Cluster_22780 |
| 400.90000 | 35.0 | Cluster_07771 |
| 401.70000 | 35.0 | Cluster_22833 |
| 402.32000 | 35.0 | Cluster_22866 |
| 402.48000 | 35.0 | Cluster_22868 |
| 402.72000 | 35.0 | Cluster_22878 |
| 403.44000 | 35.0 | Cluster_01260 |
| 403.54000 | 35.0 | Cluster_07797 |
| 404.47000 | 35.0 | Cluster_01272 |
| 405.88000 | 35.0 | Cluster_07831 |
| 406.22000 | 35.0 | Cluster_23023 |
| 406.68000 | 35.0 | Cluster_01285 |
| 406.73000 | 35.0 | Cluster_23046 |
| 406.95000 | 35.0 | Cluster_01293 |
| 407.17000 | 35.0 | Cluster_23059 |
| 407.25000 | 35.0 | Cluster_23066 |
| 407.67000 | 35.0 | Cluster_01302 |
| 407.75000 | 35.0 | Cluster_23083 |
| 407.93000 | 35.0 | Cluster_01303 |
| 408.22000 | 35.0 | Cluster_23104 |
| 408.54000 | 35.0 | Cluster_07863 |
| 408.66000 | 35.0 | Cluster_01305 |
| 408.91000 | 35.0 | Cluster_07864 |
| 408.92000 | 35.0 | Cluster_01307 |
| 409.19000 | 35.0 | Cluster_07867 |
| 409.25000 | 35.0 | Cluster_23160 |
| 409.69000 | 35.0 | Cluster_23170 |
| 410.53000 | 35.0 | Cluster_07882 |
| 411.72000 | 35.0 | Cluster_23258 |
| 412.67000 | 35.0 | Cluster_23299 |
| 412.68000 | 35.0 | Cluster_01326 |
| 413.23000 | 35.0 | Cluster_23320 |
| 413.87000 | 35.0 | Cluster_07913 |
| 413.92000 | 35.0 | Cluster_07916 |
| 413.94000 | 35.0 | Cluster_23350 |
| 413.98000 | 35.0 | Cluster_23354 |
| 414.71000 | 35.0 | Cluster_23379 |
| 414.85000 | 35.0 | Cluster_23397 |
| 414.87000 | 35.0 | Cluster_07924 |
| 415.22000 | 35.0 | Cluster_23422 |
| 415.73000 | 35.0 | Cluster_23434 |
| 416.00000 | 35.0 | Cluster_23448 |
| 416.22000 | 35.0 | Cluster_23463 |
| 416.76000 | 35.0 | Cluster_23476 |
| 416.88000 | 35.0 | Cluster_23482 |
| 416.94000 | 35.0 | Cluster_01353 |
| 418.25000 | 35.0 | Cluster_23534 |
| 418.47000 | 35.0 | Cluster_23549 |
| 419.18000 | 35.0 | Cluster_01360 |
| 419.21000 | 35.0 | Cluster_07978 |
| 419.88000 | 35.0 | Cluster_23600 |

| | | |
|---|---|---|
| 420.19000 | 35.0 | Cluster_23617 |
| 420.23000 | 35.0 | Cluster_23612 |
| 420.71000 | 35.0 | Cluster_23632 |
| 421.56000 | 35.0 | Cluster_08005 |
| 422.12000 | 35.0 | Cluster_08008 |
| 422.55000 | 35.0 | Cluster_08014 |
| 422.58000 | 35.0 | Cluster_08015 |
| 423.14000 | 35.0 | Cluster_23727 |
| 423.21000 | 35.0 | Cluster_08022 |
| 423.24000 | 35.0 | Cluster_23740 |
| 423.44000 | 35.0 | Cluster_01387 |
| 423.71000 | 35.0 | Cluster_23753 |
| 423.74000 | 35.0 | Cluster_23766 |
| 424.18000 | 35.0 | Cluster_23787 |
| 424.51000 | 35.0 | Cluster_23811 |
| 424.68000 | 35.0 | Cluster_23813 |
| 425.18000 | 35.0 | Cluster_23850 |
| 425.46000 | 35.0 | Cluster_23865 |
| 425.87000 | 35.0 | Cluster_08056 |
| 425.88000 | 35.0 | Cluster_23895 |
| 426.14000 | 35.0 | Cluster_23901 |
| 426.24000 | 35.0 | Cluster_23924 |
| 426.44000 | 35.0 | Cluster_23942 |
| 426.88000 | 35.0 | Cluster_08077 |
| 427.51000 | 35.0 | Cluster_08094 |
| 428.56000 | 35.0 | Cluster_08107 |
| 428.72000 | 35.0 | Cluster_24045 |
| 428.79000 | 35.0 | Cluster_24059 |
| 429.24000 | 35.0 | Cluster_24074 |
| 429.57000 | 35.0 | Cluster_08120 |
| 429.72000 | 35.0 | Cluster_24100 |
| 429.76000 | 35.0 | Cluster_24093 |
| 429.85000 | 35.0 | Cluster_08124 |
| 429.95000 | 35.0 | Cluster_01432 |
| 430.73000 | 35.0 | Cluster_24136 |
| 431.21000 | 35.0 | Cluster_24172 |
| 431.41000 | 35.0 | Cluster_24179 |
| 431.74000 | 35.0 | Cluster_24194 |
| 432.16000 | 35.0 | Cluster_24207 |
| 432.21000 | 35.0 | Cluster_24215 |
| 432.87000 | 35.0 | Cluster_08164 |
| 433.96000 | 35.0 | Cluster_24289 |
| 434.79000 | 35.0 | Cluster_24347 |
| 434.97000 | 35.0 | Cluster_24350 |
| 435.17000 | 35.0 | Cluster_24357 |
| 435.24000 | 35.0 | Cluster_08201 |
| 435.37000 | 35.0 | Cluster_24383 |
| 435.70000 | 35.0 | Cluster_01481 |
| 435.97000 | 35.0 | Cluster_01487 |
| 436.48000 | 35.0 | Cluster_01491 |
| 437.37000 | 35.0 | Cluster_24491 |

| | | |
|---|---|---|
| 437.46000 | 35.0 | Cluster_01497 |
| 438.22000 | 35.0 | Cluster_24532 |
| 438.30000 | 35.0 | Cluster_24543 |
| 438.96000 | 35.0 | Cluster_01503 |
| 438.97000 | 35.0 | Cluster_01504 |
| 439.82000 | 35.0 | Cluster_24632 |
| 440.21000 | 35.0 | Cluster_01512 |
| 440.56000 | 35.0 | Cluster_24655 |
| 440.74000 | 35.0 | Cluster_24659 |
| 440.86000 | 35.0 | Cluster_08277 |
| 441.21000 | 35.0 | Cluster_24682 |
| 441.24000 | 35.0 | Cluster_01516 |
| 441.73000 | 35.0 | Cluster_24703 |
| 441.87000 | 35.0 | Cluster_08293 |
| 442.00000 | 35.0 | Cluster_24716 |
| 442.28000 | 35.0 | Cluster_24730 |
| 442.57000 | 35.0 | Cluster_08303 |
| 442.77000 | 35.0 | Cluster_24758 |
| 442.89000 | 35.0 | Cluster_24763 |
| 442.98000 | 35.0 | Cluster_24761 |
| 443.12000 | 35.0 | Cluster_24771 |
| 443.22000 | 35.0 | Cluster_24782 |
| 443.28000 | 35.0 | Cluster_24785 |
| 443.29000 | 35.0 | Cluster_24795 |
| 443.89000 | 35.0 | Cluster_08322 |
| 443.92000 | 35.0 | Cluster_08324 |
| 444.72000 | 35.0 | Cluster_24866 |
| 445.18000 | 35.0 | Cluster_08343 |
| 445.55000 | 35.0 | Cluster_24900 |
| 446.27000 | 35.0 | Cluster_24941 |
| 446.73000 | 35.0 | Cluster_24959 |
| 446.91000 | 35.0 | Cluster_08369 |
| 447.17000 | 35.0 | Cluster_24974 |
| 447.25000 | 35.0 | Cluster_08376 |
| 447.28000 | 35.0 | Cluster_24982 |
| 447.29000 | 35.0 | Cluster_24988 |
| 447.60000 | 35.0 | Cluster_08380 |
| 448.30000 | 35.0 | Cluster_25022 |
| 448.79000 | 35.0 | Cluster_25051 |
| 448.97000 | 35.0 | Cluster_01573 |
| 449.25000 | 35.0 | Cluster_08416 |
| 449.74000 | 35.0 | Cluster_25092 |
| 449.84000 | 35.0 | Cluster_25102 |
| 450.23000 | 35.0 | Cluster_25125 |
| 450.93000 | 35.0 | Cluster_08435 |
| 451.14000 | 35.0 | Cluster_25153 |
| 451.17000 | 35.0 | Cluster_25168 |
| 451.57000 | 35.0 | Cluster_08446 |
| 452.13000 | 35.0 | Cluster_25195 |
| 452.26000 | 35.0 | Cluster_08462 |
| 452.75000 | 35.0 | Cluster_25235 |

| | | |
|---|---|---|
| 453.14000 | 35.0 | Cluster_25244 |
| 453.24000 | 35.0 | Cluster_25248 |
| 454.68000 | 35.0 | Cluster_25320 |
| 454.70000 | 35.0 | Cluster_25319 |
| 454.74000 | 35.0 | Cluster_25325 |
| 455.15000 | 35.0 | Cluster_25343 |
| 455.18000 | 35.0 | Cluster_01617 |
| 455.20000 | 35.0 | Cluster_08518 |
| 456.24000 | 35.0 | Cluster_25393 |
| 456.26000 | 35.0 | Cluster_08536 |
| 456.57000 | 35.0 | Cluster_08539 |
| 456.75000 | 35.0 | Cluster_25404 |
| 456.78000 | 35.0 | Cluster_25413 |
| 457.20000 | 35.0 | Cluster_25422 |
| 457.75000 | 35.0 | Cluster_25448 |
| 457.76000 | 35.0 | Cluster_25454 |
| 458.69000 | 35.0 | Cluster_25493 |
| 458.76000 | 35.0 | Cluster_25500 |
| 459.22000 | 35.0 | Cluster_08566 |
| 459.25000 | 35.0 | Cluster_25514 |
| 459.78000 | 35.0 | Cluster_25540 |
| 459.90000 | 35.0 | Cluster_08580 |
| 460.27000 | 35.0 | Cluster_25572 |
| 460.77000 | 35.0 | Cluster_25581 |
| 460.97000 | 35.0 | Cluster_01663 |
| 461.60000 | 35.0 | Cluster_08592 |
| 461.70000 | 35.0 | Cluster_25618 |
| 461.73000 | 35.0 | Cluster_25620 |
| 462.22000 | 35.0 | Cluster_08608 |
| 462.50000 | 35.0 | Cluster_25643 |
| 463.61000 | 35.0 | Cluster_25677 |
| 463.73000 | 35.0 | Cluster_25692 |
| 464.22000 | 35.0 | Cluster_08632 |
| 464.30000 | 35.0 | Cluster_25714 |
| 464.55000 | 35.0 | Cluster_08640 |
| 465.77000 | 35.0 | Cluster_25807 |
| 465.95000 | 35.0 | Cluster_08659 |
| 465.99000 | 35.0 | Cluster_01692 |
| 466.27000 | 35.0 | Cluster_25822 |
| 466.90000 | 35.0 | Cluster_08673 |
| 467.18000 | 35.0 | Cluster_08677 |
| 467.25000 | 35.0 | Cluster_08678 |
| 467.30000 | 35.0 | Cluster_25858 |
| 467.67000 | 35.0 | Cluster_00144 |
| 467.72000 | 35.0 | Cluster_25877 |
| 468.89000 | 35.0 | Cluster_08711 |
| 469.26000 | 35.0 | Cluster_08720 |
| 469.74000 | 35.0 | Cluster_25967 |
| 469.88000 | 35.0 | Cluster_08724 |
| 470.43000 | 35.0 | Cluster_26002 |
| 470.56000 | 35.0 | Cluster_00010 |

| | | |
|---|---|---|
| 470.92000 | 35.0 | Cluster_08745 |
| 471.00000 | 35.0 | Cluster_01731 |
| 471.22000 | 35.0 | Cluster_08748 |
| 471.25000 | 35.0 | Cluster_01737 |
| 471.26000 | 35.0 | Cluster_01738 |
| 471.48000 | 35.0 | Cluster_26044 |
| 471.74000 | 35.0 | Cluster_26063 |
| 471.84000 | 35.0 | Cluster_26052 |
| 472.23000 | 35.0 | Cluster_26081 |
| 472.24000 | 35.0 | Cluster_08764 |
| 472.72000 | 35.0 | Cluster_26107 |
| 472.76000 | 35.0 | Cluster_26106 |
| 472.77000 | 35.0 | Cluster_26113 |
| 473.27000 | 35.0 | Cluster_26146 |
| 473.77000 | 35.0 | Cluster_26157 |
| 474.54000 | 35.0 | Cluster_08802 |
| 474.58000 | 35.0 | Cluster_08800 |
| 474.85000 | 35.0 | Cluster_08806 |
| 475.58000 | 35.0 | Cluster_08820 |
| 475.78000 | 35.0 | Cluster_26234 |
| 476.21000 | 35.0 | Cluster_26244 |
| 476.23000 | 35.0 | Cluster_08830 |
| 476.28000 | 35.0 | Cluster_08832 |
| 477.07000 | 35.0 | Cluster_26275 |
| 477.20000 | 35.0 | Cluster_08843 |
| 477.27000 | 35.0 | Cluster_08840 |
| 477.55000 | 35.0 | Cluster_08851 |
| 477.61000 | 35.0 | Cluster_08856 |
| 477.93000 | 35.0 | Cluster_08861 |
| 478.22000 | 35.0 | Cluster_26309 |
| 478.25000 | 35.0 | Cluster_26313 |
| 478.76000 | 35.0 | Cluster_26329 |
| 478.77000 | 35.0 | Cluster_26332 |
| 478.93000 | 35.0 | Cluster_08880 |
| 478.99000 | 35.0 | Cluster_01780 |
| 479.57000 | 35.0 | Cluster_08888 |
| 479.71000 | 35.0 | Cluster_26429 |
| 479.76000 | 35.0 | Cluster_26423 |
| 480.22000 | 35.0 | Cluster_08900 |
| 480.70000 | 35.0 | Cluster_26476 |
| 481.22000 | 35.0 | Cluster_26493 |
| 481.27000 | 35.0 | Cluster_26504 |
| 481.28000 | 35.0 | Cluster_08920 |
| 481.56000 | 35.0 | Cluster_08922 |
| 481.58000 | 35.0 | Cluster_08924 |
| 481.72000 | 35.0 | Cluster_26523 |
| 481.73000 | 35.0 | Cluster_01804 |
| 481.79000 | 35.0 | Cluster_26518 |
| 482.20000 | 35.0 | Cluster_01810 |
| 482.28000 | 35.0 | Cluster_26547 |
| 482.49000 | 35.0 | Cluster_26549 |

| | | |
|---|---|---|
| 483.24000 | 35.0 | Cluster_26594 |
| 483.30000 | 35.0 | Cluster_26603 |
| 483.76000 | 35.0 | Cluster_26625 |
| 483.84000 | 35.0 | Cluster_26627 |
| 484.09000 | 35.0 | Cluster_26635 |
| 484.73000 | 35.0 | Cluster_26685 |
| 484.76000 | 35.0 | Cluster_26681 |
| 485.06000 | 35.0 | Cluster_00166 |
| 485.54000 | 35.0 | Cluster_08981 |
| 488.29000 | 35.0 | Cluster_26837 |
| 488.61000 | 35.0 | Cluster_09032 |
| 488.79000 | 35.0 | Cluster_26859 |
| 488.95000 | 35.0 | Cluster_09038 |
| 489.74000 | 35.0 | Cluster_26904 |
| 489.93000 | 35.0 | Cluster_09048 |
| 490.50000 | 35.0 | Cluster_26931 |
| 490.70000 | 35.0 | Cluster_26935 |
| 490.77000 | 35.0 | Cluster_26939 |
| 492.55000 | 35.0 | Cluster_09097 |
| 492.60000 | 35.0 | Cluster_09099 |
| 493.27000 | 35.0 | Cluster_27041 |
| 494.26000 | 35.0 | Cluster_27100 |
| 495.19000 | 35.0 | Cluster_27149 |
| 495.71000 | 35.0 | Cluster_27170 |
| 495.95000 | 35.0 | Cluster_09160 |
| 496.24000 | 35.0 | Cluster_01895 |
| 496.95000 | 35.0 | Cluster_09176 |
| 497.35000 | 35.0 | Cluster_27258 |
| 497.74000 | 35.0 | Cluster_27265 |
| 497.80000 | 35.0 | Cluster_27275 |
| 497.94000 | 35.0 | Cluster_09191 |
| 498.01000 | 35.0 | Cluster_01929 |
| 498.50000 | 35.0 | Cluster_27299 |
| 498.73000 | 35.0 | Cluster_27302 |
| 499.23000 | 35.0 | Cluster_09211 |
| 499.33000 | 35.0 | Cluster_27324 |
| 499.76000 | 35.0 | Cluster_01941 |
| 499.92000 | 35.0 | Cluster_09218 |
| 499.96000 | 35.0 | Cluster_09219 |
| 500.02000 | 35.0 | Cluster_27355 |
| 500.63000 | 35.0 | Cluster_09231 |
| 500.70000 | 35.0 | Cluster_27401 |
| 500.96000 | 35.0 | Cluster_01956 |
| 501.22000 | 35.0 | Cluster_27436 |
| 501.59000 | 35.0 | Cluster_09244 |
| 501.76000 | 35.0 | Cluster_27459 |
| 502.21000 | 35.0 | Cluster_01970 |
| 502.22000 | 35.0 | Cluster_27473 |
| 502.59000 | 35.0 | Cluster_09268 |
| 502.76000 | 35.0 | Cluster_27485 |
| 503.25000 | 35.0 | Cluster_27512 |

| | | |
|---|---|---|
| 503.28000 | 35.0 | Cluster_27518 |
| 504.21000 | 35.0 | Cluster_27547 |
| 504.56000 | 35.0 | Cluster_09288 |
| 504.98000 | 35.0 | Cluster_01984 |
| 505.01000 | 35.0 | Cluster_27567 |
| 505.24000 | 35.0 | Cluster_09298 |
| 505.52000 | 35.0 | Cluster_01990 |
| 505.60000 | 35.0 | Cluster_09305 |
| 505.79000 | 35.0 | Cluster_27593 |
| 506.73000 | 35.0 | Cluster_27611 |
| 506.74000 | 35.0 | Cluster_27621 |
| 506.76000 | 35.0 | Cluster_27614 |
| 507.25000 | 35.0 | Cluster_27638 |
| 507.97000 | 35.0 | Cluster_09350 |
| 508.62000 | 35.0 | Cluster_09366 |
| 508.70000 | 35.0 | Cluster_27686 |
| 508.71000 | 35.0 | Cluster_27692 |
| 509.20000 | 35.0 | Cluster_27715 |
| 509.26000 | 35.0 | Cluster_27718 |
| 509.87000 | 35.0 | Cluster_09377 |
| 509.91000 | 35.0 | Cluster_09383 |
| 511.23000 | 35.0 | Cluster_27820 |
| 511.79000 | 35.0 | Cluster_27852 |
| 511.91000 | 35.0 | Cluster_09420 |
| 512.25000 | 35.0 | Cluster_27867 |
| 512.27000 | 35.0 | Cluster_27865 |
| 513.09000 | 35.0 | Cluster_27898 |
| 513.77000 | 35.0 | Cluster_27929 |
| 514.72000 | 35.0 | Cluster_27964 |
| 514.74000 | 35.0 | Cluster_27971 |
| 515.25000 | 35.0 | Cluster_02053 |
| 515.48000 | 35.0 | Cluster_02061 |
| 515.90000 | 35.0 | Cluster_09478 |
| 516.74000 | 35.0 | Cluster_28062 |
| 517.92000 | 35.0 | Cluster_09503 |
| 518.00000 | 35.0 | Cluster_28115 |
| 518.26000 | 35.0 | Cluster_09508 |
| 518.31000 | 35.0 | Cluster_28130 |
| 518.51000 | 35.0 | Cluster_28138 |
| 518.59000 | 35.0 | Cluster_09521 |
| 518.78000 | 35.0 | Cluster_28152 |
| 519.59000 | 35.0 | Cluster_09531 |
| 519.81000 | 35.0 | Cluster_28200 |
| 520.04000 | 35.0 | Cluster_28202 |
| 520.51000 | 35.0 | Cluster_28225 |
| 520.80000 | 35.0 | Cluster_28240 |
| 520.81000 | 35.0 | Cluster_28232 |
| 521.02000 | 35.0 | Cluster_02095 |
| 521.26000 | 35.0 | Cluster_09553 |
| 521.66000 | 35.0 | Cluster_28271 |
| 522.22000 | 35.0 | Cluster_28286 |

| | | |
|---|---|---|
| 522.45000 | 35.0 | Cluster_02102 |
| 522.50000 | 35.0 | Cluster_28303 |
| 522.62000 | 35.0 | Cluster_09582 |
| 522.75000 | 35.0 | Cluster_28315 |
| 523.22000 | 35.0 | Cluster_28350 |
| 523.27000 | 35.0 | Cluster_28334 |
| 523.35000 | 35.0 | Cluster_28353 |
| 523.92000 | 35.0 | Cluster_09603 |
| 523.97000 | 35.0 | Cluster_28374 |
| 524.28000 | 35.0 | Cluster_28386 |
| 526.18000 | 35.0 | Cluster_28467 |
| 526.20000 | 35.0 | Cluster_09656 |
| 526.93000 | 35.0 | Cluster_09676 |
| 526.95000 | 35.0 | Cluster_09677 |
| 527.76000 | 35.0 | Cluster_28507 |
| 528.23000 | 35.0 | Cluster_28536 |
| 528.27000 | 35.0 | Cluster_00184 |
| 528.32000 | 35.0 | Cluster_28539 |
| 528.36000 | 35.0 | Cluster_28552 |
| 528.97000 | 35.0 | Cluster_09710 |
| 529.25000 | 35.0 | Cluster_09712 |
| 529.79000 | 35.0 | Cluster_28615 |
| 530.20000 | 35.0 | Cluster_28629 |
| 530.26000 | 35.0 | Cluster_02153 |
| 530.51000 | 35.0 | Cluster_28657 |
| 530.81000 | 35.0 | Cluster_28677 |
| 530.94000 | 35.0 | Cluster_09741 |
| 531.75000 | 35.0 | Cluster_28707 |
| 531.77000 | 35.0 | Cluster_28708 |
| 532.24000 | 35.0 | Cluster_09763 |
| 532.27000 | 35.0 | Cluster_09766 |
| 532.62000 | 35.0 | Cluster_09768 |
| 532.97000 | 35.0 | Cluster_09773 |
| 533.15000 | 35.0 | Cluster_28752 |
| 533.16000 | 35.0 | Cluster_28753 |
| 533.19000 | 35.0 | Cluster_09774 |
| 533.32000 | 35.0 | Cluster_09778 |
| 533.70000 | 35.0 | Cluster_28783 |
| 533.82000 | 35.0 | Cluster_28799 |
| 533.91000 | 35.0 | Cluster_09792 |
| 534.28000 | 35.0 | Cluster_28807 |
| 534.59000 | 35.0 | Cluster_09803 |
| 535.32000 | 35.0 | Cluster_09812 |
| 535.80000 | 35.0 | Cluster_28867 |
| 535.94000 | 35.0 | Cluster_09822 |
| 536.06000 | 35.0 | Cluster_28878 |
| 536.26000 | 35.0 | Cluster_09823 |
| 536.27000 | 35.0 | Cluster_28888 |
| 536.48000 | 35.0 | Cluster_28899 |
| 536.60000 | 35.0 | Cluster_09834 |
| 537.23000 | 35.0 | Cluster_09849 |

| | | |
|---|---|---|
| 537.56000 | 35.0 | Cluster_09853 |
| 537.69000 | 35.0 | Cluster_28943 |
| 537.91000 | 35.0 | Cluster_09869 |
| 538.84000 | 35.0 | Cluster_28980 |
| 538.97000 | 35.0 | Cluster_09886 |
| 538.98000 | 35.0 | Cluster_09885 |
| 539.01000 | 35.0 | Cluster_28985 |
| 539.28000 | 35.0 | Cluster_28993 |
| 539.52000 | 35.0 | Cluster_29001 |
| 540.02000 | 35.0 | Cluster_09899 |
| 540.27000 | 35.0 | Cluster_29031 |
| 540.79000 | 35.0 | Cluster_29052 |
| 541.34000 | 35.0 | Cluster_29085 |
| 541.44000 | 35.0 | Cluster_29098 |
| 541.82000 | 35.0 | Cluster_29115 |
| 542.26000 | 35.0 | Cluster_02236 |
| 542.28000 | 35.0 | Cluster_02229 |
| 542.52000 | 35.0 | Cluster_02233 |
| 542.77000 | 35.0 | Cluster_29162 |
| 543.27000 | 35.0 | Cluster_29193 |
| 544.95000 | 35.0 | Cluster_09982 |
| 545.77000 | 35.0 | Cluster_29280 |
| 545.92000 | 35.0 | Cluster_09992 |
| 546.31000 | 35.0 | Cluster_29312 |
| 547.32000 | 35.0 | Cluster_29345 |
| 548.27000 | 35.0 | Cluster_29389 |
| 548.46000 | 35.0 | Cluster_00194 |
| 548.61000 | 35.0 | Cluster_10035 |
| 548.64000 | 35.0 | Cluster_10037 |
| 549.25000 | 35.0 | Cluster_10044 |
| 549.28000 | 35.0 | Cluster_29418 |
| 549.90000 | 35.0 | Cluster_10052 |
| 549.99000 | 35.0 | Cluster_10058 |
| 550.22000 | 35.0 | Cluster_29451 |
| 550.31000 | 35.0 | Cluster_29461 |
| 550.55000 | 35.0 | Cluster_10069 |
| 550.61000 | 35.0 | Cluster_10068 |
| 550.78000 | 35.0 | Cluster_29465 |
| 550.84000 | 35.0 | Cluster_29476 |
| 551.01000 | 35.0 | Cluster_29483 |
| 551.29000 | 35.0 | Cluster_29499 |
| 551.57000 | 35.0 | Cluster_10082 |
| 551.97000 | 35.0 | Cluster_10089 |
| 552.36000 | 35.0 | Cluster_29545 |
| 552.74000 | 35.0 | Cluster_29553 |
| 552.75000 | 35.0 | Cluster_29548 |
| 552.96000 | 35.0 | Cluster_10104 |
| 553.55000 | 35.0 | Cluster_29579 |
| 553.63000 | 35.0 | Cluster_10116 |
| 554.58000 | 35.0 | Cluster_10130 |
| 554.60000 | 35.0 | Cluster_10136 |

| | | |
|---|---|---|
| 554.97000 | 35.0 | Cluster_10144 |
| 556.25000 | 35.0 | Cluster_29692 |
| 556.68000 | 35.0 | Cluster_10177 |
| 556.96000 | 35.0 | Cluster_10184 |
| 557.73000 | 35.0 | Cluster_29750 |
| 558.23000 | 35.0 | Cluster_10209 |
| 558.65000 | 35.0 | Cluster_10215 |
| 558.81000 | 35.0 | Cluster_29792 |
| 559.29000 | 35.0 | Cluster_29811 |
| 560.35000 | 35.0 | Cluster_29845 |
| 560.65000 | 35.0 | Cluster_10245 |
| 560.83000 | 35.0 | Cluster_29862 |
| 561.28000 | 35.0 | Cluster_29879 |
| 561.64000 | 35.0 | Cluster_10262 |
| 561.70000 | 35.0 | Cluster_29892 |
| 561.81000 | 35.0 | Cluster_29901 |
| 561.98000 | 35.0 | Cluster_10266 |
| 562.07000 | 35.0 | Cluster_29906 |
| 562.29000 | 35.0 | Cluster_29915 |
| 563.32000 | 35.0 | Cluster_10288 |
| 563.50000 | 35.0 | Cluster_02355 |
| 563.75000 | 35.0 | Cluster_29975 |
| 563.76000 | 35.0 | Cluster_29965 |
| 564.78000 | 35.0 | Cluster_30003 |
| 564.79000 | 35.0 | Cluster_02366 |
| 565.56000 | 35.0 | Cluster_30039 |
| 565.60000 | 35.0 | Cluster_10318 |
| 566.33000 | 35.0 | Cluster_30080 |
| 566.74000 | 35.0 | Cluster_02376 |
| 566.79000 | 35.0 | Cluster_30088 |
| 566.90000 | 35.0 | Cluster_30092 |
| 566.98000 | 35.0 | Cluster_10333 |
| 567.07000 | 35.0 | Cluster_30105 |
| 567.24000 | 35.0 | Cluster_30110 |
| 567.30000 | 35.0 | Cluster_30123 |
| 567.31000 | 35.0 | Cluster_30114 |
| 568.56000 | 35.0 | Cluster_02384 |
| 568.73000 | 35.0 | Cluster_30166 |
| 568.79000 | 35.0 | Cluster_30177 |
| 568.80000 | 35.0 | Cluster_30176 |
| 568.85000 | 35.0 | Cluster_30178 |
| 569.06000 | 35.0 | Cluster_02386 |
| 569.57000 | 35.0 | Cluster_02390 |
| 569.82000 | 35.0 | Cluster_30204 |
| 570.30000 | 35.0 | Cluster_02396 |
| 570.36000 | 35.0 | Cluster_30219 |
| 570.73000 | 35.0 | Cluster_02397 |
| 570.77000 | 35.0 | Cluster_30236 |
| 571.22000 | 35.0 | Cluster_30252 |
| 571.34000 | 35.0 | Cluster_30256 |
| 571.64000 | 35.0 | Cluster_10409 |

| | | |
|---|---|---|
| 571.77000 | 35.0 | Cluster_30264 |
| 572.13000 | 35.0 | Cluster_00213 |
| 572.30000 | 35.0 | Cluster_10420 |
| 572.79000 | 35.0 | Cluster_30301 |
| 573.35000 | 35.0 | Cluster_30317 |
| 574.26000 | 35.0 | Cluster_10456 |
| 574.80000 | 35.0 | Cluster_30355 |
| 575.25000 | 35.0 | Cluster_02433 |
| 575.33000 | 35.0 | Cluster_02435 |
| 576.55000 | 35.0 | Cluster_02439 |
| 576.77000 | 35.0 | Cluster_30417 |
| 576.98000 | 35.0 | Cluster_10494 |
| 577.30000 | 35.0 | Cluster_10500 |
| 577.81000 | 35.0 | Cluster_30446 |
| 578.32000 | 35.0 | Cluster_30460 |
| 578.55000 | 35.0 | Cluster_02454 |
| 578.86000 | 35.0 | Cluster_30486 |
| 578.98000 | 35.0 | Cluster_10523 |
| 579.33000 | 35.0 | Cluster_30494 |
| 579.55000 | 35.0 | Cluster_02460 |
| 579.61000 | 35.0 | Cluster_10531 |
| 579.63000 | 35.0 | Cluster_10533 |
| 579.77000 | 35.0 | Cluster_30512 |
| 579.81000 | 35.0 | Cluster_30517 |
| 579.83000 | 35.0 | Cluster_30515 |
| 580.36000 | 35.0 | Cluster_30555 |
| 580.86000 | 35.0 | Cluster_02466 |
| 581.28000 | 35.0 | Cluster_02468 |
| 581.31000 | 35.0 | Cluster_30577 |
| 581.96000 | 35.0 | Cluster_10563 |
| 582.31000 | 35.0 | Cluster_30648 |
| 582.54000 | 35.0 | Cluster_02478 |
| 582.65000 | 35.0 | Cluster_10574 |
| 582.70000 | 35.0 | Cluster_00226 |
| 582.74000 | 35.0 | Cluster_30654 |
| 584.05000 | 35.0 | Cluster_30718 |
| 584.31000 | 35.0 | Cluster_30724 |
| 584.54000 | 35.0 | Cluster_30733 |
| 585.02000 | 35.0 | Cluster_30763 |
| 585.29000 | 35.0 | Cluster_10612 |
| 585.35000 | 35.0 | Cluster_02501 |
| 585.80000 | 35.0 | Cluster_30791 |
| 585.84000 | 35.0 | Cluster_30792 |
| 586.62000 | 35.0 | Cluster_10630 |
| 586.75000 | 35.0 | Cluster_30815 |
| 587.30000 | 35.0 | Cluster_10642 |
| 587.33000 | 35.0 | Cluster_30848 |
| 587.34000 | 35.0 | Cluster_30837 |
| 587.65000 | 35.0 | Cluster_10646 |
| 588.69000 | 35.0 | Cluster_30892 |
| 588.82000 | 35.0 | Cluster_30900 |

| | | |
|---|---|---|
| 589.55000 | 35.0 | Cluster_30932 |
| 589.74000 | 35.0 | Cluster_30933 |
| 589.96000 | 35.0 | Cluster_10677 |
| 590.22000 | 35.0 | Cluster_30952 |
| 590.30000 | 35.0 | Cluster_30953 |
| 591.04000 | 35.0 | Cluster_30998 |
| 591.44000 | 35.0 | Cluster_31008 |
| 591.52000 | 35.0 | Cluster_02538 |
| 591.66000 | 35.0 | Cluster_10703 |
| 591.81000 | 35.0 | Cluster_31027 |
| 592.01000 | 35.0 | Cluster_10706 |
| 592.79000 | 35.0 | Cluster_31056 |
| 592.82000 | 35.0 | Cluster_31060 |
| 592.91000 | 35.0 | Cluster_31063 |
| 593.06000 | 35.0 | Cluster_02547 |
| 593.07000 | 35.0 | Cluster_02548 |
| 593.33000 | 35.0 | Cluster_10724 |
| 593.34000 | 35.0 | Cluster_31073 |
| 593.93000 | 35.0 | Cluster_10743 |
| 594.32000 | 35.0 | Cluster_10753 |
| 595.59000 | 35.0 | Cluster_31146 |
| 595.62000 | 35.0 | Cluster_10769 |
| 596.75000 | 35.0 | Cluster_31177 |
| 597.67000 | 35.0 | Cluster_10808 |
| 597.81000 | 35.0 | Cluster_31212 |
| 597.82000 | 35.0 | Cluster_31208 |
| 598.29000 | 35.0 | Cluster_10814 |
| 598.36000 | 35.0 | Cluster_10818 |
| 599.36000 | 35.0 | Cluster_31267 |
| 599.66000 | 35.0 | Cluster_10840 |
| 599.95000 | 35.0 | Cluster_10848 |
| 600.81000 | 35.0 | Cluster_31316 |
| 600.97000 | 35.0 | Cluster_10858 |
| 601.36000 | 35.0 | Cluster_31339 |
| 602.32000 | 35.0 | Cluster_31390 |
| 602.83000 | 35.0 | Cluster_31406 |
| 603.30000 | 35.0 | Cluster_31409 |
| 603.51000 | 35.0 | Cluster_02598 |
| 603.53000 | 35.0 | Cluster_31419 |
| 603.96000 | 35.0 | Cluster_10888 |
| 604.80000 | 35.0 | Cluster_31463 |
| 605.31000 | 35.0 | Cluster_31473 |
| 605.67000 | 35.0 | Cluster_10910 |
| 606.07000 | 35.0 | Cluster_31501 |
| 606.27000 | 35.0 | Cluster_31514 |
| 606.87000 | 35.0 | Cluster_31531 |
| 606.89000 | 35.0 | Cluster_10941 |
| 607.32000 | 35.0 | Cluster_31536 |
| 608.09000 | 35.0 | Cluster_02626 |
| 608.32000 | 35.0 | Cluster_31587 |
| 608.49000 | 35.0 | Cluster_31590 |

| | | |
|---|---|---|
| 608.78000 | 35.0 | Cluster_31595 |
| 608.79000 | 35.0 | Cluster_31601 |
| 609.06000 | 35.0 | Cluster_31605 |
| 609.81000 | 35.0 | Cluster_31625 |
| 610.31000 | 35.0 | Cluster_31655 |
| 611.34000 | 35.0 | Cluster_31685 |
| 612.32000 | 35.0 | Cluster_11010 |
| 613.83000 | 35.0 | Cluster_31794 |
| 614.06000 | 35.0 | Cluster_31796 |
| 614.07000 | 35.0 | Cluster_02670 |
| 614.19000 | 35.0 | Cluster_31797 |
| 614.26000 | 35.0 | Cluster_11035 |
| 615.83000 | 35.0 | Cluster_31852 |
| 616.08000 | 35.0 | Cluster_02680 |
| 616.32000 | 35.0 | Cluster_11075 |
| 616.55000 | 35.0 | Cluster_31873 |
| 616.83000 | 35.0 | Cluster_31882 |
| 617.02000 | 35.0 | Cluster_31892 |
| 617.35000 | 35.0 | Cluster_31911 |
| 617.93000 | 35.0 | Cluster_31935 |
| 618.05000 | 35.0 | Cluster_02693 |
| 618.79000 | 35.0 | Cluster_31952 |
| 619.29000 | 35.0 | Cluster_02701 |
| 619.33000 | 35.0 | Cluster_31985 |
| 619.62000 | 35.0 | Cluster_11116 |
| 620.53000 | 35.0 | Cluster_32028 |
| 620.60000 | 35.0 | Cluster_11128 |
| 621.30000 | 35.0 | Cluster_32049 |
| 621.36000 | 35.0 | Cluster_32053 |
| 621.39000 | 35.0 | Cluster_32052 |
| 621.79000 | 35.0 | Cluster_32057 |
| 621.98000 | 35.0 | Cluster_11154 |
| 622.78000 | 35.0 | Cluster_32095 |
| 622.98000 | 35.0 | Cluster_11175 |
| 623.28000 | 35.0 | Cluster_32112 |
| 623.36000 | 35.0 | Cluster_32126 |
| 623.65000 | 35.0 | Cluster_11191 |
| 623.98000 | 35.0 | Cluster_11189 |
| 624.20000 | 35.0 | Cluster_32153 |
| 624.28000 | 35.0 | Cluster_32154 |
| 624.63000 | 35.0 | Cluster_11202 |
| 625.04000 | 35.0 | Cluster_02732 |
| 625.32000 | 35.0 | Cluster_11203 |
| 625.58000 | 35.0 | Cluster_02735 |
| 626.00000 | 35.0 | Cluster_11225 |
| 626.07000 | 35.0 | Cluster_32208 |
| 626.27000 | 35.0 | Cluster_32218 |
| 626.28000 | 35.0 | Cluster_32212 |
| 626.83000 | 35.0 | Cluster_32244 |
| 627.30000 | 35.0 | Cluster_32266 |
| 627.31000 | 35.0 | Cluster_32263 |

| | | |
|---|---|---|
| 627.50000 | 35.0 | Cluster_32275 |
| 627.79000 | 35.0 | Cluster_32283 |
| 628.07000 | 35.0 | Cluster_32292 |
| 628.86000 | 35.0 | Cluster_32321 |
| 629.63000 | 35.0 | Cluster_11289 |
| 629.83000 | 35.0 | Cluster_32344 |
| 629.96000 | 35.0 | Cluster_11292 |
| 630.84000 | 35.0 | Cluster_32375 |
| 631.37000 | 35.0 | Cluster_11312 |
| 631.80000 | 35.0 | Cluster_32410 |
| 632.05000 | 35.0 | Cluster_11324 |
| 632.35000 | 35.0 | Cluster_11329 |
| 632.78000 | 35.0 | Cluster_32445 |
| 632.84000 | 35.0 | Cluster_32450 |
| 633.00000 | 35.0 | Cluster_11339 |
| 633.32000 | 35.0 | Cluster_32465 |
| 633.36000 | 35.0 | Cluster_32472 |
| 635.07000 | 35.0 | Cluster_02772 |
| 635.82000 | 35.0 | Cluster_02775 |
| 636.32000 | 35.0 | Cluster_32584 |
| 636.84000 | 35.0 | Cluster_32596 |
| 637.62000 | 35.0 | Cluster_11403 |
| 637.83000 | 35.0 | Cluster_32632 |
| 638.35000 | 35.0 | Cluster_32655 |
| 638.36000 | 35.0 | Cluster_11416 |
| 638.39000 | 35.0 | Cluster_32659 |
| 638.79000 | 35.0 | Cluster_32669 |
| 638.85000 | 35.0 | Cluster_32673 |
| 639.85000 | 35.0 | Cluster_32724 |
| 639.99000 | 35.0 | Cluster_11446 |
| 640.32000 | 35.0 | Cluster_32732 |
| 641.64000 | 35.0 | Cluster_11472 |
| 641.84000 | 35.0 | Cluster_32794 |
| 642.02000 | 35.0 | Cluster_11476 |
| 643.07000 | 35.0 | Cluster_32838 |
| 643.83000 | 35.0 | Cluster_32863 |
| 644.00000 | 35.0 | Cluster_11510 |
| 644.30000 | 35.0 | Cluster_02836 |
| 644.36000 | 35.0 | Cluster_32878 |
| 644.37000 | 35.0 | Cluster_11511 |
| 645.00000 | 35.0 | Cluster_11529 |
| 645.85000 | 35.0 | Cluster_32926 |
| 646.30000 | 35.0 | Cluster_32945 |
| 648.34000 | 35.0 | Cluster_33005 |
| 648.67000 | 35.0 | Cluster_11566 |
| 648.88000 | 35.0 | Cluster_33029 |
| 649.01000 | 35.0 | Cluster_11574 |
| 649.77000 | 35.0 | Cluster_33054 |
| 650.31000 | 35.0 | Cluster_33085 |
| 651.01000 | 35.0 | Cluster_11595 |
| 651.08000 | 35.0 | Cluster_33114 |

| | | |
|---|---|---|
| 651.32000 | 35.0 | Cluster_33124 |
| 651.36000 | 35.0 | Cluster_11600 |
| 651.86000 | 35.0 | Cluster_33158 |
| 651.99000 | 35.0 | Cluster_11614 |
| 652.06000 | 35.0 | Cluster_33159 |
| 652.35000 | 35.0 | Cluster_33162 |
| 652.86000 | 35.0 | Cluster_33187 |
| 653.33000 | 35.0 | Cluster_33199 |
| 653.57000 | 35.0 | Cluster_33207 |
| 653.60000 | 35.0 | Cluster_02883 |
| 654.05000 | 35.0 | Cluster_33221 |
| 654.68000 | 35.0 | Cluster_11655 |
| 654.78000 | 35.0 | Cluster_33237 |
| 655.99000 | 35.0 | Cluster_11675 |
| 656.63000 | 35.0 | Cluster_11676 |
| 657.90000 | 35.0 | Cluster_33331 |
| 657.95000 | 35.0 | Cluster_33350 |
| 658.07000 | 35.0 | Cluster_33340 |
| 658.66000 | 35.0 | Cluster_11699 |
| 659.28000 | 35.0 | Cluster_33386 |
| 659.60000 | 35.0 | Cluster_33396 |
| 659.63000 | 35.0 | Cluster_02907 |
| 659.98000 | 35.0 | Cluster_11725 |
| 660.33000 | 35.0 | Cluster_33416 |
| 660.46000 | 35.0 | Cluster_33423 |
| 660.84000 | 35.0 | Cluster_33430 |
| 661.01000 | 35.0 | Cluster_11741 |
| 661.31000 | 35.0 | Cluster_33436 |
| 662.02000 | 35.0 | Cluster_11748 |
| 662.12000 | 35.0 | Cluster_02928 |
| 662.30000 | 35.0 | Cluster_33472 |
| 663.37000 | 35.0 | Cluster_33509 |
| 663.55000 | 35.0 | Cluster_02941 |
| 663.60000 | 35.0 | Cluster_02940 |
| 665.32000 | 35.0 | Cluster_33570 |
| 666.35000 | 35.0 | Cluster_02964 |
| 666.68000 | 35.0 | Cluster_11814 |
| 666.95000 | 35.0 | Cluster_33617 |
| 667.82000 | 35.0 | Cluster_33644 |
| 668.33000 | 35.0 | Cluster_33651 |
| 668.35000 | 35.0 | Cluster_11848 |
| 669.10000 | 35.0 | Cluster_02977 |
| 669.33000 | 35.0 | Cluster_33680 |
| 669.78000 | 35.0 | Cluster_33695 |
| 670.01000 | 35.0 | Cluster_11876 |
| 670.40000 | 35.0 | Cluster_33725 |
| 670.80000 | 35.0 | Cluster_33734 |
| 670.82000 | 35.0 | Cluster_33742 |
| 670.87000 | 35.0 | Cluster_02986 |
| 671.44000 | 35.0 | Cluster_33760 |
| 671.82000 | 35.0 | Cluster_33770 |

| | | |
|---|---|---|
| 671.97000 | 35.0 | Cluster_11894 |
| 672.35000 | 35.0 | Cluster_33781 |
| 672.81000 | 35.0 | Cluster_33789 |
| 673.09000 | 35.0 | Cluster_03005 |
| 673.37000 | 35.0 | Cluster_33802 |
| 673.62000 | 35.0 | Cluster_33824 |
| 673.65000 | 35.0 | Cluster_11927 |
| 674.11000 | 35.0 | Cluster_03014 |
| 674.32000 | 35.0 | Cluster_11933 |
| 674.95000 | 35.0 | Cluster_00274 |
| 675.37000 | 35.0 | Cluster_33861 |
| 675.60000 | 35.0 | Cluster_33864 |
| 675.64000 | 35.0 | Cluster_03025 |
| 675.84000 | 35.0 | Cluster_03026 |
| 676.00000 | 35.0 | Cluster_11956 |
| 676.32000 | 35.0 | Cluster_11959 |
| 677.20000 | 35.0 | Cluster_33905 |
| 677.89000 | 35.0 | Cluster_33946 |
| 678.81000 | 35.0 | Cluster_33981 |
| 678.84000 | 35.0 | Cluster_33980 |
| 678.85000 | 35.0 | Cluster_03050 |
| 679.38000 | 35.0 | Cluster_33996 |
| 679.79000 | 35.0 | Cluster_34007 |
| 679.86000 | 35.0 | Cluster_34018 |
| 680.08000 | 35.0 | Cluster_03060 |
| 680.09000 | 35.0 | Cluster_34023 |
| 680.87000 | 35.0 | Cluster_34054 |
| 681.08000 | 35.0 | Cluster_03066 |
| 681.32000 | 35.0 | Cluster_12020 |
| 681.35000 | 35.0 | Cluster_34074 |
| 682.09000 | 35.0 | Cluster_34089 |
| 682.31000 | 35.0 | Cluster_34106 |
| 682.38000 | 35.0 | Cluster_03074 |
| 683.61000 | 35.0 | Cluster_12059 |
| 683.81000 | 35.0 | Cluster_34134 |
| 683.99000 | 35.0 | Cluster_12062 |
| 684.08000 | 35.0 | Cluster_03088 |
| 684.82000 | 35.0 | Cluster_34171 |
| 684.83000 | 35.0 | Cluster_34174 |
| 685.09000 | 35.0 | Cluster_03095 |
| 685.10000 | 35.0 | Cluster_03102 |
| 685.35000 | 35.0 | Cluster_34201 |
| 686.04000 | 35.0 | Cluster_12086 |
| 686.50000 | 35.0 | Cluster_00285 |
| 686.75000 | 35.0 | Cluster_00286 |
| 687.66000 | 35.0 | Cluster_34255 |
| 689.34000 | 35.0 | Cluster_34298 |
| 689.36000 | 35.0 | Cluster_34295 |
| 689.57000 | 35.0 | Cluster_03142 |
| 690.82000 | 35.0 | Cluster_34324 |
| 690.84000 | 35.0 | Cluster_34327 |

| | | |
|---|---|---|
| 691.59000 | 35.0 | Cluster_03162 |
| 691.79000 | 35.0 | Cluster_34357 |
| 691.83000 | 35.0 | Cluster_34369 |
| 692.26000 | 35.0 | Cluster_34373 |
| 692.32000 | 35.0 | Cluster_03167 |
| 692.35000 | 35.0 | Cluster_03168 |
| 692.60000 | 35.0 | Cluster_03182 |
| 692.84000 | 35.0 | Cluster_34402 |
| 693.56000 | 35.0 | Cluster_03192 |
| 693.60000 | 35.0 | Cluster_34413 |
| 693.88000 | 35.0 | Cluster_34418 |
| 694.36000 | 35.0 | Cluster_34424 |
| 694.84000 | 35.0 | Cluster_34436 |
| 694.87000 | 35.0 | Cluster_34443 |
| 695.01000 | 35.0 | Cluster_12183 |
| 695.36000 | 35.0 | Cluster_34458 |
| 695.70000 | 35.0 | Cluster_12190 |
| 695.87000 | 35.0 | Cluster_34468 |
| 696.32000 | 35.0 | Cluster_34489 |
| 696.34000 | 35.0 | Cluster_03210 |
| 696.39000 | 35.0 | Cluster_34483 |
| 696.58000 | 35.0 | Cluster_03211 |
| 697.31000 | 35.0 | Cluster_03215 |
| 697.37000 | 35.0 | Cluster_34515 |
| 697.85000 | 35.0 | Cluster_34520 |
| 697.87000 | 35.0 | Cluster_34523 |
| 698.82000 | 35.0 | Cluster_34541 |
| 698.83000 | 35.0 | Cluster_34546 |
| 699.70000 | 35.0 | Cluster_12244 |
| 699.85000 | 35.0 | Cluster_03235 |
| 700.26000 | 35.0 | Cluster_34580 |
| 700.36000 | 35.0 | Cluster_34589 |
| 700.65000 | 35.0 | Cluster_34590 |
| 700.85000 | 35.0 | Cluster_34594 |
| 701.36000 | 35.0 | Cluster_34614 |
| 702.34000 | 35.0 | Cluster_34653 |
| 702.62000 | 35.0 | Cluster_12286 |
| 703.04000 | 35.0 | Cluster_12291 |
| 703.05000 | 35.0 | Cluster_03266 |
| 703.33000 | 35.0 | Cluster_34677 |
| 703.69000 | 35.0 | Cluster_12310 |
| 703.91000 | 35.0 | Cluster_03274 |
| 704.17000 | 35.0 | Cluster_34698 |
| 704.21000 | 35.0 | Cluster_34700 |
| 704.31000 | 35.0 | Cluster_34706 |
| 704.81000 | 35.0 | Cluster_34728 |
| 704.85000 | 35.0 | Cluster_34734 |
| 705.04000 | 35.0 | Cluster_12319 |
| 705.37000 | 35.0 | Cluster_34742 |
| 705.83000 | 35.0 | Cluster_34756 |
| 705.88000 | 35.0 | Cluster_34757 |

| | | |
|---|---|---|
| 706.86000 | 35.0 | Cluster_03301 |
| 707.02000 | 35.0 | Cluster_12346 |
| 707.80000 | 35.0 | Cluster_34791 |
| 708.51000 | 35.0 | Cluster_00309 |
| 709.28000 | 35.0 | Cluster_34840 |
| 709.39000 | 35.0 | Cluster_34846 |
| 710.88000 | 35.0 | Cluster_03335 |
| 711.06000 | 35.0 | Cluster_12397 |
| 711.85000 | 35.0 | Cluster_34925 |
| 712.39000 | 35.0 | Cluster_34948 |
| 712.84000 | 35.0 | Cluster_34967 |
| 713.08000 | 35.0 | Cluster_03351 |
| 713.29000 | 35.0 | Cluster_03356 |
| 713.38000 | 35.0 | Cluster_12431 |
| 713.72000 | 35.0 | Cluster_12437 |
| 713.77000 | 35.0 | Cluster_34987 |
| 713.92000 | 35.0 | Cluster_34998 |
| 714.32000 | 35.0 | Cluster_35004 |
| 714.36000 | 35.0 | Cluster_35010 |
| 715.38000 | 35.0 | Cluster_35041 |
| 715.54000 | 35.0 | Cluster_00320 |
| 715.71000 | 35.0 | Cluster_12456 |
| 715.85000 | 35.0 | Cluster_35046 |
| 715.99000 | 35.0 | Cluster_12465 |
| 716.39000 | 35.0 | Cluster_35061 |
| 717.36000 | 35.0 | Cluster_12478 |
| 717.90000 | 35.0 | Cluster_35102 |
| 718.36000 | 35.0 | Cluster_35122 |
| 718.85000 | 35.0 | Cluster_35127 |
| 718.89000 | 35.0 | Cluster_35137 |
| 718.91000 | 35.0 | Cluster_35135 |
| 718.96000 | 35.0 | Cluster_12511 |
| 719.39000 | 35.0 | Cluster_35145 |
| 719.83000 | 35.0 | Cluster_35161 |
| 719.85000 | 35.0 | Cluster_03411 |
| 719.98000 | 35.0 | Cluster_12519 |
| 720.12000 | 35.0 | Cluster_35168 |
| 720.13000 | 35.0 | Cluster_35170 |
| 720.35000 | 35.0 | Cluster_03417 |
| 720.58000 | 35.0 | Cluster_03421 |
| 720.83000 | 35.0 | Cluster_35178 |
| 721.32000 | 35.0 | Cluster_35195 |
| 721.41000 | 35.0 | Cluster_35191 |
| 721.72000 | 35.0 | Cluster_12548 |
| 721.88000 | 35.0 | Cluster_35209 |
| 721.90000 | 35.0 | Cluster_35213 |
| 722.02000 | 35.0 | Cluster_12553 |
| 722.09000 | 35.0 | Cluster_03436 |
| 722.34000 | 35.0 | Cluster_03433 |
| 722.68000 | 35.0 | Cluster_12565 |
| 723.40000 | 35.0 | Cluster_12568 |

| | | |
|---|---|---|
| 723.42000 | 35.0 | Cluster_12570 |
| 723.95000 | 35.0 | Cluster_00326 |
| 724.04000 | 35.0 | Cluster_12575 |
| 724.90000 | 35.0 | Cluster_03465 |
| 725.06000 | 35.0 | Cluster_12593 |
| 725.32000 | 35.0 | Cluster_03469 |
| 725.41000 | 35.0 | Cluster_35286 |
| 725.42000 | 35.0 | Cluster_35293 |
| 727.38000 | 35.0 | Cluster_35317 |
| 728.10000 | 35.0 | Cluster_03487 |
| 728.13000 | 35.0 | Cluster_03484 |
| 728.42000 | 35.0 | Cluster_35338 |
| 728.98000 | 35.0 | Cluster_12653 |
| 729.15000 | 35.0 | Cluster_03497 |
| 729.84000 | 35.0 | Cluster_35373 |
| 729.89000 | 35.0 | Cluster_03507 |
| 730.91000 | 35.0 | Cluster_35408 |
| 731.10000 | 35.0 | Cluster_35410 |
| 731.83000 | 35.0 | Cluster_03518 |
| 732.11000 | 35.0 | Cluster_03522 |
| 732.64000 | 35.0 | Cluster_03534 |
| 733.03000 | 35.0 | Cluster_12708 |
| 733.13000 | 35.0 | Cluster_03548 |
| 733.83000 | 35.0 | Cluster_35471 |
| 733.85000 | 35.0 | Cluster_35474 |
| 734.58000 | 35.0 | Cluster_35493 |
| 735.04000 | 35.0 | Cluster_12729 |
| 735.74000 | 35.0 | Cluster_12741 |
| 736.37000 | 35.0 | Cluster_35541 |
| 737.04000 | 35.0 | Cluster_12754 |
| 737.16000 | 35.0 | Cluster_35560 |
| 737.82000 | 35.0 | Cluster_35593 |
| 738.39000 | 35.0 | Cluster_35611 |
| 738.63000 | 35.0 | Cluster_03590 |
| 739.38000 | 35.0 | Cluster_12791 |
| 739.39000 | 35.0 | Cluster_12789 |
| 739.84000 | 35.0 | Cluster_35645 |
| 740.37000 | 35.0 | Cluster_35657 |
| 740.38000 | 35.0 | Cluster_35661 |
| 740.93000 | 35.0 | Cluster_35676 |
| 741.85000 | 35.0 | Cluster_35683 |
| 742.37000 | 35.0 | Cluster_35697 |
| 742.67000 | 35.0 | Cluster_12831 |
| 742.84000 | 35.0 | Cluster_35707 |
| 744.37000 | 35.0 | Cluster_35750 |
| 745.38000 | 35.0 | Cluster_35777 |
| 745.68000 | 35.0 | Cluster_12857 |
| 746.06000 | 35.0 | Cluster_12863 |
| 746.15000 | 35.0 | Cluster_35792 |
| 746.30000 | 35.0 | Cluster_35797 |
| 747.12000 | 35.0 | Cluster_35817 |

| | | |
|---|---|---|
| 747.37000 | 35.0 | Cluster_35818 |
| 748.00000 | 35.0 | Cluster_12903 |
| 748.86000 | 35.0 | Cluster_03654 |
| 748.88000 | 35.0 | Cluster_35857 |
| 749.07000 | 35.0 | Cluster_12920 |
| 749.69000 | 35.0 | Cluster_12929 |
| 749.98000 | 35.0 | Cluster_12936 |
| 750.01000 | 35.0 | Cluster_12941 |
| 750.38000 | 35.0 | Cluster_35883 |
| 750.44000 | 35.0 | Cluster_35886 |
| 750.94000 | 35.0 | Cluster_35903 |
| 751.06000 | 35.0 | Cluster_12961 |
| 751.15000 | 35.0 | Cluster_03662 |
| 751.36000 | 35.0 | Cluster_35908 |
| 752.37000 | 35.0 | Cluster_35932 |
| 753.05000 | 35.0 | Cluster_12988 |
| 753.06000 | 35.0 | Cluster_12984 |
| 753.39000 | 35.0 | Cluster_35964 |
| 753.91000 | 35.0 | Cluster_35981 |
| 754.06000 | 35.0 | Cluster_13009 |
| 754.90000 | 35.0 | Cluster_36004 |
| 755.06000 | 35.0 | Cluster_13021 |
| 755.30000 | 35.0 | Cluster_13019 |
| 755.33000 | 35.0 | Cluster_13020 |
| 755.42000 | 35.0 | Cluster_36013 |
| 755.75000 | 35.0 | Cluster_13034 |
| 755.92000 | 35.0 | Cluster_36043 |
| 756.13000 | 35.0 | Cluster_03688 |
| 757.31000 | 35.0 | Cluster_36071 |
| 757.73000 | 35.0 | Cluster_13076 |
| 758.41000 | 35.0 | Cluster_36102 |
| 758.70000 | 35.0 | Cluster_13088 |
| 759.63000 | 35.0 | Cluster_03704 |
| 759.90000 | 35.0 | Cluster_36129 |
| 759.91000 | 35.0 | Cluster_36132 |
| 760.35000 | 35.0 | Cluster_36146 |
| 761.05000 | 35.0 | Cluster_00024 |
| 761.41000 | 35.0 | Cluster_36182 |
| 762.38000 | 35.0 | Cluster_13130 |
| 763.03000 | 35.0 | Cluster_13141 |
| 763.08000 | 35.0 | Cluster_13137 |
| 763.35000 | 35.0 | Cluster_03726 |
| 763.71000 | 35.0 | Cluster_13147 |
| 764.09000 | 35.0 | Cluster_13148 |
| 764.36000 | 35.0 | Cluster_36253 |
| 765.15000 | 35.0 | Cluster_03741 |
| 765.79000 | 35.0 | Cluster_36274 |
| 765.80000 | 35.0 | Cluster_36275 |
| 766.37000 | 35.0 | Cluster_36292 |
| 766.55000 | 35.0 | Cluster_00339 |
| 767.70000 | 35.0 | Cluster_13203 |

| | | |
|---|---|---|
| 767.85000 | 35.0 | Cluster_36325 |
| 768.07000 | 35.0 | Cluster_13209 |
| 768.08000 | 35.0 | Cluster_13205 |
| 768.38000 | 35.0 | Cluster_36351 |
| 769.33000 | 35.0 | Cluster_36370 |
| 769.59000 | 35.0 | Cluster_36368 |
| 769.82000 | 35.0 | Cluster_36385 |
| 772.41000 | 35.0 | Cluster_36441 |
| 772.74000 | 35.0 | Cluster_13265 |
| 772.80000 | 35.0 | Cluster_13273 |
| 772.88000 | 35.0 | Cluster_36459 |
| 772.92000 | 35.0 | Cluster_36460 |
| 773.08000 | 35.0 | Cluster_03785 |
| 773.09000 | 35.0 | Cluster_03786 |
| 773.13000 | 35.0 | Cluster_13272 |
| 773.35000 | 35.0 | Cluster_13283 |
| 773.41000 | 35.0 | Cluster_13286 |
| 774.12000 | 35.0 | Cluster_36481 |
| 774.73000 | 35.0 | Cluster_13306 |
| 775.06000 | 35.0 | Cluster_13312 |
| 775.07000 | 35.0 | Cluster_13311 |
| 775.42000 | 35.0 | Cluster_36510 |
| 776.37000 | 35.0 | Cluster_03803 |
| 777.15000 | 35.0 | Cluster_03806 |
| 777.35000 | 35.0 | Cluster_36561 |
| 777.38000 | 35.0 | Cluster_36567 |
| 777.39000 | 35.0 | Cluster_36564 |
| 778.37000 | 35.0 | Cluster_36591 |
| 778.77000 | 35.0 | Cluster_13350 |
| 778.87000 | 35.0 | Cluster_36605 |
| 780.73000 | 35.0 | Cluster_13379 |
| 780.81000 | 35.0 | Cluster_36636 |
| 780.88000 | 35.0 | Cluster_03821 |
| 781.40000 | 35.0 | Cluster_36661 |
| 782.03000 | 35.0 | Cluster_13403 |
| 783.13000 | 35.0 | Cluster_03834 |
| 783.46000 | 35.0 | Cluster_36708 |
| 783.72000 | 35.0 | Cluster_13422 |
| 784.06000 | 35.0 | Cluster_13433 |
| 784.89000 | 35.0 | Cluster_36733 |
| 784.93000 | 35.0 | Cluster_36732 |
| 785.45000 | 35.0 | Cluster_36744 |
| 785.83000 | 35.0 | Cluster_36763 |
| 787.06000 | 35.0 | Cluster_13459 |
| 787.12000 | 35.0 | Cluster_03868 |
| 787.35000 | 35.0 | Cluster_03870 |
| 788.87000 | 35.0 | Cluster_36840 |
| 789.06000 | 35.0 | Cluster_13483 |
| 790.87000 | 35.0 | Cluster_36906 |
| 791.12000 | 35.0 | Cluster_36909 |
| 791.38000 | 35.0 | Cluster_36914 |

| | | |
|---|---|---|
| 791.90000 | 35.0 | Cluster_36919 |
| 792.38000 | 35.0 | Cluster_36933 |
| 792.64000 | 35.0 | Cluster_36941 |
| 792.86000 | 35.0 | Cluster_03914 |
| 792.87000 | 35.0 | Cluster_36948 |
| 792.97000 | 35.0 | Cluster_13541 |
| 793.78000 | 35.0 | Cluster_13547 |
| 794.36000 | 35.0 | Cluster_36984 |
| 794.37000 | 35.0 | Cluster_36982 |
| 794.45000 | 35.0 | Cluster_36985 |
| 794.65000 | 35.0 | Cluster_36986 |
| 795.07000 | 35.0 | Cluster_13571 |
| 795.42000 | 35.0 | Cluster_37004 |
| 795.44000 | 35.0 | Cluster_37000 |
| 795.68000 | 35.0 | Cluster_13585 |
| 795.87000 | 35.0 | Cluster_03947 |
| 795.90000 | 35.0 | Cluster_37017 |
| 795.96000 | 35.0 | Cluster_37020 |
| 796.41000 | 35.0 | Cluster_13605 |
| 796.72000 | 35.0 | Cluster_13597 |
| 796.91000 | 35.0 | Cluster_37044 |
| 797.86000 | 35.0 | Cluster_37077 |
| 797.87000 | 35.0 | Cluster_37073 |
| 797.88000 | 35.0 | Cluster_37072 |
| 797.89000 | 35.0 | Cluster_37068 |
| 799.38000 | 35.0 | Cluster_37115 |
| 799.40000 | 35.0 | Cluster_37112 |
| 799.89000 | 35.0 | Cluster_03975 |
| 800.35000 | 35.0 | Cluster_37146 |
| 800.41000 | 35.0 | Cluster_13643 |
| 800.84000 | 35.0 | Cluster_37149 |
| 800.86000 | 35.0 | Cluster_03991 |
| 801.06000 | 35.0 | Cluster_00029 |
| 801.37000 | 35.0 | Cluster_03995 |
| 801.42000 | 35.0 | Cluster_37163 |
| 801.44000 | 35.0 | Cluster_13660 |
| 802.43000 | 35.0 | Cluster_37188 |
| 802.74000 | 35.0 | Cluster_13681 |
| 803.41000 | 35.0 | Cluster_13684 |
| 803.88000 | 35.0 | Cluster_37226 |
| 804.43000 | 35.0 | Cluster_37243 |
| 804.91000 | 35.0 | Cluster_04015 |
| 805.16000 | 35.0 | Cluster_37269 |
| 805.39000 | 35.0 | Cluster_37283 |
| 805.94000 | 35.0 | Cluster_37284 |
| 806.39000 | 35.0 | Cluster_37304 |
| 807.01000 | 35.0 | Cluster_13723 |
| 807.46000 | 35.0 | Cluster_37330 |
| 807.92000 | 35.0 | Cluster_37352 |
| 808.85000 | 35.0 | Cluster_37370 |
| 808.91000 | 35.0 | Cluster_37377 |

| | | |
|---|---|---|
| 809.40000 | 35.0 | Cluster_04037 |
| 809.91000 | 35.0 | Cluster_37401 |
| 810.39000 | 35.0 | Cluster_37407 |
| 810.44000 | 35.0 | Cluster_13774 |
| 811.74000 | 35.0 | Cluster_00036 |
| 811.86000 | 35.0 | Cluster_37446 |
| 811.91000 | 35.0 | Cluster_37441 |
| 812.39000 | 35.0 | Cluster_37458 |
| 812.41000 | 35.0 | Cluster_13796 |
| 813.82000 | 35.0 | Cluster_00352 |
| 813.90000 | 35.0 | Cluster_37483 |
| 814.10000 | 35.0 | Cluster_13827 |
| 814.16000 | 35.0 | Cluster_37492 |
| 814.17000 | 35.0 | Cluster_04067 |
| 814.90000 | 35.0 | Cluster_04074 |
| 815.06000 | 35.0 | Cluster_13832 |
| 815.79000 | 35.0 | Cluster_13844 |
| 815.92000 | 35.0 | Cluster_37530 |
| 816.38000 | 35.0 | Cluster_37540 |
| 816.43000 | 35.0 | Cluster_04088 |
| 816.72000 | 35.0 | Cluster_13857 |
| 817.34000 | 35.0 | Cluster_37561 |
| 817.88000 | 35.0 | Cluster_37570 |
| 818.42000 | 35.0 | Cluster_13883 |
| 819.05000 | 35.0 | Cluster_13897 |
| 819.42000 | 35.0 | Cluster_13906 |
| 820.17000 | 35.0 | Cluster_04104 |
| 821.23000 | 35.0 | Cluster_04111 |
| 821.38000 | 35.0 | Cluster_37637 |
| 822.06000 | 35.0 | Cluster_13934 |
| 823.42000 | 35.0 | Cluster_37669 |
| 823.91000 | 35.0 | Cluster_37674 |
| 823.94000 | 35.0 | Cluster_37688 |
| 825.32000 | 35.0 | Cluster_37717 |
| 825.39000 | 35.0 | Cluster_04137 |
| 826.38000 | 35.0 | Cluster_13981 |
| 826.86000 | 35.0 | Cluster_37765 |
| 827.04000 | 35.0 | Cluster_13986 |
| 827.06000 | 35.0 | Cluster_13988 |
| 827.09000 | 35.0 | Cluster_13989 |
| 827.88000 | 35.0 | Cluster_37794 |
| 828.80000 | 35.0 | Cluster_14000 |
| 828.98000 | 35.0 | Cluster_14004 |
| 829.16000 | 35.0 | Cluster_04165 |
| 829.36000 | 35.0 | Cluster_37821 |
| 829.43000 | 35.0 | Cluster_37825 |
| 829.91000 | 35.0 | Cluster_04169 |
| 830.74000 | 35.0 | Cluster_14020 |
| 830.89000 | 35.0 | Cluster_00042 |
| 831.45000 | 35.0 | Cluster_37853 |
| 832.41000 | 35.0 | Cluster_37873 |

| | | |
|---|---|---|
| 832.68000 | 35.0 | Cluster_37879 |
| 833.49000 | 35.0 | Cluster_37894 |
| 833.99000 | 35.0 | Cluster_37902 |
| 834.10000 | 35.0 | Cluster_14054 |
| 834.94000 | 35.0 | Cluster_37918 |
| 835.05000 | 35.0 | Cluster_14062 |
| 836.39000 | 35.0 | Cluster_14076 |
| 836.42000 | 35.0 | Cluster_14077 |
| 836.76000 | 35.0 | Cluster_14083 |
| 836.86000 | 35.0 | Cluster_37959 |
| 836.94000 | 35.0 | Cluster_04234 |
| 837.41000 | 35.0 | Cluster_37970 |
| 837.95000 | 35.0 | Cluster_37976 |
| 839.10000 | 35.0 | Cluster_14102 |
| 840.66000 | 35.0 | Cluster_04260 |
| 840.78000 | 35.0 | Cluster_14117 |
| 841.08000 | 35.0 | Cluster_14128 |
| 841.10000 | 35.0 | Cluster_14126 |
| 841.67000 | 35.0 | Cluster_00364 |
| 841.76000 | 35.0 | Cluster_38066 |
| 842.73000 | 35.0 | Cluster_14140 |
| 843.87000 | 35.0 | Cluster_38114 |
| 844.18000 | 35.0 | Cluster_04298 |
| 844.44000 | 35.0 | Cluster_14157 |
| 846.98000 | 35.0 | Cluster_38180 |
| 847.48000 | 35.0 | Cluster_38200 |
| 847.75000 | 35.0 | Cluster_14185 |
| 847.88000 | 35.0 | Cluster_38203 |
| 848.40000 | 35.0 | Cluster_04350 |
| 848.91000 | 35.0 | Cluster_04352 |
| 849.46000 | 35.0 | Cluster_38235 |
| 849.64000 | 35.0 | Cluster_04364 |
| 849.97000 | 35.0 | Cluster_38245 |
| 849.99000 | 35.0 | Cluster_38248 |
| 850.33000 | 35.0 | Cluster_38250 |
| 850.40000 | 35.0 | Cluster_04376 |
| 850.76000 | 35.0 | Cluster_14213 |
| 851.36000 | 35.0 | Cluster_14215 |
| 851.77000 | 35.0 | Cluster_38298 |
| 852.03000 | 35.0 | Cluster_38307 |
| 852.91000 | 35.0 | Cluster_04400 |
| 852.93000 | 35.0 | Cluster_04401 |
| 853.45000 | 35.0 | Cluster_38334 |
| 854.44000 | 35.0 | Cluster_38377 |
| 854.47000 | 35.0 | Cluster_38375 |
| 854.88000 | 35.0 | Cluster_38385 |
| 855.41000 | 35.0 | Cluster_38392 |
| 855.73000 | 35.0 | Cluster_14240 |
| 856.43000 | 35.0 | Cluster_14249 |
| 856.91000 | 35.0 | Cluster_04444 |
| 857.06000 | 35.0 | Cluster_14257 |

| | | |
|---|---|---|
| 857.89000 | 35.0 | Cluster_38459 |
| 857.91000 | 35.0 | Cluster_04449 |
| 858.07000 | 35.0 | Cluster_14269 |
| 858.77000 | 35.0 | Cluster_14275 |
| 859.45000 | 35.0 | Cluster_38500 |
| 859.47000 | 35.0 | Cluster_38497 |
| 860.47000 | 35.0 | Cluster_14284 |
| 861.39000 | 35.0 | Cluster_04493 |
| 862.39000 | 35.0 | Cluster_04505 |
| 862.49000 | 35.0 | Cluster_38591 |
| 863.05000 | 35.0 | Cluster_14302 |
| 863.76000 | 35.0 | Cluster_14307 |
| 863.94000 | 35.0 | Cluster_38615 |
| 864.07000 | 35.0 | Cluster_14314 |
| 865.03000 | 35.0 | Cluster_38646 |
| 865.75000 | 35.0 | Cluster_14326 |
| 865.92000 | 35.0 | Cluster_38659 |
| 865.93000 | 35.0 | Cluster_04555 |
| 866.41000 | 35.0 | Cluster_14334 |
| 869.07000 | 35.0 | Cluster_14353 |
| 869.28000 | 35.0 | Cluster_38719 |
| 869.41000 | 35.0 | Cluster_04599 |
| 869.78000 | 35.0 | Cluster_14363 |
| 871.17000 | 35.0 | Cluster_38758 |
| 871.27000 | 35.0 | Cluster_38759 |
| 871.44000 | 35.0 | Cluster_38769 |
| 871.47000 | 35.0 | Cluster_14388 |
| 872.27000 | 35.0 | Cluster_38802 |
| 874.09000 | 35.0 | Cluster_14414 |
| 874.26000 | 35.0 | Cluster_38844 |
| 874.78000 | 35.0 | Cluster_14424 |
| 875.28000 | 35.0 | Cluster_38860 |
| 875.66000 | 35.0 | Cluster_04698 |
| 876.08000 | 35.0 | Cluster_14437 |
| 876.53000 | 35.0 | Cluster_38895 |
| 877.38000 | 35.0 | Cluster_14453 |
| 877.44000 | 35.0 | Cluster_38909 |
| 878.48000 | 35.0 | Cluster_38928 |
| 880.46000 | 35.0 | Cluster_14483 |
| 881.96000 | 35.0 | Cluster_38993 |
| 882.44000 | 35.0 | Cluster_39008 |
| 882.94000 | 35.0 | Cluster_39018 |
| 884.40000 | 35.0 | Cluster_14528 |
| 884.66000 | 35.0 | Cluster_39047 |
| 885.12000 | 35.0 | Cluster_14544 |
| 885.68000 | 35.0 | Cluster_04765 |
| 885.95000 | 35.0 | Cluster_39078 |
| 886.43000 | 35.0 | Cluster_39095 |
| 886.96000 | 35.0 | Cluster_39099 |
| 887.52000 | 35.0 | Cluster_39107 |
| 887.68000 | 35.0 | Cluster_04781 |

| | | |
|---|---|---|
| 887.74000 | 35.0 | Cluster_14571 |
| 888.13000 | 35.0 | Cluster_14577 |
| 888.40000 | 35.0 | Cluster_14578 |
| 888.91000 | 35.0 | Cluster_39135 |
| 888.92000 | 35.0 | Cluster_04786 |
| 889.11000 | 35.0 | Cluster_14581 |
| 889.44000 | 35.0 | Cluster_39147 |
| 890.17000 | 35.0 | Cluster_04796 |
| 890.38000 | 35.0 | Cluster_39161 |
| 890.39000 | 35.0 | Cluster_39160 |
| 890.46000 | 35.0 | Cluster_39173 |
| 890.99000 | 35.0 | Cluster_39180 |
| 891.46000 | 35.0 | Cluster_39186 |
| 891.81000 | 35.0 | Cluster_14598 |
| 892.21000 | 35.0 | Cluster_04824 |
| 893.43000 | 35.0 | Cluster_39231 |
| 893.78000 | 35.0 | Cluster_14615 |
| 894.12000 | 35.0 | Cluster_14617 |
| 894.71000 | 35.0 | Cluster_04845 |
| 895.21000 | 35.0 | Cluster_04851 |
| 895.45000 | 35.0 | Cluster_14622 |
| 895.81000 | 35.0 | Cluster_14625 |
| 896.15000 | 35.0 | Cluster_14634 |
| 897.04000 | 35.0 | Cluster_39292 |
| 898.77000 | 35.0 | Cluster_14664 |
| 899.43000 | 35.0 | Cluster_39337 |
| 899.51000 | 35.0 | Cluster_39345 |
| 899.96000 | 35.0 | Cluster_04891 |
| 900.94000 | 35.0 | Cluster_39368 |
| 900.97000 | 35.0 | Cluster_39380 |
| 900.99000 | 35.0 | Cluster_39377 |
| 902.46000 | 35.0 | Cluster_39412 |
| 904.46000 | 35.0 | Cluster_39443 |
| 905.43000 | 35.0 | Cluster_39461 |
| 906.77000 | 35.0 | Cluster_14739 |
| 907.80000 | 35.0 | Cluster_14749 |
| 908.46000 | 35.0 | Cluster_39500 |
| 908.48000 | 35.0 | Cluster_14756 |
| 908.94000 | 35.0 | Cluster_04958 |
| 908.95000 | 35.0 | Cluster_39504 |
| 909.44000 | 35.0 | Cluster_14760 |
| 909.50000 | 35.0 | Cluster_14765 |
| 909.89000 | 35.0 | Cluster_39520 |
| 910.11000 | 35.0 | Cluster_14769 |
| 910.19000 | 35.0 | Cluster_04966 |
| 911.96000 | 35.0 | Cluster_39564 |
| 914.56000 | 35.0 | Cluster_39615 |
| 915.13000 | 35.0 | Cluster_14823 |
| 915.93000 | 35.0 | Cluster_39626 |
| 916.44000 | 35.0 | Cluster_39640 |
| 916.49000 | 35.0 | Cluster_39633 |

| | | |
|---|---|---|
| 917.45000 | 35.0 | Cluster_14854 |
| 917.97000 | 35.0 | Cluster_39674 |
| 918.66000 | 35.0 | Cluster_39698 |
| 920.45000 | 35.0 | Cluster_39721 |
| 920.76000 | 35.0 | Cluster_14891 |
| 920.79000 | 35.0 | Cluster_39725 |
| 920.88000 | 35.0 | Cluster_39726 |
| 920.95000 | 35.0 | Cluster_39731 |
| 922.42000 | 35.0 | Cluster_14907 |
| 922.98000 | 35.0 | Cluster_39772 |
| 923.30000 | 35.0 | Cluster_14924 |
| 923.45000 | 35.0 | Cluster_14932 |
| 923.92000 | 35.0 | Cluster_39782 |
| 924.49000 | 35.0 | Cluster_39788 |
| 925.11000 | 35.0 | Cluster_14961 |
| 926.42000 | 35.0 | Cluster_14975 |
| 927.48000 | 35.0 | Cluster_39834 |
| 928.47000 | 35.0 | Cluster_39858 |
| 928.96000 | 35.0 | Cluster_39861 |
| 929.12000 | 35.0 | Cluster_15009 |
| 929.43000 | 35.0 | Cluster_39880 |
| 929.90000 | 35.0 | Cluster_39897 |
| 930.47000 | 35.0 | Cluster_15020 |
| 931.73000 | 35.0 | Cluster_15043 |
| 931.90000 | 35.0 | Cluster_39924 |
| 932.64000 | 35.0 | Cluster_15062 |
| 932.95000 | 35.0 | Cluster_39941 |
| 933.14000 | 35.0 | Cluster_15072 |
| 933.33000 | 35.0 | Cluster_05038 |
| 934.00000 | 35.0 | Cluster_39963 |
| 934.13000 | 35.0 | Cluster_15084 |
| 934.43000 | 35.0 | Cluster_15083 |
| 934.78000 | 35.0 | Cluster_15089 |
| 935.78000 | 35.0 | Cluster_15100 |
| 936.45000 | 35.0 | Cluster_15116 |
| 936.80000 | 35.0 | Cluster_15129 |
| 937.00000 | 35.0 | Cluster_39997 |
| 937.44000 | 35.0 | Cluster_15144 |
| 937.49000 | 35.0 | Cluster_05051 |
| 938.07000 | 35.0 | Cluster_15154 |
| 938.13000 | 35.0 | Cluster_15156 |
| 938.21000 | 35.0 | Cluster_15160 |
| 939.69000 | 35.0 | Cluster_05058 |
| 940.46000 | 35.0 | Cluster_15180 |
| 941.72000 | 35.0 | Cluster_40098 |
| 942.00000 | 35.0 | Cluster_40094 |
| 943.45000 | 35.0 | Cluster_15229 |
| 945.13000 | 35.0 | Cluster_15244 |
| 945.39000 | 35.0 | Cluster_15242 |
| 945.72000 | 35.0 | Cluster_05075 |
| 945.73000 | 35.0 | Cluster_05078 |

| | | |
|---|---|---|
| 945.78000 | 35.0 | Cluster_15250 |
| 945.98000 | 35.0 | Cluster_40152 |
| 947.16000 | 35.0 | Cluster_15264 |
| 947.22000 | 35.0 | Cluster_05084 |
| 947.79000 | 35.0 | Cluster_15273 |
| 948.19000 | 35.0 | Cluster_15277 |
| 948.52000 | 35.0 | Cluster_40180 |
| 949.14000 | 35.0 | Cluster_15301 |
| 949.44000 | 35.0 | Cluster_40190 |
| 949.46000 | 35.0 | Cluster_15302 |
| 950.44000 | 35.0 | Cluster_15314 |
| 951.17000 | 35.0 | Cluster_15324 |
| 951.96000 | 35.0 | Cluster_40227 |
| 952.14000 | 35.0 | Cluster_15337 |
| 952.15000 | 35.0 | Cluster_15339 |
| 952.44000 | 35.0 | Cluster_15342 |
| 952.53000 | 35.0 | Cluster_40233 |
| 953.98000 | 35.0 | Cluster_40255 |
| 954.82000 | 35.0 | Cluster_15379 |
| 954.99000 | 35.0 | Cluster_40262 |
| 955.13000 | 35.0 | Cluster_15394 |
| 955.47000 | 35.0 | Cluster_15404 |
| 956.48000 | 35.0 | Cluster_15426 |
| 957.07000 | 35.0 | Cluster_40290 |
| 957.48000 | 35.0 | Cluster_40291 |
| 960.15000 | 35.0 | Cluster_15462 |
| 960.98000 | 35.0 | Cluster_40317 |
| 961.48000 | 35.0 | Cluster_15478 |
| 962.14000 | 35.0 | Cluster_15497 |
| 962.38000 | 35.0 | Cluster_40344 |
| 962.79000 | 35.0 | Cluster_15505 |
| 964.13000 | 35.0 | Cluster_15523 |
| 964.43000 | 35.0 | Cluster_40365 |
| 965.29000 | 35.0 | Cluster_40377 |
| 966.15000 | 35.0 | Cluster_15549 |
| 966.49000 | 35.0 | Cluster_40387 |
| 966.50000 | 35.0 | Cluster_05172 |
| 966.85000 | 35.0 | Cluster_15552 |
| 967.00000 | 35.0 | Cluster_40392 |
| 967.56000 | 35.0 | Cluster_15565 |
| 968.03000 | 35.0 | Cluster_40402 |
| 968.25000 | 35.0 | Cluster_05187 |
| 971.97000 | 35.0 | Cluster_40451 |
| 972.01000 | 35.0 | Cluster_40467 |
| 972.46000 | 35.0 | Cluster_15627 |
| 972.49000 | 35.0 | Cluster_15626 |
| 973.51000 | 35.0 | Cluster_40476 |
| 974.47000 | 35.0 | Cluster_15651 |
| 974.98000 | 35.0 | Cluster_40495 |
| 975.82000 | 35.0 | Cluster_15672 |
| 975.98000 | 35.0 | Cluster_05210 |

| | | |
|---|---|---|
| 976.00000 | 35.0 | Cluster_05212 |
| 976.03000 | 35.0 | Cluster_40504 |
| 977.14000 | 35.0 | Cluster_15696 |
| 977.62000 | 35.0 | Cluster_40522 |
| 979.45000 | 35.0 | Cluster_40546 |
| 979.80000 | 35.0 | Cluster_15720 |
| 979.83000 | 35.0 | Cluster_15721 |
| 979.98000 | 35.0 | Cluster_40568 |
| 980.00000 | 35.0 | Cluster_40562 |
| 980.50000 | 35.0 | Cluster_40565 |
| 980.70000 | 35.0 | Cluster_40578 |
| 981.45000 | 35.0 | Cluster_15750 |
| 983.25000 | 35.0 | Cluster_05237 |
| 983.95000 | 35.0 | Cluster_40617 |
| 985.14000 | 35.0 | Cluster_40628 |
| 986.02000 | 35.0 | Cluster_40630 |
| 986.52000 | 35.0 | Cluster_40639 |
| 986.83000 | 35.0 | Cluster_15810 |
| 987.24000 | 35.0 | Cluster_40654 |
| 987.80000 | 35.0 | Cluster_15825 |
| 988.44000 | 35.0 | Cluster_15835 |
| 991.93000 | 35.0 | Cluster_40717 |
| 993.50000 | 35.0 | Cluster_15895 |
| 997.19000 | 35.0 | Cluster_15954 |
| 1000.06000 | 35.0 | Cluster_40851 |
| 1001.03000 | 35.0 | Cluster_40878 |
| 1001.16000 | 35.0 | Cluster_15996 |
| 1001.49000 | 35.0 | Cluster_40884 |
| 1003.52000 | 35.0 | Cluster_05294 |
| 1004.00000 | 35.0 | Cluster_40937 |
| 1004.31000 | 35.0 | Cluster_05296 |
| 1004.57000 | 35.0 | Cluster_40940 |
| 1007.81000 | 35.0 | Cluster_16072 |
| 1008.78000 | 35.0 | Cluster_16080 |
| 1012.26000 | 35.0 | Cluster_05329 |
| 1012.27000 | 35.0 | Cluster_41044 |
| 1012.93000 | 35.0 | Cluster_41054 |
| 1013.01000 | 35.0 | Cluster_05341 |
| 1013.47000 | 35.0 | Cluster_16138 |
| 1013.49000 | 35.0 | Cluster_16137 |
| 1013.76000 | 35.0 | Cluster_05344 |
| 1013.90000 | 35.0 | Cluster_41056 |
| 1015.92000 | 35.0 | Cluster_41088 |
| 1016.78000 | 35.0 | Cluster_05351 |
| 1017.27000 | 35.0 | Cluster_41099 |
| 1019.25000 | 35.0 | Cluster_05360 |
| 1020.15000 | 35.0 | Cluster_16210 |
| 1022.50000 | 35.0 | Cluster_16231 |
| 1023.82000 | 35.0 | Cluster_41185 |
| 1024.48000 | 35.0 | Cluster_16254 |
| 1024.50000 | 35.0 | Cluster_16256 |

| | | |
|---|---|---|
| 1024.93000 | 35.0 | Cluster_41195 |
| 1025.46000 | 35.0 | Cluster_16264 |
| 1025.81000 | 35.0 | Cluster_41211 |
| 1026.83000 | 35.0 | Cluster_16274 |
| 1027.13000 | 35.0 | Cluster_16285 |
| 1027.14000 | 35.0 | Cluster_16284 |
| 1027.88000 | 35.0 | Cluster_16291 |
| 1028.18000 | 35.0 | Cluster_16295 |
| 1028.84000 | 35.0 | Cluster_16300 |
| 1029.99000 | 35.0 | Cluster_41262 |
| 1031.20000 | 35.0 | Cluster_16323 |
| 1032.82000 | 35.0 | Cluster_16335 |
| 1033.20000 | 35.0 | Cluster_16342 |
| 1036.52000 | 35.0 | Cluster_16382 |
| 1037.50000 | 35.0 | Cluster_16400 |
| 1039.05000 | 35.0 | Cluster_41371 |
| 1039.52000 | 35.0 | Cluster_41374 |
| 1039.81000 | 35.0 | Cluster_16429 |
| 1040.83000 | 35.0 | Cluster_05437 |
| 1041.01000 | 35.0 | Cluster_41401 |
| 1042.17000 | 35.0 | Cluster_16457 |
| 1044.49000 | 35.0 | Cluster_41432 |
| 1046.13000 | 35.0 | Cluster_16488 |
| 1046.22000 | 35.0 | Cluster_16495 |
| 1047.16000 | 35.0 | Cluster_16507 |
| 1047.83000 | 35.0 | Cluster_41470 |
| 1053.50000 | 35.0 | Cluster_16581 |
| 1053.98000 | 35.0 | Cluster_41534 |
| 1054.99000 | 35.0 | Cluster_05476 |
| 1055.22000 | 35.0 | Cluster_16595 |
| 1055.50000 | 35.0 | Cluster_16606 |
| 1056.52000 | 35.0 | Cluster_16616 |
| 1057.06000 | 35.0 | Cluster_41567 |
| 1057.33000 | 35.0 | Cluster_05486 |
| 1057.34000 | 35.0 | Cluster_05487 |
| 1057.58000 | 35.0 | Cluster_41574 |
| 1057.99000 | 35.0 | Cluster_41578 |
| 1058.02000 | 35.0 | Cluster_41582 |
| 1058.81000 | 35.0 | Cluster_16658 |
| 1058.85000 | 35.0 | Cluster_16649 |
| 1058.96000 | 35.0 | Cluster_41595 |
| 1058.97000 | 35.0 | Cluster_41597 |
| 1059.01000 | 35.0 | Cluster_41596 |
| 1060.32000 | 35.0 | Cluster_41612 |
| 1060.75000 | 35.0 | Cluster_05502 |
| 1060.82000 | 35.0 | Cluster_16679 |
| 1061.26000 | 35.0 | Cluster_05510 |
| 1061.33000 | 35.0 | Cluster_05506 |
| 1061.82000 | 35.0 | Cluster_41625 |
| 1062.83000 | 35.0 | Cluster_16700 |
| 1063.27000 | 35.0 | Cluster_05516 |

| | | |
|---|---|---|
| 1063.82000 | 35.0 | Cluster_41655 |
| 1063.99000 | 35.0 | Cluster_41663 |
| 1064.72000 | 35.0 | Cluster_05521 |
| 1064.73000 | 35.0 | Cluster_41675 |
| 1064.99000 | 35.0 | Cluster_41673 |
| 1066.26000 | 35.0 | Cluster_41702 |
| 1067.52000 | 35.0 | Cluster_41715 |
| 1068.05000 | 35.0 | Cluster_41719 |
| 1068.75000 | 35.0 | Cluster_41734 |
| 1069.51000 | 35.0 | Cluster_41748 |
| 1070.16000 | 35.0 | Cluster_16781 |
| 1070.51000 | 35.0 | Cluster_05546 |
| 1072.55000 | 35.0 | Cluster_16804 |
| 1072.76000 | 35.0 | Cluster_05555 |
| 1074.19000 | 35.0 | Cluster_16831 |
| 1074.51000 | 35.0 | Cluster_41810 |
| 1075.53000 | 35.0 | Cluster_41816 |
| 1076.75000 | 35.0 | Cluster_05574 |
| 1078.01000 | 35.0 | Cluster_05578 |
| 1079.49000 | 35.0 | Cluster_05582 |
| 1080.75000 | 35.0 | Cluster_00593 |
| 1082.00000 | 35.0 | Cluster_05588 |
| 1083.50000 | 35.0 | Cluster_05592 |
| 1086.59000 | 35.0 | Cluster_41940 |
| 1091.58000 | 35.0 | Cluster_41998 |
| 1092.90000 | 35.0 | Cluster_17025 |
| 1093.21000 | 35.0 | Cluster_17031 |
| 1093.55000 | 35.0 | Cluster_17035 |
| 1093.58000 | 35.0 | Cluster_42024 |
| 1093.99000 | 35.0 | Cluster_42029 |
| 1094.92000 | 35.0 | Cluster_00625 |
| 1096.55000 | 35.0 | Cluster_42051 |
| 1100.23000 | 35.0 | Cluster_17091 |
| 1100.56000 | 35.0 | Cluster_17097 |
| 1101.96000 | 35.0 | Cluster_17105 |
| 1102.54000 | 35.0 | Cluster_17111 |
| 1103.12000 | 35.0 | Cluster_42104 |
| 1108.23000 | 35.0 | Cluster_17176 |
| 1108.37000 | 35.0 | Cluster_42148 |
| 1108.88000 | 35.0 | Cluster_17175 |
| 1109.86000 | 35.0 | Cluster_17199 |
| 1113.50000 | 35.0 | Cluster_42184 |
| 1113.53000 | 35.0 | Cluster_17241 |
| 1115.18000 | 35.0 | Cluster_17258 |
| 1115.77000 | 35.0 | Cluster_00660 |
| 1117.22000 | 35.0 | Cluster_17276 |
| 1118.21000 | 35.0 | Cluster_17277 |
| 1118.22000 | 35.0 | Cluster_17280 |
| 1119.87000 | 35.0 | Cluster_42234 |
| 1121.54000 | 35.0 | Cluster_17322 |
| 1123.04000 | 35.0 | Cluster_42270 |

| | | |
|---|---|---|
| 1123.97000 | 35.0 | Cluster_42271 |
| 1124.25000 | 35.0 | Cluster_17359 |
| 1125.57000 | 35.0 | Cluster_42283 |
| 1126.07000 | 35.0 | Cluster_42291 |
| 1127.88000 | 35.0 | Cluster_17398 |
| 1128.57000 | 35.0 | Cluster_17409 |
| 1129.21000 | 35.0 | Cluster_17418 |
| 1129.55000 | 35.0 | Cluster_17428 |
| 1129.90000 | 35.0 | Cluster_17433 |
| 1130.23000 | 35.0 | Cluster_17434 |
| 1131.58000 | 35.0 | Cluster_17448 |
| 1133.13000 | 35.0 | Cluster_42337 |
| 1133.20000 | 35.0 | Cluster_17479 |
| 1133.22000 | 35.0 | Cluster_17484 |
| 1134.90000 | 35.0 | Cluster_17503 |
| 1135.23000 | 35.0 | Cluster_17509 |
| 1135.51000 | 35.0 | Cluster_17513 |
| 1135.54000 | 35.0 | Cluster_17519 |
| 1135.55000 | 35.0 | Cluster_17517 |
| 1136.11000 | 35.0 | Cluster_42363 |
| 1136.88000 | 35.0 | Cluster_17525 |
| 1136.89000 | 35.0 | Cluster_17538 |
| 1138.13000 | 35.0 | Cluster_42377 |
| 1140.22000 | 35.0 | Cluster_17567 |
| 1143.18000 | 35.0 | Cluster_00674 |
| 1143.56000 | 35.0 | Cluster_17607 |
| 1143.60000 | 35.0 | Cluster_42409 |
| 1144.20000 | 35.0 | Cluster_17626 |
| 1144.24000 | 35.0 | Cluster_17620 |
| 1145.21000 | 35.0 | Cluster_17645 |
| 1146.57000 | 35.0 | Cluster_00680 |
| 1149.29000 | 35.0 | Cluster_42461 |
| 1151.05000 | 35.0 | Cluster_42474 |
| 1152.54000 | 35.0 | Cluster_17735 |
| 1152.88000 | 35.0 | Cluster_17750 |
| 1153.03000 | 35.0 | Cluster_05790 |
| 1153.76000 | 35.0 | Cluster_00695 |
| 1154.35000 | 35.0 | Cluster_42485 |
| 1155.34000 | 35.0 | Cluster_42501 |
| 1156.84000 | 35.0 | Cluster_42511 |
| 1158.84000 | 35.0 | Cluster_42519 |
| 1159.61000 | 35.0 | Cluster_42529 |
| 1160.34000 | 35.0 | Cluster_42534 |
| 1162.11000 | 35.0 | Cluster_42551 |
| 1162.64000 | 35.0 | Cluster_42555 |
| 1163.09000 | 35.0 | Cluster_05817 |
| 1166.08000 | 35.0 | Cluster_05827 |
| 1166.14000 | 35.0 | Cluster_42592 |
| 1166.21000 | 35.0 | Cluster_17894 |
| 1167.19000 | 35.0 | Cluster_17908 |
| 1167.57000 | 35.0 | Cluster_17921 |

| | | |
|---|---|---|
| 1168.21000 | 35.0 | Cluster_17931 |
| 1168.33000 | 35.0 | Cluster_42608 |
| 1168.58000 | 35.0 | Cluster_05832 |
| 1173.07000 | 35.0 | Cluster_42645 |
| 1180.80000 | 35.0 | Cluster_05888 |
| 1183.23000 | 35.0 | Cluster_18040 |
| 1187.59000 | 35.0 | Cluster_18075 |
| 1190.56000 | 35.0 | Cluster_18091 |
| 1192.81000 | 35.0 | Cluster_42765 |
| 1192.92000 | 35.0 | Cluster_18117 |
| 1195.88000 | 35.0 | Cluster_18133 |
| 1196.63000 | 35.0 | Cluster_42787 |
| 1197.59000 | 35.0 | Cluster_18145 |
| 1197.90000 | 35.0 | Cluster_18140 |
| 1198.64000 | 35.0 | Cluster_42803 |
| 1199.32000 | 35.0 | Cluster_05982 |
| 1199.57000 | 35.0 | Cluster_18161 |
| 1200.26000 | 35.0 | Cluster_18168 |
| 1200.56000 | 35.0 | Cluster_18177 |
| 1201.89000 | 35.0 | Cluster_18192 |
| 1203.44000 | 35.0 | Cluster_42830 |
| 1206.62000 | 35.0 | Cluster_42853 |
| 1208.56000 | 35.0 | Cluster_18237 |
| 1209.51000 | 35.0 | Cluster_42887 |
| 1213.12000 | 35.0 | Cluster_42910 |
| 1215.15000 | 35.0 | Cluster_42924 |
| 1218.11000 | 35.0 | Cluster_42950 |
| 1219.09000 | 35.0 | Cluster_42956 |
| 1221.25000 | 35.0 | Cluster_18299 |
| 1222.10000 | 35.0 | Cluster_42968 |
| 1224.08000 | 35.0 | Cluster_42977 |
| 1224.60000 | 35.0 | Cluster_42983 |
| 1225.59000 | 35.0 | Cluster_42994 |
| 1228.07000 | 35.0 | Cluster_43008 |
| 1228.11000 | 35.0 | Cluster_43007 |
| 1230.63000 | 35.0 | Cluster_43026 |
| 1232.57000 | 35.0 | Cluster_43045 |
| 1234.58000 | 35.0 | Cluster_43065 |
| 1238.57000 | 35.0 | Cluster_43079 |
| 1245.28000 | 35.0 | Cluster_18390 |
| 1249.16000 | 35.0 | Cluster_43135 |
| 1251.87000 | 35.0 | Cluster_43149 |
| 1253.87000 | 35.0 | Cluster_43162 |
| 1255.86000 | 35.0 | Cluster_43176 |
| 1257.08000 | 35.0 | Cluster_06282 |
| 1264.27000 | 35.0 | Cluster_18463 |
| 1267.20000 | 35.0 | Cluster_43224 |
| 1267.68000 | 35.0 | Cluster_43228 |
| 1274.29000 | 35.0 | Cluster_18502 |
| 1284.15000 | 35.0 | Cluster_43287 |
| 1300.69000 | 35.0 | Cluster_43366 |

| | | |
|---|---|---|
| 1300.70000 | 35.0 | Cluster_43360 |
| 1300.73000 | 35.0 | Cluster_43361 |
| 1300.89000 | 35.0 | Cluster_43362 |
| 1301.68000 | 35.0 | Cluster_43365 |
| 1320.21000 | 35.0 | Cluster_43449 |
| 1329.73000 | 35.0 | Cluster_43484 |
| 1330.45000 | 35.0 | Cluster_06499 |
| 1331.69000 | 35.0 | Cluster_43492 |
| 1334.68000 | 35.0 | Cluster_43505 |
| 1349.89000 | 35.0 | Cluster_43562 |
| 1353.71000 | 35.0 | Cluster_43578 |
| 1354.21000 | 35.0 | Cluster_43579 |
| 1354.66000 | 35.0 | Cluster_06571 |
| 1361.03000 | 35.0 | Cluster_18681 |
| 1367.16000 | 35.0 | Cluster_43645 |
| 1372.75000 | 35.0 | Cluster_43667 |
| 1376.18000 | 35.0 | Cluster_43680 |
| 1376.34000 | 35.0 | Cluster_18707 |
| 1383.67000 | 35.0 | Cluster_18716 |
| 1394.65000 | 35.0 | Cluster_43739 |
| 1396.44000 | 35.0 | Cluster_18732 |
| 1402.12000 | 35.0 | Cluster_18741 |
| 1406.24000 | 35.0 | Cluster_43793 |
| 1407.03000 | 35.0 | Cluster_18752 |
| 1414.35000 | 35.0 | Cluster_18768 |
| 1417.37000 | 35.0 | Cluster_18772 |
| 1420.01000 | 35.0 | Cluster_18783 |
| 1422.96000 | 35.0 | Cluster_43843 |
| 1423.47000 | 35.0 | Cluster_43844 |
| 1424.69000 | 35.0 | Cluster_18789 |
| 1424.79000 | 35.0 | Cluster_43851 |
| 1425.27000 | 35.0 | Cluster_43853 |
| 1427.03000 | 35.0 | Cluster_18794 |
| 1428.82000 | 35.0 | Cluster_43862 |
| 1429.02000 | 35.0 | Cluster_18795 |
| 1430.01000 | 35.0 | Cluster_18796 |
| 1431.68000 | 35.0 | Cluster_18799 |
| 1437.03000 | 35.0 | Cluster_18803 |
| 1439.35000 | 35.0 | Cluster_18807 |
| 1459.27000 | 35.0 | Cluster_43959 |
| 1477.30000 | 35.0 | Cluster_44015 |
| 1489.22000 | 35.0 | Cluster_44043 |
| 1492.94000 | 35.0 | Cluster_44054 |
| 1495.14000 | 35.0 | Cluster_44065 |
| 1511.80000 | 35.0 | Cluster_44102 |
| 1518.72000 | 35.0 | Cluster_44112 |
| 1530.33000 | 35.0 | Cluster_44125 |
| 1560.76000 | 35.0 | Cluster_44175 |
| 1564.32000 | 35.0 | Cluster_44184 |
| 1571.76000 | 35.0 | Cluster_44194 |
| 1582.85000 | 35.0 | Cluster_44218 |

| | | |
|---|---|---|
| 1616.85000 | 35.0 | Cluster_44250 |
| 1633.97000 | 35.0 | Cluster_44257 |
| 1635.40000 | 35.0 | Cluster_44260 |
| 1676.82000 | 35.0 | Cluster_44320 |
| 1683.84000 | 35.0 | Cluster_44341 |
| 1687.92000 | 35.0 | Cluster_44353 |
| 1739.94000 | 35.0 | Cluster_44403 |
| 1792.99000 | 35.0 | Cluster_44425 |

Reject Mass List: (none)
Neutral Loss Mass List: (none)
Product Mass List: (none)
Neutral loss in top: 3
Product in top: 3
Most intense if no parent masses found enabled
Add/subtract mass not enabled
FT master scan preview mode enabled
Charge state screening enabled
Charge state dependent ETD time not enabled
Monoisotopic precursor selection enabled
Charge state rejection enabled
Unassigned charge states : rejected
Charge state 1 : rejected
Charge state 2 : not rejected
Charge state 3 : not rejected
Charge states 4+ : not rejected
Chromatography mode is disabled

Global Data Dependent Settings:
Predict ion injection time enabled
Use global parent and reject mass lists not enabled
Exclude parent mass from data dependent selection not enabled
Exclusion mass width by mass
Exclusion mass width low: 0.50000
Exclusion mass width high: 0.50000
Parent mass width by mass
Parent mass width low: 0.50000
Parent mass width high: 0.50000
Reject mass width by mass
Reject mass width low: 0.50000
Reject mass width high: 0.50000
Zoom/UltraZoom scan mass width by mass
Zoom/UltraZoom scan mass width low: 5.00
Zoom/UltraZoom scan mass width high: 5.00
FT SIM scan mass width low: 5.00
FT SIM scan mass width high: 5.00
Neutral Loss candidates processed by decreasing intensity
Neutral Loss mass width by mass
Neutral Loss mass width low: 0.50000
Neutral Loss mass width high: 0.50000
Product candidates processed by decreasing intensity

Product mass width by mass
Product mass width low: 0.50000
Product mass width high: 0.50000
MS mass range: 0.00-1000000.00
MSn mass range by mass
MSn mass range: 0.00-1000000.00
Use m/z values as masses not enabled
Analog UV data dep. not enabled
Dynamic exclusion not enabled
Isotopic data dependence not enabled
Mass Tags data dependence not enabled
Custom Data Dependent Settings:
Not enabled

---

Pass 9 (wheat-mixed-digests_MS2_TO-USE-inclusion-01_2.raw):

Creator: Orbi_30393
Last modified: 10/19/2021 by Orbi_30393
MS Run Time (min): 43.00
Sequence override of method parameters not enabled.
Divert Valve: not used during run
Contact Closure: not used during run
Syringe Pump: not used during run
MS Detector Settings:
Real-time modifications to method not enabled
Stepped collision energy not enabled
Additional Microscans:
MS2 0 0
MS3 0 0
MS4 0 0
MS5 0 0
MS6 0 0
MS7 0 0
MS8 0 0
MS9 0 0
MS10 0 0
Segment 1 Information
Duration (min): 43.00
Number of Scan Events: 2
Tune Method: Orbitrap-tune-file_2020-03-13_HESI
Scan Event Details:
1: FTMS + p norm res=15000 o(300.0-2000.0)
CV = 0.0V
2: ITMS + c norm Dep MS/MS Most intense ion from (1)
Activation Type: CID
Min. Signal Required: 500.0
Isolation Width: 1.00
Normalized Coll. Energy: 35.0
Default Charge State: 2
Activation Q: 0.250
Activation Time: 10.000

CV = 0.0V

Lock Masses:
Pos List Name: N/A
Source: API Source
Mass List: (none)
Neg List Name: N/A
Source: API Source
Mass List: (none)

Data Dependent Settings:
Use separate polarity settings disabled
Parent Mass List:

| MS Mass | MS FAIMS CV | MS Normalized Collision Energy | MS2 Mass | MS2 Normalized Collision Energy | Name |
|---|---|---|---|---|---|
| 300.13000 | | 35.0 | | | Cluster_06844 |
| 300.14000 | | 35.0 | | | Cluster_18928 |
| 300.71000 | | 35.0 | | | Cluster_18955 |
| 301.50000 | | 35.0 | | | Cluster_06851 |
| 301.65000 | | 35.0 | | | Cluster_18984 |
| 302.64000 | | 35.0 | | | Cluster_19016 |
| 302.65000 | | 35.0 | | | Cluster_19017 |
| 303.99000 | | 35.0 | | | Cluster_19040 |
| 304.11000 | | 35.0 | | | Cluster_19042 |
| 304.14000 | | 35.0 | | | Cluster_19049 |
| 304.16000 | | 35.0 | | | Cluster_19050 |
| 304.20000 | | 35.0 | | | Cluster_19052 |
| 304.49000 | | 35.0 | | | Cluster_06863 |
| 304.64000 | | 35.0 | | | Cluster_19062 |
| 306.13000 | | 35.0 | | | Cluster_06873 |
| 306.67000 | | 35.0 | | | Cluster_19109 |
| 307.12000 | | 35.0 | | | Cluster_19119 |
| 307.13000 | | 35.0 | | | Cluster_06880 |
| 308.65000 | | 35.0 | | | Cluster_19177 |
| 308.66000 | | 35.0 | | | Cluster_19179 |
| 309.53000 | | 35.0 | | | Cluster_06901 |
| 310.58000 | | 35.0 | | | Cluster_19217 |
| 310.65000 | | 35.0 | | | Cluster_19222 |
| 310.66000 | | 35.0 | | | Cluster_19221 |
| 311.17000 | | 35.0 | | | Cluster_19246 |
| 311.68000 | | 35.0 | | | Cluster_19257 |
| 313.65000 | | 35.0 | | | Cluster_19315 |
| 314.82000 | | 35.0 | | | Cluster_06941 |
| 315.65000 | | 35.0 | | | Cluster_19384 |
| 316.69000 | | 35.0 | | | Cluster_19442 |
| 317.06000 | | 35.0 | | | Cluster_19459 |
| 317.16000 | | 35.0 | | | Cluster_19466 |
| 317.64000 | | 35.0 | | | Cluster_19490 |
| 318.64000 | | 35.0 | | | Cluster_19532 |

| | | |
|---|---|---|
| 318.85000 | 35.0 | Cluster_06973 |
| 319.17000 | 35.0 | Cluster_19552 |
| 319.67000 | 35.0 | Cluster_19560 |
| 319.94000 | 35.0 | Cluster_00850 |
| 320.49000 | 35.0 | Cluster_06993 |
| 320.80000 | 35.0 | Cluster_06994 |
| 321.15000 | 35.0 | Cluster_06995 |
| 321.17000 | 35.0 | Cluster_00869 |
| 321.52000 | 35.0 | Cluster_19594 |
| 322.15000 | 35.0 | Cluster_19619 |
| 322.17000 | 35.0 | Cluster_19617 |
| 322.50000 | 35.0 | Cluster_06998 |
| 323.65000 | 35.0 | Cluster_19672 |
| 323.69000 | 35.0 | Cluster_19677 |
| 323.83000 | 35.0 | Cluster_19683 |
| 324.42000 | 35.0 | Cluster_00878 |
| 324.60000 | 35.0 | Cluster_19711 |
| 324.65000 | 35.0 | Cluster_19713 |
| 324.83000 | 35.0 | Cluster_07017 |
| 325.14000 | 35.0 | Cluster_19730 |
| 325.82000 | 35.0 | Cluster_07023 |
| 325.85000 | 35.0 | Cluster_07028 |
| 326.17000 | 35.0 | Cluster_19753 |
| 326.40000 | 35.0 | Cluster_00887 |
| 326.90000 | 35.0 | Cluster_00888 |
| 327.21000 | 35.0 | Cluster_19791 |
| 327.96000 | 35.0 | Cluster_07043 |
| 328.48000 | 35.0 | Cluster_07051 |
| 329.18000 | 35.0 | Cluster_19855 |
| 329.82000 | 35.0 | Cluster_07063 |
| 329.93000 | 35.0 | Cluster_19873 |
| 330.18000 | 35.0 | Cluster_19886 |
| 330.90000 | 35.0 | Cluster_19916 |
| 331.17000 | 35.0 | Cluster_19926 |
| 331.20000 | 35.0 | Cluster_19928 |
| 331.21000 | 35.0 | Cluster_19930 |
| 331.22000 | 35.0 | Cluster_19925 |
| 331.66000 | 35.0 | Cluster_19936 |
| 332.14000 | 35.0 | Cluster_19948 |
| 333.47000 | 35.0 | Cluster_07081 |
| 333.82000 | 35.0 | Cluster_07087 |
| 333.87000 | 35.0 | Cluster_07090 |
| 335.12000 | 35.0 | Cluster_20034 |
| 335.83000 | 35.0 | Cluster_07104 |
| 336.19000 | 35.0 | Cluster_07109 |
| 336.23000 | 35.0 | Cluster_20093 |
| 336.91000 | 35.0 | Cluster_00920 |
| 337.19000 | 35.0 | Cluster_20132 |
| 337.21000 | 35.0 | Cluster_20126 |
| 337.83000 | 35.0 | Cluster_07126 |
| 338.11000 | 35.0 | Cluster_20150 |

| | | |
|---|---|---|
| 338.67000 | 35.0 | Cluster_20172 |
| 339.16000 | 35.0 | Cluster_20192 |
| 339.47000 | 35.0 | Cluster_07139 |
| 339.81000 | 35.0 | Cluster_07150 |
| 339.94000 | 35.0 | Cluster_00929 |
| 340.18000 | 35.0 | Cluster_20223 |
| 340.26000 | 35.0 | Cluster_20229 |
| 341.15000 | 35.0 | Cluster_20257 |
| 341.16000 | 35.0 | Cluster_20261 |
| 342.16000 | 35.0 | Cluster_07167 |
| 343.11000 | 35.0 | Cluster_20316 |
| 343.18000 | 35.0 | Cluster_20318 |
| 343.66000 | 35.0 | Cluster_20344 |
| 345.19000 | 35.0 | Cluster_20390 |
| 345.67000 | 35.0 | Cluster_20402 |
| 346.15000 | 35.0 | Cluster_07199 |
| 346.67000 | 35.0 | Cluster_20444 |
| 347.20000 | 35.0 | Cluster_20457 |
| 347.71000 | 35.0 | Cluster_20471 |
| 348.44000 | 35.0 | Cluster_20495 |
| 348.87000 | 35.0 | Cluster_07234 |
| 348.92000 | 35.0 | Cluster_00975 |
| 348.93000 | 35.0 | Cluster_00976 |
| 349.20000 | 35.0 | Cluster_20517 |
| 349.21000 | 35.0 | Cluster_20520 |
| 349.64000 | 35.0 | Cluster_20543 |
| 349.73000 | 35.0 | Cluster_20549 |
| 350.42000 | 35.0 | Cluster_00982 |
| 351.18000 | 35.0 | Cluster_20611 |
| 351.19000 | 35.0 | Cluster_20622 |
| 352.66000 | 35.0 | Cluster_20669 |
| 353.17000 | 35.0 | Cluster_01002 |
| 353.21000 | 35.0 | Cluster_20705 |
| 353.66000 | 35.0 | Cluster_01003 |
| 353.80000 | 35.0 | Cluster_07278 |
| 353.87000 | 35.0 | Cluster_20720 |
| 353.91000 | 35.0 | Cluster_01004 |
| 354.16000 | 35.0 | Cluster_20732 |
| 354.20000 | 35.0 | Cluster_20739 |
| 354.40000 | 35.0 | Cluster_01009 |
| 354.42000 | 35.0 | Cluster_01011 |
| 355.14000 | 35.0 | Cluster_20781 |
| 355.16000 | 35.0 | Cluster_20782 |
| 355.22000 | 35.0 | Cluster_20787 |
| 355.36000 | 35.0 | Cluster_20789 |
| 355.70000 | 35.0 | Cluster_20796 |
| 356.24000 | 35.0 | Cluster_20824 |
| 356.55000 | 35.0 | Cluster_07307 |
| 358.42000 | 35.0 | Cluster_20933 |
| 359.13000 | 35.0 | Cluster_07326 |
| 359.17000 | 35.0 | Cluster_00118 |

| | | |
|---|---|---|
| 359.37000 | 35.0 | Cluster_20960 |
| 360.16000 | 35.0 | Cluster_20986 |
| 360.42000 | 35.0 | Cluster_01047 |
| 360.83000 | 35.0 | Cluster_21008 |
| 361.18000 | 35.0 | Cluster_21024 |
| 361.42000 | 35.0 | Cluster_01055 |
| 361.66000 | 35.0 | Cluster_21038 |
| 361.67000 | 35.0 | Cluster_21036 |
| 361.69000 | 35.0 | Cluster_21039 |
| 361.84000 | 35.0 | Cluster_07347 |
| 362.10000 | 35.0 | Cluster_21048 |
| 362.42000 | 35.0 | Cluster_01059 |
| 362.43000 | 35.0 | Cluster_21075 |
| 362.60000 | 35.0 | Cluster_21077 |
| 362.65000 | 35.0 | Cluster_21076 |
| 363.11000 | 35.0 | Cluster_21089 |
| 363.19000 | 35.0 | Cluster_21093 |
| 363.20000 | 35.0 | Cluster_21094 |
| 363.77000 | 35.0 | Cluster_21121 |
| 364.16000 | 35.0 | Cluster_21135 |
| 364.50000 | 35.0 | Cluster_21161 |
| 364.65000 | 35.0 | Cluster_21162 |
| 365.24000 | 35.0 | Cluster_21221 |
| 365.79000 | 35.0 | Cluster_07396 |
| 365.90000 | 35.0 | Cluster_21241 |
| 366.27000 | 35.0 | Cluster_21267 |
| 366.74000 | 35.0 | Cluster_21285 |
| 367.22000 | 35.0 | Cluster_21306 |
| 367.50000 | 35.0 | Cluster_07416 |
| 367.53000 | 35.0 | Cluster_21324 |
| 368.16000 | 35.0 | Cluster_21368 |
| 368.20000 | 35.0 | Cluster_21366 |
| 368.47000 | 35.0 | Cluster_07428 |
| 369.67000 | 35.0 | Cluster_21437 |
| 370.08000 | 35.0 | Cluster_21453 |
| 370.50000 | 35.0 | Cluster_07445 |
| 370.74000 | 35.0 | Cluster_21476 |
| 370.93000 | 35.0 | Cluster_21486 |
| 371.18000 | 35.0 | Cluster_07449 |
| 371.24000 | 35.0 | Cluster_21501 |
| 371.71000 | 35.0 | Cluster_21517 |
| 371.89000 | 35.0 | Cluster_21531 |
| 372.17000 | 35.0 | Cluster_21537 |
| 372.23000 | 35.0 | Cluster_07464 |
| 372.67000 | 35.0 | Cluster_21575 |
| 372.69000 | 35.0 | Cluster_21572 |
| 373.12000 | 35.0 | Cluster_21589 |
| 373.14000 | 35.0 | Cluster_21585 |
| 373.16000 | 35.0 | Cluster_07470 |
| 373.19000 | 35.0 | Cluster_07471 |
| 373.22000 | 35.0 | Cluster_21597 |

| | | |
|---|---|---|
| 373.90000 | 35.0 | Cluster_21628 |
| 374.19000 | 35.0 | Cluster_07480 |
| 374.40000 | 35.0 | Cluster_21650 |
| 374.68000 | 35.0 | Cluster_21656 |
| 375.15000 | 35.0 | Cluster_21668 |
| 375.89000 | 35.0 | Cluster_07496 |
| 377.48000 | 35.0 | Cluster_07512 |
| 377.77000 | 35.0 | Cluster_21780 |
| 378.17000 | 35.0 | Cluster_01122 |
| 378.18000 | 35.0 | Cluster_21788 |
| 378.20000 | 35.0 | Cluster_07523 |
| 378.67000 | 35.0 | Cluster_01125 |
| 379.26000 | 35.0 | Cluster_21841 |
| 380.45000 | 35.0 | Cluster_21886 |
| 380.68000 | 35.0 | Cluster_21891 |
| 381.22000 | 35.0 | Cluster_07550 |
| 381.70000 | 35.0 | Cluster_01141 |
| 382.18000 | 35.0 | Cluster_21953 |
| 382.66000 | 35.0 | Cluster_01144 |
| 383.63000 | 35.0 | Cluster_22024 |
| 383.68000 | 35.0 | Cluster_22025 |
| 384.16000 | 35.0 | Cluster_01146 |
| 384.21000 | 35.0 | Cluster_22055 |
| 384.22000 | 35.0 | Cluster_22065 |
| 385.52000 | 35.0 | Cluster_07597 |
| 385.54000 | 35.0 | Cluster_07596 |
| 385.86000 | 35.0 | Cluster_22123 |
| 386.66000 | 35.0 | Cluster_22155 |
| 386.68000 | 35.0 | Cluster_22158 |
| 386.90000 | 35.0 | Cluster_22172 |
| 387.11000 | 35.0 | Cluster_22177 |
| 387.63000 | 35.0 | Cluster_22217 |
| 387.93000 | 35.0 | Cluster_22226 |
| 388.23000 | 35.0 | Cluster_22233 |
| 388.50000 | 35.0 | Cluster_07621 |
| 388.64000 | 35.0 | Cluster_22244 |
| 388.67000 | 35.0 | Cluster_22245 |
| 388.83000 | 35.0 | Cluster_07623 |
| 389.25000 | 35.0 | Cluster_22269 |
| 389.51000 | 35.0 | Cluster_07627 |
| 389.91000 | 35.0 | Cluster_22284 |
| 390.71000 | 35.0 | Cluster_22316 |
| 390.87000 | 35.0 | Cluster_07651 |
| 390.88000 | 35.0 | Cluster_07646 |
| 391.50000 | 35.0 | Cluster_07654 |
| 391.65000 | 35.0 | Cluster_22334 |
| 391.75000 | 35.0 | Cluster_22344 |
| 391.82000 | 35.0 | Cluster_07658 |
| 392.22000 | 35.0 | Cluster_07665 |
| 392.25000 | 35.0 | Cluster_22358 |
| 392.73000 | 35.0 | Cluster_22381 |

| | | |
|---|---|---|
| 392.77000 | 35.0 | Cluster_22377 |
| 393.19000 | 35.0 | Cluster_01195 |
| 393.53000 | 35.0 | Cluster_07674 |
| 393.69000 | 35.0 | Cluster_01197 |
| 393.71000 | 35.0 | Cluster_22428 |
| 393.98000 | 35.0 | Cluster_22479 |
| 394.21000 | 35.0 | Cluster_22490 |
| 394.71000 | 35.0 | Cluster_22522 |
| 394.72000 | 35.0 | Cluster_22521 |
| 395.89000 | 35.0 | Cluster_07703 |
| 396.23000 | 35.0 | Cluster_22589 |
| 396.30000 | 35.0 | Cluster_22595 |
| 397.39000 | 35.0 | Cluster_22628 |
| 397.67000 | 35.0 | Cluster_22634 |
| 397.72000 | 35.0 | Cluster_22641 |
| 397.90000 | 35.0 | Cluster_07736 |
| 398.18000 | 35.0 | Cluster_01226 |
| 398.44000 | 35.0 | Cluster_01230 |
| 400.24000 | 35.0 | Cluster_22747 |
| 400.68000 | 35.0 | Cluster_22780 |
| 400.90000 | 35.0 | Cluster_07771 |
| 401.70000 | 35.0 | Cluster_22833 |
| 402.32000 | 35.0 | Cluster_22866 |
| 402.48000 | 35.0 | Cluster_22868 |
| 402.72000 | 35.0 | Cluster_22878 |
| 403.44000 | 35.0 | Cluster_01260 |
| 403.54000 | 35.0 | Cluster_07797 |
| 404.47000 | 35.0 | Cluster_01272 |
| 405.88000 | 35.0 | Cluster_07831 |
| 406.22000 | 35.0 | Cluster_23023 |
| 406.68000 | 35.0 | Cluster_01285 |
| 406.73000 | 35.0 | Cluster_23046 |
| 406.95000 | 35.0 | Cluster_01293 |
| 407.17000 | 35.0 | Cluster_23059 |
| 407.25000 | 35.0 | Cluster_23066 |
| 407.67000 | 35.0 | Cluster_01302 |
| 407.75000 | 35.0 | Cluster_23083 |
| 407.93000 | 35.0 | Cluster_01303 |
| 408.22000 | 35.0 | Cluster_23104 |
| 408.54000 | 35.0 | Cluster_07863 |
| 408.66000 | 35.0 | Cluster_01305 |
| 408.91000 | 35.0 | Cluster_07864 |
| 408.92000 | 35.0 | Cluster_01307 |
| 409.19000 | 35.0 | Cluster_07867 |
| 409.25000 | 35.0 | Cluster_23160 |
| 409.69000 | 35.0 | Cluster_23170 |
| 410.53000 | 35.0 | Cluster_07882 |
| 411.72000 | 35.0 | Cluster_23258 |
| 412.67000 | 35.0 | Cluster_23299 |
| 412.68000 | 35.0 | Cluster_01326 |
| 413.23000 | 35.0 | Cluster_23320 |

| | | |
|---|---|---|
| 413.87000 | 35.0 | Cluster_07913 |
| 413.92000 | 35.0 | Cluster_07916 |
| 413.94000 | 35.0 | Cluster_23350 |
| 413.98000 | 35.0 | Cluster_23354 |
| 414.71000 | 35.0 | Cluster_23379 |
| 414.85000 | 35.0 | Cluster_23397 |
| 414.87000 | 35.0 | Cluster_07924 |
| 415.22000 | 35.0 | Cluster_23422 |
| 415.73000 | 35.0 | Cluster_23434 |
| 416.00000 | 35.0 | Cluster_23448 |
| 416.22000 | 35.0 | Cluster_23463 |
| 416.76000 | 35.0 | Cluster_23476 |
| 416.88000 | 35.0 | Cluster_23482 |
| 416.94000 | 35.0 | Cluster_01353 |
| 418.25000 | 35.0 | Cluster_23534 |
| 418.47000 | 35.0 | Cluster_23549 |
| 419.18000 | 35.0 | Cluster_01360 |
| 419.21000 | 35.0 | Cluster_07978 |
| 419.88000 | 35.0 | Cluster_23600 |
| 420.19000 | 35.0 | Cluster_23617 |
| 420.23000 | 35.0 | Cluster_23612 |
| 420.71000 | 35.0 | Cluster_23632 |
| 421.56000 | 35.0 | Cluster_08005 |
| 422.12000 | 35.0 | Cluster_08008 |
| 422.55000 | 35.0 | Cluster_08014 |
| 422.58000 | 35.0 | Cluster_08015 |
| 423.14000 | 35.0 | Cluster_23727 |
| 423.21000 | 35.0 | Cluster_08022 |
| 423.24000 | 35.0 | Cluster_23740 |
| 423.44000 | 35.0 | Cluster_01387 |
| 423.71000 | 35.0 | Cluster_23753 |
| 423.74000 | 35.0 | Cluster_23766 |
| 424.18000 | 35.0 | Cluster_23787 |
| 424.51000 | 35.0 | Cluster_23811 |
| 424.68000 | 35.0 | Cluster_23813 |
| 425.18000 | 35.0 | Cluster_23850 |
| 425.46000 | 35.0 | Cluster_23865 |
| 425.87000 | 35.0 | Cluster_08056 |
| 425.88000 | 35.0 | Cluster_23895 |
| 426.14000 | 35.0 | Cluster_23901 |
| 426.24000 | 35.0 | Cluster_23924 |
| 426.44000 | 35.0 | Cluster_23942 |
| 426.88000 | 35.0 | Cluster_08077 |
| 427.51000 | 35.0 | Cluster_08094 |
| 428.56000 | 35.0 | Cluster_08107 |
| 428.72000 | 35.0 | Cluster_24045 |
| 428.79000 | 35.0 | Cluster_24059 |
| 429.24000 | 35.0 | Cluster_24074 |
| 429.57000 | 35.0 | Cluster_08120 |
| 429.72000 | 35.0 | Cluster_24100 |
| 429.76000 | 35.0 | Cluster_24093 |

| | | |
|---|---|---|
| 429.85000 | 35.0 | Cluster_08124 |
| 429.95000 | 35.0 | Cluster_01432 |
| 430.73000 | 35.0 | Cluster_24136 |
| 431.21000 | 35.0 | Cluster_24172 |
| 431.41000 | 35.0 | Cluster_24179 |
| 431.74000 | 35.0 | Cluster_24194 |
| 432.16000 | 35.0 | Cluster_24207 |
| 432.21000 | 35.0 | Cluster_24215 |
| 432.87000 | 35.0 | Cluster_08164 |
| 433.96000 | 35.0 | Cluster_24289 |
| 434.79000 | 35.0 | Cluster_24347 |
| 434.97000 | 35.0 | Cluster_24350 |
| 435.17000 | 35.0 | Cluster_24357 |
| 435.24000 | 35.0 | Cluster_08201 |
| 435.37000 | 35.0 | Cluster_24383 |
| 435.70000 | 35.0 | Cluster_01481 |
| 435.97000 | 35.0 | Cluster_01487 |
| 436.48000 | 35.0 | Cluster_01491 |
| 437.37000 | 35.0 | Cluster_24491 |
| 437.46000 | 35.0 | Cluster_01497 |
| 438.22000 | 35.0 | Cluster_24532 |
| 438.30000 | 35.0 | Cluster_24543 |
| 438.96000 | 35.0 | Cluster_01503 |
| 438.97000 | 35.0 | Cluster_01504 |
| 439.82000 | 35.0 | Cluster_24632 |
| 440.21000 | 35.0 | Cluster_01512 |
| 440.56000 | 35.0 | Cluster_24655 |
| 440.74000 | 35.0 | Cluster_24659 |
| 440.86000 | 35.0 | Cluster_08277 |
| 441.21000 | 35.0 | Cluster_24682 |
| 441.24000 | 35.0 | Cluster_01516 |
| 441.73000 | 35.0 | Cluster_24703 |
| 441.87000 | 35.0 | Cluster_08293 |
| 442.00000 | 35.0 | Cluster_24716 |
| 442.28000 | 35.0 | Cluster_24730 |
| 442.57000 | 35.0 | Cluster_08303 |
| 442.77000 | 35.0 | Cluster_24758 |
| 442.89000 | 35.0 | Cluster_24763 |
| 442.98000 | 35.0 | Cluster_24761 |
| 443.12000 | 35.0 | Cluster_24771 |
| 443.22000 | 35.0 | Cluster_24782 |
| 443.28000 | 35.0 | Cluster_24785 |
| 443.29000 | 35.0 | Cluster_24795 |
| 443.89000 | 35.0 | Cluster_08322 |
| 443.92000 | 35.0 | Cluster_08324 |
| 444.72000 | 35.0 | Cluster_24866 |
| 445.18000 | 35.0 | Cluster_08343 |
| 445.55000 | 35.0 | Cluster_24900 |
| 446.27000 | 35.0 | Cluster_24941 |
| 446.73000 | 35.0 | Cluster_24959 |
| 446.91000 | 35.0 | Cluster_08369 |

| | | |
|---|---|---|
| 447.17000 | 35.0 | Cluster_24974 |
| 447.25000 | 35.0 | Cluster_08376 |
| 447.28000 | 35.0 | Cluster_24982 |
| 447.29000 | 35.0 | Cluster_24988 |
| 447.60000 | 35.0 | Cluster_08380 |
| 448.30000 | 35.0 | Cluster_25022 |
| 448.79000 | 35.0 | Cluster_25051 |
| 448.97000 | 35.0 | Cluster_01573 |
| 449.25000 | 35.0 | Cluster_08416 |
| 449.74000 | 35.0 | Cluster_25092 |
| 449.84000 | 35.0 | Cluster_25102 |
| 450.23000 | 35.0 | Cluster_25125 |
| 450.93000 | 35.0 | Cluster_08435 |
| 451.14000 | 35.0 | Cluster_25153 |
| 451.17000 | 35.0 | Cluster_25168 |
| 451.57000 | 35.0 | Cluster_08446 |
| 452.13000 | 35.0 | Cluster_25195 |
| 452.26000 | 35.0 | Cluster_08462 |
| 452.75000 | 35.0 | Cluster_25235 |
| 453.14000 | 35.0 | Cluster_25244 |
| 453.24000 | 35.0 | Cluster_25248 |
| 454.68000 | 35.0 | Cluster_25320 |
| 454.70000 | 35.0 | Cluster_25319 |
| 454.74000 | 35.0 | Cluster_25325 |
| 455.15000 | 35.0 | Cluster_25343 |
| 455.18000 | 35.0 | Cluster_01617 |
| 455.20000 | 35.0 | Cluster_08518 |
| 456.24000 | 35.0 | Cluster_25393 |
| 456.26000 | 35.0 | Cluster_08536 |
| 456.57000 | 35.0 | Cluster_08539 |
| 456.75000 | 35.0 | Cluster_25404 |
| 456.78000 | 35.0 | Cluster_25413 |
| 457.20000 | 35.0 | Cluster_25422 |
| 457.75000 | 35.0 | Cluster_25448 |
| 457.76000 | 35.0 | Cluster_25454 |
| 458.69000 | 35.0 | Cluster_25493 |
| 458.76000 | 35.0 | Cluster_25500 |
| 459.22000 | 35.0 | Cluster_08566 |
| 459.25000 | 35.0 | Cluster_25514 |
| 459.78000 | 35.0 | Cluster_25540 |
| 459.90000 | 35.0 | Cluster_08580 |
| 460.27000 | 35.0 | Cluster_25572 |
| 460.77000 | 35.0 | Cluster_25581 |
| 460.97000 | 35.0 | Cluster_01663 |
| 461.60000 | 35.0 | Cluster_08592 |
| 461.70000 | 35.0 | Cluster_25618 |
| 461.73000 | 35.0 | Cluster_25620 |
| 462.22000 | 35.0 | Cluster_08608 |
| 462.50000 | 35.0 | Cluster_25643 |
| 463.61000 | 35.0 | Cluster_25677 |
| 463.73000 | 35.0 | Cluster_25692 |

| | | |
|---|---|---|
| 464.22000 | 35.0 | Cluster_08632 |
| 464.30000 | 35.0 | Cluster_25714 |
| 464.55000 | 35.0 | Cluster_08640 |
| 465.77000 | 35.0 | Cluster_25807 |
| 465.95000 | 35.0 | Cluster_08659 |
| 465.99000 | 35.0 | Cluster_01692 |
| 466.27000 | 35.0 | Cluster_25822 |
| 466.90000 | 35.0 | Cluster_08673 |
| 467.18000 | 35.0 | Cluster_08677 |
| 467.25000 | 35.0 | Cluster_08678 |
| 467.30000 | 35.0 | Cluster_25858 |
| 467.67000 | 35.0 | Cluster_00144 |
| 467.72000 | 35.0 | Cluster_25877 |
| 468.89000 | 35.0 | Cluster_08711 |
| 469.26000 | 35.0 | Cluster_08720 |
| 469.74000 | 35.0 | Cluster_25967 |
| 469.88000 | 35.0 | Cluster_08724 |
| 470.43000 | 35.0 | Cluster_26002 |
| 470.56000 | 35.0 | Cluster_00010 |
| 470.92000 | 35.0 | Cluster_08745 |
| 471.00000 | 35.0 | Cluster_01731 |
| 471.22000 | 35.0 | Cluster_08748 |
| 471.25000 | 35.0 | Cluster_01737 |
| 471.26000 | 35.0 | Cluster_01738 |
| 471.48000 | 35.0 | Cluster_26044 |
| 471.74000 | 35.0 | Cluster_26063 |
| 471.84000 | 35.0 | Cluster_26052 |
| 472.23000 | 35.0 | Cluster_26081 |
| 472.24000 | 35.0 | Cluster_08764 |
| 472.72000 | 35.0 | Cluster_26107 |
| 472.76000 | 35.0 | Cluster_26106 |
| 472.77000 | 35.0 | Cluster_26113 |
| 473.27000 | 35.0 | Cluster_26146 |
| 473.77000 | 35.0 | Cluster_26157 |
| 474.54000 | 35.0 | Cluster_08802 |
| 474.58000 | 35.0 | Cluster_08800 |
| 474.85000 | 35.0 | Cluster_08806 |
| 475.58000 | 35.0 | Cluster_08820 |
| 475.78000 | 35.0 | Cluster_26234 |
| 476.21000 | 35.0 | Cluster_26244 |
| 476.23000 | 35.0 | Cluster_08830 |
| 476.28000 | 35.0 | Cluster_08832 |
| 477.07000 | 35.0 | Cluster_26275 |
| 477.20000 | 35.0 | Cluster_08843 |
| 477.27000 | 35.0 | Cluster_08840 |
| 477.55000 | 35.0 | Cluster_08851 |
| 477.61000 | 35.0 | Cluster_08856 |
| 477.93000 | 35.0 | Cluster_08861 |
| 478.22000 | 35.0 | Cluster_26309 |
| 478.25000 | 35.0 | Cluster_26313 |
| 478.76000 | 35.0 | Cluster_26329 |

| | | |
|---|---|---|
| 478.77000 | 35.0 | Cluster_26332 |
| 478.93000 | 35.0 | Cluster_08880 |
| 478.99000 | 35.0 | Cluster_01780 |
| 479.57000 | 35.0 | Cluster_08888 |
| 479.71000 | 35.0 | Cluster_26429 |
| 479.76000 | 35.0 | Cluster_26423 |
| 480.22000 | 35.0 | Cluster_08900 |
| 480.70000 | 35.0 | Cluster_26476 |
| 481.22000 | 35.0 | Cluster_26493 |
| 481.27000 | 35.0 | Cluster_26504 |
| 481.28000 | 35.0 | Cluster_08920 |
| 481.56000 | 35.0 | Cluster_08922 |
| 481.58000 | 35.0 | Cluster_08924 |
| 481.72000 | 35.0 | Cluster_26523 |
| 481.73000 | 35.0 | Cluster_01804 |
| 481.79000 | 35.0 | Cluster_26518 |
| 482.20000 | 35.0 | Cluster_01810 |
| 482.28000 | 35.0 | Cluster_26547 |
| 482.49000 | 35.0 | Cluster_26549 |
| 483.24000 | 35.0 | Cluster_26594 |
| 483.30000 | 35.0 | Cluster_26603 |
| 483.76000 | 35.0 | Cluster_26625 |
| 483.84000 | 35.0 | Cluster_26627 |
| 484.09000 | 35.0 | Cluster_26635 |
| 484.73000 | 35.0 | Cluster_26685 |
| 484.76000 | 35.0 | Cluster_26681 |
| 485.06000 | 35.0 | Cluster_00166 |
| 485.54000 | 35.0 | Cluster_08981 |
| 488.29000 | 35.0 | Cluster_26837 |
| 488.61000 | 35.0 | Cluster_09032 |
| 488.79000 | 35.0 | Cluster_26859 |
| 488.95000 | 35.0 | Cluster_09038 |
| 489.74000 | 35.0 | Cluster_26904 |
| 489.93000 | 35.0 | Cluster_09048 |
| 490.50000 | 35.0 | Cluster_26931 |
| 490.70000 | 35.0 | Cluster_26935 |
| 490.77000 | 35.0 | Cluster_26939 |
| 492.55000 | 35.0 | Cluster_09097 |
| 492.60000 | 35.0 | Cluster_09099 |
| 493.27000 | 35.0 | Cluster_27041 |
| 494.26000 | 35.0 | Cluster_27100 |
| 495.19000 | 35.0 | Cluster_27149 |
| 495.71000 | 35.0 | Cluster_27170 |
| 495.95000 | 35.0 | Cluster_09160 |
| 496.24000 | 35.0 | Cluster_01895 |
| 496.95000 | 35.0 | Cluster_09176 |
| 497.35000 | 35.0 | Cluster_27258 |
| 497.74000 | 35.0 | Cluster_27265 |
| 497.80000 | 35.0 | Cluster_27275 |
| 497.94000 | 35.0 | Cluster_09191 |
| 498.01000 | 35.0 | Cluster_01929 |

| | | |
|---|---|---|
| 498.50000 | 35.0 | Cluster_27299 |
| 498.73000 | 35.0 | Cluster_27302 |
| 499.23000 | 35.0 | Cluster_09211 |
| 499.33000 | 35.0 | Cluster_27324 |
| 499.76000 | 35.0 | Cluster_01941 |
| 499.92000 | 35.0 | Cluster_09218 |
| 499.96000 | 35.0 | Cluster_09219 |
| 500.02000 | 35.0 | Cluster_27355 |
| 500.63000 | 35.0 | Cluster_09231 |
| 500.70000 | 35.0 | Cluster_27401 |
| 500.96000 | 35.0 | Cluster_01956 |
| 501.22000 | 35.0 | Cluster_27436 |
| 501.59000 | 35.0 | Cluster_09244 |
| 501.76000 | 35.0 | Cluster_27459 |
| 502.21000 | 35.0 | Cluster_01970 |
| 502.22000 | 35.0 | Cluster_27473 |
| 502.59000 | 35.0 | Cluster_09268 |
| 502.76000 | 35.0 | Cluster_27485 |
| 503.25000 | 35.0 | Cluster_27512 |
| 503.28000 | 35.0 | Cluster_27518 |
| 504.21000 | 35.0 | Cluster_27547 |
| 504.56000 | 35.0 | Cluster_09288 |
| 504.98000 | 35.0 | Cluster_01984 |
| 505.01000 | 35.0 | Cluster_27567 |
| 505.24000 | 35.0 | Cluster_09298 |
| 505.52000 | 35.0 | Cluster_01990 |
| 505.60000 | 35.0 | Cluster_09305 |
| 505.79000 | 35.0 | Cluster_27593 |
| 506.73000 | 35.0 | Cluster_27611 |
| 506.74000 | 35.0 | Cluster_27621 |
| 506.76000 | 35.0 | Cluster_27614 |
| 507.25000 | 35.0 | Cluster_27638 |
| 507.97000 | 35.0 | Cluster_09350 |
| 508.62000 | 35.0 | Cluster_09366 |
| 508.70000 | 35.0 | Cluster_27686 |
| 508.71000 | 35.0 | Cluster_27692 |
| 509.20000 | 35.0 | Cluster_27715 |
| 509.26000 | 35.0 | Cluster_27718 |
| 509.87000 | 35.0 | Cluster_09377 |
| 509.91000 | 35.0 | Cluster_09383 |
| 511.23000 | 35.0 | Cluster_27820 |
| 511.79000 | 35.0 | Cluster_27852 |
| 511.91000 | 35.0 | Cluster_09420 |
| 512.25000 | 35.0 | Cluster_27867 |
| 512.27000 | 35.0 | Cluster_27865 |
| 513.09000 | 35.0 | Cluster_27898 |
| 513.77000 | 35.0 | Cluster_27929 |
| 514.72000 | 35.0 | Cluster_27964 |
| 514.74000 | 35.0 | Cluster_27971 |
| 515.25000 | 35.0 | Cluster_02053 |
| 515.48000 | 35.0 | Cluster_02061 |

| | | |
|---|---|---|
| 515.90000 | 35.0 | Cluster_09478 |
| 516.74000 | 35.0 | Cluster_28062 |
| 517.92000 | 35.0 | Cluster_09503 |
| 518.00000 | 35.0 | Cluster_28115 |
| 518.26000 | 35.0 | Cluster_09508 |
| 518.31000 | 35.0 | Cluster_28130 |
| 518.51000 | 35.0 | Cluster_28138 |
| 518.59000 | 35.0 | Cluster_09521 |
| 518.78000 | 35.0 | Cluster_28152 |
| 519.59000 | 35.0 | Cluster_09531 |
| 519.81000 | 35.0 | Cluster_28200 |
| 520.04000 | 35.0 | Cluster_28202 |
| 520.51000 | 35.0 | Cluster_28225 |
| 520.80000 | 35.0 | Cluster_28240 |
| 520.81000 | 35.0 | Cluster_28232 |
| 521.02000 | 35.0 | Cluster_02095 |
| 521.26000 | 35.0 | Cluster_09553 |
| 521.66000 | 35.0 | Cluster_28271 |
| 522.22000 | 35.0 | Cluster_28286 |
| 522.45000 | 35.0 | Cluster_02102 |
| 522.50000 | 35.0 | Cluster_28303 |
| 522.62000 | 35.0 | Cluster_09582 |
| 522.75000 | 35.0 | Cluster_28315 |
| 523.22000 | 35.0 | Cluster_28350 |
| 523.27000 | 35.0 | Cluster_28334 |
| 523.35000 | 35.0 | Cluster_28353 |
| 523.92000 | 35.0 | Cluster_09603 |
| 523.97000 | 35.0 | Cluster_28374 |
| 524.28000 | 35.0 | Cluster_28386 |
| 526.18000 | 35.0 | Cluster_28467 |
| 526.20000 | 35.0 | Cluster_09656 |
| 526.93000 | 35.0 | Cluster_09676 |
| 526.95000 | 35.0 | Cluster_09677 |
| 527.76000 | 35.0 | Cluster_28507 |
| 528.23000 | 35.0 | Cluster_28536 |
| 528.27000 | 35.0 | Cluster_00184 |
| 528.32000 | 35.0 | Cluster_28539 |
| 528.36000 | 35.0 | Cluster_28552 |
| 528.97000 | 35.0 | Cluster_09710 |
| 529.25000 | 35.0 | Cluster_09712 |
| 529.79000 | 35.0 | Cluster_28615 |
| 530.20000 | 35.0 | Cluster_28629 |
| 530.26000 | 35.0 | Cluster_02153 |
| 530.51000 | 35.0 | Cluster_28657 |
| 530.81000 | 35.0 | Cluster_28677 |
| 530.94000 | 35.0 | Cluster_09741 |
| 531.75000 | 35.0 | Cluster_28707 |
| 531.77000 | 35.0 | Cluster_28708 |
| 532.24000 | 35.0 | Cluster_09763 |
| 532.27000 | 35.0 | Cluster_09766 |
| 532.62000 | 35.0 | Cluster_09768 |

| | | |
|---|---|---|
| 532.97000 | 35.0 | Cluster_09773 |
| 533.15000 | 35.0 | Cluster_28752 |
| 533.16000 | 35.0 | Cluster_28753 |
| 533.19000 | 35.0 | Cluster_09774 |
| 533.32000 | 35.0 | Cluster_09778 |
| 533.70000 | 35.0 | Cluster_28783 |
| 533.82000 | 35.0 | Cluster_28799 |
| 533.91000 | 35.0 | Cluster_09792 |
| 534.28000 | 35.0 | Cluster_28807 |
| 534.59000 | 35.0 | Cluster_09803 |
| 535.32000 | 35.0 | Cluster_09812 |
| 535.80000 | 35.0 | Cluster_28867 |
| 535.94000 | 35.0 | Cluster_09822 |
| 536.06000 | 35.0 | Cluster_28878 |
| 536.26000 | 35.0 | Cluster_09823 |
| 536.27000 | 35.0 | Cluster_28888 |
| 536.48000 | 35.0 | Cluster_28899 |
| 536.60000 | 35.0 | Cluster_09834 |
| 537.23000 | 35.0 | Cluster_09849 |
| 537.56000 | 35.0 | Cluster_09853 |
| 537.69000 | 35.0 | Cluster_28943 |
| 537.91000 | 35.0 | Cluster_09869 |
| 538.84000 | 35.0 | Cluster_28980 |
| 538.97000 | 35.0 | Cluster_09886 |
| 538.98000 | 35.0 | Cluster_09885 |
| 539.01000 | 35.0 | Cluster_28985 |
| 539.28000 | 35.0 | Cluster_28993 |
| 539.52000 | 35.0 | Cluster_29001 |
| 540.02000 | 35.0 | Cluster_09899 |
| 540.27000 | 35.0 | Cluster_29031 |
| 540.79000 | 35.0 | Cluster_29052 |
| 541.34000 | 35.0 | Cluster_29085 |
| 541.44000 | 35.0 | Cluster_29098 |
| 541.82000 | 35.0 | Cluster_29115 |
| 542.26000 | 35.0 | Cluster_02236 |
| 542.28000 | 35.0 | Cluster_02229 |
| 542.52000 | 35.0 | Cluster_02233 |
| 542.77000 | 35.0 | Cluster_29162 |
| 543.27000 | 35.0 | Cluster_29193 |
| 544.95000 | 35.0 | Cluster_09982 |
| 545.77000 | 35.0 | Cluster_29280 |
| 545.92000 | 35.0 | Cluster_09992 |
| 546.31000 | 35.0 | Cluster_29312 |
| 547.32000 | 35.0 | Cluster_29345 |
| 548.27000 | 35.0 | Cluster_29389 |
| 548.46000 | 35.0 | Cluster_00194 |
| 548.61000 | 35.0 | Cluster_10035 |
| 548.64000 | 35.0 | Cluster_10037 |
| 549.25000 | 35.0 | Cluster_10044 |
| 549.28000 | 35.0 | Cluster_29418 |
| 549.90000 | 35.0 | Cluster_10052 |

| | | |
|---|---|---|
| 549.99000 | 35.0 | Cluster_10058 |
| 550.22000 | 35.0 | Cluster_29451 |
| 550.31000 | 35.0 | Cluster_29461 |
| 550.55000 | 35.0 | Cluster_10069 |
| 550.61000 | 35.0 | Cluster_10068 |
| 550.78000 | 35.0 | Cluster_29465 |
| 550.84000 | 35.0 | Cluster_29476 |
| 551.01000 | 35.0 | Cluster_29483 |
| 551.29000 | 35.0 | Cluster_29499 |
| 551.57000 | 35.0 | Cluster_10082 |
| 551.97000 | 35.0 | Cluster_10089 |
| 552.36000 | 35.0 | Cluster_29545 |
| 552.74000 | 35.0 | Cluster_29553 |
| 552.75000 | 35.0 | Cluster_29548 |
| 552.96000 | 35.0 | Cluster_10104 |
| 553.55000 | 35.0 | Cluster_29579 |
| 553.63000 | 35.0 | Cluster_10116 |
| 554.58000 | 35.0 | Cluster_10130 |
| 554.60000 | 35.0 | Cluster_10136 |
| 554.97000 | 35.0 | Cluster_10144 |
| 556.25000 | 35.0 | Cluster_29692 |
| 556.68000 | 35.0 | Cluster_10177 |
| 556.96000 | 35.0 | Cluster_10184 |
| 557.73000 | 35.0 | Cluster_29750 |
| 558.23000 | 35.0 | Cluster_10209 |
| 558.65000 | 35.0 | Cluster_10215 |
| 558.81000 | 35.0 | Cluster_29792 |
| 559.29000 | 35.0 | Cluster_29811 |
| 560.35000 | 35.0 | Cluster_29845 |
| 560.65000 | 35.0 | Cluster_10245 |
| 560.83000 | 35.0 | Cluster_29862 |
| 561.28000 | 35.0 | Cluster_29879 |
| 561.64000 | 35.0 | Cluster_10262 |
| 561.70000 | 35.0 | Cluster_29892 |
| 561.81000 | 35.0 | Cluster_29901 |
| 561.98000 | 35.0 | Cluster_10266 |
| 562.07000 | 35.0 | Cluster_29906 |
| 562.29000 | 35.0 | Cluster_29915 |
| 563.32000 | 35.0 | Cluster_10288 |
| 563.50000 | 35.0 | Cluster_02355 |
| 563.75000 | 35.0 | Cluster_29975 |
| 563.76000 | 35.0 | Cluster_29965 |
| 564.78000 | 35.0 | Cluster_30003 |
| 564.79000 | 35.0 | Cluster_02366 |
| 565.56000 | 35.0 | Cluster_30039 |
| 565.60000 | 35.0 | Cluster_10318 |
| 566.33000 | 35.0 | Cluster_30080 |
| 566.74000 | 35.0 | Cluster_02376 |
| 566.79000 | 35.0 | Cluster_30088 |
| 566.90000 | 35.0 | Cluster_30092 |
| 566.98000 | 35.0 | Cluster_10333 |

| | | |
|---|---|---|
| 567.07000 | 35.0 | Cluster_30105 |
| 567.24000 | 35.0 | Cluster_30110 |
| 567.30000 | 35.0 | Cluster_30123 |
| 567.31000 | 35.0 | Cluster_30114 |
| 568.56000 | 35.0 | Cluster_02384 |
| 568.73000 | 35.0 | Cluster_30166 |
| 568.79000 | 35.0 | Cluster_30177 |
| 568.80000 | 35.0 | Cluster_30176 |
| 568.85000 | 35.0 | Cluster_30178 |
| 569.06000 | 35.0 | Cluster_02386 |
| 569.57000 | 35.0 | Cluster_02390 |
| 569.82000 | 35.0 | Cluster_30204 |
| 570.30000 | 35.0 | Cluster_02396 |
| 570.36000 | 35.0 | Cluster_30219 |
| 570.73000 | 35.0 | Cluster_02397 |
| 570.77000 | 35.0 | Cluster_30236 |
| 571.22000 | 35.0 | Cluster_30252 |
| 571.34000 | 35.0 | Cluster_30256 |
| 571.64000 | 35.0 | Cluster_10409 |
| 571.77000 | 35.0 | Cluster_30264 |
| 572.13000 | 35.0 | Cluster_00213 |
| 572.30000 | 35.0 | Cluster_10420 |
| 572.79000 | 35.0 | Cluster_30301 |
| 573.35000 | 35.0 | Cluster_30317 |
| 574.26000 | 35.0 | Cluster_10456 |
| 574.80000 | 35.0 | Cluster_30355 |
| 575.25000 | 35.0 | Cluster_02433 |
| 575.33000 | 35.0 | Cluster_02435 |
| 576.55000 | 35.0 | Cluster_02439 |
| 576.77000 | 35.0 | Cluster_30417 |
| 576.98000 | 35.0 | Cluster_10494 |
| 577.30000 | 35.0 | Cluster_10500 |
| 577.81000 | 35.0 | Cluster_30446 |
| 578.32000 | 35.0 | Cluster_30460 |
| 578.55000 | 35.0 | Cluster_02454 |
| 578.86000 | 35.0 | Cluster_30486 |
| 578.98000 | 35.0 | Cluster_10523 |
| 579.33000 | 35.0 | Cluster_30494 |
| 579.55000 | 35.0 | Cluster_02460 |
| 579.61000 | 35.0 | Cluster_10531 |
| 579.63000 | 35.0 | Cluster_10533 |
| 579.77000 | 35.0 | Cluster_30512 |
| 579.81000 | 35.0 | Cluster_30517 |
| 579.83000 | 35.0 | Cluster_30515 |
| 580.36000 | 35.0 | Cluster_30555 |
| 580.86000 | 35.0 | Cluster_02466 |
| 581.28000 | 35.0 | Cluster_02468 |
| 581.31000 | 35.0 | Cluster_30577 |
| 581.96000 | 35.0 | Cluster_10563 |
| 582.31000 | 35.0 | Cluster_30648 |
| 582.54000 | 35.0 | Cluster_02478 |

| | | |
|---|---|---|
| 582.65000 | 35.0 | Cluster_10574 |
| 582.70000 | 35.0 | Cluster_00226 |
| 582.74000 | 35.0 | Cluster_30654 |
| 584.05000 | 35.0 | Cluster_30718 |
| 584.31000 | 35.0 | Cluster_30724 |
| 584.54000 | 35.0 | Cluster_30733 |
| 585.02000 | 35.0 | Cluster_30763 |
| 585.29000 | 35.0 | Cluster_10612 |
| 585.35000 | 35.0 | Cluster_02501 |
| 585.80000 | 35.0 | Cluster_30791 |
| 585.84000 | 35.0 | Cluster_30792 |
| 586.62000 | 35.0 | Cluster_10630 |
| 586.75000 | 35.0 | Cluster_30815 |
| 587.30000 | 35.0 | Cluster_10642 |
| 587.33000 | 35.0 | Cluster_30848 |
| 587.34000 | 35.0 | Cluster_30837 |
| 587.65000 | 35.0 | Cluster_10646 |
| 588.69000 | 35.0 | Cluster_30892 |
| 588.82000 | 35.0 | Cluster_30900 |
| 589.55000 | 35.0 | Cluster_30932 |
| 589.74000 | 35.0 | Cluster_30933 |
| 589.96000 | 35.0 | Cluster_10677 |
| 590.22000 | 35.0 | Cluster_30952 |
| 590.30000 | 35.0 | Cluster_30953 |
| 591.04000 | 35.0 | Cluster_30998 |
| 591.44000 | 35.0 | Cluster_31008 |
| 591.52000 | 35.0 | Cluster_02538 |
| 591.66000 | 35.0 | Cluster_10703 |
| 591.81000 | 35.0 | Cluster_31027 |
| 592.01000 | 35.0 | Cluster_10706 |
| 592.79000 | 35.0 | Cluster_31056 |
| 592.82000 | 35.0 | Cluster_31060 |
| 592.91000 | 35.0 | Cluster_31063 |
| 593.06000 | 35.0 | Cluster_02547 |
| 593.07000 | 35.0 | Cluster_02548 |
| 593.33000 | 35.0 | Cluster_10724 |
| 593.34000 | 35.0 | Cluster_31073 |
| 593.93000 | 35.0 | Cluster_10743 |
| 594.32000 | 35.0 | Cluster_10753 |
| 595.59000 | 35.0 | Cluster_31146 |
| 595.62000 | 35.0 | Cluster_10769 |
| 596.75000 | 35.0 | Cluster_31177 |
| 597.67000 | 35.0 | Cluster_10808 |
| 597.81000 | 35.0 | Cluster_31212 |
| 597.82000 | 35.0 | Cluster_31208 |
| 598.29000 | 35.0 | Cluster_10814 |
| 598.36000 | 35.0 | Cluster_10818 |
| 599.36000 | 35.0 | Cluster_31267 |
| 599.66000 | 35.0 | Cluster_10840 |
| 599.95000 | 35.0 | Cluster_10848 |
| 600.81000 | 35.0 | Cluster_31316 |

| | | |
|---|---|---|
| 600.97000 | 35.0 | Cluster_10858 |
| 601.36000 | 35.0 | Cluster_31339 |
| 602.32000 | 35.0 | Cluster_31390 |
| 602.83000 | 35.0 | Cluster_31406 |
| 603.30000 | 35.0 | Cluster_31409 |
| 603.51000 | 35.0 | Cluster_02598 |
| 603.53000 | 35.0 | Cluster_31419 |
| 603.96000 | 35.0 | Cluster_10888 |
| 604.80000 | 35.0 | Cluster_31463 |
| 605.31000 | 35.0 | Cluster_31473 |
| 605.67000 | 35.0 | Cluster_10910 |
| 606.07000 | 35.0 | Cluster_31501 |
| 606.27000 | 35.0 | Cluster_31514 |
| 606.87000 | 35.0 | Cluster_31531 |
| 606.89000 | 35.0 | Cluster_10941 |
| 607.32000 | 35.0 | Cluster_31536 |
| 608.09000 | 35.0 | Cluster_02626 |
| 608.32000 | 35.0 | Cluster_31587 |
| 608.49000 | 35.0 | Cluster_31590 |
| 608.78000 | 35.0 | Cluster_31595 |
| 608.79000 | 35.0 | Cluster_31601 |
| 609.06000 | 35.0 | Cluster_31605 |
| 609.81000 | 35.0 | Cluster_31625 |
| 610.31000 | 35.0 | Cluster_31655 |
| 611.34000 | 35.0 | Cluster_31685 |
| 612.32000 | 35.0 | Cluster_11010 |
| 613.83000 | 35.0 | Cluster_31794 |
| 614.06000 | 35.0 | Cluster_31796 |
| 614.07000 | 35.0 | Cluster_02670 |
| 614.19000 | 35.0 | Cluster_31797 |
| 614.26000 | 35.0 | Cluster_11035 |
| 615.83000 | 35.0 | Cluster_31852 |
| 616.08000 | 35.0 | Cluster_02680 |
| 616.32000 | 35.0 | Cluster_11075 |
| 616.55000 | 35.0 | Cluster_31873 |
| 616.83000 | 35.0 | Cluster_31882 |
| 617.02000 | 35.0 | Cluster_31892 |
| 617.35000 | 35.0 | Cluster_31911 |
| 617.93000 | 35.0 | Cluster_31935 |
| 618.05000 | 35.0 | Cluster_02693 |
| 618.79000 | 35.0 | Cluster_31952 |
| 619.29000 | 35.0 | Cluster_02701 |
| 619.33000 | 35.0 | Cluster_31985 |
| 619.62000 | 35.0 | Cluster_11116 |
| 620.53000 | 35.0 | Cluster_32028 |
| 620.60000 | 35.0 | Cluster_11128 |
| 621.30000 | 35.0 | Cluster_32049 |
| 621.36000 | 35.0 | Cluster_32053 |
| 621.39000 | 35.0 | Cluster_32052 |
| 621.79000 | 35.0 | Cluster_32057 |
| 621.98000 | 35.0 | Cluster_11154 |

| | | |
|---|---|---|
| 622.78000 | 35.0 | Cluster_32095 |
| 622.98000 | 35.0 | Cluster_11175 |
| 623.28000 | 35.0 | Cluster_32112 |
| 623.36000 | 35.0 | Cluster_32126 |
| 623.65000 | 35.0 | Cluster_11191 |
| 623.98000 | 35.0 | Cluster_11189 |
| 624.20000 | 35.0 | Cluster_32153 |
| 624.28000 | 35.0 | Cluster_32154 |
| 624.63000 | 35.0 | Cluster_11202 |
| 625.04000 | 35.0 | Cluster_02732 |
| 625.32000 | 35.0 | Cluster_11203 |
| 625.58000 | 35.0 | Cluster_02735 |
| 626.00000 | 35.0 | Cluster_11225 |
| 626.07000 | 35.0 | Cluster_32208 |
| 626.27000 | 35.0 | Cluster_32218 |
| 626.28000 | 35.0 | Cluster_32212 |
| 626.83000 | 35.0 | Cluster_32244 |
| 627.30000 | 35.0 | Cluster_32266 |
| 627.31000 | 35.0 | Cluster_32263 |
| 627.50000 | 35.0 | Cluster_32275 |
| 627.79000 | 35.0 | Cluster_32283 |
| 628.07000 | 35.0 | Cluster_32292 |
| 628.86000 | 35.0 | Cluster_32321 |
| 629.63000 | 35.0 | Cluster_11289 |
| 629.83000 | 35.0 | Cluster_32344 |
| 629.96000 | 35.0 | Cluster_11292 |
| 630.84000 | 35.0 | Cluster_32375 |
| 631.37000 | 35.0 | Cluster_11312 |
| 631.80000 | 35.0 | Cluster_32410 |
| 632.05000 | 35.0 | Cluster_11324 |
| 632.35000 | 35.0 | Cluster_11329 |
| 632.78000 | 35.0 | Cluster_32445 |
| 632.84000 | 35.0 | Cluster_32450 |
| 633.00000 | 35.0 | Cluster_11339 |
| 633.32000 | 35.0 | Cluster_32465 |
| 633.36000 | 35.0 | Cluster_32472 |
| 635.07000 | 35.0 | Cluster_02772 |
| 635.82000 | 35.0 | Cluster_02775 |
| 636.32000 | 35.0 | Cluster_32584 |
| 636.84000 | 35.0 | Cluster_32596 |
| 637.62000 | 35.0 | Cluster_11403 |
| 637.83000 | 35.0 | Cluster_32632 |
| 638.35000 | 35.0 | Cluster_32655 |
| 638.36000 | 35.0 | Cluster_11416 |
| 638.39000 | 35.0 | Cluster_32659 |
| 638.79000 | 35.0 | Cluster_32669 |
| 638.85000 | 35.0 | Cluster_32673 |
| 639.85000 | 35.0 | Cluster_32724 |
| 639.99000 | 35.0 | Cluster_11446 |
| 640.32000 | 35.0 | Cluster_32732 |
| 641.64000 | 35.0 | Cluster_11472 |

| | | |
|---|---|---|
| 641.84000 | 35.0 | Cluster_32794 |
| 642.02000 | 35.0 | Cluster_11476 |
| 643.07000 | 35.0 | Cluster_32838 |
| 643.83000 | 35.0 | Cluster_32863 |
| 644.00000 | 35.0 | Cluster_11510 |
| 644.30000 | 35.0 | Cluster_02836 |
| 644.36000 | 35.0 | Cluster_32878 |
| 644.37000 | 35.0 | Cluster_11511 |
| 645.00000 | 35.0 | Cluster_11529 |
| 645.85000 | 35.0 | Cluster_32926 |
| 646.30000 | 35.0 | Cluster_32945 |
| 648.34000 | 35.0 | Cluster_33005 |
| 648.67000 | 35.0 | Cluster_11566 |
| 648.88000 | 35.0 | Cluster_33029 |
| 649.01000 | 35.0 | Cluster_11574 |
| 649.77000 | 35.0 | Cluster_33054 |
| 650.31000 | 35.0 | Cluster_33085 |
| 651.01000 | 35.0 | Cluster_11595 |
| 651.08000 | 35.0 | Cluster_33114 |
| 651.32000 | 35.0 | Cluster_33124 |
| 651.36000 | 35.0 | Cluster_11600 |
| 651.86000 | 35.0 | Cluster_33158 |
| 651.99000 | 35.0 | Cluster_11614 |
| 652.06000 | 35.0 | Cluster_33159 |
| 652.35000 | 35.0 | Cluster_33162 |
| 652.86000 | 35.0 | Cluster_33187 |
| 653.33000 | 35.0 | Cluster_33199 |
| 653.57000 | 35.0 | Cluster_33207 |
| 653.60000 | 35.0 | Cluster_02883 |
| 654.05000 | 35.0 | Cluster_33221 |
| 654.68000 | 35.0 | Cluster_11655 |
| 654.78000 | 35.0 | Cluster_33237 |
| 655.99000 | 35.0 | Cluster_11675 |
| 656.63000 | 35.0 | Cluster_11676 |
| 657.90000 | 35.0 | Cluster_33331 |
| 657.95000 | 35.0 | Cluster_33350 |
| 658.07000 | 35.0 | Cluster_33340 |
| 658.66000 | 35.0 | Cluster_11699 |
| 659.28000 | 35.0 | Cluster_33386 |
| 659.60000 | 35.0 | Cluster_33396 |
| 659.63000 | 35.0 | Cluster_02907 |
| 659.98000 | 35.0 | Cluster_11725 |
| 660.33000 | 35.0 | Cluster_33416 |
| 660.46000 | 35.0 | Cluster_33423 |
| 660.84000 | 35.0 | Cluster_33430 |
| 661.01000 | 35.0 | Cluster_11741 |
| 661.31000 | 35.0 | Cluster_33436 |
| 662.02000 | 35.0 | Cluster_11748 |
| 662.12000 | 35.0 | Cluster_02928 |
| 662.30000 | 35.0 | Cluster_33472 |
| 663.37000 | 35.0 | Cluster_33509 |

| | | |
|---|---|---|
| 663.55000 | 35.0 | Cluster_02941 |
| 663.60000 | 35.0 | Cluster_02940 |
| 665.32000 | 35.0 | Cluster_33570 |
| 666.35000 | 35.0 | Cluster_02964 |
| 666.68000 | 35.0 | Cluster_11814 |
| 666.95000 | 35.0 | Cluster_33617 |
| 667.82000 | 35.0 | Cluster_33644 |
| 668.33000 | 35.0 | Cluster_33651 |
| 668.35000 | 35.0 | Cluster_11848 |
| 669.10000 | 35.0 | Cluster_02977 |
| 669.33000 | 35.0 | Cluster_33680 |
| 669.78000 | 35.0 | Cluster_33695 |
| 670.01000 | 35.0 | Cluster_11876 |
| 670.40000 | 35.0 | Cluster_33725 |
| 670.80000 | 35.0 | Cluster_33734 |
| 670.82000 | 35.0 | Cluster_33742 |
| 670.87000 | 35.0 | Cluster_02986 |
| 671.44000 | 35.0 | Cluster_33760 |
| 671.82000 | 35.0 | Cluster_33770 |
| 671.97000 | 35.0 | Cluster_11894 |
| 672.35000 | 35.0 | Cluster_33781 |
| 672.81000 | 35.0 | Cluster_33789 |
| 673.09000 | 35.0 | Cluster_03005 |
| 673.37000 | 35.0 | Cluster_33802 |
| 673.62000 | 35.0 | Cluster_33824 |
| 673.65000 | 35.0 | Cluster_11927 |
| 674.11000 | 35.0 | Cluster_03014 |
| 674.32000 | 35.0 | Cluster_11933 |
| 674.95000 | 35.0 | Cluster_00274 |
| 675.37000 | 35.0 | Cluster_33861 |
| 675.60000 | 35.0 | Cluster_33864 |
| 675.64000 | 35.0 | Cluster_03025 |
| 675.84000 | 35.0 | Cluster_03026 |
| 676.00000 | 35.0 | Cluster_11956 |
| 676.32000 | 35.0 | Cluster_11959 |
| 677.20000 | 35.0 | Cluster_33905 |
| 677.89000 | 35.0 | Cluster_33946 |
| 678.81000 | 35.0 | Cluster_33981 |
| 678.84000 | 35.0 | Cluster_33980 |
| 678.85000 | 35.0 | Cluster_03050 |
| 679.38000 | 35.0 | Cluster_33996 |
| 679.79000 | 35.0 | Cluster_34007 |
| 679.86000 | 35.0 | Cluster_34018 |
| 680.08000 | 35.0 | Cluster_03060 |
| 680.09000 | 35.0 | Cluster_34023 |
| 680.87000 | 35.0 | Cluster_34054 |
| 681.08000 | 35.0 | Cluster_03066 |
| 681.32000 | 35.0 | Cluster_12020 |
| 681.35000 | 35.0 | Cluster_34074 |
| 682.09000 | 35.0 | Cluster_34089 |
| 682.31000 | 35.0 | Cluster_34106 |

| | | |
|---|---|---|
| 682.38000 | 35.0 | Cluster_03074 |
| 683.61000 | 35.0 | Cluster_12059 |
| 683.81000 | 35.0 | Cluster_34134 |
| 683.99000 | 35.0 | Cluster_12062 |
| 684.08000 | 35.0 | Cluster_03088 |
| 684.82000 | 35.0 | Cluster_34171 |
| 684.83000 | 35.0 | Cluster_34174 |
| 685.09000 | 35.0 | Cluster_03095 |
| 685.10000 | 35.0 | Cluster_03102 |
| 685.35000 | 35.0 | Cluster_34201 |
| 686.04000 | 35.0 | Cluster_12086 |
| 686.50000 | 35.0 | Cluster_00285 |
| 686.75000 | 35.0 | Cluster_00286 |
| 687.66000 | 35.0 | Cluster_34255 |
| 689.34000 | 35.0 | Cluster_34298 |
| 689.36000 | 35.0 | Cluster_34295 |
| 689.57000 | 35.0 | Cluster_03142 |
| 690.82000 | 35.0 | Cluster_34324 |
| 690.84000 | 35.0 | Cluster_34327 |
| 691.59000 | 35.0 | Cluster_03162 |
| 691.79000 | 35.0 | Cluster_34357 |
| 691.83000 | 35.0 | Cluster_34369 |
| 692.26000 | 35.0 | Cluster_34373 |
| 692.32000 | 35.0 | Cluster_03167 |
| 692.35000 | 35.0 | Cluster_03168 |
| 692.60000 | 35.0 | Cluster_03182 |
| 692.84000 | 35.0 | Cluster_34402 |
| 693.56000 | 35.0 | Cluster_03192 |
| 693.60000 | 35.0 | Cluster_34413 |
| 693.88000 | 35.0 | Cluster_34418 |
| 694.36000 | 35.0 | Cluster_34424 |
| 694.84000 | 35.0 | Cluster_34436 |
| 694.87000 | 35.0 | Cluster_34443 |
| 695.01000 | 35.0 | Cluster_12183 |
| 695.36000 | 35.0 | Cluster_34458 |
| 695.70000 | 35.0 | Cluster_12190 |
| 695.87000 | 35.0 | Cluster_34468 |
| 696.32000 | 35.0 | Cluster_34489 |
| 696.34000 | 35.0 | Cluster_03210 |
| 696.39000 | 35.0 | Cluster_34483 |
| 696.58000 | 35.0 | Cluster_03211 |
| 697.31000 | 35.0 | Cluster_03215 |
| 697.37000 | 35.0 | Cluster_34515 |
| 697.85000 | 35.0 | Cluster_34520 |
| 697.87000 | 35.0 | Cluster_34523 |
| 698.82000 | 35.0 | Cluster_34541 |
| 698.83000 | 35.0 | Cluster_34546 |
| 699.70000 | 35.0 | Cluster_12244 |
| 699.85000 | 35.0 | Cluster_03235 |
| 700.26000 | 35.0 | Cluster_34580 |
| 700.36000 | 35.0 | Cluster_34589 |

| | | |
|---|---|---|
| 700.65000 | 35.0 | Cluster_34590 |
| 700.85000 | 35.0 | Cluster_34594 |
| 701.36000 | 35.0 | Cluster_34614 |
| 702.34000 | 35.0 | Cluster_34653 |
| 702.62000 | 35.0 | Cluster_12286 |
| 703.04000 | 35.0 | Cluster_12291 |
| 703.05000 | 35.0 | Cluster_03266 |
| 703.33000 | 35.0 | Cluster_34677 |
| 703.69000 | 35.0 | Cluster_12310 |
| 703.91000 | 35.0 | Cluster_03274 |
| 704.17000 | 35.0 | Cluster_34698 |
| 704.21000 | 35.0 | Cluster_34700 |
| 704.31000 | 35.0 | Cluster_34706 |
| 704.81000 | 35.0 | Cluster_34728 |
| 704.85000 | 35.0 | Cluster_34734 |
| 705.04000 | 35.0 | Cluster_12319 |
| 705.37000 | 35.0 | Cluster_34742 |
| 705.83000 | 35.0 | Cluster_34756 |
| 705.88000 | 35.0 | Cluster_34757 |
| 706.86000 | 35.0 | Cluster_03301 |
| 707.02000 | 35.0 | Cluster_12346 |
| 707.80000 | 35.0 | Cluster_34791 |
| 708.51000 | 35.0 | Cluster_00309 |
| 709.28000 | 35.0 | Cluster_34840 |
| 709.39000 | 35.0 | Cluster_34846 |
| 710.88000 | 35.0 | Cluster_03335 |
| 711.06000 | 35.0 | Cluster_12397 |
| 711.85000 | 35.0 | Cluster_34925 |
| 712.39000 | 35.0 | Cluster_34948 |
| 712.84000 | 35.0 | Cluster_34967 |
| 713.08000 | 35.0 | Cluster_03351 |
| 713.29000 | 35.0 | Cluster_03356 |
| 713.38000 | 35.0 | Cluster_12431 |
| 713.72000 | 35.0 | Cluster_12437 |
| 713.77000 | 35.0 | Cluster_34987 |
| 713.92000 | 35.0 | Cluster_34998 |
| 714.32000 | 35.0 | Cluster_35004 |
| 714.36000 | 35.0 | Cluster_35010 |
| 715.38000 | 35.0 | Cluster_35041 |
| 715.54000 | 35.0 | Cluster_00320 |
| 715.71000 | 35.0 | Cluster_12456 |
| 715.85000 | 35.0 | Cluster_35046 |
| 715.99000 | 35.0 | Cluster_12465 |
| 716.39000 | 35.0 | Cluster_35061 |
| 717.36000 | 35.0 | Cluster_12478 |
| 717.90000 | 35.0 | Cluster_35102 |
| 718.36000 | 35.0 | Cluster_35122 |
| 718.85000 | 35.0 | Cluster_35127 |
| 718.89000 | 35.0 | Cluster_35137 |
| 718.91000 | 35.0 | Cluster_35135 |
| 718.96000 | 35.0 | Cluster_12511 |

| | | |
|---|---|---|
| 719.39000 | 35.0 | Cluster_35145 |
| 719.83000 | 35.0 | Cluster_35161 |
| 719.85000 | 35.0 | Cluster_03411 |
| 719.98000 | 35.0 | Cluster_12519 |
| 720.12000 | 35.0 | Cluster_35168 |
| 720.13000 | 35.0 | Cluster_35170 |
| 720.35000 | 35.0 | Cluster_03417 |
| 720.58000 | 35.0 | Cluster_03421 |
| 720.83000 | 35.0 | Cluster_35178 |
| 721.32000 | 35.0 | Cluster_35195 |
| 721.41000 | 35.0 | Cluster_35191 |
| 721.72000 | 35.0 | Cluster_12548 |
| 721.88000 | 35.0 | Cluster_35209 |
| 721.90000 | 35.0 | Cluster_35213 |
| 722.02000 | 35.0 | Cluster_12553 |
| 722.09000 | 35.0 | Cluster_03436 |
| 722.34000 | 35.0 | Cluster_03433 |
| 722.68000 | 35.0 | Cluster_12565 |
| 723.40000 | 35.0 | Cluster_12568 |
| 723.42000 | 35.0 | Cluster_12570 |
| 723.95000 | 35.0 | Cluster_00326 |
| 724.04000 | 35.0 | Cluster_12575 |
| 724.90000 | 35.0 | Cluster_03465 |
| 725.06000 | 35.0 | Cluster_12593 |
| 725.32000 | 35.0 | Cluster_03469 |
| 725.41000 | 35.0 | Cluster_35286 |
| 725.42000 | 35.0 | Cluster_35293 |
| 727.38000 | 35.0 | Cluster_35317 |
| 728.10000 | 35.0 | Cluster_03487 |
| 728.13000 | 35.0 | Cluster_03484 |
| 728.42000 | 35.0 | Cluster_35338 |
| 728.98000 | 35.0 | Cluster_12653 |
| 729.15000 | 35.0 | Cluster_03497 |
| 729.84000 | 35.0 | Cluster_35373 |
| 729.89000 | 35.0 | Cluster_03507 |
| 730.91000 | 35.0 | Cluster_35408 |
| 731.10000 | 35.0 | Cluster_35410 |
| 731.83000 | 35.0 | Cluster_03518 |
| 732.11000 | 35.0 | Cluster_03522 |
| 732.64000 | 35.0 | Cluster_03534 |
| 733.03000 | 35.0 | Cluster_12708 |
| 733.13000 | 35.0 | Cluster_03548 |
| 733.83000 | 35.0 | Cluster_35471 |
| 733.85000 | 35.0 | Cluster_35474 |
| 734.58000 | 35.0 | Cluster_35493 |
| 735.04000 | 35.0 | Cluster_12729 |
| 735.74000 | 35.0 | Cluster_12741 |
| 736.37000 | 35.0 | Cluster_35541 |
| 737.04000 | 35.0 | Cluster_12754 |
| 737.16000 | 35.0 | Cluster_35560 |
| 737.82000 | 35.0 | Cluster_35593 |

| | | |
|---|---|---|
| 738.39000 | 35.0 | Cluster_35611 |
| 738.63000 | 35.0 | Cluster_03590 |
| 739.38000 | 35.0 | Cluster_12791 |
| 739.39000 | 35.0 | Cluster_12789 |
| 739.84000 | 35.0 | Cluster_35645 |
| 740.37000 | 35.0 | Cluster_35657 |
| 740.38000 | 35.0 | Cluster_35661 |
| 740.93000 | 35.0 | Cluster_35676 |
| 741.85000 | 35.0 | Cluster_35683 |
| 742.37000 | 35.0 | Cluster_35697 |
| 742.67000 | 35.0 | Cluster_12831 |
| 742.84000 | 35.0 | Cluster_35707 |
| 744.37000 | 35.0 | Cluster_35750 |
| 745.38000 | 35.0 | Cluster_35777 |
| 745.68000 | 35.0 | Cluster_12857 |
| 746.06000 | 35.0 | Cluster_12863 |
| 746.15000 | 35.0 | Cluster_35792 |
| 746.30000 | 35.0 | Cluster_35797 |
| 747.12000 | 35.0 | Cluster_35817 |
| 747.37000 | 35.0 | Cluster_35818 |
| 748.00000 | 35.0 | Cluster_12903 |
| 748.86000 | 35.0 | Cluster_03654 |
| 748.88000 | 35.0 | Cluster_35857 |
| 749.07000 | 35.0 | Cluster_12920 |
| 749.69000 | 35.0 | Cluster_12929 |
| 749.98000 | 35.0 | Cluster_12936 |
| 750.01000 | 35.0 | Cluster_12941 |
| 750.38000 | 35.0 | Cluster_35883 |
| 750.44000 | 35.0 | Cluster_35886 |
| 750.94000 | 35.0 | Cluster_35903 |
| 751.06000 | 35.0 | Cluster_12961 |
| 751.15000 | 35.0 | Cluster_03662 |
| 751.36000 | 35.0 | Cluster_35908 |
| 752.37000 | 35.0 | Cluster_35932 |
| 753.05000 | 35.0 | Cluster_12988 |
| 753.06000 | 35.0 | Cluster_12984 |
| 753.39000 | 35.0 | Cluster_35964 |
| 753.91000 | 35.0 | Cluster_35981 |
| 754.06000 | 35.0 | Cluster_13009 |
| 754.90000 | 35.0 | Cluster_36004 |
| 755.06000 | 35.0 | Cluster_13021 |
| 755.30000 | 35.0 | Cluster_13019 |
| 755.33000 | 35.0 | Cluster_13020 |
| 755.42000 | 35.0 | Cluster_36013 |
| 755.75000 | 35.0 | Cluster_13034 |
| 755.92000 | 35.0 | Cluster_36043 |
| 756.13000 | 35.0 | Cluster_03688 |
| 757.31000 | 35.0 | Cluster_36071 |
| 757.73000 | 35.0 | Cluster_13076 |
| 758.41000 | 35.0 | Cluster_36102 |
| 758.70000 | 35.0 | Cluster_13088 |

| | | |
|---|---|---|
| 759.63000 | 35.0 | Cluster_03704 |
| 759.90000 | 35.0 | Cluster_36129 |
| 759.91000 | 35.0 | Cluster_36132 |
| 760.35000 | 35.0 | Cluster_36146 |
| 761.05000 | 35.0 | Cluster_00024 |
| 761.41000 | 35.0 | Cluster_36182 |
| 762.38000 | 35.0 | Cluster_13130 |
| 763.03000 | 35.0 | Cluster_13141 |
| 763.08000 | 35.0 | Cluster_13137 |
| 763.35000 | 35.0 | Cluster_03726 |
| 763.71000 | 35.0 | Cluster_13147 |
| 764.09000 | 35.0 | Cluster_13148 |
| 764.36000 | 35.0 | Cluster_36253 |
| 765.15000 | 35.0 | Cluster_03741 |
| 765.79000 | 35.0 | Cluster_36274 |
| 765.80000 | 35.0 | Cluster_36275 |
| 766.37000 | 35.0 | Cluster_36292 |
| 766.55000 | 35.0 | Cluster_00339 |
| 767.70000 | 35.0 | Cluster_13203 |
| 767.85000 | 35.0 | Cluster_36325 |
| 768.07000 | 35.0 | Cluster_13209 |
| 768.08000 | 35.0 | Cluster_13205 |
| 768.38000 | 35.0 | Cluster_36351 |
| 769.33000 | 35.0 | Cluster_36370 |
| 769.59000 | 35.0 | Cluster_36368 |
| 769.82000 | 35.0 | Cluster_36385 |
| 772.41000 | 35.0 | Cluster_36441 |
| 772.74000 | 35.0 | Cluster_13265 |
| 772.80000 | 35.0 | Cluster_13273 |
| 772.88000 | 35.0 | Cluster_36459 |
| 772.92000 | 35.0 | Cluster_36460 |
| 773.08000 | 35.0 | Cluster_03785 |
| 773.09000 | 35.0 | Cluster_03786 |
| 773.13000 | 35.0 | Cluster_13272 |
| 773.35000 | 35.0 | Cluster_13283 |
| 773.41000 | 35.0 | Cluster_13286 |
| 774.12000 | 35.0 | Cluster_36481 |
| 774.73000 | 35.0 | Cluster_13306 |
| 775.06000 | 35.0 | Cluster_13312 |
| 775.07000 | 35.0 | Cluster_13311 |
| 775.42000 | 35.0 | Cluster_36510 |
| 776.37000 | 35.0 | Cluster_03803 |
| 777.15000 | 35.0 | Cluster_03806 |
| 777.35000 | 35.0 | Cluster_36561 |
| 777.38000 | 35.0 | Cluster_36567 |
| 777.39000 | 35.0 | Cluster_36564 |
| 778.37000 | 35.0 | Cluster_36591 |
| 778.77000 | 35.0 | Cluster_13350 |
| 778.87000 | 35.0 | Cluster_36605 |
| 780.73000 | 35.0 | Cluster_13379 |
| 780.81000 | 35.0 | Cluster_36636 |

| | | |
|---|---|---|
| 780.88000 | 35.0 | Cluster_03821 |
| 781.40000 | 35.0 | Cluster_36661 |
| 782.03000 | 35.0 | Cluster_13403 |
| 783.13000 | 35.0 | Cluster_03834 |
| 783.46000 | 35.0 | Cluster_36708 |
| 783.72000 | 35.0 | Cluster_13422 |
| 784.06000 | 35.0 | Cluster_13433 |
| 784.89000 | 35.0 | Cluster_36733 |
| 784.93000 | 35.0 | Cluster_36732 |
| 785.45000 | 35.0 | Cluster_36744 |
| 785.83000 | 35.0 | Cluster_36763 |
| 787.06000 | 35.0 | Cluster_13459 |
| 787.12000 | 35.0 | Cluster_03868 |
| 787.35000 | 35.0 | Cluster_03870 |
| 788.87000 | 35.0 | Cluster_36840 |
| 789.06000 | 35.0 | Cluster_13483 |
| 790.87000 | 35.0 | Cluster_36906 |
| 791.12000 | 35.0 | Cluster_36909 |
| 791.38000 | 35.0 | Cluster_36914 |
| 791.90000 | 35.0 | Cluster_36919 |
| 792.38000 | 35.0 | Cluster_36933 |
| 792.64000 | 35.0 | Cluster_36941 |
| 792.86000 | 35.0 | Cluster_03914 |
| 792.87000 | 35.0 | Cluster_36948 |
| 792.97000 | 35.0 | Cluster_13541 |
| 793.78000 | 35.0 | Cluster_13547 |
| 794.36000 | 35.0 | Cluster_36984 |
| 794.37000 | 35.0 | Cluster_36982 |
| 794.45000 | 35.0 | Cluster_36985 |
| 794.65000 | 35.0 | Cluster_36986 |
| 795.07000 | 35.0 | Cluster_13571 |
| 795.42000 | 35.0 | Cluster_37004 |
| 795.44000 | 35.0 | Cluster_37000 |
| 795.68000 | 35.0 | Cluster_13585 |
| 795.87000 | 35.0 | Cluster_03947 |
| 795.90000 | 35.0 | Cluster_37017 |
| 795.96000 | 35.0 | Cluster_37020 |
| 796.41000 | 35.0 | Cluster_13605 |
| 796.72000 | 35.0 | Cluster_13597 |
| 796.91000 | 35.0 | Cluster_37044 |
| 797.86000 | 35.0 | Cluster_37077 |
| 797.87000 | 35.0 | Cluster_37073 |
| 797.88000 | 35.0 | Cluster_37072 |
| 797.89000 | 35.0 | Cluster_37068 |
| 799.38000 | 35.0 | Cluster_37115 |
| 799.40000 | 35.0 | Cluster_37112 |
| 799.89000 | 35.0 | Cluster_03975 |
| 800.35000 | 35.0 | Cluster_37146 |
| 800.41000 | 35.0 | Cluster_13643 |
| 800.84000 | 35.0 | Cluster_37149 |
| 800.86000 | 35.0 | Cluster_03991 |

| | | |
|---|---|---|
| 801.06000 | 35.0 | Cluster_00029 |
| 801.37000 | 35.0 | Cluster_03995 |
| 801.42000 | 35.0 | Cluster_37163 |
| 801.44000 | 35.0 | Cluster_13660 |
| 802.43000 | 35.0 | Cluster_37188 |
| 802.74000 | 35.0 | Cluster_13681 |
| 803.41000 | 35.0 | Cluster_13684 |
| 803.88000 | 35.0 | Cluster_37226 |
| 804.43000 | 35.0 | Cluster_37243 |
| 804.91000 | 35.0 | Cluster_04015 |
| 805.16000 | 35.0 | Cluster_37269 |
| 805.39000 | 35.0 | Cluster_37283 |
| 805.94000 | 35.0 | Cluster_37284 |
| 806.39000 | 35.0 | Cluster_37304 |
| 807.01000 | 35.0 | Cluster_13723 |
| 807.46000 | 35.0 | Cluster_37330 |
| 807.92000 | 35.0 | Cluster_37352 |
| 808.85000 | 35.0 | Cluster_37370 |
| 808.91000 | 35.0 | Cluster_37377 |
| 809.40000 | 35.0 | Cluster_04037 |
| 809.91000 | 35.0 | Cluster_37401 |
| 810.39000 | 35.0 | Cluster_37407 |
| 810.44000 | 35.0 | Cluster_13774 |
| 811.74000 | 35.0 | Cluster_00036 |
| 811.86000 | 35.0 | Cluster_37446 |
| 811.91000 | 35.0 | Cluster_37441 |
| 812.39000 | 35.0 | Cluster_37458 |
| 812.41000 | 35.0 | Cluster_13796 |
| 813.82000 | 35.0 | Cluster_00352 |
| 813.90000 | 35.0 | Cluster_37483 |
| 814.10000 | 35.0 | Cluster_13827 |
| 814.16000 | 35.0 | Cluster_37492 |
| 814.17000 | 35.0 | Cluster_04067 |
| 814.90000 | 35.0 | Cluster_04074 |
| 815.06000 | 35.0 | Cluster_13832 |
| 815.79000 | 35.0 | Cluster_13844 |
| 815.92000 | 35.0 | Cluster_37530 |
| 816.38000 | 35.0 | Cluster_37540 |
| 816.43000 | 35.0 | Cluster_04088 |
| 816.72000 | 35.0 | Cluster_13857 |
| 817.34000 | 35.0 | Cluster_37561 |
| 817.88000 | 35.0 | Cluster_37570 |
| 818.42000 | 35.0 | Cluster_13883 |
| 819.05000 | 35.0 | Cluster_13897 |
| 819.42000 | 35.0 | Cluster_13906 |
| 820.17000 | 35.0 | Cluster_04104 |
| 821.23000 | 35.0 | Cluster_04111 |
| 821.38000 | 35.0 | Cluster_37637 |
| 822.06000 | 35.0 | Cluster_13934 |
| 823.42000 | 35.0 | Cluster_37669 |
| 823.91000 | 35.0 | Cluster_37674 |

| | | |
|---|---|---|
| 823.94000 | 35.0 | Cluster_37688 |
| 825.32000 | 35.0 | Cluster_37717 |
| 825.39000 | 35.0 | Cluster_04137 |
| 826.38000 | 35.0 | Cluster_13981 |
| 826.86000 | 35.0 | Cluster_37765 |
| 827.04000 | 35.0 | Cluster_13986 |
| 827.06000 | 35.0 | Cluster_13988 |
| 827.09000 | 35.0 | Cluster_13989 |
| 827.88000 | 35.0 | Cluster_37794 |
| 828.80000 | 35.0 | Cluster_14000 |
| 828.98000 | 35.0 | Cluster_14004 |
| 829.16000 | 35.0 | Cluster_04165 |
| 829.36000 | 35.0 | Cluster_37821 |
| 829.43000 | 35.0 | Cluster_37825 |
| 829.91000 | 35.0 | Cluster_04169 |
| 830.74000 | 35.0 | Cluster_14020 |
| 830.89000 | 35.0 | Cluster_00042 |
| 831.45000 | 35.0 | Cluster_37853 |
| 832.41000 | 35.0 | Cluster_37873 |
| 832.68000 | 35.0 | Cluster_37879 |
| 833.49000 | 35.0 | Cluster_37894 |
| 833.99000 | 35.0 | Cluster_37902 |
| 834.10000 | 35.0 | Cluster_14054 |
| 834.94000 | 35.0 | Cluster_37918 |
| 835.05000 | 35.0 | Cluster_14062 |
| 836.39000 | 35.0 | Cluster_14076 |
| 836.42000 | 35.0 | Cluster_14077 |
| 836.76000 | 35.0 | Cluster_14083 |
| 836.86000 | 35.0 | Cluster_37959 |
| 836.94000 | 35.0 | Cluster_04234 |
| 837.41000 | 35.0 | Cluster_37970 |
| 837.95000 | 35.0 | Cluster_37976 |
| 839.10000 | 35.0 | Cluster_14102 |
| 840.66000 | 35.0 | Cluster_04260 |
| 840.78000 | 35.0 | Cluster_14117 |
| 841.08000 | 35.0 | Cluster_14128 |
| 841.10000 | 35.0 | Cluster_14126 |
| 841.67000 | 35.0 | Cluster_00364 |
| 841.76000 | 35.0 | Cluster_38066 |
| 842.73000 | 35.0 | Cluster_14140 |
| 843.87000 | 35.0 | Cluster_38114 |
| 844.18000 | 35.0 | Cluster_04298 |
| 844.44000 | 35.0 | Cluster_14157 |
| 846.98000 | 35.0 | Cluster_38180 |
| 847.48000 | 35.0 | Cluster_38200 |
| 847.75000 | 35.0 | Cluster_14185 |
| 847.88000 | 35.0 | Cluster_38203 |
| 848.40000 | 35.0 | Cluster_04350 |
| 848.91000 | 35.0 | Cluster_04352 |
| 849.46000 | 35.0 | Cluster_38235 |
| 849.64000 | 35.0 | Cluster_04364 |

| | | |
|---|---|---|
| 849.97000 | 35.0 | Cluster_38245 |
| 849.99000 | 35.0 | Cluster_38248 |
| 850.33000 | 35.0 | Cluster_38250 |
| 850.40000 | 35.0 | Cluster_04376 |
| 850.76000 | 35.0 | Cluster_14213 |
| 851.36000 | 35.0 | Cluster_14215 |
| 851.77000 | 35.0 | Cluster_38298 |
| 852.03000 | 35.0 | Cluster_38307 |
| 852.91000 | 35.0 | Cluster_04400 |
| 852.93000 | 35.0 | Cluster_04401 |
| 853.45000 | 35.0 | Cluster_38334 |
| 854.44000 | 35.0 | Cluster_38377 |
| 854.47000 | 35.0 | Cluster_38375 |
| 854.88000 | 35.0 | Cluster_38385 |
| 855.41000 | 35.0 | Cluster_38392 |
| 855.73000 | 35.0 | Cluster_14240 |
| 856.43000 | 35.0 | Cluster_14249 |
| 856.91000 | 35.0 | Cluster_04444 |
| 857.06000 | 35.0 | Cluster_14257 |
| 857.89000 | 35.0 | Cluster_38459 |
| 857.91000 | 35.0 | Cluster_04449 |
| 858.07000 | 35.0 | Cluster_14269 |
| 858.77000 | 35.0 | Cluster_14275 |
| 859.45000 | 35.0 | Cluster_38500 |
| 859.47000 | 35.0 | Cluster_38497 |
| 860.47000 | 35.0 | Cluster_14284 |
| 861.39000 | 35.0 | Cluster_04493 |
| 862.39000 | 35.0 | Cluster_04505 |
| 862.49000 | 35.0 | Cluster_38591 |
| 863.05000 | 35.0 | Cluster_14302 |
| 863.76000 | 35.0 | Cluster_14307 |
| 863.94000 | 35.0 | Cluster_38615 |
| 864.07000 | 35.0 | Cluster_14314 |
| 865.03000 | 35.0 | Cluster_38646 |
| 865.75000 | 35.0 | Cluster_14326 |
| 865.92000 | 35.0 | Cluster_38659 |
| 865.93000 | 35.0 | Cluster_04555 |
| 866.41000 | 35.0 | Cluster_14334 |
| 869.07000 | 35.0 | Cluster_14353 |
| 869.28000 | 35.0 | Cluster_38719 |
| 869.41000 | 35.0 | Cluster_04599 |
| 869.78000 | 35.0 | Cluster_14363 |
| 871.17000 | 35.0 | Cluster_38758 |
| 871.27000 | 35.0 | Cluster_38759 |
| 871.44000 | 35.0 | Cluster_38769 |
| 871.47000 | 35.0 | Cluster_14388 |
| 872.27000 | 35.0 | Cluster_38802 |
| 874.09000 | 35.0 | Cluster_14414 |
| 874.26000 | 35.0 | Cluster_38844 |
| 874.78000 | 35.0 | Cluster_14424 |
| 875.28000 | 35.0 | Cluster_38860 |

| | | |
|---|---|---|
| 875.66000 | 35.0 | Cluster_04698 |
| 876.08000 | 35.0 | Cluster_14437 |
| 876.53000 | 35.0 | Cluster_38895 |
| 877.38000 | 35.0 | Cluster_14453 |
| 877.44000 | 35.0 | Cluster_38909 |
| 878.48000 | 35.0 | Cluster_38928 |
| 880.46000 | 35.0 | Cluster_14483 |
| 881.96000 | 35.0 | Cluster_38993 |
| 882.44000 | 35.0 | Cluster_39008 |
| 882.94000 | 35.0 | Cluster_39018 |
| 884.40000 | 35.0 | Cluster_14528 |
| 884.66000 | 35.0 | Cluster_39047 |
| 885.12000 | 35.0 | Cluster_14544 |
| 885.68000 | 35.0 | Cluster_04765 |
| 885.95000 | 35.0 | Cluster_39078 |
| 886.43000 | 35.0 | Cluster_39095 |
| 886.96000 | 35.0 | Cluster_39099 |
| 887.52000 | 35.0 | Cluster_39107 |
| 887.68000 | 35.0 | Cluster_04781 |
| 887.74000 | 35.0 | Cluster_14571 |
| 888.13000 | 35.0 | Cluster_14577 |
| 888.40000 | 35.0 | Cluster_14578 |
| 888.91000 | 35.0 | Cluster_39135 |
| 888.92000 | 35.0 | Cluster_04786 |
| 889.11000 | 35.0 | Cluster_14581 |
| 889.44000 | 35.0 | Cluster_39147 |
| 890.17000 | 35.0 | Cluster_04796 |
| 890.38000 | 35.0 | Cluster_39161 |
| 890.39000 | 35.0 | Cluster_39160 |
| 890.46000 | 35.0 | Cluster_39173 |
| 890.99000 | 35.0 | Cluster_39180 |
| 891.46000 | 35.0 | Cluster_39186 |
| 891.81000 | 35.0 | Cluster_14598 |
| 892.21000 | 35.0 | Cluster_04824 |
| 893.43000 | 35.0 | Cluster_39231 |
| 893.78000 | 35.0 | Cluster_14615 |
| 894.12000 | 35.0 | Cluster_14617 |
| 894.71000 | 35.0 | Cluster_04845 |
| 895.21000 | 35.0 | Cluster_04851 |
| 895.45000 | 35.0 | Cluster_14622 |
| 895.81000 | 35.0 | Cluster_14625 |
| 896.15000 | 35.0 | Cluster_14634 |
| 897.04000 | 35.0 | Cluster_39292 |
| 898.77000 | 35.0 | Cluster_14664 |
| 899.43000 | 35.0 | Cluster_39337 |
| 899.51000 | 35.0 | Cluster_39345 |
| 899.96000 | 35.0 | Cluster_04891 |
| 900.94000 | 35.0 | Cluster_39368 |
| 900.97000 | 35.0 | Cluster_39380 |
| 900.99000 | 35.0 | Cluster_39377 |
| 902.46000 | 35.0 | Cluster_39412 |

| | | |
|---|---|---|
| 904.46000 | 35.0 | Cluster_39443 |
| 905.43000 | 35.0 | Cluster_39461 |
| 906.77000 | 35.0 | Cluster_14739 |
| 907.80000 | 35.0 | Cluster_14749 |
| 908.46000 | 35.0 | Cluster_39500 |
| 908.48000 | 35.0 | Cluster_14756 |
| 908.94000 | 35.0 | Cluster_04958 |
| 908.95000 | 35.0 | Cluster_39504 |
| 909.44000 | 35.0 | Cluster_14760 |
| 909.50000 | 35.0 | Cluster_14765 |
| 909.89000 | 35.0 | Cluster_39520 |
| 910.11000 | 35.0 | Cluster_14769 |
| 910.19000 | 35.0 | Cluster_04966 |
| 911.96000 | 35.0 | Cluster_39564 |
| 914.56000 | 35.0 | Cluster_39615 |
| 915.13000 | 35.0 | Cluster_14823 |
| 915.93000 | 35.0 | Cluster_39626 |
| 916.44000 | 35.0 | Cluster_39640 |
| 916.49000 | 35.0 | Cluster_39633 |
| 917.45000 | 35.0 | Cluster_14854 |
| 917.97000 | 35.0 | Cluster_39674 |
| 918.66000 | 35.0 | Cluster_39698 |
| 920.45000 | 35.0 | Cluster_39721 |
| 920.76000 | 35.0 | Cluster_14891 |
| 920.79000 | 35.0 | Cluster_39725 |
| 920.88000 | 35.0 | Cluster_39726 |
| 920.95000 | 35.0 | Cluster_39731 |
| 922.42000 | 35.0 | Cluster_14907 |
| 922.98000 | 35.0 | Cluster_39772 |
| 923.30000 | 35.0 | Cluster_14924 |
| 923.45000 | 35.0 | Cluster_14932 |
| 923.92000 | 35.0 | Cluster_39782 |
| 924.49000 | 35.0 | Cluster_39788 |
| 925.11000 | 35.0 | Cluster_14961 |
| 926.42000 | 35.0 | Cluster_14975 |
| 927.48000 | 35.0 | Cluster_39834 |
| 928.47000 | 35.0 | Cluster_39858 |
| 928.96000 | 35.0 | Cluster_39861 |
| 929.12000 | 35.0 | Cluster_15009 |
| 929.43000 | 35.0 | Cluster_39880 |
| 929.90000 | 35.0 | Cluster_39897 |
| 930.47000 | 35.0 | Cluster_15020 |
| 931.73000 | 35.0 | Cluster_15043 |
| 931.90000 | 35.0 | Cluster_39924 |
| 932.64000 | 35.0 | Cluster_15062 |
| 932.95000 | 35.0 | Cluster_39941 |
| 933.14000 | 35.0 | Cluster_15072 |
| 933.33000 | 35.0 | Cluster_05038 |
| 934.00000 | 35.0 | Cluster_39963 |
| 934.13000 | 35.0 | Cluster_15084 |
| 934.43000 | 35.0 | Cluster_15083 |

| | | |
|---|---|---|
| 934.78000 | 35.0 | Cluster_15089 |
| 935.78000 | 35.0 | Cluster_15100 |
| 936.45000 | 35.0 | Cluster_15116 |
| 936.80000 | 35.0 | Cluster_15129 |
| 937.00000 | 35.0 | Cluster_39997 |
| 937.44000 | 35.0 | Cluster_15144 |
| 937.49000 | 35.0 | Cluster_05051 |
| 938.07000 | 35.0 | Cluster_15154 |
| 938.13000 | 35.0 | Cluster_15156 |
| 938.21000 | 35.0 | Cluster_15160 |
| 939.69000 | 35.0 | Cluster_05058 |
| 940.46000 | 35.0 | Cluster_15180 |
| 941.72000 | 35.0 | Cluster_40098 |
| 942.00000 | 35.0 | Cluster_40094 |
| 943.45000 | 35.0 | Cluster_15229 |
| 945.13000 | 35.0 | Cluster_15244 |
| 945.39000 | 35.0 | Cluster_15242 |
| 945.72000 | 35.0 | Cluster_05075 |
| 945.73000 | 35.0 | Cluster_05078 |
| 945.78000 | 35.0 | Cluster_15250 |
| 945.98000 | 35.0 | Cluster_40152 |
| 947.16000 | 35.0 | Cluster_15264 |
| 947.22000 | 35.0 | Cluster_05084 |
| 947.79000 | 35.0 | Cluster_15273 |
| 948.19000 | 35.0 | Cluster_15277 |
| 948.52000 | 35.0 | Cluster_40180 |
| 949.14000 | 35.0 | Cluster_15301 |
| 949.44000 | 35.0 | Cluster_40190 |
| 949.46000 | 35.0 | Cluster_15302 |
| 950.44000 | 35.0 | Cluster_15314 |
| 951.17000 | 35.0 | Cluster_15324 |
| 951.96000 | 35.0 | Cluster_40227 |
| 952.14000 | 35.0 | Cluster_15337 |
| 952.15000 | 35.0 | Cluster_15339 |
| 952.44000 | 35.0 | Cluster_15342 |
| 952.53000 | 35.0 | Cluster_40233 |
| 953.98000 | 35.0 | Cluster_40255 |
| 954.82000 | 35.0 | Cluster_15379 |
| 954.99000 | 35.0 | Cluster_40262 |
| 955.13000 | 35.0 | Cluster_15394 |
| 955.47000 | 35.0 | Cluster_15404 |
| 956.48000 | 35.0 | Cluster_15426 |
| 957.07000 | 35.0 | Cluster_40290 |
| 957.48000 | 35.0 | Cluster_40291 |
| 960.15000 | 35.0 | Cluster_15462 |
| 960.98000 | 35.0 | Cluster_40317 |
| 961.48000 | 35.0 | Cluster_15478 |
| 962.14000 | 35.0 | Cluster_15497 |
| 962.38000 | 35.0 | Cluster_40344 |
| 962.79000 | 35.0 | Cluster_15505 |
| 964.13000 | 35.0 | Cluster_15523 |

| | | |
|---|---|---|
| 964.43000 | 35.0 | Cluster_40365 |
| 965.29000 | 35.0 | Cluster_40377 |
| 966.15000 | 35.0 | Cluster_15549 |
| 966.49000 | 35.0 | Cluster_40387 |
| 966.50000 | 35.0 | Cluster_05172 |
| 966.85000 | 35.0 | Cluster_15552 |
| 967.00000 | 35.0 | Cluster_40392 |
| 967.56000 | 35.0 | Cluster_15565 |
| 968.03000 | 35.0 | Cluster_40402 |
| 968.25000 | 35.0 | Cluster_05187 |
| 971.97000 | 35.0 | Cluster_40451 |
| 972.01000 | 35.0 | Cluster_40467 |
| 972.46000 | 35.0 | Cluster_15627 |
| 972.49000 | 35.0 | Cluster_15626 |
| 973.51000 | 35.0 | Cluster_40476 |
| 974.47000 | 35.0 | Cluster_15651 |
| 974.98000 | 35.0 | Cluster_40495 |
| 975.82000 | 35.0 | Cluster_15672 |
| 975.98000 | 35.0 | Cluster_05210 |
| 976.00000 | 35.0 | Cluster_05212 |
| 976.03000 | 35.0 | Cluster_40504 |
| 977.14000 | 35.0 | Cluster_15696 |
| 977.62000 | 35.0 | Cluster_40522 |
| 979.45000 | 35.0 | Cluster_40546 |
| 979.80000 | 35.0 | Cluster_15720 |
| 979.83000 | 35.0 | Cluster_15721 |
| 979.98000 | 35.0 | Cluster_40568 |
| 980.00000 | 35.0 | Cluster_40562 |
| 980.50000 | 35.0 | Cluster_40565 |
| 980.70000 | 35.0 | Cluster_40578 |
| 981.45000 | 35.0 | Cluster_15750 |
| 983.25000 | 35.0 | Cluster_05237 |
| 983.95000 | 35.0 | Cluster_40617 |
| 985.14000 | 35.0 | Cluster_40628 |
| 986.02000 | 35.0 | Cluster_40630 |
| 986.52000 | 35.0 | Cluster_40639 |
| 986.83000 | 35.0 | Cluster_15810 |
| 987.24000 | 35.0 | Cluster_40654 |
| 987.80000 | 35.0 | Cluster_15825 |
| 988.44000 | 35.0 | Cluster_15835 |
| 991.93000 | 35.0 | Cluster_40717 |
| 993.50000 | 35.0 | Cluster_15895 |
| 997.19000 | 35.0 | Cluster_15954 |
| 1000.06000 | 35.0 | Cluster_40851 |
| 1001.03000 | 35.0 | Cluster_40878 |
| 1001.16000 | 35.0 | Cluster_15996 |
| 1001.49000 | 35.0 | Cluster_40884 |
| 1003.52000 | 35.0 | Cluster_05294 |
| 1004.00000 | 35.0 | Cluster_40937 |
| 1004.31000 | 35.0 | Cluster_05296 |
| 1004.57000 | 35.0 | Cluster_40940 |

| | | |
|---|---|---|
| 1007.81000 | 35.0 | Cluster_16072 |
| 1008.78000 | 35.0 | Cluster_16080 |
| 1012.26000 | 35.0 | Cluster_05329 |
| 1012.27000 | 35.0 | Cluster_41044 |
| 1012.93000 | 35.0 | Cluster_41054 |
| 1013.01000 | 35.0 | Cluster_05341 |
| 1013.47000 | 35.0 | Cluster_16138 |
| 1013.49000 | 35.0 | Cluster_16137 |
| 1013.76000 | 35.0 | Cluster_05344 |
| 1013.90000 | 35.0 | Cluster_41056 |
| 1015.92000 | 35.0 | Cluster_41088 |
| 1016.78000 | 35.0 | Cluster_05351 |
| 1017.27000 | 35.0 | Cluster_41099 |
| 1019.25000 | 35.0 | Cluster_05360 |
| 1020.15000 | 35.0 | Cluster_16210 |
| 1022.50000 | 35.0 | Cluster_16231 |
| 1023.82000 | 35.0 | Cluster_41185 |
| 1024.48000 | 35.0 | Cluster_16254 |
| 1024.50000 | 35.0 | Cluster_16256 |
| 1024.93000 | 35.0 | Cluster_41195 |
| 1025.46000 | 35.0 | Cluster_16264 |
| 1025.81000 | 35.0 | Cluster_41211 |
| 1026.83000 | 35.0 | Cluster_16274 |
| 1027.13000 | 35.0 | Cluster_16285 |
| 1027.14000 | 35.0 | Cluster_16284 |
| 1027.88000 | 35.0 | Cluster_16291 |
| 1028.18000 | 35.0 | Cluster_16295 |
| 1028.84000 | 35.0 | Cluster_16300 |
| 1029.99000 | 35.0 | Cluster_41262 |
| 1031.20000 | 35.0 | Cluster_16323 |
| 1032.82000 | 35.0 | Cluster_16335 |
| 1033.20000 | 35.0 | Cluster_16342 |
| 1036.52000 | 35.0 | Cluster_16382 |
| 1037.50000 | 35.0 | Cluster_16400 |
| 1039.05000 | 35.0 | Cluster_41371 |
| 1039.52000 | 35.0 | Cluster_41374 |
| 1039.81000 | 35.0 | Cluster_16429 |
| 1040.83000 | 35.0 | Cluster_05437 |
| 1041.01000 | 35.0 | Cluster_41401 |
| 1042.17000 | 35.0 | Cluster_16457 |
| 1044.49000 | 35.0 | Cluster_41432 |
| 1046.13000 | 35.0 | Cluster_16488 |
| 1046.22000 | 35.0 | Cluster_16495 |
| 1047.16000 | 35.0 | Cluster_16507 |
| 1047.83000 | 35.0 | Cluster_41470 |
| 1053.50000 | 35.0 | Cluster_16581 |
| 1053.98000 | 35.0 | Cluster_41534 |
| 1054.99000 | 35.0 | Cluster_05476 |
| 1055.22000 | 35.0 | Cluster_16595 |
| 1055.50000 | 35.0 | Cluster_16606 |
| 1056.52000 | 35.0 | Cluster_16616 |

| | | |
|---|---|---|
| 1057.06000 | 35.0 | Cluster_41567 |
| 1057.33000 | 35.0 | Cluster_05486 |
| 1057.34000 | 35.0 | Cluster_05487 |
| 1057.58000 | 35.0 | Cluster_41574 |
| 1057.99000 | 35.0 | Cluster_41578 |
| 1058.02000 | 35.0 | Cluster_41582 |
| 1058.81000 | 35.0 | Cluster_16658 |
| 1058.85000 | 35.0 | Cluster_16649 |
| 1058.96000 | 35.0 | Cluster_41595 |
| 1058.97000 | 35.0 | Cluster_41597 |
| 1059.01000 | 35.0 | Cluster_41596 |
| 1060.32000 | 35.0 | Cluster_41612 |
| 1060.75000 | 35.0 | Cluster_05502 |
| 1060.82000 | 35.0 | Cluster_16679 |
| 1061.26000 | 35.0 | Cluster_05510 |
| 1061.33000 | 35.0 | Cluster_05506 |
| 1061.82000 | 35.0 | Cluster_41625 |
| 1062.83000 | 35.0 | Cluster_16700 |
| 1063.27000 | 35.0 | Cluster_05516 |
| 1063.82000 | 35.0 | Cluster_41655 |
| 1063.99000 | 35.0 | Cluster_41663 |
| 1064.72000 | 35.0 | Cluster_05521 |
| 1064.73000 | 35.0 | Cluster_41675 |
| 1064.99000 | 35.0 | Cluster_41673 |
| 1066.26000 | 35.0 | Cluster_41702 |
| 1067.52000 | 35.0 | Cluster_41715 |
| 1068.05000 | 35.0 | Cluster_41719 |
| 1068.75000 | 35.0 | Cluster_41734 |
| 1069.51000 | 35.0 | Cluster_41748 |
| 1070.16000 | 35.0 | Cluster_16781 |
| 1070.51000 | 35.0 | Cluster_05546 |
| 1072.55000 | 35.0 | Cluster_16804 |
| 1072.76000 | 35.0 | Cluster_05555 |
| 1074.19000 | 35.0 | Cluster_16831 |
| 1074.51000 | 35.0 | Cluster_41810 |
| 1075.53000 | 35.0 | Cluster_41816 |
| 1076.75000 | 35.0 | Cluster_05574 |
| 1078.01000 | 35.0 | Cluster_05578 |
| 1079.49000 | 35.0 | Cluster_05582 |
| 1080.75000 | 35.0 | Cluster_00593 |
| 1082.00000 | 35.0 | Cluster_05588 |
| 1083.50000 | 35.0 | Cluster_05592 |
| 1086.59000 | 35.0 | Cluster_41940 |
| 1091.58000 | 35.0 | Cluster_41998 |
| 1092.90000 | 35.0 | Cluster_17025 |
| 1093.21000 | 35.0 | Cluster_17031 |
| 1093.55000 | 35.0 | Cluster_17035 |
| 1093.58000 | 35.0 | Cluster_42024 |
| 1093.99000 | 35.0 | Cluster_42029 |
| 1094.92000 | 35.0 | Cluster_00625 |
| 1096.55000 | 35.0 | Cluster_42051 |

| | | |
|---|---|---|
| 1100.23000 | 35.0 | Cluster_17091 |
| 1100.56000 | 35.0 | Cluster_17097 |
| 1101.96000 | 35.0 | Cluster_17105 |
| 1102.54000 | 35.0 | Cluster_17111 |
| 1103.12000 | 35.0 | Cluster_42104 |
| 1108.23000 | 35.0 | Cluster_17176 |
| 1108.37000 | 35.0 | Cluster_42148 |
| 1108.88000 | 35.0 | Cluster_17175 |
| 1109.86000 | 35.0 | Cluster_17199 |
| 1113.50000 | 35.0 | Cluster_42184 |
| 1113.53000 | 35.0 | Cluster_17241 |
| 1115.18000 | 35.0 | Cluster_17258 |
| 1115.77000 | 35.0 | Cluster_00660 |
| 1117.22000 | 35.0 | Cluster_17276 |
| 1118.21000 | 35.0 | Cluster_17277 |
| 1118.22000 | 35.0 | Cluster_17280 |
| 1119.87000 | 35.0 | Cluster_42234 |
| 1121.54000 | 35.0 | Cluster_17322 |
| 1123.04000 | 35.0 | Cluster_42270 |
| 1123.97000 | 35.0 | Cluster_42271 |
| 1124.25000 | 35.0 | Cluster_17359 |
| 1125.57000 | 35.0 | Cluster_42283 |
| 1126.07000 | 35.0 | Cluster_42291 |
| 1127.88000 | 35.0 | Cluster_17398 |
| 1128.57000 | 35.0 | Cluster_17409 |
| 1129.21000 | 35.0 | Cluster_17418 |
| 1129.55000 | 35.0 | Cluster_17428 |
| 1129.90000 | 35.0 | Cluster_17433 |
| 1130.23000 | 35.0 | Cluster_17434 |
| 1131.58000 | 35.0 | Cluster_17448 |
| 1133.13000 | 35.0 | Cluster_42337 |
| 1133.20000 | 35.0 | Cluster_17479 |
| 1133.22000 | 35.0 | Cluster_17484 |
| 1134.90000 | 35.0 | Cluster_17503 |
| 1135.23000 | 35.0 | Cluster_17509 |
| 1135.51000 | 35.0 | Cluster_17513 |
| 1135.54000 | 35.0 | Cluster_17519 |
| 1135.55000 | 35.0 | Cluster_17517 |
| 1136.11000 | 35.0 | Cluster_42363 |
| 1136.88000 | 35.0 | Cluster_17525 |
| 1136.89000 | 35.0 | Cluster_17538 |
| 1138.13000 | 35.0 | Cluster_42377 |
| 1140.22000 | 35.0 | Cluster_17567 |
| 1143.18000 | 35.0 | Cluster_00674 |
| 1143.56000 | 35.0 | Cluster_17607 |
| 1143.60000 | 35.0 | Cluster_42409 |
| 1144.20000 | 35.0 | Cluster_17626 |
| 1144.24000 | 35.0 | Cluster_17620 |
| 1145.21000 | 35.0 | Cluster_17645 |
| 1146.57000 | 35.0 | Cluster_00680 |
| 1149.29000 | 35.0 | Cluster_42461 |

| | | |
|---|---|---|
| 1151.05000 | 35.0 | Cluster_42474 |
| 1152.54000 | 35.0 | Cluster_17735 |
| 1152.88000 | 35.0 | Cluster_17750 |
| 1153.03000 | 35.0 | Cluster_05790 |
| 1153.76000 | 35.0 | Cluster_00695 |
| 1154.35000 | 35.0 | Cluster_42485 |
| 1155.34000 | 35.0 | Cluster_42501 |
| 1156.84000 | 35.0 | Cluster_42511 |
| 1158.84000 | 35.0 | Cluster_42519 |
| 1159.61000 | 35.0 | Cluster_42529 |
| 1160.34000 | 35.0 | Cluster_42534 |
| 1162.11000 | 35.0 | Cluster_42551 |
| 1162.64000 | 35.0 | Cluster_42555 |
| 1163.09000 | 35.0 | Cluster_05817 |
| 1166.08000 | 35.0 | Cluster_05827 |
| 1166.14000 | 35.0 | Cluster_42592 |
| 1166.21000 | 35.0 | Cluster_17894 |
| 1167.19000 | 35.0 | Cluster_17908 |
| 1167.57000 | 35.0 | Cluster_17921 |
| 1168.21000 | 35.0 | Cluster_17931 |
| 1168.33000 | 35.0 | Cluster_42608 |
| 1168.58000 | 35.0 | Cluster_05832 |
| 1173.07000 | 35.0 | Cluster_42645 |
| 1180.80000 | 35.0 | Cluster_05888 |
| 1183.23000 | 35.0 | Cluster_18040 |
| 1187.59000 | 35.0 | Cluster_18075 |
| 1190.56000 | 35.0 | Cluster_18091 |
| 1192.81000 | 35.0 | Cluster_42765 |
| 1192.92000 | 35.0 | Cluster_18117 |
| 1195.88000 | 35.0 | Cluster_18133 |
| 1196.63000 | 35.0 | Cluster_42787 |
| 1197.59000 | 35.0 | Cluster_18145 |
| 1197.90000 | 35.0 | Cluster_18140 |
| 1198.64000 | 35.0 | Cluster_42803 |
| 1199.32000 | 35.0 | Cluster_05982 |
| 1199.57000 | 35.0 | Cluster_18161 |
| 1200.26000 | 35.0 | Cluster_18168 |
| 1200.56000 | 35.0 | Cluster_18177 |
| 1201.89000 | 35.0 | Cluster_18192 |
| 1203.44000 | 35.0 | Cluster_42830 |
| 1206.62000 | 35.0 | Cluster_42853 |
| 1208.56000 | 35.0 | Cluster_18237 |
| 1209.51000 | 35.0 | Cluster_42887 |
| 1213.12000 | 35.0 | Cluster_42910 |
| 1215.15000 | 35.0 | Cluster_42924 |
| 1218.11000 | 35.0 | Cluster_42950 |
| 1219.09000 | 35.0 | Cluster_42956 |
| 1221.25000 | 35.0 | Cluster_18299 |
| 1222.10000 | 35.0 | Cluster_42968 |
| 1224.08000 | 35.0 | Cluster_42977 |
| 1224.60000 | 35.0 | Cluster_42983 |

| | | |
|---|---|---|
| 1225.59000 | 35.0 | Cluster_42994 |
| 1228.07000 | 35.0 | Cluster_43008 |
| 1228.11000 | 35.0 | Cluster_43007 |
| 1230.63000 | 35.0 | Cluster_43026 |
| 1232.57000 | 35.0 | Cluster_43045 |
| 1234.58000 | 35.0 | Cluster_43065 |
| 1238.57000 | 35.0 | Cluster_43079 |
| 1245.28000 | 35.0 | Cluster_18390 |
| 1249.16000 | 35.0 | Cluster_43135 |
| 1251.87000 | 35.0 | Cluster_43149 |
| 1253.87000 | 35.0 | Cluster_43162 |
| 1255.86000 | 35.0 | Cluster_43176 |
| 1257.08000 | 35.0 | Cluster_06282 |
| 1264.27000 | 35.0 | Cluster_18463 |
| 1267.20000 | 35.0 | Cluster_43224 |
| 1267.68000 | 35.0 | Cluster_43228 |
| 1274.29000 | 35.0 | Cluster_18502 |
| 1284.15000 | 35.0 | Cluster_43287 |
| 1300.69000 | 35.0 | Cluster_43366 |
| 1300.70000 | 35.0 | Cluster_43360 |
| 1300.73000 | 35.0 | Cluster_43361 |
| 1300.89000 | 35.0 | Cluster_43362 |
| 1301.68000 | 35.0 | Cluster_43365 |
| 1320.21000 | 35.0 | Cluster_43449 |
| 1329.73000 | 35.0 | Cluster_43484 |
| 1330.45000 | 35.0 | Cluster_06499 |
| 1331.69000 | 35.0 | Cluster_43492 |
| 1334.68000 | 35.0 | Cluster_43505 |
| 1349.89000 | 35.0 | Cluster_43562 |
| 1353.71000 | 35.0 | Cluster_43578 |
| 1354.21000 | 35.0 | Cluster_43579 |
| 1354.66000 | 35.0 | Cluster_06571 |
| 1361.03000 | 35.0 | Cluster_18681 |
| 1367.16000 | 35.0 | Cluster_43645 |
| 1372.75000 | 35.0 | Cluster_43667 |
| 1376.18000 | 35.0 | Cluster_43680 |
| 1376.34000 | 35.0 | Cluster_18707 |
| 1383.67000 | 35.0 | Cluster_18716 |
| 1394.65000 | 35.0 | Cluster_43739 |
| 1396.44000 | 35.0 | Cluster_18732 |
| 1402.12000 | 35.0 | Cluster_18741 |
| 1406.24000 | 35.0 | Cluster_43793 |
| 1407.03000 | 35.0 | Cluster_18752 |
| 1414.35000 | 35.0 | Cluster_18768 |
| 1417.37000 | 35.0 | Cluster_18772 |
| 1420.01000 | 35.0 | Cluster_18783 |
| 1422.96000 | 35.0 | Cluster_43843 |
| 1423.47000 | 35.0 | Cluster_43844 |
| 1424.69000 | 35.0 | Cluster_18789 |
| 1424.79000 | 35.0 | Cluster_43851 |
| 1425.27000 | 35.0 | Cluster_43853 |

| | | |
|---|---|---|
| 1427.03000 | 35.0 | Cluster_18794 |
| 1428.82000 | 35.0 | Cluster_43862 |
| 1429.02000 | 35.0 | Cluster_18795 |
| 1430.01000 | 35.0 | Cluster_18796 |
| 1431.68000 | 35.0 | Cluster_18799 |
| 1437.03000 | 35.0 | Cluster_18803 |
| 1439.35000 | 35.0 | Cluster_18807 |
| 1459.27000 | 35.0 | Cluster_43959 |
| 1477.30000 | 35.0 | Cluster_44015 |
| 1489.22000 | 35.0 | Cluster_44043 |
| 1492.94000 | 35.0 | Cluster_44054 |
| 1495.14000 | 35.0 | Cluster_44065 |
| 1511.80000 | 35.0 | Cluster_44102 |
| 1518.72000 | 35.0 | Cluster_44112 |
| 1530.33000 | 35.0 | Cluster_44125 |
| 1560.76000 | 35.0 | Cluster_44175 |
| 1564.32000 | 35.0 | Cluster_44184 |
| 1571.76000 | 35.0 | Cluster_44194 |
| 1582.85000 | 35.0 | Cluster_44218 |
| 1616.85000 | 35.0 | Cluster_44250 |
| 1633.97000 | 35.0 | Cluster_44257 |
| 1635.40000 | 35.0 | Cluster_44260 |
| 1676.82000 | 35.0 | Cluster_44320 |
| 1683.84000 | 35.0 | Cluster_44341 |
| 1687.92000 | 35.0 | Cluster_44353 |
| 1739.94000 | 35.0 | Cluster_44403 |
| 1792.99000 | 35.0 | Cluster_44425 |

Reject Mass List: (none)
Neutral Loss Mass List: (none)
Product Mass List: (none)
Neutral loss in top: 3
Product in top: 3
Most intense if no parent masses found enabled
Add/subtract mass not enabled
FT master scan preview mode enabled
Charge state screening enabled
Charge state dependent ETD time not enabled
Monoisotopic precursor selection enabled
Charge state rejection enabled
Unassigned charge states : rejected
Charge state 1 : rejected
Charge state 2 : not rejected
Charge state 3 : not rejected
Charge states 4+ : not rejected
Chromatography mode is disabled

Global Data Dependent Settings:
Predict ion injection time enabled
Use global parent and reject mass lists not enabled
Exclude parent mass from data dependent selection not enabled
Exclusion mass width by mass

Exclusion mass width low: 0.50000
Exclusion mass width high: 0.50000
Parent mass width by mass
Parent mass width low: 0.50000
Parent mass width high: 0.50000
Reject mass width by mass
Reject mass width low: 0.50000
Reject mass width high: 0.50000
Zoom/UltraZoom scan mass width by mass
Zoom/UltraZoom scan mass width low: 5.00
Zoom/UltraZoom scan mass width high: 5.00
FT SIM scan mass width low: 5.00
FT SIM scan mass width high: 5.00
Neutral Loss candidates processed by decreasing intensity
Neutral Loss mass width by mass
Neutral Loss mass width low: 0.50000
Neutral Loss mass width high: 0.50000
Product candidates processed by decreasing intensity
Product mass width by mass
Product mass width low: 0.50000
Product mass width high: 0.50000
MS mass range: 0.00-1000000.00
MSn mass range by mass
MSn mass range: 0.00-1000000.00
Use m/z values as masses not enabled
Analog UV data dep. not enabled
Dynamic exclusion not enabled
Isotopic data dependence not enabled
Mass Tags data dependence not enabled

Custom Data Dependent Settings:
Not enabled

---

Pass 10 (wheat-mixed-digests_MS2_TO-USE-inclusion-01_3.raw):

Creator: Orbi_30393
Last modified: 10/19/2021 by Orbi_30393
MS Run Time (min): 43.00
Sequence override of method parameters not enabled.
Divert Valve: not used during run
Contact Closure: not used during run
Syringe Pump: not used during run
MS Detector Settings:
Real-time modifications to method not enabled
Stepped collision energy not enabled
Additional Microscans:

| | | |
|---|---|---|
| MS2 | 0 | 0 |
| MS3 | 0 | 0 |
| MS4 | 0 | 0 |
| MS5 | 0 | 0 |
| MS6 | 0 | 0 |
| MS7 | 0 | 0 |

MS8 0 0
MS9 0 0
MS10 0 0
Segment 1 Information
Duration (min): 43.00
Number of Scan Events: 2
Tune Method: Orbitrap-tune-file_2020-03-13_HESI

Scan Event Details:
1: FTMS + p norm res=15000 o(300.0-2000.0)
CV = 0.0V
2: ITMS + c norm Dep MS/MS Most intense ion from (1)
Activation Type: CID
Min. Signal Required: 500.0
Isolation Width: 0.50
Normalized Coll. Energy: 35.0
Default Charge State: 2
Activation Q: 0.250
Activation Time: 10.000
CV = 0.0V

Lock Masses:
Pos List Name: N/A
Source: API Source
Mass List: (none)
Neg List Name: N/A
Source: API Source
Mass List: (none)

Data Dependent Settings:
Use separate polarity settings disabled
Parent Mass List:

| MS Mass | MS FAIMS CV | MS Normalized Collision Energy | MS2 Mass | MS2 Normalized Collision Energy | Name |
|---|---|---|---|---|---|
| 300.13000 | | 35.0 | | | Cluster_06844 |
| 300.14000 | | 35.0 | | | Cluster_18928 |
| 300.71000 | | 35.0 | | | Cluster_18955 |
| 301.50000 | | 35.0 | | | Cluster_06851 |
| 301.65000 | | 35.0 | | | Cluster_18984 |
| 302.64000 | | 35.0 | | | Cluster_19016 |
| 302.65000 | | 35.0 | | | Cluster_19017 |
| 303.99000 | | 35.0 | | | Cluster_19040 |
| 304.11000 | | 35.0 | | | Cluster_19042 |
| 304.14000 | | 35.0 | | | Cluster_19049 |
| 304.16000 | | 35.0 | | | Cluster_19050 |
| 304.20000 | | 35.0 | | | Cluster_19052 |
| 304.49000 | | 35.0 | | | Cluster_06863 |
| 304.64000 | | 35.0 | | | Cluster_19062 |
| 306.13000 | | 35.0 | | | Cluster_06873 |

| | | |
|---|---|---|
| 306.67000 | 35.0 | Cluster_19109 |
| 307.12000 | 35.0 | Cluster_19119 |
| 307.13000 | 35.0 | Cluster_06880 |
| 308.65000 | 35.0 | Cluster_19177 |
| 308.66000 | 35.0 | Cluster_19179 |
| 309.53000 | 35.0 | Cluster_06901 |
| 310.58000 | 35.0 | Cluster_19217 |
| 310.65000 | 35.0 | Cluster_19222 |
| 310.66000 | 35.0 | Cluster_19221 |
| 311.17000 | 35.0 | Cluster_19246 |
| 311.68000 | 35.0 | Cluster_19257 |
| 313.65000 | 35.0 | Cluster_19315 |
| 314.82000 | 35.0 | Cluster_06941 |
| 315.65000 | 35.0 | Cluster_19384 |
| 316.69000 | 35.0 | Cluster_19442 |
| 317.06000 | 35.0 | Cluster_19459 |
| 317.16000 | 35.0 | Cluster_19466 |
| 317.64000 | 35.0 | Cluster_19490 |
| 318.64000 | 35.0 | Cluster_19532 |
| 318.85000 | 35.0 | Cluster_06973 |
| 319.17000 | 35.0 | Cluster_19552 |
| 319.67000 | 35.0 | Cluster_19560 |
| 319.94000 | 35.0 | Cluster_00850 |
| 320.49000 | 35.0 | Cluster_06993 |
| 320.80000 | 35.0 | Cluster_06994 |
| 321.15000 | 35.0 | Cluster_06995 |
| 321.17000 | 35.0 | Cluster_00869 |
| 321.52000 | 35.0 | Cluster_19594 |
| 322.15000 | 35.0 | Cluster_19619 |
| 322.17000 | 35.0 | Cluster_19617 |
| 322.50000 | 35.0 | Cluster_06998 |
| 323.65000 | 35.0 | Cluster_19672 |
| 323.69000 | 35.0 | Cluster_19677 |
| 323.83000 | 35.0 | Cluster_19683 |
| 324.42000 | 35.0 | Cluster_00878 |
| 324.60000 | 35.0 | Cluster_19711 |
| 324.65000 | 35.0 | Cluster_19713 |
| 324.83000 | 35.0 | Cluster_07017 |
| 325.14000 | 35.0 | Cluster_19730 |
| 325.82000 | 35.0 | Cluster_07023 |
| 325.85000 | 35.0 | Cluster_07028 |
| 326.17000 | 35.0 | Cluster_19753 |
| 326.40000 | 35.0 | Cluster_00887 |
| 326.90000 | 35.0 | Cluster_00888 |
| 327.21000 | 35.0 | Cluster_19791 |
| 327.96000 | 35.0 | Cluster_07043 |
| 328.48000 | 35.0 | Cluster_07051 |
| 329.18000 | 35.0 | Cluster_19855 |
| 329.82000 | 35.0 | Cluster_07063 |
| 329.93000 | 35.0 | Cluster_19873 |
| 330.18000 | 35.0 | Cluster_19886 |

| | | |
|---|---|---|
| 330.90000 | 35.0 | Cluster_19916 |
| 331.17000 | 35.0 | Cluster_19926 |
| 331.20000 | 35.0 | Cluster_19928 |
| 331.21000 | 35.0 | Cluster_19930 |
| 331.22000 | 35.0 | Cluster_19925 |
| 331.66000 | 35.0 | Cluster_19936 |
| 332.14000 | 35.0 | Cluster_19948 |
| 333.47000 | 35.0 | Cluster_07081 |
| 333.82000 | 35.0 | Cluster_07087 |
| 333.87000 | 35.0 | Cluster_07090 |
| 335.12000 | 35.0 | Cluster_20034 |
| 335.83000 | 35.0 | Cluster_07104 |
| 336.19000 | 35.0 | Cluster_07109 |
| 336.23000 | 35.0 | Cluster_20093 |
| 336.91000 | 35.0 | Cluster_00920 |
| 337.19000 | 35.0 | Cluster_20132 |
| 337.21000 | 35.0 | Cluster_20126 |
| 337.83000 | 35.0 | Cluster_07126 |
| 338.11000 | 35.0 | Cluster_20150 |
| 338.67000 | 35.0 | Cluster_20172 |
| 339.16000 | 35.0 | Cluster_20192 |
| 339.47000 | 35.0 | Cluster_07139 |
| 339.81000 | 35.0 | Cluster_07150 |
| 339.94000 | 35.0 | Cluster_00929 |
| 340.18000 | 35.0 | Cluster_20223 |
| 340.26000 | 35.0 | Cluster_20229 |
| 341.15000 | 35.0 | Cluster_20257 |
| 341.16000 | 35.0 | Cluster_20261 |
| 342.16000 | 35.0 | Cluster_07167 |
| 343.11000 | 35.0 | Cluster_20316 |
| 343.18000 | 35.0 | Cluster_20318 |
| 343.66000 | 35.0 | Cluster_20344 |
| 345.19000 | 35.0 | Cluster_20390 |
| 345.67000 | 35.0 | Cluster_20402 |
| 346.15000 | 35.0 | Cluster_07199 |
| 346.67000 | 35.0 | Cluster_20444 |
| 347.20000 | 35.0 | Cluster_20457 |
| 347.71000 | 35.0 | Cluster_20471 |
| 348.44000 | 35.0 | Cluster_20495 |
| 348.87000 | 35.0 | Cluster_07234 |
| 348.92000 | 35.0 | Cluster_00975 |
| 348.93000 | 35.0 | Cluster_00976 |
| 349.20000 | 35.0 | Cluster_20517 |
| 349.21000 | 35.0 | Cluster_20520 |
| 349.64000 | 35.0 | Cluster_20543 |
| 349.73000 | 35.0 | Cluster_20549 |
| 350.42000 | 35.0 | Cluster_00982 |
| 351.18000 | 35.0 | Cluster_20611 |
| 351.19000 | 35.0 | Cluster_20622 |
| 352.66000 | 35.0 | Cluster_20669 |
| 353.17000 | 35.0 | Cluster_01002 |

| | | |
|---|---|---|
| 353.21000 | 35.0 | Cluster_20705 |
| 353.66000 | 35.0 | Cluster_01003 |
| 353.80000 | 35.0 | Cluster_07278 |
| 353.87000 | 35.0 | Cluster_20720 |
| 353.91000 | 35.0 | Cluster_01004 |
| 354.16000 | 35.0 | Cluster_20732 |
| 354.20000 | 35.0 | Cluster_20739 |
| 354.40000 | 35.0 | Cluster_01009 |
| 354.42000 | 35.0 | Cluster_01011 |
| 355.14000 | 35.0 | Cluster_20781 |
| 355.16000 | 35.0 | Cluster_20782 |
| 355.22000 | 35.0 | Cluster_20787 |
| 355.36000 | 35.0 | Cluster_20789 |
| 355.70000 | 35.0 | Cluster_20796 |
| 356.24000 | 35.0 | Cluster_20824 |
| 356.55000 | 35.0 | Cluster_07307 |
| 358.42000 | 35.0 | Cluster_20933 |
| 359.13000 | 35.0 | Cluster_07326 |
| 359.17000 | 35.0 | Cluster_00118 |
| 359.37000 | 35.0 | Cluster_20960 |
| 360.16000 | 35.0 | Cluster_20986 |
| 360.42000 | 35.0 | Cluster_01047 |
| 360.83000 | 35.0 | Cluster_21008 |
| 361.18000 | 35.0 | Cluster_21024 |
| 361.42000 | 35.0 | Cluster_01055 |
| 361.66000 | 35.0 | Cluster_21038 |
| 361.67000 | 35.0 | Cluster_21036 |
| 361.69000 | 35.0 | Cluster_21039 |
| 361.84000 | 35.0 | Cluster_07347 |
| 362.10000 | 35.0 | Cluster_21048 |
| 362.42000 | 35.0 | Cluster_01059 |
| 362.43000 | 35.0 | Cluster_21075 |
| 362.60000 | 35.0 | Cluster_21077 |
| 362.65000 | 35.0 | Cluster_21076 |
| 363.11000 | 35.0 | Cluster_21089 |
| 363.19000 | 35.0 | Cluster_21093 |
| 363.20000 | 35.0 | Cluster_21094 |
| 363.77000 | 35.0 | Cluster_21121 |
| 364.16000 | 35.0 | Cluster_21135 |
| 364.50000 | 35.0 | Cluster_21161 |
| 364.65000 | 35.0 | Cluster_21162 |
| 365.24000 | 35.0 | Cluster_21221 |
| 365.79000 | 35.0 | Cluster_07396 |
| 365.90000 | 35.0 | Cluster_21241 |
| 366.27000 | 35.0 | Cluster_21267 |
| 366.74000 | 35.0 | Cluster_21285 |
| 367.22000 | 35.0 | Cluster_21306 |
| 367.50000 | 35.0 | Cluster_07416 |
| 367.53000 | 35.0 | Cluster_21324 |
| 368.16000 | 35.0 | Cluster_21368 |
| 368.20000 | 35.0 | Cluster_21366 |

| | | |
|---|---|---|
| 368.47000 | 35.0 | Cluster_07428 |
| 369.67000 | 35.0 | Cluster_21437 |
| 370.08000 | 35.0 | Cluster_21453 |
| 370.50000 | 35.0 | Cluster_07445 |
| 370.74000 | 35.0 | Cluster_21476 |
| 370.93000 | 35.0 | Cluster_21486 |
| 371.18000 | 35.0 | Cluster_07449 |
| 371.24000 | 35.0 | Cluster_21501 |
| 371.71000 | 35.0 | Cluster_21517 |
| 371.89000 | 35.0 | Cluster_21531 |
| 372.17000 | 35.0 | Cluster_21537 |
| 372.23000 | 35.0 | Cluster_07464 |
| 372.67000 | 35.0 | Cluster_21575 |
| 372.69000 | 35.0 | Cluster_21572 |
| 373.12000 | 35.0 | Cluster_21589 |
| 373.14000 | 35.0 | Cluster_21585 |
| 373.16000 | 35.0 | Cluster_07470 |
| 373.19000 | 35.0 | Cluster_07471 |
| 373.22000 | 35.0 | Cluster_21597 |
| 373.90000 | 35.0 | Cluster_21628 |
| 374.19000 | 35.0 | Cluster_07480 |
| 374.40000 | 35.0 | Cluster_21650 |
| 374.68000 | 35.0 | Cluster_21656 |
| 375.15000 | 35.0 | Cluster_21668 |
| 375.89000 | 35.0 | Cluster_07496 |
| 377.48000 | 35.0 | Cluster_07512 |
| 377.77000 | 35.0 | Cluster_21780 |
| 378.17000 | 35.0 | Cluster_01122 |
| 378.18000 | 35.0 | Cluster_21788 |
| 378.20000 | 35.0 | Cluster_07523 |
| 378.67000 | 35.0 | Cluster_01125 |
| 379.26000 | 35.0 | Cluster_21841 |
| 380.45000 | 35.0 | Cluster_21886 |
| 380.68000 | 35.0 | Cluster_21891 |
| 381.22000 | 35.0 | Cluster_07550 |
| 381.70000 | 35.0 | Cluster_01141 |
| 382.18000 | 35.0 | Cluster_21953 |
| 382.66000 | 35.0 | Cluster_01144 |
| 383.63000 | 35.0 | Cluster_22024 |
| 383.68000 | 35.0 | Cluster_22025 |
| 384.16000 | 35.0 | Cluster_01146 |
| 384.21000 | 35.0 | Cluster_22055 |
| 384.22000 | 35.0 | Cluster_22065 |
| 385.52000 | 35.0 | Cluster_07597 |
| 385.54000 | 35.0 | Cluster_07596 |
| 385.86000 | 35.0 | Cluster_22123 |
| 386.66000 | 35.0 | Cluster_22155 |
| 386.68000 | 35.0 | Cluster_22158 |
| 386.90000 | 35.0 | Cluster_22172 |
| 387.11000 | 35.0 | Cluster_22177 |
| 387.63000 | 35.0 | Cluster_22217 |

| | | |
|---|---|---|
| 387.93000 | 35.0 | Cluster_22226 |
| 388.23000 | 35.0 | Cluster_22233 |
| 388.50000 | 35.0 | Cluster_07621 |
| 388.64000 | 35.0 | Cluster_22244 |
| 388.67000 | 35.0 | Cluster_22245 |
| 388.83000 | 35.0 | Cluster_07623 |
| 389.25000 | 35.0 | Cluster_22269 |
| 389.51000 | 35.0 | Cluster_07627 |
| 389.91000 | 35.0 | Cluster_22284 |
| 390.71000 | 35.0 | Cluster_22316 |
| 390.87000 | 35.0 | Cluster_07651 |
| 390.88000 | 35.0 | Cluster_07646 |
| 391.50000 | 35.0 | Cluster_07654 |
| 391.65000 | 35.0 | Cluster_22334 |
| 391.75000 | 35.0 | Cluster_22344 |
| 391.82000 | 35.0 | Cluster_07658 |
| 392.22000 | 35.0 | Cluster_07665 |
| 392.25000 | 35.0 | Cluster_22358 |
| 392.73000 | 35.0 | Cluster_22381 |
| 392.77000 | 35.0 | Cluster_22377 |
| 393.19000 | 35.0 | Cluster_01195 |
| 393.53000 | 35.0 | Cluster_07674 |
| 393.69000 | 35.0 | Cluster_01197 |
| 393.71000 | 35.0 | Cluster_22428 |
| 393.98000 | 35.0 | Cluster_22479 |
| 394.21000 | 35.0 | Cluster_22490 |
| 394.71000 | 35.0 | Cluster_22522 |
| 394.72000 | 35.0 | Cluster_22521 |
| 395.89000 | 35.0 | Cluster_07703 |
| 396.23000 | 35.0 | Cluster_22589 |
| 396.30000 | 35.0 | Cluster_22595 |
| 397.39000 | 35.0 | Cluster_22628 |
| 397.67000 | 35.0 | Cluster_22634 |
| 397.72000 | 35.0 | Cluster_22641 |
| 397.90000 | 35.0 | Cluster_07736 |
| 398.18000 | 35.0 | Cluster_01226 |
| 398.44000 | 35.0 | Cluster_01230 |
| 400.24000 | 35.0 | Cluster_22747 |
| 400.68000 | 35.0 | Cluster_22780 |
| 400.90000 | 35.0 | Cluster_07771 |
| 401.70000 | 35.0 | Cluster_22833 |
| 402.32000 | 35.0 | Cluster_22866 |
| 402.48000 | 35.0 | Cluster_22868 |
| 402.72000 | 35.0 | Cluster_22878 |
| 403.44000 | 35.0 | Cluster_01260 |
| 403.54000 | 35.0 | Cluster_07797 |
| 404.47000 | 35.0 | Cluster_01272 |
| 405.88000 | 35.0 | Cluster_07831 |
| 406.22000 | 35.0 | Cluster_23023 |
| 406.68000 | 35.0 | Cluster_01285 |
| 406.73000 | 35.0 | Cluster_23046 |

| | | |
|---|---|---|
| 406.95000 | 35.0 | Cluster_01293 |
| 407.17000 | 35.0 | Cluster_23059 |
| 407.25000 | 35.0 | Cluster_23066 |
| 407.67000 | 35.0 | Cluster_01302 |
| 407.75000 | 35.0 | Cluster_23083 |
| 407.93000 | 35.0 | Cluster_01303 |
| 408.22000 | 35.0 | Cluster_23104 |
| 408.54000 | 35.0 | Cluster_07863 |
| 408.66000 | 35.0 | Cluster_01305 |
| 408.91000 | 35.0 | Cluster_07864 |
| 408.92000 | 35.0 | Cluster_01307 |
| 409.19000 | 35.0 | Cluster_07867 |
| 409.25000 | 35.0 | Cluster_23160 |
| 409.69000 | 35.0 | Cluster_23170 |
| 410.53000 | 35.0 | Cluster_07882 |
| 411.72000 | 35.0 | Cluster_23258 |
| 412.67000 | 35.0 | Cluster_23299 |
| 412.68000 | 35.0 | Cluster_01326 |
| 413.23000 | 35.0 | Cluster_23320 |
| 413.87000 | 35.0 | Cluster_07913 |
| 413.92000 | 35.0 | Cluster_07916 |
| 413.94000 | 35.0 | Cluster_23350 |
| 413.98000 | 35.0 | Cluster_23354 |
| 414.71000 | 35.0 | Cluster_23379 |
| 414.85000 | 35.0 | Cluster_23397 |
| 414.87000 | 35.0 | Cluster_07924 |
| 415.22000 | 35.0 | Cluster_23422 |
| 415.73000 | 35.0 | Cluster_23434 |
| 416.00000 | 35.0 | Cluster_23448 |
| 416.22000 | 35.0 | Cluster_23463 |
| 416.76000 | 35.0 | Cluster_23476 |
| 416.88000 | 35.0 | Cluster_23482 |
| 416.94000 | 35.0 | Cluster_01353 |
| 418.25000 | 35.0 | Cluster_23534 |
| 418.47000 | 35.0 | Cluster_23549 |
| 419.18000 | 35.0 | Cluster_01360 |
| 419.21000 | 35.0 | Cluster_07978 |
| 419.88000 | 35.0 | Cluster_23600 |
| 420.19000 | 35.0 | Cluster_23617 |
| 420.23000 | 35.0 | Cluster_23612 |
| 420.71000 | 35.0 | Cluster_23632 |
| 421.56000 | 35.0 | Cluster_08005 |
| 422.12000 | 35.0 | Cluster_08008 |
| 422.55000 | 35.0 | Cluster_08014 |
| 422.58000 | 35.0 | Cluster_08015 |
| 423.14000 | 35.0 | Cluster_23727 |
| 423.21000 | 35.0 | Cluster_08022 |
| 423.24000 | 35.0 | Cluster_23740 |
| 423.44000 | 35.0 | Cluster_01387 |
| 423.71000 | 35.0 | Cluster_23753 |
| 423.74000 | 35.0 | Cluster_23766 |

| | | |
|---|---|---|
| 424.18000 | 35.0 | Cluster_23787 |
| 424.51000 | 35.0 | Cluster_23811 |
| 424.68000 | 35.0 | Cluster_23813 |
| 425.18000 | 35.0 | Cluster_23850 |
| 425.46000 | 35.0 | Cluster_23865 |
| 425.87000 | 35.0 | Cluster_08056 |
| 425.88000 | 35.0 | Cluster_23895 |
| 426.14000 | 35.0 | Cluster_23901 |
| 426.24000 | 35.0 | Cluster_23924 |
| 426.44000 | 35.0 | Cluster_23942 |
| 426.88000 | 35.0 | Cluster_08077 |
| 427.51000 | 35.0 | Cluster_08094 |
| 428.56000 | 35.0 | Cluster_08107 |
| 428.72000 | 35.0 | Cluster_24045 |
| 428.79000 | 35.0 | Cluster_24059 |
| 429.24000 | 35.0 | Cluster_24074 |
| 429.57000 | 35.0 | Cluster_08120 |
| 429.72000 | 35.0 | Cluster_24100 |
| 429.76000 | 35.0 | Cluster_24093 |
| 429.85000 | 35.0 | Cluster_08124 |
| 429.95000 | 35.0 | Cluster_01432 |
| 430.73000 | 35.0 | Cluster_24136 |
| 431.21000 | 35.0 | Cluster_24172 |
| 431.41000 | 35.0 | Cluster_24179 |
| 431.74000 | 35.0 | Cluster_24194 |
| 432.16000 | 35.0 | Cluster_24207 |
| 432.21000 | 35.0 | Cluster_24215 |
| 432.87000 | 35.0 | Cluster_08164 |
| 433.96000 | 35.0 | Cluster_24289 |
| 434.79000 | 35.0 | Cluster_24347 |
| 434.97000 | 35.0 | Cluster_24350 |
| 435.17000 | 35.0 | Cluster_24357 |
| 435.24000 | 35.0 | Cluster_08201 |
| 435.37000 | 35.0 | Cluster_24383 |
| 435.70000 | 35.0 | Cluster_01481 |
| 435.97000 | 35.0 | Cluster_01487 |
| 436.48000 | 35.0 | Cluster_01491 |
| 437.37000 | 35.0 | Cluster_24491 |
| 437.46000 | 35.0 | Cluster_01497 |
| 438.22000 | 35.0 | Cluster_24532 |
| 438.30000 | 35.0 | Cluster_24543 |
| 438.96000 | 35.0 | Cluster_01503 |
| 438.97000 | 35.0 | Cluster_01504 |
| 439.82000 | 35.0 | Cluster_24632 |
| 440.21000 | 35.0 | Cluster_01512 |
| 440.56000 | 35.0 | Cluster_24655 |
| 440.74000 | 35.0 | Cluster_24659 |
| 440.86000 | 35.0 | Cluster_08277 |
| 441.21000 | 35.0 | Cluster_24682 |
| 441.24000 | 35.0 | Cluster_01516 |
| 441.73000 | 35.0 | Cluster_24703 |

| | | |
|---|---|---|
| 441.87000 | 35.0 | Cluster_08293 |
| 442.00000 | 35.0 | Cluster_24716 |
| 442.28000 | 35.0 | Cluster_24730 |
| 442.57000 | 35.0 | Cluster_08303 |
| 442.77000 | 35.0 | Cluster_24758 |
| 442.89000 | 35.0 | Cluster_24763 |
| 442.98000 | 35.0 | Cluster_24761 |
| 443.12000 | 35.0 | Cluster_24771 |
| 443.22000 | 35.0 | Cluster_24782 |
| 443.28000 | 35.0 | Cluster_24785 |
| 443.29000 | 35.0 | Cluster_24795 |
| 443.89000 | 35.0 | Cluster_08322 |
| 443.92000 | 35.0 | Cluster_08324 |
| 444.72000 | 35.0 | Cluster_24866 |
| 445.18000 | 35.0 | Cluster_08343 |
| 445.55000 | 35.0 | Cluster_24900 |
| 446.27000 | 35.0 | Cluster_24941 |
| 446.73000 | 35.0 | Cluster_24959 |
| 446.91000 | 35.0 | Cluster_08369 |
| 447.17000 | 35.0 | Cluster_24974 |
| 447.25000 | 35.0 | Cluster_08376 |
| 447.28000 | 35.0 | Cluster_24982 |
| 447.29000 | 35.0 | Cluster_24988 |
| 447.60000 | 35.0 | Cluster_08380 |
| 448.30000 | 35.0 | Cluster_25022 |
| 448.79000 | 35.0 | Cluster_25051 |
| 448.97000 | 35.0 | Cluster_01573 |
| 449.25000 | 35.0 | Cluster_08416 |
| 449.74000 | 35.0 | Cluster_25092 |
| 449.84000 | 35.0 | Cluster_25102 |
| 450.23000 | 35.0 | Cluster_25125 |
| 450.93000 | 35.0 | Cluster_08435 |
| 451.14000 | 35.0 | Cluster_25153 |
| 451.17000 | 35.0 | Cluster_25168 |
| 451.57000 | 35.0 | Cluster_08446 |
| 452.13000 | 35.0 | Cluster_25195 |
| 452.26000 | 35.0 | Cluster_08462 |
| 452.75000 | 35.0 | Cluster_25235 |
| 453.14000 | 35.0 | Cluster_25244 |
| 453.24000 | 35.0 | Cluster_25248 |
| 454.68000 | 35.0 | Cluster_25320 |
| 454.70000 | 35.0 | Cluster_25319 |
| 454.74000 | 35.0 | Cluster_25325 |
| 455.15000 | 35.0 | Cluster_25343 |
| 455.18000 | 35.0 | Cluster_01617 |
| 455.20000 | 35.0 | Cluster_08518 |
| 456.24000 | 35.0 | Cluster_25393 |
| 456.26000 | 35.0 | Cluster_08536 |
| 456.57000 | 35.0 | Cluster_08539 |
| 456.75000 | 35.0 | Cluster_25404 |
| 456.78000 | 35.0 | Cluster_25413 |

| | | |
|---|---|---|
| 457.20000 | 35.0 | Cluster_25422 |
| 457.75000 | 35.0 | Cluster_25448 |
| 457.76000 | 35.0 | Cluster_25454 |
| 458.69000 | 35.0 | Cluster_25493 |
| 458.76000 | 35.0 | Cluster_25500 |
| 459.22000 | 35.0 | Cluster_08566 |
| 459.25000 | 35.0 | Cluster_25514 |
| 459.78000 | 35.0 | Cluster_25540 |
| 459.90000 | 35.0 | Cluster_08580 |
| 460.27000 | 35.0 | Cluster_25572 |
| 460.77000 | 35.0 | Cluster_25581 |
| 460.97000 | 35.0 | Cluster_01663 |
| 461.60000 | 35.0 | Cluster_08592 |
| 461.70000 | 35.0 | Cluster_25618 |
| 461.73000 | 35.0 | Cluster_25620 |
| 462.22000 | 35.0 | Cluster_08608 |
| 462.50000 | 35.0 | Cluster_25643 |
| 463.61000 | 35.0 | Cluster_25677 |
| 463.73000 | 35.0 | Cluster_25692 |
| 464.22000 | 35.0 | Cluster_08632 |
| 464.30000 | 35.0 | Cluster_25714 |
| 464.55000 | 35.0 | Cluster_08640 |
| 465.77000 | 35.0 | Cluster_25807 |
| 465.95000 | 35.0 | Cluster_08659 |
| 465.99000 | 35.0 | Cluster_01692 |
| 466.27000 | 35.0 | Cluster_25822 |
| 466.90000 | 35.0 | Cluster_08673 |
| 467.18000 | 35.0 | Cluster_08677 |
| 467.25000 | 35.0 | Cluster_08678 |
| 467.30000 | 35.0 | Cluster_25858 |
| 467.67000 | 35.0 | Cluster_00144 |
| 467.72000 | 35.0 | Cluster_25877 |
| 468.89000 | 35.0 | Cluster_08711 |
| 469.26000 | 35.0 | Cluster_08720 |
| 469.74000 | 35.0 | Cluster_25967 |
| 469.88000 | 35.0 | Cluster_08724 |
| 470.43000 | 35.0 | Cluster_26002 |
| 470.56000 | 35.0 | Cluster_00010 |
| 470.92000 | 35.0 | Cluster_08745 |
| 471.00000 | 35.0 | Cluster_01731 |
| 471.22000 | 35.0 | Cluster_08748 |
| 471.25000 | 35.0 | Cluster_01737 |
| 471.26000 | 35.0 | Cluster_01738 |
| 471.48000 | 35.0 | Cluster_26044 |
| 471.74000 | 35.0 | Cluster_26063 |
| 471.84000 | 35.0 | Cluster_26052 |
| 472.23000 | 35.0 | Cluster_26081 |
| 472.24000 | 35.0 | Cluster_08764 |
| 472.72000 | 35.0 | Cluster_26107 |
| 472.76000 | 35.0 | Cluster_26106 |
| 472.77000 | 35.0 | Cluster_26113 |

| | | |
|---|---|---|
| 473.27000 | 35.0 | Cluster_26146 |
| 473.77000 | 35.0 | Cluster_26157 |
| 474.54000 | 35.0 | Cluster_08802 |
| 474.58000 | 35.0 | Cluster_08800 |
| 474.85000 | 35.0 | Cluster_08806 |
| 475.58000 | 35.0 | Cluster_08820 |
| 475.78000 | 35.0 | Cluster_26234 |
| 476.21000 | 35.0 | Cluster_26244 |
| 476.23000 | 35.0 | Cluster_08830 |
| 476.28000 | 35.0 | Cluster_08832 |
| 477.07000 | 35.0 | Cluster_26275 |
| 477.20000 | 35.0 | Cluster_08843 |
| 477.27000 | 35.0 | Cluster_08840 |
| 477.55000 | 35.0 | Cluster_08851 |
| 477.61000 | 35.0 | Cluster_08856 |
| 477.93000 | 35.0 | Cluster_08861 |
| 478.22000 | 35.0 | Cluster_26309 |
| 478.25000 | 35.0 | Cluster_26313 |
| 478.76000 | 35.0 | Cluster_26329 |
| 478.77000 | 35.0 | Cluster_26332 |
| 478.93000 | 35.0 | Cluster_08880 |
| 478.99000 | 35.0 | Cluster_01780 |
| 479.57000 | 35.0 | Cluster_08888 |
| 479.71000 | 35.0 | Cluster_26429 |
| 479.76000 | 35.0 | Cluster_26423 |
| 480.22000 | 35.0 | Cluster_08900 |
| 480.70000 | 35.0 | Cluster_26476 |
| 481.22000 | 35.0 | Cluster_26493 |
| 481.27000 | 35.0 | Cluster_26504 |
| 481.28000 | 35.0 | Cluster_08920 |
| 481.56000 | 35.0 | Cluster_08922 |
| 481.58000 | 35.0 | Cluster_08924 |
| 481.72000 | 35.0 | Cluster_26523 |
| 481.73000 | 35.0 | Cluster_01804 |
| 481.79000 | 35.0 | Cluster_26518 |
| 482.20000 | 35.0 | Cluster_01810 |
| 482.28000 | 35.0 | Cluster_26547 |
| 482.49000 | 35.0 | Cluster_26549 |
| 483.24000 | 35.0 | Cluster_26594 |
| 483.30000 | 35.0 | Cluster_26603 |
| 483.76000 | 35.0 | Cluster_26625 |
| 483.84000 | 35.0 | Cluster_26627 |
| 484.09000 | 35.0 | Cluster_26635 |
| 484.73000 | 35.0 | Cluster_26685 |
| 484.76000 | 35.0 | Cluster_26681 |
| 485.06000 | 35.0 | Cluster_00166 |
| 485.54000 | 35.0 | Cluster_08981 |
| 488.29000 | 35.0 | Cluster_26837 |
| 488.61000 | 35.0 | Cluster_09032 |
| 488.79000 | 35.0 | Cluster_26859 |
| 488.95000 | 35.0 | Cluster_09038 |

| | | |
|---|---|---|
| 489.74000 | 35.0 | Cluster_26904 |
| 489.93000 | 35.0 | Cluster_09048 |
| 490.50000 | 35.0 | Cluster_26931 |
| 490.70000 | 35.0 | Cluster_26935 |
| 490.77000 | 35.0 | Cluster_26939 |
| 492.55000 | 35.0 | Cluster_09097 |
| 492.60000 | 35.0 | Cluster_09099 |
| 493.27000 | 35.0 | Cluster_27041 |
| 494.26000 | 35.0 | Cluster_27100 |
| 495.19000 | 35.0 | Cluster_27149 |
| 495.71000 | 35.0 | Cluster_27170 |
| 495.95000 | 35.0 | Cluster_09160 |
| 496.24000 | 35.0 | Cluster_01895 |
| 496.95000 | 35.0 | Cluster_09176 |
| 497.35000 | 35.0 | Cluster_27258 |
| 497.74000 | 35.0 | Cluster_27265 |
| 497.80000 | 35.0 | Cluster_27275 |
| 497.94000 | 35.0 | Cluster_09191 |
| 498.01000 | 35.0 | Cluster_01929 |
| 498.50000 | 35.0 | Cluster_27299 |
| 498.73000 | 35.0 | Cluster_27302 |
| 499.23000 | 35.0 | Cluster_09211 |
| 499.33000 | 35.0 | Cluster_27324 |
| 499.76000 | 35.0 | Cluster_01941 |
| 499.92000 | 35.0 | Cluster_09218 |
| 499.96000 | 35.0 | Cluster_09219 |
| 500.02000 | 35.0 | Cluster_27355 |
| 500.63000 | 35.0 | Cluster_09231 |
| 500.70000 | 35.0 | Cluster_27401 |
| 500.96000 | 35.0 | Cluster_01956 |
| 501.22000 | 35.0 | Cluster_27436 |
| 501.59000 | 35.0 | Cluster_09244 |
| 501.76000 | 35.0 | Cluster_27459 |
| 502.21000 | 35.0 | Cluster_01970 |
| 502.22000 | 35.0 | Cluster_27473 |
| 502.59000 | 35.0 | Cluster_09268 |
| 502.76000 | 35.0 | Cluster_27485 |
| 503.25000 | 35.0 | Cluster_27512 |
| 503.28000 | 35.0 | Cluster_27518 |
| 504.21000 | 35.0 | Cluster_27547 |
| 504.56000 | 35.0 | Cluster_09288 |
| 504.98000 | 35.0 | Cluster_01984 |
| 505.01000 | 35.0 | Cluster_27567 |
| 505.24000 | 35.0 | Cluster_09298 |
| 505.52000 | 35.0 | Cluster_01990 |
| 505.60000 | 35.0 | Cluster_09305 |
| 505.79000 | 35.0 | Cluster_27593 |
| 506.73000 | 35.0 | Cluster_27611 |
| 506.74000 | 35.0 | Cluster_27621 |
| 506.76000 | 35.0 | Cluster_27614 |
| 507.25000 | 35.0 | Cluster_27638 |

| | | |
|---|---|---|
| 507.97000 | 35.0 | Cluster_09350 |
| 508.62000 | 35.0 | Cluster_09366 |
| 508.70000 | 35.0 | Cluster_27686 |
| 508.71000 | 35.0 | Cluster_27692 |
| 509.20000 | 35.0 | Cluster_27715 |
| 509.26000 | 35.0 | Cluster_27718 |
| 509.87000 | 35.0 | Cluster_09377 |
| 509.91000 | 35.0 | Cluster_09383 |
| 511.23000 | 35.0 | Cluster_27820 |
| 511.79000 | 35.0 | Cluster_27852 |
| 511.91000 | 35.0 | Cluster_09420 |
| 512.25000 | 35.0 | Cluster_27867 |
| 512.27000 | 35.0 | Cluster_27865 |
| 513.09000 | 35.0 | Cluster_27898 |
| 513.77000 | 35.0 | Cluster_27929 |
| 514.72000 | 35.0 | Cluster_27964 |
| 514.74000 | 35.0 | Cluster_27971 |
| 515.25000 | 35.0 | Cluster_02053 |
| 515.48000 | 35.0 | Cluster_02061 |
| 515.90000 | 35.0 | Cluster_09478 |
| 516.74000 | 35.0 | Cluster_28062 |
| 517.92000 | 35.0 | Cluster_09503 |
| 518.00000 | 35.0 | Cluster_28115 |
| 518.26000 | 35.0 | Cluster_09508 |
| 518.31000 | 35.0 | Cluster_28130 |
| 518.51000 | 35.0 | Cluster_28138 |
| 518.59000 | 35.0 | Cluster_09521 |
| 518.78000 | 35.0 | Cluster_28152 |
| 519.59000 | 35.0 | Cluster_09531 |
| 519.81000 | 35.0 | Cluster_28200 |
| 520.04000 | 35.0 | Cluster_28202 |
| 520.51000 | 35.0 | Cluster_28225 |
| 520.80000 | 35.0 | Cluster_28240 |
| 520.81000 | 35.0 | Cluster_28232 |
| 521.02000 | 35.0 | Cluster_02095 |
| 521.26000 | 35.0 | Cluster_09553 |
| 521.66000 | 35.0 | Cluster_28271 |
| 522.22000 | 35.0 | Cluster_28286 |
| 522.45000 | 35.0 | Cluster_02102 |
| 522.50000 | 35.0 | Cluster_28303 |
| 522.62000 | 35.0 | Cluster_09582 |
| 522.75000 | 35.0 | Cluster_28315 |
| 523.22000 | 35.0 | Cluster_28350 |
| 523.27000 | 35.0 | Cluster_28334 |
| 523.35000 | 35.0 | Cluster_28353 |
| 523.92000 | 35.0 | Cluster_09603 |
| 523.97000 | 35.0 | Cluster_28374 |
| 524.28000 | 35.0 | Cluster_28386 |
| 526.18000 | 35.0 | Cluster_28467 |
| 526.20000 | 35.0 | Cluster_09656 |
| 526.93000 | 35.0 | Cluster_09676 |

| | | |
|---|---|---|
| 526.95000 | 35.0 | Cluster_09677 |
| 527.76000 | 35.0 | Cluster_28507 |
| 528.23000 | 35.0 | Cluster_28536 |
| 528.27000 | 35.0 | Cluster_00184 |
| 528.32000 | 35.0 | Cluster_28539 |
| 528.36000 | 35.0 | Cluster_28552 |
| 528.97000 | 35.0 | Cluster_09710 |
| 529.25000 | 35.0 | Cluster_09712 |
| 529.79000 | 35.0 | Cluster_28615 |
| 530.20000 | 35.0 | Cluster_28629 |
| 530.26000 | 35.0 | Cluster_02153 |
| 530.51000 | 35.0 | Cluster_28657 |
| 530.81000 | 35.0 | Cluster_28677 |
| 530.94000 | 35.0 | Cluster_09741 |
| 531.75000 | 35.0 | Cluster_28707 |
| 531.77000 | 35.0 | Cluster_28708 |
| 532.24000 | 35.0 | Cluster_09763 |
| 532.27000 | 35.0 | Cluster_09766 |
| 532.62000 | 35.0 | Cluster_09768 |
| 532.97000 | 35.0 | Cluster_09773 |
| 533.15000 | 35.0 | Cluster_28752 |
| 533.16000 | 35.0 | Cluster_28753 |
| 533.19000 | 35.0 | Cluster_09774 |
| 533.32000 | 35.0 | Cluster_09778 |
| 533.70000 | 35.0 | Cluster_28783 |
| 533.82000 | 35.0 | Cluster_28799 |
| 533.91000 | 35.0 | Cluster_09792 |
| 534.28000 | 35.0 | Cluster_28807 |
| 534.59000 | 35.0 | Cluster_09803 |
| 535.32000 | 35.0 | Cluster_09812 |
| 535.80000 | 35.0 | Cluster_28867 |
| 535.94000 | 35.0 | Cluster_09822 |
| 536.06000 | 35.0 | Cluster_28878 |
| 536.26000 | 35.0 | Cluster_09823 |
| 536.27000 | 35.0 | Cluster_28888 |
| 536.48000 | 35.0 | Cluster_28899 |
| 536.60000 | 35.0 | Cluster_09834 |
| 537.23000 | 35.0 | Cluster_09849 |
| 537.56000 | 35.0 | Cluster_09853 |
| 537.69000 | 35.0 | Cluster_28943 |
| 537.91000 | 35.0 | Cluster_09869 |
| 538.84000 | 35.0 | Cluster_28980 |
| 538.97000 | 35.0 | Cluster_09886 |
| 538.98000 | 35.0 | Cluster_09885 |
| 539.01000 | 35.0 | Cluster_28985 |
| 539.28000 | 35.0 | Cluster_28993 |
| 539.52000 | 35.0 | Cluster_29001 |
| 540.02000 | 35.0 | Cluster_09899 |
| 540.27000 | 35.0 | Cluster_29031 |
| 540.79000 | 35.0 | Cluster_29052 |
| 541.34000 | 35.0 | Cluster_29085 |

| | | |
|---|---|---|
| 541.44000 | 35.0 | Cluster_29098 |
| 541.82000 | 35.0 | Cluster_29115 |
| 542.26000 | 35.0 | Cluster_02236 |
| 542.28000 | 35.0 | Cluster_02229 |
| 542.52000 | 35.0 | Cluster_02233 |
| 542.77000 | 35.0 | Cluster_29162 |
| 543.27000 | 35.0 | Cluster_29193 |
| 544.95000 | 35.0 | Cluster_09982 |
| 545.77000 | 35.0 | Cluster_29280 |
| 545.92000 | 35.0 | Cluster_09992 |
| 546.31000 | 35.0 | Cluster_29312 |
| 547.32000 | 35.0 | Cluster_29345 |
| 548.27000 | 35.0 | Cluster_29389 |
| 548.46000 | 35.0 | Cluster_00194 |
| 548.61000 | 35.0 | Cluster_10035 |
| 548.64000 | 35.0 | Cluster_10037 |
| 549.25000 | 35.0 | Cluster_10044 |
| 549.28000 | 35.0 | Cluster_29418 |
| 549.90000 | 35.0 | Cluster_10052 |
| 549.99000 | 35.0 | Cluster_10058 |
| 550.22000 | 35.0 | Cluster_29451 |
| 550.31000 | 35.0 | Cluster_29461 |
| 550.55000 | 35.0 | Cluster_10069 |
| 550.61000 | 35.0 | Cluster_10068 |
| 550.78000 | 35.0 | Cluster_29465 |
| 550.84000 | 35.0 | Cluster_29476 |
| 551.01000 | 35.0 | Cluster_29483 |
| 551.29000 | 35.0 | Cluster_29499 |
| 551.57000 | 35.0 | Cluster_10082 |
| 551.97000 | 35.0 | Cluster_10089 |
| 552.36000 | 35.0 | Cluster_29545 |
| 552.74000 | 35.0 | Cluster_29553 |
| 552.75000 | 35.0 | Cluster_29548 |
| 552.96000 | 35.0 | Cluster_10104 |
| 553.55000 | 35.0 | Cluster_29579 |
| 553.63000 | 35.0 | Cluster_10116 |
| 554.58000 | 35.0 | Cluster_10130 |
| 554.60000 | 35.0 | Cluster_10136 |
| 554.97000 | 35.0 | Cluster_10144 |
| 556.25000 | 35.0 | Cluster_29692 |
| 556.68000 | 35.0 | Cluster_10177 |
| 556.96000 | 35.0 | Cluster_10184 |
| 557.73000 | 35.0 | Cluster_29750 |
| 558.23000 | 35.0 | Cluster_10209 |
| 558.65000 | 35.0 | Cluster_10215 |
| 558.81000 | 35.0 | Cluster_29792 |
| 559.29000 | 35.0 | Cluster_29811 |
| 560.35000 | 35.0 | Cluster_29845 |
| 560.65000 | 35.0 | Cluster_10245 |
| 560.83000 | 35.0 | Cluster_29862 |
| 561.28000 | 35.0 | Cluster_29879 |

| | | |
|---|---|---|
| 561.64000 | 35.0 | Cluster_10262 |
| 561.70000 | 35.0 | Cluster_29892 |
| 561.81000 | 35.0 | Cluster_29901 |
| 561.98000 | 35.0 | Cluster_10266 |
| 562.07000 | 35.0 | Cluster_29906 |
| 562.29000 | 35.0 | Cluster_29915 |
| 563.32000 | 35.0 | Cluster_10288 |
| 563.50000 | 35.0 | Cluster_02355 |
| 563.75000 | 35.0 | Cluster_29975 |
| 563.76000 | 35.0 | Cluster_29965 |
| 564.78000 | 35.0 | Cluster_30003 |
| 564.79000 | 35.0 | Cluster_02366 |
| 565.56000 | 35.0 | Cluster_30039 |
| 565.60000 | 35.0 | Cluster_10318 |
| 566.33000 | 35.0 | Cluster_30080 |
| 566.74000 | 35.0 | Cluster_02376 |
| 566.79000 | 35.0 | Cluster_30088 |
| 566.90000 | 35.0 | Cluster_30092 |
| 566.98000 | 35.0 | Cluster_10333 |
| 567.07000 | 35.0 | Cluster_30105 |
| 567.24000 | 35.0 | Cluster_30110 |
| 567.30000 | 35.0 | Cluster_30123 |
| 567.31000 | 35.0 | Cluster_30114 |
| 568.56000 | 35.0 | Cluster_02384 |
| 568.73000 | 35.0 | Cluster_30166 |
| 568.79000 | 35.0 | Cluster_30177 |
| 568.80000 | 35.0 | Cluster_30176 |
| 568.85000 | 35.0 | Cluster_30178 |
| 569.06000 | 35.0 | Cluster_02386 |
| 569.57000 | 35.0 | Cluster_02390 |
| 569.82000 | 35.0 | Cluster_30204 |
| 570.30000 | 35.0 | Cluster_02396 |
| 570.36000 | 35.0 | Cluster_30219 |
| 570.73000 | 35.0 | Cluster_02397 |
| 570.77000 | 35.0 | Cluster_30236 |
| 571.22000 | 35.0 | Cluster_30252 |
| 571.34000 | 35.0 | Cluster_30256 |
| 571.64000 | 35.0 | Cluster_10409 |
| 571.77000 | 35.0 | Cluster_30264 |
| 572.13000 | 35.0 | Cluster_00213 |
| 572.30000 | 35.0 | Cluster_10420 |
| 572.79000 | 35.0 | Cluster_30301 |
| 573.35000 | 35.0 | Cluster_30317 |
| 574.26000 | 35.0 | Cluster_10456 |
| 574.80000 | 35.0 | Cluster_30355 |
| 575.25000 | 35.0 | Cluster_02433 |
| 575.33000 | 35.0 | Cluster_02435 |
| 576.55000 | 35.0 | Cluster_02439 |
| 576.77000 | 35.0 | Cluster_30417 |
| 576.98000 | 35.0 | Cluster_10494 |
| 577.30000 | 35.0 | Cluster_10500 |

| | | |
|---|---|---|
| 577.81000 | 35.0 | Cluster_30446 |
| 578.32000 | 35.0 | Cluster_30460 |
| 578.55000 | 35.0 | Cluster_02454 |
| 578.86000 | 35.0 | Cluster_30486 |
| 578.98000 | 35.0 | Cluster_10523 |
| 579.33000 | 35.0 | Cluster_30494 |
| 579.55000 | 35.0 | Cluster_02460 |
| 579.61000 | 35.0 | Cluster_10531 |
| 579.63000 | 35.0 | Cluster_10533 |
| 579.77000 | 35.0 | Cluster_30512 |
| 579.81000 | 35.0 | Cluster_30517 |
| 579.83000 | 35.0 | Cluster_30515 |
| 580.36000 | 35.0 | Cluster_30555 |
| 580.86000 | 35.0 | Cluster_02466 |
| 581.28000 | 35.0 | Cluster_02468 |
| 581.31000 | 35.0 | Cluster_30577 |
| 581.96000 | 35.0 | Cluster_10563 |
| 582.31000 | 35.0 | Cluster_30648 |
| 582.54000 | 35.0 | Cluster_02478 |
| 582.65000 | 35.0 | Cluster_10574 |
| 582.70000 | 35.0 | Cluster_00226 |
| 582.74000 | 35.0 | Cluster_30654 |
| 584.05000 | 35.0 | Cluster_30718 |
| 584.31000 | 35.0 | Cluster_30724 |
| 584.54000 | 35.0 | Cluster_30733 |
| 585.02000 | 35.0 | Cluster_30763 |
| 585.29000 | 35.0 | Cluster_10612 |
| 585.35000 | 35.0 | Cluster_02501 |
| 585.80000 | 35.0 | Cluster_30791 |
| 585.84000 | 35.0 | Cluster_30792 |
| 586.62000 | 35.0 | Cluster_10630 |
| 586.75000 | 35.0 | Cluster_30815 |
| 587.30000 | 35.0 | Cluster_10642 |
| 587.33000 | 35.0 | Cluster_30848 |
| 587.34000 | 35.0 | Cluster_30837 |
| 587.65000 | 35.0 | Cluster_10646 |
| 588.69000 | 35.0 | Cluster_30892 |
| 588.82000 | 35.0 | Cluster_30900 |
| 589.55000 | 35.0 | Cluster_30932 |
| 589.74000 | 35.0 | Cluster_30933 |
| 589.96000 | 35.0 | Cluster_10677 |
| 590.22000 | 35.0 | Cluster_30952 |
| 590.30000 | 35.0 | Cluster_30953 |
| 591.04000 | 35.0 | Cluster_30998 |
| 591.44000 | 35.0 | Cluster_31008 |
| 591.52000 | 35.0 | Cluster_02538 |
| 591.66000 | 35.0 | Cluster_10703 |
| 591.81000 | 35.0 | Cluster_31027 |
| 592.01000 | 35.0 | Cluster_10706 |
| 592.79000 | 35.0 | Cluster_31056 |
| 592.82000 | 35.0 | Cluster_31060 |

| | | |
|---|---|---|
| 592.91000 | 35.0 | Cluster_31063 |
| 593.06000 | 35.0 | Cluster_02547 |
| 593.07000 | 35.0 | Cluster_02548 |
| 593.33000 | 35.0 | Cluster_10724 |
| 593.34000 | 35.0 | Cluster_31073 |
| 593.93000 | 35.0 | Cluster_10743 |
| 594.32000 | 35.0 | Cluster_10753 |
| 595.59000 | 35.0 | Cluster_31146 |
| 595.62000 | 35.0 | Cluster_10769 |
| 596.75000 | 35.0 | Cluster_31177 |
| 597.67000 | 35.0 | Cluster_10808 |
| 597.81000 | 35.0 | Cluster_31212 |
| 597.82000 | 35.0 | Cluster_31208 |
| 598.29000 | 35.0 | Cluster_10814 |
| 598.36000 | 35.0 | Cluster_10818 |
| 599.36000 | 35.0 | Cluster_31267 |
| 599.66000 | 35.0 | Cluster_10840 |
| 599.95000 | 35.0 | Cluster_10848 |
| 600.81000 | 35.0 | Cluster_31316 |
| 600.97000 | 35.0 | Cluster_10858 |
| 601.36000 | 35.0 | Cluster_31339 |
| 602.32000 | 35.0 | Cluster_31390 |
| 602.83000 | 35.0 | Cluster_31406 |
| 603.30000 | 35.0 | Cluster_31409 |
| 603.51000 | 35.0 | Cluster_02598 |
| 603.53000 | 35.0 | Cluster_31419 |
| 603.96000 | 35.0 | Cluster_10888 |
| 604.80000 | 35.0 | Cluster_31463 |
| 605.31000 | 35.0 | Cluster_31473 |
| 605.67000 | 35.0 | Cluster_10910 |
| 606.07000 | 35.0 | Cluster_31501 |
| 606.27000 | 35.0 | Cluster_31514 |
| 606.87000 | 35.0 | Cluster_31531 |
| 606.89000 | 35.0 | Cluster_10941 |
| 607.32000 | 35.0 | Cluster_31536 |
| 608.09000 | 35.0 | Cluster_02626 |
| 608.32000 | 35.0 | Cluster_31587 |
| 608.49000 | 35.0 | Cluster_31590 |
| 608.78000 | 35.0 | Cluster_31595 |
| 608.79000 | 35.0 | Cluster_31601 |
| 609.06000 | 35.0 | Cluster_31605 |
| 609.81000 | 35.0 | Cluster_31625 |
| 610.31000 | 35.0 | Cluster_31655 |
| 611.34000 | 35.0 | Cluster_31685 |
| 612.32000 | 35.0 | Cluster_11010 |
| 613.83000 | 35.0 | Cluster_31794 |
| 614.06000 | 35.0 | Cluster_31796 |
| 614.07000 | 35.0 | Cluster_02670 |
| 614.19000 | 35.0 | Cluster_31797 |
| 614.26000 | 35.0 | Cluster_11035 |
| 615.83000 | 35.0 | Cluster_31852 |

| | | |
|---|---|---|
| 616.08000 | 35.0 | Cluster_02680 |
| 616.32000 | 35.0 | Cluster_11075 |
| 616.55000 | 35.0 | Cluster_31873 |
| 616.83000 | 35.0 | Cluster_31882 |
| 617.02000 | 35.0 | Cluster_31892 |
| 617.35000 | 35.0 | Cluster_31911 |
| 617.93000 | 35.0 | Cluster_31935 |
| 618.05000 | 35.0 | Cluster_02693 |
| 618.79000 | 35.0 | Cluster_31952 |
| 619.29000 | 35.0 | Cluster_02701 |
| 619.33000 | 35.0 | Cluster_31985 |
| 619.62000 | 35.0 | Cluster_11116 |
| 620.53000 | 35.0 | Cluster_32028 |
| 620.60000 | 35.0 | Cluster_11128 |
| 621.30000 | 35.0 | Cluster_32049 |
| 621.36000 | 35.0 | Cluster_32053 |
| 621.39000 | 35.0 | Cluster_32052 |
| 621.79000 | 35.0 | Cluster_32057 |
| 621.98000 | 35.0 | Cluster_11154 |
| 622.78000 | 35.0 | Cluster_32095 |
| 622.98000 | 35.0 | Cluster_11175 |
| 623.28000 | 35.0 | Cluster_32112 |
| 623.36000 | 35.0 | Cluster_32126 |
| 623.65000 | 35.0 | Cluster_11191 |
| 623.98000 | 35.0 | Cluster_11189 |
| 624.20000 | 35.0 | Cluster_32153 |
| 624.28000 | 35.0 | Cluster_32154 |
| 624.63000 | 35.0 | Cluster_11202 |
| 625.04000 | 35.0 | Cluster_02732 |
| 625.32000 | 35.0 | Cluster_11203 |
| 625.58000 | 35.0 | Cluster_02735 |
| 626.00000 | 35.0 | Cluster_11225 |
| 626.07000 | 35.0 | Cluster_32208 |
| 626.27000 | 35.0 | Cluster_32218 |
| 626.28000 | 35.0 | Cluster_32212 |
| 626.83000 | 35.0 | Cluster_32244 |
| 627.30000 | 35.0 | Cluster_32266 |
| 627.31000 | 35.0 | Cluster_32263 |
| 627.50000 | 35.0 | Cluster_32275 |
| 627.79000 | 35.0 | Cluster_32283 |
| 628.07000 | 35.0 | Cluster_32292 |
| 628.86000 | 35.0 | Cluster_32321 |
| 629.63000 | 35.0 | Cluster_11289 |
| 629.83000 | 35.0 | Cluster_32344 |
| 629.96000 | 35.0 | Cluster_11292 |
| 630.84000 | 35.0 | Cluster_32375 |
| 631.37000 | 35.0 | Cluster_11312 |
| 631.80000 | 35.0 | Cluster_32410 |
| 632.05000 | 35.0 | Cluster_11324 |
| 632.35000 | 35.0 | Cluster_11329 |
| 632.78000 | 35.0 | Cluster_32445 |

| | | |
|---|---|---|
| 632.84000 | 35.0 | Cluster_32450 |
| 633.00000 | 35.0 | Cluster_11339 |
| 633.32000 | 35.0 | Cluster_32465 |
| 633.36000 | 35.0 | Cluster_32472 |
| 635.07000 | 35.0 | Cluster_02772 |
| 635.82000 | 35.0 | Cluster_02775 |
| 636.32000 | 35.0 | Cluster_32584 |
| 636.84000 | 35.0 | Cluster_32596 |
| 637.62000 | 35.0 | Cluster_11403 |
| 637.83000 | 35.0 | Cluster_32632 |
| 638.35000 | 35.0 | Cluster_32655 |
| 638.36000 | 35.0 | Cluster_11416 |
| 638.39000 | 35.0 | Cluster_32659 |
| 638.79000 | 35.0 | Cluster_32669 |
| 638.85000 | 35.0 | Cluster_32673 |
| 639.85000 | 35.0 | Cluster_32724 |
| 639.99000 | 35.0 | Cluster_11446 |
| 640.32000 | 35.0 | Cluster_32732 |
| 641.64000 | 35.0 | Cluster_11472 |
| 641.84000 | 35.0 | Cluster_32794 |
| 642.02000 | 35.0 | Cluster_11476 |
| 643.07000 | 35.0 | Cluster_32838 |
| 643.83000 | 35.0 | Cluster_32863 |
| 644.00000 | 35.0 | Cluster_11510 |
| 644.30000 | 35.0 | Cluster_02836 |
| 644.36000 | 35.0 | Cluster_32878 |
| 644.37000 | 35.0 | Cluster_11511 |
| 645.00000 | 35.0 | Cluster_11529 |
| 645.85000 | 35.0 | Cluster_32926 |
| 646.30000 | 35.0 | Cluster_32945 |
| 648.34000 | 35.0 | Cluster_33005 |
| 648.67000 | 35.0 | Cluster_11566 |
| 648.88000 | 35.0 | Cluster_33029 |
| 649.01000 | 35.0 | Cluster_11574 |
| 649.77000 | 35.0 | Cluster_33054 |
| 650.31000 | 35.0 | Cluster_33085 |
| 651.01000 | 35.0 | Cluster_11595 |
| 651.08000 | 35.0 | Cluster_33114 |
| 651.32000 | 35.0 | Cluster_33124 |
| 651.36000 | 35.0 | Cluster_11600 |
| 651.86000 | 35.0 | Cluster_33158 |
| 651.99000 | 35.0 | Cluster_11614 |
| 652.06000 | 35.0 | Cluster_33159 |
| 652.35000 | 35.0 | Cluster_33162 |
| 652.86000 | 35.0 | Cluster_33187 |
| 653.33000 | 35.0 | Cluster_33199 |
| 653.57000 | 35.0 | Cluster_33207 |
| 653.60000 | 35.0 | Cluster_02883 |
| 654.05000 | 35.0 | Cluster_33221 |
| 654.68000 | 35.0 | Cluster_11655 |
| 654.78000 | 35.0 | Cluster_33237 |

| | | |
|---|---|---|
| 655.99000 | 35.0 | Cluster_11675 |
| 656.63000 | 35.0 | Cluster_11676 |
| 657.90000 | 35.0 | Cluster_33331 |
| 657.95000 | 35.0 | Cluster_33350 |
| 658.07000 | 35.0 | Cluster_33340 |
| 658.66000 | 35.0 | Cluster_11699 |
| 659.28000 | 35.0 | Cluster_33386 |
| 659.60000 | 35.0 | Cluster_33396 |
| 659.63000 | 35.0 | Cluster_02907 |
| 659.98000 | 35.0 | Cluster_11725 |
| 660.33000 | 35.0 | Cluster_33416 |
| 660.46000 | 35.0 | Cluster_33423 |
| 660.84000 | 35.0 | Cluster_33430 |
| 661.01000 | 35.0 | Cluster_11741 |
| 661.31000 | 35.0 | Cluster_33436 |
| 662.02000 | 35.0 | Cluster_11748 |
| 662.12000 | 35.0 | Cluster_02928 |
| 662.30000 | 35.0 | Cluster_33472 |
| 663.37000 | 35.0 | Cluster_33509 |
| 663.55000 | 35.0 | Cluster_02941 |
| 663.60000 | 35.0 | Cluster_02940 |
| 665.32000 | 35.0 | Cluster_33570 |
| 666.35000 | 35.0 | Cluster_02964 |
| 666.68000 | 35.0 | Cluster_11814 |
| 666.95000 | 35.0 | Cluster_33617 |
| 667.82000 | 35.0 | Cluster_33644 |
| 668.33000 | 35.0 | Cluster_33651 |
| 668.35000 | 35.0 | Cluster_11848 |
| 669.10000 | 35.0 | Cluster_02977 |
| 669.33000 | 35.0 | Cluster_33680 |
| 669.78000 | 35.0 | Cluster_33695 |
| 670.01000 | 35.0 | Cluster_11876 |
| 670.40000 | 35.0 | Cluster_33725 |
| 670.80000 | 35.0 | Cluster_33734 |
| 670.82000 | 35.0 | Cluster_33742 |
| 670.87000 | 35.0 | Cluster_02986 |
| 671.44000 | 35.0 | Cluster_33760 |
| 671.82000 | 35.0 | Cluster_33770 |
| 671.97000 | 35.0 | Cluster_11894 |
| 672.35000 | 35.0 | Cluster_33781 |
| 672.81000 | 35.0 | Cluster_33789 |
| 673.09000 | 35.0 | Cluster_03005 |
| 673.37000 | 35.0 | Cluster_33802 |
| 673.62000 | 35.0 | Cluster_33824 |
| 673.65000 | 35.0 | Cluster_11927 |
| 674.11000 | 35.0 | Cluster_03014 |
| 674.32000 | 35.0 | Cluster_11933 |
| 674.95000 | 35.0 | Cluster_00274 |
| 675.37000 | 35.0 | Cluster_33861 |
| 675.60000 | 35.0 | Cluster_33864 |
| 675.64000 | 35.0 | Cluster_03025 |

| | | |
|---|---|---|
| 675.84000 | 35.0 | Cluster_03026 |
| 676.00000 | 35.0 | Cluster_11956 |
| 676.32000 | 35.0 | Cluster_11959 |
| 677.20000 | 35.0 | Cluster_33905 |
| 677.89000 | 35.0 | Cluster_33946 |
| 678.81000 | 35.0 | Cluster_33981 |
| 678.84000 | 35.0 | Cluster_33980 |
| 678.85000 | 35.0 | Cluster_03050 |
| 679.38000 | 35.0 | Cluster_33996 |
| 679.79000 | 35.0 | Cluster_34007 |
| 679.86000 | 35.0 | Cluster_34018 |
| 680.08000 | 35.0 | Cluster_03060 |
| 680.09000 | 35.0 | Cluster_34023 |
| 680.87000 | 35.0 | Cluster_34054 |
| 681.08000 | 35.0 | Cluster_03066 |
| 681.32000 | 35.0 | Cluster_12020 |
| 681.35000 | 35.0 | Cluster_34074 |
| 682.09000 | 35.0 | Cluster_34089 |
| 682.31000 | 35.0 | Cluster_34106 |
| 682.38000 | 35.0 | Cluster_03074 |
| 683.61000 | 35.0 | Cluster_12059 |
| 683.81000 | 35.0 | Cluster_34134 |
| 683.99000 | 35.0 | Cluster_12062 |
| 684.08000 | 35.0 | Cluster_03088 |
| 684.82000 | 35.0 | Cluster_34171 |
| 684.83000 | 35.0 | Cluster_34174 |
| 685.09000 | 35.0 | Cluster_03095 |
| 685.10000 | 35.0 | Cluster_03102 |
| 685.35000 | 35.0 | Cluster_34201 |
| 686.04000 | 35.0 | Cluster_12086 |
| 686.50000 | 35.0 | Cluster_00285 |
| 686.75000 | 35.0 | Cluster_00286 |
| 687.66000 | 35.0 | Cluster_34255 |
| 689.34000 | 35.0 | Cluster_34298 |
| 689.36000 | 35.0 | Cluster_34295 |
| 689.57000 | 35.0 | Cluster_03142 |
| 690.82000 | 35.0 | Cluster_34324 |
| 690.84000 | 35.0 | Cluster_34327 |
| 691.59000 | 35.0 | Cluster_03162 |
| 691.79000 | 35.0 | Cluster_34357 |
| 691.83000 | 35.0 | Cluster_34369 |
| 692.26000 | 35.0 | Cluster_34373 |
| 692.32000 | 35.0 | Cluster_03167 |
| 692.35000 | 35.0 | Cluster_03168 |
| 692.60000 | 35.0 | Cluster_03182 |
| 692.84000 | 35.0 | Cluster_34402 |
| 693.56000 | 35.0 | Cluster_03192 |
| 693.60000 | 35.0 | Cluster_34413 |
| 693.88000 | 35.0 | Cluster_34418 |
| 694.36000 | 35.0 | Cluster_34424 |
| 694.84000 | 35.0 | Cluster_34436 |

| | | |
|---|---|---|
| 694.87000 | 35.0 | Cluster_34443 |
| 695.01000 | 35.0 | Cluster_12183 |
| 695.36000 | 35.0 | Cluster_34458 |
| 695.70000 | 35.0 | Cluster_12190 |
| 695.87000 | 35.0 | Cluster_34468 |
| 696.32000 | 35.0 | Cluster_34489 |
| 696.34000 | 35.0 | Cluster_03210 |
| 696.39000 | 35.0 | Cluster_34483 |
| 696.58000 | 35.0 | Cluster_03211 |
| 697.31000 | 35.0 | Cluster_03215 |
| 697.37000 | 35.0 | Cluster_34515 |
| 697.85000 | 35.0 | Cluster_34520 |
| 697.87000 | 35.0 | Cluster_34523 |
| 698.82000 | 35.0 | Cluster_34541 |
| 698.83000 | 35.0 | Cluster_34546 |
| 699.70000 | 35.0 | Cluster_12244 |
| 699.85000 | 35.0 | Cluster_03235 |
| 700.26000 | 35.0 | Cluster_34580 |
| 700.36000 | 35.0 | Cluster_34589 |
| 700.65000 | 35.0 | Cluster_34590 |
| 700.85000 | 35.0 | Cluster_34594 |
| 701.36000 | 35.0 | Cluster_34614 |
| 702.34000 | 35.0 | Cluster_34653 |
| 702.62000 | 35.0 | Cluster_12286 |
| 703.04000 | 35.0 | Cluster_12291 |
| 703.05000 | 35.0 | Cluster_03266 |
| 703.33000 | 35.0 | Cluster_34677 |
| 703.69000 | 35.0 | Cluster_12310 |
| 703.91000 | 35.0 | Cluster_03274 |
| 704.17000 | 35.0 | Cluster_34698 |
| 704.21000 | 35.0 | Cluster_34700 |
| 704.31000 | 35.0 | Cluster_34706 |
| 704.81000 | 35.0 | Cluster_34728 |
| 704.85000 | 35.0 | Cluster_34734 |
| 705.04000 | 35.0 | Cluster_12319 |
| 705.37000 | 35.0 | Cluster_34742 |
| 705.83000 | 35.0 | Cluster_34756 |
| 705.88000 | 35.0 | Cluster_34757 |
| 706.86000 | 35.0 | Cluster_03301 |
| 707.02000 | 35.0 | Cluster_12346 |
| 707.80000 | 35.0 | Cluster_34791 |
| 708.51000 | 35.0 | Cluster_00309 |
| 709.28000 | 35.0 | Cluster_34840 |
| 709.39000 | 35.0 | Cluster_34846 |
| 710.88000 | 35.0 | Cluster_03335 |
| 711.06000 | 35.0 | Cluster_12397 |
| 711.85000 | 35.0 | Cluster_34925 |
| 712.39000 | 35.0 | Cluster_34948 |
| 712.84000 | 35.0 | Cluster_34967 |
| 713.08000 | 35.0 | Cluster_03351 |
| 713.29000 | 35.0 | Cluster_03356 |

| | | |
|---|---|---|
| 713.38000 | 35.0 | Cluster_12431 |
| 713.72000 | 35.0 | Cluster_12437 |
| 713.77000 | 35.0 | Cluster_34987 |
| 713.92000 | 35.0 | Cluster_34998 |
| 714.32000 | 35.0 | Cluster_35004 |
| 714.36000 | 35.0 | Cluster_35010 |
| 715.38000 | 35.0 | Cluster_35041 |
| 715.54000 | 35.0 | Cluster_00320 |
| 715.71000 | 35.0 | Cluster_12456 |
| 715.85000 | 35.0 | Cluster_35046 |
| 715.99000 | 35.0 | Cluster_12465 |
| 716.39000 | 35.0 | Cluster_35061 |
| 717.36000 | 35.0 | Cluster_12478 |
| 717.90000 | 35.0 | Cluster_35102 |
| 718.36000 | 35.0 | Cluster_35122 |
| 718.85000 | 35.0 | Cluster_35127 |
| 718.89000 | 35.0 | Cluster_35137 |
| 718.91000 | 35.0 | Cluster_35135 |
| 718.96000 | 35.0 | Cluster_12511 |
| 719.39000 | 35.0 | Cluster_35145 |
| 719.83000 | 35.0 | Cluster_35161 |
| 719.85000 | 35.0 | Cluster_03411 |
| 719.98000 | 35.0 | Cluster_12519 |
| 720.12000 | 35.0 | Cluster_35168 |
| 720.13000 | 35.0 | Cluster_35170 |
| 720.35000 | 35.0 | Cluster_03417 |
| 720.58000 | 35.0 | Cluster_03421 |
| 720.83000 | 35.0 | Cluster_35178 |
| 721.32000 | 35.0 | Cluster_35195 |
| 721.41000 | 35.0 | Cluster_35191 |
| 721.72000 | 35.0 | Cluster_12548 |
| 721.88000 | 35.0 | Cluster_35209 |
| 721.90000 | 35.0 | Cluster_35213 |
| 722.02000 | 35.0 | Cluster_12553 |
| 722.09000 | 35.0 | Cluster_03436 |
| 722.34000 | 35.0 | Cluster_03433 |
| 722.68000 | 35.0 | Cluster_12565 |
| 723.40000 | 35.0 | Cluster_12568 |
| 723.42000 | 35.0 | Cluster_12570 |
| 723.95000 | 35.0 | Cluster_00326 |
| 724.04000 | 35.0 | Cluster_12575 |
| 724.90000 | 35.0 | Cluster_03465 |
| 725.06000 | 35.0 | Cluster_12593 |
| 725.32000 | 35.0 | Cluster_03469 |
| 725.41000 | 35.0 | Cluster_35286 |
| 725.42000 | 35.0 | Cluster_35293 |
| 727.38000 | 35.0 | Cluster_35317 |
| 728.10000 | 35.0 | Cluster_03487 |
| 728.13000 | 35.0 | Cluster_03484 |
| 728.42000 | 35.0 | Cluster_35338 |
| 728.98000 | 35.0 | Cluster_12653 |

| | | |
|---|---|---|
| 729.15000 | 35.0 | Cluster_03497 |
| 729.84000 | 35.0 | Cluster_35373 |
| 729.89000 | 35.0 | Cluster_03507 |
| 730.91000 | 35.0 | Cluster_35408 |
| 731.10000 | 35.0 | Cluster_35410 |
| 731.83000 | 35.0 | Cluster_03518 |
| 732.11000 | 35.0 | Cluster_03522 |
| 732.64000 | 35.0 | Cluster_03534 |
| 733.03000 | 35.0 | Cluster_12708 |
| 733.13000 | 35.0 | Cluster_03548 |
| 733.83000 | 35.0 | Cluster_35471 |
| 733.85000 | 35.0 | Cluster_35474 |
| 734.58000 | 35.0 | Cluster_35493 |
| 735.04000 | 35.0 | Cluster_12729 |
| 735.74000 | 35.0 | Cluster_12741 |
| 736.37000 | 35.0 | Cluster_35541 |
| 737.04000 | 35.0 | Cluster_12754 |
| 737.16000 | 35.0 | Cluster_35560 |
| 737.82000 | 35.0 | Cluster_35593 |
| 738.39000 | 35.0 | Cluster_35611 |
| 738.63000 | 35.0 | Cluster_03590 |
| 739.38000 | 35.0 | Cluster_12791 |
| 739.39000 | 35.0 | Cluster_12789 |
| 739.84000 | 35.0 | Cluster_35645 |
| 740.37000 | 35.0 | Cluster_35657 |
| 740.38000 | 35.0 | Cluster_35661 |
| 740.93000 | 35.0 | Cluster_35676 |
| 741.85000 | 35.0 | Cluster_35683 |
| 742.37000 | 35.0 | Cluster_35697 |
| 742.67000 | 35.0 | Cluster_12831 |
| 742.84000 | 35.0 | Cluster_35707 |
| 744.37000 | 35.0 | Cluster_35750 |
| 745.38000 | 35.0 | Cluster_35777 |
| 745.68000 | 35.0 | Cluster_12857 |
| 746.06000 | 35.0 | Cluster_12863 |
| 746.15000 | 35.0 | Cluster_35792 |
| 746.30000 | 35.0 | Cluster_35797 |
| 747.12000 | 35.0 | Cluster_35817 |
| 747.37000 | 35.0 | Cluster_35818 |
| 748.00000 | 35.0 | Cluster_12903 |
| 748.86000 | 35.0 | Cluster_03654 |
| 748.88000 | 35.0 | Cluster_35857 |
| 749.07000 | 35.0 | Cluster_12920 |
| 749.69000 | 35.0 | Cluster_12929 |
| 749.98000 | 35.0 | Cluster_12936 |
| 750.01000 | 35.0 | Cluster_12941 |
| 750.38000 | 35.0 | Cluster_35883 |
| 750.44000 | 35.0 | Cluster_35886 |
| 750.94000 | 35.0 | Cluster_35903 |
| 751.06000 | 35.0 | Cluster_12961 |
| 751.15000 | 35.0 | Cluster_03662 |

| | | |
|---|---|---|
| 751.36000 | 35.0 | Cluster_35908 |
| 752.37000 | 35.0 | Cluster_35932 |
| 753.05000 | 35.0 | Cluster_12988 |
| 753.06000 | 35.0 | Cluster_12984 |
| 753.39000 | 35.0 | Cluster_35964 |
| 753.91000 | 35.0 | Cluster_35981 |
| 754.06000 | 35.0 | Cluster_13009 |
| 754.90000 | 35.0 | Cluster_36004 |
| 755.06000 | 35.0 | Cluster_13021 |
| 755.30000 | 35.0 | Cluster_13019 |
| 755.33000 | 35.0 | Cluster_13020 |
| 755.42000 | 35.0 | Cluster_36013 |
| 755.75000 | 35.0 | Cluster_13034 |
| 755.92000 | 35.0 | Cluster_36043 |
| 756.13000 | 35.0 | Cluster_03688 |
| 757.31000 | 35.0 | Cluster_36071 |
| 757.73000 | 35.0 | Cluster_13076 |
| 758.41000 | 35.0 | Cluster_36102 |
| 758.70000 | 35.0 | Cluster_13088 |
| 759.63000 | 35.0 | Cluster_03704 |
| 759.90000 | 35.0 | Cluster_36129 |
| 759.91000 | 35.0 | Cluster_36132 |
| 760.35000 | 35.0 | Cluster_36146 |
| 761.05000 | 35.0 | Cluster_00024 |
| 761.41000 | 35.0 | Cluster_36182 |
| 762.38000 | 35.0 | Cluster_13130 |
| 763.03000 | 35.0 | Cluster_13141 |
| 763.08000 | 35.0 | Cluster_13137 |
| 763.35000 | 35.0 | Cluster_03726 |
| 763.71000 | 35.0 | Cluster_13147 |
| 764.09000 | 35.0 | Cluster_13148 |
| 764.36000 | 35.0 | Cluster_36253 |
| 765.15000 | 35.0 | Cluster_03741 |
| 765.79000 | 35.0 | Cluster_36274 |
| 765.80000 | 35.0 | Cluster_36275 |
| 766.37000 | 35.0 | Cluster_36292 |
| 766.55000 | 35.0 | Cluster_00339 |
| 767.70000 | 35.0 | Cluster_13203 |
| 767.85000 | 35.0 | Cluster_36325 |
| 768.07000 | 35.0 | Cluster_13209 |
| 768.08000 | 35.0 | Cluster_13205 |
| 768.38000 | 35.0 | Cluster_36351 |
| 769.33000 | 35.0 | Cluster_36370 |
| 769.59000 | 35.0 | Cluster_36368 |
| 769.82000 | 35.0 | Cluster_36385 |
| 772.41000 | 35.0 | Cluster_36441 |
| 772.74000 | 35.0 | Cluster_13265 |
| 772.80000 | 35.0 | Cluster_13273 |
| 772.88000 | 35.0 | Cluster_36459 |
| 772.92000 | 35.0 | Cluster_36460 |
| 773.08000 | 35.0 | Cluster_03785 |

| | | |
|---|---|---|
| 773.09000 | 35.0 | Cluster_03786 |
| 773.13000 | 35.0 | Cluster_13272 |
| 773.35000 | 35.0 | Cluster_13283 |
| 773.41000 | 35.0 | Cluster_13286 |
| 774.12000 | 35.0 | Cluster_36481 |
| 774.73000 | 35.0 | Cluster_13306 |
| 775.06000 | 35.0 | Cluster_13312 |
| 775.07000 | 35.0 | Cluster_13311 |
| 775.42000 | 35.0 | Cluster_36510 |
| 776.37000 | 35.0 | Cluster_03803 |
| 777.15000 | 35.0 | Cluster_03806 |
| 777.35000 | 35.0 | Cluster_36561 |
| 777.38000 | 35.0 | Cluster_36567 |
| 777.39000 | 35.0 | Cluster_36564 |
| 778.37000 | 35.0 | Cluster_36591 |
| 778.77000 | 35.0 | Cluster_13350 |
| 778.87000 | 35.0 | Cluster_36605 |
| 780.73000 | 35.0 | Cluster_13379 |
| 780.81000 | 35.0 | Cluster_36636 |
| 780.88000 | 35.0 | Cluster_03821 |
| 781.40000 | 35.0 | Cluster_36661 |
| 782.03000 | 35.0 | Cluster_13403 |
| 783.13000 | 35.0 | Cluster_03834 |
| 783.46000 | 35.0 | Cluster_36708 |
| 783.72000 | 35.0 | Cluster_13422 |
| 784.06000 | 35.0 | Cluster_13433 |
| 784.89000 | 35.0 | Cluster_36733 |
| 784.93000 | 35.0 | Cluster_36732 |
| 785.45000 | 35.0 | Cluster_36744 |
| 785.83000 | 35.0 | Cluster_36763 |
| 787.06000 | 35.0 | Cluster_13459 |
| 787.12000 | 35.0 | Cluster_03868 |
| 787.35000 | 35.0 | Cluster_03870 |
| 788.87000 | 35.0 | Cluster_36840 |
| 789.06000 | 35.0 | Cluster_13483 |
| 790.87000 | 35.0 | Cluster_36906 |
| 791.12000 | 35.0 | Cluster_36909 |
| 791.38000 | 35.0 | Cluster_36914 |
| 791.90000 | 35.0 | Cluster_36919 |
| 792.38000 | 35.0 | Cluster_36933 |
| 792.64000 | 35.0 | Cluster_36941 |
| 792.86000 | 35.0 | Cluster_03914 |
| 792.87000 | 35.0 | Cluster_36948 |
| 792.97000 | 35.0 | Cluster_13541 |
| 793.78000 | 35.0 | Cluster_13547 |
| 794.36000 | 35.0 | Cluster_36984 |
| 794.37000 | 35.0 | Cluster_36982 |
| 794.45000 | 35.0 | Cluster_36985 |
| 794.65000 | 35.0 | Cluster_36986 |
| 795.07000 | 35.0 | Cluster_13571 |
| 795.42000 | 35.0 | Cluster_37004 |

| | | |
|---|---|---|
| 795.44000 | 35.0 | Cluster_37000 |
| 795.68000 | 35.0 | Cluster_13585 |
| 795.87000 | 35.0 | Cluster_03947 |
| 795.90000 | 35.0 | Cluster_37017 |
| 795.96000 | 35.0 | Cluster_37020 |
| 796.41000 | 35.0 | Cluster_13605 |
| 796.72000 | 35.0 | Cluster_13597 |
| 796.91000 | 35.0 | Cluster_37044 |
| 797.86000 | 35.0 | Cluster_37077 |
| 797.87000 | 35.0 | Cluster_37073 |
| 797.88000 | 35.0 | Cluster_37072 |
| 797.89000 | 35.0 | Cluster_37068 |
| 799.38000 | 35.0 | Cluster_37115 |
| 799.40000 | 35.0 | Cluster_37112 |
| 799.89000 | 35.0 | Cluster_03975 |
| 800.35000 | 35.0 | Cluster_37146 |
| 800.41000 | 35.0 | Cluster_13643 |
| 800.84000 | 35.0 | Cluster_37149 |
| 800.86000 | 35.0 | Cluster_03991 |
| 801.06000 | 35.0 | Cluster_00029 |
| 801.37000 | 35.0 | Cluster_03995 |
| 801.42000 | 35.0 | Cluster_37163 |
| 801.44000 | 35.0 | Cluster_13660 |
| 802.43000 | 35.0 | Cluster_37188 |
| 802.74000 | 35.0 | Cluster_13681 |
| 803.41000 | 35.0 | Cluster_13684 |
| 803.88000 | 35.0 | Cluster_37226 |
| 804.43000 | 35.0 | Cluster_37243 |
| 804.91000 | 35.0 | Cluster_04015 |
| 805.16000 | 35.0 | Cluster_37269 |
| 805.39000 | 35.0 | Cluster_37283 |
| 805.94000 | 35.0 | Cluster_37284 |
| 806.39000 | 35.0 | Cluster_37304 |
| 807.01000 | 35.0 | Cluster_13723 |
| 807.46000 | 35.0 | Cluster_37330 |
| 807.92000 | 35.0 | Cluster_37352 |
| 808.85000 | 35.0 | Cluster_37370 |
| 808.91000 | 35.0 | Cluster_37377 |
| 809.40000 | 35.0 | Cluster_04037 |
| 809.91000 | 35.0 | Cluster_37401 |
| 810.39000 | 35.0 | Cluster_37407 |
| 810.44000 | 35.0 | Cluster_13774 |
| 811.74000 | 35.0 | Cluster_00036 |
| 811.86000 | 35.0 | Cluster_37446 |
| 811.91000 | 35.0 | Cluster_37441 |
| 812.39000 | 35.0 | Cluster_37458 |
| 812.41000 | 35.0 | Cluster_13796 |
| 813.82000 | 35.0 | Cluster_00352 |
| 813.90000 | 35.0 | Cluster_37483 |
| 814.10000 | 35.0 | Cluster_13827 |
| 814.16000 | 35.0 | Cluster_37492 |

| | | |
|---|---|---|
| 814.17000 | 35.0 | Cluster_04067 |
| 814.90000 | 35.0 | Cluster_04074 |
| 815.06000 | 35.0 | Cluster_13832 |
| 815.79000 | 35.0 | Cluster_13844 |
| 815.92000 | 35.0 | Cluster_37530 |
| 816.38000 | 35.0 | Cluster_37540 |
| 816.43000 | 35.0 | Cluster_04088 |
| 816.72000 | 35.0 | Cluster_13857 |
| 817.34000 | 35.0 | Cluster_37561 |
| 817.88000 | 35.0 | Cluster_37570 |
| 818.42000 | 35.0 | Cluster_13883 |
| 819.05000 | 35.0 | Cluster_13897 |
| 819.42000 | 35.0 | Cluster_13906 |
| 820.17000 | 35.0 | Cluster_04104 |
| 821.23000 | 35.0 | Cluster_04111 |
| 821.38000 | 35.0 | Cluster_37637 |
| 822.06000 | 35.0 | Cluster_13934 |
| 823.42000 | 35.0 | Cluster_37669 |
| 823.91000 | 35.0 | Cluster_37674 |
| 823.94000 | 35.0 | Cluster_37688 |
| 825.32000 | 35.0 | Cluster_37717 |
| 825.39000 | 35.0 | Cluster_04137 |
| 826.38000 | 35.0 | Cluster_13981 |
| 826.86000 | 35.0 | Cluster_37765 |
| 827.04000 | 35.0 | Cluster_13986 |
| 827.06000 | 35.0 | Cluster_13988 |
| 827.09000 | 35.0 | Cluster_13989 |
| 827.88000 | 35.0 | Cluster_37794 |
| 828.80000 | 35.0 | Cluster_14000 |
| 828.98000 | 35.0 | Cluster_14004 |
| 829.16000 | 35.0 | Cluster_04165 |
| 829.36000 | 35.0 | Cluster_37821 |
| 829.43000 | 35.0 | Cluster_37825 |
| 829.91000 | 35.0 | Cluster_04169 |
| 830.74000 | 35.0 | Cluster_14020 |
| 830.89000 | 35.0 | Cluster_00042 |
| 831.45000 | 35.0 | Cluster_37853 |
| 832.41000 | 35.0 | Cluster_37873 |
| 832.68000 | 35.0 | Cluster_37879 |
| 833.49000 | 35.0 | Cluster_37894 |
| 833.99000 | 35.0 | Cluster_37902 |
| 834.10000 | 35.0 | Cluster_14054 |
| 834.94000 | 35.0 | Cluster_37918 |
| 835.05000 | 35.0 | Cluster_14062 |
| 836.39000 | 35.0 | Cluster_14076 |
| 836.42000 | 35.0 | Cluster_14077 |
| 836.76000 | 35.0 | Cluster_14083 |
| 836.86000 | 35.0 | Cluster_37959 |
| 836.94000 | 35.0 | Cluster_04234 |
| 837.41000 | 35.0 | Cluster_37970 |
| 837.95000 | 35.0 | Cluster_37976 |

| | | |
|---|---|---|
| 839.10000 | 35.0 | Cluster_14102 |
| 840.66000 | 35.0 | Cluster_04260 |
| 840.78000 | 35.0 | Cluster_14117 |
| 841.08000 | 35.0 | Cluster_14128 |
| 841.10000 | 35.0 | Cluster_14126 |
| 841.67000 | 35.0 | Cluster_00364 |
| 841.76000 | 35.0 | Cluster_38066 |
| 842.73000 | 35.0 | Cluster_14140 |
| 843.87000 | 35.0 | Cluster_38114 |
| 844.18000 | 35.0 | Cluster_04298 |
| 844.44000 | 35.0 | Cluster_14157 |
| 846.98000 | 35.0 | Cluster_38180 |
| 847.48000 | 35.0 | Cluster_38200 |
| 847.75000 | 35.0 | Cluster_14185 |
| 847.88000 | 35.0 | Cluster_38203 |
| 848.40000 | 35.0 | Cluster_04350 |
| 848.91000 | 35.0 | Cluster_04352 |
| 849.46000 | 35.0 | Cluster_38235 |
| 849.64000 | 35.0 | Cluster_04364 |
| 849.97000 | 35.0 | Cluster_38245 |
| 849.99000 | 35.0 | Cluster_38248 |
| 850.33000 | 35.0 | Cluster_38250 |
| 850.40000 | 35.0 | Cluster_04376 |
| 850.76000 | 35.0 | Cluster_14213 |
| 851.36000 | 35.0 | Cluster_14215 |
| 851.77000 | 35.0 | Cluster_38298 |
| 852.03000 | 35.0 | Cluster_38307 |
| 852.91000 | 35.0 | Cluster_04400 |
| 852.93000 | 35.0 | Cluster_04401 |
| 853.45000 | 35.0 | Cluster_38334 |
| 854.44000 | 35.0 | Cluster_38377 |
| 854.47000 | 35.0 | Cluster_38375 |
| 854.88000 | 35.0 | Cluster_38385 |
| 855.41000 | 35.0 | Cluster_38392 |
| 855.73000 | 35.0 | Cluster_14240 |
| 856.43000 | 35.0 | Cluster_14249 |
| 856.91000 | 35.0 | Cluster_04444 |
| 857.06000 | 35.0 | Cluster_14257 |
| 857.89000 | 35.0 | Cluster_38459 |
| 857.91000 | 35.0 | Cluster_04449 |
| 858.07000 | 35.0 | Cluster_14269 |
| 858.77000 | 35.0 | Cluster_14275 |
| 859.45000 | 35.0 | Cluster_38500 |
| 859.47000 | 35.0 | Cluster_38497 |
| 860.47000 | 35.0 | Cluster_14284 |
| 861.39000 | 35.0 | Cluster_04493 |
| 862.39000 | 35.0 | Cluster_04505 |
| 862.49000 | 35.0 | Cluster_38591 |
| 863.05000 | 35.0 | Cluster_14302 |
| 863.76000 | 35.0 | Cluster_14307 |
| 863.94000 | 35.0 | Cluster_38615 |

| | | |
|---|---|---|
| 864.07000 | 35.0 | Cluster_14314 |
| 865.03000 | 35.0 | Cluster_38646 |
| 865.75000 | 35.0 | Cluster_14326 |
| 865.92000 | 35.0 | Cluster_38659 |
| 865.93000 | 35.0 | Cluster_04555 |
| 866.41000 | 35.0 | Cluster_14334 |
| 869.07000 | 35.0 | Cluster_14353 |
| 869.28000 | 35.0 | Cluster_38719 |
| 869.41000 | 35.0 | Cluster_04599 |
| 869.78000 | 35.0 | Cluster_14363 |
| 871.17000 | 35.0 | Cluster_38758 |
| 871.27000 | 35.0 | Cluster_38759 |
| 871.44000 | 35.0 | Cluster_38769 |
| 871.47000 | 35.0 | Cluster_14388 |
| 872.27000 | 35.0 | Cluster_38802 |
| 874.09000 | 35.0 | Cluster_14414 |
| 874.26000 | 35.0 | Cluster_38844 |
| 874.78000 | 35.0 | Cluster_14424 |
| 875.28000 | 35.0 | Cluster_38860 |
| 875.66000 | 35.0 | Cluster_04698 |
| 876.08000 | 35.0 | Cluster_14437 |
| 876.53000 | 35.0 | Cluster_38895 |
| 877.38000 | 35.0 | Cluster_14453 |
| 877.44000 | 35.0 | Cluster_38909 |
| 878.48000 | 35.0 | Cluster_38928 |
| 880.46000 | 35.0 | Cluster_14483 |
| 881.96000 | 35.0 | Cluster_38993 |
| 882.44000 | 35.0 | Cluster_39008 |
| 882.94000 | 35.0 | Cluster_39018 |
| 884.40000 | 35.0 | Cluster_14528 |
| 884.66000 | 35.0 | Cluster_39047 |
| 885.12000 | 35.0 | Cluster_14544 |
| 885.68000 | 35.0 | Cluster_04765 |
| 885.95000 | 35.0 | Cluster_39078 |
| 886.43000 | 35.0 | Cluster_39095 |
| 886.96000 | 35.0 | Cluster_39099 |
| 887.52000 | 35.0 | Cluster_39107 |
| 887.68000 | 35.0 | Cluster_04781 |
| 887.74000 | 35.0 | Cluster_14571 |
| 888.13000 | 35.0 | Cluster_14577 |
| 888.40000 | 35.0 | Cluster_14578 |
| 888.91000 | 35.0 | Cluster_39135 |
| 888.92000 | 35.0 | Cluster_04786 |
| 889.11000 | 35.0 | Cluster_14581 |
| 889.44000 | 35.0 | Cluster_39147 |
| 890.17000 | 35.0 | Cluster_04796 |
| 890.38000 | 35.0 | Cluster_39161 |
| 890.39000 | 35.0 | Cluster_39160 |
| 890.46000 | 35.0 | Cluster_39173 |
| 890.99000 | 35.0 | Cluster_39180 |
| 891.46000 | 35.0 | Cluster_39186 |

| | | |
|---|---|---|
| 891.81000 | 35.0 | Cluster_14598 |
| 892.21000 | 35.0 | Cluster_04824 |
| 893.43000 | 35.0 | Cluster_39231 |
| 893.78000 | 35.0 | Cluster_14615 |
| 894.12000 | 35.0 | Cluster_14617 |
| 894.71000 | 35.0 | Cluster_04845 |
| 895.21000 | 35.0 | Cluster_04851 |
| 895.45000 | 35.0 | Cluster_14622 |
| 895.81000 | 35.0 | Cluster_14625 |
| 896.15000 | 35.0 | Cluster_14634 |
| 897.04000 | 35.0 | Cluster_39292 |
| 898.77000 | 35.0 | Cluster_14664 |
| 899.43000 | 35.0 | Cluster_39337 |
| 899.51000 | 35.0 | Cluster_39345 |
| 899.96000 | 35.0 | Cluster_04891 |
| 900.94000 | 35.0 | Cluster_39368 |
| 900.97000 | 35.0 | Cluster_39380 |
| 900.99000 | 35.0 | Cluster_39377 |
| 902.46000 | 35.0 | Cluster_39412 |
| 904.46000 | 35.0 | Cluster_39443 |
| 905.43000 | 35.0 | Cluster_39461 |
| 906.77000 | 35.0 | Cluster_14739 |
| 907.80000 | 35.0 | Cluster_14749 |
| 908.46000 | 35.0 | Cluster_39500 |
| 908.48000 | 35.0 | Cluster_14756 |
| 908.94000 | 35.0 | Cluster_04958 |
| 908.95000 | 35.0 | Cluster_39504 |
| 909.44000 | 35.0 | Cluster_14760 |
| 909.50000 | 35.0 | Cluster_14765 |
| 909.89000 | 35.0 | Cluster_39520 |
| 910.11000 | 35.0 | Cluster_14769 |
| 910.19000 | 35.0 | Cluster_04966 |
| 911.96000 | 35.0 | Cluster_39564 |
| 914.56000 | 35.0 | Cluster_39615 |
| 915.13000 | 35.0 | Cluster_14823 |
| 915.93000 | 35.0 | Cluster_39626 |
| 916.44000 | 35.0 | Cluster_39640 |
| 916.49000 | 35.0 | Cluster_39633 |
| 917.45000 | 35.0 | Cluster_14854 |
| 917.97000 | 35.0 | Cluster_39674 |
| 918.66000 | 35.0 | Cluster_39698 |
| 920.45000 | 35.0 | Cluster_39721 |
| 920.76000 | 35.0 | Cluster_14891 |
| 920.79000 | 35.0 | Cluster_39725 |
| 920.88000 | 35.0 | Cluster_39726 |
| 920.95000 | 35.0 | Cluster_39731 |
| 922.42000 | 35.0 | Cluster_14907 |
| 922.98000 | 35.0 | Cluster_39772 |
| 923.30000 | 35.0 | Cluster_14924 |
| 923.45000 | 35.0 | Cluster_14932 |
| 923.92000 | 35.0 | Cluster_39782 |

| | | |
|---|---|---|
| 924.49000 | 35.0 | Cluster_39788 |
| 925.11000 | 35.0 | Cluster_14961 |
| 926.42000 | 35.0 | Cluster_14975 |
| 927.48000 | 35.0 | Cluster_39834 |
| 928.47000 | 35.0 | Cluster_39858 |
| 928.96000 | 35.0 | Cluster_39861 |
| 929.12000 | 35.0 | Cluster_15009 |
| 929.43000 | 35.0 | Cluster_39880 |
| 929.90000 | 35.0 | Cluster_39897 |
| 930.47000 | 35.0 | Cluster_15020 |
| 931.73000 | 35.0 | Cluster_15043 |
| 931.90000 | 35.0 | Cluster_39924 |
| 932.64000 | 35.0 | Cluster_15062 |
| 932.95000 | 35.0 | Cluster_39941 |
| 933.14000 | 35.0 | Cluster_15072 |
| 933.33000 | 35.0 | Cluster_05038 |
| 934.00000 | 35.0 | Cluster_39963 |
| 934.13000 | 35.0 | Cluster_15084 |
| 934.43000 | 35.0 | Cluster_15083 |
| 934.78000 | 35.0 | Cluster_15089 |
| 935.78000 | 35.0 | Cluster_15100 |
| 936.45000 | 35.0 | Cluster_15116 |
| 936.80000 | 35.0 | Cluster_15129 |
| 937.00000 | 35.0 | Cluster_39997 |
| 937.44000 | 35.0 | Cluster_15144 |
| 937.49000 | 35.0 | Cluster_05051 |
| 938.07000 | 35.0 | Cluster_15154 |
| 938.13000 | 35.0 | Cluster_15156 |
| 938.21000 | 35.0 | Cluster_15160 |
| 939.69000 | 35.0 | Cluster_05058 |
| 940.46000 | 35.0 | Cluster_15180 |
| 941.72000 | 35.0 | Cluster_40098 |
| 942.00000 | 35.0 | Cluster_40094 |
| 943.45000 | 35.0 | Cluster_15229 |
| 945.13000 | 35.0 | Cluster_15244 |
| 945.39000 | 35.0 | Cluster_15242 |
| 945.72000 | 35.0 | Cluster_05075 |
| 945.73000 | 35.0 | Cluster_05078 |
| 945.78000 | 35.0 | Cluster_15250 |
| 945.98000 | 35.0 | Cluster_40152 |
| 947.16000 | 35.0 | Cluster_15264 |
| 947.22000 | 35.0 | Cluster_05084 |
| 947.79000 | 35.0 | Cluster_15273 |
| 948.19000 | 35.0 | Cluster_15277 |
| 948.52000 | 35.0 | Cluster_40180 |
| 949.14000 | 35.0 | Cluster_15301 |
| 949.44000 | 35.0 | Cluster_40190 |
| 949.46000 | 35.0 | Cluster_15302 |
| 950.44000 | 35.0 | Cluster_15314 |
| 951.17000 | 35.0 | Cluster_15324 |
| 951.96000 | 35.0 | Cluster_40227 |

| | | |
|---|---|---|
| 952.14000 | 35.0 | Cluster_15337 |
| 952.15000 | 35.0 | Cluster_15339 |
| 952.44000 | 35.0 | Cluster_15342 |
| 952.53000 | 35.0 | Cluster_40233 |
| 953.98000 | 35.0 | Cluster_40255 |
| 954.82000 | 35.0 | Cluster_15379 |
| 954.99000 | 35.0 | Cluster_40262 |
| 955.13000 | 35.0 | Cluster_15394 |
| 955.47000 | 35.0 | Cluster_15404 |
| 956.48000 | 35.0 | Cluster_15426 |
| 957.07000 | 35.0 | Cluster_40290 |
| 957.48000 | 35.0 | Cluster_40291 |
| 960.15000 | 35.0 | Cluster_15462 |
| 960.98000 | 35.0 | Cluster_40317 |
| 961.48000 | 35.0 | Cluster_15478 |
| 962.14000 | 35.0 | Cluster_15497 |
| 962.38000 | 35.0 | Cluster_40344 |
| 962.79000 | 35.0 | Cluster_15505 |
| 964.13000 | 35.0 | Cluster_15523 |
| 964.43000 | 35.0 | Cluster_40365 |
| 965.29000 | 35.0 | Cluster_40377 |
| 966.15000 | 35.0 | Cluster_15549 |
| 966.49000 | 35.0 | Cluster_40387 |
| 966.50000 | 35.0 | Cluster_05172 |
| 966.85000 | 35.0 | Cluster_15552 |
| 967.00000 | 35.0 | Cluster_40392 |
| 967.56000 | 35.0 | Cluster_15565 |
| 968.03000 | 35.0 | Cluster_40402 |
| 968.25000 | 35.0 | Cluster_05187 |
| 971.97000 | 35.0 | Cluster_40451 |
| 972.01000 | 35.0 | Cluster_40467 |
| 972.46000 | 35.0 | Cluster_15627 |
| 972.49000 | 35.0 | Cluster_15626 |
| 973.51000 | 35.0 | Cluster_40476 |
| 974.47000 | 35.0 | Cluster_15651 |
| 974.98000 | 35.0 | Cluster_40495 |
| 975.82000 | 35.0 | Cluster_15672 |
| 975.98000 | 35.0 | Cluster_05210 |
| 976.00000 | 35.0 | Cluster_05212 |
| 976.03000 | 35.0 | Cluster_40504 |
| 977.14000 | 35.0 | Cluster_15696 |
| 977.62000 | 35.0 | Cluster_40522 |
| 979.45000 | 35.0 | Cluster_40546 |
| 979.80000 | 35.0 | Cluster_15720 |
| 979.83000 | 35.0 | Cluster_15721 |
| 979.98000 | 35.0 | Cluster_40568 |
| 980.00000 | 35.0 | Cluster_40562 |
| 980.50000 | 35.0 | Cluster_40565 |
| 980.70000 | 35.0 | Cluster_40578 |
| 981.45000 | 35.0 | Cluster_15750 |
| 983.25000 | 35.0 | Cluster_05237 |

| | | |
|---|---|---|
| 983.95000 | 35.0 | Cluster_40617 |
| 985.14000 | 35.0 | Cluster_40628 |
| 986.02000 | 35.0 | Cluster_40630 |
| 986.52000 | 35.0 | Cluster_40639 |
| 986.83000 | 35.0 | Cluster_15810 |
| 987.24000 | 35.0 | Cluster_40654 |
| 987.80000 | 35.0 | Cluster_15825 |
| 988.44000 | 35.0 | Cluster_15835 |
| 991.93000 | 35.0 | Cluster_40717 |
| 993.50000 | 35.0 | Cluster_15895 |
| 997.19000 | 35.0 | Cluster_15954 |
| 1000.06000 | 35.0 | Cluster_40851 |
| 1001.03000 | 35.0 | Cluster_40878 |
| 1001.16000 | 35.0 | Cluster_15996 |
| 1001.49000 | 35.0 | Cluster_40884 |
| 1003.52000 | 35.0 | Cluster_05294 |
| 1004.00000 | 35.0 | Cluster_40937 |
| 1004.31000 | 35.0 | Cluster_05296 |
| 1004.57000 | 35.0 | Cluster_40940 |
| 1007.81000 | 35.0 | Cluster_16072 |
| 1008.78000 | 35.0 | Cluster_16080 |
| 1012.26000 | 35.0 | Cluster_05329 |
| 1012.27000 | 35.0 | Cluster_41044 |
| 1012.93000 | 35.0 | Cluster_41054 |
| 1013.01000 | 35.0 | Cluster_05341 |
| 1013.47000 | 35.0 | Cluster_16138 |
| 1013.49000 | 35.0 | Cluster_16137 |
| 1013.76000 | 35.0 | Cluster_05344 |
| 1013.90000 | 35.0 | Cluster_41056 |
| 1015.92000 | 35.0 | Cluster_41088 |
| 1016.78000 | 35.0 | Cluster_05351 |
| 1017.27000 | 35.0 | Cluster_41099 |
| 1019.25000 | 35.0 | Cluster_05360 |
| 1020.15000 | 35.0 | Cluster_16210 |
| 1022.50000 | 35.0 | Cluster_16231 |
| 1023.82000 | 35.0 | Cluster_41185 |
| 1024.48000 | 35.0 | Cluster_16254 |
| 1024.50000 | 35.0 | Cluster_16256 |
| 1024.93000 | 35.0 | Cluster_41195 |
| 1025.46000 | 35.0 | Cluster_16264 |
| 1025.81000 | 35.0 | Cluster_41211 |
| 1026.83000 | 35.0 | Cluster_16274 |
| 1027.13000 | 35.0 | Cluster_16285 |
| 1027.14000 | 35.0 | Cluster_16284 |
| 1027.88000 | 35.0 | Cluster_16291 |
| 1028.18000 | 35.0 | Cluster_16295 |
| 1028.84000 | 35.0 | Cluster_16300 |
| 1029.99000 | 35.0 | Cluster_41262 |
| 1031.20000 | 35.0 | Cluster_16323 |
| 1032.82000 | 35.0 | Cluster_16335 |
| 1033.20000 | 35.0 | Cluster_16342 |

| | | |
|---|---|---|
| 1036.52000 | 35.0 | Cluster_16382 |
| 1037.50000 | 35.0 | Cluster_16400 |
| 1039.05000 | 35.0 | Cluster_41371 |
| 1039.52000 | 35.0 | Cluster_41374 |
| 1039.81000 | 35.0 | Cluster_16429 |
| 1040.83000 | 35.0 | Cluster_05437 |
| 1041.01000 | 35.0 | Cluster_41401 |
| 1042.17000 | 35.0 | Cluster_16457 |
| 1044.49000 | 35.0 | Cluster_41432 |
| 1046.13000 | 35.0 | Cluster_16488 |
| 1046.22000 | 35.0 | Cluster_16495 |
| 1047.16000 | 35.0 | Cluster_16507 |
| 1047.83000 | 35.0 | Cluster_41470 |
| 1053.50000 | 35.0 | Cluster_16581 |
| 1053.98000 | 35.0 | Cluster_41534 |
| 1054.99000 | 35.0 | Cluster_05476 |
| 1055.22000 | 35.0 | Cluster_16595 |
| 1055.50000 | 35.0 | Cluster_16606 |
| 1056.52000 | 35.0 | Cluster_16616 |
| 1057.06000 | 35.0 | Cluster_41567 |
| 1057.33000 | 35.0 | Cluster_05486 |
| 1057.34000 | 35.0 | Cluster_05487 |
| 1057.58000 | 35.0 | Cluster_41574 |
| 1057.99000 | 35.0 | Cluster_41578 |
| 1058.02000 | 35.0 | Cluster_41582 |
| 1058.81000 | 35.0 | Cluster_16658 |
| 1058.85000 | 35.0 | Cluster_16649 |
| 1058.96000 | 35.0 | Cluster_41595 |
| 1058.97000 | 35.0 | Cluster_41597 |
| 1059.01000 | 35.0 | Cluster_41596 |
| 1060.32000 | 35.0 | Cluster_41612 |
| 1060.75000 | 35.0 | Cluster_05502 |
| 1060.82000 | 35.0 | Cluster_16679 |
| 1061.26000 | 35.0 | Cluster_05510 |
| 1061.33000 | 35.0 | Cluster_05506 |
| 1061.82000 | 35.0 | Cluster_41625 |
| 1062.83000 | 35.0 | Cluster_16700 |
| 1063.27000 | 35.0 | Cluster_05516 |
| 1063.82000 | 35.0 | Cluster_41655 |
| 1063.99000 | 35.0 | Cluster_41663 |
| 1064.72000 | 35.0 | Cluster_05521 |
| 1064.73000 | 35.0 | Cluster_41675 |
| 1064.99000 | 35.0 | Cluster_41673 |
| 1066.26000 | 35.0 | Cluster_41702 |
| 1067.52000 | 35.0 | Cluster_41715 |
| 1068.05000 | 35.0 | Cluster_41719 |
| 1068.75000 | 35.0 | Cluster_41734 |
| 1069.51000 | 35.0 | Cluster_41748 |
| 1070.16000 | 35.0 | Cluster_16781 |
| 1070.51000 | 35.0 | Cluster_05546 |
| 1072.55000 | 35.0 | Cluster_16804 |

| | | |
|---|---|---|
| 1072.76000 | 35.0 | Cluster_05555 |
| 1074.19000 | 35.0 | Cluster_16831 |
| 1074.51000 | 35.0 | Cluster_41810 |
| 1075.53000 | 35.0 | Cluster_41816 |
| 1076.75000 | 35.0 | Cluster_05574 |
| 1078.01000 | 35.0 | Cluster_05578 |
| 1079.49000 | 35.0 | Cluster_05582 |
| 1080.75000 | 35.0 | Cluster_00593 |
| 1082.00000 | 35.0 | Cluster_05588 |
| 1083.50000 | 35.0 | Cluster_05592 |
| 1086.59000 | 35.0 | Cluster_41940 |
| 1091.58000 | 35.0 | Cluster_41998 |
| 1092.90000 | 35.0 | Cluster_17025 |
| 1093.21000 | 35.0 | Cluster_17031 |
| 1093.55000 | 35.0 | Cluster_17035 |
| 1093.58000 | 35.0 | Cluster_42024 |
| 1093.99000 | 35.0 | Cluster_42029 |
| 1094.92000 | 35.0 | Cluster_00625 |
| 1096.55000 | 35.0 | Cluster_42051 |
| 1100.23000 | 35.0 | Cluster_17091 |
| 1100.56000 | 35.0 | Cluster_17097 |
| 1101.96000 | 35.0 | Cluster_17105 |
| 1102.54000 | 35.0 | Cluster_17111 |
| 1103.12000 | 35.0 | Cluster_42104 |
| 1108.23000 | 35.0 | Cluster_17176 |
| 1108.37000 | 35.0 | Cluster_42148 |
| 1108.88000 | 35.0 | Cluster_17175 |
| 1109.86000 | 35.0 | Cluster_17199 |
| 1113.50000 | 35.0 | Cluster_42184 |
| 1113.53000 | 35.0 | Cluster_17241 |
| 1115.18000 | 35.0 | Cluster_17258 |
| 1115.77000 | 35.0 | Cluster_00660 |
| 1117.22000 | 35.0 | Cluster_17276 |
| 1118.21000 | 35.0 | Cluster_17277 |
| 1118.22000 | 35.0 | Cluster_17280 |
| 1119.87000 | 35.0 | Cluster_42234 |
| 1121.54000 | 35.0 | Cluster_17322 |
| 1123.04000 | 35.0 | Cluster_42270 |
| 1123.97000 | 35.0 | Cluster_42271 |
| 1124.25000 | 35.0 | Cluster_17359 |
| 1125.57000 | 35.0 | Cluster_42283 |
| 1126.07000 | 35.0 | Cluster_42291 |
| 1127.88000 | 35.0 | Cluster_17398 |
| 1128.57000 | 35.0 | Cluster_17409 |
| 1129.21000 | 35.0 | Cluster_17418 |
| 1129.55000 | 35.0 | Cluster_17428 |
| 1129.90000 | 35.0 | Cluster_17433 |
| 1130.23000 | 35.0 | Cluster_17434 |
| 1131.58000 | 35.0 | Cluster_17448 |
| 1133.13000 | 35.0 | Cluster_42337 |
| 1133.20000 | 35.0 | Cluster_17479 |

| | | |
|---|---|---|
| 1133.22000 | 35.0 | Cluster_17484 |
| 1134.90000 | 35.0 | Cluster_17503 |
| 1135.23000 | 35.0 | Cluster_17509 |
| 1135.51000 | 35.0 | Cluster_17513 |
| 1135.54000 | 35.0 | Cluster_17519 |
| 1135.55000 | 35.0 | Cluster_17517 |
| 1136.11000 | 35.0 | Cluster_42363 |
| 1136.88000 | 35.0 | Cluster_17525 |
| 1136.89000 | 35.0 | Cluster_17538 |
| 1138.13000 | 35.0 | Cluster_42377 |
| 1140.22000 | 35.0 | Cluster_17567 |
| 1143.18000 | 35.0 | Cluster_00674 |
| 1143.56000 | 35.0 | Cluster_17607 |
| 1143.60000 | 35.0 | Cluster_42409 |
| 1144.20000 | 35.0 | Cluster_17626 |
| 1144.24000 | 35.0 | Cluster_17620 |
| 1145.21000 | 35.0 | Cluster_17645 |
| 1146.57000 | 35.0 | Cluster_00680 |
| 1149.29000 | 35.0 | Cluster_42461 |
| 1151.05000 | 35.0 | Cluster_42474 |
| 1152.54000 | 35.0 | Cluster_17735 |
| 1152.88000 | 35.0 | Cluster_17750 |
| 1153.03000 | 35.0 | Cluster_05790 |
| 1153.76000 | 35.0 | Cluster_00695 |
| 1154.35000 | 35.0 | Cluster_42485 |
| 1155.34000 | 35.0 | Cluster_42501 |
| 1156.84000 | 35.0 | Cluster_42511 |
| 1158.84000 | 35.0 | Cluster_42519 |
| 1159.61000 | 35.0 | Cluster_42529 |
| 1160.34000 | 35.0 | Cluster_42534 |
| 1162.11000 | 35.0 | Cluster_42551 |
| 1162.64000 | 35.0 | Cluster_42555 |
| 1163.09000 | 35.0 | Cluster_05817 |
| 1166.08000 | 35.0 | Cluster_05827 |
| 1166.14000 | 35.0 | Cluster_42592 |
| 1166.21000 | 35.0 | Cluster_17894 |
| 1167.19000 | 35.0 | Cluster_17908 |
| 1167.57000 | 35.0 | Cluster_17921 |
| 1168.21000 | 35.0 | Cluster_17931 |
| 1168.33000 | 35.0 | Cluster_42608 |
| 1168.58000 | 35.0 | Cluster_05832 |
| 1173.07000 | 35.0 | Cluster_42645 |
| 1180.80000 | 35.0 | Cluster_05888 |
| 1183.23000 | 35.0 | Cluster_18040 |
| 1187.59000 | 35.0 | Cluster_18075 |
| 1190.56000 | 35.0 | Cluster_18091 |
| 1192.81000 | 35.0 | Cluster_42765 |
| 1192.92000 | 35.0 | Cluster_18117 |
| 1195.88000 | 35.0 | Cluster_18133 |
| 1196.63000 | 35.0 | Cluster_42787 |
| 1197.59000 | 35.0 | Cluster_18145 |

| | | |
|---|---|---|
| 1197.90000 | 35.0 | Cluster_18140 |
| 1198.64000 | 35.0 | Cluster_42803 |
| 1199.32000 | 35.0 | Cluster_05982 |
| 1199.57000 | 35.0 | Cluster_18161 |
| 1200.26000 | 35.0 | Cluster_18168 |
| 1200.56000 | 35.0 | Cluster_18177 |
| 1201.89000 | 35.0 | Cluster_18192 |
| 1203.44000 | 35.0 | Cluster_42830 |
| 1206.62000 | 35.0 | Cluster_42853 |
| 1208.56000 | 35.0 | Cluster_18237 |
| 1209.51000 | 35.0 | Cluster_42887 |
| 1213.12000 | 35.0 | Cluster_42910 |
| 1215.15000 | 35.0 | Cluster_42924 |
| 1218.11000 | 35.0 | Cluster_42950 |
| 1219.09000 | 35.0 | Cluster_42956 |
| 1221.25000 | 35.0 | Cluster_18299 |
| 1222.10000 | 35.0 | Cluster_42968 |
| 1224.08000 | 35.0 | Cluster_42977 |
| 1224.60000 | 35.0 | Cluster_42983 |
| 1225.59000 | 35.0 | Cluster_42994 |
| 1228.07000 | 35.0 | Cluster_43008 |
| 1228.11000 | 35.0 | Cluster_43007 |
| 1230.63000 | 35.0 | Cluster_43026 |
| 1232.57000 | 35.0 | Cluster_43045 |
| 1234.58000 | 35.0 | Cluster_43065 |
| 1238.57000 | 35.0 | Cluster_43079 |
| 1245.28000 | 35.0 | Cluster_18390 |
| 1249.16000 | 35.0 | Cluster_43135 |
| 1251.87000 | 35.0 | Cluster_43149 |
| 1253.87000 | 35.0 | Cluster_43162 |
| 1255.86000 | 35.0 | Cluster_43176 |
| 1257.08000 | 35.0 | Cluster_06282 |
| 1264.27000 | 35.0 | Cluster_18463 |
| 1267.20000 | 35.0 | Cluster_43224 |
| 1267.68000 | 35.0 | Cluster_43228 |
| 1274.29000 | 35.0 | Cluster_18502 |
| 1284.15000 | 35.0 | Cluster_43287 |
| 1300.69000 | 35.0 | Cluster_43366 |
| 1300.70000 | 35.0 | Cluster_43360 |
| 1300.73000 | 35.0 | Cluster_43361 |
| 1300.89000 | 35.0 | Cluster_43362 |
| 1301.68000 | 35.0 | Cluster_43365 |
| 1320.21000 | 35.0 | Cluster_43449 |
| 1329.73000 | 35.0 | Cluster_43484 |
| 1330.45000 | 35.0 | Cluster_06499 |
| 1331.69000 | 35.0 | Cluster_43492 |
| 1334.68000 | 35.0 | Cluster_43505 |
| 1349.89000 | 35.0 | Cluster_43562 |
| 1353.71000 | 35.0 | Cluster_43578 |
| 1354.21000 | 35.0 | Cluster_43579 |
| 1354.66000 | 35.0 | Cluster_06571 |

| | | |
|---|---|---|
| 1361.03000 | 35.0 | Cluster_18681 |
| 1367.16000 | 35.0 | Cluster_43645 |
| 1372.75000 | 35.0 | Cluster_43667 |
| 1376.18000 | 35.0 | Cluster_43680 |
| 1376.34000 | 35.0 | Cluster_18707 |
| 1383.67000 | 35.0 | Cluster_18716 |
| 1394.65000 | 35.0 | Cluster_43739 |
| 1396.44000 | 35.0 | Cluster_18732 |
| 1402.12000 | 35.0 | Cluster_18741 |
| 1406.24000 | 35.0 | Cluster_43793 |
| 1407.03000 | 35.0 | Cluster_18752 |
| 1414.35000 | 35.0 | Cluster_18768 |
| 1417.37000 | 35.0 | Cluster_18772 |
| 1420.01000 | 35.0 | Cluster_18783 |
| 1422.96000 | 35.0 | Cluster_43843 |
| 1423.47000 | 35.0 | Cluster_43844 |
| 1424.69000 | 35.0 | Cluster_18789 |
| 1424.79000 | 35.0 | Cluster_43851 |
| 1425.27000 | 35.0 | Cluster_43853 |
| 1427.03000 | 35.0 | Cluster_18794 |
| 1428.82000 | 35.0 | Cluster_43862 |
| 1429.02000 | 35.0 | Cluster_18795 |
| 1430.01000 | 35.0 | Cluster_18796 |
| 1431.68000 | 35.0 | Cluster_18799 |
| 1437.03000 | 35.0 | Cluster_18803 |
| 1439.35000 | 35.0 | Cluster_18807 |
| 1459.27000 | 35.0 | Cluster_43959 |
| 1477.30000 | 35.0 | Cluster_44015 |
| 1489.22000 | 35.0 | Cluster_44043 |
| 1492.94000 | 35.0 | Cluster_44054 |
| 1495.14000 | 35.0 | Cluster_44065 |
| 1511.80000 | 35.0 | Cluster_44102 |
| 1518.72000 | 35.0 | Cluster_44112 |
| 1530.33000 | 35.0 | Cluster_44125 |
| 1560.76000 | 35.0 | Cluster_44175 |
| 1564.32000 | 35.0 | Cluster_44184 |
| 1571.76000 | 35.0 | Cluster_44194 |
| 1582.85000 | 35.0 | Cluster_44218 |
| 1616.85000 | 35.0 | Cluster_44250 |
| 1633.97000 | 35.0 | Cluster_44257 |
| 1635.40000 | 35.0 | Cluster_44260 |
| 1676.82000 | 35.0 | Cluster_44320 |
| 1683.84000 | 35.0 | Cluster_44341 |
| 1687.92000 | 35.0 | Cluster_44353 |
| 1739.94000 | 35.0 | Cluster_44403 |
| 1792.99000 | 35.0 | Cluster_44425 |

Reject Mass List: (none)
Neutral Loss Mass List: (none)
Product Mass List: (none)
Neutral loss in top: 3
Product in top: 3

Most intense if no parent masses found enabled
Add/subtract mass not enabled
FT master scan preview mode enabled
Charge state screening enabled
Charge state dependent ETD time not enabled
Monoisotopic precursor selection enabled
Charge state rejection enabled
Unassigned charge states : rejected
Charge state 1 : rejected
Charge state 2 : not rejected
Charge state 3 : not rejected
Charge states 4+ : not rejected
Chromatography mode is disabled

Global Data Dependent Settings:
Predict ion injection time enabled
Use global parent and reject mass lists not enabled
Exclude parent mass from data dependent selection not enabled
Exclusion mass width by mass
Exclusion mass width low: 0.50000
Exclusion mass width high: 0.50000
Parent mass width by mass
Parent mass width low: 0.50000
Parent mass width high: 0.50000
Reject mass width by mass
Reject mass width low: 0.50000
Reject mass width high: 0.50000
Zoom/UltraZoom scan mass width by mass
Zoom/UltraZoom scan mass width low: 5.00
Zoom/UltraZoom scan mass width high: 5.00
FT SIM scan mass width low: 5.00
FT SIM scan mass width high: 5.00
Neutral Loss candidates processed by decreasing intensity
Neutral Loss mass width by mass
Neutral Loss mass width low: 0.50000
Neutral Loss mass width high: 0.50000
Product candidates processed by decreasing intensity
Product mass width by mass
Product mass width low: 0.50000
Product mass width high: 0.50000
MS mass range: 0.00-1000000.00
MSn mass range by mass
MSn mass range: 0.00-1000000.00
Use m/z values as masses not enabled
Analog UV data dep. not enabled
Dynamic exclusion not enabled
Isotopic data dependence not enabled
Mass Tags data dependence not enabled
Custom Data Dependent Settings:
Not enabled

Pass 11 (wheat-mixed-digests_MS2_TO-USE-inclusion-01_4.raw):

Creator: Orbi_30393
Last modified: 10/19/2021 by Orbi_30393
MS Run Time (min): 43.00
Sequence override of method parameters not enabled.
Divert Valve: not used during run
Contact Closure: not used during run
Syringe Pump: not used during run
MS Detector Settings:
Real-time modifications to method not enabled
Stepped collision energy not enabled
Additional Microscans:

| | | |
|---|---|---|
| MS2 | 0 | 0 |
| MS3 | 0 | 0 |
| MS4 | 0 | 0 |
| MS5 | 0 | 0 |
| MS6 | 0 | 0 |
| MS7 | 0 | 0 |
| MS8 | 0 | 0 |
| MS9 | 0 | 0 |
| MS10 | 0 | 0 |

Segment 1 Information

Duration (min): 43.00
Number of Scan Events: 2
Tune Method: Orbitrap-tune-file_2020-03-13_HESI

Scan Event Details:
1: FTMS + p norm res=15000 o(300.0-2000.0)
CV = 0.0V
2: ITMS + c norm Dep MS/MS Most intense ion from (1)
Activation Type: CID
Min. Signal Required: 500.0
Isolation Width: 0.20
Normalized Coll. Energy: 35.0
Default Charge State: 2
Activation Q: 0.250
Activation Time: 10.000
CV = 0.0V

Lock Masses:
Pos List Name: N/A
Source: API Source
Mass List: (none)
Neg List Name: N/A
Source: API Source
Mass List: (none)

Data Dependent Settings:

Use separate polarity settings disabled
Parent Mass List:

| MS Mass | MS FAIMS CV | MS Normalized Collision Energy | MS2 Mass | MS2 Normalized Collision Energy | Name |
|---|---|---|---|---|---|
| 300.13000 | | 35.0 | | | Cluster_06844 |
| 300.14000 | | 35.0 | | | Cluster_18928 |
| 300.71000 | | 35.0 | | | Cluster_18955 |
| 301.50000 | | 35.0 | | | Cluster_06851 |
| 301.65000 | | 35.0 | | | Cluster_18984 |
| 302.64000 | | 35.0 | | | Cluster_19016 |
| 302.65000 | | 35.0 | | | Cluster_19017 |
| 303.99000 | | 35.0 | | | Cluster_19040 |
| 304.11000 | | 35.0 | | | Cluster_19042 |
| 304.14000 | | 35.0 | | | Cluster_19049 |
| 304.16000 | | 35.0 | | | Cluster_19050 |
| 304.20000 | | 35.0 | | | Cluster_19052 |
| 304.49000 | | 35.0 | | | Cluster_06863 |
| 304.64000 | | 35.0 | | | Cluster_19062 |
| 306.13000 | | 35.0 | | | Cluster_06873 |
| 306.67000 | | 35.0 | | | Cluster_19109 |
| 307.12000 | | 35.0 | | | Cluster_19119 |
| 307.13000 | | 35.0 | | | Cluster_06880 |
| 308.65000 | | 35.0 | | | Cluster_19177 |
| 308.66000 | | 35.0 | | | Cluster_19179 |
| 309.53000 | | 35.0 | | | Cluster_06901 |
| 310.58000 | | 35.0 | | | Cluster_19217 |
| 310.65000 | | 35.0 | | | Cluster_19222 |
| 310.66000 | | 35.0 | | | Cluster_19221 |
| 311.17000 | | 35.0 | | | Cluster_19246 |
| 311.68000 | | 35.0 | | | Cluster_19257 |
| 313.65000 | | 35.0 | | | Cluster_19315 |
| 314.82000 | | 35.0 | | | Cluster_06941 |
| 315.65000 | | 35.0 | | | Cluster_19384 |
| 316.69000 | | 35.0 | | | Cluster_19442 |
| 317.06000 | | 35.0 | | | Cluster_19459 |
| 317.16000 | | 35.0 | | | Cluster_19466 |
| 317.64000 | | 35.0 | | | Cluster_19490 |
| 318.64000 | | 35.0 | | | Cluster_19532 |
| 318.85000 | | 35.0 | | | Cluster_06973 |
| 319.17000 | | 35.0 | | | Cluster_19552 |
| 319.67000 | | 35.0 | | | Cluster_19560 |
| 319.94000 | | 35.0 | | | Cluster_00850 |
| 320.49000 | | 35.0 | | | Cluster_06993 |
| 320.80000 | | 35.0 | | | Cluster_06994 |
| 321.15000 | | 35.0 | | | Cluster_06995 |
| 321.17000 | | 35.0 | | | Cluster_00869 |
| 321.52000 | | 35.0 | | | Cluster_19594 |
| 322.15000 | | 35.0 | | | Cluster_19619 |
| 322.17000 | | 35.0 | | | Cluster_19617 |

| | | |
|---|---|---|
| 322.50000 | 35.0 | Cluster_06998 |
| 323.65000 | 35.0 | Cluster_19672 |
| 323.69000 | 35.0 | Cluster_19677 |
| 323.83000 | 35.0 | Cluster_19683 |
| 324.42000 | 35.0 | Cluster_00878 |
| 324.60000 | 35.0 | Cluster_19711 |
| 324.65000 | 35.0 | Cluster_19713 |
| 324.83000 | 35.0 | Cluster_07017 |
| 325.14000 | 35.0 | Cluster_19730 |
| 325.82000 | 35.0 | Cluster_07023 |
| 325.85000 | 35.0 | Cluster_07028 |
| 326.17000 | 35.0 | Cluster_19753 |
| 326.40000 | 35.0 | Cluster_00887 |
| 326.90000 | 35.0 | Cluster_00888 |
| 327.21000 | 35.0 | Cluster_19791 |
| 327.96000 | 35.0 | Cluster_07043 |
| 328.48000 | 35.0 | Cluster_07051 |
| 329.18000 | 35.0 | Cluster_19855 |
| 329.82000 | 35.0 | Cluster_07063 |
| 329.93000 | 35.0 | Cluster_19873 |
| 330.18000 | 35.0 | Cluster_19886 |
| 330.90000 | 35.0 | Cluster_19916 |
| 331.17000 | 35.0 | Cluster_19926 |
| 331.20000 | 35.0 | Cluster_19928 |
| 331.21000 | 35.0 | Cluster_19930 |
| 331.22000 | 35.0 | Cluster_19925 |
| 331.66000 | 35.0 | Cluster_19936 |
| 332.14000 | 35.0 | Cluster_19948 |
| 333.47000 | 35.0 | Cluster_07081 |
| 333.82000 | 35.0 | Cluster_07087 |
| 333.87000 | 35.0 | Cluster_07090 |
| 335.12000 | 35.0 | Cluster_20034 |
| 335.83000 | 35.0 | Cluster_07104 |
| 336.19000 | 35.0 | Cluster_07109 |
| 336.23000 | 35.0 | Cluster_20093 |
| 336.91000 | 35.0 | Cluster_00920 |
| 337.19000 | 35.0 | Cluster_20132 |
| 337.21000 | 35.0 | Cluster_20126 |
| 337.83000 | 35.0 | Cluster_07126 |
| 338.11000 | 35.0 | Cluster_20150 |
| 338.67000 | 35.0 | Cluster_20172 |
| 339.16000 | 35.0 | Cluster_20192 |
| 339.47000 | 35.0 | Cluster_07139 |
| 339.81000 | 35.0 | Cluster_07150 |
| 339.94000 | 35.0 | Cluster_00929 |
| 340.18000 | 35.0 | Cluster_20223 |
| 340.26000 | 35.0 | Cluster_20229 |
| 341.15000 | 35.0 | Cluster_20257 |
| 341.16000 | 35.0 | Cluster_20261 |
| 342.16000 | 35.0 | Cluster_07167 |
| 343.11000 | 35.0 | Cluster_20316 |

| | | |
|---|---|---|
| 343.18000 | 35.0 | Cluster_20318 |
| 343.66000 | 35.0 | Cluster_20344 |
| 345.19000 | 35.0 | Cluster_20390 |
| 345.67000 | 35.0 | Cluster_20402 |
| 346.15000 | 35.0 | Cluster_07199 |
| 346.67000 | 35.0 | Cluster_20444 |
| 347.20000 | 35.0 | Cluster_20457 |
| 347.71000 | 35.0 | Cluster_20471 |
| 348.44000 | 35.0 | Cluster_20495 |
| 348.87000 | 35.0 | Cluster_07234 |
| 348.92000 | 35.0 | Cluster_00975 |
| 348.93000 | 35.0 | Cluster_00976 |
| 349.20000 | 35.0 | Cluster_20517 |
| 349.21000 | 35.0 | Cluster_20520 |
| 349.64000 | 35.0 | Cluster_20543 |
| 349.73000 | 35.0 | Cluster_20549 |
| 350.42000 | 35.0 | Cluster_00982 |
| 351.18000 | 35.0 | Cluster_20611 |
| 351.19000 | 35.0 | Cluster_20622 |
| 352.66000 | 35.0 | Cluster_20669 |
| 353.17000 | 35.0 | Cluster_01002 |
| 353.21000 | 35.0 | Cluster_20705 |
| 353.66000 | 35.0 | Cluster_01003 |
| 353.80000 | 35.0 | Cluster_07278 |
| 353.87000 | 35.0 | Cluster_20720 |
| 353.91000 | 35.0 | Cluster_01004 |
| 354.16000 | 35.0 | Cluster_20732 |
| 354.20000 | 35.0 | Cluster_20739 |
| 354.40000 | 35.0 | Cluster_01009 |
| 354.42000 | 35.0 | Cluster_01011 |
| 355.14000 | 35.0 | Cluster_20781 |
| 355.16000 | 35.0 | Cluster_20782 |
| 355.22000 | 35.0 | Cluster_20787 |
| 355.36000 | 35.0 | Cluster_20789 |
| 355.70000 | 35.0 | Cluster_20796 |
| 356.24000 | 35.0 | Cluster_20824 |
| 356.55000 | 35.0 | Cluster_07307 |
| 358.42000 | 35.0 | Cluster_20933 |
| 359.13000 | 35.0 | Cluster_07326 |
| 359.17000 | 35.0 | Cluster_00118 |
| 359.37000 | 35.0 | Cluster_20960 |
| 360.16000 | 35.0 | Cluster_20986 |
| 360.42000 | 35.0 | Cluster_01047 |
| 360.83000 | 35.0 | Cluster_21008 |
| 361.18000 | 35.0 | Cluster_21024 |
| 361.42000 | 35.0 | Cluster_01055 |
| 361.66000 | 35.0 | Cluster_21038 |
| 361.67000 | 35.0 | Cluster_21036 |
| 361.69000 | 35.0 | Cluster_21039 |
| 361.84000 | 35.0 | Cluster_07347 |
| 362.10000 | 35.0 | Cluster_21048 |

| | | |
|---|---|---|
| 362.42000 | 35.0 | Cluster_01059 |
| 362.43000 | 35.0 | Cluster_21075 |
| 362.60000 | 35.0 | Cluster_21077 |
| 362.65000 | 35.0 | Cluster_21076 |
| 363.11000 | 35.0 | Cluster_21089 |
| 363.19000 | 35.0 | Cluster_21093 |
| 363.20000 | 35.0 | Cluster_21094 |
| 363.77000 | 35.0 | Cluster_21121 |
| 364.16000 | 35.0 | Cluster_21135 |
| 364.50000 | 35.0 | Cluster_21161 |
| 364.65000 | 35.0 | Cluster_21162 |
| 365.24000 | 35.0 | Cluster_21221 |
| 365.79000 | 35.0 | Cluster_07396 |
| 365.90000 | 35.0 | Cluster_21241 |
| 366.27000 | 35.0 | Cluster_21267 |
| 366.74000 | 35.0 | Cluster_21285 |
| 367.22000 | 35.0 | Cluster_21306 |
| 367.50000 | 35.0 | Cluster_07416 |
| 367.53000 | 35.0 | Cluster_21324 |
| 368.16000 | 35.0 | Cluster_21368 |
| 368.20000 | 35.0 | Cluster_21366 |
| 368.47000 | 35.0 | Cluster_07428 |
| 369.67000 | 35.0 | Cluster_21437 |
| 370.08000 | 35.0 | Cluster_21453 |
| 370.50000 | 35.0 | Cluster_07445 |
| 370.74000 | 35.0 | Cluster_21476 |
| 370.93000 | 35.0 | Cluster_21486 |
| 371.18000 | 35.0 | Cluster_07449 |
| 371.24000 | 35.0 | Cluster_21501 |
| 371.71000 | 35.0 | Cluster_21517 |
| 371.89000 | 35.0 | Cluster_21531 |
| 372.17000 | 35.0 | Cluster_21537 |
| 372.23000 | 35.0 | Cluster_07464 |
| 372.67000 | 35.0 | Cluster_21575 |
| 372.69000 | 35.0 | Cluster_21572 |
| 373.12000 | 35.0 | Cluster_21589 |
| 373.14000 | 35.0 | Cluster_21585 |
| 373.16000 | 35.0 | Cluster_07470 |
| 373.19000 | 35.0 | Cluster_07471 |
| 373.22000 | 35.0 | Cluster_21597 |
| 373.90000 | 35.0 | Cluster_21628 |
| 374.19000 | 35.0 | Cluster_07480 |
| 374.40000 | 35.0 | Cluster_21650 |
| 374.68000 | 35.0 | Cluster_21656 |
| 375.15000 | 35.0 | Cluster_21668 |
| 375.89000 | 35.0 | Cluster_07496 |
| 377.48000 | 35.0 | Cluster_07512 |
| 377.77000 | 35.0 | Cluster_21780 |
| 378.17000 | 35.0 | Cluster_01122 |
| 378.18000 | 35.0 | Cluster_21788 |
| 378.20000 | 35.0 | Cluster_07523 |

| | | |
|---|---|---|
| 378.67000 | 35.0 | Cluster_01125 |
| 379.26000 | 35.0 | Cluster_21841 |
| 380.45000 | 35.0 | Cluster_21886 |
| 380.68000 | 35.0 | Cluster_21891 |
| 381.22000 | 35.0 | Cluster_07550 |
| 381.70000 | 35.0 | Cluster_01141 |
| 382.18000 | 35.0 | Cluster_21953 |
| 382.66000 | 35.0 | Cluster_01144 |
| 383.63000 | 35.0 | Cluster_22024 |
| 383.68000 | 35.0 | Cluster_22025 |
| 384.16000 | 35.0 | Cluster_01146 |
| 384.21000 | 35.0 | Cluster_22055 |
| 384.22000 | 35.0 | Cluster_22065 |
| 385.52000 | 35.0 | Cluster_07597 |
| 385.54000 | 35.0 | Cluster_07596 |
| 385.86000 | 35.0 | Cluster_22123 |
| 386.66000 | 35.0 | Cluster_22155 |
| 386.68000 | 35.0 | Cluster_22158 |
| 386.90000 | 35.0 | Cluster_22172 |
| 387.11000 | 35.0 | Cluster_22177 |
| 387.63000 | 35.0 | Cluster_22217 |
| 387.93000 | 35.0 | Cluster_22226 |
| 388.23000 | 35.0 | Cluster_22233 |
| 388.50000 | 35.0 | Cluster_07621 |
| 388.64000 | 35.0 | Cluster_22244 |
| 388.67000 | 35.0 | Cluster_22245 |
| 388.83000 | 35.0 | Cluster_07623 |
| 389.25000 | 35.0 | Cluster_22269 |
| 389.51000 | 35.0 | Cluster_07627 |
| 389.91000 | 35.0 | Cluster_22284 |
| 390.71000 | 35.0 | Cluster_22316 |
| 390.87000 | 35.0 | Cluster_07651 |
| 390.88000 | 35.0 | Cluster_07646 |
| 391.50000 | 35.0 | Cluster_07654 |
| 391.65000 | 35.0 | Cluster_22334 |
| 391.75000 | 35.0 | Cluster_22344 |
| 391.82000 | 35.0 | Cluster_07658 |
| 392.22000 | 35.0 | Cluster_07665 |
| 392.25000 | 35.0 | Cluster_22358 |
| 392.73000 | 35.0 | Cluster_22381 |
| 392.77000 | 35.0 | Cluster_22377 |
| 393.19000 | 35.0 | Cluster_01195 |
| 393.53000 | 35.0 | Cluster_07674 |
| 393.69000 | 35.0 | Cluster_01197 |
| 393.71000 | 35.0 | Cluster_22428 |
| 393.98000 | 35.0 | Cluster_22479 |
| 394.21000 | 35.0 | Cluster_22490 |
| 394.71000 | 35.0 | Cluster_22522 |
| 394.72000 | 35.0 | Cluster_22521 |
| 395.89000 | 35.0 | Cluster_07703 |
| 396.23000 | 35.0 | Cluster_22589 |

| | | |
|---|---|---|
| 396.30000 | 35.0 | Cluster_22595 |
| 397.39000 | 35.0 | Cluster_22628 |
| 397.67000 | 35.0 | Cluster_22634 |
| 397.72000 | 35.0 | Cluster_22641 |
| 397.90000 | 35.0 | Cluster_07736 |
| 398.18000 | 35.0 | Cluster_01226 |
| 398.44000 | 35.0 | Cluster_01230 |
| 400.24000 | 35.0 | Cluster_22747 |
| 400.68000 | 35.0 | Cluster_22780 |
| 400.90000 | 35.0 | Cluster_07771 |
| 401.70000 | 35.0 | Cluster_22833 |
| 402.32000 | 35.0 | Cluster_22866 |
| 402.48000 | 35.0 | Cluster_22868 |
| 402.72000 | 35.0 | Cluster_22878 |
| 403.44000 | 35.0 | Cluster_01260 |
| 403.54000 | 35.0 | Cluster_07797 |
| 404.47000 | 35.0 | Cluster_01272 |
| 405.88000 | 35.0 | Cluster_07831 |
| 406.22000 | 35.0 | Cluster_23023 |
| 406.68000 | 35.0 | Cluster_01285 |
| 406.73000 | 35.0 | Cluster_23046 |
| 406.95000 | 35.0 | Cluster_01293 |
| 407.17000 | 35.0 | Cluster_23059 |
| 407.25000 | 35.0 | Cluster_23066 |
| 407.67000 | 35.0 | Cluster_01302 |
| 407.75000 | 35.0 | Cluster_23083 |
| 407.93000 | 35.0 | Cluster_01303 |
| 408.22000 | 35.0 | Cluster_23104 |
| 408.54000 | 35.0 | Cluster_07863 |
| 408.66000 | 35.0 | Cluster_01305 |
| 408.91000 | 35.0 | Cluster_07864 |
| 408.92000 | 35.0 | Cluster_01307 |
| 409.19000 | 35.0 | Cluster_07867 |
| 409.25000 | 35.0 | Cluster_23160 |
| 409.69000 | 35.0 | Cluster_23170 |
| 410.53000 | 35.0 | Cluster_07882 |
| 411.72000 | 35.0 | Cluster_23258 |
| 412.67000 | 35.0 | Cluster_23299 |
| 412.68000 | 35.0 | Cluster_01326 |
| 413.23000 | 35.0 | Cluster_23320 |
| 413.87000 | 35.0 | Cluster_07913 |
| 413.92000 | 35.0 | Cluster_07916 |
| 413.94000 | 35.0 | Cluster_23350 |
| 413.98000 | 35.0 | Cluster_23354 |
| 414.71000 | 35.0 | Cluster_23379 |
| 414.85000 | 35.0 | Cluster_23397 |
| 414.87000 | 35.0 | Cluster_07924 |
| 415.22000 | 35.0 | Cluster_23422 |
| 415.73000 | 35.0 | Cluster_23434 |
| 416.00000 | 35.0 | Cluster_23448 |
| 416.22000 | 35.0 | Cluster_23463 |

| | | |
|---|---|---|
| 416.76000 | 35.0 | Cluster_23476 |
| 416.88000 | 35.0 | Cluster_23482 |
| 416.94000 | 35.0 | Cluster_01353 |
| 418.25000 | 35.0 | Cluster_23534 |
| 418.47000 | 35.0 | Cluster_23549 |
| 419.18000 | 35.0 | Cluster_01360 |
| 419.21000 | 35.0 | Cluster_07978 |
| 419.88000 | 35.0 | Cluster_23600 |
| 420.19000 | 35.0 | Cluster_23617 |
| 420.23000 | 35.0 | Cluster_23612 |
| 420.71000 | 35.0 | Cluster_23632 |
| 421.56000 | 35.0 | Cluster_08005 |
| 422.12000 | 35.0 | Cluster_08008 |
| 422.55000 | 35.0 | Cluster_08014 |
| 422.58000 | 35.0 | Cluster_08015 |
| 423.14000 | 35.0 | Cluster_23727 |
| 423.21000 | 35.0 | Cluster_08022 |
| 423.24000 | 35.0 | Cluster_23740 |
| 423.44000 | 35.0 | Cluster_01387 |
| 423.71000 | 35.0 | Cluster_23753 |
| 423.74000 | 35.0 | Cluster_23766 |
| 424.18000 | 35.0 | Cluster_23787 |
| 424.51000 | 35.0 | Cluster_23811 |
| 424.68000 | 35.0 | Cluster_23813 |
| 425.18000 | 35.0 | Cluster_23850 |
| 425.46000 | 35.0 | Cluster_23865 |
| 425.87000 | 35.0 | Cluster_08056 |
| 425.88000 | 35.0 | Cluster_23895 |
| 426.14000 | 35.0 | Cluster_23901 |
| 426.24000 | 35.0 | Cluster_23924 |
| 426.44000 | 35.0 | Cluster_23942 |
| 426.88000 | 35.0 | Cluster_08077 |
| 427.51000 | 35.0 | Cluster_08094 |
| 428.56000 | 35.0 | Cluster_08107 |
| 428.72000 | 35.0 | Cluster_24045 |
| 428.79000 | 35.0 | Cluster_24059 |
| 429.24000 | 35.0 | Cluster_24074 |
| 429.57000 | 35.0 | Cluster_08120 |
| 429.72000 | 35.0 | Cluster_24100 |
| 429.76000 | 35.0 | Cluster_24093 |
| 429.85000 | 35.0 | Cluster_08124 |
| 429.95000 | 35.0 | Cluster_01432 |
| 430.73000 | 35.0 | Cluster_24136 |
| 431.21000 | 35.0 | Cluster_24172 |
| 431.41000 | 35.0 | Cluster_24179 |
| 431.74000 | 35.0 | Cluster_24194 |
| 432.16000 | 35.0 | Cluster_24207 |
| 432.21000 | 35.0 | Cluster_24215 |
| 432.87000 | 35.0 | Cluster_08164 |
| 433.96000 | 35.0 | Cluster_24289 |
| 434.79000 | 35.0 | Cluster_24347 |

| | | |
|---|---|---|
| 434.97000 | 35.0 | Cluster_24350 |
| 435.17000 | 35.0 | Cluster_24357 |
| 435.24000 | 35.0 | Cluster_08201 |
| 435.37000 | 35.0 | Cluster_24383 |
| 435.70000 | 35.0 | Cluster_01481 |
| 435.97000 | 35.0 | Cluster_01487 |
| 436.48000 | 35.0 | Cluster_01491 |
| 437.37000 | 35.0 | Cluster_24491 |
| 437.46000 | 35.0 | Cluster_01497 |
| 438.22000 | 35.0 | Cluster_24532 |
| 438.30000 | 35.0 | Cluster_24543 |
| 438.96000 | 35.0 | Cluster_01503 |
| 438.97000 | 35.0 | Cluster_01504 |
| 439.82000 | 35.0 | Cluster_24632 |
| 440.21000 | 35.0 | Cluster_01512 |
| 440.56000 | 35.0 | Cluster_24655 |
| 440.74000 | 35.0 | Cluster_24659 |
| 440.86000 | 35.0 | Cluster_08277 |
| 441.21000 | 35.0 | Cluster_24682 |
| 441.24000 | 35.0 | Cluster_01516 |
| 441.73000 | 35.0 | Cluster_24703 |
| 441.87000 | 35.0 | Cluster_08293 |
| 442.00000 | 35.0 | Cluster_24716 |
| 442.28000 | 35.0 | Cluster_24730 |
| 442.57000 | 35.0 | Cluster_08303 |
| 442.77000 | 35.0 | Cluster_24758 |
| 442.89000 | 35.0 | Cluster_24763 |
| 442.98000 | 35.0 | Cluster_24761 |
| 443.12000 | 35.0 | Cluster_24771 |
| 443.22000 | 35.0 | Cluster_24782 |
| 443.28000 | 35.0 | Cluster_24785 |
| 443.29000 | 35.0 | Cluster_24795 |
| 443.89000 | 35.0 | Cluster_08322 |
| 443.92000 | 35.0 | Cluster_08324 |
| 444.72000 | 35.0 | Cluster_24866 |
| 445.18000 | 35.0 | Cluster_08343 |
| 445.55000 | 35.0 | Cluster_24900 |
| 446.27000 | 35.0 | Cluster_24941 |
| 446.73000 | 35.0 | Cluster_24959 |
| 446.91000 | 35.0 | Cluster_08369 |
| 447.17000 | 35.0 | Cluster_24974 |
| 447.25000 | 35.0 | Cluster_08376 |
| 447.28000 | 35.0 | Cluster_24982 |
| 447.29000 | 35.0 | Cluster_24988 |
| 447.60000 | 35.0 | Cluster_08380 |
| 448.30000 | 35.0 | Cluster_25022 |
| 448.79000 | 35.0 | Cluster_25051 |
| 448.97000 | 35.0 | Cluster_01573 |
| 449.25000 | 35.0 | Cluster_08416 |
| 449.74000 | 35.0 | Cluster_25092 |
| 449.84000 | 35.0 | Cluster_25102 |

| | | |
|---|---|---|
| 450.23000 | 35.0 | Cluster_25125 |
| 450.93000 | 35.0 | Cluster_08435 |
| 451.14000 | 35.0 | Cluster_25153 |
| 451.17000 | 35.0 | Cluster_25168 |
| 451.57000 | 35.0 | Cluster_08446 |
| 452.13000 | 35.0 | Cluster_25195 |
| 452.26000 | 35.0 | Cluster_08462 |
| 452.75000 | 35.0 | Cluster_25235 |
| 453.14000 | 35.0 | Cluster_25244 |
| 453.24000 | 35.0 | Cluster_25248 |
| 454.68000 | 35.0 | Cluster_25320 |
| 454.70000 | 35.0 | Cluster_25319 |
| 454.74000 | 35.0 | Cluster_25325 |
| 455.15000 | 35.0 | Cluster_25343 |
| 455.18000 | 35.0 | Cluster_01617 |
| 455.20000 | 35.0 | Cluster_08518 |
| 456.24000 | 35.0 | Cluster_25393 |
| 456.26000 | 35.0 | Cluster_08536 |
| 456.57000 | 35.0 | Cluster_08539 |
| 456.75000 | 35.0 | Cluster_25404 |
| 456.78000 | 35.0 | Cluster_25413 |
| 457.20000 | 35.0 | Cluster_25422 |
| 457.75000 | 35.0 | Cluster_25448 |
| 457.76000 | 35.0 | Cluster_25454 |
| 458.69000 | 35.0 | Cluster_25493 |
| 458.76000 | 35.0 | Cluster_25500 |
| 459.22000 | 35.0 | Cluster_08566 |
| 459.25000 | 35.0 | Cluster_25514 |
| 459.78000 | 35.0 | Cluster_25540 |
| 459.90000 | 35.0 | Cluster_08580 |
| 460.27000 | 35.0 | Cluster_25572 |
| 460.77000 | 35.0 | Cluster_25581 |
| 460.97000 | 35.0 | Cluster_01663 |
| 461.60000 | 35.0 | Cluster_08592 |
| 461.70000 | 35.0 | Cluster_25618 |
| 461.73000 | 35.0 | Cluster_25620 |
| 462.22000 | 35.0 | Cluster_08608 |
| 462.50000 | 35.0 | Cluster_25643 |
| 463.61000 | 35.0 | Cluster_25677 |
| 463.73000 | 35.0 | Cluster_25692 |
| 464.22000 | 35.0 | Cluster_08632 |
| 464.30000 | 35.0 | Cluster_25714 |
| 464.55000 | 35.0 | Cluster_08640 |
| 465.77000 | 35.0 | Cluster_25807 |
| 465.95000 | 35.0 | Cluster_08659 |
| 465.99000 | 35.0 | Cluster_01692 |
| 466.27000 | 35.0 | Cluster_25822 |
| 466.90000 | 35.0 | Cluster_08673 |
| 467.18000 | 35.0 | Cluster_08677 |
| 467.25000 | 35.0 | Cluster_08678 |
| 467.30000 | 35.0 | Cluster_25858 |

| | | |
|---|---|---|
| 467.67000 | 35.0 | Cluster_00144 |
| 467.72000 | 35.0 | Cluster_25877 |
| 468.89000 | 35.0 | Cluster_08711 |
| 469.26000 | 35.0 | Cluster_08720 |
| 469.74000 | 35.0 | Cluster_25967 |
| 469.88000 | 35.0 | Cluster_08724 |
| 470.43000 | 35.0 | Cluster_26002 |
| 470.56000 | 35.0 | Cluster_00010 |
| 470.92000 | 35.0 | Cluster_08745 |
| 471.00000 | 35.0 | Cluster_01731 |
| 471.22000 | 35.0 | Cluster_08748 |
| 471.25000 | 35.0 | Cluster_01737 |
| 471.26000 | 35.0 | Cluster_01738 |
| 471.48000 | 35.0 | Cluster_26044 |
| 471.74000 | 35.0 | Cluster_26063 |
| 471.84000 | 35.0 | Cluster_26052 |
| 472.23000 | 35.0 | Cluster_26081 |
| 472.24000 | 35.0 | Cluster_08764 |
| 472.72000 | 35.0 | Cluster_26107 |
| 472.76000 | 35.0 | Cluster_26106 |
| 472.77000 | 35.0 | Cluster_26113 |
| 473.27000 | 35.0 | Cluster_26146 |
| 473.77000 | 35.0 | Cluster_26157 |
| 474.54000 | 35.0 | Cluster_08802 |
| 474.58000 | 35.0 | Cluster_08800 |
| 474.85000 | 35.0 | Cluster_08806 |
| 475.58000 | 35.0 | Cluster_08820 |
| 475.78000 | 35.0 | Cluster_26234 |
| 476.21000 | 35.0 | Cluster_26244 |
| 476.23000 | 35.0 | Cluster_08830 |
| 476.28000 | 35.0 | Cluster_08832 |
| 477.07000 | 35.0 | Cluster_26275 |
| 477.20000 | 35.0 | Cluster_08843 |
| 477.27000 | 35.0 | Cluster_08840 |
| 477.55000 | 35.0 | Cluster_08851 |
| 477.61000 | 35.0 | Cluster_08856 |
| 477.93000 | 35.0 | Cluster_08861 |
| 478.22000 | 35.0 | Cluster_26309 |
| 478.25000 | 35.0 | Cluster_26313 |
| 478.76000 | 35.0 | Cluster_26329 |
| 478.77000 | 35.0 | Cluster_26332 |
| 478.93000 | 35.0 | Cluster_08880 |
| 478.99000 | 35.0 | Cluster_01780 |
| 479.57000 | 35.0 | Cluster_08888 |
| 479.71000 | 35.0 | Cluster_26429 |
| 479.76000 | 35.0 | Cluster_26423 |
| 480.22000 | 35.0 | Cluster_08900 |
| 480.70000 | 35.0 | Cluster_26476 |
| 481.22000 | 35.0 | Cluster_26493 |
| 481.27000 | 35.0 | Cluster_26504 |
| 481.28000 | 35.0 | Cluster_08920 |

| | | |
|---|---|---|
| 481.56000 | 35.0 | Cluster_08922 |
| 481.58000 | 35.0 | Cluster_08924 |
| 481.72000 | 35.0 | Cluster_26523 |
| 481.73000 | 35.0 | Cluster_01804 |
| 481.79000 | 35.0 | Cluster_26518 |
| 482.20000 | 35.0 | Cluster_01810 |
| 482.28000 | 35.0 | Cluster_26547 |
| 482.49000 | 35.0 | Cluster_26549 |
| 483.24000 | 35.0 | Cluster_26594 |
| 483.30000 | 35.0 | Cluster_26603 |
| 483.76000 | 35.0 | Cluster_26625 |
| 483.84000 | 35.0 | Cluster_26627 |
| 484.09000 | 35.0 | Cluster_26635 |
| 484.73000 | 35.0 | Cluster_26685 |
| 484.76000 | 35.0 | Cluster_26681 |
| 485.06000 | 35.0 | Cluster_00166 |
| 485.54000 | 35.0 | Cluster_08981 |
| 488.29000 | 35.0 | Cluster_26837 |
| 488.61000 | 35.0 | Cluster_09032 |
| 488.79000 | 35.0 | Cluster_26859 |
| 488.95000 | 35.0 | Cluster_09038 |
| 489.74000 | 35.0 | Cluster_26904 |
| 489.93000 | 35.0 | Cluster_09048 |
| 490.50000 | 35.0 | Cluster_26931 |
| 490.70000 | 35.0 | Cluster_26935 |
| 490.77000 | 35.0 | Cluster_26939 |
| 492.55000 | 35.0 | Cluster_09097 |
| 492.60000 | 35.0 | Cluster_09099 |
| 493.27000 | 35.0 | Cluster_27041 |
| 494.26000 | 35.0 | Cluster_27100 |
| 495.19000 | 35.0 | Cluster_27149 |
| 495.71000 | 35.0 | Cluster_27170 |
| 495.95000 | 35.0 | Cluster_09160 |
| 496.24000 | 35.0 | Cluster_01895 |
| 496.95000 | 35.0 | Cluster_09176 |
| 497.35000 | 35.0 | Cluster_27258 |
| 497.74000 | 35.0 | Cluster_27265 |
| 497.80000 | 35.0 | Cluster_27275 |
| 497.94000 | 35.0 | Cluster_09191 |
| 498.01000 | 35.0 | Cluster_01929 |
| 498.50000 | 35.0 | Cluster_27299 |
| 498.73000 | 35.0 | Cluster_27302 |
| 499.23000 | 35.0 | Cluster_09211 |
| 499.33000 | 35.0 | Cluster_27324 |
| 499.76000 | 35.0 | Cluster_01941 |
| 499.92000 | 35.0 | Cluster_09218 |
| 499.96000 | 35.0 | Cluster_09219 |
| 500.02000 | 35.0 | Cluster_27355 |
| 500.63000 | 35.0 | Cluster_09231 |
| 500.70000 | 35.0 | Cluster_27401 |
| 500.96000 | 35.0 | Cluster_01956 |

| | | |
|---|---|---|
| 501.22000 | 35.0 | Cluster_27436 |
| 501.59000 | 35.0 | Cluster_09244 |
| 501.76000 | 35.0 | Cluster_27459 |
| 502.21000 | 35.0 | Cluster_01970 |
| 502.22000 | 35.0 | Cluster_27473 |
| 502.59000 | 35.0 | Cluster_09268 |
| 502.76000 | 35.0 | Cluster_27485 |
| 503.25000 | 35.0 | Cluster_27512 |
| 503.28000 | 35.0 | Cluster_27518 |
| 504.21000 | 35.0 | Cluster_27547 |
| 504.56000 | 35.0 | Cluster_09288 |
| 504.98000 | 35.0 | Cluster_01984 |
| 505.01000 | 35.0 | Cluster_27567 |
| 505.24000 | 35.0 | Cluster_09298 |
| 505.52000 | 35.0 | Cluster_01990 |
| 505.60000 | 35.0 | Cluster_09305 |
| 505.79000 | 35.0 | Cluster_27593 |
| 506.73000 | 35.0 | Cluster_27611 |
| 506.74000 | 35.0 | Cluster_27621 |
| 506.76000 | 35.0 | Cluster_27614 |
| 507.25000 | 35.0 | Cluster_27638 |
| 507.97000 | 35.0 | Cluster_09350 |
| 508.62000 | 35.0 | Cluster_09366 |
| 508.70000 | 35.0 | Cluster_27686 |
| 508.71000 | 35.0 | Cluster_27692 |
| 509.20000 | 35.0 | Cluster_27715 |
| 509.26000 | 35.0 | Cluster_27718 |
| 509.87000 | 35.0 | Cluster_09377 |
| 509.91000 | 35.0 | Cluster_09383 |
| 511.23000 | 35.0 | Cluster_27820 |
| 511.79000 | 35.0 | Cluster_27852 |
| 511.91000 | 35.0 | Cluster_09420 |
| 512.25000 | 35.0 | Cluster_27867 |
| 512.27000 | 35.0 | Cluster_27865 |
| 513.09000 | 35.0 | Cluster_27898 |
| 513.77000 | 35.0 | Cluster_27929 |
| 514.72000 | 35.0 | Cluster_27964 |
| 514.74000 | 35.0 | Cluster_27971 |
| 515.25000 | 35.0 | Cluster_02053 |
| 515.48000 | 35.0 | Cluster_02061 |
| 515.90000 | 35.0 | Cluster_09478 |
| 516.74000 | 35.0 | Cluster_28062 |
| 517.92000 | 35.0 | Cluster_09503 |
| 518.00000 | 35.0 | Cluster_28115 |
| 518.26000 | 35.0 | Cluster_09508 |
| 518.31000 | 35.0 | Cluster_28130 |
| 518.51000 | 35.0 | Cluster_28138 |
| 518.59000 | 35.0 | Cluster_09521 |
| 518.78000 | 35.0 | Cluster_28152 |
| 519.59000 | 35.0 | Cluster_09531 |
| 519.81000 | 35.0 | Cluster_28200 |

| | | |
|---|---|---|
| 520.04000 | 35.0 | Cluster_28202 |
| 520.51000 | 35.0 | Cluster_28225 |
| 520.80000 | 35.0 | Cluster_28240 |
| 520.81000 | 35.0 | Cluster_28232 |
| 521.02000 | 35.0 | Cluster_02095 |
| 521.26000 | 35.0 | Cluster_09553 |
| 521.66000 | 35.0 | Cluster_28271 |
| 522.22000 | 35.0 | Cluster_28286 |
| 522.45000 | 35.0 | Cluster_02102 |
| 522.50000 | 35.0 | Cluster_28303 |
| 522.62000 | 35.0 | Cluster_09582 |
| 522.75000 | 35.0 | Cluster_28315 |
| 523.22000 | 35.0 | Cluster_28350 |
| 523.27000 | 35.0 | Cluster_28334 |
| 523.35000 | 35.0 | Cluster_28353 |
| 523.92000 | 35.0 | Cluster_09603 |
| 523.97000 | 35.0 | Cluster_28374 |
| 524.28000 | 35.0 | Cluster_28386 |
| 526.18000 | 35.0 | Cluster_28467 |
| 526.20000 | 35.0 | Cluster_09656 |
| 526.93000 | 35.0 | Cluster_09676 |
| 526.95000 | 35.0 | Cluster_09677 |
| 527.76000 | 35.0 | Cluster_28507 |
| 528.23000 | 35.0 | Cluster_28536 |
| 528.27000 | 35.0 | Cluster_00184 |
| 528.32000 | 35.0 | Cluster_28539 |
| 528.36000 | 35.0 | Cluster_28552 |
| 528.97000 | 35.0 | Cluster_09710 |
| 529.25000 | 35.0 | Cluster_09712 |
| 529.79000 | 35.0 | Cluster_28615 |
| 530.20000 | 35.0 | Cluster_28629 |
| 530.26000 | 35.0 | Cluster_02153 |
| 530.51000 | 35.0 | Cluster_28657 |
| 530.81000 | 35.0 | Cluster_28677 |
| 530.94000 | 35.0 | Cluster_09741 |
| 531.75000 | 35.0 | Cluster_28707 |
| 531.77000 | 35.0 | Cluster_28708 |
| 532.24000 | 35.0 | Cluster_09763 |
| 532.27000 | 35.0 | Cluster_09766 |
| 532.62000 | 35.0 | Cluster_09768 |
| 532.97000 | 35.0 | Cluster_09773 |
| 533.15000 | 35.0 | Cluster_28752 |
| 533.16000 | 35.0 | Cluster_28753 |
| 533.19000 | 35.0 | Cluster_09774 |
| 533.32000 | 35.0 | Cluster_09778 |
| 533.70000 | 35.0 | Cluster_28783 |
| 533.82000 | 35.0 | Cluster_28799 |
| 533.91000 | 35.0 | Cluster_09792 |
| 534.28000 | 35.0 | Cluster_28807 |
| 534.59000 | 35.0 | Cluster_09803 |
| 535.32000 | 35.0 | Cluster_09812 |

| | | |
|---|---|---|
| 535.80000 | 35.0 | Cluster_28867 |
| 535.94000 | 35.0 | Cluster_09822 |
| 536.06000 | 35.0 | Cluster_28878 |
| 536.26000 | 35.0 | Cluster_09823 |
| 536.27000 | 35.0 | Cluster_28888 |
| 536.48000 | 35.0 | Cluster_28899 |
| 536.60000 | 35.0 | Cluster_09834 |
| 537.23000 | 35.0 | Cluster_09849 |
| 537.56000 | 35.0 | Cluster_09853 |
| 537.69000 | 35.0 | Cluster_28943 |
| 537.91000 | 35.0 | Cluster_09869 |
| 538.84000 | 35.0 | Cluster_28980 |
| 538.97000 | 35.0 | Cluster_09886 |
| 538.98000 | 35.0 | Cluster_09885 |
| 539.01000 | 35.0 | Cluster_28985 |
| 539.28000 | 35.0 | Cluster_28993 |
| 539.52000 | 35.0 | Cluster_29001 |
| 540.02000 | 35.0 | Cluster_09899 |
| 540.27000 | 35.0 | Cluster_29031 |
| 540.79000 | 35.0 | Cluster_29052 |
| 541.34000 | 35.0 | Cluster_29085 |
| 541.44000 | 35.0 | Cluster_29098 |
| 541.82000 | 35.0 | Cluster_29115 |
| 542.26000 | 35.0 | Cluster_02236 |
| 542.28000 | 35.0 | Cluster_02229 |
| 542.52000 | 35.0 | Cluster_02233 |
| 542.77000 | 35.0 | Cluster_29162 |
| 543.27000 | 35.0 | Cluster_29193 |
| 544.95000 | 35.0 | Cluster_09982 |
| 545.77000 | 35.0 | Cluster_29280 |
| 545.92000 | 35.0 | Cluster_09992 |
| 546.31000 | 35.0 | Cluster_29312 |
| 547.32000 | 35.0 | Cluster_29345 |
| 548.27000 | 35.0 | Cluster_29389 |
| 548.46000 | 35.0 | Cluster_00194 |
| 548.61000 | 35.0 | Cluster_10035 |
| 548.64000 | 35.0 | Cluster_10037 |
| 549.25000 | 35.0 | Cluster_10044 |
| 549.28000 | 35.0 | Cluster_29418 |
| 549.90000 | 35.0 | Cluster_10052 |
| 549.99000 | 35.0 | Cluster_10058 |
| 550.22000 | 35.0 | Cluster_29451 |
| 550.31000 | 35.0 | Cluster_29461 |
| 550.55000 | 35.0 | Cluster_10069 |
| 550.61000 | 35.0 | Cluster_10068 |
| 550.78000 | 35.0 | Cluster_29465 |
| 550.84000 | 35.0 | Cluster_29476 |
| 551.01000 | 35.0 | Cluster_29483 |
| 551.29000 | 35.0 | Cluster_29499 |
| 551.57000 | 35.0 | Cluster_10082 |
| 551.97000 | 35.0 | Cluster_10089 |

| | | |
|---|---|---|
| 552.36000 | 35.0 | Cluster_29545 |
| 552.74000 | 35.0 | Cluster_29553 |
| 552.75000 | 35.0 | Cluster_29548 |
| 552.96000 | 35.0 | Cluster_10104 |
| 553.55000 | 35.0 | Cluster_29579 |
| 553.63000 | 35.0 | Cluster_10116 |
| 554.58000 | 35.0 | Cluster_10130 |
| 554.60000 | 35.0 | Cluster_10136 |
| 554.97000 | 35.0 | Cluster_10144 |
| 556.25000 | 35.0 | Cluster_29692 |
| 556.68000 | 35.0 | Cluster_10177 |
| 556.96000 | 35.0 | Cluster_10184 |
| 557.73000 | 35.0 | Cluster_29750 |
| 558.23000 | 35.0 | Cluster_10209 |
| 558.65000 | 35.0 | Cluster_10215 |
| 558.81000 | 35.0 | Cluster_29792 |
| 559.29000 | 35.0 | Cluster_29811 |
| 560.35000 | 35.0 | Cluster_29845 |
| 560.65000 | 35.0 | Cluster_10245 |
| 560.83000 | 35.0 | Cluster_29862 |
| 561.28000 | 35.0 | Cluster_29879 |
| 561.64000 | 35.0 | Cluster_10262 |
| 561.70000 | 35.0 | Cluster_29892 |
| 561.81000 | 35.0 | Cluster_29901 |
| 561.98000 | 35.0 | Cluster_10266 |
| 562.07000 | 35.0 | Cluster_29906 |
| 562.29000 | 35.0 | Cluster_29915 |
| 563.32000 | 35.0 | Cluster_10288 |
| 563.50000 | 35.0 | Cluster_02355 |
| 563.75000 | 35.0 | Cluster_29975 |
| 563.76000 | 35.0 | Cluster_29965 |
| 564.78000 | 35.0 | Cluster_30003 |
| 564.79000 | 35.0 | Cluster_02366 |
| 565.56000 | 35.0 | Cluster_30039 |
| 565.60000 | 35.0 | Cluster_10318 |
| 566.33000 | 35.0 | Cluster_30080 |
| 566.74000 | 35.0 | Cluster_02376 |
| 566.79000 | 35.0 | Cluster_30088 |
| 566.90000 | 35.0 | Cluster_30092 |
| 566.98000 | 35.0 | Cluster_10333 |
| 567.07000 | 35.0 | Cluster_30105 |
| 567.24000 | 35.0 | Cluster_30110 |
| 567.30000 | 35.0 | Cluster_30123 |
| 567.31000 | 35.0 | Cluster_30114 |
| 568.56000 | 35.0 | Cluster_02384 |
| 568.73000 | 35.0 | Cluster_30166 |
| 568.79000 | 35.0 | Cluster_30177 |
| 568.80000 | 35.0 | Cluster_30176 |
| 568.85000 | 35.0 | Cluster_30178 |
| 569.06000 | 35.0 | Cluster_02386 |
| 569.57000 | 35.0 | Cluster_02390 |

| | | |
|---|---|---|
| 569.82000 | 35.0 | Cluster_30204 |
| 570.30000 | 35.0 | Cluster_02396 |
| 570.36000 | 35.0 | Cluster_30219 |
| 570.73000 | 35.0 | Cluster_02397 |
| 570.77000 | 35.0 | Cluster_30236 |
| 571.22000 | 35.0 | Cluster_30252 |
| 571.34000 | 35.0 | Cluster_30256 |
| 571.64000 | 35.0 | Cluster_10409 |
| 571.77000 | 35.0 | Cluster_30264 |
| 572.13000 | 35.0 | Cluster_00213 |
| 572.30000 | 35.0 | Cluster_10420 |
| 572.79000 | 35.0 | Cluster_30301 |
| 573.35000 | 35.0 | Cluster_30317 |
| 574.26000 | 35.0 | Cluster_10456 |
| 574.80000 | 35.0 | Cluster_30355 |
| 575.25000 | 35.0 | Cluster_02433 |
| 575.33000 | 35.0 | Cluster_02435 |
| 576.55000 | 35.0 | Cluster_02439 |
| 576.77000 | 35.0 | Cluster_30417 |
| 576.98000 | 35.0 | Cluster_10494 |
| 577.30000 | 35.0 | Cluster_10500 |
| 577.81000 | 35.0 | Cluster_30446 |
| 578.32000 | 35.0 | Cluster_30460 |
| 578.55000 | 35.0 | Cluster_02454 |
| 578.86000 | 35.0 | Cluster_30486 |
| 578.98000 | 35.0 | Cluster_10523 |
| 579.33000 | 35.0 | Cluster_30494 |
| 579.55000 | 35.0 | Cluster_02460 |
| 579.61000 | 35.0 | Cluster_10531 |
| 579.63000 | 35.0 | Cluster_10533 |
| 579.77000 | 35.0 | Cluster_30512 |
| 579.81000 | 35.0 | Cluster_30517 |
| 579.83000 | 35.0 | Cluster_30515 |
| 580.36000 | 35.0 | Cluster_30555 |
| 580.86000 | 35.0 | Cluster_02466 |
| 581.28000 | 35.0 | Cluster_02468 |
| 581.31000 | 35.0 | Cluster_30577 |
| 581.96000 | 35.0 | Cluster_10563 |
| 582.31000 | 35.0 | Cluster_30648 |
| 582.54000 | 35.0 | Cluster_02478 |
| 582.65000 | 35.0 | Cluster_10574 |
| 582.70000 | 35.0 | Cluster_00226 |
| 582.74000 | 35.0 | Cluster_30654 |
| 584.05000 | 35.0 | Cluster_30718 |
| 584.31000 | 35.0 | Cluster_30724 |
| 584.54000 | 35.0 | Cluster_30733 |
| 585.02000 | 35.0 | Cluster_30763 |
| 585.29000 | 35.0 | Cluster_10612 |
| 585.35000 | 35.0 | Cluster_02501 |
| 585.80000 | 35.0 | Cluster_30791 |
| 585.84000 | 35.0 | Cluster_30792 |

| | | |
|---|---|---|
| 586.62000 | 35.0 | Cluster_10630 |
| 586.75000 | 35.0 | Cluster_30815 |
| 587.30000 | 35.0 | Cluster_10642 |
| 587.33000 | 35.0 | Cluster_30848 |
| 587.34000 | 35.0 | Cluster_30837 |
| 587.65000 | 35.0 | Cluster_10646 |
| 588.69000 | 35.0 | Cluster_30892 |
| 588.82000 | 35.0 | Cluster_30900 |
| 589.55000 | 35.0 | Cluster_30932 |
| 589.74000 | 35.0 | Cluster_30933 |
| 589.96000 | 35.0 | Cluster_10677 |
| 590.22000 | 35.0 | Cluster_30952 |
| 590.30000 | 35.0 | Cluster_30953 |
| 591.04000 | 35.0 | Cluster_30998 |
| 591.44000 | 35.0 | Cluster_31008 |
| 591.52000 | 35.0 | Cluster_02538 |
| 591.66000 | 35.0 | Cluster_10703 |
| 591.81000 | 35.0 | Cluster_31027 |
| 592.01000 | 35.0 | Cluster_10706 |
| 592.79000 | 35.0 | Cluster_31056 |
| 592.82000 | 35.0 | Cluster_31060 |
| 592.91000 | 35.0 | Cluster_31063 |
| 593.06000 | 35.0 | Cluster_02547 |
| 593.07000 | 35.0 | Cluster_02548 |
| 593.33000 | 35.0 | Cluster_10724 |
| 593.34000 | 35.0 | Cluster_31073 |
| 593.93000 | 35.0 | Cluster_10743 |
| 594.32000 | 35.0 | Cluster_10753 |
| 595.59000 | 35.0 | Cluster_31146 |
| 595.62000 | 35.0 | Cluster_10769 |
| 596.75000 | 35.0 | Cluster_31177 |
| 597.67000 | 35.0 | Cluster_10808 |
| 597.81000 | 35.0 | Cluster_31212 |
| 597.82000 | 35.0 | Cluster_31208 |
| 598.29000 | 35.0 | Cluster_10814 |
| 598.36000 | 35.0 | Cluster_10818 |
| 599.36000 | 35.0 | Cluster_31267 |
| 599.66000 | 35.0 | Cluster_10840 |
| 599.95000 | 35.0 | Cluster_10848 |
| 600.81000 | 35.0 | Cluster_31316 |
| 600.97000 | 35.0 | Cluster_10858 |
| 601.36000 | 35.0 | Cluster_31339 |
| 602.32000 | 35.0 | Cluster_31390 |
| 602.83000 | 35.0 | Cluster_31406 |
| 603.30000 | 35.0 | Cluster_31409 |
| 603.51000 | 35.0 | Cluster_02598 |
| 603.53000 | 35.0 | Cluster_31419 |
| 603.96000 | 35.0 | Cluster_10888 |
| 604.80000 | 35.0 | Cluster_31463 |
| 605.31000 | 35.0 | Cluster_31473 |
| 605.67000 | 35.0 | Cluster_10910 |

| | | |
|---|---|---|
| 606.07000 | 35.0 | Cluster_31501 |
| 606.27000 | 35.0 | Cluster_31514 |
| 606.87000 | 35.0 | Cluster_31531 |
| 606.89000 | 35.0 | Cluster_10941 |
| 607.32000 | 35.0 | Cluster_31536 |
| 608.09000 | 35.0 | Cluster_02626 |
| 608.32000 | 35.0 | Cluster_31587 |
| 608.49000 | 35.0 | Cluster_31590 |
| 608.78000 | 35.0 | Cluster_31595 |
| 608.79000 | 35.0 | Cluster_31601 |
| 609.06000 | 35.0 | Cluster_31605 |
| 609.81000 | 35.0 | Cluster_31625 |
| 610.31000 | 35.0 | Cluster_31655 |
| 611.34000 | 35.0 | Cluster_31685 |
| 612.32000 | 35.0 | Cluster_11010 |
| 613.83000 | 35.0 | Cluster_31794 |
| 614.06000 | 35.0 | Cluster_31796 |
| 614.07000 | 35.0 | Cluster_02670 |
| 614.19000 | 35.0 | Cluster_31797 |
| 614.26000 | 35.0 | Cluster_11035 |
| 615.83000 | 35.0 | Cluster_31852 |
| 616.08000 | 35.0 | Cluster_02680 |
| 616.32000 | 35.0 | Cluster_11075 |
| 616.55000 | 35.0 | Cluster_31873 |
| 616.83000 | 35.0 | Cluster_31882 |
| 617.02000 | 35.0 | Cluster_31892 |
| 617.35000 | 35.0 | Cluster_31911 |
| 617.93000 | 35.0 | Cluster_31935 |
| 618.05000 | 35.0 | Cluster_02693 |
| 618.79000 | 35.0 | Cluster_31952 |
| 619.29000 | 35.0 | Cluster_02701 |
| 619.33000 | 35.0 | Cluster_31985 |
| 619.62000 | 35.0 | Cluster_11116 |
| 620.53000 | 35.0 | Cluster_32028 |
| 620.60000 | 35.0 | Cluster_11128 |
| 621.30000 | 35.0 | Cluster_32049 |
| 621.36000 | 35.0 | Cluster_32053 |
| 621.39000 | 35.0 | Cluster_32052 |
| 621.79000 | 35.0 | Cluster_32057 |
| 621.98000 | 35.0 | Cluster_11154 |
| 622.78000 | 35.0 | Cluster_32095 |
| 622.98000 | 35.0 | Cluster_11175 |
| 623.28000 | 35.0 | Cluster_32112 |
| 623.36000 | 35.0 | Cluster_32126 |
| 623.65000 | 35.0 | Cluster_11191 |
| 623.98000 | 35.0 | Cluster_11189 |
| 624.20000 | 35.0 | Cluster_32153 |
| 624.28000 | 35.0 | Cluster_32154 |
| 624.63000 | 35.0 | Cluster_11202 |
| 625.04000 | 35.0 | Cluster_02732 |
| 625.32000 | 35.0 | Cluster_11203 |

| | | |
|---|---|---|
| 625.58000 | 35.0 | Cluster_02735 |
| 626.00000 | 35.0 | Cluster_11225 |
| 626.07000 | 35.0 | Cluster_32208 |
| 626.27000 | 35.0 | Cluster_32218 |
| 626.28000 | 35.0 | Cluster_32212 |
| 626.83000 | 35.0 | Cluster_32244 |
| 627.30000 | 35.0 | Cluster_32266 |
| 627.31000 | 35.0 | Cluster_32263 |
| 627.50000 | 35.0 | Cluster_32275 |
| 627.79000 | 35.0 | Cluster_32283 |
| 628.07000 | 35.0 | Cluster_32292 |
| 628.86000 | 35.0 | Cluster_32321 |
| 629.63000 | 35.0 | Cluster_11289 |
| 629.83000 | 35.0 | Cluster_32344 |
| 629.96000 | 35.0 | Cluster_11292 |
| 630.84000 | 35.0 | Cluster_32375 |
| 631.37000 | 35.0 | Cluster_11312 |
| 631.80000 | 35.0 | Cluster_32410 |
| 632.05000 | 35.0 | Cluster_11324 |
| 632.35000 | 35.0 | Cluster_11329 |
| 632.78000 | 35.0 | Cluster_32445 |
| 632.84000 | 35.0 | Cluster_32450 |
| 633.00000 | 35.0 | Cluster_11339 |
| 633.32000 | 35.0 | Cluster_32465 |
| 633.36000 | 35.0 | Cluster_32472 |
| 635.07000 | 35.0 | Cluster_02772 |
| 635.82000 | 35.0 | Cluster_02775 |
| 636.32000 | 35.0 | Cluster_32584 |
| 636.84000 | 35.0 | Cluster_32596 |
| 637.62000 | 35.0 | Cluster_11403 |
| 637.83000 | 35.0 | Cluster_32632 |
| 638.35000 | 35.0 | Cluster_32655 |
| 638.36000 | 35.0 | Cluster_11416 |
| 638.39000 | 35.0 | Cluster_32659 |
| 638.79000 | 35.0 | Cluster_32669 |
| 638.85000 | 35.0 | Cluster_32673 |
| 639.85000 | 35.0 | Cluster_32724 |
| 639.99000 | 35.0 | Cluster_11446 |
| 640.32000 | 35.0 | Cluster_32732 |
| 641.64000 | 35.0 | Cluster_11472 |
| 641.84000 | 35.0 | Cluster_32794 |
| 642.02000 | 35.0 | Cluster_11476 |
| 643.07000 | 35.0 | Cluster_32838 |
| 643.83000 | 35.0 | Cluster_32863 |
| 644.00000 | 35.0 | Cluster_11510 |
| 644.30000 | 35.0 | Cluster_02836 |
| 644.36000 | 35.0 | Cluster_32878 |
| 644.37000 | 35.0 | Cluster_11511 |
| 645.00000 | 35.0 | Cluster_11529 |
| 645.85000 | 35.0 | Cluster_32926 |
| 646.30000 | 35.0 | Cluster_32945 |

| | | |
|---|---|---|
| 648.34000 | 35.0 | Cluster_33005 |
| 648.67000 | 35.0 | Cluster_11566 |
| 648.88000 | 35.0 | Cluster_33029 |
| 649.01000 | 35.0 | Cluster_11574 |
| 649.77000 | 35.0 | Cluster_33054 |
| 650.31000 | 35.0 | Cluster_33085 |
| 651.01000 | 35.0 | Cluster_11595 |
| 651.08000 | 35.0 | Cluster_33114 |
| 651.32000 | 35.0 | Cluster_33124 |
| 651.36000 | 35.0 | Cluster_11600 |
| 651.86000 | 35.0 | Cluster_33158 |
| 651.99000 | 35.0 | Cluster_11614 |
| 652.06000 | 35.0 | Cluster_33159 |
| 652.35000 | 35.0 | Cluster_33162 |
| 652.86000 | 35.0 | Cluster_33187 |
| 653.33000 | 35.0 | Cluster_33199 |
| 653.57000 | 35.0 | Cluster_33207 |
| 653.60000 | 35.0 | Cluster_02883 |
| 654.05000 | 35.0 | Cluster_33221 |
| 654.68000 | 35.0 | Cluster_11655 |
| 654.78000 | 35.0 | Cluster_33237 |
| 655.99000 | 35.0 | Cluster_11675 |
| 656.63000 | 35.0 | Cluster_11676 |
| 657.90000 | 35.0 | Cluster_33331 |
| 657.95000 | 35.0 | Cluster_33350 |
| 658.07000 | 35.0 | Cluster_33340 |
| 658.66000 | 35.0 | Cluster_11699 |
| 659.28000 | 35.0 | Cluster_33386 |
| 659.60000 | 35.0 | Cluster_33396 |
| 659.63000 | 35.0 | Cluster_02907 |
| 659.98000 | 35.0 | Cluster_11725 |
| 660.33000 | 35.0 | Cluster_33416 |
| 660.46000 | 35.0 | Cluster_33423 |
| 660.84000 | 35.0 | Cluster_33430 |
| 661.01000 | 35.0 | Cluster_11741 |
| 661.31000 | 35.0 | Cluster_33436 |
| 662.02000 | 35.0 | Cluster_11748 |
| 662.12000 | 35.0 | Cluster_02928 |
| 662.30000 | 35.0 | Cluster_33472 |
| 663.37000 | 35.0 | Cluster_33509 |
| 663.55000 | 35.0 | Cluster_02941 |
| 663.60000 | 35.0 | Cluster_02940 |
| 665.32000 | 35.0 | Cluster_33570 |
| 666.35000 | 35.0 | Cluster_02964 |
| 666.68000 | 35.0 | Cluster_11814 |
| 666.95000 | 35.0 | Cluster_33617 |
| 667.82000 | 35.0 | Cluster_33644 |
| 668.33000 | 35.0 | Cluster_33651 |
| 668.35000 | 35.0 | Cluster_11848 |
| 669.10000 | 35.0 | Cluster_02977 |
| 669.33000 | 35.0 | Cluster_33680 |

| | | |
|---|---|---|
| 669.78000 | 35.0 | Cluster_33695 |
| 670.01000 | 35.0 | Cluster_11876 |
| 670.40000 | 35.0 | Cluster_33725 |
| 670.80000 | 35.0 | Cluster_33734 |
| 670.82000 | 35.0 | Cluster_33742 |
| 670.87000 | 35.0 | Cluster_02986 |
| 671.44000 | 35.0 | Cluster_33760 |
| 671.82000 | 35.0 | Cluster_33770 |
| 671.97000 | 35.0 | Cluster_11894 |
| 672.35000 | 35.0 | Cluster_33781 |
| 672.81000 | 35.0 | Cluster_33789 |
| 673.09000 | 35.0 | Cluster_03005 |
| 673.37000 | 35.0 | Cluster_33802 |
| 673.62000 | 35.0 | Cluster_33824 |
| 673.65000 | 35.0 | Cluster_11927 |
| 674.11000 | 35.0 | Cluster_03014 |
| 674.32000 | 35.0 | Cluster_11933 |
| 674.95000 | 35.0 | Cluster_00274 |
| 675.37000 | 35.0 | Cluster_33861 |
| 675.60000 | 35.0 | Cluster_33864 |
| 675.64000 | 35.0 | Cluster_03025 |
| 675.84000 | 35.0 | Cluster_03026 |
| 676.00000 | 35.0 | Cluster_11956 |
| 676.32000 | 35.0 | Cluster_11959 |
| 677.20000 | 35.0 | Cluster_33905 |
| 677.89000 | 35.0 | Cluster_33946 |
| 678.81000 | 35.0 | Cluster_33981 |
| 678.84000 | 35.0 | Cluster_33980 |
| 678.85000 | 35.0 | Cluster_03050 |
| 679.38000 | 35.0 | Cluster_33996 |
| 679.79000 | 35.0 | Cluster_34007 |
| 679.86000 | 35.0 | Cluster_34018 |
| 680.08000 | 35.0 | Cluster_03060 |
| 680.09000 | 35.0 | Cluster_34023 |
| 680.87000 | 35.0 | Cluster_34054 |
| 681.08000 | 35.0 | Cluster_03066 |
| 681.32000 | 35.0 | Cluster_12020 |
| 681.35000 | 35.0 | Cluster_34074 |
| 682.09000 | 35.0 | Cluster_34089 |
| 682.31000 | 35.0 | Cluster_34106 |
| 682.38000 | 35.0 | Cluster_03074 |
| 683.61000 | 35.0 | Cluster_12059 |
| 683.81000 | 35.0 | Cluster_34134 |
| 683.99000 | 35.0 | Cluster_12062 |
| 684.08000 | 35.0 | Cluster_03088 |
| 684.82000 | 35.0 | Cluster_34171 |
| 684.83000 | 35.0 | Cluster_34174 |
| 685.09000 | 35.0 | Cluster_03095 |
| 685.10000 | 35.0 | Cluster_03102 |
| 685.35000 | 35.0 | Cluster_34201 |
| 686.04000 | 35.0 | Cluster_12086 |

| | | |
|---|---|---|
| 686.50000 | 35.0 | Cluster_00285 |
| 686.75000 | 35.0 | Cluster_00286 |
| 687.66000 | 35.0 | Cluster_34255 |
| 689.34000 | 35.0 | Cluster_34298 |
| 689.36000 | 35.0 | Cluster_34295 |
| 689.57000 | 35.0 | Cluster_03142 |
| 690.82000 | 35.0 | Cluster_34324 |
| 690.84000 | 35.0 | Cluster_34327 |
| 691.59000 | 35.0 | Cluster_03162 |
| 691.79000 | 35.0 | Cluster_34357 |
| 691.83000 | 35.0 | Cluster_34369 |
| 692.26000 | 35.0 | Cluster_34373 |
| 692.32000 | 35.0 | Cluster_03167 |
| 692.35000 | 35.0 | Cluster_03168 |
| 692.60000 | 35.0 | Cluster_03182 |
| 692.84000 | 35.0 | Cluster_34402 |
| 693.56000 | 35.0 | Cluster_03192 |
| 693.60000 | 35.0 | Cluster_34413 |
| 693.88000 | 35.0 | Cluster_34418 |
| 694.36000 | 35.0 | Cluster_34424 |
| 694.84000 | 35.0 | Cluster_34436 |
| 694.87000 | 35.0 | Cluster_34443 |
| 695.01000 | 35.0 | Cluster_12183 |
| 695.36000 | 35.0 | Cluster_34458 |
| 695.70000 | 35.0 | Cluster_12190 |
| 695.87000 | 35.0 | Cluster_34468 |
| 696.32000 | 35.0 | Cluster_34489 |
| 696.34000 | 35.0 | Cluster_03210 |
| 696.39000 | 35.0 | Cluster_34483 |
| 696.58000 | 35.0 | Cluster_03211 |
| 697.31000 | 35.0 | Cluster_03215 |
| 697.37000 | 35.0 | Cluster_34515 |
| 697.85000 | 35.0 | Cluster_34520 |
| 697.87000 | 35.0 | Cluster_34523 |
| 698.82000 | 35.0 | Cluster_34541 |
| 698.83000 | 35.0 | Cluster_34546 |
| 699.70000 | 35.0 | Cluster_12244 |
| 699.85000 | 35.0 | Cluster_03235 |
| 700.26000 | 35.0 | Cluster_34580 |
| 700.36000 | 35.0 | Cluster_34589 |
| 700.65000 | 35.0 | Cluster_34590 |
| 700.85000 | 35.0 | Cluster_34594 |
| 701.36000 | 35.0 | Cluster_34614 |
| 702.34000 | 35.0 | Cluster_34653 |
| 702.62000 | 35.0 | Cluster_12286 |
| 703.04000 | 35.0 | Cluster_12291 |
| 703.05000 | 35.0 | Cluster_03266 |
| 703.33000 | 35.0 | Cluster_34677 |
| 703.69000 | 35.0 | Cluster_12310 |
| 703.91000 | 35.0 | Cluster_03274 |
| 704.17000 | 35.0 | Cluster_34698 |

| | | |
|---|---|---|
| 704.21000 | 35.0 | Cluster_34700 |
| 704.31000 | 35.0 | Cluster_34706 |
| 704.81000 | 35.0 | Cluster_34728 |
| 704.85000 | 35.0 | Cluster_34734 |
| 705.04000 | 35.0 | Cluster_12319 |
| 705.37000 | 35.0 | Cluster_34742 |
| 705.83000 | 35.0 | Cluster_34756 |
| 705.88000 | 35.0 | Cluster_34757 |
| 706.86000 | 35.0 | Cluster_03301 |
| 707.02000 | 35.0 | Cluster_12346 |
| 707.80000 | 35.0 | Cluster_34791 |
| 708.51000 | 35.0 | Cluster_00309 |
| 709.28000 | 35.0 | Cluster_34840 |
| 709.39000 | 35.0 | Cluster_34846 |
| 710.88000 | 35.0 | Cluster_03335 |
| 711.06000 | 35.0 | Cluster_12397 |
| 711.85000 | 35.0 | Cluster_34925 |
| 712.39000 | 35.0 | Cluster_34948 |
| 712.84000 | 35.0 | Cluster_34967 |
| 713.08000 | 35.0 | Cluster_03351 |
| 713.29000 | 35.0 | Cluster_03356 |
| 713.38000 | 35.0 | Cluster_12431 |
| 713.72000 | 35.0 | Cluster_12437 |
| 713.77000 | 35.0 | Cluster_34987 |
| 713.92000 | 35.0 | Cluster_34998 |
| 714.32000 | 35.0 | Cluster_35004 |
| 714.36000 | 35.0 | Cluster_35010 |
| 715.38000 | 35.0 | Cluster_35041 |
| 715.54000 | 35.0 | Cluster_00320 |
| 715.71000 | 35.0 | Cluster_12456 |
| 715.85000 | 35.0 | Cluster_35046 |
| 715.99000 | 35.0 | Cluster_12465 |
| 716.39000 | 35.0 | Cluster_35061 |
| 717.36000 | 35.0 | Cluster_12478 |
| 717.90000 | 35.0 | Cluster_35102 |
| 718.36000 | 35.0 | Cluster_35122 |
| 718.85000 | 35.0 | Cluster_35127 |
| 718.89000 | 35.0 | Cluster_35137 |
| 718.91000 | 35.0 | Cluster_35135 |
| 718.96000 | 35.0 | Cluster_12511 |
| 719.39000 | 35.0 | Cluster_35145 |
| 719.83000 | 35.0 | Cluster_35161 |
| 719.85000 | 35.0 | Cluster_03411 |
| 719.98000 | 35.0 | Cluster_12519 |
| 720.12000 | 35.0 | Cluster_35168 |
| 720.13000 | 35.0 | Cluster_35170 |
| 720.35000 | 35.0 | Cluster_03417 |
| 720.58000 | 35.0 | Cluster_03421 |
| 720.83000 | 35.0 | Cluster_35178 |
| 721.32000 | 35.0 | Cluster_35195 |
| 721.41000 | 35.0 | Cluster_35191 |

| | | |
|---|---|---|
| 721.72000 | 35.0 | Cluster_12548 |
| 721.88000 | 35.0 | Cluster_35209 |
| 721.90000 | 35.0 | Cluster_35213 |
| 722.02000 | 35.0 | Cluster_12553 |
| 722.09000 | 35.0 | Cluster_03436 |
| 722.34000 | 35.0 | Cluster_03433 |
| 722.68000 | 35.0 | Cluster_12565 |
| 723.40000 | 35.0 | Cluster_12568 |
| 723.42000 | 35.0 | Cluster_12570 |
| 723.95000 | 35.0 | Cluster_00326 |
| 724.04000 | 35.0 | Cluster_12575 |
| 724.90000 | 35.0 | Cluster_03465 |
| 725.06000 | 35.0 | Cluster_12593 |
| 725.32000 | 35.0 | Cluster_03469 |
| 725.41000 | 35.0 | Cluster_35286 |
| 725.42000 | 35.0 | Cluster_35293 |
| 727.38000 | 35.0 | Cluster_35317 |
| 728.10000 | 35.0 | Cluster_03487 |
| 728.13000 | 35.0 | Cluster_03484 |
| 728.42000 | 35.0 | Cluster_35338 |
| 728.98000 | 35.0 | Cluster_12653 |
| 729.15000 | 35.0 | Cluster_03497 |
| 729.84000 | 35.0 | Cluster_35373 |
| 729.89000 | 35.0 | Cluster_03507 |
| 730.91000 | 35.0 | Cluster_35408 |
| 731.10000 | 35.0 | Cluster_35410 |
| 731.83000 | 35.0 | Cluster_03518 |
| 732.11000 | 35.0 | Cluster_03522 |
| 732.64000 | 35.0 | Cluster_03534 |
| 733.03000 | 35.0 | Cluster_12708 |
| 733.13000 | 35.0 | Cluster_03548 |
| 733.83000 | 35.0 | Cluster_35471 |
| 733.85000 | 35.0 | Cluster_35474 |
| 734.58000 | 35.0 | Cluster_35493 |
| 735.04000 | 35.0 | Cluster_12729 |
| 735.74000 | 35.0 | Cluster_12741 |
| 736.37000 | 35.0 | Cluster_35541 |
| 737.04000 | 35.0 | Cluster_12754 |
| 737.16000 | 35.0 | Cluster_35560 |
| 737.82000 | 35.0 | Cluster_35593 |
| 738.39000 | 35.0 | Cluster_35611 |
| 738.63000 | 35.0 | Cluster_03590 |
| 739.38000 | 35.0 | Cluster_12791 |
| 739.39000 | 35.0 | Cluster_12789 |
| 739.84000 | 35.0 | Cluster_35645 |
| 740.37000 | 35.0 | Cluster_35657 |
| 740.38000 | 35.0 | Cluster_35661 |
| 740.93000 | 35.0 | Cluster_35676 |
| 741.85000 | 35.0 | Cluster_35683 |
| 742.37000 | 35.0 | Cluster_35697 |
| 742.67000 | 35.0 | Cluster_12831 |

| | | |
|---|---|---|
| 742.84000 | 35.0 | Cluster_35707 |
| 744.37000 | 35.0 | Cluster_35750 |
| 745.38000 | 35.0 | Cluster_35777 |
| 745.68000 | 35.0 | Cluster_12857 |
| 746.06000 | 35.0 | Cluster_12863 |
| 746.15000 | 35.0 | Cluster_35792 |
| 746.30000 | 35.0 | Cluster_35797 |
| 747.12000 | 35.0 | Cluster_35817 |
| 747.37000 | 35.0 | Cluster_35818 |
| 748.00000 | 35.0 | Cluster_12903 |
| 748.86000 | 35.0 | Cluster_03654 |
| 748.88000 | 35.0 | Cluster_35857 |
| 749.07000 | 35.0 | Cluster_12920 |
| 749.69000 | 35.0 | Cluster_12929 |
| 749.98000 | 35.0 | Cluster_12936 |
| 750.01000 | 35.0 | Cluster_12941 |
| 750.38000 | 35.0 | Cluster_35883 |
| 750.44000 | 35.0 | Cluster_35886 |
| 750.94000 | 35.0 | Cluster_35903 |
| 751.06000 | 35.0 | Cluster_12961 |
| 751.15000 | 35.0 | Cluster_03662 |
| 751.36000 | 35.0 | Cluster_35908 |
| 752.37000 | 35.0 | Cluster_35932 |
| 753.05000 | 35.0 | Cluster_12988 |
| 753.06000 | 35.0 | Cluster_12984 |
| 753.39000 | 35.0 | Cluster_35964 |
| 753.91000 | 35.0 | Cluster_35981 |
| 754.06000 | 35.0 | Cluster_13009 |
| 754.90000 | 35.0 | Cluster_36004 |
| 755.06000 | 35.0 | Cluster_13021 |
| 755.30000 | 35.0 | Cluster_13019 |
| 755.33000 | 35.0 | Cluster_13020 |
| 755.42000 | 35.0 | Cluster_36013 |
| 755.75000 | 35.0 | Cluster_13034 |
| 755.92000 | 35.0 | Cluster_36043 |
| 756.13000 | 35.0 | Cluster_03688 |
| 757.31000 | 35.0 | Cluster_36071 |
| 757.73000 | 35.0 | Cluster_13076 |
| 758.41000 | 35.0 | Cluster_36102 |
| 758.70000 | 35.0 | Cluster_13088 |
| 759.63000 | 35.0 | Cluster_03704 |
| 759.90000 | 35.0 | Cluster_36129 |
| 759.91000 | 35.0 | Cluster_36132 |
| 760.35000 | 35.0 | Cluster_36146 |
| 761.05000 | 35.0 | Cluster_00024 |
| 761.41000 | 35.0 | Cluster_36182 |
| 762.38000 | 35.0 | Cluster_13130 |
| 763.03000 | 35.0 | Cluster_13141 |
| 763.08000 | 35.0 | Cluster_13137 |
| 763.35000 | 35.0 | Cluster_03726 |
| 763.71000 | 35.0 | Cluster_13147 |

| | | |
|---|---|---|
| 764.09000 | 35.0 | Cluster_13148 |
| 764.36000 | 35.0 | Cluster_36253 |
| 765.15000 | 35.0 | Cluster_03741 |
| 765.79000 | 35.0 | Cluster_36274 |
| 765.80000 | 35.0 | Cluster_36275 |
| 766.37000 | 35.0 | Cluster_36292 |
| 766.55000 | 35.0 | Cluster_00339 |
| 767.70000 | 35.0 | Cluster_13203 |
| 767.85000 | 35.0 | Cluster_36325 |
| 768.07000 | 35.0 | Cluster_13209 |
| 768.08000 | 35.0 | Cluster_13205 |
| 768.38000 | 35.0 | Cluster_36351 |
| 769.33000 | 35.0 | Cluster_36370 |
| 769.59000 | 35.0 | Cluster_36368 |
| 769.82000 | 35.0 | Cluster_36385 |
| 772.41000 | 35.0 | Cluster_36441 |
| 772.74000 | 35.0 | Cluster_13265 |
| 772.80000 | 35.0 | Cluster_13273 |
| 772.88000 | 35.0 | Cluster_36459 |
| 772.92000 | 35.0 | Cluster_36460 |
| 773.08000 | 35.0 | Cluster_03785 |
| 773.09000 | 35.0 | Cluster_03786 |
| 773.13000 | 35.0 | Cluster_13272 |
| 773.35000 | 35.0 | Cluster_13283 |
| 773.41000 | 35.0 | Cluster_13286 |
| 774.12000 | 35.0 | Cluster_36481 |
| 774.73000 | 35.0 | Cluster_13306 |
| 775.06000 | 35.0 | Cluster_13312 |
| 775.07000 | 35.0 | Cluster_13311 |
| 775.42000 | 35.0 | Cluster_36510 |
| 776.37000 | 35.0 | Cluster_03803 |
| 777.15000 | 35.0 | Cluster_03806 |
| 777.35000 | 35.0 | Cluster_36561 |
| 777.38000 | 35.0 | Cluster_36567 |
| 777.39000 | 35.0 | Cluster_36564 |
| 778.37000 | 35.0 | Cluster_36591 |
| 778.77000 | 35.0 | Cluster_13350 |
| 778.87000 | 35.0 | Cluster_36605 |
| 780.73000 | 35.0 | Cluster_13379 |
| 780.81000 | 35.0 | Cluster_36636 |
| 780.88000 | 35.0 | Cluster_03821 |
| 781.40000 | 35.0 | Cluster_36661 |
| 782.03000 | 35.0 | Cluster_13403 |
| 783.13000 | 35.0 | Cluster_03834 |
| 783.46000 | 35.0 | Cluster_36708 |
| 783.72000 | 35.0 | Cluster_13422 |
| 784.06000 | 35.0 | Cluster_13433 |
| 784.89000 | 35.0 | Cluster_36733 |
| 784.93000 | 35.0 | Cluster_36732 |
| 785.45000 | 35.0 | Cluster_36744 |
| 785.83000 | 35.0 | Cluster_36763 |

| | | |
|---|---|---|
| 787.06000 | 35.0 | Cluster_13459 |
| 787.12000 | 35.0 | Cluster_03868 |
| 787.35000 | 35.0 | Cluster_03870 |
| 788.87000 | 35.0 | Cluster_36840 |
| 789.06000 | 35.0 | Cluster_13483 |
| 790.87000 | 35.0 | Cluster_36906 |
| 791.12000 | 35.0 | Cluster_36909 |
| 791.38000 | 35.0 | Cluster_36914 |
| 791.90000 | 35.0 | Cluster_36919 |
| 792.38000 | 35.0 | Cluster_36933 |
| 792.64000 | 35.0 | Cluster_36941 |
| 792.86000 | 35.0 | Cluster_03914 |
| 792.87000 | 35.0 | Cluster_36948 |
| 792.97000 | 35.0 | Cluster_13541 |
| 793.78000 | 35.0 | Cluster_13547 |
| 794.36000 | 35.0 | Cluster_36984 |
| 794.37000 | 35.0 | Cluster_36982 |
| 794.45000 | 35.0 | Cluster_36985 |
| 794.65000 | 35.0 | Cluster_36986 |
| 795.07000 | 35.0 | Cluster_13571 |
| 795.42000 | 35.0 | Cluster_37004 |
| 795.44000 | 35.0 | Cluster_37000 |
| 795.68000 | 35.0 | Cluster_13585 |
| 795.87000 | 35.0 | Cluster_03947 |
| 795.90000 | 35.0 | Cluster_37017 |
| 795.96000 | 35.0 | Cluster_37020 |
| 796.41000 | 35.0 | Cluster_13605 |
| 796.72000 | 35.0 | Cluster_13597 |
| 796.91000 | 35.0 | Cluster_37044 |
| 797.86000 | 35.0 | Cluster_37077 |
| 797.87000 | 35.0 | Cluster_37073 |
| 797.88000 | 35.0 | Cluster_37072 |
| 797.89000 | 35.0 | Cluster_37068 |
| 799.38000 | 35.0 | Cluster_37115 |
| 799.40000 | 35.0 | Cluster_37112 |
| 799.89000 | 35.0 | Cluster_03975 |
| 800.35000 | 35.0 | Cluster_37146 |
| 800.41000 | 35.0 | Cluster_13643 |
| 800.84000 | 35.0 | Cluster_37149 |
| 800.86000 | 35.0 | Cluster_03991 |
| 801.06000 | 35.0 | Cluster_00029 |
| 801.37000 | 35.0 | Cluster_03995 |
| 801.42000 | 35.0 | Cluster_37163 |
| 801.44000 | 35.0 | Cluster_13660 |
| 802.43000 | 35.0 | Cluster_37188 |
| 802.74000 | 35.0 | Cluster_13681 |
| 803.41000 | 35.0 | Cluster_13684 |
| 803.88000 | 35.0 | Cluster_37226 |
| 804.43000 | 35.0 | Cluster_37243 |
| 804.91000 | 35.0 | Cluster_04015 |
| 805.16000 | 35.0 | Cluster_37269 |

| | | |
|---|---|---|
| 805.39000 | 35.0 | Cluster_37283 |
| 805.94000 | 35.0 | Cluster_37284 |
| 806.39000 | 35.0 | Cluster_37304 |
| 807.01000 | 35.0 | Cluster_13723 |
| 807.46000 | 35.0 | Cluster_37330 |
| 807.92000 | 35.0 | Cluster_37352 |
| 808.85000 | 35.0 | Cluster_37370 |
| 808.91000 | 35.0 | Cluster_37377 |
| 809.40000 | 35.0 | Cluster_04037 |
| 809.91000 | 35.0 | Cluster_37401 |
| 810.39000 | 35.0 | Cluster_37407 |
| 810.44000 | 35.0 | Cluster_13774 |
| 811.74000 | 35.0 | Cluster_00036 |
| 811.86000 | 35.0 | Cluster_37446 |
| 811.91000 | 35.0 | Cluster_37441 |
| 812.39000 | 35.0 | Cluster_37458 |
| 812.41000 | 35.0 | Cluster_13796 |
| 813.82000 | 35.0 | Cluster_00352 |
| 813.90000 | 35.0 | Cluster_37483 |
| 814.10000 | 35.0 | Cluster_13827 |
| 814.16000 | 35.0 | Cluster_37492 |
| 814.17000 | 35.0 | Cluster_04067 |
| 814.90000 | 35.0 | Cluster_04074 |
| 815.06000 | 35.0 | Cluster_13832 |
| 815.79000 | 35.0 | Cluster_13844 |
| 815.92000 | 35.0 | Cluster_37530 |
| 816.38000 | 35.0 | Cluster_37540 |
| 816.43000 | 35.0 | Cluster_04088 |
| 816.72000 | 35.0 | Cluster_13857 |
| 817.34000 | 35.0 | Cluster_37561 |
| 817.88000 | 35.0 | Cluster_37570 |
| 818.42000 | 35.0 | Cluster_13883 |
| 819.05000 | 35.0 | Cluster_13897 |
| 819.42000 | 35.0 | Cluster_13906 |
| 820.17000 | 35.0 | Cluster_04104 |
| 821.23000 | 35.0 | Cluster_04111 |
| 821.38000 | 35.0 | Cluster_37637 |
| 822.06000 | 35.0 | Cluster_13934 |
| 823.42000 | 35.0 | Cluster_37669 |
| 823.91000 | 35.0 | Cluster_37674 |
| 823.94000 | 35.0 | Cluster_37688 |
| 825.32000 | 35.0 | Cluster_37717 |
| 825.39000 | 35.0 | Cluster_04137 |
| 826.38000 | 35.0 | Cluster_13981 |
| 826.86000 | 35.0 | Cluster_37765 |
| 827.04000 | 35.0 | Cluster_13986 |
| 827.06000 | 35.0 | Cluster_13988 |
| 827.09000 | 35.0 | Cluster_13989 |
| 827.88000 | 35.0 | Cluster_37794 |
| 828.80000 | 35.0 | Cluster_14000 |
| 828.98000 | 35.0 | Cluster_14004 |

| | | |
|---|---|---|
| 829.16000 | 35.0 | Cluster_04165 |
| 829.36000 | 35.0 | Cluster_37821 |
| 829.43000 | 35.0 | Cluster_37825 |
| 829.91000 | 35.0 | Cluster_04169 |
| 830.74000 | 35.0 | Cluster_14020 |
| 830.89000 | 35.0 | Cluster_00042 |
| 831.45000 | 35.0 | Cluster_37853 |
| 832.41000 | 35.0 | Cluster_37873 |
| 832.68000 | 35.0 | Cluster_37879 |
| 833.49000 | 35.0 | Cluster_37894 |
| 833.99000 | 35.0 | Cluster_37902 |
| 834.10000 | 35.0 | Cluster_14054 |
| 834.94000 | 35.0 | Cluster_37918 |
| 835.05000 | 35.0 | Cluster_14062 |
| 836.39000 | 35.0 | Cluster_14076 |
| 836.42000 | 35.0 | Cluster_14077 |
| 836.76000 | 35.0 | Cluster_14083 |
| 836.86000 | 35.0 | Cluster_37959 |
| 836.94000 | 35.0 | Cluster_04234 |
| 837.41000 | 35.0 | Cluster_37970 |
| 837.95000 | 35.0 | Cluster_37976 |
| 839.10000 | 35.0 | Cluster_14102 |
| 840.66000 | 35.0 | Cluster_04260 |
| 840.78000 | 35.0 | Cluster_14117 |
| 841.08000 | 35.0 | Cluster_14128 |
| 841.10000 | 35.0 | Cluster_14126 |
| 841.67000 | 35.0 | Cluster_00364 |
| 841.76000 | 35.0 | Cluster_38066 |
| 842.73000 | 35.0 | Cluster_14140 |
| 843.87000 | 35.0 | Cluster_38114 |
| 844.18000 | 35.0 | Cluster_04298 |
| 844.44000 | 35.0 | Cluster_14157 |
| 846.98000 | 35.0 | Cluster_38180 |
| 847.48000 | 35.0 | Cluster_38200 |
| 847.75000 | 35.0 | Cluster_14185 |
| 847.88000 | 35.0 | Cluster_38203 |
| 848.40000 | 35.0 | Cluster_04350 |
| 848.91000 | 35.0 | Cluster_04352 |
| 849.46000 | 35.0 | Cluster_38235 |
| 849.64000 | 35.0 | Cluster_04364 |
| 849.97000 | 35.0 | Cluster_38245 |
| 849.99000 | 35.0 | Cluster_38248 |
| 850.33000 | 35.0 | Cluster_38250 |
| 850.40000 | 35.0 | Cluster_04376 |
| 850.76000 | 35.0 | Cluster_14213 |
| 851.36000 | 35.0 | Cluster_14215 |
| 851.77000 | 35.0 | Cluster_38298 |
| 852.03000 | 35.0 | Cluster_38307 |
| 852.91000 | 35.0 | Cluster_04400 |
| 852.93000 | 35.0 | Cluster_04401 |
| 853.45000 | 35.0 | Cluster_38334 |

| | | |
|---|---|---|
| 854.44000 | 35.0 | Cluster_38377 |
| 854.47000 | 35.0 | Cluster_38375 |
| 854.88000 | 35.0 | Cluster_38385 |
| 855.41000 | 35.0 | Cluster_38392 |
| 855.73000 | 35.0 | Cluster_14240 |
| 856.43000 | 35.0 | Cluster_14249 |
| 856.91000 | 35.0 | Cluster_04444 |
| 857.06000 | 35.0 | Cluster_14257 |
| 857.89000 | 35.0 | Cluster_38459 |
| 857.91000 | 35.0 | Cluster_04449 |
| 858.07000 | 35.0 | Cluster_14269 |
| 858.77000 | 35.0 | Cluster_14275 |
| 859.45000 | 35.0 | Cluster_38500 |
| 859.47000 | 35.0 | Cluster_38497 |
| 860.47000 | 35.0 | Cluster_14284 |
| 861.39000 | 35.0 | Cluster_04493 |
| 862.39000 | 35.0 | Cluster_04505 |
| 862.49000 | 35.0 | Cluster_38591 |
| 863.05000 | 35.0 | Cluster_14302 |
| 863.76000 | 35.0 | Cluster_14307 |
| 863.94000 | 35.0 | Cluster_38615 |
| 864.07000 | 35.0 | Cluster_14314 |
| 865.03000 | 35.0 | Cluster_38646 |
| 865.75000 | 35.0 | Cluster_14326 |
| 865.92000 | 35.0 | Cluster_38659 |
| 865.93000 | 35.0 | Cluster_04555 |
| 866.41000 | 35.0 | Cluster_14334 |
| 869.07000 | 35.0 | Cluster_14353 |
| 869.28000 | 35.0 | Cluster_38719 |
| 869.41000 | 35.0 | Cluster_04599 |
| 869.78000 | 35.0 | Cluster_14363 |
| 871.17000 | 35.0 | Cluster_38758 |
| 871.27000 | 35.0 | Cluster_38759 |
| 871.44000 | 35.0 | Cluster_38769 |
| 871.47000 | 35.0 | Cluster_14388 |
| 872.27000 | 35.0 | Cluster_38802 |
| 874.09000 | 35.0 | Cluster_14414 |
| 874.26000 | 35.0 | Cluster_38844 |
| 874.78000 | 35.0 | Cluster_14424 |
| 875.28000 | 35.0 | Cluster_38860 |
| 875.66000 | 35.0 | Cluster_04698 |
| 876.08000 | 35.0 | Cluster_14437 |
| 876.53000 | 35.0 | Cluster_38895 |
| 877.38000 | 35.0 | Cluster_14453 |
| 877.44000 | 35.0 | Cluster_38909 |
| 878.48000 | 35.0 | Cluster_38928 |
| 880.46000 | 35.0 | Cluster_14483 |
| 881.96000 | 35.0 | Cluster_38993 |
| 882.44000 | 35.0 | Cluster_39008 |
| 882.94000 | 35.0 | Cluster_39018 |
| 884.40000 | 35.0 | Cluster_14528 |

| | | |
|---|---|---|
| 884.66000 | 35.0 | Cluster_39047 |
| 885.12000 | 35.0 | Cluster_14544 |
| 885.68000 | 35.0 | Cluster_04765 |
| 885.95000 | 35.0 | Cluster_39078 |
| 886.43000 | 35.0 | Cluster_39095 |
| 886.96000 | 35.0 | Cluster_39099 |
| 887.52000 | 35.0 | Cluster_39107 |
| 887.68000 | 35.0 | Cluster_04781 |
| 887.74000 | 35.0 | Cluster_14571 |
| 888.13000 | 35.0 | Cluster_14577 |
| 888.40000 | 35.0 | Cluster_14578 |
| 888.91000 | 35.0 | Cluster_39135 |
| 888.92000 | 35.0 | Cluster_04786 |
| 889.11000 | 35.0 | Cluster_14581 |
| 889.44000 | 35.0 | Cluster_39147 |
| 890.17000 | 35.0 | Cluster_04796 |
| 890.38000 | 35.0 | Cluster_39161 |
| 890.39000 | 35.0 | Cluster_39160 |
| 890.46000 | 35.0 | Cluster_39173 |
| 890.99000 | 35.0 | Cluster_39180 |
| 891.46000 | 35.0 | Cluster_39186 |
| 891.81000 | 35.0 | Cluster_14598 |
| 892.21000 | 35.0 | Cluster_04824 |
| 893.43000 | 35.0 | Cluster_39231 |
| 893.78000 | 35.0 | Cluster_14615 |
| 894.12000 | 35.0 | Cluster_14617 |
| 894.71000 | 35.0 | Cluster_04845 |
| 895.21000 | 35.0 | Cluster_04851 |
| 895.45000 | 35.0 | Cluster_14622 |
| 895.81000 | 35.0 | Cluster_14625 |
| 896.15000 | 35.0 | Cluster_14634 |
| 897.04000 | 35.0 | Cluster_39292 |
| 898.77000 | 35.0 | Cluster_14664 |
| 899.43000 | 35.0 | Cluster_39337 |
| 899.51000 | 35.0 | Cluster_39345 |
| 899.96000 | 35.0 | Cluster_04891 |
| 900.94000 | 35.0 | Cluster_39368 |
| 900.97000 | 35.0 | Cluster_39380 |
| 900.99000 | 35.0 | Cluster_39377 |
| 902.46000 | 35.0 | Cluster_39412 |
| 904.46000 | 35.0 | Cluster_39443 |
| 905.43000 | 35.0 | Cluster_39461 |
| 906.77000 | 35.0 | Cluster_14739 |
| 907.80000 | 35.0 | Cluster_14749 |
| 908.46000 | 35.0 | Cluster_39500 |
| 908.48000 | 35.0 | Cluster_14756 |
| 908.94000 | 35.0 | Cluster_04958 |
| 908.95000 | 35.0 | Cluster_39504 |
| 909.44000 | 35.0 | Cluster_14760 |
| 909.50000 | 35.0 | Cluster_14765 |
| 909.89000 | 35.0 | Cluster_39520 |

| | | |
|---|---|---|
| 910.11000 | 35.0 | Cluster_14769 |
| 910.19000 | 35.0 | Cluster_04966 |
| 911.96000 | 35.0 | Cluster_39564 |
| 914.56000 | 35.0 | Cluster_39615 |
| 915.13000 | 35.0 | Cluster_14823 |
| 915.93000 | 35.0 | Cluster_39626 |
| 916.44000 | 35.0 | Cluster_39640 |
| 916.49000 | 35.0 | Cluster_39633 |
| 917.45000 | 35.0 | Cluster_14854 |
| 917.97000 | 35.0 | Cluster_39674 |
| 918.66000 | 35.0 | Cluster_39698 |
| 920.45000 | 35.0 | Cluster_39721 |
| 920.76000 | 35.0 | Cluster_14891 |
| 920.79000 | 35.0 | Cluster_39725 |
| 920.88000 | 35.0 | Cluster_39726 |
| 920.95000 | 35.0 | Cluster_39731 |
| 922.42000 | 35.0 | Cluster_14907 |
| 922.98000 | 35.0 | Cluster_39772 |
| 923.30000 | 35.0 | Cluster_14924 |
| 923.45000 | 35.0 | Cluster_14932 |
| 923.92000 | 35.0 | Cluster_39782 |
| 924.49000 | 35.0 | Cluster_39788 |
| 925.11000 | 35.0 | Cluster_14961 |
| 926.42000 | 35.0 | Cluster_14975 |
| 927.48000 | 35.0 | Cluster_39834 |
| 928.47000 | 35.0 | Cluster_39858 |
| 928.96000 | 35.0 | Cluster_39861 |
| 929.12000 | 35.0 | Cluster_15009 |
| 929.43000 | 35.0 | Cluster_39880 |
| 929.90000 | 35.0 | Cluster_39897 |
| 930.47000 | 35.0 | Cluster_15020 |
| 931.73000 | 35.0 | Cluster_15043 |
| 931.90000 | 35.0 | Cluster_39924 |
| 932.64000 | 35.0 | Cluster_15062 |
| 932.95000 | 35.0 | Cluster_39941 |
| 933.14000 | 35.0 | Cluster_15072 |
| 933.33000 | 35.0 | Cluster_05038 |
| 934.00000 | 35.0 | Cluster_39963 |
| 934.13000 | 35.0 | Cluster_15084 |
| 934.43000 | 35.0 | Cluster_15083 |
| 934.78000 | 35.0 | Cluster_15089 |
| 935.78000 | 35.0 | Cluster_15100 |
| 936.45000 | 35.0 | Cluster_15116 |
| 936.80000 | 35.0 | Cluster_15129 |
| 937.00000 | 35.0 | Cluster_39997 |
| 937.44000 | 35.0 | Cluster_15144 |
| 937.49000 | 35.0 | Cluster_05051 |
| 938.07000 | 35.0 | Cluster_15154 |
| 938.13000 | 35.0 | Cluster_15156 |
| 938.21000 | 35.0 | Cluster_15160 |
| 939.69000 | 35.0 | Cluster_05058 |

| | | |
|---|---|---|
| 940.46000 | 35.0 | Cluster_15180 |
| 941.72000 | 35.0 | Cluster_40098 |
| 942.00000 | 35.0 | Cluster_40094 |
| 943.45000 | 35.0 | Cluster_15229 |
| 945.13000 | 35.0 | Cluster_15244 |
| 945.39000 | 35.0 | Cluster_15242 |
| 945.72000 | 35.0 | Cluster_05075 |
| 945.73000 | 35.0 | Cluster_05078 |
| 945.78000 | 35.0 | Cluster_15250 |
| 945.98000 | 35.0 | Cluster_40152 |
| 947.16000 | 35.0 | Cluster_15264 |
| 947.22000 | 35.0 | Cluster_05084 |
| 947.79000 | 35.0 | Cluster_15273 |
| 948.19000 | 35.0 | Cluster_15277 |
| 948.52000 | 35.0 | Cluster_40180 |
| 949.14000 | 35.0 | Cluster_15301 |
| 949.44000 | 35.0 | Cluster_40190 |
| 949.46000 | 35.0 | Cluster_15302 |
| 950.44000 | 35.0 | Cluster_15314 |
| 951.17000 | 35.0 | Cluster_15324 |
| 951.96000 | 35.0 | Cluster_40227 |
| 952.14000 | 35.0 | Cluster_15337 |
| 952.15000 | 35.0 | Cluster_15339 |
| 952.44000 | 35.0 | Cluster_15342 |
| 952.53000 | 35.0 | Cluster_40233 |
| 953.98000 | 35.0 | Cluster_40255 |
| 954.82000 | 35.0 | Cluster_15379 |
| 954.99000 | 35.0 | Cluster_40262 |
| 955.13000 | 35.0 | Cluster_15394 |
| 955.47000 | 35.0 | Cluster_15404 |
| 956.48000 | 35.0 | Cluster_15426 |
| 957.07000 | 35.0 | Cluster_40290 |
| 957.48000 | 35.0 | Cluster_40291 |
| 960.15000 | 35.0 | Cluster_15462 |
| 960.98000 | 35.0 | Cluster_40317 |
| 961.48000 | 35.0 | Cluster_15478 |
| 962.14000 | 35.0 | Cluster_15497 |
| 962.38000 | 35.0 | Cluster_40344 |
| 962.79000 | 35.0 | Cluster_15505 |
| 964.13000 | 35.0 | Cluster_15523 |
| 964.43000 | 35.0 | Cluster_40365 |
| 965.29000 | 35.0 | Cluster_40377 |
| 966.15000 | 35.0 | Cluster_15549 |
| 966.49000 | 35.0 | Cluster_40387 |
| 966.50000 | 35.0 | Cluster_05172 |
| 966.85000 | 35.0 | Cluster_15552 |
| 967.00000 | 35.0 | Cluster_40392 |
| 967.56000 | 35.0 | Cluster_15565 |
| 968.03000 | 35.0 | Cluster_40402 |
| 968.25000 | 35.0 | Cluster_05187 |
| 971.97000 | 35.0 | Cluster_40451 |

| | | |
|---|---|---|
| 972.01000 | 35.0 | Cluster_40467 |
| 972.46000 | 35.0 | Cluster_15627 |
| 972.49000 | 35.0 | Cluster_15626 |
| 973.51000 | 35.0 | Cluster_40476 |
| 974.47000 | 35.0 | Cluster_15651 |
| 974.98000 | 35.0 | Cluster_40495 |
| 975.82000 | 35.0 | Cluster_15672 |
| 975.98000 | 35.0 | Cluster_05210 |
| 976.00000 | 35.0 | Cluster_05212 |
| 976.03000 | 35.0 | Cluster_40504 |
| 977.14000 | 35.0 | Cluster_15696 |
| 977.62000 | 35.0 | Cluster_40522 |
| 979.45000 | 35.0 | Cluster_40546 |
| 979.80000 | 35.0 | Cluster_15720 |
| 979.83000 | 35.0 | Cluster_15721 |
| 979.98000 | 35.0 | Cluster_40568 |
| 980.00000 | 35.0 | Cluster_40562 |
| 980.50000 | 35.0 | Cluster_40565 |
| 980.70000 | 35.0 | Cluster_40578 |
| 981.45000 | 35.0 | Cluster_15750 |
| 983.25000 | 35.0 | Cluster_05237 |
| 983.95000 | 35.0 | Cluster_40617 |
| 985.14000 | 35.0 | Cluster_40628 |
| 986.02000 | 35.0 | Cluster_40630 |
| 986.52000 | 35.0 | Cluster_40639 |
| 986.83000 | 35.0 | Cluster_15810 |
| 987.24000 | 35.0 | Cluster_40654 |
| 987.80000 | 35.0 | Cluster_15825 |
| 988.44000 | 35.0 | Cluster_15835 |
| 991.93000 | 35.0 | Cluster_40717 |
| 993.50000 | 35.0 | Cluster_15895 |
| 997.19000 | 35.0 | Cluster_15954 |
| 1000.06000 | 35.0 | Cluster_40851 |
| 1001.03000 | 35.0 | Cluster_40878 |
| 1001.16000 | 35.0 | Cluster_15996 |
| 1001.49000 | 35.0 | Cluster_40884 |
| 1003.52000 | 35.0 | Cluster_05294 |
| 1004.00000 | 35.0 | Cluster_40937 |
| 1004.31000 | 35.0 | Cluster_05296 |
| 1004.57000 | 35.0 | Cluster_40940 |
| 1007.81000 | 35.0 | Cluster_16072 |
| 1008.78000 | 35.0 | Cluster_16080 |
| 1012.26000 | 35.0 | Cluster_05329 |
| 1012.27000 | 35.0 | Cluster_41044 |
| 1012.93000 | 35.0 | Cluster_41054 |
| 1013.01000 | 35.0 | Cluster_05341 |
| 1013.47000 | 35.0 | Cluster_16138 |
| 1013.49000 | 35.0 | Cluster_16137 |
| 1013.76000 | 35.0 | Cluster_05344 |
| 1013.90000 | 35.0 | Cluster_41056 |
| 1015.92000 | 35.0 | Cluster_41088 |

| | | |
|---|---|---|
| 1016.78000 | 35.0 | Cluster_05351 |
| 1017.27000 | 35.0 | Cluster_41099 |
| 1019.25000 | 35.0 | Cluster_05360 |
| 1020.15000 | 35.0 | Cluster_16210 |
| 1022.50000 | 35.0 | Cluster_16231 |
| 1023.82000 | 35.0 | Cluster_41185 |
| 1024.48000 | 35.0 | Cluster_16254 |
| 1024.50000 | 35.0 | Cluster_16256 |
| 1024.93000 | 35.0 | Cluster_41195 |
| 1025.46000 | 35.0 | Cluster_16264 |
| 1025.81000 | 35.0 | Cluster_41211 |
| 1026.83000 | 35.0 | Cluster_16274 |
| 1027.13000 | 35.0 | Cluster_16285 |
| 1027.14000 | 35.0 | Cluster_16284 |
| 1027.88000 | 35.0 | Cluster_16291 |
| 1028.18000 | 35.0 | Cluster_16295 |
| 1028.84000 | 35.0 | Cluster_16300 |
| 1029.99000 | 35.0 | Cluster_41262 |
| 1031.20000 | 35.0 | Cluster_16323 |
| 1032.82000 | 35.0 | Cluster_16335 |
| 1033.20000 | 35.0 | Cluster_16342 |
| 1036.52000 | 35.0 | Cluster_16382 |
| 1037.50000 | 35.0 | Cluster_16400 |
| 1039.05000 | 35.0 | Cluster_41371 |
| 1039.52000 | 35.0 | Cluster_41374 |
| 1039.81000 | 35.0 | Cluster_16429 |
| 1040.83000 | 35.0 | Cluster_05437 |
| 1041.01000 | 35.0 | Cluster_41401 |
| 1042.17000 | 35.0 | Cluster_16457 |
| 1044.49000 | 35.0 | Cluster_41432 |
| 1046.13000 | 35.0 | Cluster_16488 |
| 1046.22000 | 35.0 | Cluster_16495 |
| 1047.16000 | 35.0 | Cluster_16507 |
| 1047.83000 | 35.0 | Cluster_41470 |
| 1053.50000 | 35.0 | Cluster_16581 |
| 1053.98000 | 35.0 | Cluster_41534 |
| 1054.99000 | 35.0 | Cluster_05476 |
| 1055.22000 | 35.0 | Cluster_16595 |
| 1055.50000 | 35.0 | Cluster_16606 |
| 1056.52000 | 35.0 | Cluster_16616 |
| 1057.06000 | 35.0 | Cluster_41567 |
| 1057.33000 | 35.0 | Cluster_05486 |
| 1057.34000 | 35.0 | Cluster_05487 |
| 1057.58000 | 35.0 | Cluster_41574 |
| 1057.99000 | 35.0 | Cluster_41578 |
| 1058.02000 | 35.0 | Cluster_41582 |
| 1058.81000 | 35.0 | Cluster_16658 |
| 1058.85000 | 35.0 | Cluster_16649 |
| 1058.96000 | 35.0 | Cluster_41595 |
| 1058.97000 | 35.0 | Cluster_41597 |
| 1059.01000 | 35.0 | Cluster_41596 |

| | | |
|---|---|---|
| 1060.32000 | 35.0 | Cluster_41612 |
| 1060.75000 | 35.0 | Cluster_05502 |
| 1060.82000 | 35.0 | Cluster_16679 |
| 1061.26000 | 35.0 | Cluster_05510 |
| 1061.33000 | 35.0 | Cluster_05506 |
| 1061.82000 | 35.0 | Cluster_41625 |
| 1062.83000 | 35.0 | Cluster_16700 |
| 1063.27000 | 35.0 | Cluster_05516 |
| 1063.82000 | 35.0 | Cluster_41655 |
| 1063.99000 | 35.0 | Cluster_41663 |
| 1064.72000 | 35.0 | Cluster_05521 |
| 1064.73000 | 35.0 | Cluster_41675 |
| 1064.99000 | 35.0 | Cluster_41673 |
| 1066.26000 | 35.0 | Cluster_41702 |
| 1067.52000 | 35.0 | Cluster_41715 |
| 1068.05000 | 35.0 | Cluster_41719 |
| 1068.75000 | 35.0 | Cluster_41734 |
| 1069.51000 | 35.0 | Cluster_41748 |
| 1070.16000 | 35.0 | Cluster_16781 |
| 1070.51000 | 35.0 | Cluster_05546 |
| 1072.55000 | 35.0 | Cluster_16804 |
| 1072.76000 | 35.0 | Cluster_05555 |
| 1074.19000 | 35.0 | Cluster_16831 |
| 1074.51000 | 35.0 | Cluster_41810 |
| 1075.53000 | 35.0 | Cluster_41816 |
| 1076.75000 | 35.0 | Cluster_05574 |
| 1078.01000 | 35.0 | Cluster_05578 |
| 1079.49000 | 35.0 | Cluster_05582 |
| 1080.75000 | 35.0 | Cluster_00593 |
| 1082.00000 | 35.0 | Cluster_05588 |
| 1083.50000 | 35.0 | Cluster_05592 |
| 1086.59000 | 35.0 | Cluster_41940 |
| 1091.58000 | 35.0 | Cluster_41998 |
| 1092.90000 | 35.0 | Cluster_17025 |
| 1093.21000 | 35.0 | Cluster_17031 |
| 1093.55000 | 35.0 | Cluster_17035 |
| 1093.58000 | 35.0 | Cluster_42024 |
| 1093.99000 | 35.0 | Cluster_42029 |
| 1094.92000 | 35.0 | Cluster_00625 |
| 1096.55000 | 35.0 | Cluster_42051 |
| 1100.23000 | 35.0 | Cluster_17091 |
| 1100.56000 | 35.0 | Cluster_17097 |
| 1101.96000 | 35.0 | Cluster_17105 |
| 1102.54000 | 35.0 | Cluster_17111 |
| 1103.12000 | 35.0 | Cluster_42104 |
| 1108.23000 | 35.0 | Cluster_17176 |
| 1108.37000 | 35.0 | Cluster_42148 |
| 1108.88000 | 35.0 | Cluster_17175 |
| 1109.86000 | 35.0 | Cluster_17199 |
| 1113.50000 | 35.0 | Cluster_42184 |
| 1113.53000 | 35.0 | Cluster_17241 |

| | | |
|---|---|---|
| 1115.18000 | 35.0 | Cluster_17258 |
| 1115.77000 | 35.0 | Cluster_00660 |
| 1117.22000 | 35.0 | Cluster_17276 |
| 1118.21000 | 35.0 | Cluster_17277 |
| 1118.22000 | 35.0 | Cluster_17280 |
| 1119.87000 | 35.0 | Cluster_42234 |
| 1121.54000 | 35.0 | Cluster_17322 |
| 1123.04000 | 35.0 | Cluster_42270 |
| 1123.97000 | 35.0 | Cluster_42271 |
| 1124.25000 | 35.0 | Cluster_17359 |
| 1125.57000 | 35.0 | Cluster_42283 |
| 1126.07000 | 35.0 | Cluster_42291 |
| 1127.88000 | 35.0 | Cluster_17398 |
| 1128.57000 | 35.0 | Cluster_17409 |
| 1129.21000 | 35.0 | Cluster_17418 |
| 1129.55000 | 35.0 | Cluster_17428 |
| 1129.90000 | 35.0 | Cluster_17433 |
| 1130.23000 | 35.0 | Cluster_17434 |
| 1131.58000 | 35.0 | Cluster_17448 |
| 1133.13000 | 35.0 | Cluster_42337 |
| 1133.20000 | 35.0 | Cluster_17479 |
| 1133.22000 | 35.0 | Cluster_17484 |
| 1134.90000 | 35.0 | Cluster_17503 |
| 1135.23000 | 35.0 | Cluster_17509 |
| 1135.51000 | 35.0 | Cluster_17513 |
| 1135.54000 | 35.0 | Cluster_17519 |
| 1135.55000 | 35.0 | Cluster_17517 |
| 1136.11000 | 35.0 | Cluster_42363 |
| 1136.88000 | 35.0 | Cluster_17525 |
| 1136.89000 | 35.0 | Cluster_17538 |
| 1138.13000 | 35.0 | Cluster_42377 |
| 1140.22000 | 35.0 | Cluster_17567 |
| 1143.18000 | 35.0 | Cluster_00674 |
| 1143.56000 | 35.0 | Cluster_17607 |
| 1143.60000 | 35.0 | Cluster_42409 |
| 1144.20000 | 35.0 | Cluster_17626 |
| 1144.24000 | 35.0 | Cluster_17620 |
| 1145.21000 | 35.0 | Cluster_17645 |
| 1146.57000 | 35.0 | Cluster_00680 |
| 1149.29000 | 35.0 | Cluster_42461 |
| 1151.05000 | 35.0 | Cluster_42474 |
| 1152.54000 | 35.0 | Cluster_17735 |
| 1152.88000 | 35.0 | Cluster_17750 |
| 1153.03000 | 35.0 | Cluster_05790 |
| 1153.76000 | 35.0 | Cluster_00695 |
| 1154.35000 | 35.0 | Cluster_42485 |
| 1155.34000 | 35.0 | Cluster_42501 |
| 1156.84000 | 35.0 | Cluster_42511 |
| 1158.84000 | 35.0 | Cluster_42519 |
| 1159.61000 | 35.0 | Cluster_42529 |
| 1160.34000 | 35.0 | Cluster_42534 |

| | | |
|---|---|---|
| 1162.11000 | 35.0 | Cluster_42551 |
| 1162.64000 | 35.0 | Cluster_42555 |
| 1163.09000 | 35.0 | Cluster_05817 |
| 1166.08000 | 35.0 | Cluster_05827 |
| 1166.14000 | 35.0 | Cluster_42592 |
| 1166.21000 | 35.0 | Cluster_17894 |
| 1167.19000 | 35.0 | Cluster_17908 |
| 1167.57000 | 35.0 | Cluster_17921 |
| 1168.21000 | 35.0 | Cluster_17931 |
| 1168.33000 | 35.0 | Cluster_42608 |
| 1168.58000 | 35.0 | Cluster_05832 |
| 1173.07000 | 35.0 | Cluster_42645 |
| 1180.80000 | 35.0 | Cluster_05888 |
| 1183.23000 | 35.0 | Cluster_18040 |
| 1187.59000 | 35.0 | Cluster_18075 |
| 1190.56000 | 35.0 | Cluster_18091 |
| 1192.81000 | 35.0 | Cluster_42765 |
| 1192.92000 | 35.0 | Cluster_18117 |
| 1195.88000 | 35.0 | Cluster_18133 |
| 1196.63000 | 35.0 | Cluster_42787 |
| 1197.59000 | 35.0 | Cluster_18145 |
| 1197.90000 | 35.0 | Cluster_18140 |
| 1198.64000 | 35.0 | Cluster_42803 |
| 1199.32000 | 35.0 | Cluster_05982 |
| 1199.57000 | 35.0 | Cluster_18161 |
| 1200.26000 | 35.0 | Cluster_18168 |
| 1200.56000 | 35.0 | Cluster_18177 |
| 1201.89000 | 35.0 | Cluster_18192 |
| 1203.44000 | 35.0 | Cluster_42830 |
| 1206.62000 | 35.0 | Cluster_42853 |
| 1208.56000 | 35.0 | Cluster_18237 |
| 1209.51000 | 35.0 | Cluster_42887 |
| 1213.12000 | 35.0 | Cluster_42910 |
| 1215.15000 | 35.0 | Cluster_42924 |
| 1218.11000 | 35.0 | Cluster_42950 |
| 1219.09000 | 35.0 | Cluster_42956 |
| 1221.25000 | 35.0 | Cluster_18299 |
| 1222.10000 | 35.0 | Cluster_42968 |
| 1224.08000 | 35.0 | Cluster_42977 |
| 1224.60000 | 35.0 | Cluster_42983 |
| 1225.59000 | 35.0 | Cluster_42994 |
| 1228.07000 | 35.0 | Cluster_43008 |
| 1228.11000 | 35.0 | Cluster_43007 |
| 1230.63000 | 35.0 | Cluster_43026 |
| 1232.57000 | 35.0 | Cluster_43045 |
| 1234.58000 | 35.0 | Cluster_43065 |
| 1238.57000 | 35.0 | Cluster_43079 |
| 1245.28000 | 35.0 | Cluster_18390 |
| 1249.16000 | 35.0 | Cluster_43135 |
| 1251.87000 | 35.0 | Cluster_43149 |
| 1253.87000 | 35.0 | Cluster_43162 |

| | | |
|---|---|---|
| 1255.86000 | 35.0 | Cluster_43176 |
| 1257.08000 | 35.0 | Cluster_06282 |
| 1264.27000 | 35.0 | Cluster_18463 |
| 1267.20000 | 35.0 | Cluster_43224 |
| 1267.68000 | 35.0 | Cluster_43228 |
| 1274.29000 | 35.0 | Cluster_18502 |
| 1284.15000 | 35.0 | Cluster_43287 |
| 1300.69000 | 35.0 | Cluster_43366 |
| 1300.70000 | 35.0 | Cluster_43360 |
| 1300.73000 | 35.0 | Cluster_43361 |
| 1300.89000 | 35.0 | Cluster_43362 |
| 1301.68000 | 35.0 | Cluster_43365 |
| 1320.21000 | 35.0 | Cluster_43449 |
| 1329.73000 | 35.0 | Cluster_43484 |
| 1330.45000 | 35.0 | Cluster_06499 |
| 1331.69000 | 35.0 | Cluster_43492 |
| 1334.68000 | 35.0 | Cluster_43505 |
| 1349.89000 | 35.0 | Cluster_43562 |
| 1353.71000 | 35.0 | Cluster_43578 |
| 1354.21000 | 35.0 | Cluster_43579 |
| 1354.66000 | 35.0 | Cluster_06571 |
| 1361.03000 | 35.0 | Cluster_18681 |
| 1367.16000 | 35.0 | Cluster_43645 |
| 1372.75000 | 35.0 | Cluster_43667 |
| 1376.18000 | 35.0 | Cluster_43680 |
| 1376.34000 | 35.0 | Cluster_18707 |
| 1383.67000 | 35.0 | Cluster_18716 |
| 1394.65000 | 35.0 | Cluster_43739 |
| 1396.44000 | 35.0 | Cluster_18732 |
| 1402.12000 | 35.0 | Cluster_18741 |
| 1406.24000 | 35.0 | Cluster_43793 |
| 1407.03000 | 35.0 | Cluster_18752 |
| 1414.35000 | 35.0 | Cluster_18768 |
| 1417.37000 | 35.0 | Cluster_18772 |
| 1420.01000 | 35.0 | Cluster_18783 |
| 1422.96000 | 35.0 | Cluster_43843 |
| 1423.47000 | 35.0 | Cluster_43844 |
| 1424.69000 | 35.0 | Cluster_18789 |
| 1424.79000 | 35.0 | Cluster_43851 |
| 1425.27000 | 35.0 | Cluster_43853 |
| 1427.03000 | 35.0 | Cluster_18794 |
| 1428.82000 | 35.0 | Cluster_43862 |
| 1429.02000 | 35.0 | Cluster_18795 |
| 1430.01000 | 35.0 | Cluster_18796 |
| 1431.68000 | 35.0 | Cluster_18799 |
| 1437.03000 | 35.0 | Cluster_18803 |
| 1439.35000 | 35.0 | Cluster_18807 |
| 1459.27000 | 35.0 | Cluster_43959 |
| 1477.30000 | 35.0 | Cluster_44015 |
| 1489.22000 | 35.0 | Cluster_44043 |
| 1492.94000 | 35.0 | Cluster_44054 |

| | | |
|---|---|---|
| 1495.14000 | 35.0 | Cluster_44065 |
| 1511.80000 | 35.0 | Cluster_44102 |
| 1518.72000 | 35.0 | Cluster_44112 |
| 1530.33000 | 35.0 | Cluster_44125 |
| 1560.76000 | 35.0 | Cluster_44175 |
| 1564.32000 | 35.0 | Cluster_44184 |
| 1571.76000 | 35.0 | Cluster_44194 |
| 1582.85000 | 35.0 | Cluster_44218 |
| 1616.85000 | 35.0 | Cluster_44250 |
| 1633.97000 | 35.0 | Cluster_44257 |
| 1635.40000 | 35.0 | Cluster_44260 |
| 1676.82000 | 35.0 | Cluster_44320 |
| 1683.84000 | 35.0 | Cluster_44341 |
| 1687.92000 | 35.0 | Cluster_44353 |
| 1739.94000 | 35.0 | Cluster_44403 |
| 1792.99000 | 35.0 | Cluster_44425 |

Reject Mass List: (none)
Neutral Loss Mass List: (none)
Product Mass List: (none)
Neutral loss in top: 3
Product in top: 3
Most intense if no parent masses found enabled
Add/subtract mass not enabled
FT master scan preview mode enabled
Charge state screening enabled
Charge state dependent ETD time not enabled
Monoisotopic precursor selection enabled
Charge state rejection enabled
Unassigned charge states : rejected
Charge state 1 : rejected
Charge state 2 : not rejected
Charge state 3 : not rejected
Charge states 4+ : not rejected
Chromatography mode is disabled

Global Data Dependent Settings:
Predict ion injection time enabled
Use global parent and reject mass lists not enabled
Exclude parent mass from data dependent selection not enabled
Exclusion mass width by mass
Exclusion mass width low: 0.50000
Exclusion mass width high: 0.50000
Parent mass width by mass
Parent mass width low: 0.50000
Parent mass width high: 0.50000
Reject mass width by mass
Reject mass width low: 0.50000
Reject mass width high: 0.50000
Zoom/UltraZoom scan mass width by mass
Zoom/UltraZoom scan mass width low: 5.00
Zoom/UltraZoom scan mass width high: 5.00

FT SIM scan mass width low: 5.00
FT SIM scan mass width high: 5.00
Neutral Loss candidates processed by decreasing intensity
Neutral Loss mass width by mass
Neutral Loss mass width low: 0.50000
Neutral Loss mass width high: 0.50000
Product candidates processed by decreasing intensity
Product mass width by mass
Product mass width low: 0.50000
Product mass width high: 0.50000
MS mass range: 0.00-1000000.00
MSn mass range by mass
MSn mass range: 0.00-1000000.00
Use m/z values as masses not enabled
Analog UV data dep. not enabled
Dynamic exclusion not enabled
Isotopic data dependence not enabled
Mass Tags data dependence not enabled

Custom Data Dependent Settings:

Not enabled

#### 4. Galaxy workflow to create a wheat non redundant FASTA database with contaminant and decoy sequences

To use the workflow below, simply copy/paste the command line into a text editor and save it with a .ga extension. Galaxy will then be able to recognise and import it.

```
{
    "a_galaxy_workflow": "true",
    "annotation": "",
    "format-version": "0.1",
    "name": "Wheat - creating non redundant FASTA DBs with decoy",
    "steps": {
        "0": {
            "annotation": "",
            "content_id": null,
            "errors": null,
            "id": 0,
            "input_connections": {},
            "inputs": [
                {
                    "description": "",
                    "name": "from-Gabriel_alpha-amylase-blast_2022-
03-08.fasta"
                }
            ],
            "label": "from-Gabriel_alpha-amylase-blast_2022-03-
08.fasta",
            "name": "Input dataset",
            "outputs": [],
            "position": {
                "bottom": 207.96875,
                "height": 102.5625,
                "left": 360.25,
                "right": 560.25,
                "top": 105.40625,
                "width": 200,
                "x": 360.25,
                "y": 105.40625
            },
            "tool_id": null,
            "tool_state": "{\"optional\": false}",
            "tool_version": null,
            "type": "data_input",
            "uuid": "b4d34c10-e494-478e-b73a-e0672b3431f6",
            "workflow_outputs": []
        },
        "1": {
            "annotation": "",
            "content_id": null,
            "errors": null,
            "id": 1,
            "input_connections": {},
            "inputs": [
                {
                    "description": "",
```

```
                    "name": "Triticum_aestivum.IWGSC.pep.all.fa"
                }
            ],
            "label": "Triticum_aestivum.IWGSC.pep.all.fa",
            "name": "Input dataset",
            "outputs": [],
            "position": {
                "bottom": 318.96875,
                "height": 102.5625,
                "left": 358.25,
                "right": 558.25,
                "top": 216.40625,
                "width": 200,
                "x": 358.25,
                "y": 216.40625
            },
            "tool_id": null,
            "tool_state": "{\"optional\": false}",
            "tool_version": null,
            "type": "data_input",
            "uuid": "7152c491-219f-47f0-ad77-cad307e81d00",
            "workflow_outputs": []
        },
        "2": {
            "annotation": "",
            "content_id": null,
            "errors": null,
            "id": 2,
            "input_connections": {},
            "inputs": [
                {
                    "description": "",
                    "name": "uniprot_Triticum-
aestivum_142969entries_2020-02-26.fasta"
                }
            ],
            "label": "uniprot_Triticum-aestivum_142969entries_2020-
02-26.fasta",
            "name": "Input dataset",
            "outputs": [],
            "position": {
                "bottom": 452.359375,
                "height": 122.953125,
                "left": 355.25,
                "right": 555.25,
                "top": 329.40625,
                "width": 200,
                "x": 355.25,
                "y": 329.40625
            },
            "tool_id": null,
            "tool_state": "{\"optional\": false}",
            "tool_version": null,
            "type": "data_input",
            "uuid": "fda351ea-f08d-426d-a946-4eaefa86dd0a",
            "workflow_outputs": []
```

```
        },
        "3": {
            "annotation": "",
            "content_id": null,
            "errors": null,
            "id": 3,
            "input_connections": {},
            "inputs": [
                {
                    "description": "",
                    "name": "1-uniprot_Triticum-
aestivum_142969entries_2020-02-26.fasta"
                }
            ],
            "label": "1-uniprot_Triticum-
aestivum_142969entries_2020-02-26.fasta",
            "name": "Input dataset",
            "outputs": [],
            "position": {
                "bottom": 633.359375,
                "height": 122.953125,
                "left": 353.25,
                "right": 553.25,
                "top": 510.40625,
                "width": 200,
                "x": 353.25,
                "y": 510.40625
            },
            "tool_id": null,
            "tool_state": "{\"optional\": false}",
            "tool_version": null,
            "type": "data_input",
            "uuid": "f2f13af5-a14a-4d43-9ef1-e26b0eb88597",
            "workflow_outputs": []
        },
        "4": {
            "annotation": "",
            "content_id": null,
            "errors": null,
            "id": 4,
            "input_connections": {},
            "inputs": [
                {
                    "description": "",
                    "name": "2-
Triticum_aestivum.IWGSC.pep.all.fasta"
                }
            ],
            "label": "2-Triticum_aestivum.IWGSC.pep.all.fasta",
            "name": "Input dataset",
            "outputs": [],
            "position": {
                "bottom": 741.96875,
                "height": 102.5625,
                "left": 349.25,
                "right": 549.25,
```

```
                "top": 639.40625,
                "width": 200,
                "x": 349.25,
                "y": 639.40625
            },
            "tool_id": null,
            "tool_state": "{\"optional\": false}",
            "tool_version": null,
            "type": "data_input",
            "uuid": "274c7c93-e1f0-428d-9fce-75e7d69ef118",
            "workflow_outputs": []
        },
        "5": {
            "annotation": "",
            "content_id":
"toolshed.g2.bx.psu.edu/repos/galaxyp/dbbuilder/dbbuilder/0.3.1",
            "errors": null,
            "id": 5,
            "input_connections": {},
            "inputs": [],
            "label": null,
            "name": "Protein Database Downloader",
            "outputs": [
                {
                    "name": "output_database",
                    "type": "fasta"
                }
            ],
            "position": {
                "bottom": 557.96875,
                "height": 102.5625,
                "left": 607.25,
                "right": 807.25,
                "top": 455.40625,
                "width": 200,
                "x": 607.25,
                "y": 455.40625
            },
            "post_job_actions": {},
            "tool_id":
"toolshed.g2.bx.psu.edu/repos/galaxyp/dbbuilder/dbbuilder/0.3.1",
            "tool_shed_repository": {
                "changeset_revision": "c1b437242fee",
                "name": "dbbuilder",
                "owner": "galaxyp",
                "tool_shed": "toolshed.g2.bx.psu.edu"
            },
            "tool_state": "{\"source\": {\"from\": \"cRAP\",
\"__current_case__\": 1}, \"__page__\": null,
\"__rerun_remap_job_id__\": null}",
            "tool_version": "0.3.1",
            "type": "tool",
            "uuid": "0720d479-9fbe-4ba0-9535-c4d97cddcc23",
            "workflow_outputs": [
                {
                    "label": null,
```

```
                    "output_name": "output_database",
                    "uuid": "0d4d9480-f55e-40be-9f17-d5b63eb3f719"
                }
            ]
        },
        "6": {
            "annotation": "",
            "content_id": null,
            "errors": null,
            "id": 6,
            "input_connections": {},
            "inputs": [
                {
                    "description": "",
                    "name": "3-from-Gabriel_alpha-amylase-
blast_2022-03-08.fasta"
                }
            ],
            "label": "3-from-Gabriel_alpha-amylase-blast_2022-03-
08.fasta",
            "name": "Input dataset",
            "outputs": [],
            "position": {
                "bottom": 850.96875,
                "height": 102.5625,
                "left": 352.25,
                "right": 552.25,
                "top": 748.40625,
                "width": 200,
                "x": 352.25,
                "y": 748.40625
            },
            "tool_id": null,
            "tool_state": "{\"optional\": false}",
            "tool_version": null,
            "type": "data_input",
            "uuid": "ac70a284-a0f1-469f-bd75-899a0d3e9905",
            "workflow_outputs": []
        },
        "7": {
            "annotation": "",
            "content_id":
"toolshed.g2.bx.psu.edu/repos/galaxyp/fasta_merge_files_and_filter_u
nique_sequences/fasta_merge_files_and_filter_unique_sequences/1.2.0"
,
            "errors": null,
            "id": 7,
            "input_connections": {
                "batchmode|input_fastas": [
                    {
                        "id": 0,
                        "output_name": "output"
                    },
                    {
                        "id": 1,
                        "output_name": "output"
```

```
                    },
                    {
                        "id": 2,
                        "output_name": "output"
                    },
                    {
                        "id": 5,
                        "output_name": "output_database"
                    }
                ]
            },
            "inputs": [],
            "label": null,
            "name": "FASTA Merge Files and Filter Unique Sequences",
            "outputs": [
                {
                    "name": "output",
                    "type": "fasta"
                }
            ],
            "position": {
                "bottom": 455.140625,
                "height": 174.734375,
                "left": 938.25,
                "right": 1138.25,
                "top": 280.40625,
                "width": 200,
                "x": 938.25,
                "y": 280.40625
            },
            "post_job_actions": {},
            "tool_id":
"toolshed.g2.bx.psu.edu/repos/galaxyp/fasta_merge_files_and_filter_u
nique_sequences/fasta_merge_files_and_filter_unique_sequences/1.2.0"
,
            "tool_shed_repository": {
                "changeset_revision": "f546e7278f04",
                "name":
"fasta_merge_files_and_filter_unique_sequences",
                "owner": "galaxyp",
                "tool_shed": "toolshed.g2.bx.psu.edu"
            },
            "tool_state": "{\"accession_parser\": \"^>([^ ]+).*$\",
\"batchmode\": {\"processmode\": \"merge\", \"__current_case__\": 1,
\"input_fastas\": {\"__class__\": \"ConnectedValue\"}},
\"uniqueness_criterion\": \"sequence\", \"__page__\": null,
\"__rerun_remap_job_id__\": null}",
            "tool_version": "1.2.0",
            "type": "tool",
            "uuid": "acf11599-e033-46b5-8ca7-ab97ae8be58b",
            "workflow_outputs": [
                {
                    "label": null,
                    "output_name": "output",
                    "uuid": "848c5bf4-290e-4608-a129-dc565c7692d6"
                }
```

```
            ]
        },
        "8": {
            "annotation": "",
            "content_id":
"toolshed.g2.bx.psu.edu/repos/galaxyp/fasta_merge_files_and_filter_u
nique_sequences/fasta_merge_files_and_filter_unique_sequences/1.2.0"
,
            "errors": null,
            "id": 8,
            "input_connections": {
                "batchmode|input_fastas": [
                    {
                        "id": 3,
                        "output_name": "output"
                    },
                    {
                        "id": 4,
                        "output_name": "output"
                    },
                    {
                        "id": 6,
                        "output_name": "output"
                    },
                    {
                        "id": 5,
                        "output_name": "output_database"
                    }
                ]
            },
            "inputs": [],
            "label": null,
            "name": "FASTA Merge Files and Filter Unique Sequences",
            "outputs": [
                {
                    "name": "output",
                    "type": "fasta"
                }
            ],
            "position": {
                "bottom": 704.140625,
                "height": 174.734375,
                "left": 939.25,
                "right": 1139.25,
                "top": 529.40625,
                "width": 200,
                "x": 939.25,
                "y": 529.40625
            },
            "post_job_actions": {},
            "tool_id":
"toolshed.g2.bx.psu.edu/repos/galaxyp/fasta_merge_files_and_filter_u
nique_sequences/fasta_merge_files_and_filter_unique_sequences/1.2.0"
,
            "tool_shed_repository": {
                "changeset_revision": "f546e7278f04",
```

                "name":
"fasta_merge_files_and_filter_unique_sequences",
                "owner": "galaxyp",
                "tool_shed": "toolshed.g2.bx.psu.edu"
            },
            "tool_state": "{\"accession_parser\": \"^>([^ ]+).*$\",
\"batchmode\": {\"processmode\": \"merge\", \"__current_case__\": 1,
\"input_fastas\": {\"__class__\": \"ConnectedValue\"}},
\"uniqueness_criterion\": \"sequence\", \"__page__\": null,
\"__rerun_remap_job_id__\": null}",
            "tool_version": "1.2.0",
            "type": "tool",
            "uuid": "1f78efc3-aeaf-44e3-9400-97da2feaee9c",
            "workflow_outputs": [
                {
                    "label": null,
                    "output_name": "output",
                    "uuid": "08d15055-fb91-42ff-b946-e979934c55f9"
                }
            ]
        },
        "9": {
            "annotation": "",
            "content_id":
"toolshed.g2.bx.psu.edu/repos/galaxyp/openms_decoydatabase/DecoyData
base/2.6+galaxy0",
            "errors": null,
            "id": 9,
            "input_connections": {
                "in": {
                    "id": 7,
                    "output_name": "output"
                }
            },
            "inputs": [],
            "label": null,
            "name": "DecoyDatabase",
            "outputs": [
                {
                    "name": "out",
                    "type": "fasta"
                }
            ],
            "position": {
                "bottom": 445.75,
                "height": 154.34375,
                "left": 1210.25,
                "right": 1410.25,
                "top": 291.40625,
                "width": 200,
                "x": 1210.25,
                "y": 291.40625
            },
            "post_job_actions": {},

```
            "tool_id":
"toolshed.g2.bx.psu.edu/repos/galaxyp/openms_decoydatabase/DecoyData
base/2.6+galaxy0",
            "tool_shed_repository": {
                "changeset_revision": "370141bc0da3",
                "name": "openms_decoydatabase",
                "owner": "galaxyp",
                "tool_shed": "toolshed.g2.bx.psu.edu"
            },
            "tool_state": "{\"Decoy\": {\"non_shuffle_pattern\":
\"\", \"keepPeptideNTerm\": \"true\", \"keepPeptideCTerm\":
\"true\"}, \"OPTIONAL_OUTPUTS\": null, \"adv_opts_cond\":
{\"adv_opts_selector\": \"basic\", \"__current_case__\": 0},
\"decoy_string\": \"DECOY_\", \"decoy_string_position\": \"prefix\",
\"enzyme\": \"Trypsin\", \"in\": {\"__class__\":
\"ConnectedValue\"}, \"method\": \"reverse\", \"only_decoy\":
\"false\", \"type\": \"protein\", \"__page__\": null,
\"__rerun_remap_job_id__\": null}",
            "tool_version": "2.6+galaxy0",
            "type": "tool",
            "uuid": "68e37f73-0773-4ff3-895b-5286dea2efe1",
            "workflow_outputs": [
                {
                    "label": null,
                    "output_name": "out",
                    "uuid": "5ee1eec0-9464-4428-81ea-263b09602f36"
                }
            ]
        },
        "10": {
            "annotation": "",
            "content_id":
"toolshed.g2.bx.psu.edu/repos/galaxyp/openms_decoydatabase/DecoyData
base/2.6+galaxy0",
            "errors": null,
            "id": 10,
            "input_connections": {
                "in": {
                    "id": 8,
                    "output_name": "output"
                }
            },
            "inputs": [],
            "label": null,
            "name": "DecoyDatabase",
            "outputs": [
                {
                    "name": "out",
                    "type": "fasta"
                }
            ],
            "position": {
                "bottom": 695.75,
                "height": 154.34375,
                "left": 1216.25,
                "right": 1416.25,
```

```
                "top": 541.40625,
                "width": 200,
                "x": 1216.25,
                "y": 541.40625
            },
            "post_job_actions": {},
            "tool_id":
"toolshed.g2.bx.psu.edu/repos/galaxyp/openms_decoydatabase/DecoyDatabase/2.6+galaxy0",
            "tool_shed_repository": {
                "changeset_revision": "370141bc0da3",
                "name": "openms_decoydatabase",
                "owner": "galaxyp",
                "tool_shed": "toolshed.g2.bx.psu.edu"
            },
            "tool_state": "{\"Decoy\": {\"non_shuffle_pattern\":
\"\", \"keepPeptideNTerm\": \"true\", \"keepPeptideCTerm\":
\"true\"}, \"OPTIONAL_OUTPUTS\": null, \"adv_opts_cond\":
{\"adv_opts_selector\": \"basic\", \"__current_case__\": 0},
\"decoy_string\": \"DECOY_\", \"decoy_string_position\": \"prefix\",
\"enzyme\": \"Trypsin\", \"in\": {\"__class__\":
\"ConnectedValue\"}, \"method\": \"reverse\", \"only_decoy\":
\"false\", \"type\": \"protein\", \"__page__\": null,
\"__rerun_remap_job_id__\": null}",
            "tool_version": "2.6+galaxy0",
            "type": "tool",
            "uuid": "0c5ec050-869a-4657-b1f7-e8827236f495",
            "workflow_outputs": [
                {
                    "label": null,
                    "output_name": "out",
                    "uuid": "9bf5c5ff-cb8b-4dc7-809d-53f987794688"
                }
            ]
        }
    },
    "tags": [],
    "uuid": "53b7f604-35b5-41ec-b665-bedbd5c6a8f7",
    "version": 2
}
```
